# Parallel evolution of methyltransferases leads to vobasine biosynthesis in *Tabernaemontana elegans* and *Catharanthus roseus*

**DOI:** 10.1101/2024.07.28.605509

**Authors:** Maisha Farzana, Matthew Bailey Richardson, Daniel André Ramey Deschênes, Zhan Mai, Destiny Ichechi Njoku, Ghislain Deslongchamps, Yang Qu

**Affiliations:** Department of Chemistry, University of New Brunswick, Fredericton, NB, E3B 5A3, Canada

**Author notes:** Correspondence: Ghislain Deslongchamps, Yang Qu.

## Abstract

Monoterpenoid indole alkaloids (MIA) are one of the largest and most complex alkaloid class in nature, boasting many clinically significant drugs such as anticancer vinblastine and antiarrhythmic ajmaline. Many MIAs undergo nitrogen *N*-methylation, altering their reactivity and affinity to the biological targets through a straightforward reaction. Remarkably, all known MIA *N*-methyltransferases (NMT) originate from the neofunctionalization of ancestral γ-tocopherol *C*-methyltransferases (γTMTs), a phenomenon seemingly unique to the Apocynaceae family. In this study, we unveil and characterize a new γTMT-like enzyme from the plant *Tabernaemontana elegans* (toad tree): perivine *N*β-methyltransferase (TePeNMT). TePeNMT and other homologs form a distinct clade in our phylogenetic study, setting them apart from other γTMTs and γTMT-like NMTs discovered to date. Enzyme kinetic experiments and enzyme homology modeling studies reveal the significant differences in enzyme active sites between TePeNMT and CrPeNMT, a previously characterized perivine *N*β-methyltransferase from *Catharanthus roseus* (Madagascar periwinkle). Collectively, our findings suggest that parallel evolution of ancestral γTMTs may be responsible for the occurrence of perivine *N*-methylation in *T. elegans* and *C. roseus*.

## Introduction

Natural product methylation plays a key role in diversifying structures, altering compound polarity, membrane permeability and stability, and modifying their electronic properties for target interaction [1]. A prominent example lies in the frequent *O*-methylation of phenylpropanoids and flavonoids within plants, integral to cell wall biosynthesis and specialized metabolism, with profound implications for human health [2]. Similarly, nitrogen *N*-methylation, alongside *O*-methylation, significantly contribute to the diversity of plant alkaloids, such as benzylisoquinoline alkaloids (BIA) and monoterpenoid indole alkaloids (MIA). Notable instances include (*S*)-coclaurine *N*-methylation and benzylisoquinoline *O*-methylations in the biosynthesis of morphine (BIA), as well as tabersonine 16-*O*-methylation and 16-methoxy-2,3-dihydrotabersonine *N*-methylation in vinblastine (MIA) production [3,4]. Promoting van der Waals interactions and alterations in molecular electronic properties, these methylations are crucial for downstream biosynthetic enzyme recognition, underscoring their indispensability in morphine and vinblastine biosynthesis. The acquisition and retention of these methylations likely conferred adaptive advantages during the evolutionary processes of host plants opium poppy and *Catharanthus roseus* (Madagascar’s periwinkle).

Phylogenetic investigations propose that all identified MIA *N*-methyltransferases (NMT) likely originated from the neofunctionalization of ancestral γ-tocopherol (vitamin E) *C*-methyltransferases (γTMT) [3] (Fig. 1). The first characterized MIA NMT is the 16-methoxy-2,3-dihydrotabersonine *N-*methyltransferase (CrDhtNMT) from *C. roseus* [3,5,6]. Unlike the plastid-located γTMTs, CrDhtNMT lacks a chloroplast transit peptide, indicative of its novel role in cytosolic MIA biosynthesis. While CrDhtNMT does not catalyze γ-tocopherol methylation, it retains the ability to bind γ-tocopherol, as evidenced by its inhibition by γ-tocopherol [3]. Homologous γTMT-like NMTs have also been identified in other MIA-producing species within the Apocynaceae family. RsNNMT, RsANMT, RsPiNMT, VmPiNMT, and CrPeNMT are responsible for the *N*-methylation of norajmaline, ajmaline, picrinine, and perivine respectively, in *Rauwolfia serpentina* (Indian snakeroot), *Vinca minor*, and *C. roseus* [7–10] (Fig. 2). Although previous studies associated CrDhtNMT activity with the chloroplast through sucrose gradient centrifugation [6], recent findings utilizing fluorescent protein tagging suggest its localization in peroxisomes, while other MIA NMT members exhibit either cytosolic or vacuolar association [10].

**Figure 1.**
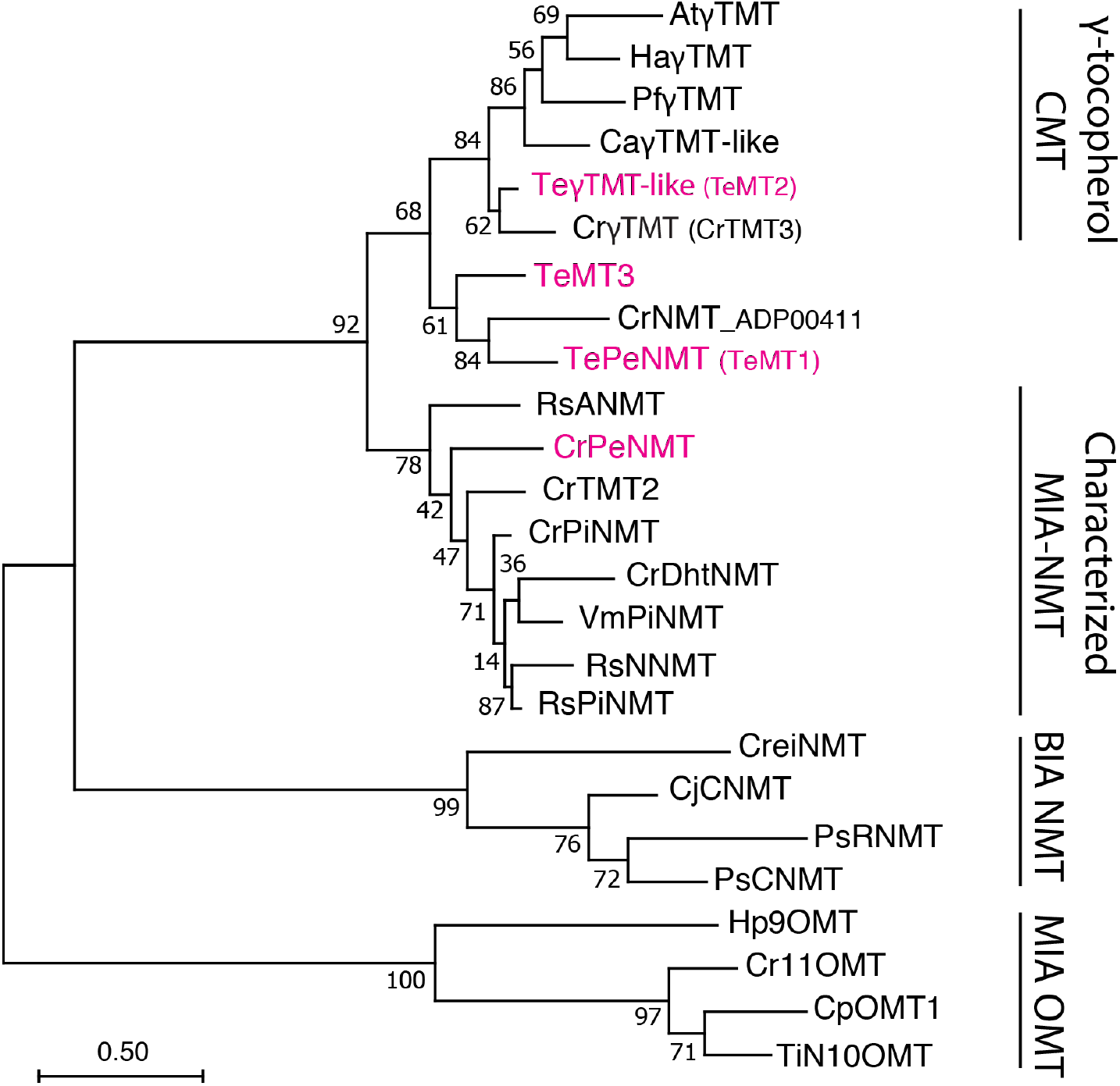
Phylogeny of γ-tocopherol *C*-methyltransferase (CMT)-like *N*-methyltransferases (NMT) in Apocynaceae family. The *Tabernaemontana elegans* perivine *N*β-NMT (TePeNMT, TeMT1) and homologs from *Catharanthus roseus* and *Vinca minor* form a new clade, distinguishing them from the *bona fide* γ-TMTs and other characterized monoterpenoid indole alkaloid (MIA) NMTs to date. The enzymes labeled in red were investigated in this study. The evolutionary history was inferred by using the Maximum Likelihood method and JTT matrix-based model. The tree with the highest log likelihood is shown. The percentage of trees in which the associated taxa clustered together is shown next to the branches (500 bootstrap replicates). Initial tree(s) for the heuristic search were obtained automatically by applying Neighbor-Join and BioNJ algorithms to a matrix of pairwise distances estimated using the JTT model, and then selecting the topology with superior log likelihood value. The tree is drawn to scale, with branch lengths measured in the number of substitutions per site (scale bar). Evolutionary analyses were conducted in MEGA11. OMT: *O*-methyltransferase; BIA: benzylisoquinoline alkaloids; PiNMT: picrinine NMT; DhtNMT: 16-methoxyl-2,3-dihydrotabersonine NMT; NNMT: norajmaline NMT; ANMT; ajmaline *N*β-NMT; CNMT: coclaurine NMT; RNMT: reticuline NMT; At: *Arabidopsis thaliana*; Ca: *Coffea arabica*; Cp: *Cinchona pubescens*; Cr: Catharanthus roseus; Cre: Chlamydomonas reinhardtii; Cj: *Coptis japonica*; Ha: *Helianthus annuus*; Hp: Hamelia patens; Pf: Perilla frutescens; Ps: *Papaver somniferum*; Rs: *Rauvolfia serpentina*; Te: *Tabernaemontana elegans*; Ti: *Tabernanthe iboga*; Vm: *Vinca minor*. The alignment for constructing the phylogenetic tree is included in Supplementary data 1.

**Figure 2.**
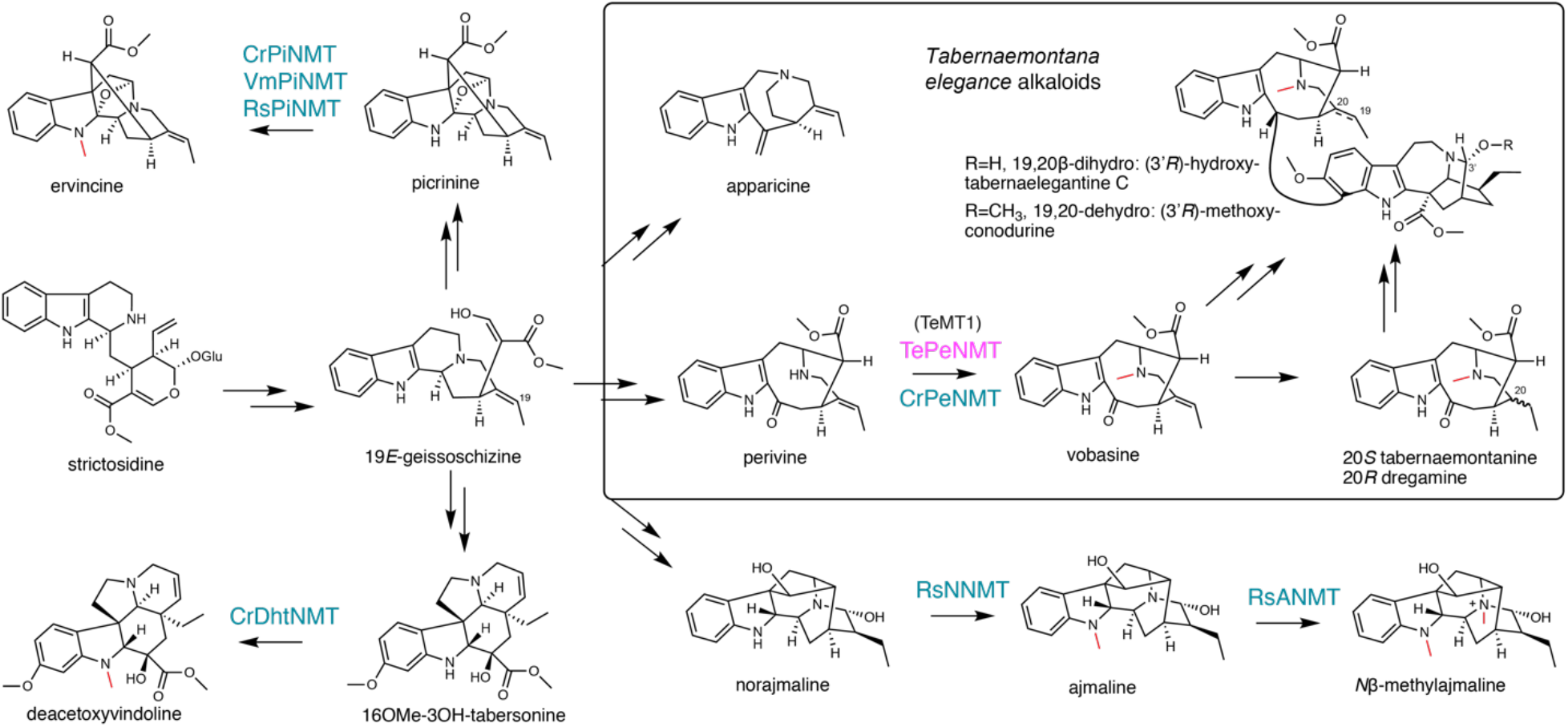
γ-tocopherol *C*-methyltransferase-like *N*-methyltransferases (NMT) catalyze diverse reactions, contributing to monoterpenoid indole alkaloid (MIA) diversity. The methyl substitutions on MIAs are labeled in red. The previously characterized NMTs are labeled in blue. The *Tabernaemontana elegans* perivine *N*β-NMT characterized in this study is labeled in magenta.

The *N*β-methylation of perivine (i.e., methylation of the non-indole nitrogen), resulting in the formation of vobasine, is frequently observed in *Tabernaemontana* genus and in other plants in the Apocynaceae family. Vobasine exhibits moderate antifungal and anticancer properties, characteristics shared by its two 19,20-reduced derivatives, tabernaemontanine, and dregamine [11–14] (Fig. 2). Notably, vobasine and vobasine-type MIAs are often detected as dimerized bisindole alkaloids, either with the same type or a different MIA type, in *Tabernaemontana* spp., demonstrating heightened anticancer activities [13–16]. For example, the iboga-vobasinyl bisindole alkaloid (3’*R*)-hydroxytabernaelegantine C (Fig. 2) has been shown to induce potent apoptosis in colon and liver cancer cells [16]. Although *N*β-methylperivine (vobasine) and *N*β-formylperivine (periformyline) have been identified in *Catharanthus* spp. [17,18], our research indicated that non-methylated perivine is the primary accumulating form for vobasine type MIAs in *C. roseus* [19]. In *Tabernaemontana* spp., *N*-methylated vobasine and its derivatives are frequently observed [11,20,21].

In this study, we discover and characterize a novel enzyme TePeNMT responsible for perivine *N*β-methylation in the plant *Tabernaemontana elegans* (toad tree). Surprisingly, despite both enzymes catalyzing the same reaction, TePeNMT exhibited a low amino acid identity of 50% when compared to CrPeNMT. Phylogenetic analysis revealed that TePeNMT, along with several other γTMT-like enzymes in the Apocynaceae family, constitutes a distinct subgroup that shares closer evolutionary ties with genuine γTMTs. Homology modeling of CrPeNMT and TePeNMT revealed drastic differences in active site conformation and perivine substrate docking positions. Our findings strongly suggest the possibility of parallel evolution being responsible for perivine *N*-methylation in *C. roseus* and *T. elegans*.

## Results

### Vobasine and apparicine are the two most abundant MIAs in *T. elegans* leaf

To study the biosynthesis of vobasine and vobasinyl bisindole alkaloids, we investigated the MIA profiles of the leaf, leaf latex, flower, and young root tissues of *T. elegans* growing in our greenhouse using liquid chromatography tandem mass spectrometry (LC-MS/MS). Two major MIAs with [M+H]^+^ *m/z* 265 and 353 accumulated in leaf and flowers (Fig. 3A). In comparison, the MIA profile in *T. elegans* young roots exhibited greater diversity, encompassing the *m/z* 265 and *m/z* 353 MIAs. Notably, leaf latex contained only minimal amounts of MIAs (Fig. 3A). This finding suggests that MIAs in *T. elegans* are not sequestered within the leaf latex, in contrast to *C. roseus* and *Rauvolfia tetraphylla*, where laticifers and idioblasts (a special type of parenchyma cells) serve as major sinks for MIAs [22–25].

**Figure 3.**
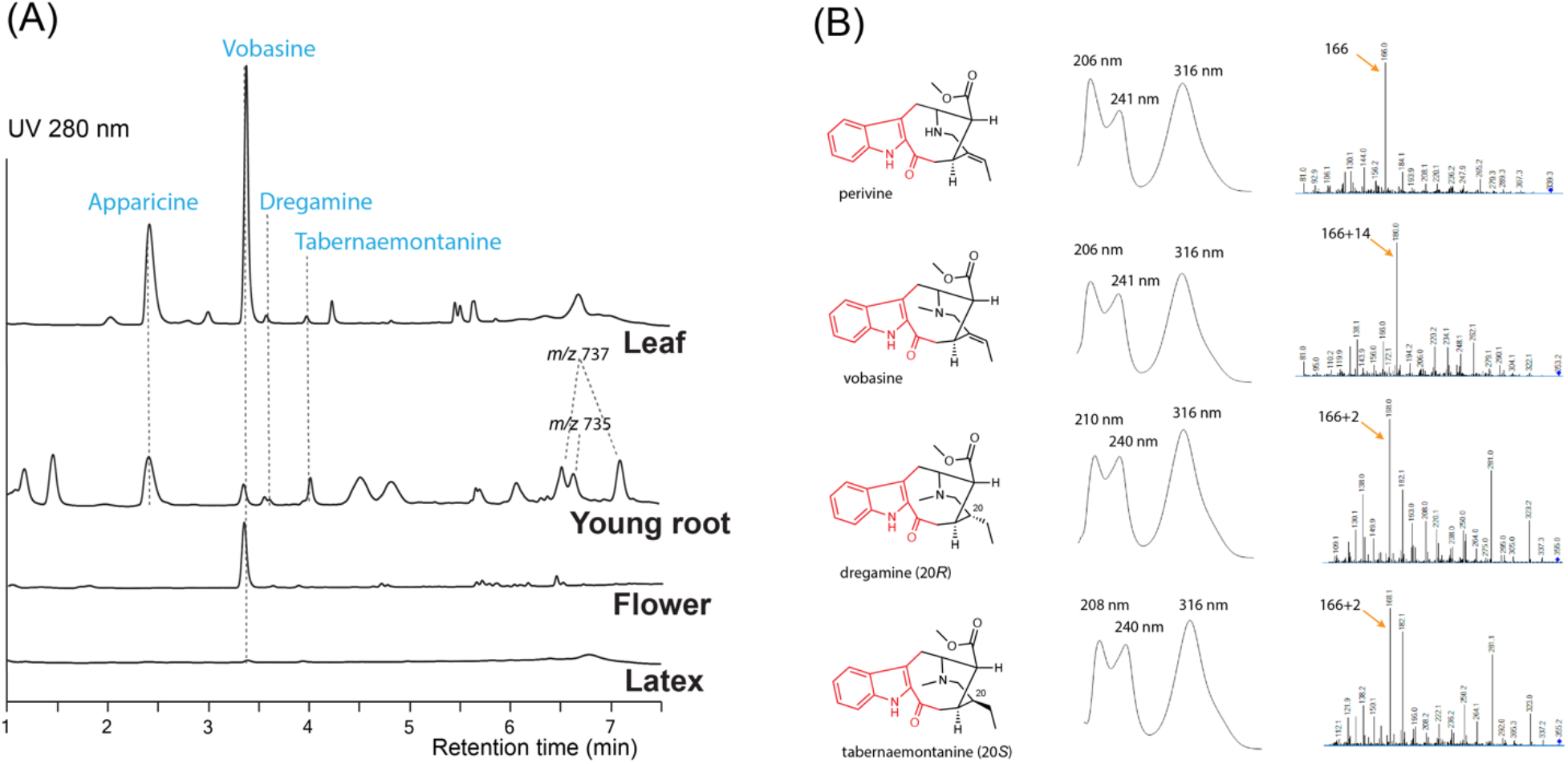
Representative LC-MS chromatograms (A) for *Tabernaemontana elegans* tissues (leaf, young root, flower, and latex) and alkaloid UV absorption and MS/MS profiles (B). Each chromatogram (UV 280 nm) was normalized by tissue weight and drawn to scale. Apparicine and vobasine were the two most abundant alkaloids in leaf tissues, while peaks tentatively identified as bisindole alkaloids (*m/z* 735 and 737) were detected in young root tissues. The chromophores are labeled in red.

We purified the *m/z* 265 and *m/z* 353 MIAs from *T. elegans* leaf by thin layer chromatography, and identified them as apparicine and vobasine, respectively, through LC-MS/MS and 1D/2D nuclear magnetic resonance (NMR) analyses. (Supplementary figure 1-12, Supplementary table 1 and 2). The identification was further corroborated by comparing the NMR chemical shifts to literature values [26–28]. Both MIAs showed UV absorption maxima shifted to longer wavelengths (302 nm for apparicine, and 316 nm for vobasine) compared to the typical indole UV absorption maxima at 280-290 nm (Fig. 3B and Supplementary fig. 1), consistent with their extended indole conjugation (Fig. 2). Additionally, we purified and identified two diastereomers (*m/z* 355), tabernaemontanine and dregamine (Fig. 2 and 3, Supplementary figure 13-22, Supplementary table 1 and 2), derived from vobasine 19,20-reduction. The disappearance of 19,20-alkene signals in both ^1^H and ^13^C NMR spectra was consistent with their 19,20-reduction, and Nuclear Overhauser Effect Spectroscopy (NOSEY) cross peaks clearly differentiated the two diastereomers. Similar to vobasine and perivine, both tabernaemontanine and dregamine showed UV absorption profiles and MS/MS fingerprints indicative of the unique indole-C3-ketone chromophore (Supplementary fig. 1). There are several bisindole alkaloids found only in young roots, which we tentatively identified as 3’-methoxytabernaelegantine A and C (*m/z* 737), and 3’-methoxyconodurine (*m/z* 735) based on *m/z* values (Fig. 2 and 3A). Structural elucidation of these bis-MIAs awaits additional NMR studies.

### γTMT-like methyltransferases identified in *T. elegans* comprise a novel clade that is distantly related to characterized MIA NMTs

To identify the enzymes for vobasine biosynthesis, we sequenced *T. elegans* leaf and root total RNA and successfully identified many putative vobasine biosynthetic enzymes using known MIA biosynthetic enzymes (Table 1). For the last step of vobasine biosynthesis, we searched the transcriptomes with six characterized MIA NMT protein sequences. Given the substantial accumulation of vobasine as a major MIA in *T. elegans* leaf, we expected the putative MT to be highly expressed in this tissue. From all hits, we selected the top three highest-expressed γTMT-like methyltransferases (MTs), namely TeMT1-3 (Genbank PP067959-PP067961), in descending order of expression levels in leaf tissue, for further investigation (Table 1). TeMT1 was the predominant MT in both leaf and root, showing transcripts per million (TPM) values 7.7 and 26.3 folds of those for TeMT2 and 3 (Table 1). In addition, TeMT1’s leaf TPM (401.8) was comparable to other vobasine biosynthetic genes such as TeSBE (450.2) and TeGS (173.9). TeMT2 showed over 80% amino acid sequence identity to various *bona fide* γTMTs, suggesting its potential status as a γTMT. The low expression of other γTMT-like transcripts preclude them from further analyses.

**Table 1.** The transcripts per million (TPM) values and sequence alignment results of TePeNMT, TeMT2/3, and ten other vobasine biosynthetic genes in *Tabernaemontana elegans* leaf and root tissues.

| enzyme | E-value | Amino acid identity (%) | Accession (E-value) | Leaf TPM | Root TPM |
| --- | --- | --- | --- | --- | --- |
| Te7DLS | 0 | 94.3 | Cr7DLS | 85.6 | 1315.0 |
| Te7DLH | 0 | 89.8 | Cr7DLH | 14.4 | 324.7 |
| Te7DLGT | 0 | 90.2 | Cr7DLGT | 19.0 | 233.7 |
| TeLAMT | 0 | 83.6 | CrLAMT | 86.6 | 310.9 |
| TeSLS | 0 | 91.0 | CrSLS | 117.3 | 1450.0 |
| TeSTR | 2.2E-163 | 72.3 | CrSTR | 20.6 | 185.4 |
| TeSGD | 0 | 70.8 | CrSGD | 35.3 | 148.8 |
| TeGS | 0 | 88.7 | CrGS | 173.9 | 1430.5 |
| TeSBE | 0 | 83.4 | RsSBE | 450.2 | 1055.4 |
| TeGO | 0 | 92.5 | CrGO | 369.9 | 1080.6 |
| TePeNMT(TeMT1) | 1.7E-104 | 52.3 | RsPiNMT | 401.8 | 574.9 |
| TeMT2 | 2.9E-110 | 53.9 | RsPiNMT | 52.5 | 20.0 |
| TeMT3 | 2.7E-109 | 52.3 | RsPiNMT | 15.3 | 77.9 |

Interestingly, both TeMT1 and 3 showed only 50% amino acid identity to the characterized CrPeNMT. In a phylogenetic analysis, TeMT2 clustered with other characterized γTMTs, while TeMT1 and 3 formed a distinct clade from both γTMTs and all other characterized MIA NMTs (Fig. 1). Notably, these γTMT-like MTs were also distinguishable from BIA NMTs and MIA *O*-methyltransferases (OMT). In contrast, eight other vobasine biosynthetic enzymes are highly conserved (83-94% amino acid sequence identity) between *T. elegans* and two other well-studied MIA-producing species, *C. roseus* and *Rauwolfia serpentina* (Indian snakeroot) (Table 1). Notable exceptions included the putative strictosidine synthase TeSTR and putative strictosidine β-glucosidase TeSGD, which were 72% and 71% identity to *C. roseus* STR and SGD at amino acid level, respectively (Table 1). It is also worth noting that TeMT1-3 lacked chloroplast transit peptides found in *bona fide* γTMT, as predicted by TargetP 2.0 (https://services.healthtech.dtu.dk/services/TargetP-2.0/).

### TeMT1 is the perivine *N*β-methyltransferase in *T. elegans*

To study their functions, we expressed *N*-terminal His-tagged TeMT1-3 along with the previously characterized CrPeNMT in *E. coli*. Additionally, we expressed the closest TeMT1 ortholog CrNMT-ADP00411 (Genbank ADP00411) from *C. roseus* in *E. coli* for comparison. This ortholog shared 61% amino acid identity, and clustered with TeMT1 and 3 in the phylogenetic analysis, suggesting a common evolution ancestry (Fig. 1). Subsequently, we supplied perivine to cultures expressing these proteins. Consistent with previous research, CrPeNMT methylated perivine to vobasine, which co-eluted with purified vobasine standard (Fig. 4A). TeMT1 also catalyzed perivine *N*β-methylation (Fig. 4A). In contrast, TeMT2 did not show detectable perivine methylation, while TeMT3 and CrNMT-ADP00411 showed negligible perivine NMT activity (Fig. 4A). Based on these results, we designated TeMT1 as *T. elegans* perivine *N*β-methyltransferase (TePeNMT). None of TeMT1-3 showed activity with 22 other MIAs encompassing multiple MIA skeleton families (Supplementary table 3),

**Figure 4.**
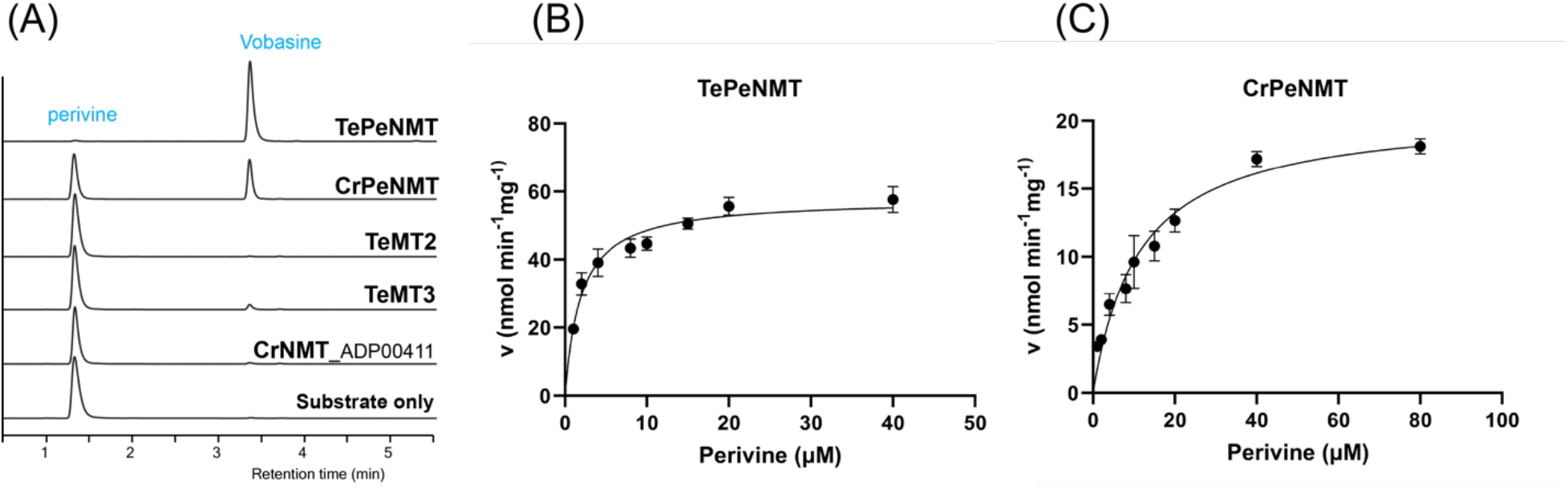
In vivo and in vitro biochemical characterizations identified TeMT1 as the perivine *N*β-methyltransferase in *Tabernaemontana elegans*. (A) *E. coli* cultures expressing CrPeNMT and TeMT1 both methylate the substrate perivine to vobasine as shown by LC-MS/MS multiple reaction monitoring (MRM) with [M+H]^+^ *m/z* 339>166 for perivine and *m/z* 353>180 for vobasine. The MS/MS fingerprints used for selecting these parameters are included in Supplementary Fig. 1. (B) and (C) The Michaelis-Menten saturation kinetics experiments for perivine substrate show superior kinetics for TePeNMT compared to CrPeNMT. Each data point shows the mean value of three technical replicates. The error bars indicate standard deviation.The curves are graphed with Prism Graphpad 9.5.0. The kinetics data are included in Table 2.

Following purification of recombinant TePeNMT and CrPeNMT through standard affinity chromatography (Supplementary fig. 25), we conducted in vitro enzyme kinetics assays using saturating co-substrate *S*-adenosylmethionine (SAM) and various perivine concentrations for comparison. Both recombinant enzymes methylated perivine substrate; however, TePeNMT showed superior enzyme kinetics (Fig. 4B, 4C and Table 2). Specifically, CrPeNMT displayed a *K*_M_ value for perivine 5.9 times greater than that of TePeNMT, indicating significantly lower binding affinity towards perivine compared to TePeNMT. Moreover, TePeNMT exhibited a *V*_max_ 2.8 times higher than that of CrPeNMT. Consequently, the catalytic efficiency (*k*_cat_/*K*_M_) of TePeNMT, calculated from *V*_max_/*K*_M_, surpassed that of CrPeNMT by a factor of 16.6.

**Table 2.** Michaelis-Menten saturation kinetics for TePeNMT and CrPeNMT.

|  | TePeNMT | CrPeNMT |
| --- | --- | --- |
| $K_M$ for perivine ($\mu\text{M}$) | 1.91 (95% CI<sup>a</sup>: 1.47-2.47) | 11.32 (95% CI: 8.58-14.78) |
| $V_{\max}$ (nmol mg protein<sup>-1</sup> min<sup>-1</sup>) | 57.83 (95% CI: 54.75-61.18) | 20.70 (95% CI: 18.83-22.82) |

<sup>a</sup> confidence interval

### Homology modeling indicates significant disparities in both active site conformation and perivine substrate docking positions between TePeNMT and CrPeNMT

After observing the low identity in enzyme primary sequences, conducting phylogenetic analysis, and noting the kinetic differences, we further continued to investigate the tertiary structures of TePeNMT, CrPeNMT, TeMT3, and CrNMT-ADP00411 for their catalytic mechanisms. Using the *Arabidopsis thaliana* phosphoethanolamine NMT (PDB: 5WP5) as template, we built homology models for all four enzymes with the Molecular Operating Environment (MOE). While all NMTs shared a high degree of tertiary structure similarity, we found significant differences in their active sites, particularly in accommodating perivine binding and its *N*β-methylation via S_N_2 displacement of the methylsulfonium group of SAM (Fig. 5A and B, Supplementary fig.26).

**Figure 5.**
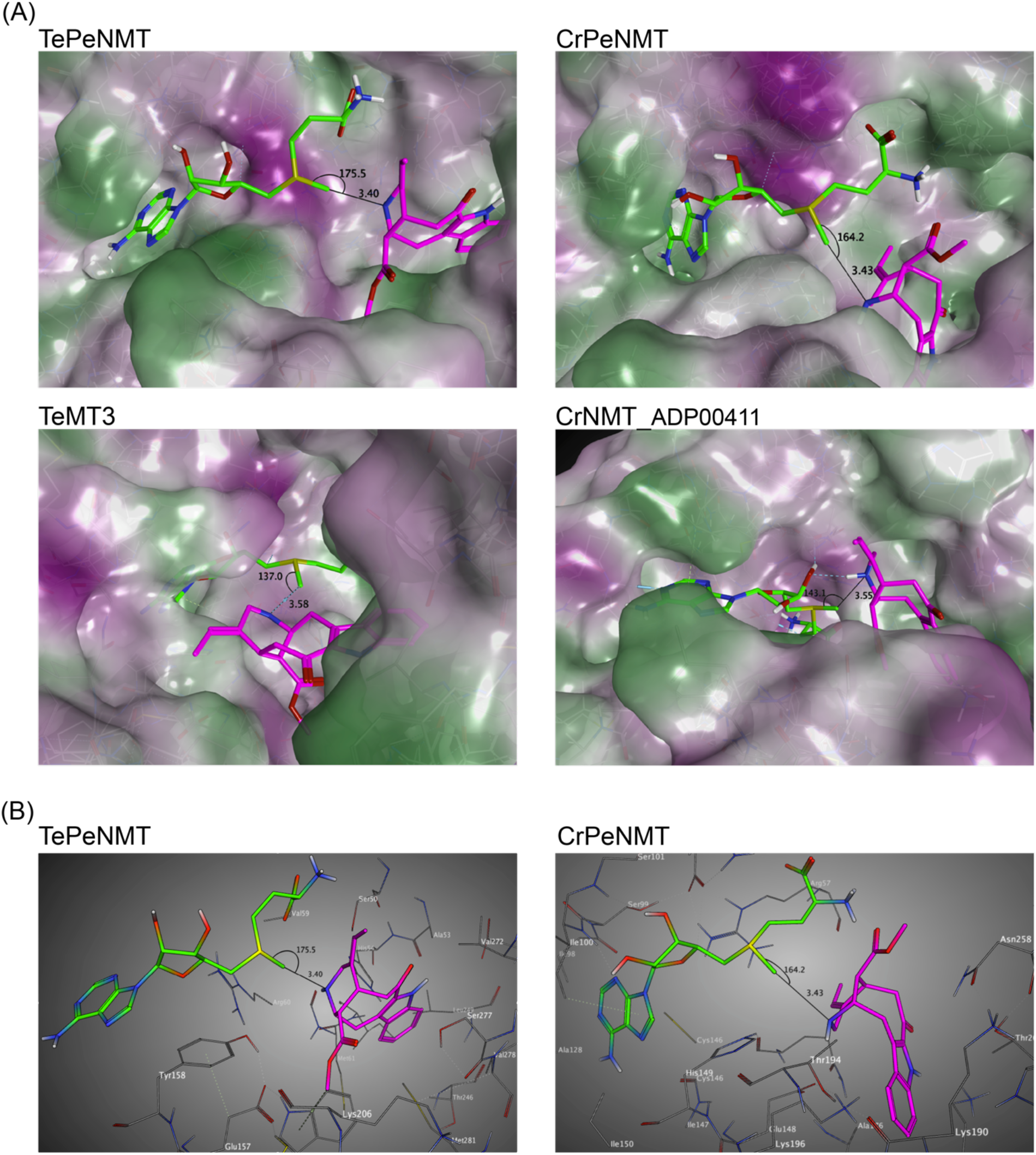
Homology modeling supports that TePeNMT’s superior enzyme efficiency is contributed by significant differences in enzyme active sites and perivine binding poses between TePeNMT and CrPeNMT. Enzyme models are generated from the template *Arabidopsis thaliana* phosphoethanolamine NMT-2 (PDB: 5WP5.A) with MOE version 2022.02. *S*-adenosylmethionine (SAM) is shown in green, and perivine is shown in magenta. The distance and angel between SAM methylsulfonium donor and perivine-*N*β are illustrated. (A) the lipophilic mapping of four NMT enzymes, in which polar regions are shaded purple and lipophilic regions are shaded green. (B) The ball-and-stick diagrams of the same enzyme active sites for TePeNMT and CrPeNMT.

For all NMTs, the SAM co-substrate was coordinated in the conserved glycine-rich region via multiple hydrogen bonds and hydrophobic interactions (Supplementary fig.26). In TePeNMT, perivine was docked in a hydrophobic pocket primarily via van der Waals interactions with residues Trp184, Ser50/277, Val59/272, and His54 (Fig. 5B and Supplementary fig. 26). The distance between SAM’s methyl group and the perivine *N*β nitrogen (**N**…**C**H_3_, 3.40 Å) and the optimal angle between the methyl and the *N*β nitrogen (**N**…**C**H_3_–**S**, 175.5°) is compatible for efficient S_N_2 displacement (Fig. 5A and B). In comparison, perivine rotated over 90° in the active site of CrPeNMT, with the indole portion situated in a deeper binding pocket. The active site consisted of residues distinct from those found in TePeNMT. Specifically, perivine was coordinated with Asp258, Lys190/196/271, Thr262, Ile198, and Arg57 (Fig. 5B and Supplementary Fig. 26). This different docking pose of perivine altered both the angle between SAM’s methyl group and the *N*β nitrogen (164.2°) and their distance (3.43 Å), likely contributing to reduced *K*_M_ and *V*_max_ for CrPeNMT. Conversely, perivine docking position was highly suboptimal for both TeMT3 and CrNMT-ADP00411 (Fig. 5A). The angles between SAM’s methyl group and the perivine *N*β nitrogen (137.0 ° and 143.1°) were not conducive for productive S_N_2 methylation, aligning with their negligible perivine NMT activity.

## Discussion

Monoterpenoid indole alkaloids (MIAs) stand as a fascinating alkaloid class found primarily in the families Apocynaceae, Loganiaceae, and Rubiaceae. Noted for their structural diversity and potent biological activities, an essential objective in this field of research is uncovering the biosynthetic pathways involved in the construction of these highly complex compounds. Notably, many of these biologically active molecules are either *N*-methylated themselves or are *N*-methylated as precursors in MIA biosynthesis. Methylation alters MIAs’ solubility, electronic property, reactivity, and binding capacity to biological targets, thus providing a direct pathway to new bioactivities through a simple chemical transformation.

The characterization of *C. roseus* DhtNMT, a γ-tocopherol *C*-methyltransferase like enzyme involved in vindoline biosynthesis [3], has been a focal point in initiating the study of this interesting methyltransferase class of γ-TMT ancestry. All MIA NMTs known to date derive from γ-TMT, a lineage that includes VmPiNMT, RsPiNMT, RsNNMT, RsANMT, and CrPeNMT. The studies on these enzymes have not only elucidated their substrate specificity but also shed light on their altered subcellular localization, providing valuable clues about their metabolic roles within plant cells [10].

By investigating the metabolomes and transcriptomes of various *T. elegans* tissues, we identified a new γ-TMT-like enzyme TePeNMT, which functions as a perivine *N*β-methyltransferase. Its abundant expression in *T. elegans* leaf correlates with the high accumulation of vobasine observed in the same tissue (Fig. 3, Table 1). Interestingly, while CrPeNMT has been previously characterized in *C. roseus* [8], this plant species is not known for the accumulation of vobasine or its further oxidized form, periformyline (*N*β-formylperivine), which have been documented in other Catharanthus species *C. trichophyllus* and *C. lanceus* [17,18]. Our previous research has also demonstrated that perivine, rather than vobasine or periformyline, accumulates in the leaf tissue of *C. roseus* [19]. Perhaps, a lower level of CrPeNMT or the cellular and subcellular compartmentalization of CrPeNMT and substrate perivine, could be a reason why perivine remains non-methylated in *C. roseus*. The physiological role of CrPeNMT remains to be elucidated.

Our phylogenetic analysis revealed a distinct evolutionary path for TePeNMT, setting it apart from all other characterized MIA NMTs (Fig. 1). Interestingly, TePeNMT exhibits a closer relationship with bona fide γ-TMTs found in both MIA and non-MIA producing species, such as *Arabidopsis thaliana*. Notably, an ortholog of TePeNMT in *C. roseus*, designated as CrNMT-ADP00411 and sharing 61% amino acid sequence identity, was identified in our phylogenetic study. However, this ortholog lacks perivine NMT activity, indicating that TePeNMT evolved specifically in *T. elegans* to catalyze perivine *N*β-methylation. TePeNMT and CrPeNMT only share 50% amino acid identity. The fact that eight other vobasine biosynthetic enzymes encompassing key enzymes such as geissoschizine synthase (GS) and sarpagan bridge enzyme (SBE) are highly conserved (83%-94% identity) between *T. elegans* and *C. roseus* (Table 1) further supports that the two PeNMTs evolved independently in these two species.

Moreover, our study on enzyme kinetics and homology modeling revealed significant catalytic disparities between TePeNMT and CrPeNMT (Fig. 5). The low amino acid identity (50%) resulted in drastic differences in their active sites for vobasine binding. They barely shared any conserved amino acids, further supporting an independent evolution path that shaped the two perivine binding pockets. This variation in binding poses of perivine and SAM results in a superior disposition of the methylsulfonium donor and perivine *N*β in TePeNMT, facilitating a more efficient S_N_2 reaction. The findings from homology modeling corroborate the kinetics data for these enzymes, wherein TePeNMT displayed superior performance over CrPeNMT in terms of both substrate affinity *K*_M_ and enzyme velocity *V*_max_ (Table 2).

Vobasine serves as a precursor to the potent anticancer drug (3’*R*)-hydroxytabernaelegantine C and stands as the most abundant MIA in *T. elegans* (Fig. 2 and 3). While it is unclear for the *in planta* role of vobasine and its derivatives tabernaemontanine and dregamine identified in this study, TePeNMT likely coevolved with the occurrence of vobasine, potentially contributing to the improved enzyme efficiency and fitness of *T. elegans*. Our phytochemical, phylogenetic, kinetic, and homology modeling data collectively suggested distinct evolutionary paths for the two PeNMTs from *T. elegans* and *C. roseus*. This presents another notable example of enzyme parallel evolution in the plant kingdom.

## Material and Methods

### Plant materials and alkaloid purification

Fresh leaves and stems (600 g total) of glasshouse grown *T. elegans* were submerged in methanol for a week. The extract was evaporated under vacuum and resuspended in 10 mL 1 M HCl. The suspension was extracted with ethyl acetate, and the aqueous phase was basified using NaOH to above pH 7. After extraction by ethyl acetate, the total alkaloids in organic phase were evaporated and dissolved in methanol. Total alkaloids were separated by thin layer chromatography (Sicica gel60 F254, Millipore Sigma, Rockville, MD, USA) with solvents ethyl acetate and methanol (9:1, v/v). After extracting the TLC bands with methanol, 14 mg vobasine, 8 mg tabernaemontanine, 6 mg dregamine, and 5 mg apparicine were recovered. For tissue alkaloid comparison, 100 mg tissues of leaf, root and flower tissues were ground in 2 ml methanol to generate total extract. For latex, 100 mg freshly collected latex was mixed with 2 ml methanol. The samples were analyzed by LC-MS/MS.

### mRNA extraction and transcriptome assembly

*T. elegans* leaf and root tissues (100 mg) from glasshouse grown plants were ground into fine powder in liquid nitrogen. RNA were extracted using a Mornarch® RNA Cleanup Kit (New England Biolabs, Ipswich, MA, USA) according to the manufacture’s protocol. The RNA was sequenced using Illumina NovaSeq 25M reads at the Atlantic Cancer Research Institute (Moncton, NB, Canada). The assembled transcriptomes were analyzed using CLC Genomic Workbench 20.0.4 (Qiagen, Redwood City, CA, USA).

### LC-MS/MS and NMR

The samples were analyzed by Ultivo Triple Quadrupole LC-MS/MS system from Agilent (Santa Clara, CA, USA), equipped with an Avantor® ACE® UltraCore C18 2.5 Super C18 column (50×3 mm, particle size 2.5 *μ*g,) as well as a photodiode array detector and a mass spectrometer. For alkaloid analysis, the following solvent systems were used: Solvent A, Methanol: Acetonitrile: Ammonium acetate 1 M, water at 29:71:2:398; solvent B, Methanol: Acetonitrile: Ammonium acetate 1 M: water at 130:320:0.25:49.7. The following linear elution gradient was used: 0-5 min 80% A, 20% B; 5-5.8 min 1% A, 99% B; 5.8-8 min 80% A, 20% B; the flow during the analysis was constant and 0.6 ml/min. The photodiode array detector range was 200 to 500 nm. The mass spectrometer was operated with the gas temperature at 300°C and gas flow of 10 L/min. Capillary voltage was 4 kV from m/z 100 to m/z 1000 with scan time 500 ms and the fragmentor performed at 135 V with positive polarity. The MRM mode was operated as same as scan mode with the adjusted precursor and product ion. ^1^H (400 MHz) and ^13^C (100 MHz) NMR spectra were recorded on an Agilent 400 MR and a Bruker Avance III HD 400 MHz spectrometers in CDCl_3_.

### Protein expression and purification

All MT sequences were codon optimized, synthesized, and subcloned into pET30a(+) vector between NotI/XhoI restriction sites (Bio Basic Inc., Toronto, ON, Canada). CrPeNMT (Genbank KC708453) was amplified from *C. roseus* leaf cDNA using primers (5’-ATAGGATCCAATGGCCTCAATGGGAGAGAAGGA and 5’-ATAGTCGACTCATTTAGTTTTGCGAAATGTAACTG) and cloned in pET30b+ vector within BamHI/SalI sites. The clones were transformed and expressed in *Escherichia coli* strain BL21-(DE3). An overnight culture was used to inoculate 200 mL LB media, which was grown at 37°C to OD_600_ between 0.6-0.8. The cultures were then induced with IPTG at a final concentration of 0.1 mM and shifted to 15°C with shaking at 200 rpm. After approximately 12 hours of growth, the cells were harvested by centrifugation. The pellet cells were sonicated and resuspended in ice-cold lysis buffer (25 mM Tris-HCl at pH 7.5, 100 mM NaCl, 10% glycerol, and 20 mM imidazole) and lysed using sonication. The lysate was centrifuged at 4°C, 6,000 g for 25 minutes. The resulting supernatant was subjected to purification using standard Ni-NTA affinity chromatography. The column was washed twice with 3 mL of wash buffer (25 mM Tris-HCl at pH 7.5, 100 mM NaCl, 10% glycerol, and 30 mM imidazole) to rid of proteins with low binding affinities. Recombinant MT proteins were eluted using buffer containing imidazole at 250 mM and desalted in the storage buffer (25 mM Tris-HCl at pH 7.5, 100 mM NaCl, 10% glycerol) using a PD-10 Desalting column (Cytiva, Marlborough, MA, USA) and stored in -80°C until analysis.

### In vivo biotransformation and in vitro kinetics assay

For in vivo biotransformation experiment, an aliquot (1 mL) of induced culture was centrifuged to collect cell pellet, which was resuspended in 0.5 mL Tris-HCl at pH 7.5 supplemented with 2 *μ*g perivine or other alkaloid substrates. The biotransformation alkaloid substrates was carried out at 37°C, 200 rpm for 5 h. An aliquot of the biotransformation broth was mixed with equal volume of methanol and analyzed by LC-MS/MS. In vitro enzyme kinetics assays (50 *μ*l) contained 20 mM Tris pH 7.5, 3 *μ*g of the purified recombinant proteins, 50*μ*M *S*-adenosyl-methionine, and perivine concentrations ranging from 1 to 80 *μ*M. After adding the enzyme, the reaction was incubated in a 30 °C water bath for 5 min and subsequently terminated by adding 150 *μ*L of methanol. The product formation was analyzed by LC-MS/MS. Triplicated experiments were graphed using GraphPad Prism (9.5.0) (Boston, MA, USA).

### Homology modeling and substrate docking studies

All computational experiments and visualization were carried out with MOE version 2022.02 on local computers or on the Digital Research Alliance of Canada (DRAC, formerly Compute Canada) Advanced Research Computing Network (alliancecan.ca). All molecular mechanics calculations and simulations were conducted using the AMBER14:EHT forcefield with Reaction Field solvation. AM1-BCC charges were used for modeling the coenzyme and the substrates while AMBER charges were used for the enzymes. Sequence homology searches for both TePeNMT and CrPeNMT on the Protein DataBank found phosphoethanolamine *N*-methyltransferase 2 (AtPMT2) from *Arabidopsis thaliana* to be the best template for homology modeling (PDB ID: 5WP5) based on best Hidden Markov Model energy scores; *S*-adenosyl-homocysteine (SAH) was co-crystallized with the AtPMT2 enzyme. Sequence identity of TePeNMT and CrPeNMT to the relevant 5WP5 domain was 19.3%. and 19.2%, respectively. Homology models were then derived in MOE using default settings and scored using the GBVI/WSA dG method. SAM was modeled and docked into each homology model using the location of SAH in the 5WP5 enzyme as general docking region and default MOE-Dock settings. Perivine was then modeled and docked to the PeNMT-SAM complexes; for each perivine docking experiment, 100,000 docking poses were initially generated via the Triangle Matcher method and scored by the London dG function. A subset comprising the best docking poses were refined by the induced fit method where the bound ligands and active site residues were submitted to local geometry optimization and rescored by the GBVI/WSA dG scoring function. The top scoring docking poses that were geometrically compatible for for β*N* methylation of perivine by SAM via S_N_2 displacement were retained for subsequent analyses. Cartesian coordinates for the TePeNMT and CrPeNMT homology models and their perivine docked complexes can be found in the Supplemental Information.

## Acknowledgements

The authors thank the Chemical Computing Group ULC (www.chemcomp.com) for MOE licenses. This research was enabled in part by support provided by ACENET (www.ace-net.ca) and the Digital Research Alliance of Canada (alliancecan.ca).

