## Supplementary information for "Parallel evolution of methyltransferases leads to vobasine biosynthesis in *Tabernaemontana elegans* and *Catharanthus roseus*"

**Supplementary Figure 1. UV absorption and MS/MS profiles for MIAs**

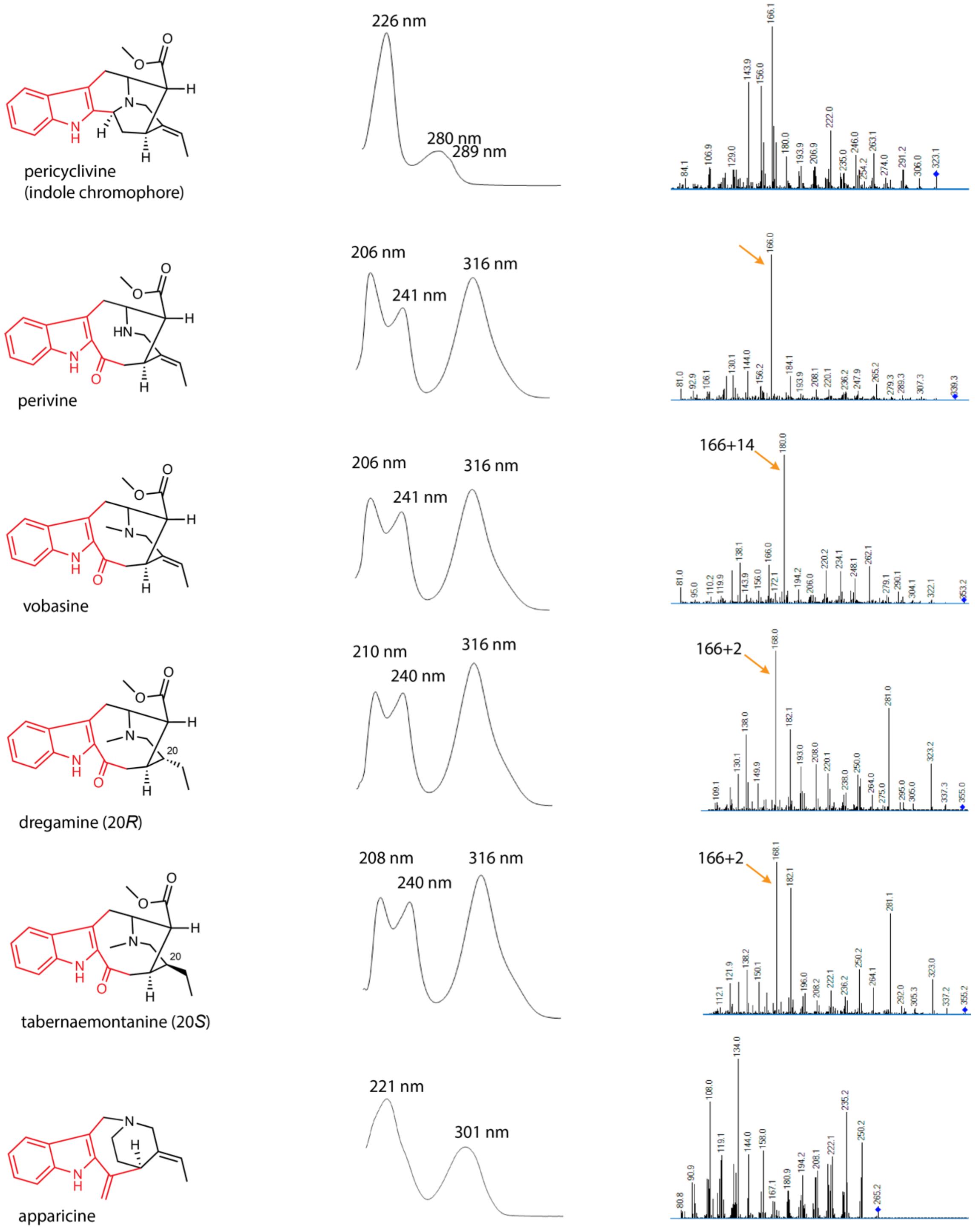

UV absorption and MS/MS profiles for pericyclivine (indole chromophore), perivine, vobasine, dregamine and tabernaemontanine, and apparicine. Chromophores are shown in red. The orange arrows indicate MS daughter ions +14 amu (methylation) from perivine for vobasine, and +2 from vobasine for dregamine and tabernaemontanine.

**Supplementary Figure 2. $^1$H NMR spectra for vobasine in $CDCl_3$.**

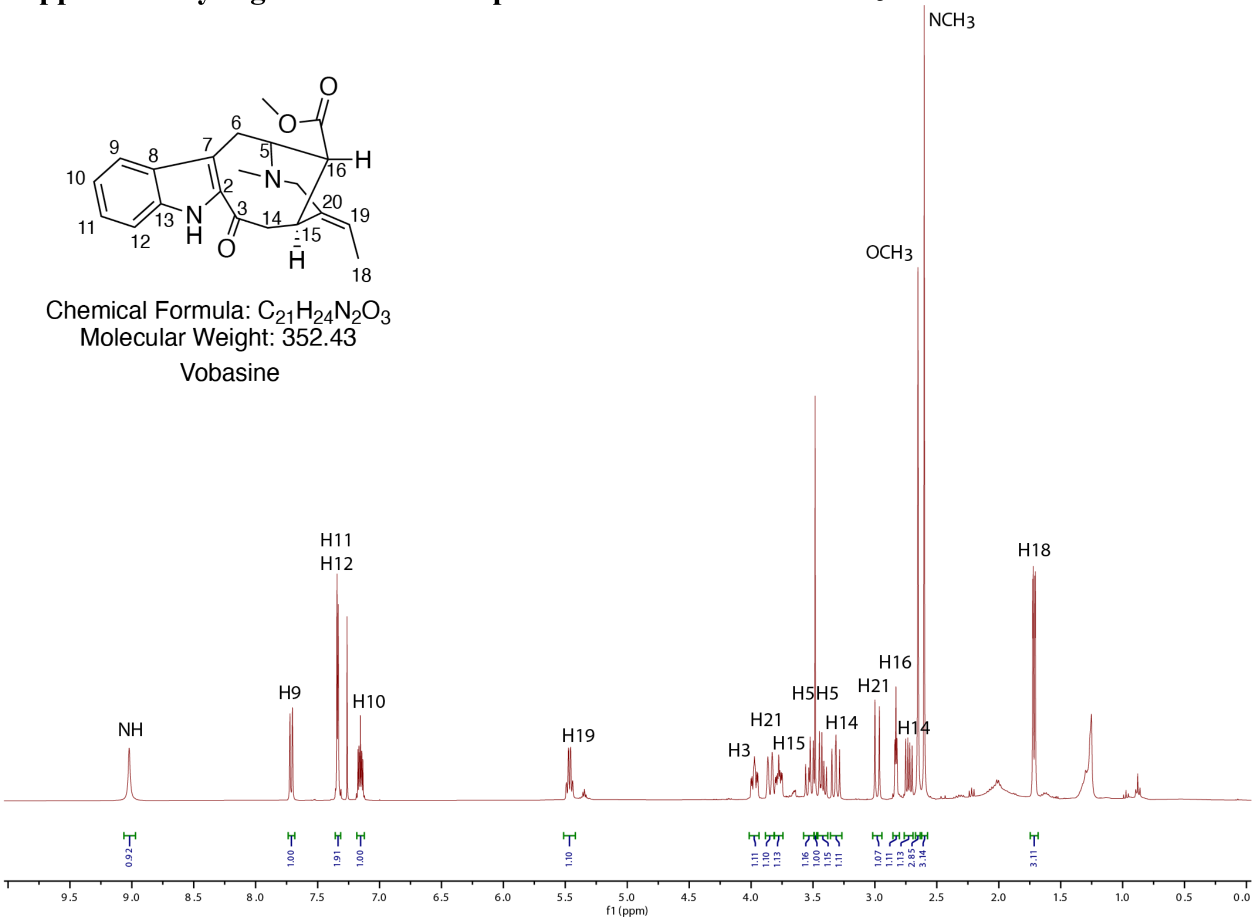

**Supplementary Figure 3. $^{13}$C NMR spectra for vobasine in $CDCl_3$.**

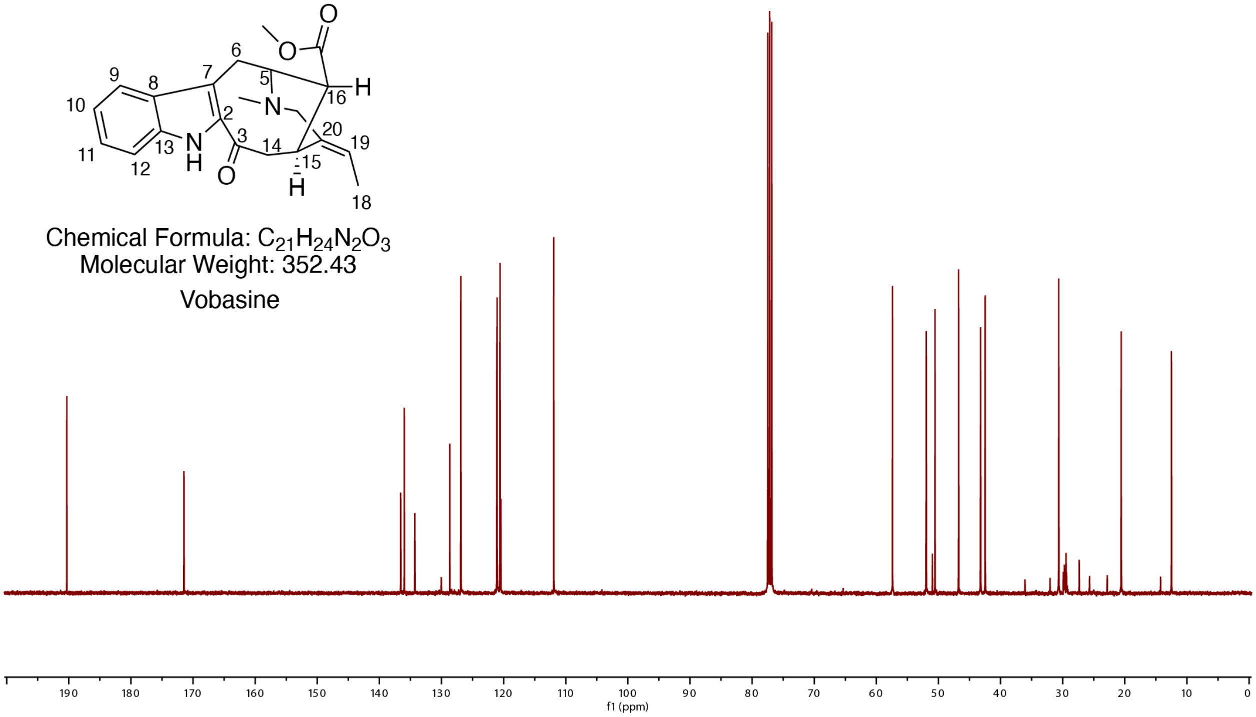

**Supplementary Figure 4. HSQC NMR spectra for vobasine in $CDCl_3$.**

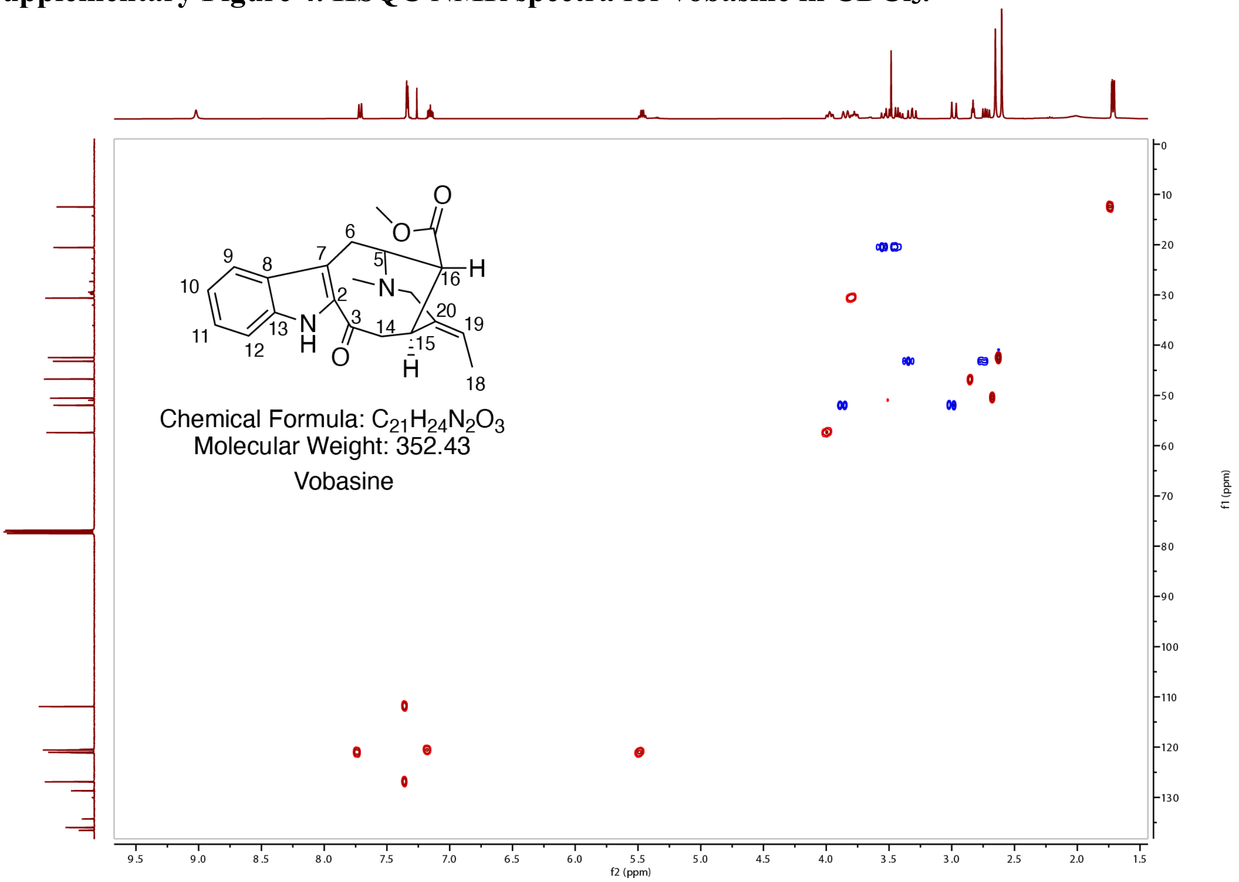

**Supplementary Figure 5. HMBC NMR spectra for vobasine in CDCl₃.**

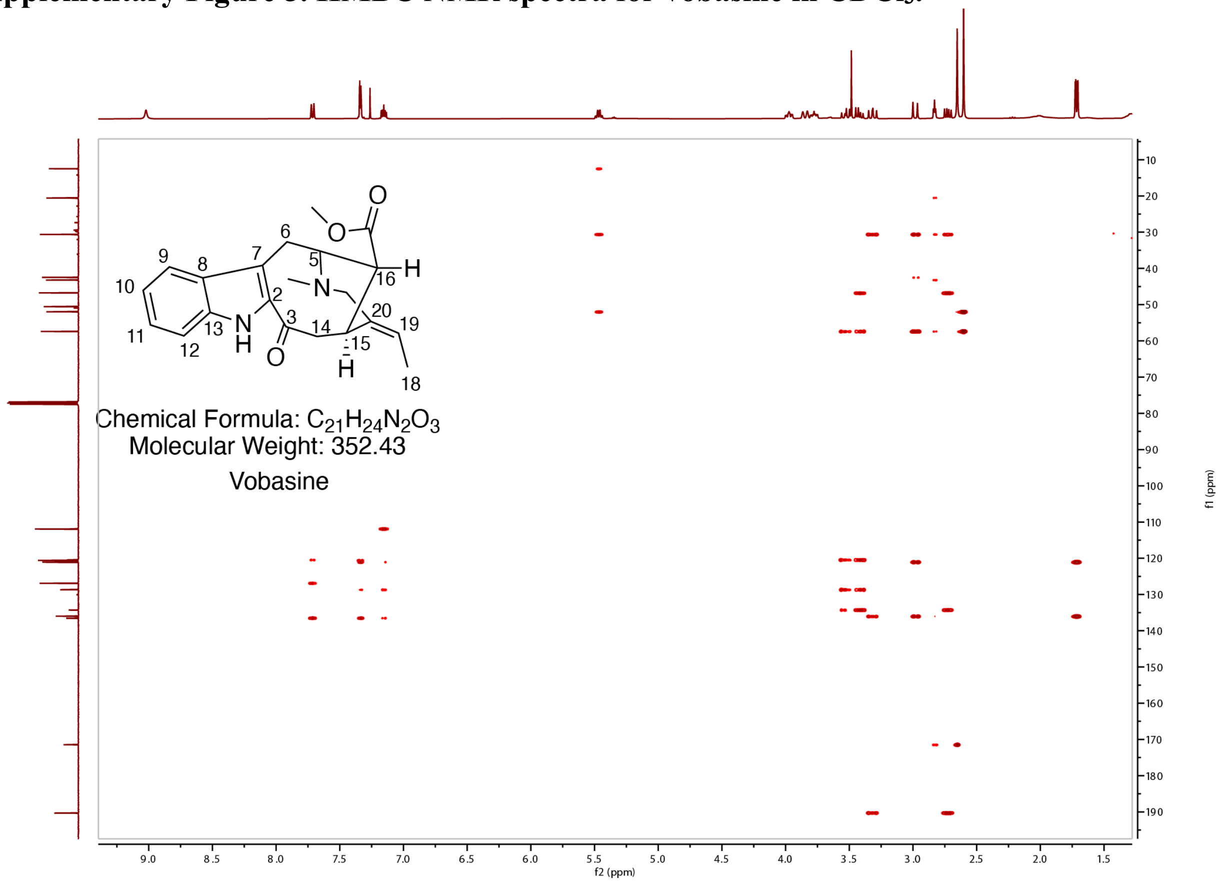

**Supplementary Figure 6. COSY NMR spectra for vobasine in $CDCl_3$.**

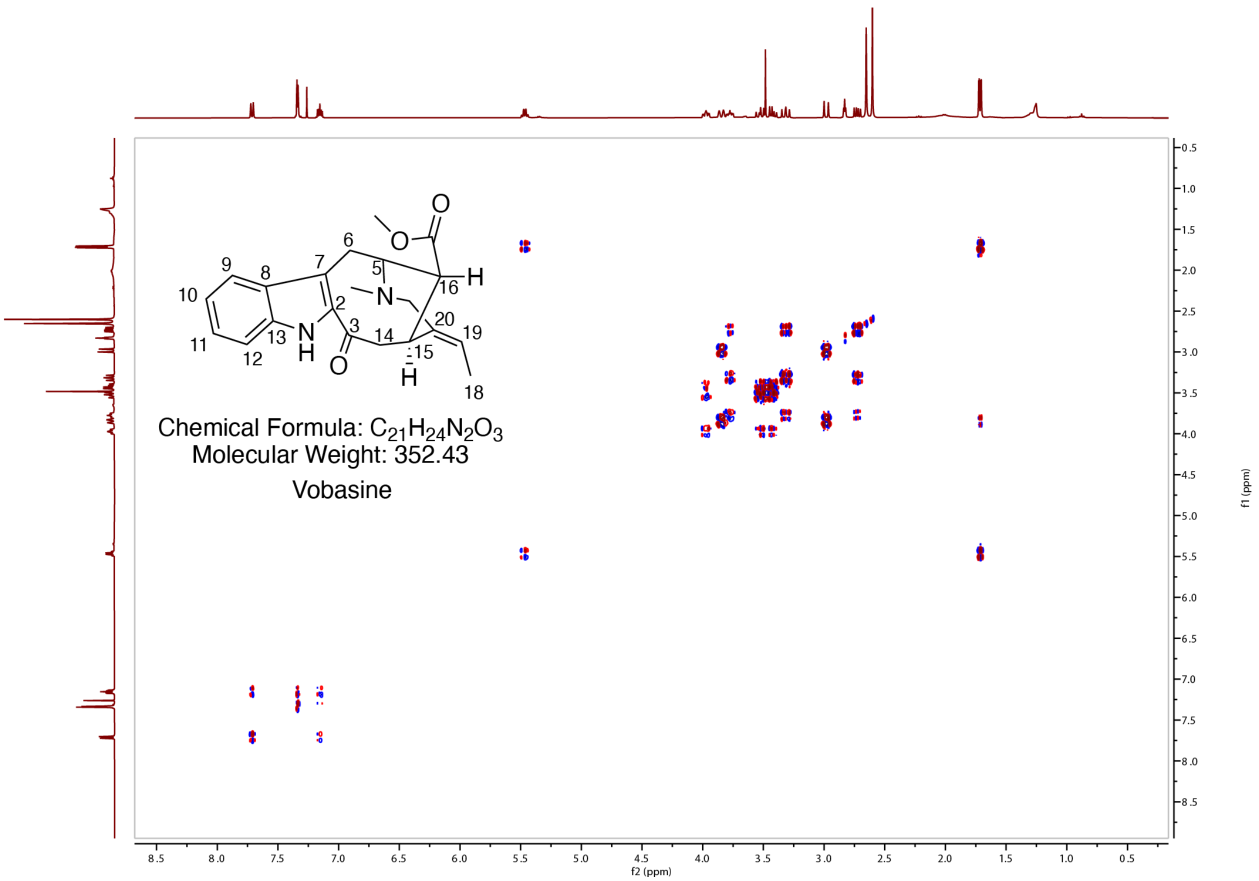

**Supplementary Figure 7. NOESY NMR spectra for vobasine in $CDCl_3$.**

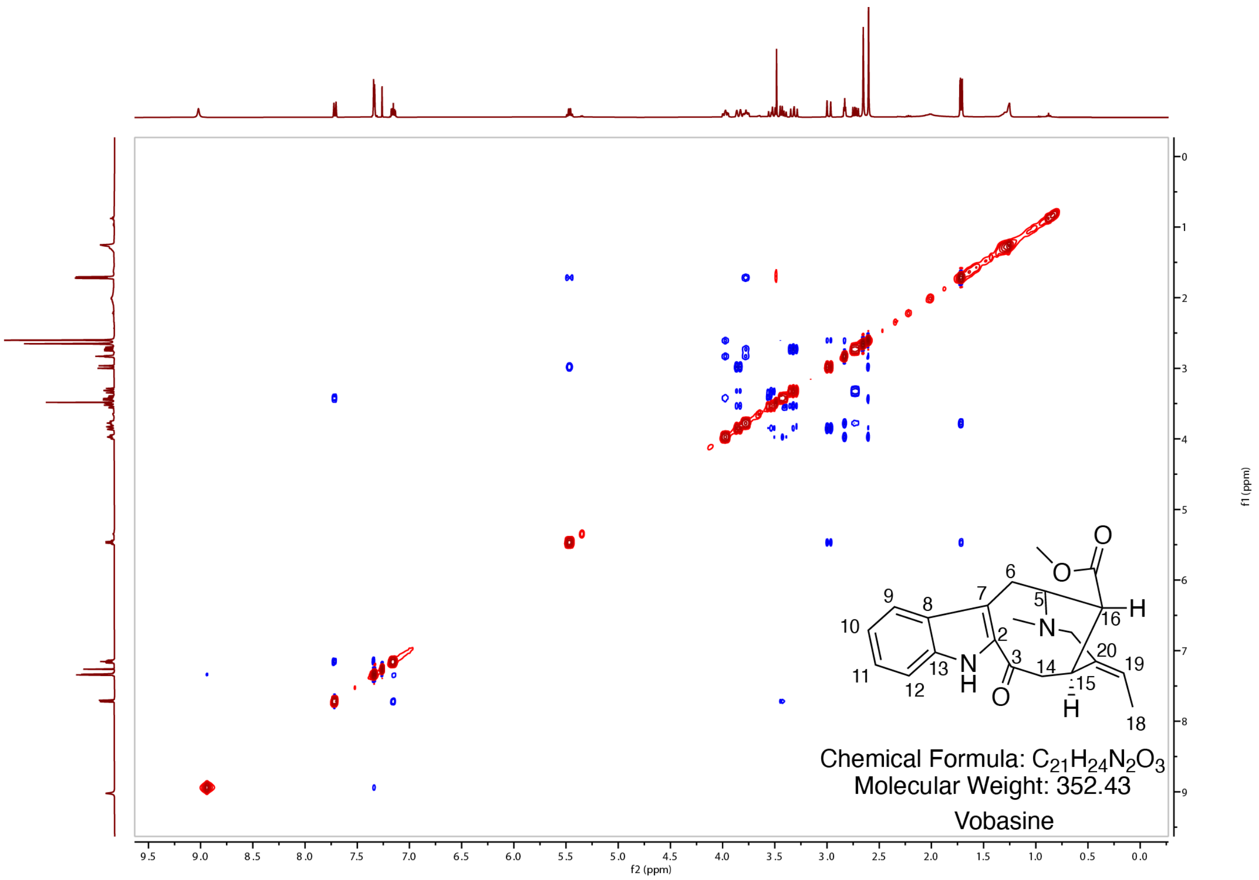

**Supplementary Figure 8. $^1$H NMR spectra for apparicine in $CDCl_3$.**

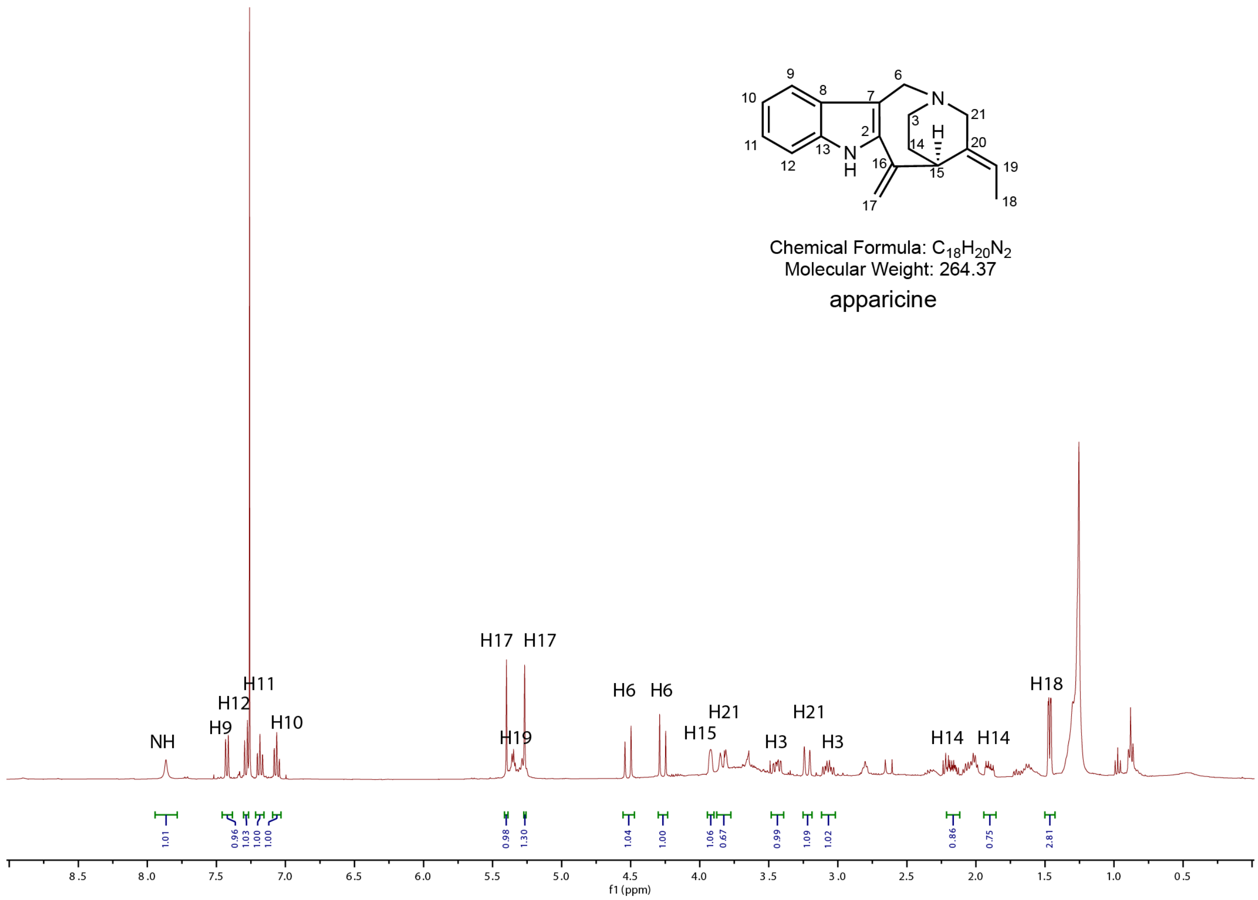

**Supplementary Figure 9. $^{13}$C NMR spectra for apparicine in CDCl$_3$.**

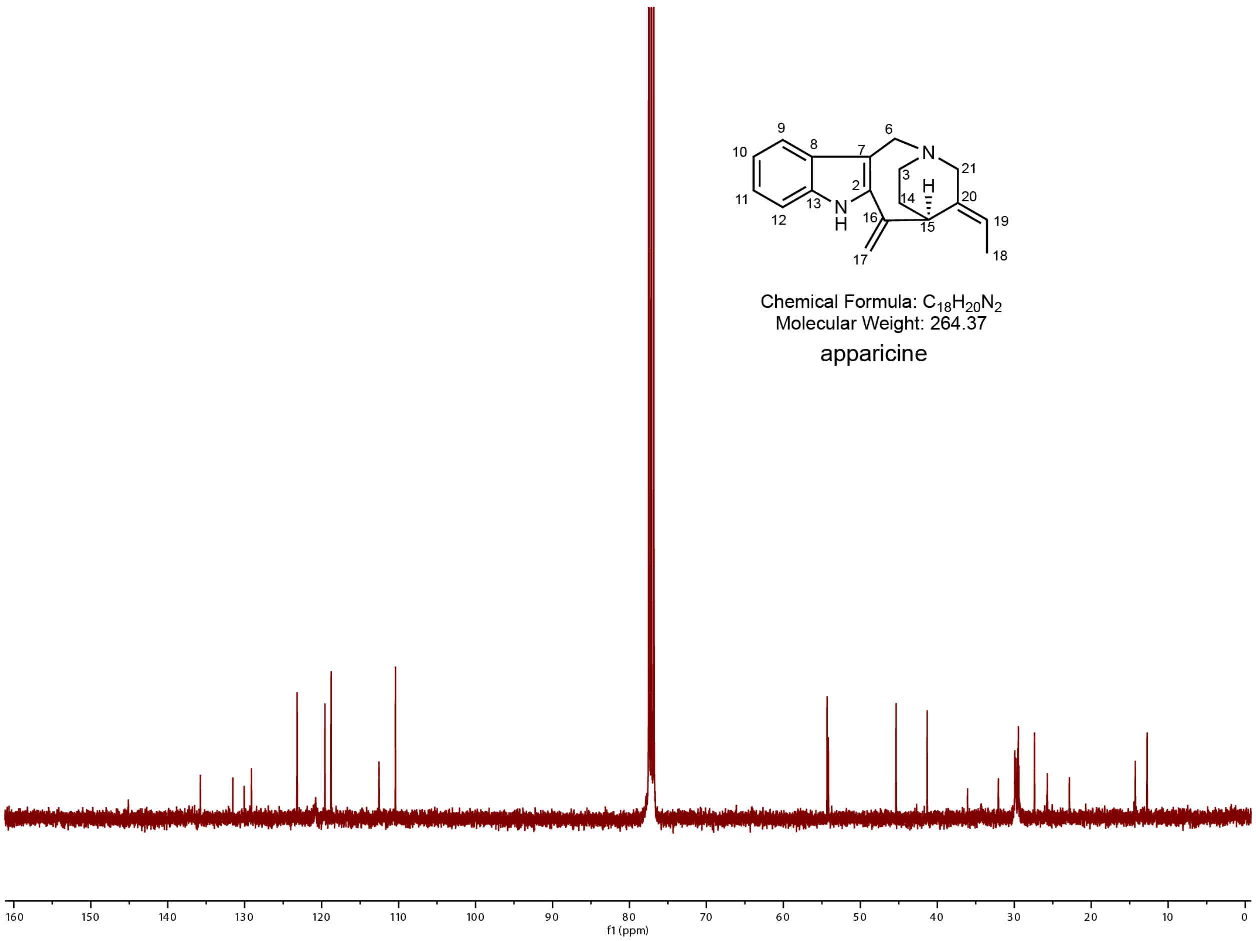

**Supplementary Figure 10. HSQC NMR spectra for apparicine in $CDCl_3$.**

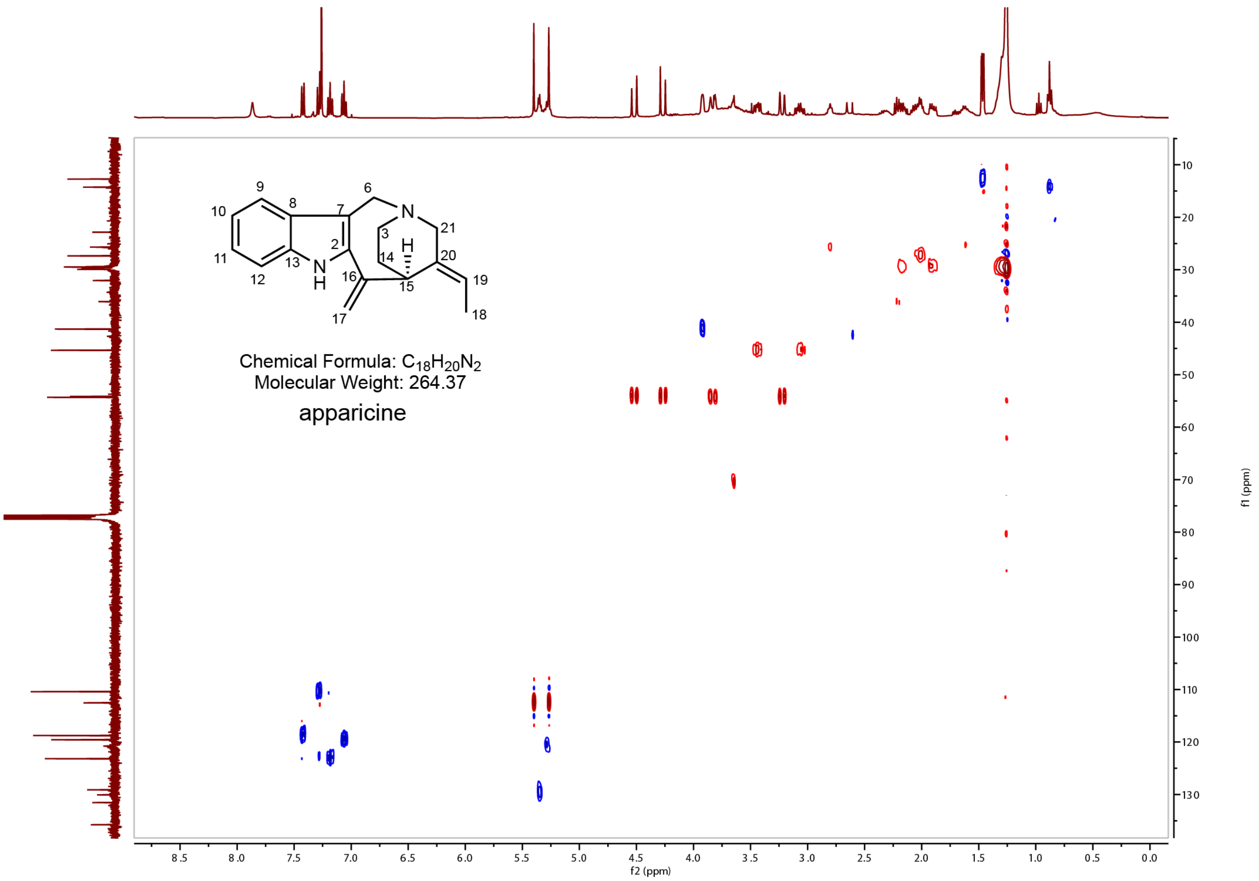

**Supplementary Figure 11. HMBC NMR spectra for apparicine in $CDCl_3$.**

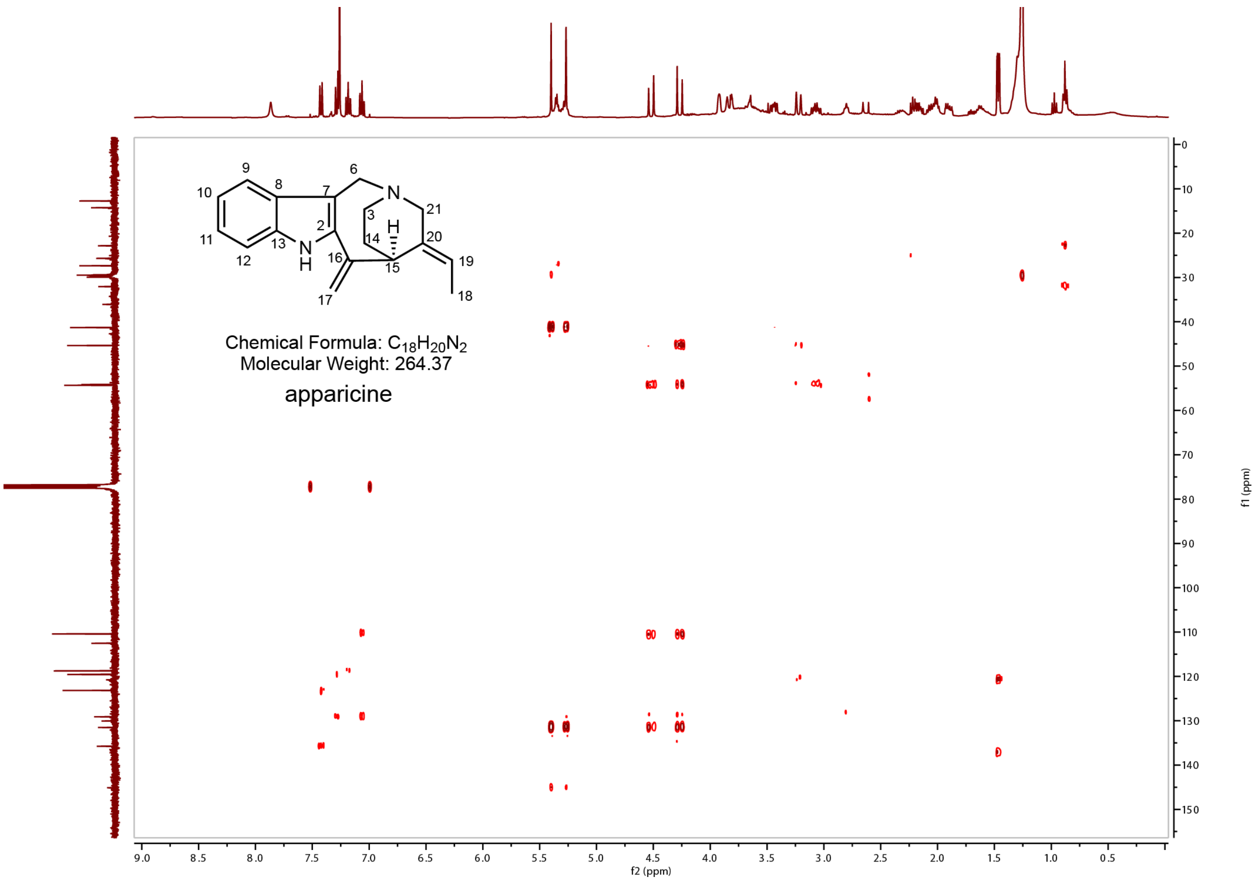

**Supplementary Figure 12. COSY NMR spectra for apparicine in $CDCl_3$.**

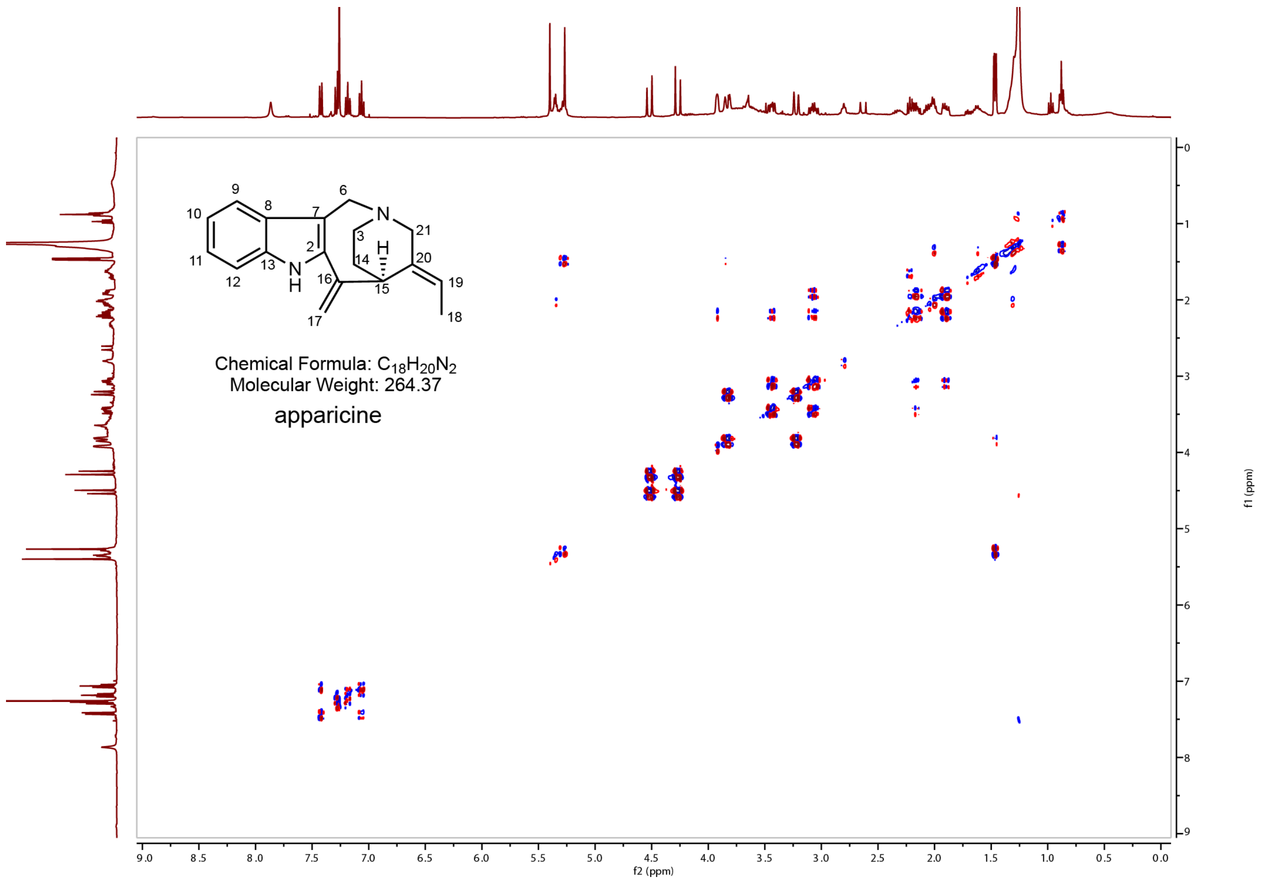

**Supplementary Figure 13. $^1$H NMR spectra for dregamine in $CDCl_3$.**

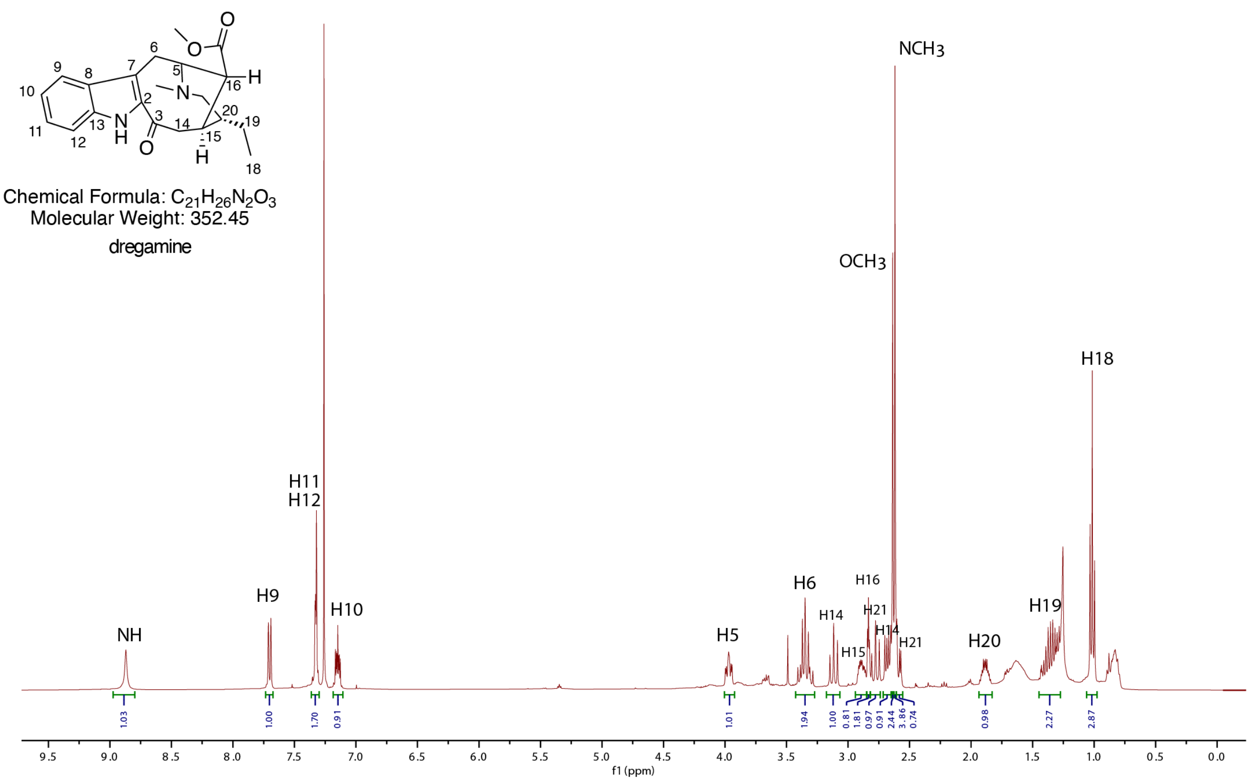

**Supplementary Figure 14. $^{13}$C NMR spectra for dregamine in $CDCl_3$.**

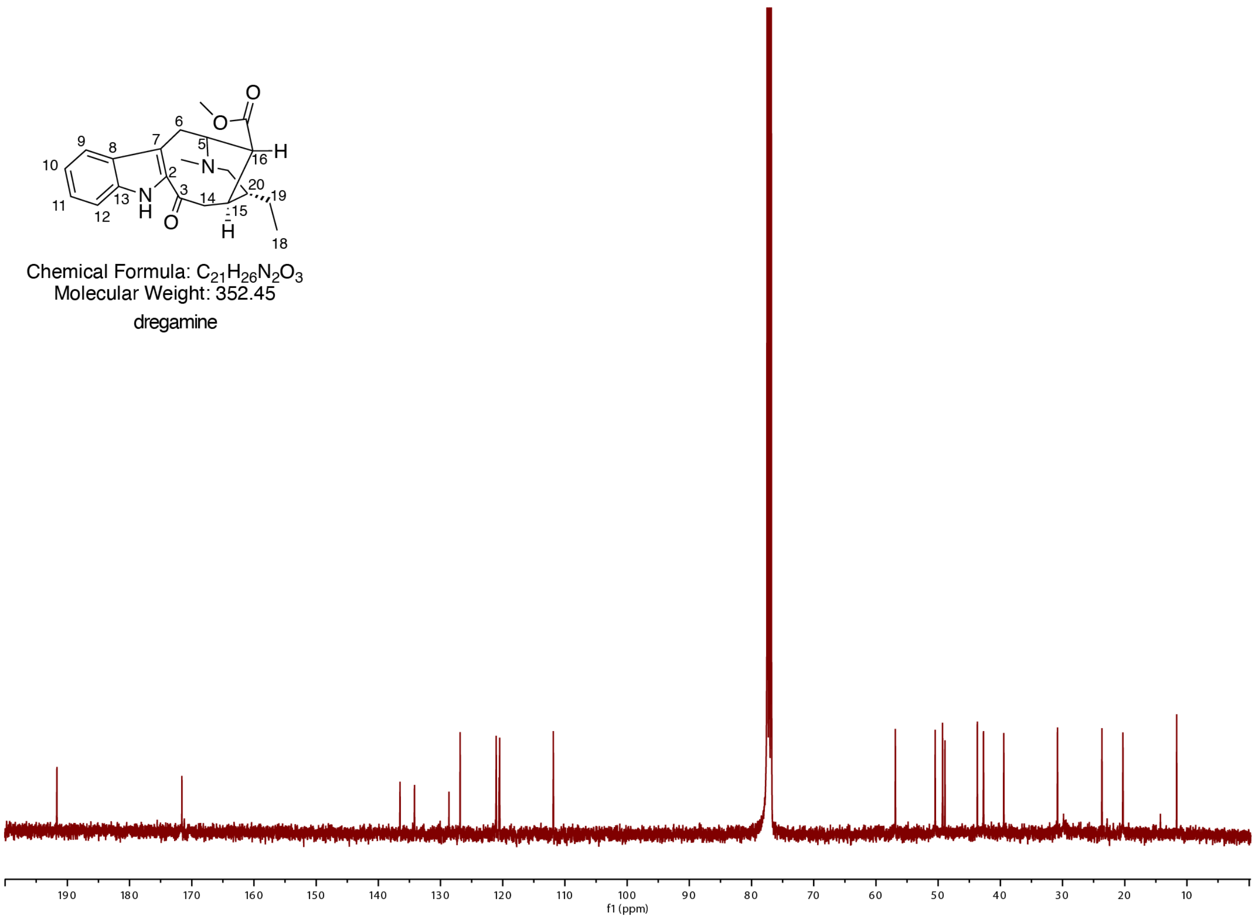

**Supplementary Figure 15. HSQC NMR spectra for dregamine in $CDCl_3$.**

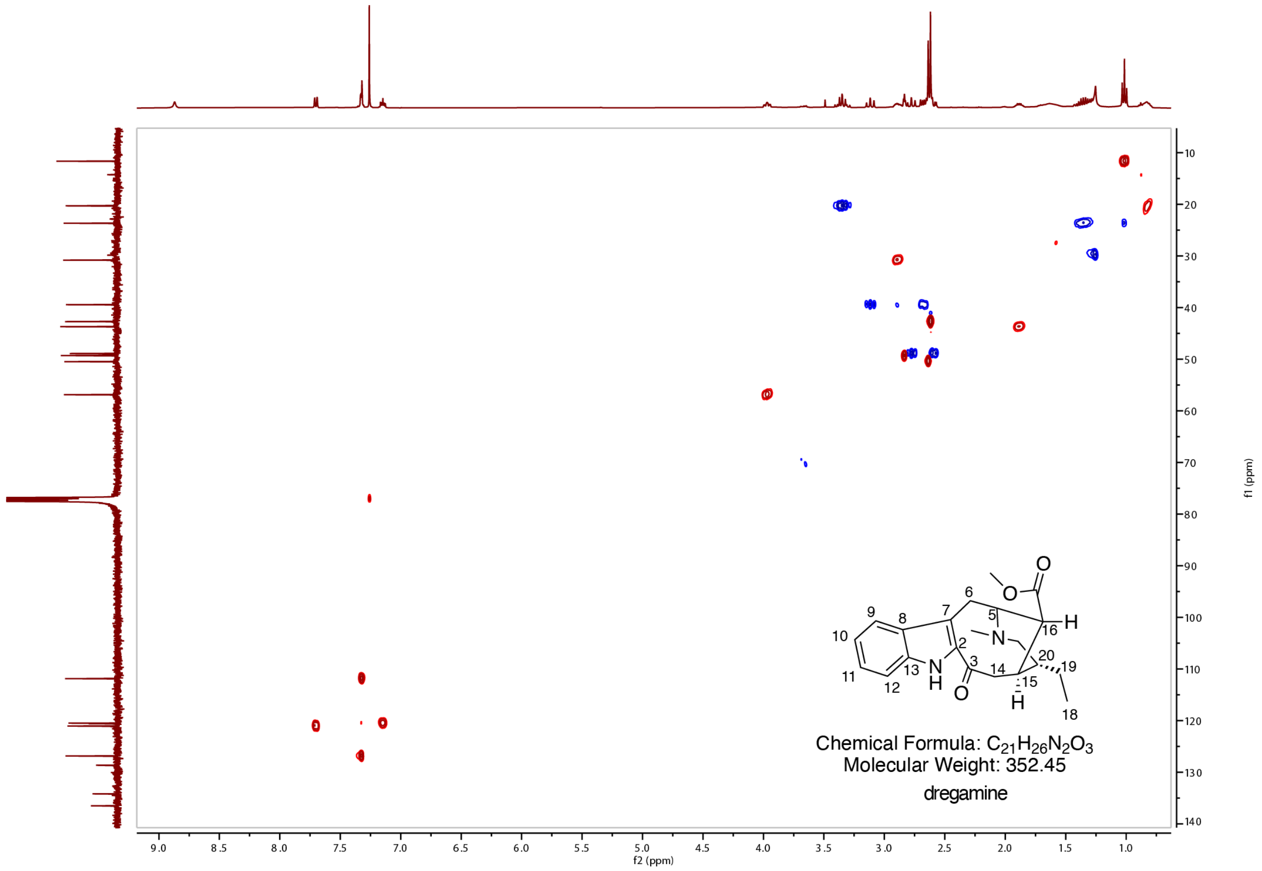

**Supplementary Figure 16. HMBC NMR spectra for dregamine in CDCl$_3$.**

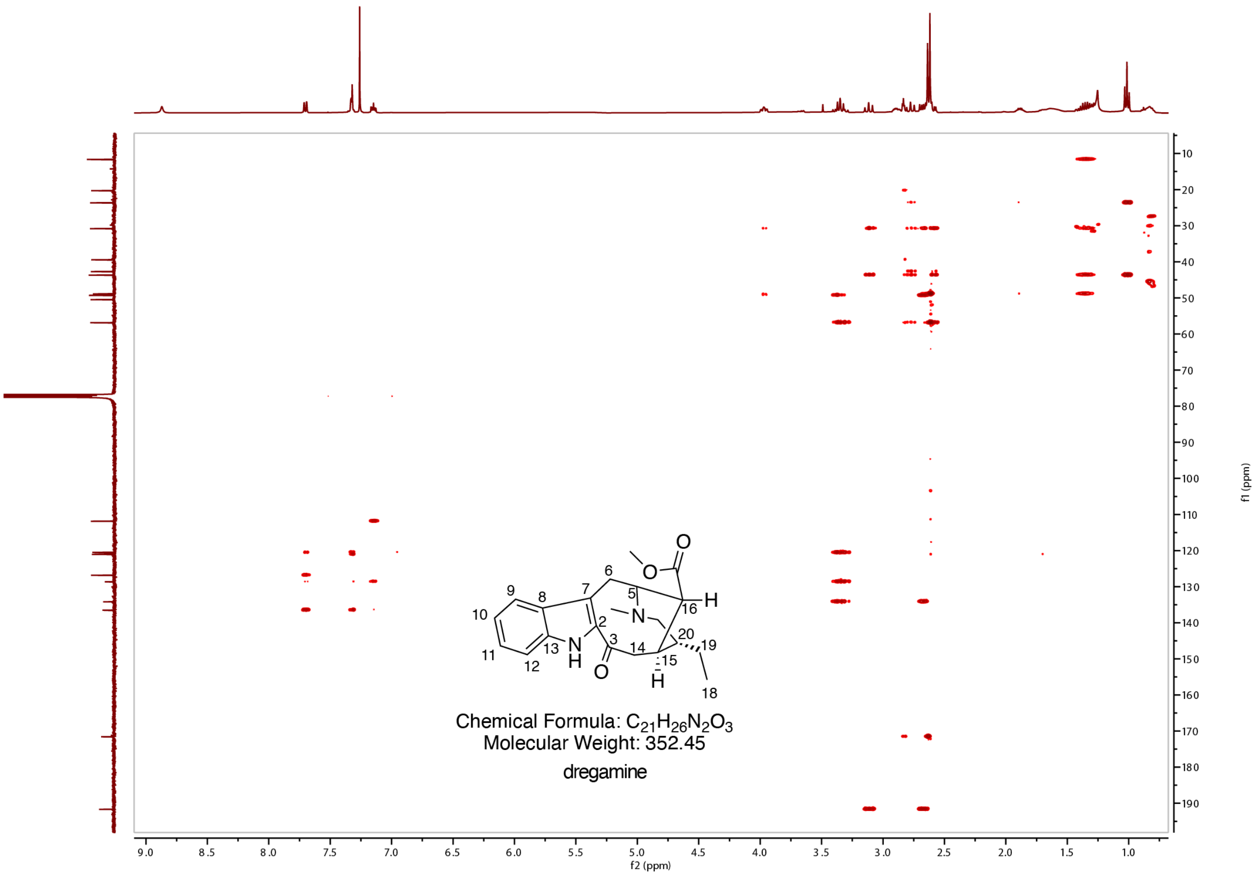

**Supplementary Figure 17. NOESY NMR spectra for dregamine in $CDCl_3$.**

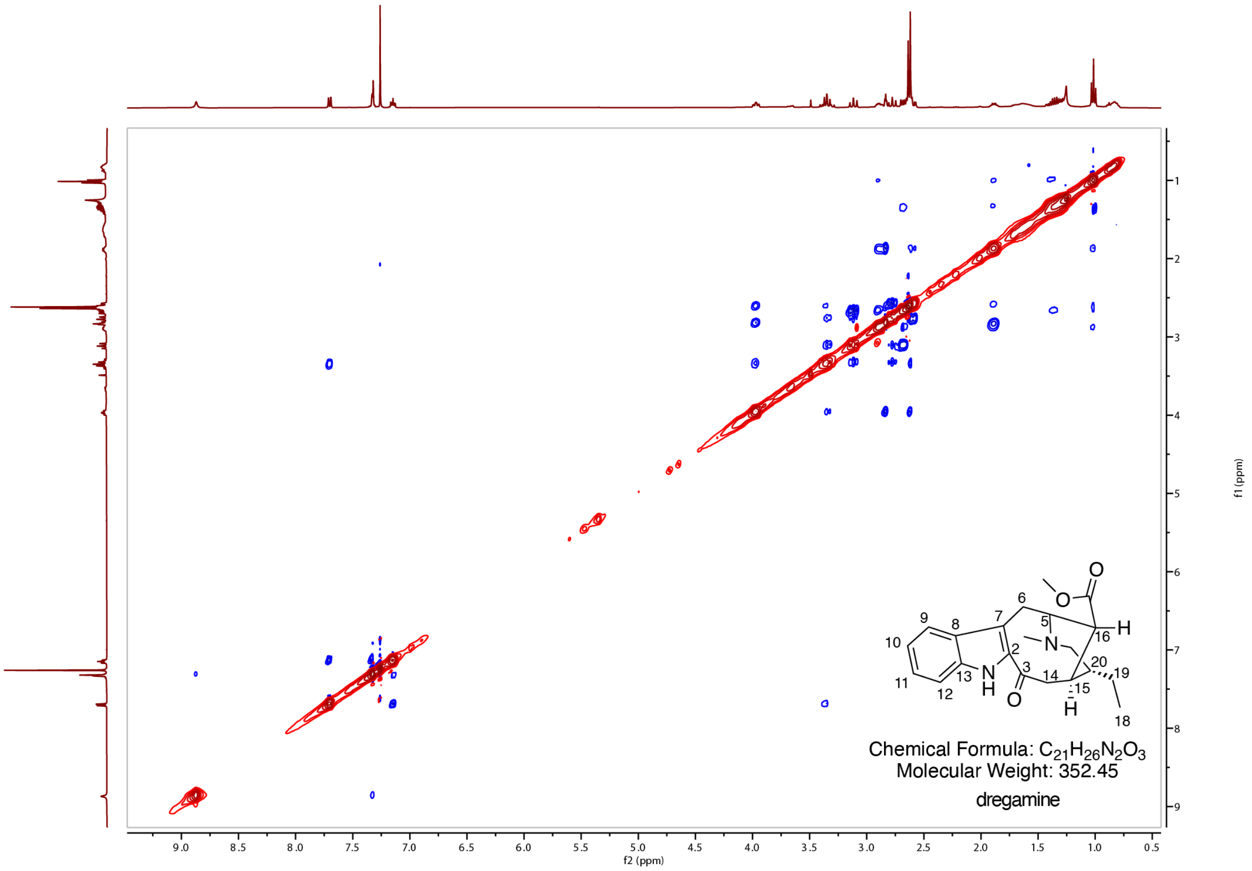

**Supplementary Figure 18. COSY NMR spectra for dregamine in $CDCl_3$.**

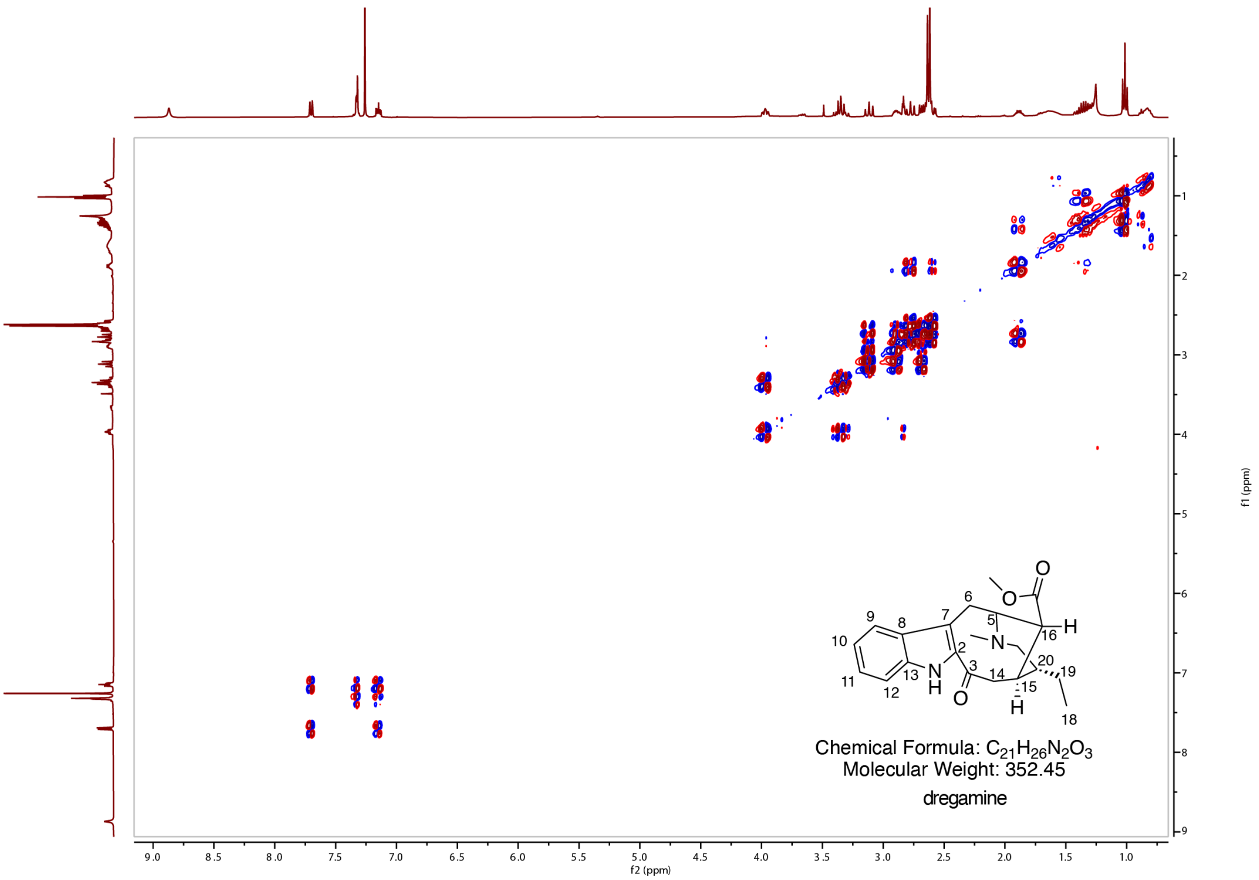

**Supplementary Figure 19. $^{13}$C NMR spectra for tabernaemontanine in $CDCl_3$.**

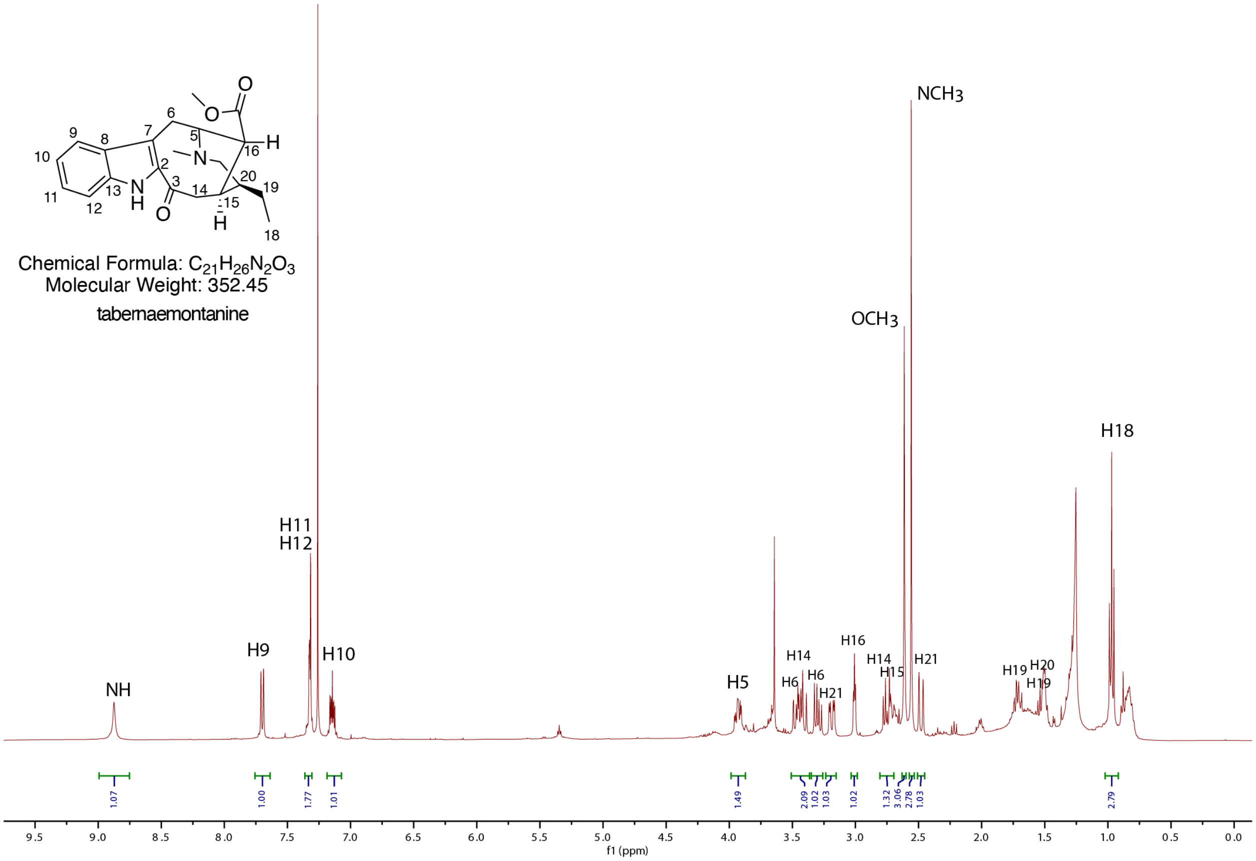

**Supplementary Figure 20.** $^{13}$**C NMR spectra for tabernaemontanine in CDCl$_3$.**

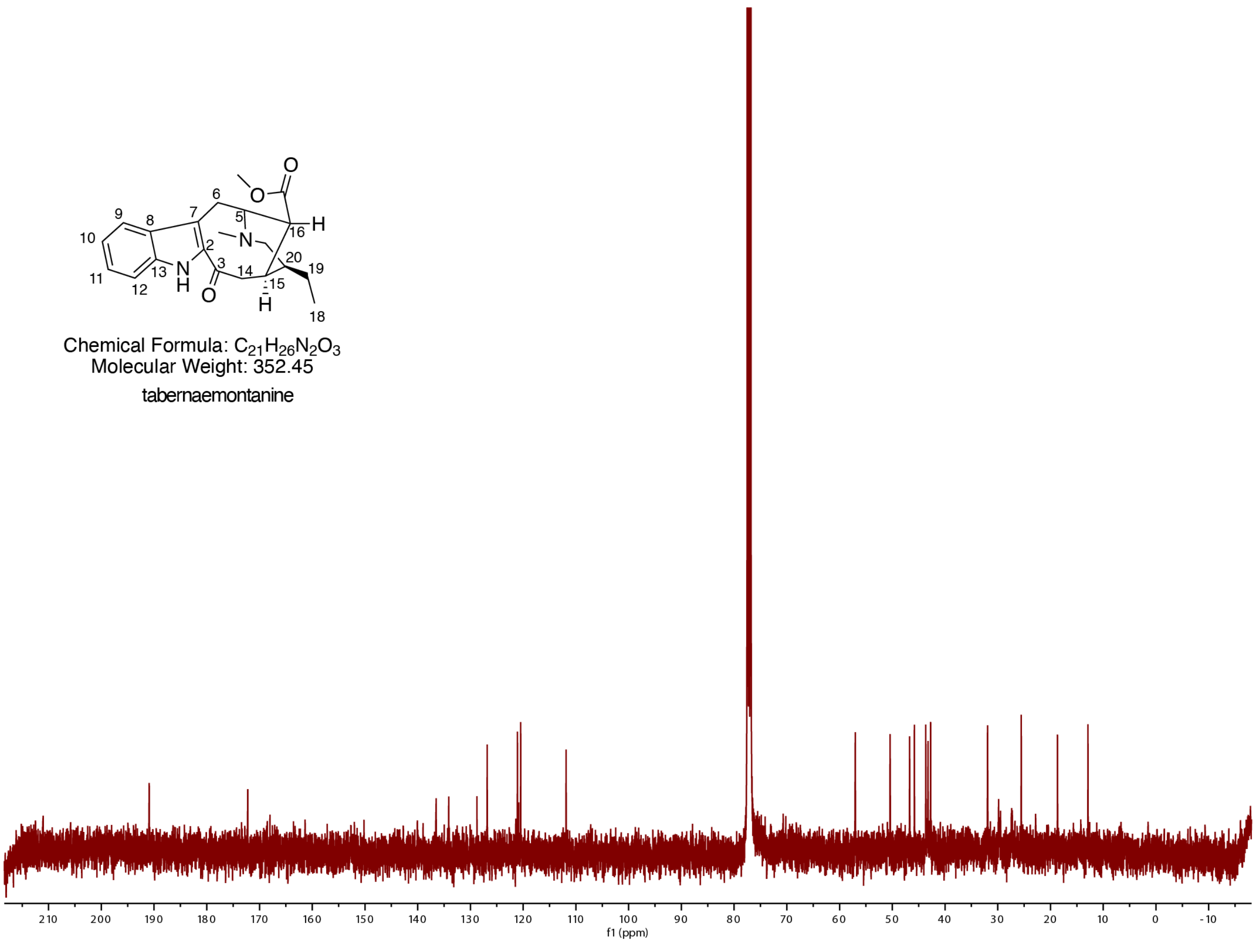

**Supplementary Figure 21. HSQC NMR spectra for tabernaemontanine in $CDCl_3$.**

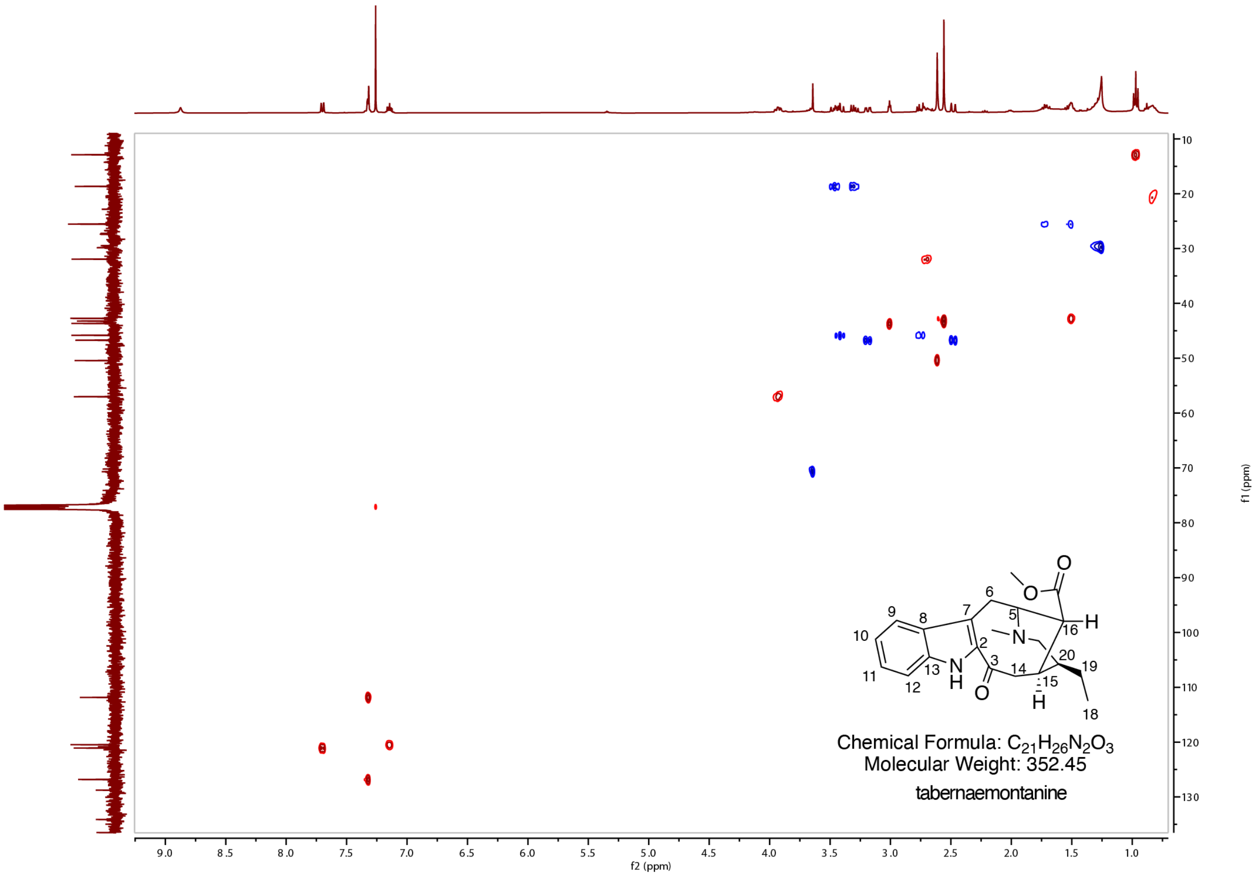

**Supplementary Figure 22. HMBC NMR spectra for tabernaemontanine in $CDCl_3$.**

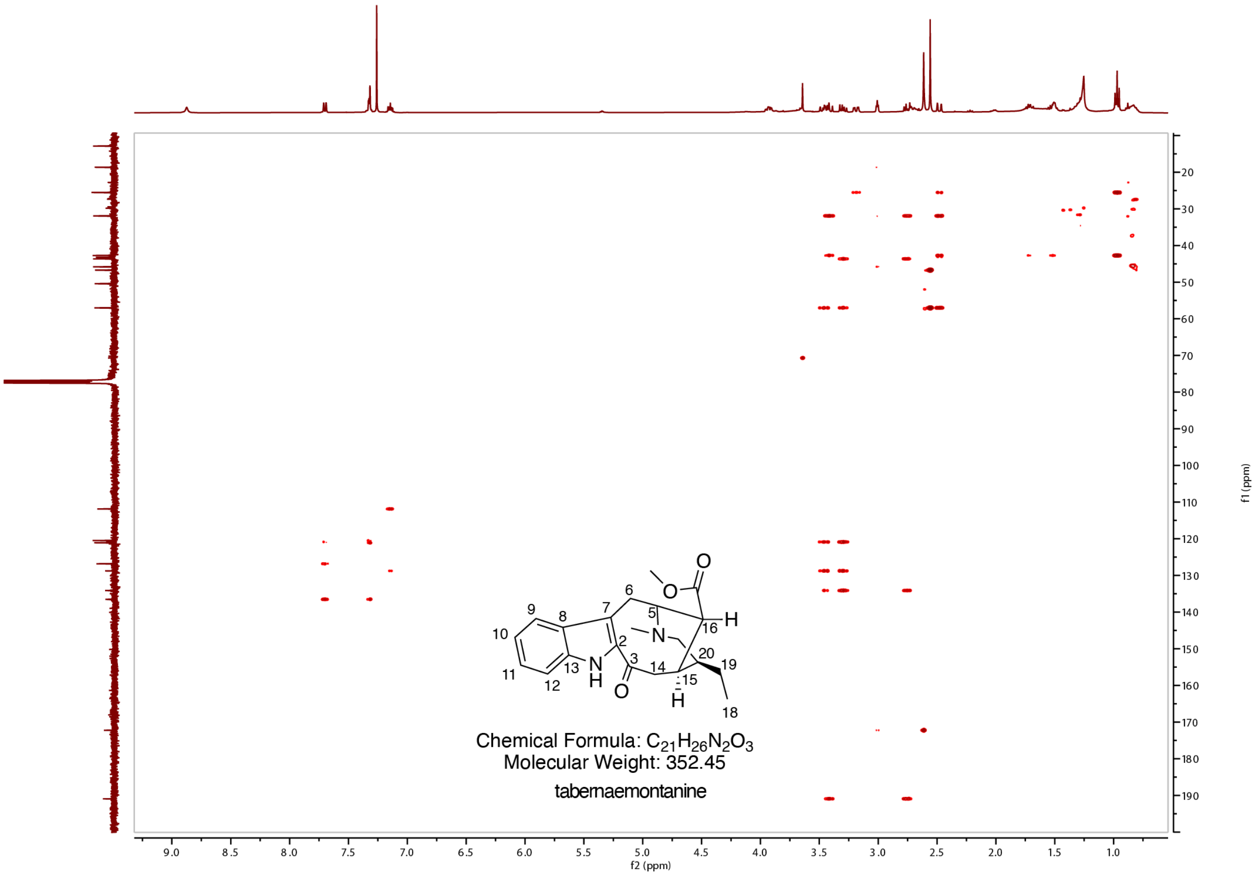

**Supplementary Figure 23. NOESY NMR spectra for tabernaemontanine in $CDCl_3$.**

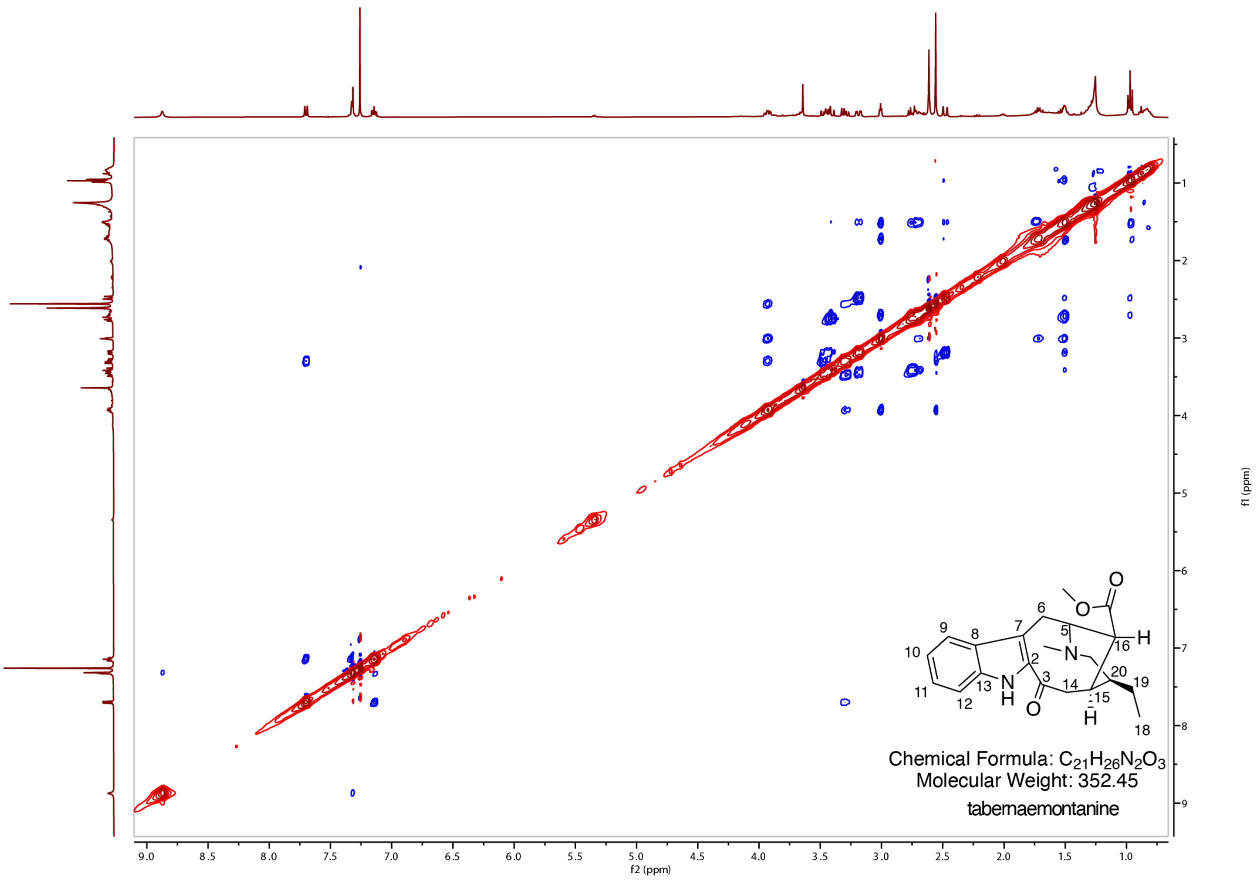

**Supplementary Figure 24. COSY NMR spectra for tabernaemontanine in CDCl₃.**

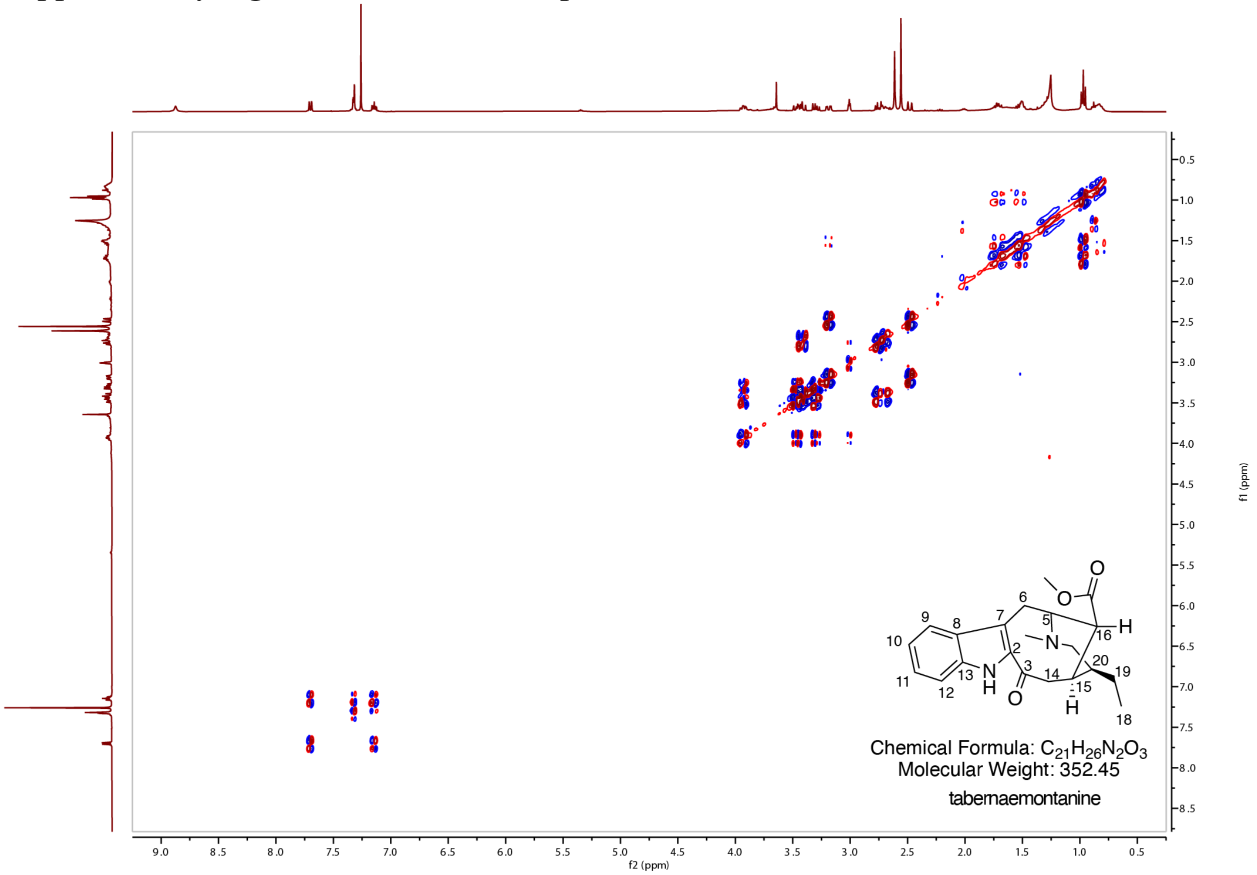

**Supplementary Figure 25. SDS-PAGE gels of N-terminally 6X-Histagged TePeNMT and CrPeNMT**

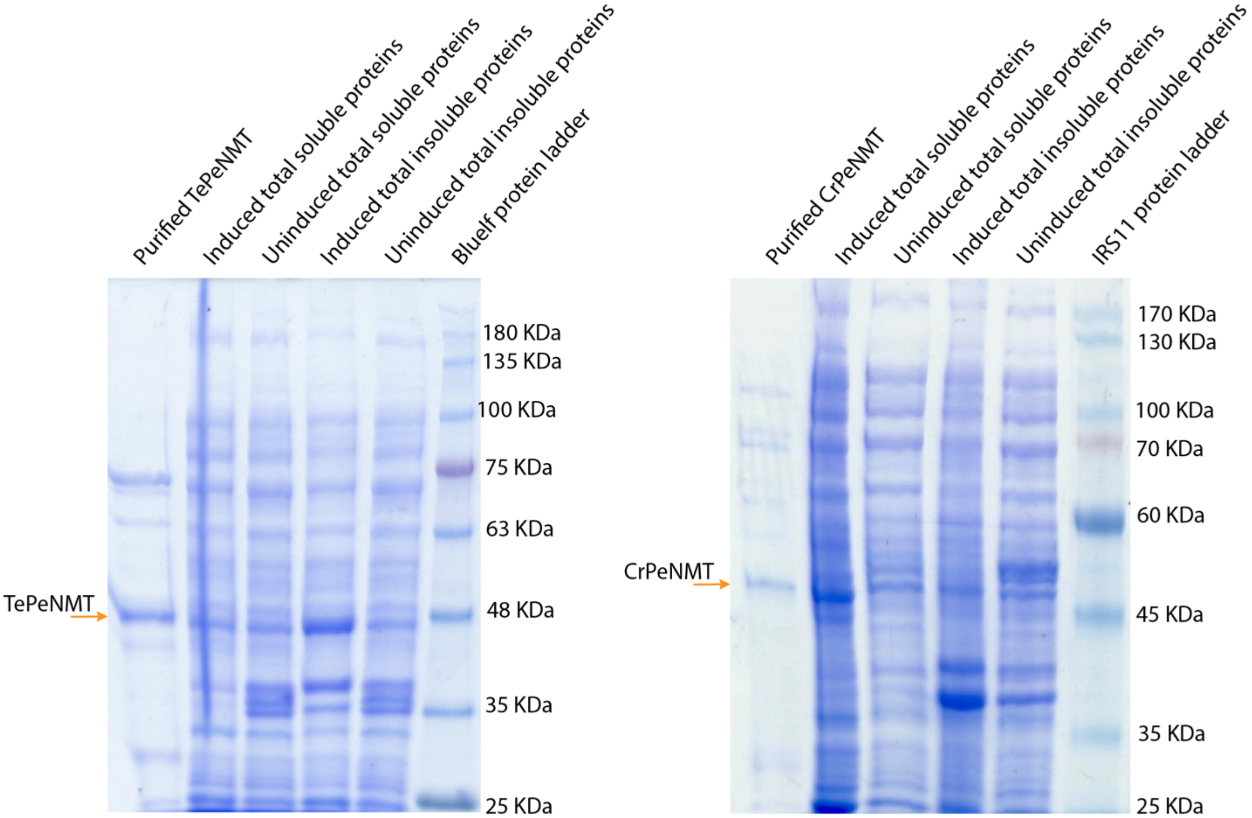

SDS-PAGE gels of *N*-terminally 6X-Histagged TePeNMT (left panel) and CrPeNMT (right panel) proteins purified by Ni-NTA affinity Chromatography. Orange arrows indicate recombinant enzymes.

**Supplementary Figure 26. 2D ligand interaction maps for *S*-adenosylmethionine (SAM) and perivine at the enzyme active sites of TePeNMT and CrPeNMT.**

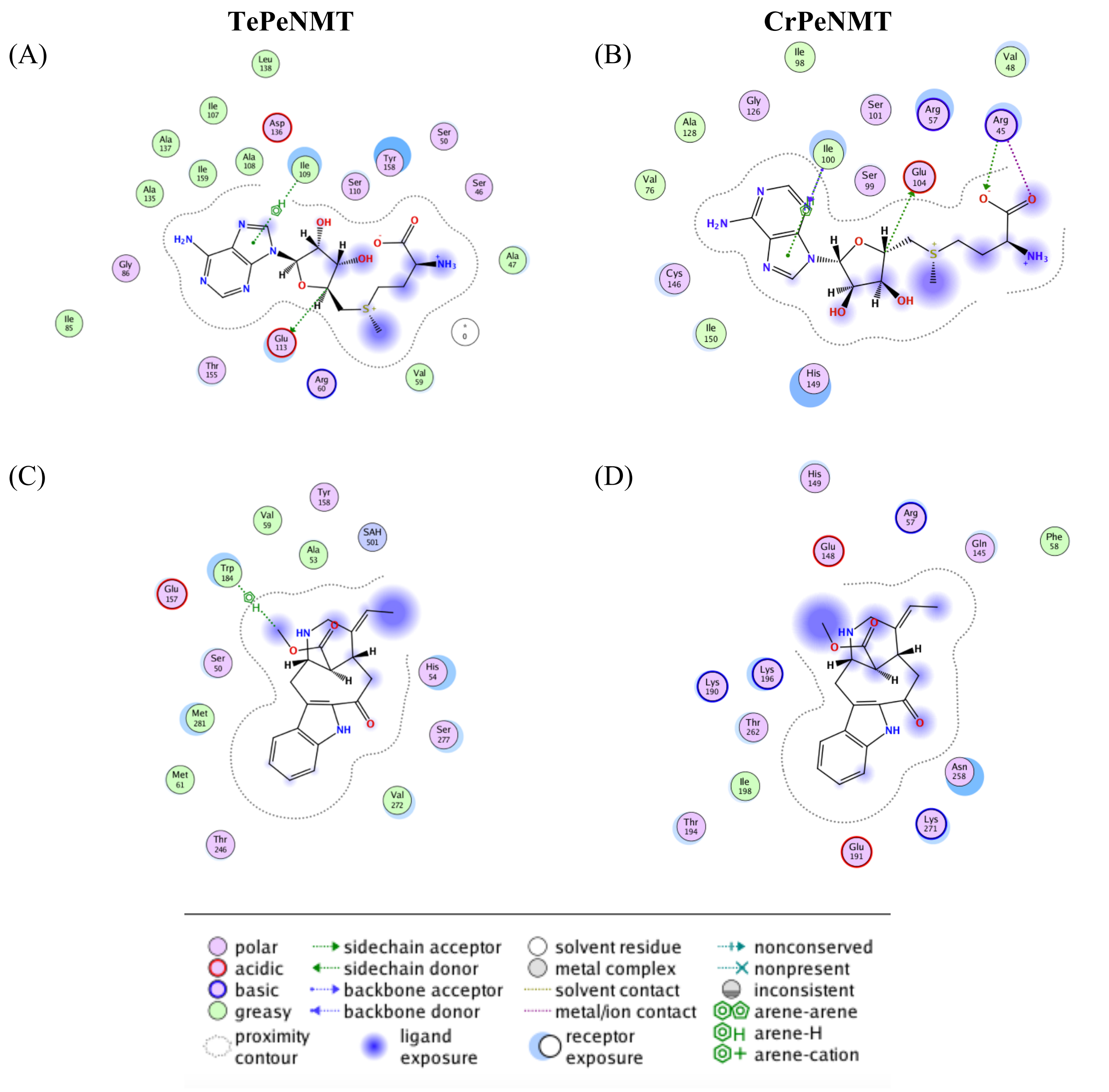

(A) and (B): active site amino acid residues interacting with SAM. (C) and (D): active site amino acid residues interaction with perivine, (A) and (C): enzyme active sites of TePeNMT. (B) and (D): enzyme active sites of CrPeNMT.

**Supplementary Table 1. $^1$H NMR chemical shifts of vobasine, dregamine, tabernaemontanine, and apparicine in $CDCl_3$.**

| | Vobasine This study | Vobasine Reference[1] | Dregamine This study | Tabernaemontanine This study | Apparicine This study | Apparicine Reference[2] |
|---|---|---|---|---|---|---|
| NH | 9.02 s | - | 8.87 s | 8.88 s | 7.87 br s | 7.84 s |
| 3 | - | - | - | | 3.07 dddd 3.44 ddd | 3.07 dddd 3.42 ddd |
| 5 | 3.97 ddd | - | 3.97 ddd | 3.93 ddd | - | - |
| 6 | 3.44 dd 3.51 dd | - | 3.35 m 3.35 m | 3.30 dd 3.46 m | 4.27 d 4.52 d | 4.28 d 4.51 d |
| 9 | 7.71 d | 7.71 d | 7.70 d | 7.70 d | 7.42 d | 7.42 d |
| 10 | 7.15 ddd | 7.15 m | 7.15 ddd | 7.14 ddd | 7.06 ddd | 7.06 ddd |
| 11 | 7.34 dt | 7.15 m | 7.33 ddd | 7.32 ddd | 7.18 ddd | 7.18 ddd |
| 12 | 7.34 dt | 7.33 d | 7.32 d | 7.32 d | 7.28 dd | 7.28 d |
| 14 | 2.73 dd 3.32 dd | - | 2.68 dd 3.12 t | 2.75 dd 3.42 dd | 1.90 dddd 2.18 dddd | 1.89 ddt 2.16 dddd |
| 15 | 3.78 ddd | - | 2.89 dddd | 2.70 m | 3.92 br s | 3.92 br s |
| 16 | 2.83 dd | - | 2.83 t | 3.01 t | - | - |
| 17 | - | - | 1.02 t | 0.97 t | 5.27 s 5.40 s | 5.26 s 5.39 s |
| 18 | 1.71 dd | 1.71 dd | 1.35 m 1.35 m | 1.53 dd 1.71 dd | 1.47 dd | 1.46 dd |
| 19 | 5.46 q | 5.45 q | 1.89 dddd | 1.51 m | 5.28 q | 5.25 q |
| 21 | 2.98 d 3.85 d | - | 2.59 dd 2.76 d | 2.48 d 3.19 dd | 3.22 d 3.83 dt | 3.20 d 3.82 dt |
| OMe | 2.60 s | 2.61 s | 2.64 s | 2.61 s | - | - |
| NMe | 2.65 s | 2.65 s | 2.62 s | 2.56 s | - | - |

**Supplementary Table 2. $^{13}C$ NMR chemical shifts of vobasine, dregamine, tabernaemontanine, and apparicine in $CDCl_3$.**

| | Vobasine This study | Vobasine Reference[3] | Dregamine This study | Tabernaemontanine This study | Apparicine This study | Apparicine Reference[2] |
|---|---|---|---|---|---|---|
| 2 | 134.3 | 134.1 | 134.2 | 134.1 | 145.1 | 145.2 |
| 3 | 190.3 | 190.1 | 191.7 | 190.9 | 45.3 | 45.3 |
| 5 | 57.4 | 57.2 | 56.9 | 57.0 | - | - |
| 6 | 20.6 | 20.3 | 20.3 | 18.7 | 54.3 | 54.2 |
| 7 | 120.5 | 120.4 | 120.5 | 120.9 | 110.1 | 111.5 |
| 8 | 128.7 | 128.5 | 128.7 | 128.8 | 129.1 | 129.0 |
| 9 | 121.1 | 120.8 | 121.1 | 121.1 | 118.8 | 118.6 |
| 10 | 120.6 | 120.4 | 120.6 | 120.5 | 123.2 | 119.3 |
| 11 | 126.9 | 126.7 | 126.9 | 126.8 | 119.6 | 123.0 |
| 12 | 111.9 | 111.7 | 111.9 | 111.9 | 110.4 | 110.2 |
| 13 | 136.6 | 136.3 | 136.5 | 136.5 | 137.0 | 137.4 |
| 14 | 43.2 | 43.0 | 39.4 | 45.8 | 29.6 | 29.6 |
| 15 | 30.6 | 30.4 | 30.8 | 32.0 | 41.3 | 41.2 |
| 16 | 46.7 | 46.6 | 49.3 | 43.7 | 135.8 | 135.6 |
| 17 | - | - | - | - | 112.5 | 112.2 |
| 18 | 12.5 | 12.3 | 11.7 | 12.9 | 12.7 | 12.6 |
| 19 | 121.0 | 120.8 | 23.7 | 25.5 | 120.8 | 120.1 |
| 20 | 136.0 | 135.8 | 43.7 | 42.6 | 131.5 | 131.3 |
| 21 | 52.0 | 51.8 | 48.9 | 46.7 | 54.2 | 54.3 |
| C=O | 171.5 | 171.2 | 171.6 | 172.2 | - | - |
| OMe | 50.6 | 50.3 | 50.5 | 50.4 | - | - |
| NMe | 42.5 | 42.3 | 42.7 | 43.2 | - | - |

### Supplementary Table 3. MIA substrates tested for TeMT1-3 in this study

| MIA type | heteroyohimbine | corynanthe | yohimbe | iboga | Aspidosperma | sarpagan | akuammiline | Vincamine |
|---|---|---|---|---|---|---|---|---|
| MIA tested for TeMT1-3 in this study | Ajmalicine<br>Reserpiline | Corynantheidine<br>Paynantheine<br>Speciociliatine | Yohimbine | Catharanthine<br>Coronaridine<br>Heyneanine<br>Voacangine<br>Voacangarine | Tabersonine<br>Vincadifformine<br>19-hydroxytabersonine<br>2, 3-dihydrotabersonine<br>Minovincinine | Pericyclivine<br>Strictamine<br>Vomilenine<br>Ajmaline | Picrinine | Vincamine |

### Supplementary Data 1. Multiple alignment for NMTs used in the phylogenetic analysis of this study.

>Hp9OMT
-------------------------------MYVVISHRYICHPSKVSLSSMFSNQTHPLAMDDAQQNYLRVLEM
GCTQILHAVFTTVIE-LNVFEIIAKAGPEAQLSAAEISS--HL--PTQN--------QQAPAILERMLQLLASYSV
LKCVHVSNQDGRGTRL--YGLTPMCRYLVAD-----TMGISM-GPAML----------CYTDKAMADSWSYLKDAV
L--EGK-IPFNKANKMDLFEYFG--KSSTLNETFNQAMHSETFFVLRAVLQNY-KGFEALKELVDVGG-------G
LGITLNAIISKYPGIRGINFDLPQVIKDAPIRTGVENLPGDMFEYVPKGEAILLKNILHDWTDEHCLKLLKNCYNA
LPEH---GKVIVIEMILPTSPENDLLSRAVFFVDIMMLALTSGGRERTLKEFDALAKGAGFIACKLVCQTFGYGIL
EFYKSSTLNSSEIP
>CpOMT1_MW456557
-----------------------------------------------------MATSENSTKLLRAQAHIWNQ
TFIFKNSASLKCAIQ-LGIPDVIQ--KHGKPITLSDLIS--AL--PINP--------SKALYI--DRLMRILVNNG
F---LAQEKEGY------YTLTSAGRLLLKDDETLSAREFVL----------------MVLDPALVKPWSVLTEWF
K--NDDRSPFDTAHGKSFWEYMA--DDPKLGKLFNDAMASDSQLITKVLITECRYVFEELTSLVDVGG-------G
TGTVARSIAKMLPHLNCIVFDLPHVVANQEGTENLDFVAGDMFEKVPPTNAILLKWILHDWSDEDCVKILKNCKKA
IPGRDKGGKVIVIDMVMDSQLIKDDESVEAQICFDMEMLVLFRSKERTEKELATLFWDAGFSRYKVLPVLGTRGLI
EVY----------P
>Cr16OMT
-------------------------------------------------------MDVQSEEFRGAQAQIWSQ
SCSFITSASLKCAVK-LGIPDTID--NHGKPITLSELTN--ALVPPVHP--------SKAPFI--YRLMRVLAKNG
F-CSEEQLDGETEPL---YSLTPSSRILLKKEPLN-LRGIVL----------------TMADPVQLKAWESLSDWY
QNEDDSSTAFETAHGKNFWGYSS--EHMEHAEFFNEAMASDSQLISKLLIGEYKFLFEGLASLVDIGG-------G
TGTIAKAIAKNFPQLKCTVFDLPHVVANLESKENVEFVAGDMFEKIPSANAIFLKWILHDWNDEDCVKILKSCKKA
IPAK----------------------------------------------------------------------
--------------
>TiN10OMT_MH454075
-------------------------------------------------------DAMKSAELFKAQAHIFKQ
VFCFTNGASLKCAVQ-LGIPDAID--NHGKAMTLSELTD--AL--PINP--------SKAPHI--HRLMRILVTAG
F-FVEERLGNGKEEKANGYALTPSSRLLLKNKPLS-LRASAL----------------TMLDPVTVKTWNALSEWF
Q--NEDQTAFETAHGKNMWDFFA--EDPGLSKKFNESMASDSQLVTEVLVTKCKFVFEGLTSMVDVGG-------G
TGTVAGAIAKTFPSLRCTVFDLPHVVANLEPTENLDFVAGDMFGKIPPANAIFLKWVLHDWNDEDCVKILKNCKRA
IPGKEKGGKVIIVDIIMETEKHDIDEFDYAKMCMDMEMLVLCNSKERTEKELAMLVSEAGFSGYKIFPVLGIRSLI
EVY----------P
>RsANMT_KC708445
--------------------------------------------------------------------------
------MAENQEALA--EFYDKGV--GVWDNLSREHMHF--GYYAPGAT-ATIGG--HRASLV--RLIDEALCFAE
F-PD-DPEKKPRNMLDVGCGIGGTCLHVAKKYDIQ-CKGINI-SPEQVKIAQGLAAAQGLESKVSFDVGDALDMPY
P--DGA---FDLVLSIHCIEHLQ--DKEKFIREMVRVAASGATII---ILSHVHRDLS-----------------P
SEQSLKPQEERVLRKIGSSVQ-AWFCPL------SNYV--SLLAPLP----------VEVIKIADWSRNIDPSSRL
MLKV-------AFSVKGIVSNLMKGVQGWTAIKNVLPMKLLHKALHDGLVKFVVLTCRKSN---------------
--------------
>CrPeNMT_TMT1_KC708453
--------------------------------------------------------------------------
-------MGEKEAVA--ELYDKVTSNGILEELFGEHLHD--GYYEVGTV-ATISA--HRAAVV--RIIDEALRFAD
V-FTDDQAKKPRNMLDVGCGKGGTCVHIARKYDIQ-CTGISI-SPDEIQCAKHLAASQGLENKVSFDVGDALNMRY
S--DGS---FELIFVIQCIEHIQ--DKEKFIREIVRVAAPGAQIV---IISTACRNLS-----------------P
SEKSLKPKEEKTLKKICNYLHLSGFCSL------SDYS--NWLTPLP----------IEDMKIADWTQNAAPFYTL
LLRE-------AFSIKGFISLLMNG--GWTAVKVILGMKTIHEAIENDLLKIVAVTFRKTK---------------
--------------
>CrDhtNMT_ADP00410
--------------------------------------------------------------------------
------MEEKQEKVA--EFYDKVT--GAWDLFYGVHLHD--GYYEPGTT-ATMAI--SQDAVI--RMIDELLRFAG
V-SE-DPAKKPRSMLDVGSGLGGTCVYVAKKYDIQ-CTGITI-SPNQVKYAQDYAATEGVENKVSFDCGDALDMPY
S--DGK---FDVVFTINCIKHVH--DKEKFIREMVRVAAPGAAII---IASQAHPNLS-----------------P
GE-SLKPRDKKILQKICDGAGAVSLCSS------DDYV--RWLTPLP----------VKEIKAADWTQNITPLYPL
LMKE-------AFTWKGFTSIVLKG--GWRAINLINAVRLVAKAANDGILKFAVVTGRKSI---------------
--------------
>CrPiNMT_TMT2_A0A8X8M4T9
--------------------------------------------------------------------------
---MAAVVEKQEAVA--EFYDNST--GAWEELFGEHLHD--GYYEPGTT-ATIPA--HRAAVV--RMIDEALRFAG
V-STDDPAKKPRNLLDVGCGLGGTCLYLAKKYDIK-CTGITI-SPEQVKCAEDLAAAQGLENKVSFDVGDALDMPY
Q--DGE---FDVVFTLQCIDHVQ--DKEKFIREMVRVGSPGAAIV---VITYTHRDLS-----------------P
TEQSLKPHEIKTLKKICDNIVLSSISST------HDYV--NWMTSLS----------LKDIKTADWTQNIIPFYPL
LFKV-------SFSMKGFISLLMKG--GWSAIKVVLAVKMMSKAIDDGLLYYTAVSGRKPN---------------
--------------
>RsNNMT_KC708449

----------------------------------------------------------------------------
------MAEKQQAVT--EFYNNTSPRGAWEFLLGDHLHE--GFYDPGTT-ATISG--SQAAAA--RMIDEALRFAN
I-YD-DPSKKPKNMLDIGCGVGGTCVHVAKQYGIQ-CKGITL-SPEEVKCAQGIAKAQGLEEKVSFDVGDALNLPY
K--DGT---FDLVLTIECIEHVQ--DKEKFIREMIRVAAPGAPIV---ILSYAHRNLS-----------------P
SAESLKPDEKKVLKKICDNLALSCLCSS------ADFV--RWLTQLP----------AEDIKTADWTQNTSPFFPL
LMKE-------TFFTWKGFTSLLMKG--GWTAIKELLALRMMSKAADDGLLKFVAITCRKSK---------------
--------------
>VmPiNMT_KC708450
--------------------------------------------------MYTCSIIIYILTFWQLSKIKKQVAAA
EKQVMTVTEKQEAVA--EFYDKST--DAWEVFFGEHLHD--GFYEPGTT-ATIPG--SKVAVV--RMIDELLRFAG
I-SD-DPEKKPKTMLDVGCGLGGTCLHVAKKYDIK-CTGITI-SPEQVKCAQDLAATQGLESKVSFDVGDALDMPY
K--DGT---FDLVFTIQCIEHIQ--DKEKFIREMVRVAAPGAPVV---IAGYAARNLS-----------------P
SEESLKPEEKMVLEKICDHIVLSWLCST------GDYV--KWLTPLP----------VQDIKVWDLTQNITPFYPL
CIKE-------AFTWKSFTSLLKMG--GWSAIKVVFAVKMMAMAAEEGLLKFAAVTCRKSK---------------
--------------
>CrPiNMT_TMT4_A0A8X8M501
----------------------------------------------------------------------------
------MAEKQQAVA--EFYDNST--GAWEVFFGDHLHD--GFYDPGTT-ATIPA--SRAAVV--RMIDEALRFAN
V-ST-DPAKKPRNMLDVGCGIGGTCLYVAKKYDIQ-CTGITI-SPEQVKCAQGFAAAQGLENKATFDCGDALNMPY
K--DGT---FDVVFTIQCIEHIQ--DKEKFIREMVRVAAPGAAIV---IVSYGHRNLS-----------------P
GEESLKPEEKKTLKKICDNIVLSWLCSS------ADYV--RWLTPLP----------VQDIKTADWTQNIQPFYPL
LFKE-------AFTWRGFTSLLMKG--GWSAIKVVLAVKVMAKAADDGLLKFMAVTCKKSK---------------
--------------
>RsPiNMT_KC708448
----------------------------------------------------------------------------
------MAEKQQAVA--EFYDNST--GAWEVFFGDHLHD--GFYDPGTT-ATIAG--SRAAVV--RMIDEALRFAN
I-SD-DPAKKPKTMLDVGCGIGGTCLHVAKKYGIQ-CKGITI-SSEQVKCAQGFAEEQGLEKKVSFDVGDALDMPY
K--DGT---FDLVFTIQCIEHIQ--DKEKFIREMVRVAAPGAPIV---IVSYAHRNLS-----------------P
SEGSLKPEEKKVLKKICDNIVLSWVCSS------ADYV--RWLTPLP----------VEDIKAADWTQNITPFYPL
LMKE-------AFTWKGFTSLLMKG--GWSAIKVVLAVRMMAKAADDGVLKFVAVTCRKSK---------------
--------------
>CrNMT_ADP00411
---------------------------------------------------------------------MAKKSSV
EQLQEKDQEKFKGVA--DLYGGTS--RAWEDILGDHWHH--GYYDPGST-VSKSD--NAAALI--RMIDEVLRFGS
VFSAENQENKPKRILDIGCGIGGTCTYLARKYGAH-CTGITI-SSGEVERAQALATAQGLQEKVSFEVANALALPF
P--DGQ---FDLVWCMETAEHIP--EKEQLVKEIVRVAAPGGQII---LTSWCHRNLL-----------------P
SEQSLPLDEQKFIKKMCDLVLMHPFCSV------NEYI--NLFQSHH----------VEDMKTDDWCEFAKPFWTA
MVGS-------ACTIKGFFSLLWIG--GWNSLKFVMGSNMMIKAHKKGLVKLLVLSCRKPR---------------
--------------
>TePeNMT_PP067959
------------------------------------------------------------------------MAET
SKSAKADQERVNGVA--EIYNETA--GTMERMCGEHIHH--GYYEPGSAPASMSA--HPAAQV--RMIDEVLRSAS
V-SV-DPEKRPKSILDIGCGIGGTCLHLARKYGAQ-CKGIAI-SPVEVERAQALAAAQGLQGQVSFAVADALSLPF
P--EGQ---FDLIWCMETSEYIA--DKEKLVSEFVRVAAPGGTII---LASWCHRDLS-----------------P
SEKSMKPDEEKLIKKICDLNLFPPFCSS------SDYI--KLFKSHP----------LEDIKAEEWSQYASPTWPL
LIGE-------TFTIKGFISLLWTG--GWKAVKLVTSVQTMMKAHQKGALKFAVISCRKPK---------------
--------------
>TeNMT3_PP067961
----------------------------------------------------------------------------
----------MKGTR--ESYDEKY--GRWEALWGEHLHN--GYYEPGSI-VYLSD--HLAAQV--RMIEEVLRFAS
V-SD-DPERKPRSILDVGCGVGGSSLYLASKYDAL-CKGINI-SPFEVERAQALAAAQGLQSKVSFQVADALAQPF
S--DGQ---FDLVWCLESAEYVA--DKEKLVSEMVRVAASGATIV---IASWCHRDLS-----------------P
SEQSLKPDEEKLMKKLCDSMLLSPLCST------ADYI--KLLKSHS----------FEDIKVADWSEHASPFWPV
MYRS-------TFTWKGFVSLLRTG--GWKAMKLVLAFKTMMEGQKKGLFKLAIIACQKPENAHKDFATFDVSAK-
--------------
>AthγTMT_AAD02882
--------------------------MKATLAAPSSLTSLPYRTNSSFGSKSSLLFRSPSSSSSVSMTTTRGNVAV
AAAATSTEALRKGIA--EFYNETS--GLWEEIWGDHMHH--GFYDPDSS-VQLSDSGHKEAQI--RMIEESLRFAG
V-TDEEEEKKIKKVVDVGCGIGGSSRYLASKFGAE-CIGITL-SPVQAKRANDLAAAQSLSHKASFQVADALDQPF
E--DGK---FDLVWSMESGEHMP--DKAKFVKELVRVAAPGGRII---IVTWCHRNLS-----------------A
GEEALQPWEQNILDKICKTFYLPAWCST------DDYV--NLLQSHS----------LQDIKCADWSENVAPFWPA
VIRT-------ALTWKGLVSLLRSG---MKSIKGALTMPLMIEGYKKGVIKFGIITCQKPL---------------
--------------
>HaγTMT_ABB52798
---------------------------------------------------------MATTAVGVSATPMTEKLTA
ADDDQQQQKLKKGIA--EFYDESS--GMWENIWGEHMHH--GYYNSDDV-VELSD--HRSAQI--RMIEQALTFAS
V-SD-DPEKKPKTIVDVGCGIGGSSRYLARKYGAE-CHGITL-SPVQAERANALAAAQGLADKVSFQVADALNQPF
S--DGK---FDLVWSMESGEHMP--DKLKFVSELTRVAAPGATII---IVTWCHRDLN-----------------P
GEKSLRPEEEKILNKICSSFYLPAWCST------ADYV--KLLESLS----------LQDIKSADWSGNVAPFWPA
VIKT-------ALSWKGITSLLRSG---WKSIRGAMVMPLMIEGFKKDVIKFSIITCKKPE---------------

```
-------------
>PfγTMT_AAL36933
------MAEAVTPGICTTGWRRGGVHAPTYNISIKPATALLVGCTTKTKSITSFSTDSLRTRGRARRPTMSLNAAA
AEMETEMETLRKGIA--EFYDESS--GVWENIWGDHMHH--GFYEPAAD-VSISD--HRAAQI--RMIEESLRFAS
F-SPITTTEKPKNIVDVGCGIGGSSRYLARKYGAKLSRAITLSSPVQAQRAQQLADAQGLNGKVSFEVADALNQPF
P--EGK---FDLVWSMESGEHMP--DKKKFVNELVRVAAPGGRII---IVTWCHRDLS----------------P
SEESLRQEEKDLLNKICSAYYLPAWCST------ADYV--KLLDSLS----------MEDIKSADWSDHVAPFWPA
VIKS-------ALTWKGITSLLRSG---WKTIRGAMVMPLMIEGYKKGVIKFAIITCRKPAS-------------
-------------
>CaγTMT-like_XP_027097246
----------------------------------MLQILSTKQLSTRFIHTGARILNKKEMAVRQASLGGVDVD
LETKTNVEKLKRGIA--EFYDESS--GIWEDIWGDHMHH--GFYDPNST-VSLSD--HRAAQV--RMIEEALKFAS
V-PE-DPMKKPKSIVDVGCGIGGSSQYLARKYGTE-CTGITL-SPVQAERARALAAAQGLENKVSFEVADALNQPF
P--DGK---FDLVWSMESGEHMP--EKAKFVNELARVAAPGARII---IVTWCHRNLS----------------S
SEQSLNPDEKKLLDKICDAYYLPTWCST------DDYV--KLLQPLS----------LQDIKAADWSEYVAPFWPA
VIRS-------ALTWKGITSLILSG---WKTIKGAMVMPLMMKGYKKGLIKFAIITCRKPE--------------
-------------
>TeNMT2_ PP067960
------------------------------------------------------------------------
------------------------------MWGDHMHH--GYYEPGTN-VSVID--HQAAQI--RLIEEVLRFAS
V-SD-DPIKKPKSIVDVGCGIGGSSRYLARKYGAK-CKGITL-SPVQAQRAQALAVAQGLQSKVSFEVADALAQPF
P--DGQ---FDLVWSMESGEHMP--DKEKFVRELARVAAPGATII---IVTWCHRDLS----------------P
SEQSLKPEEKKLLNKICDSYYLPAWCST------ADYV--KLLESVS----------LVDIKTADWSEYVAPFWPA
VIRS-------ALTWKGFASLLRSG---WKTIKGALVMPLMIEGFKKDLIKFAVITCRKPE--------------
-------------
>CrγTMT_TMT3_A0A8X8M4W6
MAAAPVFFPSSVVVSVYRAHGYAFSCSSLLSQLPAIKLSSYRPTRTDHTAITLLPNSNRIASRLQAMQSPTFEVNG
DGKTEEVEKLQKGIA--EFYDESS--GIWEEIWGDHMHH--GFYDPGTT-VSLAD--HRTAQI--RMIEEALRFAS
V-SD-DPMKKPKRIVDVGCGIGGSSRYFARKYDAE-CKGITL-SPVQAKRAQALAVAEGLQGTVSFEVADALAQPF
P--HGQ---FDLVWSMESGEHMP--DKEKFVRELVRVAAPGATII---IVTWCHRDLS----------------P
SEHSLKPQEQKLLNKICDSYYLPAWCST------ADYV--KILESLF----------LEDIKRADWSEYVAPFWPA
VIRS-------ALTWKGFTSLIRSG---WKTIKGAMVMPLMIEGFKKDVIKFSVITCRKPE--------------
-------------
>CreiNMT_XP_001695187
-----------------YNALVKTFIGAVENGYVPDFILRTGIRFLLSNRVAEPFSPSLPSDLNGQLQQTVAFVN
DLKTMPIAVNTSEAN-EQHYEIPT--PYYLLVLGSHLKYSSCLYRSPRE-------SLEQAEY--NMLDLYCERAQ
L-------RPGQRVLELGCGWGSFSLFAAARYPASTFFAVSN-SATQKAFIDGEAAKRGITN-LTVLTANMVDFEA
PTANGL---FDRVVSIEMFEHMK--NYQRLLRKVSGWLAAEGKLF---VHIFLHRTTPYHFEVQSEEDWMSKYFFT
GGTM--PSSDLLLYFQDDLVIRNHWYVN---GRHYSRTLEDWLVRH-------------DRNSREVRKILTTAYTG
SKDS-------ANVWFHR----------WRIFYIACRRLFNYKGGEEWGVGHFLFEQRG----------------
-------------
>PsRNMT_KX369612
--------------STTMETTKISQQDDLWKNMELGQISDEEVRRLMKIGIEKRIKWGTKPTQQEQLAQLLDFNK
SLRGMKMATEIDTLENHKIYETPE---SFNQIIGGKESA--GLFTDETT-TTMEE-----ANT--KMMDLYCERAG
L-------KDGHTILDLGCGAGLLVLHLAKKYKKSKITGITN-TSSHKEYILKQCKNLNLSN-VEIILADVTKVDI
---EST---FDRVFVIGLIEHMK--NFELFLRKISKWMKDDGLLL---LEHLCHKSFSDHWEPLSEDDWYAKNFFP
SGTLVIPSATCLLYFQEDVTVIDHWILS---GNNFARSNEVILKRI-------------DGKIEEVKDIFMSFYGI
GREE-------AVKLINW----------WRLLCITANELFKYNNGEEWLISQLLFKKKLMTCI-------------
-------------
>CjCNMT_BAB71802
-------------MAVEAKQTKKAAIVELLKQLELGLVPYDDIKQLIRRELARRLQWGYKPTYEEQIAEIQNLTH
SLRQMKIATEVETLD-SQLYEIPI--EFLKIMNGSNLKGSCCYFKEDST--TLDE-----AEI--AMLDLYCERAQ
I-------QDGQSVLDLGCGQGALTLHVAQKYKNCRVTAVTN-SVSQKEYIEEESRRRNLLN-VEVKLADITTHEM
A--ET----YDRILVIELFEHMK--NYELLLRKISEWISKDGLLF---LEHICHKTFAYHYEPLDDDDWFTEYVFP
AGTMIIPSASFFLYFQDDVSVVNHWTLS---GKHFSRTNEEWLKRL-------------DANLDVI----KPMFET
LMGN----EEEAVKLINY----------WRGFCLSGMEMFGYNNGEEWMASHVLFKKK-----------------
-------------
>PsCNMT_AAP45316
---------------------MQLKAKEELLRNMELGLIPDQEIRQLIRVELEKRLQWGYKETHEEQLSQLLDLVH
SLKGMKMATEMENLD-LKLYEAPM--EFLKIQHGSNMKQSAGYYTDEST--TLDE-----AEI--AMLDLYMERAQ
I-------KDGQSVLDLGCGLGAVALFGANKFKKCQFTGVTS-SVEQKDYIEGKCKELKLTN-VKVLLADITTYET
---EER---FDRIFAVELIEHMK--NYQLLLKKISEWMKDDGLLF---VEHVCHKTLAYHYEPVDAEDWYTNYIFP
AGTLTLSSASMLLYFQDDVSVVNQWTLS---GKHYSRSHEEWLKNM-------------DKNIVEFKEIMRSITK-
TEKE-------AIKLLNF----------WRIFCMCGAELFGYKNGEEWMLTHLLFKKK-----------------
-------------
```

**Supplementary Data 2. Cartesian coordinates for CrPeNMT homology model.**

N -2.1385667 21.7753992 -15.9844882
H -2.8881317 22.3451992 -15.612073
H -2.4187043 21.4412796 -16.898962
H -1.952445 20.9940613 -15.3721405
C -0.8983927 22.6008534 -16.1048954
H -1.0805973 23.3575681 -16.8709468
C 0.33849 21.7897622 -16.5659561
O 1.2553163 22.327476 -17.1874333
C -0.6053818 23.3116885 -14.7771474
H 0.3668547 23.80077 -14.8479421
H -0.5567763 22.5711725 -13.9778747
C -1.639947 24.3835703 -14.4160168
H -1.396285 24.7758699 -13.426708
H -2.6273748 23.9290305 -14.3587376
S -1.7285502 25.7740286 -15.5733039
C -2.9534829 26.7844932 -14.6928753
H -3.1342375 27.7045542 -15.2488035
H -2.5798462 27.0332981 -13.6984196
H -3.8891334 26.2323095 -14.5953778
N 0.3673335 20.4886197 -16.2657559
H -0.4109708 20.1080689 -15.7405706
C 1.3938537 19.5859753 -16.7771625
H 1.3580658 19.6103407 -17.8672226
H 2.3753087 19.9422459 -16.4680576
C 1.2496557 18.1385969 -16.3224611
O 0.7986647 17.8537127 -15.2154758
N 1.7628938 17.2390447 -17.1522057
H 2.1121366 17.5432004 -18.0501647
C 2.0288584 15.8382482 -16.8338954
H 1.9028599 15.7027836 -15.7604402
C 3.4957816 15.5076051 -17.1394012
O 4.119702 16.1812749 -17.9609391
C 0.9990926 14.9223526 -17.5032517
H 1.2572628 13.8928438 -17.2749979
H 0.0423459 15.1358366 -17.026009
C 0.8149397 15.0455348 -19.020348
H 1.6157817 14.5050359 -19.5288645
H 0.8699863 16.0954136 -19.3176114
C -0.5554566 14.4754363 -19.4180847
O -1.297864 15.2053148 -20.1119062
O -0.875195 13.3478418 -18.9747065
N 4.1061184 14.7016468 -16.2632362
H 3.5442127 14.1102083 -15.6715657
C 5.5629684 14.6122266 -16.0983634

H 6.0446756 14.8186955 -17.0600251
C 5.9695982 13.2129132 -15.6736854
O 5.3463653 12.639753 -14.786269
C 6.0594875 15.5900786 -15.0182406
H 5.5966651 15.3285324 -14.0641537
H 7.1389711 15.4636292 -14.915048
C 5.7688522 17.0627619 -15.3210483
H 6.199365 17.3109927 -16.2919241
H 4.6914859 17.2284646 -15.351451
C 6.3694499 17.9615862 -14.2310701
H 7.4516987 17.8220036 -14.192695
H 5.9514537 17.6743696 -13.2638997
C 6.0413128 19.4366002 -14.4861744
H 6.4275873 20.029751 -13.6516003
H 4.9522862 19.5499365 -14.5026758
N 6.6253332 19.9107507 -15.7656037
H 7.6331593 19.818804 -15.7546612
H 6.3843249 20.8792368 -15.933991
H 6.2683797 19.3550584 -16.5356419
N 7.106455 12.7593095 -16.1705568
H 7.4956956 13.2759729 -16.9574346
C 7.5639831 11.383838 -16.027934
H 6.7220861 10.7521143 -15.7410336
C 8.6256294 11.3080201 -14.9245199
O 9.6775993 11.9537076 -14.9983873
C 8.0743326 10.8706348 -17.3888252
H 8.9570019 11.4416081 -17.6791626
H 8.3787829 9.8302572 -17.2664366
C 7.0169986 10.9370419 -18.5172246
H 6.1405201 10.371064 -18.1914962
H 7.409044 10.4393792 -19.4038394
C 6.5974706 12.3719046 -18.896483
O 7.4785217 13.2681533 -18.9084406
O 5.3745041 12.6072441 -18.9936412
N 8.3366279 10.5700175 -13.8495991
H 7.428984 10.1185224 -13.7997984
C 9.2537472 10.4628666 -12.7165948
H 9.411715 11.4704627 -12.3299929
C 10.6243327 9.9036971 -13.153139
O 10.6986872 8.9385328 -13.9069108
C 8.6037136 9.6350831 -11.6036772
H 8.4204313 8.619799 -11.9600852
H 7.655759 10.0896643 -11.3114457
H 9.2647124 9.595986 -10.737355
N 11.703524 10.5367378 -12.6721267
H 11.5296608 11.3030954 -12.0418231

C 13.1142625 10.3142897 -13.0702562
H 13.6516325 11.146085 -12.6173586
C 13.4243279 10.5119185 -14.5675031
O 14.4681525 10.0599575 -15.0339048
C 13.7765064 9.0619071 -12.4395625
H 14.845682 9.122573 -12.6445486
C 13.6265139 9.0675566 -10.9118445
H 12.5803642 8.9423089 -10.6318339
H 14.0018322 10.0070241 -10.506304
H 14.2061243 8.2486758 -10.485005
C 13.2962742 7.7038875 -12.9652798
H 12.2890587 7.4911381 -12.6080123
H 13.9637118 6.9192992 -12.6112727
H 13.2950871 7.7094786 -14.0550769
N 12.605294 11.3349009 -15.2378421
H 11.7369915 11.5814565 -14.7805248
C 12.7314974 11.833814 -16.6111805
H 11.8724211 12.4921825 -16.7495073
C 12.6154246 10.7757957 -17.7237248
O 13.1625965 9.6728211 -17.644985
C 13.9716254 12.728998 -16.7542726
H 14.876663 12.1276992 -16.6566322
H 13.9650252 13.5017634 -15.9855672
H 13.9695782 13.2008787 -17.7373648
N 11.9102878 11.1457732 -18.7984009
H 11.4901366 12.0606529 -18.8311174
C 11.6113386 10.2161504 -19.8849453
H 11.4683386 9.2575284 -19.3922061
C 12.7697246 10.041093 -20.8749029
O 13.4841971 10.9883786 -21.2103005
C 10.2956217 10.5834444 -20.6018401
H 9.5642602 10.8737195 -19.8541547
H 10.4613373 11.4376453 -21.2598264
C 9.7172444 9.4047201 -21.4196476
H 10.350257 9.2350742 -22.2929309
H 8.7362979 9.7002982 -21.7986705
C 9.5899082 8.089336 -20.6183328
O 8.4815808 7.7830257 -20.1380566
O 10.6205886 7.3866881 -20.4498584
N 12.8901662 8.8305339 -21.4204041
H 12.2017952 8.1247666 -21.1470783
C 13.7998552 8.4972353 -22.5092701
H 14.1771403 9.4236569 -22.9434863
C 13.0399482 7.741936 -23.6088019
O 12.2666108 6.8259826 -23.3460136
C 14.9936922 7.7225667 -21.9245386

H 15.5173435 8.3745069 -21.2235556
H 14.6001376 6.8863099 -21.3570609
C 16.0029798 7.189281 -22.9591753
H 15.4878043 6.568367 -23.6890142
C 16.7306364 8.3210255 -23.6896215
H 17.2504768 8.9574053 -22.9726436
H 16.0235264 8.9250527 -24.2541046
H 17.4549132 7.90003 -24.3877077
C 17.0509741 6.3236455 -22.2584349
H 17.5894864 6.9110609 -21.5149428
H 17.7563171 5.9395483 -22.9960493
H 16.5630007 5.4766885 -21.778768
N 13.2962296 8.1160457 -24.8611726
H 13.9627613 8.8548464 -25.0049569
C 12.7297022 7.4720004 -26.0467334
H 11.6552424 7.6561114 -26.0668372
C 12.963355 5.9483119 -26.0268958
O 14.0825531 5.4964434 -25.7932219
C 13.3669937 8.1475173 -27.2671617
H 13.1403425 9.2139322 -27.2266732
H 14.4513269 8.049724 -27.187207
C 12.9656811 7.6189319 -28.6307211
C 12.1494424 8.3971049 -29.4761888
H 11.769383 9.3526089 -29.1454154
C 11.8565891 7.9520163 -30.7798108
H 11.2562024 8.555452 -31.4472005
C 12.3673248 6.7178848 -31.2374061
O 12.1160781 6.2857259 -32.4969706
H 11.5991475 6.9488647 -33.0185918
C 13.1697677 5.9340538 -30.3847744
H 13.5492799 4.9896701 -30.7459872
C 13.4827043 6.3907892 -29.091865
H 14.1273567 5.791667 -28.4621886
N 11.8951892 5.1788313 -26.2599184
H 11.0304658 5.6427895 -26.4800882
C 11.8426092 3.7036502 -26.2137578
H 10.7811114 3.4599108 -26.2631227
C 12.3180866 3.0493723 -24.893297
O 12.588105 1.8486882 -24.8229072
C 12.4769853 3.1028593 -27.4818833
H 13.5610504 3.0598399 -27.3602348
H 12.2582179 3.7523311 -28.3311342
C 11.9188234 1.7098089 -27.8129528
O 10.6734795 1.5608478 -27.7641107
O 12.7303627 0.8270589 -28.169355
N 12.3786583 3.8077385 -23.7892188

H 12.1567416 4.7960764 -23.8565606
C 12.7236866 3.2518266 -22.4780661
H 13.5786598 2.5902866 -22.6181944
C 11.5733328 2.4149249 -21.9209309
O 10.5345585 2.9233827 -21.5059758
C 13.1586237 4.3717878 -21.5322934
H 14.0067916 4.8753226 -21.9927235
H 12.3398627 5.0838206 -21.4074609
C 13.5708913 3.8411374 -20.1519667
H 12.6903613 3.4453154 -19.6452956
H 14.3110593 3.0473288 -20.2626016
C 14.1595742 4.9730826 -19.3074994
H 13.477531 5.8273082 -19.3386531
H 15.1255757 5.265841 -19.7198963
C 14.3246592 4.5251755 -17.8558798
H 14.9647772 3.6393135 -17.8237746
H 13.3343658 4.2459127 -17.4769636
N 14.894404 5.6206571 -17.034387
H 15.8026652 5.9022239 -17.3743342
H 14.9585333 5.3479973 -16.0632593
H 14.281514 6.4323719 -17.0835563
N 11.8145313 1.1129027 -21.796752
H 12.636759 0.7428888 -22.2469702
C 10.915173 0.206206 -21.075752
H 9.9261211 0.3116122 -21.5233934
C 10.7879049 0.6022152 -19.5986625
O 11.7565775 1.0128309 -18.9601126
C 11.3345322 -1.270057 -21.22834
H 10.6296722 -1.8783407 -20.6606041
C 11.2418011 -1.7098514 -22.6941182
H 11.9608367 -1.1650701 -23.3087444
H 10.2395993 -1.5174528 -23.0780593
H 11.4468521 -2.7773516 -22.7753339
C 12.7458793 -1.5768076 -20.7095885
H 13.4907476 -1.0077613 -21.2656222
H 12.9570051 -2.6395233 -20.8284199
H 12.8214824 -1.3296274 -19.6506744
N 9.6090573 0.3641778 -19.0190877
H 8.8636369 0.042936 -19.61778
C 9.3030641 0.4995601 -17.5765515
H 8.2498576 0.2378134 -17.4888618
C 9.3464409 1.9102799 -16.9589934
O 9.2888891 2.018324 -15.7386553
C 10.0341763 -0.545724 -16.7016046
H 9.6693134 -0.4676256 -15.6770594
C 9.7794235 -1.9797159 -17.169852

H 10.215899 -2.1450359 -18.1540049
H 8.7079161 -2.1740235 -17.2032576
H 10.239578 -2.6699839 -16.4624216
O 11.4299275 -0.3613229 -16.691399
H 11.6667592 0.1678767 -17.4727227
N 9.2779939 2.9909397 -17.7450031
H 9.3809 2.8980378 -18.7455964
C 8.8924366 4.3186524 -17.2226461
H 9.484174 4.5459203 -16.3361664
C 7.3950813 4.3363773 -16.8420819
O 6.54833 4.1563548 -17.7157283
C 9.1737252 5.4060425 -18.2724269
H 8.8412563 6.3707971 -17.8859208
H 8.6096819 5.1895781 -19.1816057
O 10.5517975 5.4957348 -18.5958262
H 10.5894134 6.1754387 -19.3178923
N 7.0367162 4.4791013 -15.5515566
H 7.7637049 4.4744686 -14.8485576
C 5.6335974 4.372326 -15.0850094
H 5.0070584 4.6318068 -15.9400818
C 5.0839267 5.3356751 -13.9496445
O 4.1802507 4.8921545 -13.2441249
C 5.3501324 2.8742188 -14.7954481
H 5.6601937 2.2734319 -15.6495461
H 4.2770818 2.7385852 -14.6680957
C 6.0481724 2.3356799 -13.5563096
O 6.9355157 2.9397106 -12.9802825
N 5.6839717 1.1536912 -13.1130988
H 6.1721809 0.8278604 -12.2984982
H 4.9434468 0.6468731 -13.5561149
N 5.4209234 6.6581869 -13.7954432
H 6.1313111 7.0377768 -14.4037847
C 4.6727308 7.6690857 -12.8730958
H 3.6722803 7.2716519 -12.7031805
H 5.1665915 7.6425688 -11.902207
C 4.4605241 9.2667223 -13.2399755
O 5.4811072 9.6663177 -13.812121
N 3.2536409 10.1109433 -12.9872607
H 2.6474233 9.6614132 -12.3206553
C 2.3771526 11.4343133 -13.6922496
H 3.1658733 12.0715482 -14.0792353
C 1.0505375 12.8198719 -12.978889
O 0.738086 12.1140384 -12.0071596
C 1.9540762 10.4352165 -14.9544058
H 0.9329988 10.1097298 -14.746017
C 2.7417697 9.1177858 -15.5124699

H 3.7821969 9.3932306 -15.6797871
H 2.6711718 8.3582525 -14.7352443
C 1.9181844 11.302737 -16.1856385
H 2.9290002 11.590086 -16.4899791
H 1.3394983 12.1952761 -15.9637221
H 1.4171187 10.8031861 -17.0150758
C 2.3461044 8.3062146 -16.7841532
H 1.2838958 8.043668 -16.7411571
H 2.9411495 7.3919643 -16.8207085
H 2.5328117 8.8854332 -17.695951
N 0.2333932 14.4003043 -13.1834479
H 0.7437184 14.7653437 -13.9704046
C -1.2123799 15.955379 -12.9597387
H -1.0028175 16.3292374 -13.960482
C -2.5312399 15.1695335 -13.1010222
H -2.3258053 14.1092915 -13.2484796
H -3.0618878 15.2689053 -12.1555772
C -3.4175086 15.6977339 -14.2585728
H -3.2432872 16.7657087 -14.3985759
C -3.0783778 14.9714564 -15.5675978
H -3.2811317 13.903087 -15.4809576
H -2.0269104 15.1037593 -15.8111044
H -3.6757972 15.3801559 -16.3812018
C -4.9110054 15.512951 -13.9555661
H -5.1400087 14.4586235 -13.8201664
H -5.5040244 15.9052164 -14.7807417
H -5.172629 16.0607214 -13.0540348
C -2.0596333 17.9254177 -12.3430272
O -1.3832877 18.2475852 -11.374691
N -3.271554 19.2249776 -12.5340521
H -3.8017849 18.9469697 -13.3433045
C -4.1853409 20.9390259 -11.9780666
H -4.1201797 20.8378689 -10.8941409
C -5.8142116 21.9798425 -12.0685304
O -6.4577917 21.990306 -13.1238488
C -3.3553264 22.2522131 -12.3368714
H -2.7508435 21.9810248 -13.199742
H -3.9431639 23.1195955 -12.6376299
C -2.3935577 22.6639575 -11.2141059
H -1.5875927 23.2551301 -11.6539897
H -1.9417954 21.7737149 -10.7710014
C -3.1027948 23.4921229 -10.1365626
O -4.1338848 23.0043489 -9.6189633
O -2.6283562 24.6174618 -9.8712537
N -6.4475046 23.0719258 -11.2698861
H -5.8681384 23.316696 -10.4654281

C -7.9445496 23.8116708 -11.2481781
H -8.5313975 23.1447577 -11.8786809
C -8.4505552 25.3183528 -11.7282815
O -7.9562339 26.3541144 -11.2608184
C -8.5827225 23.7319299 -9.8142739
H -8.3098143 22.7644516 -9.3910452
H -8.1237068 24.4929663 -9.1820639
C -10.1128287 23.8553926 -9.699581
H -10.4295997 24.8665053 -9.9608487
H -10.3884539 23.7049408 -8.6526486
C -10.8425833 22.8299823 -10.5754585
O -10.9583405 23.1023724 -11.793048
O -11.2574013 21.7859035 -10.0265513
N -9.602907 25.5151302 -12.4609812
H -10.1970189 24.6947314 -12.6038089
C -10.0093518 26.7968804 -13.139427
H -9.6033406 27.6153906 -12.5465371
C -11.5373224 27.0770783 -13.3129338
O -12.3430523 26.1660776 -13.4651874
C -9.3158021 26.7963794 -14.5218177
H -8.2652178 26.5407625 -14.3723088
H -9.7616841 25.9988297 -15.1187812
C -9.3656862 28.1006173 -15.3396433
H -10.3980501 28.3527717 -15.5746801
C -8.7151959 29.2767986 -14.6053596
H -7.6855865 29.0272619 -14.3458923
H -9.2616884 29.5103425 -13.6946444
H -8.721151 30.1580805 -15.2468748
C -8.6202634 27.902131 -16.6598573
H -7.5714266 27.6723762 -16.4704162
H -8.6904418 28.808201 -17.2612677
H -9.0736317 27.0808258 -17.214682
N -11.9412789 28.3572898 -13.4297045
H -11.2467902 29.0828588 -13.3712445
C -13.3321225 28.7589476 -13.708797
H -13.9922764 28.0787705 -13.1669539
C -13.6973559 28.668285 -15.20173
O -13.2212429 29.4599915 -16.0166546
C -13.5976231 30.1744778 -13.1694646
H -13.3339687 30.2032292 -12.1112647
H -12.9526566 30.8857408 -13.6884859
C -15.0439728 30.6091374 -13.3172145
C -16.0007793 30.2075446 -12.3654889
H -15.7026398 29.6095012 -11.5164327
C -17.3497155 30.5734271 -12.5229045
H -18.084102 30.2600018 -11.7942258

C -17.7459906 31.3401123 -13.632773
H -18.784204 31.6156339 -13.7569758
C -16.7930155 31.742311 -14.5848431
H -17.0984245 32.326439 -15.4416514
C -15.4440982 31.3771052 -14.4276294
H -14.7165289 31.6734974 -15.1713742
N -14.6326318 27.780819 -15.5396291
H -14.9521194 27.1150618 -14.8471413
C -15.2390964 27.6886577 -16.8666391
H -15.735075 28.6312197 -17.0994544
H -14.4624243 27.5134405 -17.6111349
C -16.2666536 26.5596064 -16.957317
O -16.3953891 25.7524785 -16.0401148
N -16.9764725 26.4878918 -18.0819378
H -16.8289854 27.176356 -18.8044725
C -17.9575033 25.4408391 -18.3831007
H -17.7899232 24.5827215 -17.7304426
C -17.7792012 24.9927546 -19.8397808
O -17.7959146 25.8188285 -20.750926
C -19.3742105 25.9740925 -18.1133224
H -19.4072767 26.3626889 -17.0943744
H -19.5911057 26.7896238 -18.8053619
C -20.4418095 24.8784784 -18.2588574
H -20.2170614 24.0779836 -17.5490798
H -20.3816661 24.4589508 -19.2655484
C -21.873747 25.3850101 -18.0153021
O -22.0329252 26.4946208 -17.4568844
O -22.8044396 24.6367793 -18.3888393
N -17.5576422 23.6953514 -20.0649298
H -17.6282623 23.0479037 -19.293315
C -17.3269048 23.1256682 -21.396853
H -17.6295227 23.8524687 -22.1539443
C -18.173276 21.8705255 -21.6133325
O -18.4492201 21.1344351 -20.6665456
C -15.8303326 22.833808 -21.5971231
H -15.5031065 22.1074362 -20.8521036
H -15.2664698 23.755845 -21.4523805
C -15.5235885 22.3038595 -22.9769314
N -16.0573017 22.8083599 -24.1656877
C -15.6125519 22.0028993 -25.140073
H -15.854249 22.1181749 -26.1888167
N -14.8449621 21.02673 -24.6277346
H -14.429788 20.2755673 -25.1579691
C -14.7816493 21.1953241 -23.2621255
H -14.2791179 20.561661 -22.5447457
N -18.5185024 21.5818415 -22.8701808

H -18.1432063 22.1738025 -23.6009505
C -19.2288161 20.3661674 -23.257331
H -19.9116933 20.1067755 -22.4457962
C -18.2431347 19.2015468 -23.4437471
O -17.5034608 19.1367934 -24.4310812
C -20.0753688 20.6536823 -24.5101802
H -20.7622502 21.4721288 -24.2876809
H -19.4111275 20.9769363 -25.3134978
C -20.8899068 19.4416545 -25.0031907
H -20.2061397 18.632868 -25.252551
C -21.8900833 18.9305006 -23.9653095
H -22.5664824 19.7301243 -23.6638583
H -21.3716057 18.543429 -23.0898099
H -22.4721546 18.1090986 -24.387707
C -21.6558409 19.8178908 -26.2704486
H -22.3670983 20.6156827 -26.0549008
H -22.1966695 18.9460329 -26.6408658
H -20.9593117 20.149503 -27.0399621
N -18.305586 18.2264973 -22.5363312
H -18.9848484 18.3084584 -21.7879638
C -17.4987295 17.0022879 -22.56464
H -16.4445225 17.2767669 -22.5352472
C -17.7136686 16.1851165 -23.8479637
O -16.7324037 15.7061853 -24.4089253
C -17.8183622 16.1676056 -21.3118248
H -17.6392762 16.7679502 -20.4189096
H -17.1541192 15.3031549 -21.279748
C -19.2469521 15.6902349 -21.2993199
N -20.3555822 16.5291505 -21.1770985
C -21.409785 15.8002387 -21.5780685
H -22.404937 16.1941265 -21.7444035
N -21.0189659 14.5690479 -21.940586
H -21.5461555 13.999855 -22.611493
C -19.6585309 14.4792161 -21.7690455
H -19.0246639 13.6887216 -22.1457658
N -18.9342631 16.1416122 -24.3967885
H -19.7139306 16.4659898 -23.8383305
C -19.2306073 15.4622993 -25.6698397
H -18.9191364 14.4191048 -25.5943322
C -18.4740459 16.0915514 -26.8531044
O -17.994577 15.3816032 -27.732199
C -20.7422481 15.4827005 -25.9605596
H -20.9140981 15.0341016 -26.9402223
H -21.0845436 16.5175317 -26.0055241
C -21.5715667 14.7083567 -24.9337949
O -21.1731487 13.5796127 -24.57134

O -22.5721805 15.2767049 -24.4473625
N -18.2496141 17.4112082 -26.8323886
H -18.5716361 17.9329869 -26.0288096
C -17.4531297 18.1047217 -27.8489587
H -17.7977212 17.8135617 -28.8424323
H -17.5879439 19.1789559 -27.7304308
C -15.956921 17.7942072 -27.7454374
O -15.2888172 17.6280449 -28.7627028
N -15.439025 17.6201509 -26.5247796
H -16.0609943 17.7043393 -25.7294654
C -14.0743246 17.1266898 -26.3031017
H -13.3826248 17.7038566 -26.9186169
C -13.9256189 15.6534948 -26.724003
O -12.9564171 15.2990751 -27.3953963
C -13.702554 17.3422062 -24.8291262
H -13.7609572 18.4063683 -24.6038623
H -14.431762 16.8391893 -24.1953113
C -12.3186415 16.8515113 -24.4543145
C -11.189853 17.6482056 -24.7279804
H -11.3040053 18.609059 -25.2098121
C -9.9051194 17.1911998 -24.3777829
H -9.0312386 17.792005 -24.5808524
C -9.7460546 15.9368061 -23.7535044
O -8.5019852 15.5094553 -23.4137942
H -8.5099345 14.6569228 -22.9779248
C -10.8775819 15.1370117 -23.4808996
H -10.7593813 14.1747477 -23.0085044
C -12.161335 15.5959257 -23.8337612
H -13.0275541 14.9793858 -23.6353157
N -14.9268359 14.8201123 -26.423841
H -15.7014839 15.176822 -25.8741997
C -14.9828394 13.4197593 -26.8408227
H -14.0769896 12.9181517 -26.4986893
C -15.0310294 13.2711907 -28.3685764
O -14.316041 12.4302505 -28.9112801
C -16.1878629 12.7409781 -26.1724631
H -16.0987507 12.8425083 -25.0898935
H -17.1015123 13.2506904 -26.4752072
C -16.3194695 11.2716038 -26.5208461
C -15.679925 10.2960364 -25.7306851
H -15.1296517 10.5903453 -24.8487413
C -15.7427155 8.9385754 -26.1018896
H -15.2470437 8.1817352 -25.5143806
C -16.4327316 8.5582085 -27.2723703
O -16.4601685 7.2539103 -27.6505408
H -16.8895214 7.1591939 -28.5014005

C -17.087672 9.5346431 -28.0532165
H -17.603424 9.2540987 -28.9595907
C -17.035941 10.887789 -27.6716777
H -17.5222426 11.6367727 -28.2851494
N -15.7781968 14.1209507 -29.0785084
H -16.4213335 14.735133 -28.58639
C -15.8302485 14.0991621 -30.5458371
H -16.023302 13.0723458 -30.857743
C -14.4935853 14.5210283 -31.1785911
O -13.967663 13.81329 -32.0333803
C -16.994066 14.9750367 -31.0424011
H -17.90762 14.6865268 -30.519727
H -16.7819786 16.0183928 -30.8031416
C -17.238316 14.8508303 -32.5555673
H -16.3290961 15.1177248 -33.0992785
H -18.0038016 15.5779021 -32.8375267
C -17.6986865 13.4395358 -32.9540066
O -18.9264795 13.2539991 -33.097649
O -16.8225467 12.5548429 -33.0888592
N -13.837982 15.5729289 -30.6701081
H -14.2892367 16.1201696 -29.9440593
C -12.4898729 15.9648231 -31.1351682
H -12.5346458 16.1337827 -32.2113024
C -11.4694612 14.8360737 -30.9194148
O -10.6875397 14.5295424 -31.8238719
C -12.0510367 17.2817301 -30.4618222
H -12.2135606 17.2075124 -29.385962
C -10.5745282 17.6261432 -30.7001145
H -10.360449 17.6415137 -31.7693089
H -9.9326989 16.8899406 -30.2164525
H -10.3471552 18.6032195 -30.2737645
C -12.8768666 18.454303 -31.009272
H -12.6844 18.5843449 -32.0745506
H -12.6143381 19.3701809 -30.4802161
H -13.9403605 18.2682826 -30.8668087
N -11.5418689 14.1369177 -29.7810758
H -12.2068198 14.4377171 -29.0741423
C -10.7457172 12.934077 -29.5229422
H -9.6880299 13.1653715 -29.6509979
H -10.9128238 12.6231638 -28.4919327
C -11.1015855 11.7534031 -30.4368387
O -10.2052046 11.0525872 -30.9091058
N -12.3806552 11.5703557 -30.7665667
H -13.0669852 12.1981408 -30.3596135
C -12.8830576 10.4904933 -31.6335773
H -12.4536585 9.5465298 -31.299618

C -12.4546029 10.6930136 -33.0856704
O -11.8379689 9.7967565 -33.662114
C -14.4115139 10.3664975 -31.5108397
H -14.8837716 11.3195435 -31.7430037
C -15.0075931 9.281508 -32.405127
H -14.497612 8.3332624 -32.2385487
H -14.9184434 9.5764776 -33.4497101
H -16.0681325 9.1738969 -32.1753859
O -14.7191285 9.9990169 -30.1858573
H -14.6611214 10.8041515 -29.6445041
N -12.6061399 11.9050074 -33.6278135
H -13.1277568 12.5991368 -33.0943308
C -12.1108057 12.2899864 -34.96069
H -12.5626445 11.6330904 -35.7039491
C -10.5924574 12.1086371 -35.0606404
O -10.1118234 11.5017037 -36.0194828
C -12.5253388 13.7419652 -35.2846723
H -12.2047179 14.3950538 -34.4716908
C -11.9014053 14.2583629 -36.5896139
H -12.1485647 13.587198 -37.4129321
H -10.8182307 14.3273054 -36.4922998
H -12.2846337 15.2548982 -36.8119629
C -14.0473735 13.8569901 -35.4397281
H -14.390893 13.2481522 -36.2760117
H -14.3288191 14.8958381 -35.6094773
H -14.5551193 13.5168148 -34.5364626
N -9.8327613 12.5183483 -34.0376294
H -10.2691124 13.0158676 -33.2677802
C -8.3866883 12.2947826 -33.9964652
H -7.94803 12.713808 -34.9032965
C -8.0253687 10.7951613 -33.964585
O -7.1750255 10.3535353 -34.7385592
C -7.808095 13.0498299 -32.7940855
H -8.2322251 12.6640796 -31.8665257
H -8.0425363 14.1119177 -32.8768195
H -6.7249933 12.9258191 -32.7693519
N -8.7383063 9.9928143 -33.1690346
H -9.4399586 10.4216383 -32.5757625
C -8.5261026 8.5373236 -33.0519002
H -7.476906 8.3564513 -32.8190178
C -8.836892 7.7959867 -34.3557814
O -8.0721417 6.9214099 -34.7582541
C -9.3671494 7.960609 -31.9009351
H -10.4270834 8.1175921 -32.1029293
C -9.1210937 6.4714297 -31.6602582
H -8.0569718 6.2924769 -31.5037895

H -9.4663804 5.8883818 -32.5134126
H -9.6722286 6.1524028 -30.7752524
O -9.0287859 8.6085484 -30.6970051
H -9.4300372 9.4946216 -30.711502
N -9.8765834 8.2071453 -35.0894348
H -10.4751493 8.9342993 -34.7066042
C -10.218649 7.6484065 -36.4087586
H -10.3078933 6.565189 -36.3149444
C -9.0961985 7.9031619 -37.4291875
O -8.8107229 7.0319327 -38.2457573
C -11.5883493 8.2058925 -36.8724934
H -11.5800876 9.2895272 -36.7418409
C -11.8649231 7.9080697 -38.3605028
H -11.8182208 6.8336763 -38.5433383
H -11.1330206 8.4106419 -38.9928817
H -12.8465962 8.2794081 -38.6492608
C -12.7194695 7.6055139 -36.002287
H -12.8725628 6.5606242 -36.275951
H -12.4291906 7.6247014 -34.9527788
C -14.0547499 8.3529914 -36.1158443
H -14.469557 8.2558855 -37.1180749
H -13.9111854 9.4081356 -35.8800284
H -14.765007 7.9268607 -35.4065953
N -8.3937191 9.0388911 -37.3367375
H -8.6846259 9.7176446 -36.6427142
C -7.265239 9.3812153 -38.2229573
H -7.5533014 9.1179589 -39.2408252
C -5.9497281 8.6286735 -37.9283472
O -4.9741865 8.7465544 -38.6840646
C -7.0390962 10.899473 -38.1899562
H -8.0014138 11.4136505 -38.2093944
H -6.4800146 11.1924024 -39.0771632
O -6.2990547 11.3175345 -37.05805
H -6.6658562 10.914714 -36.2473093
N -5.8900761 7.8921141 -36.8100548
H -6.7249579 7.8243468 -36.2397125
C -4.6565738 7.3302429 -36.256341
H -3.8362244 8.0007177 -36.5123651
C -4.2800623 5.9408626 -36.8074929
O -3.1352983 5.5205919 -36.6491747
C -4.7709993 7.3240925 -34.7269169
H -5.5236596 6.6026775 -34.4107831
H -5.0474193 8.3167635 -34.3677705
H -3.8122191 7.0434425 -34.2890576
N -5.1800992 5.2659515 -37.5310566
H -6.0659794 5.7270073 -37.7010793

C -4.9863126 3.9392569 -38.1512939
H -5.0038412 3.1815968 -37.3671395
C -3.6467069 3.7571432 -38.8886477
O -2.9731921 2.7386014 -38.7264792
C -6.1697208 3.6740353 -39.1048049
H -7.0402153 3.4328757 -38.4935022
H -5.9499646 2.7970377 -39.7150907
C -6.5637775 4.8197352 -40.0201111
N -5.7699764 5.9207787 -40.3641479
C -6.5674691 6.752385 -41.049963
H -6.277414 7.7318812 -41.4028678
N -7.7926363 6.2244398 -41.1826337
H -8.5854946 6.6964794 -41.5940588
C -7.8129371 5.0107498 -40.5332027
H -8.6720512 4.3747877 -40.3736965
N -3.1845526 4.7824287 -39.6094995
H -3.8397485 5.5490973 -39.7402001
C -1.9094941 4.7790629 -40.3503428
H -1.9058746 3.9210138 -41.0259016
C -0.6755178 4.6000095 -39.4590681
O 0.3500762 4.1523355 -39.9634635
C -1.8108003 6.0655976 -41.1818839
H -1.9802055 6.9149351 -40.5182176
H -0.8101594 6.155225 -41.6088882
C -2.8366963 6.0737128 -42.3295833
H -3.7805566 5.6467642 -41.9998432
H -2.4675716 5.4691415 -43.158521
C -3.090666 7.5062857 -42.7975055
H -2.21557 7.8834898 -43.328123
H -3.2604295 8.1357064 -41.9227019
N -4.2898586 7.6146451 -43.6353146
H -5.0985667 8.0752973 -43.2163844
C -4.3632322 7.5927761 -44.9465972
N -3.4118636 7.0991039 -45.680707
H -3.2172373 7.5904414 -46.5573116
H -2.6560279 6.644648 -45.2205878
N -5.4028833 8.1065841 -45.5234574
H -6.0523492 8.6180683 -44.9202338
H -5.4423454 8.1830156 -46.5185052
N -0.7546542 4.8641941 -38.1528525
H -1.6478001 5.1514024 -37.7665882
C 0.3441051 4.6164829 -37.2169392
H 1.2323201 5.1264964 -37.5885598
C 0.6996833 3.1210416 -37.0906775
O 1.8637562 2.7984179 -36.8587498
C -0.0133202 5.2297426 -35.858726

H -0.9017486 4.748926 -35.4486878
H -0.2065543 6.2971976 -35.9704315
H 0.8169899 5.0909097 -35.1647239
N -0.2423564 2.2117153 -37.3765593
H -1.1831707 2.5200405 -37.5942453
C 0.0441412 0.7786215 -37.4595675
H 0.5673747 0.4724402 -36.5515695
C 0.9552045 0.4341165 -38.6550826
O 1.7906982 -0.4590169 -38.5465271
C -1.2872208 0.0221727 -37.522223
H -1.8167315 0.2672544 -38.4447155
H -1.9079066 0.2944651 -36.668599
H -1.1024213 -1.0504495 -37.4916568
N 0.8781965 1.1977947 -39.7544587
H 0.2102942 1.9555281 -39.7611379
C 1.8112123 1.0873253 -40.8905783
H 2.0039892 0.0301488 -41.0804667
C 3.1532747 1.7347408 -40.5479506
O 4.1918602 1.1211358 -40.7786454
C 1.2242149 1.7004207 -42.1796402
H 1.0712918 2.7691655 -42.0362597
C 2.1680245 1.5163423 -43.3761676
H 2.3959256 0.458749 -43.5151264
H 3.0968456 2.0616945 -43.2111974
H 1.7047064 1.9050683 -44.2829471
C -0.1270984 1.0684292 -42.5415657
H -0.0134202 -0.0094525 -42.6618157
H -0.4995357 1.4948188 -43.4727939
H -0.8607535 1.2658962 -41.7604837
N 3.1497115 2.9075864 -39.8983128
H 2.2584563 3.3646896 -39.7477886
C 4.3829764 3.5769137 -39.4271644
H 5.0316617 3.7532182 -40.287047
C 5.1714679 2.6870662 -38.4572727
O 6.3914124 2.6466122 -38.5342007
C 4.0854947 4.9447377 -38.7734971
H 3.4492565 4.7868126 -37.9038288
C 5.3579813 5.667574 -38.3077099
H 6.0473034 5.7700222 -39.1479344
H 5.8520541 5.0988091 -37.5198768
H 5.1091859 6.6515776 -37.9116408
C 3.3657123 5.9094878 -39.7292705
H 4.0142422 6.146524 -40.5715909
H 3.1117469 6.8290265 -39.2030367
H 2.4524687 5.4645623 -40.1099666
N 4.5099699 1.8691594 -37.6261136

H 3.501018 1.9661057 -37.5679173
C 5.1768327 0.8861925 -36.7517641
H 5.9189905 1.4157616 -36.1524491
C 5.9655105 -0.1805465 -37.5278367
O 6.9655235 -0.6580613 -37.005066
C 4.1151826 0.2654933 -35.821984
H 3.6872668 1.0584642 -35.2070059
H 3.3115988 -0.1409395 -36.4366679
C 4.6026419 -0.8663175 -34.8987054
H 4.9701494 -1.6946528 -35.5041773
H 3.7425868 -1.2253198 -34.3369632
C 5.691348 -0.4487287 -33.8979481
H 5.305648 0.3329929 -33.2411631
H 6.5347315 -0.0299299 -34.4515579
N 6.1852786 -1.5988657 -33.1065998
H 7.0302672 -2.023504 -33.4565587
C 5.6931212 -2.0929878 -31.9796119
N 4.6137617 -1.6238221 -31.4121932
H 4.2496792 -2.0315972 -30.5644291
H 4.1500759 -0.8388203 -31.8268032
N 6.2826626 -3.0960025 -31.392016
H 7.158683 -3.4460252 -31.7437805
H 5.9220268 -3.4384356 -30.5210906
N 5.5130667 -0.5786026 -38.7196586
H 4.6989518 -0.1098716 -39.0946508
C 6.155844 -1.6215128 -39.5477068
H 6.7171868 -2.2989085 -38.9044692
C 7.1870424 -1.0471198 -40.5141252
O 8.1672071 -1.7115576 -40.8379074
C 5.0729423 -2.429844 -40.2970744
H 4.4719049 -1.7392024 -40.8919679
C 5.6505611 -3.4778044 -41.2668926
H 6.3652394 -4.1194898 -40.7503349
H 6.1577817 -2.9777053 -42.0922651
H 4.8537515 -4.0898952 -41.6888069
C 4.1272498 -3.1381766 -39.3001663
H 3.4521743 -2.3981917 -38.8745679
H 3.519856 -3.8567523 -39.8472324
C 4.8022508 -3.8790266 -38.1377464
H 5.1620025 -3.1674568 -37.3956499
H 5.6444629 -4.4611102 -38.5038849
H 4.0797422 -4.5405924 -37.6627072
N 6.9634562 0.1827065 -40.9660073
H 6.1059518 0.645179 -40.6903566
C 7.8918005 0.9019409 -41.8307362
H 8.3560989 0.1968688 -42.5199967

C 9.0541694 1.5083209 -41.0392823
O 10.1751688 1.4175308 -41.5113396
C 7.0910835 1.9372281 -42.6425295
H 6.4755174 2.5141194 -41.9494156
C 8.015055 2.9405889 -43.3353853
H 8.8170607 2.4130698 -43.8517664
H 8.4436482 3.6079035 -42.5859543
H 7.4537262 3.5413332 -44.0376991
C 6.1329575 1.2547909 -43.6511673
H 5.3388 0.7519175 -43.1009127
H 5.6559528 2.0240523 -44.2518133
C 6.7607953 0.2206656 -44.5975052
H 7.0017951 -0.6915304 -44.0477814
H 7.6692296 0.6123557 -45.0487245
H 6.0481605 -0.0291608 -45.3837918
N 8.798992 1.9819869 -39.8195457
H 7.8269675 1.9509433 -39.5448266
C 9.6149082 2.8267209 -38.9312898
H 9.2170332 2.5797365 -37.9459199
C 9.3250434 4.3409519 -39.0306445
O 8.5238253 4.7993794 -39.8542185
C 11.099113 2.4020543 -38.8279642
H 11.7082873 3.1408157 -39.3434734
H 11.2510327 1.4269978 -39.2904088
C 11.5965131 2.3038259 -37.3833269
O 11.4492724 3.3205405 -36.6703128
O 12.1249463 1.2330349 -37.0094698
N 9.8588958 5.1058401 -38.0754761
H 10.516748 4.6466534 -37.4396315
C 9.3325992 6.4041998 -37.6512809
H 8.3526531 6.2042132 -37.2199303
C 9.1395831 7.4234129 -38.7896545
O 10.0494829 7.7492656 -39.5515223
C 10.2024572 6.9760756 -36.5200309
H 10.3373002 6.2059069 -35.7593743
H 11.1814432 7.2466722 -36.9189787
C 9.5601829 8.2096529 -35.8657287
H 8.5933201 7.9247972 -35.443284
H 9.3792816 8.9717041 -36.6261681
C 10.4478246 8.8008571 -34.7647909
O 10.8020372 8.0456266 -33.8314376
O 10.7428137 10.012847 -34.8410345
N 7.9356494 8.0039627 -38.8202964
H 7.2747294 7.6949909 -38.1276181
C 7.4979327 9.066309 -39.7246913
H 6.4475177 9.2219085 -39.4784392

C 7.4607978 8.7250306 -41.2328113
O 7.2182063 9.6195186 -42.0501456
C 8.189382 10.3821855 -39.339387
H 9.2533861 10.3291555 -39.5703739
H 8.0772996 10.5585665 -38.2684137
H 7.7299772 11.2135753 -39.8729389
N 7.5456185 7.4423073 -41.6155564
H 7.7541 6.7354562 -40.9135248
C 7.344726 6.9801426 -42.997353
H 6.9802142 7.8289962 -43.5714683
C 6.2228111 5.9398072 -43.1423331
O 5.8136015 5.298277 -42.1811172
C 8.7040589 6.5417867 -43.5937623
H 9.3769214 6.2789443 -42.781539
H 8.5857406 5.6690598 -44.2357264
C 9.2911549 7.7100855 -44.425368
H 9.1907816 8.6410518 -43.8675988
C 10.7676023 7.5379856 -44.7641775
H 10.9094523 6.6899523 -45.4326336
H 11.347273 7.4053009 -43.8535032
H 11.1408564 8.4293545 -45.2677901
C 8.5546316 7.8484516 -45.7653518
H 8.6226347 6.9085588 -46.3141622
H 9.0244539 8.6267692 -46.3653165
H 7.5122898 8.121164 -45.6183874
N 5.6747873 5.8134553 -44.3622881
H 6.051826 6.3868802 -45.1032308
C 4.5891328 4.8649351 -44.6980872
H 4.5631062 4.1139725 -43.9051263
C 4.8447935 4.0459821 -45.973334
O 3.9017478 3.6498304 -46.6520897
C 3.2105171 5.5603428 -44.666108
H 2.4533252 4.7738203 -44.6831793
H 3.1054279 6.0864467 -43.7153432
C 2.9161759 6.5454592 -45.8179066
H 3.1261552 6.0697121 -46.7735315
H 3.5448542 7.4315362 -45.7241825
C 1.4308664 6.9484155 -45.7776237
H 1.2590212 7.5693356 -44.8966521
H 0.8212512 6.0482975 -45.6740764
N 0.981203 7.6811794 -46.9780385
H 0.7524881 8.6673509 -46.8923419
C 0.6228584 7.177358 -48.1443321
N 0.8186153 5.929421 -48.4685675
H 0.5752193 5.6132175 -49.3870961
H 1.388143 5.3577715 -47.8677134

N 0.0406873 7.949872 -49.0092621
H -0.2933387 8.855375 -48.6610897
H -0.2623347 7.6163466 -49.9005672
N 6.1190758 3.8578277 -46.3209588
H 6.8301691 4.1363819 -45.6635574
C 6.5958218 3.2428633 -47.5626859
H 5.830005 2.5719029 -47.9542416
C 7.8916732 2.4367347 -47.343168
O 8.5668267 2.64766 -46.3401455
C 6.8674228 4.3491416 -48.5851797
H 7.7970464 4.8464069 -48.304794
H 7.032437 3.8770175 -49.5533758
C 5.793543 5.4093771 -48.7466088
C 4.5694425 5.071977 -49.3430479
H 4.3617759 4.0434651 -49.5976387
C 3.6563468 6.0786253 -49.6922932
H 2.7837729 5.8310304 -50.2772817
C 3.9157208 7.4151039 -49.3467661
H 3.2349086 8.196869 -49.6514519
C 5.0976506 7.7468283 -48.6644446
H 5.3156042 8.7854837 -48.4512446
C 6.0470079 6.7471097 -48.3865945
H 6.9997683 7.0245668 -47.966154
N 8.2414517 1.5676536 -48.3001291
H 7.6597218 1.5224626 -49.118932
C 9.3979198 0.6578697 -48.2914649
H 9.1016447 -0.2646165 -47.791358
C 10.6602547 1.1969945 -47.5742744
O 11.1786285 2.245083 -47.9427162
C 9.7087369 0.3224341 -49.75895
H 9.967396 1.234773 -50.3010873
H 8.8483582 -0.1504811 -50.2328802
H 10.5608699 -0.3531544 -49.8204047
N 11.1058984 0.4683517 -46.5404187
H 10.6020477 -0.3756197 -46.3181145
C 12.3498292 0.6411544 -45.7620579
H 12.0842348 0.2321049 -44.78769
C 12.7677959 2.0582833 -45.3295639
O 13.9257997 2.2966503 -44.965395
C 13.488643 -0.2542563 -46.2966031
H 14.4106292 0.3125754 -46.3885813
H 13.233996 -0.5667301 -47.3062372
C 13.7404825 -1.5012609 -45.4212878
O 12.782663 -1.9446545 -44.7479203
O 14.8558588 -2.0616043 -45.4835954
N 11.7676098 2.9221305 -45.1175289

H 10.8663841 2.6705479 -45.5032949
C 11.8434355 4.1137479 -44.2565654
H 10.9702733 4.7131008 -44.4953178
C 13.0474841 5.0013958 -44.5690478
O 13.2566867 5.4146228 -45.7033659
C 11.6839069 3.7215239 -42.7654516
H 10.7042933 3.2772 -42.7433744
C 12.7172875 2.6995277 -42.243958
H 13.7411237 3.021697 -42.4103113
H 12.5627012 1.7344082 -42.7242798
H 12.5786212 2.5252549 -41.1832717
C 11.6078686 4.8907969 -41.7666777
H 12.5088229 4.963613 -41.1569353
H 10.7748965 4.7152793 -41.0967139
H 11.4403098 5.837781 -42.2673589
N 13.7997814 5.3551081 -43.5357305
H 13.5427582 4.9757027 -42.6392564
C 14.9722896 6.1999847 -43.5773004
H 14.7513394 7.1316738 -44.1038787
C 16.1325397 5.5277022 -44.3124657
O 16.971215 6.2121722 -44.8791594
C 15.3363167 6.5059358 -42.1167684
H 14.5993989 7.1982355 -41.7073855
H 15.2826306 5.5845827 -41.5328418
C 16.7149966 7.0988042 -41.9408267
C 16.9780411 8.3984896 -42.403626
H 16.1908624 8.9809401 -42.8654804
C 17.7536204 6.3218351 -41.3924162
H 17.5600729 5.3123153 -41.0611771
C 18.2759221 8.9240865 -42.299878
H 18.4802673 9.9218835 -42.6403416
C 19.0546992 6.8452967 -41.3078177
H 19.8555147 6.2433117 -40.90308
C 19.314826 8.149128 -41.7601691
H 20.3159865 8.5514257 -41.6963891
N 16.2058513 4.196207 -44.3268838
H 15.3924863 3.6449569 -44.0809636
C 17.3365519 3.5317345 -44.9880782
H 18.247824 4.0148928 -44.6328091
C 17.3362498 3.7162578 -46.5202865
O 18.411362 3.6889561 -47.1212283
C 17.4139431 2.0551039 -44.544089
H 17.2640812 2.0186564 -43.4640312
C 18.7685605 1.4097662 -44.8475106
H 18.7772956 0.3945679 -44.4540022
H 19.5699411 1.9849547 -44.3903993

H 18.9200973 1.3641078 -45.9264033
O 16.4573901 1.2286298 -45.1599023
H 15.5577683 1.5969757 -45.0914068
N 16.1818698 4.0690221 -47.1078517
H 15.3465728 4.1192239 -46.5422669
C 16.0387294 4.3659216 -48.539581
H 16.9320402 3.9392428 -48.9945931
C 16.0907108 5.8966401 -49.0803814
O 16.0794458 5.9767522 -50.2908172
C 14.8556147 3.4957892 -49.0773819
H 13.9798023 4.1366509 -49.1933825
H 14.5968292 2.7346661 -48.3378472
C 15.1275989 2.7525254 -50.4024108
O 14.1400315 2.2286328 -50.965854
O 16.307982 2.6299838 -50.8096799
N 16.2900343 7.1238414 -48.4203981
H 16.4497442 7.0131143 -47.4277553
C 16.1613041 8.688626 -48.8951202
H 15.5643498 8.6270493 -49.8051702
C 17.4543756 9.8556596 -49.3389581
O 18.4787587 9.2741032 -49.0634247
C 15.1839224 9.2297703 -47.7945255
H 14.6070166 8.4267525 -47.3302822
H 14.4664933 9.8869038 -48.2885934
C 15.9020718 10.0680588 -46.713811
O 16.3872989 11.1886415 -47.009515
O 15.9809798 9.6471257 -45.537687
N 17.6582507 11.2710311 -50.0187799
H 16.7432841 11.6473397 -50.215925
C 18.9196021 12.4999217 -50.4996121
H 19.4239745 12.7072473 -49.557024
C 19.8910586 11.6284874 -51.3214552
H 19.3040143 10.926469 -51.916006
H 20.4598406 12.2416738 -52.0211191
C 20.8996192 10.8577388 -50.4511839
H 20.4908512 10.6758339 -49.4595413
H 21.799341 11.4600084 -50.3269145
C 21.2793501 9.5187183 -51.0747255
O 22.2065281 9.4215954 -51.8603748
N 20.5592735 8.459898 -50.7803827
H 20.794535 7.5910496 -51.2218482
H 19.7724068 8.5434667 -50.1355936
C 19.092366 14.182846 -51.3913657
O 18.0971829 14.350783 -52.0856275
N 20.1085432 15.3856543 -51.5493382
H 20.9260687 15.212857 -50.986138

C 20.2948739 16.9461427 -52.3131094
H 20.2227503 16.7196393 -53.3770916
C 21.4078328 18.3179152 -52.2406162
O 22.5468109 18.183726 -51.7988277
C 19.0686253 17.7464106 -51.9697146
H 19.2338989 18.2728335 -51.0391949
H 18.2005998 17.0932304 -51.9204808
H 18.8667864 18.4802587 -52.7438545
N 21.1702238 19.6749261 -52.6514902
H 20.2345851 19.8379585 -52.9935248
C 22.0024176 21.0115075 -52.4479166
H 22.8683875 20.7431257 -51.8441611
C 21.2832516 22.1579803 -51.6588977
O 20.098186 22.3880363 -51.8856444
C 22.5113401 21.7661131 -53.7349204
H 21.6390627 22.0773934 -54.3111731
H 23.0283835 22.6774737 -53.4283394
C 23.4507222 21.0203275 -54.6889031
H 22.9115143 20.1707727 -55.1106251
H 23.7236569 21.6915994 -55.5045992
C 24.7266258 20.5101083 -54.0066815
H 24.4530278 19.8220364 -53.2042071
H 25.2791937 21.3544152 -53.5879837
C 25.5984771 19.7681775 -55.0268131
H 25.909443 20.4728439 -55.8051529
H 24.9888066 18.9920389 -55.5006992
N 26.7854235 19.1515691 -54.3842775
H 27.3653918 19.8580064 -53.9512649
H 27.3408378 18.6543916 -55.0695705
H 26.4940335 18.4923601 -53.6723818
N 22.0141776 22.9790716 -50.8776924
H 22.9848334 22.7561847 -50.7481297
C 21.4363151 24.0595014 -50.0406279
H 20.6651534 23.5880877 -49.4217957
C 20.6932301 25.1571828 -50.8434591
O 21.1123859 25.4928957 -51.9501055
C 22.4954675 24.6021902 -49.0576117
H 21.9681595 25.0697176 -48.2281703
H 23.0446883 23.7611733 -48.6298199
C 23.4967813 25.6136439 -49.6395768
H 24.1308012 25.1185841 -50.3761876
H 22.9616084 26.426131 -50.1294037
C 24.3600993 26.2058559 -48.5122502
H 25.0695733 25.4548231 -48.159759
H 23.7129745 26.4715583 -47.6719088
C 25.1007745 27.47991 -48.9414417

H 25.6487684 27.8643149 -48.0751497
H 24.3538345 28.231996 -49.2207631
N 26.033051 27.2429212 -50.0717632
H 26.729265 26.5557402 -49.8134378
H 26.5058074 28.1019389 -50.3270792
H 25.5216444 26.9054137 -50.8778729
N 19.6653497 25.7902966 -50.2502701
C 18.5260935 26.4199977 -50.9302971
H 17.683172 26.2611953 -50.25573
C 17.9666354 25.9230463 -52.2825018
O 17.2120042 26.6858309 -52.8855268
C 18.8245867 27.9163968 -50.8994079
H 19.6064095 28.1600803 -51.6210341
H 17.9311673 28.5198536 -51.0658878
C 19.9982801 26.7186724 -49.1631419
H 19.6497361 26.3469128 -48.204326
H 21.0706298 26.8855675 -49.103357
C 19.3411503 28.0678737 -49.4726293
H 18.5032263 28.2293829 -48.7937525
H 20.0551531 28.8885143 -49.395854
N 18.1444302 24.6585658 -52.7170806
H 18.7575415 24.0313991 -52.2081714
C 17.2497154 24.0944877 -53.7603078
H 17.1809009 24.8305569 -54.5644565
C 15.8108259 23.9536811 -53.2184155
O 15.6116559 23.7733122 -52.0168994
C 17.7905208 22.7718561 -54.367871
H 18.2164828 22.1394345 -53.5915213
H 16.9504542 22.2274091 -54.799801
C 18.8275157 22.9966047 -55.490398
H 19.7490875 23.3914824 -55.0618205
H 18.4292468 23.7618966 -56.1540239
C 19.1471149 21.7322105 -56.331149
H 18.3107319 21.0347568 -56.2604309
H 20.0375569 21.2460875 -55.9308658
N 19.3493069 22.0762304 -57.7577649
H 19.2235844 23.0639289 -58.0004847
C 19.3902273 21.3013719 -58.8329725
N 19.5688706 20.0105413 -58.789513
H 19.5868354 19.4947518 -59.6476506
H 19.6328429 19.5190873 -57.9186205
N 19.2411043 21.8219077 -60.01574
H 18.7989704 22.7503794 -60.07748
H 19.2658129 21.2691169 -60.8598066
N 14.821947 23.9635607 -54.1171727
H 15.0640462 24.1295175 -55.0795782

C 13.4070457 23.7053603 -53.8023649
H 13.2318405 23.8314889 -52.7303655
C 13.0148305 22.2690653 -54.1901151
O 13.2650034 21.8554559 -55.3268438
C 12.5042714 24.70102 -54.554355
H 11.4657565 24.4018961 -54.4176821
H 12.7363243 24.6603459 -55.6166903
C 12.5918951 26.14774 -54.1111584
O 13.1869008 26.5106661 -53.1113018
N 11.9857542 27.0281791 -54.8750885
H 12.0663734 27.990201 -54.6088372
H 11.474667 26.7304553 -55.7012412
N 12.3104354 21.5618859 -53.3039725
H 12.1319232 21.9882772 -52.3959269
C 11.7798987 20.2088839 -53.5218785
H 12.1354009 19.8236533 -54.4750295
C 10.2499869 20.2006078 -53.5304971
O 9.636071 20.6928764 -52.5837385
C 12.2939867 19.2814334 -52.4104324
H 12.0003572 19.7111831 -51.4550013
H 13.3811827 19.2399207 -52.459261
C 11.7331357 17.8531895 -52.451972
H 10.6473428 17.8812916 -52.5280063
H 11.9562163 17.3702826 -51.5054196
S 12.3827862 16.813984 -53.7779941
C 13.8931229 16.2105975 -52.9728887
H 14.4627188 15.602152 -53.6752431
H 13.6285852 15.6018071 -52.1075691
H 14.5036321 17.0516099 -52.6454799
N 9.6561252 19.476022 -54.4810838
H 10.2356146 19.1014098 -55.228018
C 8.2618632 19.0294284 -54.4110718
H 7.7432147 19.5950109 -53.6411374
C 8.2012683 17.5367186 -54.0391612
O 8.6091906 16.6916482 -54.8315145
C 7.5848191 19.3377223 -55.7563497
H 7.6533027 20.404727 -55.9472126
H 8.1407942 18.8235216 -56.5354205
C 6.1094319 18.9161937 -55.8471269
H 6.0223168 17.8621598 -55.5987058
C 5.2008171 19.7105303 -54.9111302
H 5.2842959 20.7773402 -55.1160026
H 5.4789999 19.5191252 -53.8783082
H 4.1706631 19.3882301 -55.0433692
C 5.5958432 19.1157678 -57.2735893
H 5.6369121 20.1715894 -57.5396105

H 4.5696247 18.7588489 -57.3505647
H 6.2159663 18.5490014 -57.9673902
N 7.650806 17.207511 -52.8705797
H 7.2809473 17.9542867 -52.2913677
C 7.3710433 15.8279812 -52.4464654
H 8.0060151 15.1508264 -53.0126345
C 5.9027388 15.4793458 -52.7611271
O 4.9709185 16.075372 -52.2065813
C 7.75986 15.6405981 -50.9667627
H 7.1203425 16.2573012 -50.3370099
H 8.7862314 15.9905422 -50.8370111
C 7.6824443 14.1863395 -50.4760692
O 6.8708591 13.4090086 -51.019936
O 8.4087115 13.8653462 -49.5055914
N 5.6864177 14.5711102 -53.7189983
H 6.4974395 14.0713565 -54.0777662
C 4.3447916 14.1842873 -54.1843736
H 3.6345714 14.9349283 -53.8406842
C 3.9351687 12.8544228 -53.5605881
O 4.3232699 11.7831128 -54.0192536
C 4.2336867 14.1575708 -55.7201635
H 4.8568936 13.3737071 -56.1278179
C 2.7974381 13.8629456 -56.1479142
H 2.1426664 14.6095088 -55.703318
H 2.4969621 12.8683791 -55.8186253
H 2.7229068 13.8922377 -57.2343704
C 4.6461988 15.4961607 -56.3443168
H 4.0765308 16.3124174 -55.897042
H 4.4772019 15.482627 -57.4198195
H 5.7094427 15.6583488 -56.1699027
N 3.0257182 12.9205146 -52.5909151
H 2.717312 13.8404901 -52.2952782
C 2.6511322 11.7941466 -51.7378099
H 1.5919382 11.8736081 -51.495824
H 2.826334 10.8515972 -52.2505211
C 3.4378837 11.764253 -50.4292475
O 3.7783718 10.6859284 -49.9559292
N 3.6725557 12.9262538 -49.8125931
H 3.3218807 13.7634739 -50.2589932
C 4.5639542 13.1282033 -48.6574106
H 5.5763021 12.8435635 -48.9560907
C 4.2394702 12.317265 -47.379758
O 4.9944892 12.3378076 -46.4012804
C 4.557135 14.6320349 -48.3562011
H 4.9178299 15.1702423 -49.2356233
H 5.2327209 14.8285762 -47.5267004

S 2.8894722 15.2074225 -47.9074698
H 2.7061254 14.4032963 -46.8479299
N 3.0995178 11.6227276 -47.3420058
H 2.5537956 11.6005599 -48.1895828
C 2.6973692 10.7695608 -46.2284903
H 1.7296098 10.3306215 -46.4382888
H 3.4252333 9.964224 -46.1301078
C 2.5964225 11.5335775 -44.9096616
O 2.0756425 12.6458024 -44.8617237
N 3.1475683 10.9405853 -43.8409104
H 3.5995258 10.050913 -43.9813798
C 3.2644505 11.5676352 -42.5110625
H 2.5363332 12.3721322 -42.4922009
C 4.6024855 12.3215117 -42.2989558
O 4.9330438 12.7035656 -41.1837733
C 2.7685976 10.6219858 -41.3876643
H 3.329892 9.6866489 -41.3966698
H 2.9110664 11.1133101 -40.4234859
C 1.2482459 10.3497499 -41.5881163
H 0.7919812 11.3134804 -41.7977461
H 1.1127473 9.7337692 -42.4784474
C 0.4227669 9.7161131 -40.4391496
H 0.7314226 10.1512193 -39.4901884
H 0.5960085 8.639396 -40.4025918
C -1.0758004 10.0217816 -40.7095582
H -1.41602 9.4220592 -41.5561837
H -1.1388727 11.0716698 -41.0127608
N -2.0107233 9.8625097 -39.5598994
H -2.0996365 8.9127489 -39.2334567
H -2.9406933 10.1949484 -39.8135329
H -1.7309402 10.4529809 -38.7839952
N 5.3030928 12.6456133 -43.3932679
H 4.9509657 12.3158047 -44.2839729
C 6.263606 13.7539353 -43.4685514
H 6.339384 14.0373157 -44.5168193
H 5.8664254 14.6025745 -42.9108326
C 7.6966603 13.5217271 -42.9844836
O 8.5229441 14.4189115 -43.1446022
N 8.052298 12.3272621 -42.5032093
H 7.3432049 11.6119669 -42.3818921
C 9.4128024 12.0200965 -42.040962
H 9.6646061 12.6654394 -41.1981862
H 9.4474178 10.9834761 -41.7123813
C 10.481707 12.1834795 -43.1192423
O 11.5222019 12.7773601 -42.8556136
N 10.1750316 11.7843984 -44.3589289

H 9.2846575 11.3332788 -44.4955374
C 11.0186339 12.0413754 -45.5425943
H 11.9980506 11.588156 -45.3790969
C 11.2701527 13.5389522 -45.7486745
O 12.4236188 13.9518113 -45.8260238
C 10.4258214 11.3832015 -46.8083101
H 10.5905208 10.3099959 -46.7269139
C 8.9312833 11.6033541 -47.0463739
H 8.7099762 12.6632397 -47.1442538
H 8.3393666 11.1827496 -46.2344072
H 8.6361123 11.1131843 -47.9758533
O 11.0895781 11.8306891 -47.9597574
H 10.484578 12.3844576 -48.4701324
N 10.2252136 14.371186 -45.663101
H 9.3074535 13.9722939 -45.5446904
C 10.2857366 15.8236971 -45.8547703
H 10.7235008 16.0265557 -46.8268567
C 11.1106349 16.537419 -44.7650356
O 11.8863183 17.4526382 -45.0535196
C 8.8657689 16.3970441 -45.8856242
H 8.8986116 17.4030429 -46.3069571
H 8.4867539 16.4648006 -44.8668808
S 7.7163401 15.3759861 -46.8510702
H 8.4533831 15.1086649 -47.9512639
N 10.9632892 16.0999518 -43.508704
H 10.2825801 15.3646959 -43.3383485
C 11.7666912 16.5933009 -42.3777083
H 11.7210304 17.682167 -42.3558897
C 13.2337723 16.2149077 -42.5646649
O 14.1041332 17.0794444 -42.4631432
C 11.2181531 16.0596538 -41.0385742
H 11.0788588 14.9803609 -41.1088303
C 12.1583111 16.3454275 -39.8608403
H 12.413162 17.4044263 -39.8365978
H 13.0713173 15.7583224 -39.9544092
H 11.6747709 16.0658247 -38.9240883
C 9.8619149 16.7101419 -40.7311641
H 9.9738266 17.7906836 -40.6408162
H 9.465485 16.313601 -39.7953368
H 9.147258 16.4904463 -41.5236473
N 13.5086787 14.9554922 -42.9189931
H 12.7374487 14.300883 -43.0121126
C 14.8616324 14.4600437 -43.1763215
H 15.4605874 14.5795315 -42.2714439
C 15.5717593 15.257738 -44.2626702
O 16.6984701 15.6980688 -44.0520232

C 14.8119003 12.9696725 -43.5454865
H 15.47017 12.778969 -44.3885111
H 13.8159203 12.6800994 -43.8734669
C 15.2350372 12.1081566 -42.3887418
N 16.5503927 11.9810507 -41.9354251
C 16.4753185 11.2196844 -40.8299497
H 17.3253826 10.9192111 -40.2306378
N 15.2029599 10.8627996 -40.5836198
H 14.890693 10.2906702 -39.8085624
C 14.4037633 11.4233475 -41.5538219
H 13.3257702 11.3738953 -41.6274226
N 14.9078435 15.5075468 -45.387301
H 13.989292 15.091668 -45.5152898
C 15.5112271 16.2350314 -46.5011762
H 16.4944499 15.8005517 -46.6810838
C 15.788866 17.702243 -46.1455768
O 16.9395239 18.1374501 -46.2012692
C 14.6897345 16.0179734 -47.7882799
H 15.1327128 16.6236197 -48.5603665
C 14.7908711 14.5234165 -48.182398
H 14.1962195 13.8987777 -47.5198436
H 15.8220997 14.1781548 -48.1262874
H 14.4372692 14.3716629 -49.2011021
C 13.2366266 16.4975189 -47.7320752
H 12.7646062 16.0195718 -46.8976035
H 13.2059957 17.5711979 -47.5585925
C 12.4407379 16.1837724 -49.0033466
H 12.2244151 15.1164961 -49.062049
H 13.0189562 16.48687 -49.8741816
H 11.4970485 16.7230964 -48.9888032
N 14.8187234 18.4245126 -45.5793613
H 13.9008259 18.0120409 -45.4473034
C 15.0314858 19.8172963 -45.1857533
H 15.4248899 20.3646793 -46.0422485
C 16.0654803 19.9815299 -44.052507
O 16.775709 20.9872676 -44.0197683
C 13.6739593 20.4215669 -44.8288207
H 13.2487701 19.8884107 -43.9775321
H 12.9997992 20.3346726 -45.682005
H 13.7950387 21.4777533 -44.588073
N 16.1945128 19.0031707 -43.1417661
H 15.5550477 18.2125874 -43.1960906
C 17.1426889 19.0652184 -42.0165262
H 17.2682905 20.115006 -41.7540704
C 18.5555244 18.5806105 -42.3633946
O 19.5022617 19.2022987 -41.8920977

C 16.5356863 18.3408313 -40.8014493
H 15.5546198 18.7709715 -40.5929376
H 16.3868088 17.2899072 -41.0560764
C 17.3783728 18.4041283 -39.5137768
H 18.2761985 17.8007418 -39.655826
H 16.803368 17.9420041 -38.7110996
C 17.8159365 19.8120997 -39.0658109
H 18.4907804 20.2290409 -39.816017
H 18.3932875 19.7217441 -38.1440225
N 16.6897875 20.7491832 -38.8609668
H 16.5665419 21.4502419 -39.5716769
C 15.8807065 20.8193544 -37.8169131
N 14.9853566 21.7589121 -37.7435133
H 14.3645694 21.8061651 -36.956886
H 14.8726153 22.4170928 -38.4946404
N 15.9472023 19.9843426 -36.8189475
H 16.6241288 19.2455779 -36.8502505
H 15.2949912 20.0477172 -36.0589826
N 18.7300829 17.5290883 -43.1764286
H 17.9068943 17.0488226 -43.5306661
C 20.0684893 17.0390531 -43.5746573
H 20.7065707 16.9291063 -42.6964889
C 20.7865705 18.0002679 -44.5366372
O 22.0134416 18.0053269 -44.5586972
C 19.9813346 15.699411 -44.3219547
H 19.304658 15.8747004 -45.1616404
H 20.9671976 15.5209637 -44.7569981
C 19.5628823 14.3578569 -43.6784448
H 20.3290906 14.0366348 -42.971993
H 18.6129878 14.4437749 -43.1562454
C 19.4543242 13.3380914 -44.844005
H 20.418487 13.3271226 -45.354375
H 18.7167883 13.694725 -45.5669937
C 19.1362141 11.8730891 -44.4980398
H 19.490443 11.6480878 -43.4892399
H 19.686472 11.2395421 -45.2001729
N 17.6955314 11.5486037 -44.6343123
H 17.1272324 12.0169392 -43.9401613
H 17.4806865 10.5531329 -44.5600195
H 17.3202713 11.7308229 -45.5705452
N 20.0478271 18.6624719 -45.4364741
H 19.046923 18.5093921 -45.436498
C 20.6358853 19.3309775 -46.6085273
H 21.6749389 19.5607893 -46.3684505
C 20.0541746 20.7091617 -46.9622581
O 20.3152965 21.2210728 -48.0515675

C 20.6868561 18.354292 -47.7957786
H 21.0473326 18.879063 -48.6814314
H 21.4566581 17.6177888 -47.5613333
C 19.4373287 17.602644 -48.1667651
C 19.394892 16.1940631 -48.1271936
H 20.196105 15.6434021 -47.6560693
C 18.3451897 15.5113149 -48.7728262
H 18.2923324 14.4372359 -48.7484014
C 17.3896775 16.2497781 -49.5044677
O 16.5031357 15.6745726 -50.3589183
H 16.8175994 14.8393012 -50.723428
C 17.3695605 17.6457992 -49.4114359
H 16.6398896 18.1806343 -49.9827419
C 18.3848271 18.32481 -48.7295873
H 18.4231516 19.3990316 -48.7319403
N 19.3517895 21.3539171 -46.024161
H 19.1651048 20.8775033 -45.1539639
C 19.0384741 22.7911583 -46.0750765
H 18.3539854 22.9771147 -45.2457682
C 18.2660689 23.2374827 -47.3416898
O 18.3498463 24.3844502 -47.7803859
C 20.3254224 23.5929801 -45.7682459
H 21.0190659 23.4837652 -46.6030665
H 20.8007875 23.1682607 -44.8823023
C 20.0831759 25.0814136 -45.500918
O 19.0655738 25.4065578 -44.8420731
O 20.8655128 25.9146396 -46.0124582
N 17.4548563 22.3556176 -47.9295338
H 17.4238769 21.424189 -47.5465261
C 16.4823397 22.7281121 -48.9710723
H 16.9464183 23.4477338 -49.6473765
C 15.24281 23.3977265 -48.3725054
O 14.9522023 23.2457856 -47.1836795
C 16.0422905 21.5162724 -49.8130752
H 15.2603537 21.8499514 -50.4920367
C 17.2011516 21.0558416 -50.7125561
H 18.1143336 20.9147098 -50.1404907
H 17.4026366 21.8357731 -51.4439333
H 16.943333 20.1436236 -51.2459316
C 15.398919 20.4182088 -48.9405732
H 16.1665696 19.8238888 -48.4571155
H 14.7955406 20.8598812 -48.151009
C 14.4493492 19.5350595 -49.7478417
H 14.8862927 19.2531794 -50.7036183
H 13.5268763 20.0795328 -49.9194258
H 14.2201392 18.6408149 -49.1834476

N 14.4360211 24.010578 -49.2375354
H 14.7402083 24.0446147 -50.2085445
C 12.9993631 24.158699 -49.0076405
H 12.8175786 24.2573771 -47.9397338
C 12.2662528 22.9046082 -49.5103378
O 12.6107608 22.3634176 -50.5604784
C 12.5234879 25.4517901 -49.6859483
H 13.0406288 26.283309 -49.2048236
H 12.830272 25.44303 -50.731898
C 11.0112581 25.7338392 -49.6079951
H 10.8575365 26.7986213 -49.7828324
H 10.658697 25.5126149 -48.6002629
C 10.1424015 24.9839405 -50.6247752
O 10.5934761 24.3664138 -51.5757024
N 8.8434225 24.9954737 -50.4307784
H 8.2650384 24.4828781 -51.0906918
H 8.4331033 25.5005896 -49.6701504
N 11.2359635 22.4621827 -48.7844512
H 11.0578115 22.9049529 -47.8891113
C 10.3429926 21.383246 -49.214284
H 10.5949631 21.0920705 -50.235015
C 8.8711568 21.8264889 -49.2272665
O 8.3637704 22.3460571 -48.2286926
C 10.5732244 20.1618264 -48.309774
H 10.3332883 20.4190762 -47.2777976
H 11.6227969 19.8672222 -48.3609254
S 9.5400098 18.7650291 -48.8429061
H 8.3402798 19.3259352 -48.6493615
N 8.1565084 21.4877271 -50.299685
H 8.6580875 21.1050509 -51.0997332
C 6.6915622 21.4723115 -50.3796701
H 6.2622286 21.8728319 -49.4665579
C 6.2201094 20.031677 -50.5303368
O 6.4399128 19.4158451 -51.5664035
C 6.1943466 22.3420754 -51.5434893
H 6.7074003 22.060723 -52.4640156
C 4.6826819 22.2560971 -51.7615391
H 4.1511206 22.4652032 -50.8329504
H 4.408085 21.2625488 -52.117419
H 4.3832472 22.9787312 -52.5207642
O 6.4597851 23.693421 -51.2481149
H 6.0035097 23.908674 -50.423727
N 5.5750189 19.4858299 -49.5003679
H 5.4489048 20.0421054 -48.6609263
C 4.8934822 18.1915638 -49.5707935
H 5.3805521 17.5583986 -50.3075649

H 4.9561912 17.6920084 -48.6044006
C 3.4178491 18.3381219 -49.9451647
O 2.7499778 19.2457229 -49.4508098
N 2.8759735 17.4192718 -50.7447628
H 3.489359 16.728848 -51.172119
C 1.4218424 17.2768095 -50.9304672
H 0.9352328 17.8212694 -50.1261785
C 0.9560668 15.8305307 -50.7697592
O 1.6406314 14.8966804 -51.1867693
C 0.9136146 17.8905378 -52.2565359
H -0.1448096 17.6360649 -52.3435319
C 1.6387051 17.3248981 -53.5004349
H 2.6843131 17.6363338 -53.4916327
H 1.6106378 16.2363657 -53.4623999
C 0.9961788 19.4249524 -52.1858548
H 2.0365676 19.7521869 -52.1926341
H 0.5247676 19.7783184 -51.266841
H 0.4642133 19.8755679 -53.0218273
C 1.0052803 17.7519525 -54.8316522
H 1.1088672 18.8260369 -54.9781292
H -0.0515642 17.481976 -54.8436029
H 1.510511 17.2397663 -55.65089
N -0.2477491 15.6495421 -50.2260329
H -0.7465903 16.4668413 -49.8863459
C -0.9656272 14.3704334 -50.212057
H -0.6470092 13.784003 -51.0743246
C -2.4736961 14.6046495 -50.3361702
O -2.9923013 15.6108318 -49.855645
C -0.6345655 13.5626088 -48.9469876
H -1.035592 14.063649 -48.0667914
H 0.4483293 13.4718768 -48.8486547
O -1.2039403 12.2647033 -49.0661658
H -0.8230701 11.6633302 -48.3915009
N -3.189996 13.6624306 -50.9545782
H -2.693636 12.8593069 -51.3085801
C -4.6570956 13.7038033 -51.1109514
H -4.9453094 14.6681594 -51.5324737
C -5.3659256 13.5807981 -49.7505522
O -6.4750307 14.0756237 -49.5707605
C -5.0838402 12.5810381 -52.0946599
H -4.5868338 11.656628 -51.7934535
C -6.60297 12.3174183 -52.0927149
H -7.1427278 13.2367823 -52.3255887
H -6.9278242 11.951214 -51.1188279
H -6.8636599 11.5560362 -52.8261546
C -4.6265523 12.9470809 -53.5247128

H -5.2699191 13.744181 -53.8956398
H -3.6030739 13.3215718 -53.5000957
C -4.6555746 11.788384 -54.5300713
H -5.6777838 11.458972 -54.7113472
H -4.0625365 10.9539342 -54.1543668
H -4.2321241 12.1260136 -55.4769672
N -4.7271601 12.913471 -48.788543
H -3.775143 12.6272275 -48.9723444
C -5.2999645 12.5558781 -47.4919011
H -6.3581933 12.3426238 -47.6238231
C -5.12327 13.6827605 -46.4548777
O -3.9915082 13.9538372 -46.0403971
C -4.6452439 11.2446981 -47.0565748
H -3.587913 11.2809028 -47.3174804
H -5.1004324 10.4159779 -47.6001787
O -4.7366918 11.0128571 -45.6723862
H -5.6758702 11.0025864 -45.3613888
N -6.2181431 14.2734505 -45.9265974
C -6.1448261 15.2535177 -44.8392723
H -5.5037262 16.0768951 -45.1469994
C -5.5994107 14.6593758 -43.5365247
O -4.990551 15.3639187 -42.7414758
C -7.5767455 15.7683423 -44.6357585
H -8.056633 15.2366621 -43.8112928
H -7.5919532 16.8425494 -44.4489233
C -7.6005661 14.1249832 -46.3585443
H -7.6892693 14.0050037 -47.4364082
H -8.051605 13.2737373 -45.845865
C -8.2932888 15.4153499 -45.9362787
H -8.111043 16.1900229 -46.680453
H -9.3633916 15.2698172 -45.7853592
N -5.798928 13.3603836 -43.3244477
H -6.3334307 12.8472974 -44.0199305
C -5.3819279 12.5909009 -42.1533687
H -5.5086175 13.2194111 -41.2710266
C -3.889537 12.1953296 -42.1819728
O -3.2267231 12.2061379 -41.1417984
C -6.3412377 11.3905832 -41.9922851
H -7.3113502 11.7703061 -41.6675729
H -5.9575394 10.7498656 -41.1971473
C -6.5663179 10.5388814 -43.2538189
O -6.908747 11.078591 -44.3329274
O -6.3508923 9.3107568 -43.1812859
N -3.3008267 11.9683216 -43.3619274
H -3.8866016 11.9470974 -44.1933319
C -1.8430178 11.9831623 -43.5618261

H -1.3723892 11.257173 -42.9150736
C -1.2618897 13.3502526 -43.1982805
O -0.4067809 13.4546671 -42.3135894
C -1.4776396 11.6187214 -45.0103698
H -2.1218696 12.1563712 -45.7031098
H -0.4492902 11.9202012 -45.2048059
C -1.5826308 10.1109484 -45.2530965
H -2.542401 9.7508069 -44.8866851
H -0.8029225 9.6052696 -44.6769365
C -1.4265517 9.7523301 -46.7342513
O -2.1887154 8.9014817 -47.2378949
O -0.4387212 10.1870761 -47.3704002
N -1.8238071 14.3981459 -43.7998905
H -2.5652821 14.2183388 -44.4716944
C -1.3629664 15.7743681 -43.644951
H -0.2966211 15.7971953 -43.8619811
C -1.5011144 16.2671664 -42.1931932
O -0.5669668 16.8774303 -41.6842395
C -2.0937697 16.6432986 -44.6956286
H -3.1616424 16.4331521 -44.6208048
C -1.6453572 16.3062192 -46.1415492
H -1.8060816 15.2459139 -46.3228022
H -2.2857106 16.8442621 -46.8391984
C -1.9169274 18.1412204 -44.4519846
H -0.8624135 18.3648704 -44.4776171
H -2.351247 18.4235725 -43.4927015
H -2.4173335 18.7036014 -45.2397571
C -0.1850993 16.6247081 -46.5047295
H 0.502714 16.0852862 -45.8550906
H 0.005957 17.6948585 -46.4390428
H -0.0012286 16.3148047 -47.5317343
N -2.5671139 15.9256739 -41.4689525
H -3.3267024 15.4447242 -41.9411177
C -2.7735512 16.3191514 -40.0675664
H -2.7786111 17.4067291 -40.0129945
C -1.6436206 15.8404344 -39.1348382
O -1.1884872 16.6049745 -38.2795479
C -4.1712491 15.8244989 -39.6523631
H -4.8964345 16.3669418 -40.2600162
H -4.276831 14.7663523 -39.8970248
C -4.5547229 16.0502673 -38.1793378
H -5.6347657 16.1922723 -38.1337433
H -4.0852274 16.9595111 -37.8033673
C -4.2106249 14.87867 -37.2602295
O -3.478791 13.9624486 -37.5900405
N -4.7524033 14.838826 -36.0651956

H -4.5608134 14.0108061 -35.5281391
H -5.3835633 15.5549238 -35.7602172
N -1.1004638 14.6424705 -39.3708469
H -1.5013084 14.0764096 -40.106404
C 0.0418472 14.1322373 -38.6116889
H -0.0436926 14.4501244 -37.5713118
C 1.3850099 14.6518156 -39.1239408
O 2.2660308 14.8908551 -38.3035319
C 0.0435491 12.6104591 -38.6527647
H 1.0465421 12.2488474 -38.4138007
H -0.2262749 12.3017169 -39.6598596
S -1.1117245 11.9445062 -37.4241918
H -2.0986728 12.845829 -37.6105998
N 1.5575739 14.8170369 -40.4417058
H 0.8054631 14.5690666 -41.0762493
C 2.7701594 15.422793 -40.987119
H 3.6444812 14.9045317 -40.5883586
C 2.8696392 16.8863013 -40.5636503
O 3.5847836 17.2487937 -39.6283336
C 2.7907183 15.3258836 -42.5238653
H 1.9156234 15.7985274 -42.9702768
H 2.7822393 14.2994419 -42.8465516
H 3.7036186 15.7788831 -42.9060338
N 2.1002966 17.7180647 -41.2791163
H 1.1970245 17.3057366 -41.4945638
C 2.6117307 18.5517644 -42.3861192
H 3.1341734 17.8723516 -43.0516016
C 1.3970127 19.0600589 -43.2242848
H 1.358215 18.4262558 -44.0974844
H 0.5018309 18.6692998 -42.7733033
C 1.1166488 20.5727843 -43.5444475
H 1.2915895 21.151315 -42.6635274
H 1.8411256 20.9728381 -44.2381746
C -0.3230066 20.9360951 -43.9914732
H -1.0227159 20.3720331 -43.3859128
H -0.4699022 20.6178342 -45.0179328
C -0.6963571 22.4298742 -43.8025919
H 0.1995555 22.9755746 -43.4898631
H -1.4260614 22.4979107 -42.9928553
N -1.2635441 23.0865246 -45.012343
H -0.5895041 23.0906335 -45.7804691
H -1.4992876 24.0597524 -44.8902245
H -2.0834365 22.6400856 -45.4285437
C 3.7642777 19.4606364 -41.9383207
O 4.6934606 18.9814363 -41.3095185
N 3.6311882 20.7771193 -41.9989438

H 2.8876535 21.1417414 -42.5604266
C 4.0788258 21.6000937 -40.8865894
H 5.1237588 21.3834285 -40.6673247
C 3.9999911 23.0820359 -41.3327063
H 3.0493725 23.2473995 -41.8409777
H 4.7944901 23.2717171 -42.0553756
C 4.1026739 24.1179475 -40.2399803
N 3.0918861 25.0331692 -39.9409579
C 3.4223691 25.5589047 -38.7527186
H 2.8010558 26.2448663 -38.192991
N 4.5752577 25.0358408 -38.3042514
H 4.8930887 25.1113105 -37.3429746
C 5.0364284 24.1390869 -39.2428601
H 5.8685618 23.4582498 -39.1352582
C 3.2420288 21.199852 -39.6385361
O 3.0797663 22.025223 -38.7500341
N 2.530232 20.0383135 -39.640792
H 2.865367 19.2416506 -40.1623869
C 1.4823966 19.8353562 -38.7041374
H 1.2481924 20.7922532 -38.2384218
C 2.0858318 19.0933861 -37.5361898
O 3.067777 19.6165891 -37.0050026
C 0.1413169 19.4928001 -39.3442083
H 0.1442248 18.4588372 -39.6817178
H -0.6141341 19.569777 -38.5611991
C -0.267998 20.4123699 -40.5175301
H 0.1183068 19.9509151 -41.3678312
C -1.7776922 20.4353344 -40.70414
H -2.2492346 20.8376265 -39.8066394
H -2.1313518 19.4171078 -40.8620015
H -2.0806715 21.041301 -41.5528518
C 0.1944014 21.8747087 -40.5369619
H 1.2344464 22.003532 -40.7787416
H 0.035205 22.3050513 -39.5470494
H -0.3843236 22.4742456 -41.2283432
N 1.4878269 18.020571 -37.0275607
H 0.680745 17.619781 -37.490007
C 1.8370793 17.567107 -35.682473
H 1.6480608 18.3923588 -34.994206
C 3.3255548 17.1909504 -35.5105984
O 3.975249 17.6990186 -34.5966676
C 0.897452 16.4265003 -35.2938252
H 1.0844826 15.5580817 -35.9233497
H -0.1416476 16.7397962 -35.4098183
H 1.0723946 16.150676 -34.2533891
N 3.895938 16.3935091 -36.4204664

H 3.3273487 16.0340792 -37.1822979
C 5.2891756 15.9425756 -36.3358269
H 5.4806631 15.6036171 -35.3165186
C 6.3276383 17.0397494 -36.6405866
O 7.4910534 16.9063308 -36.2670223
C 5.4618095 14.744717 -37.274927
H 5.2733635 15.0489023 -38.3063018
H 4.7617187 13.9547081 -37.0015401
H 6.47971 14.3605827 -37.1961688
N 5.9285687 18.1328717 -37.2982429
H 4.962632 18.1795496 -37.6016175
C 6.8314934 19.2418052 -37.635999
H 7.8429605 18.8505207 -37.6832907
C 6.844892 20.3829303 -36.6121256
O 7.5699517 21.3645417 -36.8076774
C 6.4883151 19.7654549 -39.0249614
H 5.4763035 20.1549587 -38.9918722
H 6.5452826 18.940506 -39.7349417
O 7.3909096 20.7802827 -39.4271918
H 7.5897174 21.3108912 -38.6363837
N 6.0301942 20.3258281 -35.5583977
H 5.4633655 19.4990513 -35.4131305
C 5.9102691 21.4436743 -34.6242259
H 5.8569889 22.3420848 -35.2375565
C 7.1639527 21.593333 -33.7492034
O 7.6308825 20.6298769 -33.149975
C 4.6035516 21.3149437 -33.8160339
H 3.9610644 20.5586151 -34.2705869
H 4.8261419 20.983383 -32.8007128
C 3.8146552 22.6339158 -33.7630863
H 4.3761121 23.361526 -33.1761685
H 2.8597389 22.454182 -33.2696363
C 3.5503843 23.2164719 -35.1512038
O 4.0029425 24.3035214 -35.4804174
N 2.9488639 22.4629638 -36.0411139
H 2.950617 22.7735293 -37.0102779
H 2.7918841 21.4817724 -35.8719266
N 7.7210904 22.8081187 -33.6914249
H 7.2987774 23.5567177 -34.2190106
C 8.9972279 23.0788989 -33.0131334
H 9.0701395 24.1482844 -32.8145984
H 9.0214978 22.5526099 -32.0586359
C 10.2493908 22.6746477 -33.805537
O 11.3286693 22.5814544 -33.2268775
N 10.1271567 22.3925806 -35.1093067
H 9.2142375 22.4390954 -35.5407744

C 11.2828804 22.1739636 -35.9765919
H 12.1470555 21.9182832 -35.3639204
C 11.6311631 23.4425166 -36.7703543
O 10.8468171 23.8739616 -37.6062429
C 11.0248787 21.0051895 -36.9419587
H 10.2282802 21.2982624 -37.627223
H 11.9359448 20.8776523 -37.5163873
C 10.6521745 19.6372191 -36.3484215
H 9.6886744 19.7031776 -35.8480046
C 10.554009 18.6256642 -37.4922608
H 11.5421073 18.4392663 -37.908598
H 9.9033941 19.0025286 -38.2786216
H 10.1402736 17.6890333 -37.1141196
C 11.700169 19.1190278 -35.363519
H 12.6761902 19.0615675 -35.8409176
H 11.4049478 18.1266817 -35.0192001
H 11.7434391 19.7788958 -34.4960605
N 12.8948299 23.8661966 -36.7091582
H 13.3352564 23.7065076 -35.812184
C 13.4155394 25.0179326 -37.4790076
H 12.6681205 25.8145446 -37.4282954
C 13.6732908 24.7387133 -38.9836209
O 14.3679382 25.4938641 -39.6582383
C 14.7076514 25.5493859 -36.8171534
H 15.5330047 24.8870948 -37.0856383
H 14.9254605 26.5347873 -37.2302733
C 14.686988 25.6625318 -35.2845744
H 15.5866248 26.1902592 -34.9596257
H 14.7320807 24.6561241 -34.8586402
C 13.4378158 26.3886491 -34.7758366
O 13.4280188 27.6368107 -34.819975
O 12.4900572 25.6633847 -34.4019596
N 13.2648524 23.5772005 -39.5105643
H 12.5354699 23.085907 -39.0068225
C 13.5570177 23.1795007 -40.8976494
H 14.5254876 23.6009869 -41.1761233
C 12.542468 23.7627457 -41.8860466
O 11.3445727 23.7642826 -41.6133993
C 13.6717175 21.6505914 -41.0511484
H 14.6737933 21.3409764 -40.7633607
H 13.5409867 21.3814689 -42.0977122
C 12.6807254 20.8409029 -40.234541
O 13.0819143 19.9571612 -39.4998137
N 11.4063889 21.1652132 -40.2593715
H 10.7663929 20.6171536 -39.7086994
H 11.0835354 21.9909526 -40.7547637

N 13.0133136 24.1123106 -43.0888082
H 14.003369 23.9861169 -43.252692
C 12.2559559 24.8063065 -44.14478
H 11.634147 25.5703974 -43.6725806
C 11.2900693 23.8636724 -44.8836361
O 11.4794884 23.5296958 -46.0528063
C 13.2402964 25.5207442 -45.0935189
H 13.7974318 24.7585126 -45.6299273
H 12.6747639 26.1075278 -45.8195301
C 14.2367 26.4457212 -44.363787
H 13.6782029 27.2076148 -43.8182901
H 14.8081668 25.8651489 -43.6389009
C 15.2530414 27.1445263 -45.2802328
H 15.9662655 27.6665972 -44.6388995
H 14.7410762 27.8844194 -45.8974409
C 16.0017683 26.1590706 -46.1895801
H 15.4383225 26.0365783 -47.1185131
H 16.0485386 25.1791219 -45.7028254
N 17.3833513 26.6013149 -46.4784672
H 17.4501689 27.5257558 -46.8646949
H 17.8394313 25.9286781 -47.0934372
H 17.9672793 26.5200239 -45.6378488
N 10.2586788 23.4106932 -44.1767312
H 10.1841574 23.7178249 -43.2124954
C 9.2046284 22.5279462 -44.6902692
H 9.4646982 22.2318761 -45.7066837
C 7.8664666 23.2561846 -44.773848
O 7.5173305 24.0732073 -43.9268036
C 9.0728533 21.2239852 -43.8778855
H 8.316634 20.6091786 -44.3642886
C 10.3766276 20.4230278 -43.8853184
H 11.1563743 20.9672999 -43.3545985
H 10.6901088 20.2389319 -44.9118237
H 10.2204614 19.4614051 -43.3955968
C 8.6444678 21.4437578 -42.4225151
H 9.3296493 22.1190396 -41.9159669
H 8.6253233 20.491972 -41.8909169
H 7.6444986 21.8751438 -42.3818422
N 7.0848025 22.9020958 -45.7835561
H 7.4542566 22.2468272 -46.4660422
C 5.688535 23.2901281 -45.9603944
H 5.2592379 23.5642908 -44.9971347
C 4.9368154 22.090787 -46.5162855
O 5.5175932 21.2799915 -47.2402207
C 5.572196 24.4861637 -46.9093264
H 6.1315988 25.3254024 -46.4927753

H 4.5253632 24.7775338 -47.0057321
O 6.0920421 24.166681 -48.1936715
H 6.9019288 23.6412406 -48.069159
N 3.6649475 21.9328174 -46.1572492
H 3.1924113 22.6509743 -45.6290061
C 2.8540294 20.891737 -46.7718369
H 3.2603949 20.7420939 -47.768523
C 1.4232672 21.3469321 -46.9993605
O 0.9655742 22.2461308 -46.2958749
C 2.8723023 19.5311039 -46.0727033
H 1.8998189 19.3593649 -45.6756641
H 2.831625 18.8431272 -46.9167144
C 4.0238134 18.9905664 -45.2169582
C 4.3071869 17.6253821 -45.3860567
H 3.6888653 17.0224533 -46.0309537
C 4.8919789 19.744078 -44.3979068
H 4.7530479 20.7996534 -44.2590226
C 5.3992881 17.0235059 -44.752768
H 5.5957283 15.9801922 -44.9285487
C 5.9619852 19.1264943 -43.7177794
H 6.5828412 19.700868 -43.0462872
C 6.2154413 17.761582 -43.8888187
H 7.0295482 17.2877361 -43.3583586
N 0.7049211 20.6843602 -47.9071269
H 1.1580329 19.9917536 -48.4999263
C -0.7426341 20.8387991 -48.0115702
H -1.0824204 20.9641876 -46.9898499
C -1.5361428 19.617803 -48.4852348
O -0.9924186 18.6595985 -49.0384563
C -1.11455 22.1121096 -48.8003637
H -1.7026334 21.8618073 -49.6844732
H -0.2187602 22.6483505 -49.119513
C -1.9431406 22.9948283 -47.8556361
O -2.8487189 22.4398679 -47.1824651
O -1.4945848 24.1167787 -47.540539
N -2.8489941 19.6723543 -48.2388324
H -3.2118883 20.5386921 -47.8451067
C -3.8093695 18.7281656 -48.8164363
H -3.3973647 17.7249245 -48.7310293
C -3.9734597 19.0752603 -50.2936677
O -4.3973474 20.175993 -50.6415666
C -5.1732061 18.7435119 -48.0992768
H -5.6730347 19.6950951 -48.2825975
C -6.0637773 17.6068255 -48.6193746
H -5.6052813 16.639463 -48.4108445
H -6.2151491 17.7031978 -49.6940242

H -7.0395154 17.6576898 -48.1394967
C -5.0359728 18.5600012 -46.5826229
H -4.4763312 17.6507279 -46.373113
H -6.0235974 18.4938287 -46.1264681
H -4.5200769 19.4177832 -46.1531065
N -3.6215795 18.1436875 -51.1741785
H -3.2879382 17.2507786 -50.8282856
C -3.6115106 18.3903012 -52.6077545
H -4.5005105 18.9612831 -52.879691
H -2.7342712 18.9845956 -52.864576
C -3.5764643 17.1062873 -53.4192378
O -2.760003 16.2192524 -53.1788795
N -4.4347018 17.0516582 -54.4326559
H -5.0732818 17.8204822 -54.5581563
C -4.3864658 16.05468 -55.4956002
H -4.0915978 15.0912157 -55.0883004
C -3.3390088 16.4662952 -56.5310674
O -3.4495113 17.5140222 -57.164842
C -5.7890218 15.9175663 -56.0942179
H -6.1264015 16.9003061 -56.4288134
H -6.4684305 15.5787826 -55.3090581
C -5.884318 14.9468145 -57.2730016
O -4.8383242 14.4918336 -57.7921059
O -7.0322555 14.7422459 -57.7062923
N -2.3237965 15.6294866 -56.7222922
H -2.3348741 14.7515165 -56.2290766
C -1.1959614 15.917138 -57.5968154
H -0.8042434 16.9026035 -57.3349828
C -1.5477935 15.9520276 -59.0940706
O -0.7986989 16.5414193 -59.8719076
C -0.1261968 14.8713849 -57.2991538
H -0.4332321 13.8964363 -57.6832249
H 0.0211865 14.8034629 -56.2217764
H 0.8044678 15.171372 -57.7763393
N -2.6894224 15.3899282 -59.5099429
H -3.3057519 14.9434825 -58.8268714
C -3.1680469 15.5081525 -60.8889374
H -2.2977069 15.4764396 -61.5426236
C -3.8714604 16.8578481 -61.1353749
O -3.8881519 17.3427345 -62.2717246
C -4.0589673 14.2899356 -61.2019821
H -3.5800509 13.3944349 -60.8022451
H -5.0094949 14.4145144 -60.6792547
C -4.3336983 14.054691 -62.7013104
H -4.8066073 14.9365606 -63.1323418
C -3.0607759 13.732287 -63.4923323

H -2.5395826 12.8897156 -63.037984
H -2.3987076 14.5952794 -63.5160859
H -3.3198034 13.4669686 -64.5171777
C -5.2831101 12.8667808 -62.868187
H -4.8269732 11.9632398 -62.4652122
H -5.5106869 12.7195113 -63.9246257
H -6.2135557 13.0676634 -62.3361046
N -4.3438173 17.5317856 -60.0767708
H -4.239743 17.0913676 -59.1689351
C -5.1392953 18.7670931 -60.1508346
H -5.3101043 19.014173 -61.1954728
C -4.4652452 20.0192241 -59.553186
O -4.7247452 21.1197762 -60.036646
C -6.5275032 18.4864972 -59.5562995
H -7.1041324 19.4099779 -59.5149897
H -6.4332717 18.0886791 -58.546576
C -7.2894454 17.5033969 -60.4354308
O -7.5563357 17.7986585 -61.5883171
N -7.5704357 16.2932746 -60.0008698
H -8.0536807 15.6856149 -60.632247
H -7.3300548 15.9419432 -59.0695682
N -3.5291447 19.8666245 -58.6129897
H -3.3778296 18.9328279 -58.2480178
C -2.7161973 20.948246 -58.0426512
H -3.3974747 21.6988151 -57.6398965
C -1.8381347 21.6244684 -59.108357
O -1.3603305 20.9679296 -60.0376461
C -1.8708616 20.3801509 -56.8857777
H -1.2155043 19.5960797 -57.2688558
H -2.5403692 19.9339514 -56.1488536
C -1.0113161 21.4298223 -56.1713507
H -0.2443703 21.7912918 -56.8567282
H -0.5059511 20.9491715 -55.3327881
S -1.9402459 22.8506813 -55.5394919
C -0.5701828 23.8292679 -54.8690719
H -0.9613234 24.74235 -54.4197375
H 0.1193026 24.0926016 -55.6715556
H -0.0410525 23.252321 -54.1098706
N -1.6015408 22.9369568 -58.9811233
H -1.9423219 23.4054305 -58.1486339
C -0.8192637 23.7390643 -59.9395053
H -0.2576772 23.064157 -60.5851844
C 0.2282464 24.5957342 -59.2270807
O 0.004776 25.0706294 -58.1183271
C -1.7509353 24.6002576 -60.822006
H -2.2485006 25.3361505 -60.1875853

H -1.1470927 25.1432575 -61.5512182
C -2.8355023 23.8041089 -61.576147
H -3.4692309 24.5095959 -62.1148217
H -3.4670243 23.293783 -60.85025
C -2.285634 22.7794283 -62.5838831
H -1.469829 22.2166229 -62.1362081
H -1.8727529 23.3160784 -63.440655
N -3.3480238 21.8600974 -63.0454371
H -4.1592275 22.2924263 -63.4494333
C -3.3712553 20.539867 -62.9367282
N -4.3632524 19.8552986 -63.436589
H -4.394121 18.8635073 -63.2050823
H -5.2093213 20.3214733 -63.7056537
N -2.4470361 19.8412767 -62.3456859
H -1.8072108 20.3142319 -61.7116381
H -2.5589807 18.8412342 -62.2636225
N 1.3477479 24.8147152 -59.9086339
H 1.3980397 24.4714973 -60.8561952
C 2.4801481 25.6382775 -59.4817666
H 2.171737 26.310206 -58.6796175
C 2.9540436 26.4820188 -60.6742369
O 2.5493109 26.2124622 -61.8048482
C 3.6101897 24.7291168 -58.9715134
H 3.9556652 24.0994097 -59.7921348
H 4.4519802 25.3525014 -58.6710504
C 3.2414508 23.8502424 -57.790578
C 3.2255769 24.3988016 -56.4940977
H 3.4578802 25.4439975 -56.3432923
C 2.944317 22.484486 -57.9805326
H 2.957988 22.0603603 -58.9749413
C 2.9118273 23.5871558 -55.3886793
H 2.893409 23.9989979 -54.390721
C 2.6376701 21.6672874 -56.8746023
H 2.4174114 20.6212776 -57.0198648
C 2.6196309 22.2218223 -55.5761608
O 2.3179465 21.4526177 -54.4997302
H 2.18435 20.5319836 -54.724826
N 3.8334992 27.4628261 -60.4509498
H 4.2082822 27.584594 -59.5219213
C 4.4939213 28.1620494 -61.5616058
H 3.7309422 28.5488971 -62.2384471
C 5.3882736 27.2020676 -62.3485732
O 6.0176283 26.3114741 -61.7616727
C 5.3217809 29.3477141 -61.0436929
H 6.091829 28.9980095 -60.3569351
H 4.6612574 30.029898 -60.5071673

O 5.9393037 30.0538829 -62.1032955
H 6.7243398 29.5613144 -62.4093461
N 5.5820288 27.4940815 -63.6317522
H 5.0106585 28.1958685 -64.0746187
C 6.7307802 26.9874338 -64.3760478
H 6.6366855 25.9085016 -64.4254029
C 8.028407 27.3558119 -63.6304401
O 8.1187891 28.4400932 -63.0417593
C 6.7479946 27.5450176 -65.8100117
H 7.719706 27.3149662 -66.2505881
H 6.6485008 28.6315395 -65.7731873
C 5.6672195 26.9697377 -66.7358325
O 4.772257 26.2412801 -66.2508656
O 5.7522914 27.2543315 -67.9504335
N 9.0046514 26.4438149 -63.5827792
H 8.8767 25.5757138 -64.0999271
C 10.2928452 26.6834173 -62.9168859
H 10.9628395 25.863104 -63.1596956
H 10.726691 27.5994604 -63.3176861
C 10.2581107 26.8168125 -61.3836782
O 11.1929088 27.3706979 -60.812188
N 9.2068629 26.3463782 -60.7000323
H 8.4713915 25.8971 -61.2280263
C 9.0731089 26.4374344 -59.2283582
H 9.1851045 27.4791121 -58.9258279
C 10.1069491 25.6136815 -58.4432941
O 10.4836519 25.994552 -57.3288833
C 7.6897912 25.9495096 -58.7796666
H 7.6527518 25.9341602 -57.6888603
H 7.5223822 24.9374057 -59.148062
O 6.6580577 26.7898418 -59.2502181
H 6.5249451 26.5892104 -60.1998395
N 10.5092653 24.4553591 -58.9775317
H 10.148396 24.2047745 -59.8925566
C 11.2703098 23.4359545 -58.2511685
H 11.53296 23.831252 -57.2725145
C 12.5826537 23.0539197 -58.9343996
O 12.7237566 23.0892687 -60.153064
C 10.4016771 22.1945978 -58.0069762
H 10.1678104 21.7295215 -58.9665107
H 10.9725225 21.4720044 -57.4234963
C 9.1174901 22.4918067 -57.2616053
C 9.1440574 22.7882535 -55.8861933
H 10.0770687 22.7663086 -55.3446341
C 7.8965917 22.5106125 -57.9559699
H 7.8822044 22.2996273 -59.0114969

C 7.9532069 23.1130515 -55.2127899
H 7.9771022 23.3415634 -54.1554797
C 6.7052395 22.8277978 -57.284616
H 5.7704859 22.841606 -57.8236995
C 6.7337942 23.135257 -55.9133376
H 5.8189192 23.3828901 -55.3936062
N 13.517583 22.5839715 -58.1188466
H 13.3072881 22.5350507 -57.1278463
C 14.8179288 22.0601079 -58.5444263
H 15.0314648 22.3351386 -59.5799209
C 14.8820061 20.5283609 -58.4571595
O 15.7122647 19.8905221 -59.1081171
C 15.861854 22.6970743 -57.6264017
H 15.4958654 22.6849266 -56.6000881
H 16.7664742 22.1004187 -57.6568063
C 16.1785143 24.1454572 -58.0345006
H 16.4935678 24.6931106 -57.143149
H 15.2907343 24.6425045 -58.4312527
C 17.305602 24.1476694 -59.067901
O 17.1190339 23.5958259 -60.1692975
O 18.44763 24.4781666 -58.6847995
N 14.0008482 19.9431131 -57.6415347
H 13.375073 20.5442361 -57.11928
C 13.8557196 18.5135042 -57.3961515
H 14.3000178 17.9505765 -58.2132188
C 12.3562485 18.1809218 -57.2916426
O 11.6055412 18.9392478 -56.6695756
C 14.5613807 18.1437631 -56.0764476
H 13.9703084 18.5856413 -55.2756477
H 14.5331341 17.0604963 -55.9537592
C 16.0075358 18.6382877 -55.8749913
H 16.0774433 19.7022195 -56.0961422
C 16.4272853 18.4705187 -54.4170553
H 16.4803628 17.4129 -54.1558347
H 15.7119752 18.972793 -53.7650525
H 17.3987076 18.9361346 -54.2814031
C 17.0207935 17.8886108 -56.7338335
H 16.9321508 16.814618 -56.5697651
H 18.0321975 18.2073195 -56.4892937
H 16.8337156 18.1120355 -57.7796173
N 11.9268476 17.0254245 -57.802589
H 12.5667476 16.4679935 -58.3580903
C 10.6134922 16.4408486 -57.4727742
H 10.200647 16.9817986 -56.6240596
C 10.8460628 14.9974708 -57.0338666
O 11.4968357 14.2196056 -57.7343886

C 9.584129 16.5453119 -58.623281
H 9.8891 15.8453796 -59.384544
C 9.5544193 17.9798466 -59.2141413
H 9.2019235 18.6905423 -58.4697116
H 10.5636784 18.2729503 -59.500348
C 8.1844474 16.0604285 -58.2049168
H 7.7911132 16.6907992 -57.4099126
H 8.2346799 15.0332518 -57.8434264
H 7.5072373 16.083636 -59.0554359
C 8.7221937 18.1222675 -60.4791129
H 7.6661354 18.1114382 -60.2270789
H 8.9737782 17.3097286 -61.1536164
H 8.9770545 19.0592215 -60.9698104
N 10.3564225 14.6574374 -55.8469035
H 9.7941865 15.3397182 -55.3446681
C 10.5610505 13.361721 -55.2088406
H 11.1527113 12.721799 -55.8623133
C 9.2265463 12.6740554 -54.9607565
O 8.2716464 13.3041168 -54.5053329
C 11.33476 13.5316808 -53.9000093
H 12.3374437 13.8914798 -54.1303004
H 10.8329836 14.2807638 -53.2847405
C 11.4560679 12.2414929 -53.107116
C 12.2584577 11.1893237 -53.5913009
H 12.8093437 11.3079028 -54.5115507
C 12.3285074 9.9712448 -52.892086
H 12.935959 9.1595666 -53.2719667
C 11.6002187 9.80203 -51.7022888
H 11.6597189 8.8679635 -51.1608939
C 10.7885537 10.8448729 -51.2227153
H 10.2173261 10.7141468 -50.3149552
C 10.7115946 12.0603749 -51.9250526
H 10.0647863 12.8469327 -51.5572318
N 9.1602094 11.3777451 -55.2613215
H 9.967209 10.9333375 -55.6892098
C 7.9328766 10.5948167 -55.1189933
H 7.301889 11.0701356 -54.3646456
C 8.255057 9.1831792 -54.6477118
O 9.0891373 8.509012 -55.2435789
C 7.1597415 10.5440188 -56.4471444
H 7.7114147 9.9222563 -57.1494066
C 5.8095002 9.8629732 -56.2113163
H 5.4896513 10.0212756 -55.1882349
H 5.9140703 8.7885214 -56.3956002
H 5.0585275 10.2781186 -56.8705113
C 7.0334598 11.9183157 -57.1168015

H 6.7794035 12.6694729 -56.3779092
H 6.2773547 11.9109892 -57.8926942
H 7.9869524 12.1928344 -57.5734035
N 7.5304832 8.6922801 -53.6430702
H 6.8760449 9.3036927 -53.1726615
C 7.6681163 7.3252057 -53.1358409
H 8.2068826 6.7406371 -53.8801015
C 6.2876119 6.6673452 -52.9862792
O 5.4097723 7.1751835 -52.2984167
C 8.5623854 7.3067255 -51.8731256
H 9.5579647 7.6240205 -52.1890165
C 8.094785 8.2976582 -50.7891014
H 7.0661867 8.0877915 -50.5071326
H 8.1462975 9.3195321 -51.1659953
H 8.7354529 8.2405992 -49.9112774
C 8.6896144 5.865025 -51.3446308
H 7.7397634 5.5693468 -50.9079072
H 8.8984866 5.198318 -52.1818034
C 9.804598 5.6643442 -50.3096542
H 9.5898272 6.2220642 -49.3977556
H 10.7584064 5.9929064 -50.7211156
H 9.8773923 4.6049462 -50.0589894
N 6.0735872 5.5791504 -53.7322323
H 6.8322139 5.3057652 -54.3480851
C 4.8587987 4.7465541 -53.7745885
H 4.9169297 4.1923917 -54.7089597
C 3.516118 5.5057891 -53.8614048
O 2.5191161 5.1118456 -53.2532807
C 4.8607276 3.7088065 -52.6438668
H 4.6254463 4.2273965 -51.7263386
H 4.0580549 2.9937707 -52.8329263
C 6.1573545 2.9237375 -52.4338794
H 5.9667513 2.1123083 -51.7318169
H 6.9089625 3.5794581 -51.9982039
C 6.7097361 2.3381834 -53.7201912
O 7.8204849 2.6367907 -54.124086
N 5.9596121 1.5408023 -54.4434662
H 6.3860314 1.1601171 -55.2659575
H 4.9443976 1.5731702 -54.3602218
N 3.4922299 6.609794 -54.6151327
H 4.3734201 6.9183661 -54.9925454
C 2.3102518 7.4619524 -54.7883182
H 1.523502 7.1422596 -54.1042785
C 1.7272705 7.3825057 -56.209906
O 0.5213259 7.2247492 -56.3878293
C 2.7044591 8.890572 -54.3960494

H 3.4554001 9.2689719 -55.0839756
H 3.1257726 8.8895343 -53.3873539
S 1.267988 9.9956678 -54.455393
H 1.9581013 11.1116626 -54.1764562
N 2.5785643 7.3956426 -57.2408166
H 3.5667076 7.4519611 -57.0465178
C 2.1412431 7.4998195 -58.6453044
H 1.3706166 8.2684934 -58.6961616
C 1.455654 6.224998 -59.1639819
O 0.6846533 6.2812467 -60.1185169
C 3.3271499 7.9979964 -59.498981
H 4.2271993 7.4604443 -59.1935633
C 3.5120954 9.5030321 -59.2234801
H 2.7433385 10.070109 -59.7525158
H 3.3888159 9.7067547 -58.1618482
C 3.1360484 7.7830679 -61.0060698
H 2.1735433 8.1823495 -61.3276595
H 3.2063335 6.7193164 -61.2171017
H 3.9295564 8.2650958 -61.5743967
C 4.8911054 10.0097662 -59.6504475
H 5.0099305 9.9628237 -60.7314375
H 5.6735898 9.4236341 -59.1685517
H 4.9797716 11.0455733 -59.3443017
N 1.5717582 5.1041732 -58.4564899
H 2.1827045 5.1206104 -57.6529454
C 0.7328439 3.9149034 -58.651388
H 0.8947388 3.5462889 -59.6639284
C -0.7823536 4.1812482 -58.5331897
O -1.5480582 3.5308124 -59.2486143
C 1.1982313 2.8047712 -57.6992863
H 0.5310809 1.9524817 -57.8022436
H 2.1807305 2.4791663 -58.0396875
C 1.2976814 3.1948038 -56.2119626
H 0.3012642 3.1688227 -55.7657597
H 1.6944627 4.205053 -56.0979221
C 2.2384522 2.2254211 -55.4939914
O 3.4720441 2.4194285 -55.590334
O 1.7426194 1.2099946 -54.9540701
N -1.2002262 5.2068174 -57.7761104
H -0.5129244 5.7123035 -57.2254629
C -2.598939 5.6605511 -57.6700557
H -3.2650489 4.7962077 -57.7183515
C -3.0249164 6.6303356 -58.7861373
O -4.212663 6.7435325 -59.0897442
C -2.8084363 6.3407486 -56.3074555
H -3.8525083 6.6461592 -56.2269157

H -2.1951503 7.2411331 -56.2540386
C -2.494111 5.4648105 -55.1222584
N -3.2748369 4.3995001 -54.6813414
C -1.402258 5.5724542 -54.3123075
H -0.6083186 6.2963122 -54.4092074
C -2.6439835 3.8919381 -53.61534
H -2.9894624 3.0311948 -53.0587202
N -1.5076655 4.5702295 -53.3754836
H -0.8044458 4.3204002 -52.6953078
N -2.0794667 7.3391896 -59.4136958
H -1.1158875 7.1556197 -59.1719981
C -2.3616997 8.3287098 -60.4667779
H -3.2276859 8.9173499 -60.1602128
C -2.7464924 7.5875773 -61.7548098
O -2.001621 6.7304906 -62.2333305
C -1.169923 9.3091347 -60.6337605
H -0.2399088 8.7535995 -60.5132491
C -1.1908147 10.461597 -59.5993055
H -0.3002541 11.0742561 -59.749503
H -2.0610367 11.0934907 -59.7853356
C -1.1248122 9.968303 -62.0256172
H -2.0519598 10.5058078 -62.2273489
H -0.9656117 9.221869 -62.8030621
H -0.2929621 10.6711498 -62.0761009
C -1.2165483 10.0459058 -58.1225757
H -0.4028053 9.3553923 -57.9105932
H -2.1699243 9.5788426 -57.8756608
H -1.0975652 10.9321865 -57.4986943
N -3.9191623 7.9002296 -62.3131057
H -4.4943561 8.5906527 -61.8567087
C -4.4811881 7.1647627 -63.4544377
H -4.2521094 6.1048541 -63.3300885
C -3.8496852 7.5951677 -64.7889643
O -3.2727482 6.7640869 -65.4867586
C -6.0132403 7.3088003 -63.4796484
H -6.2694267 8.3537053 -63.6622205
H -6.4039576 6.7184002 -64.3096062
C -6.7166646 6.8608593 -62.1861027
H -7.7887787 7.0203155 -62.3018223
H -6.3872767 7.4812947 -61.3521355
C -6.4970237 5.3889815 -61.8459751
O -6.9995508 4.4963847 -62.5114945
N -5.7572628 5.0902401 -60.8042215
H -5.833909 4.14822 -60.4307188
H -5.3482376 5.8097848 -60.2156199
N -3.8569681 8.8981386 -65.0946658

H -4.3202539 9.5439063 -64.4785248
C -3.1401507 9.4583446 -66.2477274
H -3.0411782 8.6833109 -67.0107969
C -1.7236527 9.8698317 -65.8282051
O -1.4618108 10.9868417 -65.364135
C -3.9305454 10.5951731 -66.9132344
H -4.0322571 11.4261952 -66.2131145
H -4.9304066 10.2295566 -67.1538309
C -3.2659619 11.0967388 -68.2087631
O -2.0957811 10.7292587 -68.4770125
O -3.9030893 11.919111 -68.8986723
N -0.803794 8.9111941 -65.9612056
H -1.116004 8.0117478 -66.302686
C 0.6148988 9.1220086 -65.6716635
H 0.701243 9.6227611 -64.7065084
C 1.2656105 10.0741342 -66.6689175
O 2.098196 10.8648421 -66.2441968
C 1.3529267 7.7757301 -65.5926566
H 1.1126767 7.1689474 -66.4679944
H 2.4297305 7.955249 -65.5762918
C 0.9574017 7.0366377 -64.3062496
H 1.2856483 7.6316669 -63.4540904
H -0.1280695 6.9419732 -64.2625537
C 1.5717334 5.633152 -64.2128231
H 2.6407791 5.6991469 -64.4166036
H 1.1128656 4.9710833 -64.9484462
C 1.3819517 5.0770151 -62.7971826
H 1.7874912 5.8138986 -62.1066302
H 1.9538723 4.1524789 -62.6816544
N -0.0351696 4.8430507 -62.4374898
H -0.6226182 5.656999 -62.6128357
H -0.1159741 4.6848988 -61.4413045
H -0.410129 4.0257107 -62.9061462
N 0.857613 10.1083443 -67.9375912
H 0.0306158 9.5949737 -68.2147783
C 1.4627276 11.0419099 -68.8908006
H 2.5495178 10.9536548 -68.8456466
C 1.0955327 12.4971915 -68.5945729
O 1.9570158 13.368256 -68.6695282
C 1.0240044 10.7936445 -70.3246517
H -0.0383867 11.0268319 -70.4208665
H 1.5987591 11.5133366 -70.9013528
C 1.2870139 9.4334093 -70.9591885
H 0.6207275 8.6875652 -70.5204886
H 2.321262 9.1555211 -70.7495981
C 1.0670261 9.5267931 -72.4818624

O 1.5837226 8.6346019 -73.1863309
O 0.449561 10.5278113 -72.9382914
N -0.1651418 12.7909911 -68.2489103
H -0.8538143 12.0316837 -68.2624408
C -0.597046 14.1400191 -67.8500139
H -0.3606733 14.8368522 -68.6493023
C 0.1788132 14.6128788 -66.6331437
O 0.6875716 15.7306213 -66.6664212
C -2.1126775 14.1466946 -67.5941196
H -2.6227997 14.0571914 -68.5530931
H -2.3576696 13.272807 -66.9910062
C -2.6896842 15.3612165 -66.8459386
H -3.7652708 15.2087699 -66.7430743
H -2.2667664 15.3935762 -65.8416272
C -2.4521123 16.7168604 -67.5216241
H -2.901966 16.7154243 -68.5155877
H -1.3820194 16.9074539 -67.6060141
C -3.0951271 17.8042082 -66.6541248
H -2.7560098 17.6658543 -65.6223159
H -4.1813943 17.6762267 -66.6795216
N -2.71647 19.159312 -67.1134731
H -1.700728 19.2804625 -67.0277463
H -3.1123092 19.8764223 -66.5258363
H -2.9676407 19.3212056 -68.075571
N 0.3204954 13.766832 -65.6119406
H -0.1301998 12.8600258 -65.6626692
C 1.1372004 14.0988161 -64.4489276
H 0.7681291 15.0311553 -64.0186157
C 2.6010132 14.3488103 -64.8440332
O 3.1203401 15.4231067 -64.5515805
C 1.0048208 13.0052398 -63.3833035
H -0.0274297 12.9715737 -63.0338871
H 1.2416141 12.0362933 -63.8251743
C 1.9196494 13.2544928 -62.2026704
C 1.5902525 14.2330469 -61.2449909
H 0.6507624 14.7645696 -61.3091331
C 2.5054578 14.5589645 -60.2283914
H 2.2645 15.3361693 -59.5185997
C 3.7477584 13.9046379 -60.163313
H 4.4610224 14.1635952 -59.3939693
C 4.075634 12.9324819 -61.1222156
H 5.0420339 12.4538226 -61.0965527
C 3.1634757 12.6011592 -62.1355357
H 3.4384687 11.8742898 -62.8856969
N 3.2219607 13.4431795 -65.6129092
H 2.7236004 12.5935768 -65.8627977

C 4.6285742 13.5613066 -66.0272132
H 5.2190656 13.6913037 -65.136371
C 4.8554892 14.8103973 -66.9071792
O 5.7538651 15.5982238 -66.6248052
C 5.1469566 12.2423806 -66.6375441
H 4.5264298 12.0049949 -67.503892
C 6.6112232 12.3569321 -67.0848273
H 7.294004 12.1712733 -66.2581363
H 6.8394987 13.3368053 -67.4929083
H 6.78462 11.6171695 -67.8603109
C 5.0988351 11.0728698 -65.6156212
H 5.8917595 11.1882859 -64.8758281
H 4.1709873 11.0890031 -65.0529447
C 5.2309691 9.6913749 -66.2729081
H 6.195366 9.5950969 -66.7695518
H 4.4343643 9.5515472 -67.0036455
H 5.1519131 8.9179254 -65.5089451
N 3.9636351 15.1292163 -67.8520424
H 3.2265652 14.4600239 -68.0547949
C 4.0218868 16.3871815 -68.6226211
H 5.0042765 16.4596209 -69.0901891
C 3.8797531 17.6361744 -67.751329
O 4.3853571 18.6900604 -68.1393617
C 2.9423549 16.3991231 -69.7126516
H 1.9845357 16.1193492 -69.2705717
H 2.8543825 17.4158555 -70.0989016
C 3.2701847 15.4763868 -70.8965743
H 3.2834725 14.434007 -70.5792135
H 4.2600221 15.7274747 -71.2810722
C 2.2462379 15.6651832 -72.0260354
H 2.5661878 15.0769262 -72.8877034
H 2.2537104 16.7118842 -72.3344896
N 0.873631 15.2917612 -71.613203
H 0.3736673 15.9475456 -71.0440993
C 0.2606047 14.1451865 -71.8617678
N 0.8324717 13.2109762 -72.562001
H 0.4215544 12.2750905 -72.7048818
H 1.7817286 13.3170273 -72.8573687
N -0.9394563 13.9204432 -71.4047869
H -1.4254517 14.5628763 -70.815964
H -1.3235025 12.9947997 -71.5411287
N 3.1468849 17.5726836 -66.640555
H 2.7147858 16.6893069 -66.3908726
C 3.0422072 18.6861717 -65.6891141
H 2.9536692 19.6115987 -66.2544648
C 4.3172776 18.8331359 -64.8542892

O 4.818539 19.9522103 -64.7316767
C 1.7891423 18.5422215 -64.8007602
H 1.0555546 17.9044931 -65.2924164
H 2.0612306 18.0606817 -63.8602092
C 1.1195113 19.8957002 -64.5028273
H 1.8473703 20.5638183 -64.0369374
H 0.3048766 19.7346467 -63.7929669
C 0.5491297 20.5310302 -65.7790337
O -0.2282403 19.8629453 -66.4986209
O 0.9457575 21.6510811 -66.1609826
N 4.94132 17.725144 -64.4234338
H 4.4998686 16.8199543 -64.5638902
C 6.2140599 17.8019982 -63.6909489
H 6.0524835 18.5815138 -62.9562327
C 7.3890477 18.3481159 -64.518059
O 8.2732855 18.9822581 -63.9445966
C 6.5604161 16.5360761 -62.8697259
H 7.6036421 16.612572 -62.5826921
C 5.687129 16.5745079 -61.5928064
H 4.6438896 16.3754152 -61.8451636
H 5.7360653 17.5498829 -61.1123979
H 6.0256541 15.83006 -60.8757306
C 6.3611885 15.1622226 -63.5084856
H 5.298497 14.9474787 -63.5301493
H 6.74844 15.1843201 -64.5161972
C 7.0610066 14.0140301 -62.7739185
H 6.6684855 13.8981371 -61.7667884
H 8.122155 14.2216184 -62.7084251
H 6.9110273 13.0840706 -63.3228865
N 7.30301 18.2991724 -65.852446
H 6.594574 17.6978391 -66.2541671
C 8.2430857 18.9945263 -66.7521705
H 9.256973 18.7089679 -66.4899572
C 8.1690656 20.51075 -66.5818082
O 9.1990222 21.1611148 -66.460308
C 7.977772 18.6399987 -68.229651
H 6.9892592 19.0220523 -68.484297
C 8.9845327 19.2914653 -69.1874602
H 10.0014391 18.9979181 -68.9194354
H 8.9126701 20.3778295 -69.1491886
H 8.7886603 18.966429 -70.2109033
C 7.96589 17.1408008 -68.5243393
H 8.943109 16.8146886 -68.8779901
H 7.2357508 16.9438971 -69.3046957
H 7.7171778 16.551408 -67.6508645
N 6.9615765 21.0893303 -66.532496

H 6.1488449 20.486938 -66.5022801
C 6.780615 22.5474602 -66.4135413
H 7.4330732 23.0541204 -67.1286945
C 7.2202667 23.0717325 -65.0569848
O 7.912679 24.0815246 -64.9821131
C 5.3159302 22.9327156 -66.6760503
H 4.6831159 22.049465 -66.5844177
H 4.9780527 23.653042 -65.9286919
C 5.1585947 23.5769347 -68.0589042
H 5.5905341 22.9257093 -68.8152424
H 5.7275276 24.5057287 -68.0837541
C 3.6905468 23.8966561 -68.3952165
H 3.6380314 24.2920585 -69.4108383
H 3.3606373 24.6917945 -67.7218206
N 2.7656957 22.7454938 -68.2527494
H 2.0283473 22.845749 -67.5579178
C 2.88091 21.5274568 -68.7454514
N 3.8693124 21.140862 -69.4991559
H 4.0949378 20.1565639 -69.4957941
H 4.5709032 21.8214323 -69.7161739
N 1.9903342 20.6334323 -68.4586807
H 1.2955849 20.8980357 -67.7519289
H 2.1477213 19.6774308 -68.6872359
N 6.8385406 22.3850171 -63.9793491
H 6.2678414 21.5594362 -64.1188139
C 7.0762139 22.8882716 -62.6154744
H 6.7536171 23.9302454 -62.5989197
C 8.5529194 22.901266 -62.209056
O 8.8936518 23.5075295 -61.1960446
C 6.2302789 22.1490035 -61.5659052
H 6.3002834 22.7158685 -60.6375336
C 4.7406745 22.1370212 -61.9312552
H 4.5608851 21.5072413 -62.7992222
H 4.4104188 23.1524728 -62.155835
H 4.1599278 21.7513496 -61.0951958
C 6.7187884 20.740197 -61.2639461
H 6.946144 20.2365441 -62.1944734
H 5.9697426 20.1820863 -60.7022476
H 7.6299397 20.7966649 -60.6732389
N 9.4330305 22.2598839 -62.9727844
H 9.1190088 21.8290885 -63.831774
C 10.8586508 22.2157185 -62.6987607
H 11.0205601 22.3896924 -61.6385775
C 11.6587618 23.2613323 -63.4943908
O 11.250341 23.686733 -64.571033
C 11.3162717 20.8126336 -63.009183

H 11.2222436 20.673128 -64.0870845
H 10.6948542 20.0962774 -62.4706286
H 12.34619 20.6925253 -62.698318
N 12.8048385 23.6814376 -62.9622398
H 13.0708624 23.3246349 -62.0518115
C 13.7495271 24.5578181 -63.6509041
H 13.1778 25.2862108 -64.2238724
C 14.6719354 23.7794981 -64.6223379
O 14.975169 22.6149417 -64.3663321
C 14.5534374 25.3133525 -62.584314
H 15.1349368 24.6050227 -61.9900518
H 13.8784704 25.8645982 -61.9278724
H 15.2402197 26.0126728 -63.0612602
N 15.2562672 24.4346856 -65.6428965
C 16.3457623 23.8565942 -66.4329207
H 15.9645131 23.0084207 -66.9972777
C 17.5084646 23.3942187 -65.5437926
O 17.9666931 24.1420647 -64.6815619
C 16.7604415 24.9539896 -67.41675
H 17.5396302 25.5804047 -66.9808329
H 17.0916402 24.5356973 -68.368371
C 15.4800201 25.7739039 -67.5748403
H 14.8102338 25.2746548 -68.2775496
H 15.6876092 26.7949285 -67.8961341
C 14.8755827 25.7311526 -66.1737705
H 13.7917232 25.8368128 -66.2352599
H 15.3069372 26.5190763 -65.555338
N 17.9761557 22.1537945 -65.7184936
H 17.5578649 21.5750566 -66.4370437
C 18.9691881 21.5330127 -64.828799
H 19.4528712 20.7173447 -65.3653614
H 19.7268063 22.271958 -64.5676816
C 18.3945524 20.9579177 -63.5231969
O 19.1431088 20.4234825 -62.6929978
N 17.0763422 20.9995032 -63.325819
H 16.4871733 21.5126475 -63.9742723
C 16.4200515 20.282351 -62.2438022
H 16.8992307 20.5608404 -61.3042546
C 16.5430015 18.7673145 -62.4295642
O 16.5979498 18.2344241 -63.5433115
C 14.9511917 20.6876016 -62.1594518
H 14.4759675 20.4721613 -63.1160417
H 14.876111 21.7519627 -61.935609
H 14.4412873 20.1255796 -61.3775577
N 16.4551063 18.0541021 -61.3125826
H 16.2855776 18.5690471 -60.4511779

C 16.3207142 16.6066863 -61.2672556
H 16.667364 16.1436285 -62.1925943
C 14.8430527 16.4876811 -61.1714432
O 14.2703098 16.4918358 -60.0760996
C 17.1025975 16.0055349 -60.0967373
H 16.7267382 16.4254048 -59.1674263
H 16.9332383 14.927969 -60.0696969
C 18.6151246 16.2789271 -60.2035818
H 18.7934009 17.3000794 -60.5397025
H 19.0498173 15.6006874 -60.9384452
C 19.3309473 16.1098317 -58.8681021
O 19.8209138 17.063498 -58.2767724
N 19.4280626 14.9016131 -58.3528361
H 19.9160454 14.8206799 -57.47876
H 19.0269415 14.1074667 -58.8221429
N 14.2900884 16.7928196 -62.3621341
H 14.9082487 16.9578584 -63.1449686
C 12.8660165 16.8921841 -62.5786515
H 12.50313 17.8129391 -62.118628
C 12.3945187 16.8195691 -64.0583579
H 11.5182164 16.1806944 -64.1205188
C 11.927267 18.2080647 -64.4477732
H 12.7140434 18.9289267 -64.2293846
H 11.0102521 18.4103486 -63.8975376
H 11.6871596 18.2879955 -65.5028421
C 13.3973882 16.3601632 -65.1198995
H 13.1242213 16.743578 -66.1014841
H 14.3632553 16.7589904 -64.8585091
C 13.4802412 14.8541873 -65.2635705
H 12.5137973 14.4524802 -65.5470749
H 13.7913259 14.4259101 -64.3196536
H 14.1809111 14.6061911 -66.0551717
C 12.2832975 15.764129 -61.7971147
O 11.2494559 16.0014477 -61.1925983
N 12.9953431 14.6160479 -61.6922139
H 13.8487719 14.4634771 -62.2140222
C 12.4409562 13.5648977 -60.9104669
H 11.9661348 14.0429408 -60.056893
C 13.3950247 12.5287825 -60.3251963
O 14.3898634 12.1321749 -60.9337501
C 11.3641755 12.9163869 -61.7760491
H 11.69877 11.9263678 -62.0871113
C 10.1448767 12.7675339 -60.8818847
H 9.7797381 13.7482455 -60.5753726
H 10.389334 12.200937 -59.985975
H 9.3515675 12.2659959 -61.4236928

C 10.8142608 13.647437 -63.0167973
H 10.2651247 14.5505195 -62.7809551
H 10.1246692 12.9691349 -63.5158578
H 11.6027534 13.8396563 -63.7244018
N 12.992268 12.0624941 -59.1407394
H 12.2290494 12.5696896 -58.7005972
C 13.3517678 10.7991302 -58.4875107
H 13.8691189 10.1454328 -59.1901295
C 12.022295 10.1409558 -58.0722694
O 11.2124517 10.7801555 -57.3905926
C 14.2732964 11.0619765 -57.2714117
H 13.7556098 11.7467959 -56.5983282
C 14.5539489 9.7552096 -56.5040559
H 15.098085 9.0547942 -57.1393494
H 13.6255968 9.2720965 -56.1977078
H 15.1326487 9.9473315 -55.6028041
C 15.5967541 11.7351388 -57.7104154
H 16.1322448 11.0671286 -58.3834904
H 15.3704105 12.6510699 -58.2557262
C 16.5299331 12.1281646 -56.5575202
H 16.9353557 11.2398739 -56.0737671
H 15.9862613 12.7290053 -55.8275692
H 17.3598219 12.7148318 -56.9523203
N 11.7580335 8.9046503 -58.5018071
H 12.4971308 8.4049395 -59.0006782
C 10.4983895 8.1867901 -58.2471085
H 9.9675434 8.6814518 -57.4396159
C 10.7986262 6.7492106 -57.8252037
O 11.102295 5.9443846 -58.7003625
C 9.5576584 8.1569015 -59.488233
H 10.0657489 7.6118162 -60.2850846
C 9.2280089 9.5527545 -60.0531933
H 8.7489912 10.1650888 -59.2892799
H 10.1684049 10.018405 -60.3410172
C 8.2538598 7.3944316 -59.1507915
H 7.707404 7.913199 -58.3643917
H 8.4738782 6.3822778 -58.8197286
H 7.6197814 7.2880813 -60.0276344
C 8.3419087 9.5094014 -61.311817
H 7.3388524 9.1658198 -61.0701747
H 8.7868381 8.8466241 -62.055071
H 8.2449684 10.4990386 -61.7479271
N 10.3778145 6.3647875 -56.6173089
H 10.0867171 7.0875804 -55.969359
C 10.0771669 4.965996 -56.2829504
H 10.6161371 4.3342597 -56.978693

C 8.5735844 4.707958 -56.4325046
O 7.7547345 5.5273144 -56.0055968
C 10.6004787 4.5910992 -54.8968489
H 10.1061673 5.1945444 -54.1366728
H 11.6735419 4.7889482 -54.8607758
O 10.3828632 3.2140938 -54.6418497
H 9.4472056 3.0366627 -54.4338089
N 8.1607001 3.6309068 -57.1055167
H 8.8769937 2.9804449 -57.4273788
C 6.7371884 3.300688 -57.3620578
H 6.1596131 3.5083897 -56.4621738
C 6.5491433 1.8064488 -57.6597888
O 7.4787363 1.1227106 -58.0834072
C 6.1822375 4.201798 -58.4902398
H 6.9381585 4.3099942 -59.2694911
C 4.8723056 3.766805 -59.1432156
H 4.1171477 3.6139869 -58.3752925
H 5.0235349 2.8460121 -59.7068042
H 4.5357146 4.5394803 -59.8340945
O 5.8902408 5.4700017 -57.9372644
H 6.6083844 5.6589023 -57.3062582
N 5.3715908 1.2506011 -57.3572631
H 4.6553821 1.8125038 -56.8983762
C 5.0533527 -0.1448405 -57.6414244
H 5.9175855 -0.7539901 -57.3692877
C 4.7686066 -0.3651727 -59.1348527
O 3.9382459 0.3149866 -59.7426248
C 3.8790949 -0.6020956 -56.7719286
H 2.9852171 -0.0491227 -57.0590648
H 4.0992285 -0.4156114 -55.7198245
H 3.7121333 -1.6687299 -56.9224242
N 5.384449 -1.4132231 -59.6710028
H 5.9615485 -1.978817 -59.0585404
C 5.274073 -1.8707675 -61.0481607
H 4.52063 -1.2819514 -61.5709594
C 4.819635 -3.3368143 -61.111532
O 4.71484 -4.0327033 -60.0985856
C 6.6290324 -1.6468094 -61.7413333
H 6.7247861 -2.2962304 -62.6122492
H 7.4480343 -1.8571985 -61.0513851
S 6.7161264 0.0744788 -62.2988248
H 5.7073454 -0.0156779 -63.1767116
N 4.5804398 -3.8163232 -62.3306187
H 4.6770828 -3.1691214 -63.1064769
C 4.346574 -5.2294842 -62.6551505
H 4.459447 -5.8264837 -61.7510928

C 5.4032849 -5.7429896 -63.6329606
O 6.1897972 -4.9572822 -64.1649364
C 2.9059161 -5.4309908 -63.1662228
H 2.7672556 -6.4849372 -63.4143421
H 2.2061259 -5.1920836 -62.3647088
C 2.5629917 -4.5759187 -64.3962885
H 3.3538419 -4.6507815 -65.1381943
H 2.4774119 -3.5317996 -64.1082129
C 1.239358 -5.0199314 -65.0200982
H 0.4409992 -4.8935195 -64.2851521
H 1.2911458 -6.0804497 -65.2787751
N 0.9258565 -4.2126199 -66.212061
H 0.4081314 -3.3590145 -66.0351975
C 1.2259418 -4.5210722 -67.4631425
N 1.9688723 -5.5468659 -67.7776155
H 2.1152903 -5.8721481 -68.7219343
H 2.3570204 -6.1389378 -67.0480665
N 0.7801215 -3.779182 -68.4391839
H 0.1596895 -3.0162654 -68.2328983
H 0.9754474 -4.0537471 -69.3841574
N 5.3899017 -7.0384504 -63.929528
H 4.7383203 -7.6344439 -63.4363782
C 6.0942102 -7.5596355 -65.1036889
H 7.0977374 -7.1300138 -65.1006469
C 5.4232967 -7.0925274 -66.4226341
O 4.3354509 -6.5106139 -66.4170726
C 6.2677109 -9.0838745 -64.9594063
H 6.6936781 -9.2945219 -63.9792089
H 6.9785677 -9.4402086 -65.7029615
C 4.9912565 -9.885612 -65.1365704
O 4.2025192 -9.6370408 -66.0222303
N 4.7808317 -10.9165378 -64.3629691
H 3.901636 -11.390675 -64.4666077
H 5.4341878 -11.1196392 -63.6117621
N 6.087026 -7.301176 -67.5654737
H 6.97492 -7.7768733 -67.5279906
C 5.5686416 -6.8960675 -68.8824775
H 5.3268371 -5.8354456 -68.847767
C 4.2717675 -7.636028 -69.2757134
O 3.3960169 -7.0361793 -69.9063768
C 6.6966829 -7.1057223 -69.9118019
H 7.5465829 -6.4850685 -69.622889
H 7.0144693 -8.1489414 -69.8602459
C 6.3428838 -6.7968035 -71.3816833
H 5.5274226 -7.4439008 -71.7033055
C 5.9359276 -5.3387014 -71.5982807

H 6.7312629 -4.6712647 -71.263949
H 5.0185384 -5.1252485 -71.0539862
H 5.7487945 -5.1681951 -72.6580208
C 7.5553389 -7.0841982 -72.2684394
H 8.3866897 -6.4341584 -71.9947723
H 7.2949337 -6.9136733 -73.313153
H 7.8578121 -8.1251123 -72.1530906
N 4.1334798 -8.8979656 -68.8636108
H 4.8250579 -9.2749777 -68.2335885
C 3.0282568 -9.7990911 -69.2066128
H 2.994302 -9.8630138 -70.2915907
C 1.6602333 -9.3119172 -68.6927965
O 1.5945623 -8.516802 -67.7502102
C 3.3072741 -11.2092038 -68.6610331
H 2.5364612 -11.8929918 -69.0166306
H 3.2739869 -11.2045757 -67.5732222
O 4.5717364 -11.6737482 -69.1032695
H 4.7435367 -12.5260357 -68.6915725
N 0.5433211 -9.7783737 -69.2905212
C -0.7985269 -9.4881579 -68.7920022
H -0.974344 -8.4156383 -68.878059
C -0.9754439 -9.9212677 -67.3331253
O -0.527133 -10.9988821 -66.9390965
C -1.7659872 -10.2366409 -69.7171588
H -1.976375 -11.2318292 -69.3196156
H -2.693753 -9.6827823 -69.8650191
C 0.4554123 -10.5666366 -70.5124142
H 1.1583618 -10.2258453 -71.2721388
H 0.6242898 -11.6193873 -70.280253
C -0.973233 -10.3730318 -71.0152853
H -1.0393598 -9.4455678 -71.586019
H -1.3115491 -11.2200922 -71.6130074
N -1.6717958 -9.0991233 -66.547554
H -1.9774122 -8.2113845 -66.9104634
C -1.9426282 -9.3712957 -65.1363939
H -0.9869589 -9.4906116 -64.6280691
C -2.7599268 -10.6578647 -64.9656591
O -3.8400503 -10.7898855 -65.5414905
C -2.701309 -8.1901542 -64.5041285
H -2.7718945 -8.3380187 -63.4261289
H -3.7101012 -8.1606276 -64.9151862
O -2.0772418 -6.945159 -64.7763668
H -2.4140318 -6.2669389 -64.1668781
N -2.292544 -11.5662814 -64.1043462
H -1.3627723 -11.44079 -63.7365719
C -3.0207712 -12.7921241 -63.7457168

H -3.2244875 -13.323451 -64.6781217
C -4.3966937 -12.4684447 -63.1343671
O -4.6230048 -11.3825377 -62.579092
C -2.1096583 -13.72106 -62.9038127
H -1.910148 -14.6234283 -63.4834728
H -1.1418491 -13.2284864 -62.8004951
C -2.5148567 -14.1252411 -61.4840284
H -2.7946617 -13.2267165 -60.9280213
H -1.6416138 -14.553518 -60.9864657
C -3.6505368 -15.1492933 -61.4302847
O -4.2452488 -15.22482 -60.332582
O -4.0275463 -15.7087837 -62.4791995
N -5.3275999 -13.4242844 -63.2208885
H -5.0081732 -14.3620026 -63.4746751
C -6.7373853 -13.2558538 -62.8708644
H -7.1275333 -12.4809128 -63.5271843
C -6.946757 -12.7679238 -61.4410492
O -7.9117529 -12.0322895 -61.2615722
C -7.4812086 -14.5788163 -63.1367407
H -7.2969487 -14.8900304 -64.1668939
H -7.0568127 -15.3459934 -62.4846586
C -8.9995942 -14.5392484 -62.9048429
H -9.3885239 -15.5543231 -62.9959171
H -9.1947796 -14.2219061 -61.8805407
C -9.7553918 -13.6445641 -63.9063237
H -10.1326223 -14.2766743 -64.7127237
H -9.0952164 -12.9031723 -64.3565096
C -10.9343519 -12.935928 -63.2296966
H -11.4774207 -13.6713244 -62.6279936
H -11.6147183 -12.563259 -64.0023007
N -10.4659423 -11.8111498 -62.3804404
H -9.7675482 -12.1340372 -61.710333
H -11.2343898 -11.3862407 -61.8810379
H -10.0067443 -11.1094506 -62.9439105
N -6.0816248 -13.0886981 -60.4718269
H -5.3969747 -13.8110332 -60.6825751
C -6.0968226 -12.5397581 -59.1017552
H -7.1172031 -12.2357542 -58.8791371
C -5.2212816 -11.2922417 -58.8815129
O -5.4720267 -10.5585664 -57.9262851
C -5.7549487 -13.6167123 -58.0640669
H -6.3952429 -14.4869167 -58.2181221
H -5.9482444 -13.2182487 -57.0671658
O -4.3992676 -14.0098058 -58.1432093
H -4.343102 -14.5866227 -58.960205
N -4.2784466 -10.9636136 -59.7796263

H -4.191226 -11.5356199 -60.6116894
C -3.5645761 -9.675878 -59.7459851
H -3.2982635 -9.4612025 -58.710529
C -4.4764878 -8.5391038 -60.2170946
O -4.6424323 -7.5619692 -59.492832
C -2.2637066 -9.7208699 -60.5782299
H -1.5860132 -10.4576694 -60.1459091
H -2.5060962 -10.0411425 -61.5913691
C -1.54592 -8.348949 -60.6594275
H -2.2252017 -7.6004366 -61.0640231
C -1.049715 -7.869789 -59.2926931
H -0.3925492 -8.6162577 -58.8481829
H -1.895953 -7.6778135 -58.6345231
H -0.5004847 -6.9352805 -59.4146506
C -0.3461376 -8.4080241 -61.5969519
H 0.3865714 -9.1103831 -61.226032
H 0.1108929 -7.4212003 -61.6794127
H -0.6523785 -8.7230191 -62.5882788
N -5.1169134 -8.689943 -61.3849318
H -4.9457027 -9.5459942 -61.90868
C -6.0412299 -7.6860189 -61.9416269
H -5.4551489 -6.7857532 -62.1437637
C -7.1240799 -7.231914 -60.9375162
O -7.2297424 -6.0346225 -60.7273625
C -6.5934143 -8.1928127 -63.2907024
H -5.7661369 -8.2380254 -63.9984981
H -6.9541082 -9.2147834 -63.16253
C -7.7329938 -7.3712305 -63.9310273
H -7.8971692 -7.7500898 -64.9408023
H -8.6442331 -7.5659483 -63.3636415
C -7.5575872 -5.8442729 -64.0094301
H -7.4980538 -5.4352241 -63.0077227
H -8.4615509 -5.4163366 -64.4482493
C -6.3614685 -5.3390559 -64.8205827
H -6.662531 -5.2804452 -65.8695286
H -5.5195257 -6.0309988 -64.7300999
N -5.9571078 -4.0034871 -64.3263948
H -6.7679802 -3.395136 -64.17414
H -5.2630821 -3.5587778 -64.9038722
H -5.4837897 -4.0957187 -63.4295262
N -7.871645 -8.0931999 -60.2239082
C -8.873936 -7.6616363 -59.2573345
H -9.5234761 -6.9139405 -59.7159941
C -8.2649088 -7.0320422 -57.9992117
O -8.9337415 -6.198365 -57.3998078
C -9.7099499 -8.8997193 -58.9176722

H -10.0869565 -8.8805224 -57.8938592
H -10.5345133 -8.9914458 -59.6264198
C -7.9351771 -9.5147877 -60.3439147
H -8.4774898 -9.7379089 -61.262144
H -6.9490279 -9.9651637 -60.3563997
C -8.720291 -10.0246495 -59.1344312
H -9.2281338 -10.9685741 -59.3297095
H -8.0659944 -10.10908 -58.2666461
N -7.021786 -7.3606245 -57.6043131
H -6.4998668 -8.0322734 -58.1535313
C -6.3056253 -6.6353359 -56.5348897
H -6.9728801 -6.5111878 -55.6824392
C -5.9322881 -5.2270172 -57.0036012
O -6.1602659 -4.2700824 -56.2681912
C -5.0687729 -7.4390979 -56.0860549
H -5.3895788 -8.424192 -55.7418211
H -4.4305065 -7.5948128 -56.9556649
C -4.1992388 -6.7757217 -54.993123
H -3.2115868 -7.2341931 -55.0481208
H -4.0668317 -5.7140503 -55.2046993
C -4.7169933 -6.9550987 -53.5545555
H -4.8133604 -8.0239603 -53.351868
H -5.6999526 -6.4874112 -53.4551922
C -3.7314371 -6.3211879 -52.5553124
H -3.9467867 -5.2535484 -52.4822222
H -2.7197203 -6.4281609 -52.9491867
N -3.7880118 -6.9492504 -51.2117734
H -4.7008678 -6.8646728 -50.7950431
H -3.1294465 -6.5095671 -50.5629086
H -3.5314121 -7.9249485 -51.2652042
N -5.4101965 -5.0928218 -58.2204721
H -5.2513331 -5.9347831 -58.7634563
C -5.1139113 -3.8079675 -58.8626221
H -4.4172266 -3.2441876 -58.2414818
C -6.3827766 -2.9486339 -58.9977917
O -6.424878 -1.8461333 -58.4601641
C -4.4306352 -4.0925634 -60.2147685
H -3.4889227 -4.612916 -60.0251308
H -5.0681263 -4.750292 -60.8032861
C -4.1402352 -2.8329003 -61.0402102
H -3.5223312 -2.1630568 -60.4391233
H -5.0881717 -2.3339607 -61.2560169
C -3.4537445 -3.1310734 -62.3813936
O -3.6879035 -4.2298107 -62.9487496
O -2.7668995 -2.224337 -62.9038172
N -7.4664462 -3.5002571 -59.5499567

H -7.3734285 -4.4240226 -59.9575726
C -8.7116426 -2.7639545 -59.7964255
H -8.4353106 -1.8095062 -60.2480485
C -9.4638687 -2.4089424 -58.5074308
O -9.9528278 -1.288221 -58.3755535
C -9.6118555 -3.5466591 -60.7821039
H -9.0607949 -4.3693492 -61.2330581
H -10.4668038 -3.9705644 -60.2532929
C -10.1111674 -2.6392604 -61.9230753
H -10.7214885 -1.8416841 -61.4932013
H -10.7558622 -3.2309231 -62.5768496
C -8.9667387 -2.0313695 -62.7636765
O -8.2371866 -2.7994259 -63.4391515
O -8.8066446 -0.7847794 -62.7441526
N -9.47042 -3.2962327 -57.4999447
H -9.0417097 -4.20374 -57.6524768
C -10.1010431 -3.0170332 -56.1973756
H -11.0288266 -2.4764864 -56.3973745
C -9.2769901 -2.0761537 -55.3142795
O -9.8498703 -1.4157787 -54.4554675
C -10.4660358 -4.334596 -55.4944689
H -11.0058283 -4.9560219 -56.2112972
H -9.5566922 -4.8544857 -55.1890845
C -11.3704398 -4.1053174 -54.2696322
H -10.7558442 -3.807233 -53.4188719
H -12.0692675 -3.2953097 -54.4855386
C -12.2007088 -5.3376816 -53.8874421
H -12.8839548 -5.5640901 -54.7082912
H -12.7938671 -5.0865768 -53.0059178
C -11.3301939 -6.5636396 -53.5898349
H -10.6293076 -6.3072624 -52.7892185
H -10.7480368 -6.8020651 -54.4859351
N -12.1634946 -7.726805 -53.1936491
H -12.701381 -7.5095402 -52.3633894
H -11.5843428 -8.534113 -52.9993255
H -12.8073944 -7.9650457 -53.9371857
N -7.9716141 -1.9444161 -55.5661454
H -7.5758423 -2.5040646 -56.3110106
C -7.1488087 -0.8533673 -55.0040867
H -7.6222074 -0.5067725 -54.0856447
C -7.1064176 0.3871534 -55.9041949
O -6.4532318 1.363562 -55.5514579
C -5.7327836 -1.3132419 -54.6059369
H -5.1704231 -0.4493004 -54.2512706
C -5.7686133 -2.3392788 -53.4714731
H -6.3200593 -3.2274988 -53.7777969

H -6.2540956 -1.8980821 -52.6008782
H -4.7497511 -2.6174525 -53.2046511
O -5.0121386 -1.9015539 -55.6648057
H -5.5081724 -2.690785 -55.9458878
N -7.8166632 0.3654646 -57.0417343
H -8.3462446 -0.4726061 -57.2427193
C -7.8502309 1.3899778 -58.0923619
H -8.3461036 0.901136 -58.9292334
C -6.4594769 1.7892235 -58.633805
O -6.3001095 2.8344891 -59.2640218
C -8.7526115 2.5574542 -57.6322694
H -9.6187161 2.1305136 -57.1235023
H -8.1999693 3.1478449 -56.8997247
C -9.2803135 3.5032468 -58.7320415
H -8.4579137 4.0436594 -59.1952806
C -10.0659324 2.771923 -59.8264709
H -10.8596408 2.1741863 -59.3761475
H -9.4021933 2.1159166 -60.3858728
H -10.5024011 3.4948423 -60.5147045
C -10.2132689 4.5376623 -58.0986059
H -11.0746174 4.0426197 -57.649767
H -10.5543428 5.2422816 -58.85601
H -9.6730952 5.0878871 -57.3275553
N -5.4426723 0.9434196 -58.4481881
H -5.6678236 0.0355238 -58.0548346
C -4.0798935 1.185912 -58.9357536
H -3.8150141 2.2211407 -58.7113507
C -4.0051982 1.032849 -60.4518383
O -4.7678273 0.27695 -61.0434649
C -3.0949649 0.2568287 -58.2130025
H -3.5138127 -0.7490093 -58.1422002
H -2.1678002 0.2159683 -58.784929
C -2.795052 0.8141 -56.8097441
H -2.3119699 1.7866437 -56.9208895
H -3.7301621 0.9656837 -56.2690902
C -1.8893985 -0.0773955 -55.9502131
H -2.4545609 -0.9480272 -55.6154601
H -1.5950424 0.5033683 -55.0736083
C -0.6439782 -0.5463608 -56.7108241
H -0.3018319 0.268945 -57.3493463
H -0.9215663 -1.3830817 -57.3566641
N 0.4467047 -0.9339892 -55.7916971
H 0.7465728 -0.112789 -55.2498367
H 1.2748543 -1.2105249 -56.2979961
H 0.1611309 -1.6486113 -55.1440047
N -3.0487572 1.7262357 -61.0774219

H -2.4915023 2.3509615 -60.5015827
C -2.7384924 1.6213578 -62.5167704
H -3.2764944 0.767828 -62.9341887
C -1.2710996 1.2723612 -62.7035448
O -0.4066791 2.1504201 -62.6343866
C -3.1835379 2.9095297 -63.2545982
H -3.4195988 3.6894009 -62.5300726
H -2.3764105 3.2816492 -63.8842477
C -4.4157079 2.6815394 -64.1627795
H -4.7723658 3.6569256 -64.4963477
H -4.1117891 2.1108096 -65.0407656
C -5.5713926 1.9356428 -63.4710507
H -5.670715 2.3288555 -62.460506
H -5.3424209 0.8725846 -63.4125411
C -6.9189723 2.0674559 -64.1837241
H -6.9312356 1.427326 -65.0694601
H -7.0421595 3.108668 -64.4916204
N -8.0097774 1.7033435 -63.2510456
H -7.9605382 0.719054 -62.960874
H -8.919042 1.7965361 -63.6756161
H -7.9574248 2.2846703 -62.4272009
N -0.9851324 -0.0152724 -62.8685472
H -1.7507671 -0.6921228 -62.9343551
C 0.3622951 -0.5630426 -62.7563147
H 0.9805693 0.166144 -62.2307181
C 0.9623679 -0.7635467 -64.1569443
O 0.4536773 -1.5034089 -65.0035394
C 0.3662204 -1.8384641 -61.8795023
H -0.0630003 -2.6610733 -62.4548948
C 1.819163 -2.1735927 -61.5191769
H 2.2062048 -1.4454116 -60.8056847
H 2.4410637 -2.152216 -62.4072739
H 1.8767011 -3.1699076 -61.08193
C -0.4305112 -1.6738873 -60.5577608
H 0.0351023 -0.9039866 -59.9410057
H -1.442414 -1.3440835 -60.7786126
C -0.5557201 -2.9669112 -59.7395202
H 0.4082644 -3.2467547 -59.3150915
H -0.9233073 -3.7726035 -60.3764866
H -1.265722 -2.8133331 -58.927048
N 2.0820627 -0.0851478 -64.4029156
H 2.4655832 0.475942 -63.6575331
C 2.9068083 -0.2770199 -65.5962123
H 2.3018658 -0.6952979 -66.4020926
C 4.0396154 -1.2643317 -65.292497
O 4.4307098 -1.4269569 -64.1332412

C 3.4495725 1.0814523 -66.061202
H 4.0958151 0.9525892 -66.9322466
H 4.0432311 1.5292122 -65.261646
S 2.0709612 2.1846743 -66.4876026
H 1.3003227 1.8921115 -65.4342484
N 4.6448883 -1.8495044 -66.3268338
H 4.3466771 -1.6176306 -67.261478
C 5.9932657 -2.3856146 -66.1595621
H 6.0491473 -2.8706147 -65.1886549
C 7.0088041 -1.2347704 -66.1249314
O 6.737833 -0.132612 -66.6086652
C 6.3073177 -3.4524623 -67.2163512
H 5.4957053 -4.1766962 -67.2517385
H 7.2191217 -3.9815768 -66.9375472
C 6.5230252 -2.8364934 -68.5773457
O 7.5678347 -2.280934 -68.8635157
N 5.5086591 -2.8222407 -69.403895
H 5.6807276 -2.3748843 -70.3032807
H 4.6772054 -3.3460722 -69.2191437
N 8.1801732 -1.5131329 -65.5618081
H 8.3470506 -2.4508453 -65.2320084
C 9.2452561 -0.5331003 -65.3748175
H 8.8589516 0.286887 -64.7647094
C 9.6957767 0.098359 -66.7072817
O 9.7789872 1.3222783 -66.8014925
C 10.3873618 -1.2029495 -64.5780293
H 11.2574185 -0.549599 -64.5671677
H 10.0590078 -1.2929866 -63.5414857
C 10.8177317 -2.5865022 -65.0490904
C 11.9181375 -2.7365552 -65.9169673
H 12.4819757 -1.8755661 -66.2390297
C 12.3053206 -4.0136478 -66.363435
H 13.1452796 -4.1187607 -67.0335377
C 11.597087 -5.1547648 -65.9319127
O 11.9592972 -6.3881262 -66.3699191
H 12.7274721 -6.3595775 -66.9421564
C 10.5129368 -5.0118722 -65.040923
H 9.9908157 -5.8930459 -64.6965091
C 10.1282528 -3.7322316 -64.5973171
H 9.308215 -3.6379075 -63.8983316
N 9.914116 -0.6953223 -67.7660937
H 9.7185 -1.6813804 -67.6780474
C 10.4258322 -0.1969978 -69.0526211
H 11.3531195 0.3334907 -68.8684677
C 9.4744436 0.8147866 -69.7068052
O 9.9050969 1.903903 -70.0918449

C 10.7555701 -1.3866897 -69.9704476
H 11.4799712 -2.0264374 -69.4643361
H 9.8468928 -1.968167 -70.1290859
C 11.319076 -0.9912917 -71.3499645
H 10.5886703 -0.3825456 -71.8825983
C 12.633967 -0.2142643 -71.2488822
H 13.3625122 -0.779567 -70.6682336
H 12.462991 0.7532524 -70.7772734
H 13.0319279 -0.0350601 -72.2476385
C 11.5735498 -2.2547509 -72.1719436
H 12.3160219 -2.8816896 -71.6783682
H 11.9344282 -1.9816748 -73.1636924
H 10.6437644 -2.8131246 -72.2827093
N 8.1721243 0.5219023 -69.7300654
H 7.8781855 -0.3931025 -69.4008782
C 7.1627271 1.4694778 -70.216607
H 7.465206 1.8258281 -71.2019721
C 7.0485684 2.7218684 -69.344085
O 6.7977963 3.7974389 -69.8845881
C 5.8037654 0.7721156 -70.3591403
H 5.0142317 1.5249066 -70.3314356
H 5.6424419 0.0910384 -69.5229777
C 5.6722935 0.0300481 -71.6647418
N 5.9532087 -1.3209948 -71.8725283
C 5.7499517 -1.545272 -73.178947
H 5.897551 -2.4993019 -73.6695688
N 5.3500598 -0.4176791 -73.790252
H 5.1471925 -0.3273254 -74.7758818
C 5.2980715 0.5887433 -72.851667
H 5.0341347 1.6258243 -73.0140616
N 7.3164396 2.6332608 -68.0360767
H 7.5599259 1.7314697 -67.6420576
C 7.2614763 3.7968527 -67.1509479
H 6.3620564 4.3393293 -67.4291629
C 8.4261241 4.7738874 -67.3880553
O 8.1695259 5.9696373 -67.5199989
C 7.0948289 3.3363918 -65.6891448
H 6.3288467 2.5601209 -65.6683109
H 8.0271337 2.8928056 -65.3390609
C 6.6681347 4.4549309 -64.7117129
H 7.5189438 5.109165 -64.5258907
C 5.4997377 5.3156355 -65.2074584
H 4.6757884 4.679795 -65.5306189
H 5.8301918 5.9439793 -66.0348046
H 5.1677863 5.9791257 -64.4099764
C 6.2145039 3.8387536 -63.3851094

H 5.3016056 3.2618588 -63.5260214
H 6.0445249 4.6237662 -62.6487135
H 6.9882306 3.1722436 -63.0041928
N 9.6480168 4.2850584 -67.6533181
H 9.7997559 3.2846604 -67.5630426
C 10.7262591 5.1507473 -68.1801541
H 10.8557976 5.9933419 -67.499146
C 10.3952471 5.6900104 -69.5631661
O 10.7418516 6.824877 -69.8713061
C 12.0605472 4.4240518 -68.3016877
H 12.1166467 3.8610945 -69.2350101
H 12.121121 3.7293563 -67.4803064
O 13.1491082 5.3247477 -68.1948499
H 13.269569 5.8510611 -69.0056855
N 9.6793356 4.9112594 -70.3797769
H 9.4739327 3.9631262 -70.084374
C 9.1529553 5.3651188 -71.6612905
H 9.9800387 5.6169965 -72.3253587
H 8.5721268 4.5597261 -72.1096179
C 8.2436502 6.5889815 -71.5206075
O 8.4964877 7.5881455 -72.1829673
N 7.271716 6.5802667 -70.6017955
H 7.1008963 5.7179054 -70.0934582
C 6.3753098 7.7248641 -70.3564864
H 5.913338 8.0175862 -71.2986071
C 7.1064366 8.967594 -69.8279953
O 6.7901523 10.0960769 -70.2162708
C 5.2706065 7.3106618 -69.3736671
H 5.7289851 6.9026424 -68.4718638
H 4.7203521 8.2063148 -69.0832355
C 4.2527546 6.3259864 -69.9274463
C 3.9569052 5.1377987 -69.2312677
H 4.470828 4.90886 -68.3126257
C 2.9868958 4.2463839 -69.7247382
H 2.7656118 3.3367646 -69.1875547
C 2.2932897 4.5473782 -70.9100298
H 1.5351606 3.8746796 -71.2849157
C 2.5749518 5.7375725 -71.6014384
H 2.0297721 5.995317 -72.5016579
C 3.5527959 6.6201892 -71.1146242
H 3.7383022 7.5375435 -71.6559592
N 8.1333138 8.7500029 -69.0098282
H 8.2811018 7.8069615 -68.6678179
C 9.07562 9.7716512 -68.5730746
H 8.5547353 10.5922679 -68.0912188
C 9.8605926 10.3853852 -69.7513712

O 9.7566054 11.5847949 -70.0065046
C 9.9597189 9.1076336 -67.5219712
H 10.8339721 9.7058895 -67.3759088
H 10.2895408 8.1279684 -67.8552867
S 9.0625811 8.9471298 -65.9503501
H 9.8351265 9.8474364 -65.3019252
N 10.4805271 9.5504548 -70.5898632
H 10.5030738 8.5688311 -70.3340113
C 11.2121939 9.9579073 -71.8004392
H 12.0155281 10.6371849 -71.5108146
C 10.3189662 10.6842638 -72.8220458
O 10.6706195 11.7496569 -73.3218699
C 11.8287769 8.7012708 -72.4308687
H 11.0787146 8.1829872 -73.0301747
H 12.1581597 8.0264929 -71.641054
O 12.9535704 9.016732 -73.2287619
H 13.2151123 8.2260534 -73.7115009
N 9.0917743 10.1931924 -73.0341901
H 8.8462209 9.3288767 -72.56349
C 8.0758771 10.7997598 -73.908273
H 8.5230395 10.9933282 -74.8827677
C 7.5601152 12.1572996 -73.4059811
O 6.9709603 12.9057875 -74.1813275
C 6.8980951 9.8177845 -74.0665242
H 6.5963806 9.4797408 -73.0746319
H 6.0469815 10.3468555 -74.4990079
C 7.2010356 8.596359 -74.9559661
H 8.1359891 8.1311667 -74.6495739
C 6.078985 7.5667695 -74.8172612
H 5.1265064 8.0041828 -75.1217947
H 6.002105 7.244648 -73.7790986
H 6.2918725 6.6974123 -75.4373065
C 7.3195826 8.9722824 -76.4354788
H 6.4050387 9.4644876 -76.7684964
H 7.4802548 8.0744735 -77.031481
H 8.1643764 9.6422728 -76.586479
N 7.8083199 12.5056056 -72.1428918
H 8.3267403 11.8644839 -71.5540116
C 7.4700463 13.8201725 -71.5844624
H 6.6249218 14.2248713 -72.140381
C 8.5965015 14.8537852 -71.7283394
O 8.4412966 15.9715581 -71.2569983
C 7.0096966 13.6702722 -70.1392581
H 6.7278152 14.646009 -69.7435907
H 7.8241777 13.2648047 -69.5378654
O 5.8818799 12.8084382 -70.0899695

H 6.1934362 11.8939971 -70.2199382
N 9.6527789 14.5194905 -72.4726668
H 9.6609405 13.583641 -72.8564542
C 10.8450282 15.3307806 -72.7823003
H 11.3215018 14.7681748 -73.5857787
C 11.9223128 15.3449434 -71.6844862
O 12.7208264 16.2703293 -71.5460031
C 10.4885248 16.6983789 -73.4123986
H 10.3052822 17.4216129 -72.6154718
H 9.5630553 16.5886969 -73.9810148
C 11.5483248 17.2610639 -74.3780375
O 12.3958398 16.4764423 -74.8659835
O 11.4371665 18.4657644 -74.7049311
N 11.9834183 14.26542 -70.8998863
H 11.310854 13.5258562 -71.0438426
C 13.0028759 14.0912496 -69.8714545
H 13.306885 15.0728525 -69.5156226
C 14.2829326 13.4339092 -70.3807713
O 14.2769208 12.3465484 -70.963321
C 12.4106606 13.3349922 -68.6923727
H 12.0504773 12.3665639 -69.0383334
H 13.215576 13.1470367 -67.9823596
C 11.3089208 14.0511588 -67.9372688
C 10.4143586 13.2895275 -67.1652494
H 10.4207411 12.2129036 -67.2452246
C 9.622269 13.9020251 -66.183757
H 9.0420531 13.2912381 -65.5123696
C 9.6896914 15.2939169 -66.0117228
O 9.0688287 15.8810816 -64.9659394
H 8.6517885 15.2197967 -64.4135997
C 10.476267 16.0688853 -66.8777947
H 10.5008894 17.1334103 -66.7732363
C 11.2869235 15.4568867 -67.8389113
H 11.9309688 16.084664 -68.4366982
N 15.413457 14.0666489 -70.0648519
H 15.3470006 14.8954175 -69.4865486
C 16.7466287 13.6072225 -70.4479379
H 16.6439033 13.0148765 -71.3578561
C 17.3845581 12.7089696 -69.3861672
O 17.0498046 12.785335 -68.2019845
C 17.6340751 14.8021464 -70.8021027
H 17.0888067 15.4656922 -71.4759625
H 18.5250258 14.4458703 -71.3186039
O 18.0387931 15.5263655 -69.6592734
H 17.3011331 15.5682376 -69.0191956
N 18.3085085 11.8526167 -69.839663

H 18.540374 11.8840546 -70.8182419
C 19.0346307 10.8940314 -69.0057114
H 19.4675616 10.1472293 -69.6713046
C 18.1121144 10.1118048 -68.0476025
O 18.3664568 10.0593789 -66.8463913
C 20.2060694 11.6112569 -68.3167241
H 19.8233584 12.3411857 -67.6012615
H 20.822286 12.1241624 -69.055329
H 20.8190484 10.8836954 -67.7848943
N 17.0082147 9.5547067 -68.5680509
H 16.8205954 9.6340163 -69.5538031
C 16.1258953 8.7216485 -67.7571872
H 16.0455573 9.2076642 -66.7871337
C 16.7726785 7.3640399 -67.4612841
O 17.037384 6.5613184 -68.3551552
C 14.6792575 8.6592675 -68.293561
H 14.2122908 9.6269136 -68.1101076
H 14.1203773 7.931987 -67.704949
C 14.479784 8.3334479 -69.7606206
O 14.1690558 7.2130685 -70.1375173
N 14.5042034 9.3331716 -70.6191546
H 14.1908225 9.1433714 -71.5603483
H 14.5384129 10.2922772 -70.2963576
N 17.0392665 7.1424113 -66.177116
H 16.830673 7.8928146 -65.5251381
C 17.5157217 5.8978433 -65.5954028
H 17.9177789 5.2616605 -66.384763
C 16.3357946 5.207524 -64.9288189
O 15.6159122 5.8225069 -64.1466295
C 18.6212937 6.1737034 -64.5672921
H 18.1711266 6.6156535 -63.6769974
H 19.0702786 5.2249355 -64.2716113
C 19.7093446 7.0913046 -65.0380035
C 19.7931824 8.4044157 -64.7311366
H 19.0720161 8.9404149 -64.124824
N 20.8504143 8.9783652 -65.4059943
H 20.9953366 9.9773354 -65.4322483
C 21.4912532 8.0630289 -66.2111008
C 22.5607112 8.1678796 -67.1126483
H 23.0547657 9.1163029 -67.2628505
C 22.976687 7.0234162 -67.8144749
H 23.7994497 7.0845023 -68.5131482
C 22.3164877 5.7977862 -67.6094385
H 22.6337933 4.9207785 -68.1562422
C 21.2368826 5.7069224 -66.707767
H 20.7248034 4.766601 -66.5741431

C 20.7952953 6.8365602 -65.9837824
N 16.1848563 3.912817 -65.176291
H 16.821799 3.469261 -65.8205527
C 15.0796393 3.1104436 -64.6775838
H 14.54308 3.6333506 -63.8833365
C 15.6247004 1.7985474 -64.1325112
O 16.0577958 0.9402323 -64.9020713
C 14.131694 2.8887033 -65.8487103
H 13.64995 3.8395378 -66.0623428
H 14.7289034 2.6128478 -66.7202805
C 13.0901348 1.8002141 -65.5654381
H 13.4487714 1.0489044 -64.8679001
C 11.8698551 2.4330043 -64.9143906
H 11.3898081 3.1481945 -65.575671
H 12.1699034 2.9421886 -63.9979327
H 11.1508884 1.6662489 -64.6483251
C 12.8433832 1.0456084 -66.8611406
H 12.3756177 1.6885177 -67.5980209
H 12.2143535 0.1981272 -66.6495824
H 13.7858171 0.6635186 -67.2549475
N 15.5022455 1.6123315 -62.8227831
H 15.0176951 2.3212374 -62.2749974
C 15.9819101 0.4156113 -62.1354336
H 16.5767406 -0.1688088 -62.8309248
C 14.7948239 -0.412656 -61.638592
O 14.0877462 0.0314062 -60.7298387
C 16.9228566 0.7713672 -60.9737509
H 16.3487296 1.1324943 -60.119959
C 17.7625409 -0.435787 -60.5611816
H 18.3692173 -0.7740933 -61.4018301
H 17.1180753 -1.2500821 -60.2345302
H 18.4224701 -0.1512396 -59.7425426
O 17.83844 1.7718527 -61.3585353
H 17.3314862 2.5714806 -61.5236629
N 14.5757734 -1.6381667 -62.1522123
C 13.7832369 -2.6243789 -61.4326093
H 12.8078798 -2.2156713 -61.1651937
C 14.5459745 -3.0172483 -60.1610187
O 15.6270337 -3.5983735 -60.2304277
C 13.5987399 -3.7952014 -62.405327
H 13.5382697 -4.7567764 -61.8952256
H 12.6985773 -3.6284725 -62.9974839
C 15.1494966 -2.2202696 -63.3610773
H 14.6833669 -1.7644417 -64.236083
H 16.2312348 -2.0961457 -63.4033807
C 14.823558 -3.7137458 -63.3158808

H 14.6132543 -4.1093829 -64.3096971
H 15.6551597 -4.2543635 -62.8618106
N 13.9662979 -2.746838 -58.9908767
H 13.0661721 -2.2754632 -59.0090904
C 14.5182136 -3.1102436 -57.6789229
H 15.4998097 -3.5591397 -57.8229977
C 13.7014467 -4.2044761 -56.9428997
O 13.6238671 -4.1604091 -55.7146881
C 14.714941 -1.81596 -56.8514598
H 13.7756341 -1.2643591 -56.8431172
H 14.9534945 -2.0737327 -55.819355
C 15.8229594 -0.873091 -57.3541501
H 15.6404761 -0.6146515 -58.3959849
C 15.8120489 0.4127376 -56.5303236
H 16.028931 0.1980959 -55.4826737
H 14.8309305 0.8833728 -56.593059
H 16.5554212 1.1109018 -56.9145674
C 17.2136708 -1.5018124 -57.2217831
H 17.3755269 -1.8389114 -56.1975999
H 17.9750578 -0.7631874 -57.470298
H 17.3151205 -2.3433035 -57.9050428
N 13.0948036 -5.214763 -57.6151661
C 12.2053006 -6.1721577 -56.9514439
H 11.3888186 -5.644905 -56.4617025
C 12.9249789 -7.0386965 -55.9075064
O 12.3195788 -7.4085346 -54.9111168
C 11.6162382 -7.0363963 -58.0682371
H 11.3764525 -8.0460584 -57.7312578
H 10.7270492 -6.5549761 -58.4702904
C 13.2765494 -5.6211828 -59.0043124
H 12.7053484 -4.9577645 -59.6524598
H 14.33053 -5.6213029 -59.2835882
C 12.7120005 -7.0340446 -59.1282169
H 12.3119336 -7.2235629 -60.1252091
H 13.4796364 -7.7662537 -58.8743209
N 14.2193922 -7.3309605 -56.0875591
H 14.6758111 -6.9951267 -56.9187628
C 15.0103271 -8.0990672 -55.112049
H 14.434692 -8.9910839 -54.8655987
C 15.2070223 -7.332927 -53.7964256
O 15.0766997 -7.9116657 -52.717904
C 16.3497863 -8.552893 -55.7413196
H 16.1067111 -9.1238207 -56.6393262
C 17.2457079 -7.3784555 -56.1790683
H 17.6104389 -6.833258 -55.3064419
H 16.7096291 -6.695959 -56.8363059

H 18.1109972 -7.7622999 -56.7213065
C 17.1758962 -9.4774166 -54.8192359
H 18.0864798 -9.7670527 -55.3455807
H 17.4716894 -8.935527 -53.9193684
C 16.4479887 -10.7630839 -54.4053729
H 16.0929498 -11.2919093 -55.2902479
H 15.605773 -10.5327737 -53.7532482
H 17.1389073 -11.4070984 -53.8607548
N 15.3935547 -6.0141343 -53.87863
H 15.4044782 -5.57695 -54.785056
C 15.4989416 -5.1573755 -52.7005961
H 16.1260958 -5.6549509 -51.9609007
C 14.1275042 -4.979588 -52.0493859
O 14.0041547 -5.2410058 -50.8625342
C 16.1529791 -3.8128398 -53.0612849
H 15.5453294 -3.3096024 -53.8136461
H 16.1749842 -3.1849001 -52.1697456
C 17.5888162 -3.9565456 -53.5969701
H 17.9604463 -2.9636573 -53.8583359
H 17.5825816 -4.5500643 -54.513358
C 18.5239075 -4.5941443 -52.560969
O 18.5957494 -5.8428049 -52.4662396
O 19.14147 -3.8633975 -51.7551244
N 13.0718701 -4.7526557 -52.834066
H 13.246795 -4.5814834 -53.8192455
C 11.6674641 -4.7573474 -52.3915481
H 11.5110988 -3.9424476 -51.6838844
C 11.2638044 -6.055375 -51.6705936
O 10.7000723 -5.9916155 -50.5815018
C 10.8225721 -4.4658085 -53.6373247
H 11.2703018 -4.9725609 -54.4847155
H 10.8980451 -3.3996945 -53.8162933
C 9.3400869 -4.8422281 -53.6032018
O 8.5960027 -4.4336681 -52.6834481
O 8.8625046 -5.4577929 -54.5871172
N 11.6917214 -7.2238873 -52.1577427
H 12.1378338 -7.2256039 -53.0701622
C 11.4803921 -8.5084706 -51.4804795
H 10.4192842 -8.5902366 -51.2439924
C 12.2581264 -8.616735 -50.1576347
O 11.7173718 -9.1125472 -49.1729519
C 11.8590517 -9.6523727 -52.4351888
H 12.9441046 -9.6885066 -52.5437256
H 11.4310558 -9.4628766 -53.4176041
C 11.3583046 -11.0199242 -51.9562242
H 11.7482812 -11.2164564 -50.9570363

H 11.7634607 -11.7769659 -52.6281464
S 9.5513028 -11.2041661 -51.9296248
C 9.4280423 -12.9497713 -51.463499
H 8.3802362 -13.2482151 -51.4521366
H 9.8585708 -13.0948903 -50.4720088
H 9.9653893 -13.5614913 -52.1881289
N 13.4992124 -8.1155487 -50.1013757
H 13.8751445 -7.7085546 -50.9531837
C 14.3462607 -8.0992723 -48.8917458
H 14.3261654 -9.0880359 -48.4302966
C 13.8414349 -7.109273 -47.8321094
O 13.8696885 -7.4088854 -46.6438106
C 15.7803139 -7.7771225 -49.3386629
H 16.0728206 -8.4953921 -50.1072518
H 15.78842 -6.7853287 -49.7877521
C 16.8146607 -7.8333564 -48.2019743
H 16.5459291 -7.1366515 -47.4067893
H 16.8280532 -8.8437172 -47.7908348
C 18.2187971 -7.4873246 -48.7193819
H 18.4184365 -8.0829343 -49.6128579
H 18.9536731 -7.7499724 -47.9570584
C 18.3487833 -5.992413 -49.0458139
H 18.4319706 -5.4217815 -48.1165061
H 17.4451803 -5.6561907 -49.5619324
N 19.5031294 -5.7293883 -49.9309239
H 20.3954323 -5.9966135 -49.5549232
H 19.5174901 -4.749167 -50.2346865
H 19.3464276 -6.1609146 -50.848426
N 13.3322727 -5.9630881 -48.2710839
H 13.3916351 -5.7966786 -49.2700395
C 12.6752334 -4.9259791 -47.4652083
H 13.3066394 -4.639705 -46.6204001
C 11.3514425 -5.4659331 -46.907204
O 11.0801258 -5.3692947 -45.708615
C 12.463281 -3.6936637 -48.3766892
H 12.0638674 -4.0566426 -49.3241162
C 11.3939495 -2.735481 -47.8251092
H 11.5699158 -2.5482408 -46.7698325
H 10.4027027 -3.1721763 -47.9439616
H 11.4167651 -1.795219 -48.3641339
C 13.8090055 -3.0011842 -48.7056933
H 14.1943818 -2.5034339 -47.8276267
H 14.5553025 -3.7476582 -48.9678682
C 13.7415405 -1.9940272 -49.8643963
H 13.1388901 -1.128049 -49.5957665
H 13.3172396 -2.4665957 -50.7493734

H 14.7489847 -1.6485579 -50.0979748
N 10.5569209 -6.1251239 -47.7575478
H 10.8058184 -6.1502801 -48.7417588
C 9.3406062 -6.8155937 -47.3499945
H 8.687436 -6.0980664 -46.8515223
C 9.6451199 -7.9320667 -46.3426887
O 8.9789309 -7.9857565 -45.3181334
C 8.6090851 -7.3439192 -48.5900398
H 9.2347282 -8.0646011 -49.1166765
H 8.3743305 -6.5167723 -49.2610325
H 7.6825766 -7.8315087 -48.286617
N 10.6967849 -8.7327052 -46.5459848
H 11.1877227 -8.6812436 -47.4322827
C 11.1410652 -9.7413254 -45.5766499
H 10.3095113 -10.4188421 -45.3765601
C 11.5417736 -9.1181235 -44.2316211
O 11.0027769 -9.5243077 -43.2065835
C 12.2879134 -10.5821685 -46.1574739
H 13.1383181 -9.9410307 -46.3900585
H 11.9539012 -11.054893 -47.0827182
C 12.7241101 -11.6616207 -45.1622198
O 13.5224159 -11.3337599 -44.252894
O 12.2018212 -12.793503 -45.265959
N 12.3829782 -8.0769071 -44.2118859
H 12.8033958 -7.7736289 -45.0842588
C 12.8418208 -7.4756884 -42.9545116
H 13.20788 -8.2795145 -42.315361
C 11.7105569 -6.7843906 -42.179033
O 11.5105007 -7.0762136 -40.995856
C 14.0107534 -6.5158331 -43.2086705
H 14.8199205 -7.0620265 -43.6953675
H 13.6867322 -5.7238326 -43.8863098
C 14.5352889 -5.889822 -41.9505213
C 15.3513898 -6.4920806 -41.0560262
H 15.7532765 -7.4912347 -41.1718495
N 15.5335855 -5.6654725 -39.9629168
H 16.0897329 -5.9158626 -39.160673
C 14.8148192 -4.49376 -40.0872147
C 14.6321954 -3.3866221 -39.242576
H 15.1032791 -3.3500841 -38.2735638
C 13.819259 -2.3236695 -39.676404
H 13.6551516 -1.4597462 -39.0434416
C 13.2231146 -2.3758491 -40.9490175
H 12.61592 -1.5480294 -41.2963996
C 13.413713 -3.4925454 -41.785488
H 12.9690569 -3.4997165 -42.7718436

C 14.1985453 -4.5894945 -41.3712467
N 10.9043925 -5.9527161 -42.8492866
H 11.1134384 -5.7630804 -43.8240366
C 9.7438538 -5.2921988 -42.2212211
H 10.0844229 -4.773692 -41.3238945
C 8.6828352 -6.2978319 -41.7797415
O 8.1475363 -6.1684389 -40.6765001
C 9.0796136 -4.2487583 -43.1304959
H 8.177974 -3.8895772 -42.6371761
C 9.9720763 -3.0467225 -43.4104666
H 10.8669468 -3.3593763 -43.9445606
H 10.2661708 -2.5763767 -42.4714854
H 9.4314004 -2.3171273 -44.0117579
O 8.7181591 -4.8066588 -44.3726664
H 9.5496823 -4.9479732 -44.8577988
N 8.4358032 -7.3581704 -42.5633335
H 8.8963046 -7.4314515 -43.466883
C 7.5723311 -8.4495715 -42.1268535
H 6.6350766 -8.0155664 -41.7795424
C 8.1700975 -9.1517994 -40.9247922
O 7.4758181 -9.174459 -39.9330543
C 7.2448461 -9.4620492 -43.2330206
H 8.1639771 -9.7685181 -43.7320877
H 6.8107539 -10.3548548 -42.779457
C 6.239774 -8.9211213 -44.2597927
H 6.2060887 -9.6067299 -45.1067549
H 6.5649357 -7.9487869 -44.6272582
C 4.8301142 -8.7907822 -43.6962677
O 4.0185217 -9.6936336 -43.7800733
N 4.4373901 -7.6329512 -43.2092787
H 3.4549186 -7.5555601 -42.9669857
H 5.0762198 -6.8618807 -43.140931
N 9.4273277 -9.6006942 -40.9284614
H 9.9686587 -9.4839231 -41.780828
C 10.0214635 -10.412134 -39.8567427
H 9.3084547 -11.1990181 -39.609992
C 10.253828 -9.6601678 -38.5291229
O 10.2250454 -10.2893742 -37.4666527
C 11.2944858 -11.0964365 -40.3945858
H 11.8618845 -11.5134649 -39.5643313
H 11.9244235 -10.3674132 -40.904203
C 10.9694015 -12.2521054 -41.3360884
O 10.0165345 -12.9890257 -41.1245311
N 11.7728694 -12.4937022 -42.343769
H 11.4995019 -13.1733068 -43.0362986
H 12.499172 -11.8467642 -42.6449554

N 10.2929531 -8.3240194 -38.537167
H 10.3624872 -7.8437779 -39.4304161
C 10.2075646 -7.5261957 -37.311539
H 10.9370343 -7.9083139 -36.5955343
C 8.8173012 -7.6078404 -36.6366541
O 8.7224682 -7.6976754 -35.4119872
C 10.5911494 -6.0827475 -37.6571958
H 9.8848154 -5.6694188 -38.3793169
H 11.5923603 -6.0574983 -38.0909011
H 10.5793001 -5.471428 -36.7549126
N 7.7281362 -7.648853 -37.4093658
H 7.8464977 -7.6620426 -38.4175533
C 6.3615845 -7.7065981 -36.8837774
H 6.2525433 -6.8732437 -36.1891368
C 5.9991926 -8.9928891 -36.0905116
O 5.3798625 -8.8602907 -35.0312186
C 5.3875801 -7.4605357 -38.0417105
H 5.317507 -8.3466364 -38.6717451
H 5.7360643 -6.6360806 -38.6607618
H 4.3964641 -7.2320804 -37.651936
N 6.3537059 -10.227742 -36.5201682
C 6.0921003 -11.4546801 -35.7992746
H 5.0595046 -11.4728627 -35.4552056
C 7.0141917 -11.5868196 -34.5825291
O 6.5442037 -12.022288 -33.5387212
C 6.2944393 -12.5878507 -36.8162142
H 6.5796654 -13.5239315 -36.3340225
H 5.3790469 -12.7278175 -37.3944718
C 6.9710996 -10.6108314 -37.7595143
H 6.2311127 -10.4906961 -38.55036
H 7.8621115 -10.0271148 -37.9027458
C 7.3852351 -12.0701274 -37.7139111
H 7.3632158 -12.5333561 -38.7012058
H 8.3603036 -12.1870356 -37.2408137
N 8.2600128 -11.0993172 -34.6473682
H 8.5974122 -10.7351963 -35.529587
C 9.1638845 -11.0324667 -33.493428
H 9.2559856 -12.0264496 -33.0531509
C 8.6246533 -10.1030979 -32.3918366
O 8.5898426 -10.4804776 -31.2234502
C 10.5463395 -10.5906214 -33.9929249
H 10.9400176 -11.3588206 -34.6604347
H 10.4349591 -9.6795107 -34.5821898
C 11.5645544 -10.3336599 -32.8982995
C 12.1676299 -11.4139774 -32.226674
H 11.9007571 -12.4282594 -32.4853161

C 13.115578 -11.1764532 -31.2147571
H 13.5750579 -12.007349 -30.6985902
C 13.464218 -9.8578058 -30.873536
H 14.194 -9.6750549 -30.0974929
C 12.8624226 -8.7769348 -31.541472
H 13.1324783 -7.7636193 -31.281157
C 11.9107974 -9.0133361 -32.5496745
H 11.4482941 -8.1777601 -33.0568941
N 8.0603085 -8.9495652 -32.7631566
H 8.1434311 -8.665565 -33.7337643
C 7.4168677 -8.0211198 -31.8271501
H 7.9214617 -8.1082216 -30.8649975
C 5.9315139 -8.326438 -31.5456683
O 5.2535744 -7.4801421 -30.961817
C 7.5941536 -6.574922 -32.324633
H 7.124499 -6.4941971 -33.3064549
H 7.0538484 -5.9068789 -31.6550215
C 9.0136829 -6.0386909 -32.3987931
C 9.4463363 -5.359517 -33.5561945
H 8.7912065 -5.2787379 -34.4129759
C 10.7433545 -4.8150995 -33.6203321
H 11.075635 -4.3075551 -34.5142454
C 11.6148391 -4.9453172 -32.5180674
O 12.8717441 -4.435554 -32.5709646
H 13.0345853 -3.9454995 -33.3800472
C 11.174962 -5.6029107 -31.3511048
H 11.8453005 -5.6829437 -30.507972
C 9.8769556 -6.144496 -31.2893049
H 9.5563376 -6.6474406 -30.3872537
N 5.3871766 -9.481238 -31.956426
H 6.0033601 -10.1538206 -32.4006634
C 3.9274016 -9.6951489 -32.0875758
H 3.5788743 -9.0167038 -32.8656313
C 3.101727 -9.3795924 -30.8315989
O 1.9946272 -8.8568 -30.9597617
C 3.6184426 -11.1199467 -32.5819641
H 4.1373467 -11.2775613 -33.5237867
C 4.0262775 -12.229422 -31.6076246
H 3.4924745 -12.148523 -30.6645446
H 5.0959923 -12.1726016 -31.405786
H 3.820187 -13.1996915 -32.0601955
O 2.2341245 -11.2757929 -32.8256162
H 2.1031661 -11.0808149 -33.7675107
N 3.6312656 -9.6287895 -29.6257628
H 4.5649006 -10.0097299 -29.5872965
C 2.9405287 -9.3177676 -28.3687039

H 1.9221669 -9.6969241 -28.445873
C 2.8276563 -7.8017421 -28.1507249
O 1.7167741 -7.2833488 -28.1555535
C 3.6362754 -10.0368499 -27.1973246
H 3.6353806 -11.1098669 -27.3948457
H 4.6745595 -9.7054275 -27.1474315
C 2.978396 -9.7839054 -25.8233671
H 2.9966347 -8.7178673 -25.6005677
C 1.5310448 -10.2812004 -25.7697653
H 1.4817257 -11.3377576 -26.0293787
H 0.9096754 -9.6938931 -26.4446131
H 1.1425322 -10.1385847 -24.7604909
C 3.7720642 -10.514862 -24.7416685
H 3.7563083 -11.5903824 -24.9153329
H 3.3368301 -10.2987848 -23.7649057
H 4.8026104 -10.1599394 -24.7388168
N 3.9616382 -7.0915666 -28.1298216
H 4.8305866 -7.5964343 -28.1841824
C 4.0327294 -5.6286734 -27.9990707
H 3.5475118 -5.3373421 -27.066531
C 3.3223149 -4.9052738 -29.1588863
O 2.7887892 -3.8112358 -29.0000078
C 5.522688 -5.2420034 -27.9073454
H 5.9417744 -5.7125461 -27.0159418
H 6.0387796 -5.6508386 -28.77724
C 5.8148372 -3.7290084 -27.839953
H 5.4302001 -3.2507583 -28.740477
C 5.2052008 -3.0369301 -26.6265279
H 5.5270028 -3.5308572 -25.7079944
H 4.1165442 -3.0902357 -26.6670599
H 5.5002705 -1.9890255 -26.5938106
C 7.3263404 -3.5054407 -27.8182893
H 7.7487717 -3.9367071 -26.9075049
H 7.5460048 -2.4366408 -27.8259105
H 7.7977514 -3.9704279 -28.681937
N 3.3240981 -5.4922525 -30.3565642
H 3.8200164 -6.3736162 -30.4481514
C 2.6217824 -4.9657189 -31.5266407
H 2.8835245 -3.9148142 -31.6417372
C 1.0943839 -5.029999 -31.3514762
O 0.4108227 -4.0473604 -31.6319395
C 3.127867 -5.7370606 -32.7582961
H 4.209855 -5.607543 -32.8206349
H 2.9292394 -6.7975108 -32.6056175
C 2.5111026 -5.3318787 -34.1077532
H 1.4532277 -5.5947615 -34.1197899

C 2.6686143 -3.8435188 -34.4097948
H 3.7203921 -3.5715168 -34.3494023
H 2.0853599 -3.2554452 -33.704076
H 2.2981698 -3.6320755 -35.4139741
C 3.226156 -6.095398 -35.2194507
H 4.2560045 -5.7514367 -35.3113525
H 2.7083162 -5.937219 -36.1650524
H 3.2404669 -7.1588567 -34.9842874
N 0.5617721 -6.1321914 -30.8100106
H 1.1854655 -6.8955181 -30.565335
C -0.8774119 -6.2932057 -30.5294087
H -1.4450707 -5.8133694 -31.3273322
C -1.3387857 -5.5989339 -29.2472027
O -2.4972833 -5.2107496 -29.1799714
C -1.2259879 -7.7857952 -30.5059816
H -0.5395652 -8.3161481 -29.8425807
H -2.2423577 -7.9080194 -30.1304602
C -1.1550235 -8.3653732 -31.9235204
H -0.1595575 -8.2283015 -32.3457716
H -1.8592111 -7.825137 -32.5503861
C -1.517416 -9.8499764 -31.9535904
H -1.7789138 -10.1101266 -32.9786297
H -2.3926915 -10.0247935 -31.3275004
N -0.3815104 -10.6888054 -31.5472774
H 0.4686067 -10.5891001 -32.0922565
C -0.2819976 -11.5284697 -30.5397689
N 0.7987728 -12.2399241 -30.4453776
H 0.9213052 -12.9191248 -29.7195572
H 1.4926744 -12.1243647 -31.1788985
N -1.2140364 -11.691179 -29.6413828
H -2.0524106 -11.1461476 -29.7056909
H -1.0837421 -12.3425586 -28.8909044
N -0.4359914 -5.4137744 -28.291648
H 0.4565919 -5.878876 -28.4106519
C -0.6121594 -4.6637386 -27.0405899
H -1.5484464 -4.9448234 -26.5574101
C -0.6419177 -3.1442925 -27.2699944
O -1.4719593 -2.454789 -26.6849815
C 0.5604857 -5.0831334 -26.1452238
H 0.4619065 -6.1502756 -25.9371324
H 1.4675594 -4.941078 -26.7285774
C 0.7664583 -4.3591921 -24.8161269
H -0.1072872 -4.5150072 -24.1788573
H 0.8875321 -3.2879945 -24.9917974
C 2.035639 -4.9226683 -24.1581527
O 1.8929473 -5.6917077 -23.1831504

O 3.1384206 -4.6239782 -24.6732221
N 0.1917402 -2.6286023 -28.1822229
H 0.9142711 -3.2259607 -28.5641261
C 0.1832659 -1.2150978 -28.5658876
H 0.0037262 -0.6137299 -27.6722889
C -0.9246664 -0.8558517 -29.5768898
O -1.3427934 0.2986789 -29.6289793
C 1.5712314 -0.8568396 -29.111109
H 1.7755088 -1.4407739 -30.0082925
H 2.3275326 -1.0683264 -28.3535512
H 1.5991124 0.2052893 -29.3567835
N -1.3690744 -1.8151607 -30.399589
H -0.9570034 -2.7323259 -30.3107115
C -2.4122034 -1.6197109 -31.4130676
H -2.8401389 -0.6237023 -31.2850044
C -3.5838278 -2.5891446 -31.1989809
O -4.4984085 -2.2893246 -30.4387788
C -1.8158924 -1.6685056 -32.833723
H -1.3145361 -2.624073 -32.987596
H -2.63415 -1.6141187 -33.553035
C -0.8409829 -0.5505884 -33.1512774
C -1.3096435 0.649862 -33.7150379
H -2.3647469 0.7833783 -33.9051926
C 0.531843 -0.7025696 -32.8802269
H 0.8864084 -1.6149005 -32.4260483
C -0.4110316 1.6879453 -34.01348
H -0.7806818 2.6123909 -34.4308109
C 1.4300197 0.3388147 -33.1757978
H 2.4760809 0.2347951 -32.9481857
C 0.9602995 1.5313944 -33.7512922
H 1.6466328 2.3356076 -33.9776088
N -3.6205628 -3.7132496 -31.9236746
H -2.8479066 -3.9034345 -32.5436994
C -4.6046085 -4.7861899 -31.7255437
H -4.6477881 -5.0219769 -30.6606922
C -4.2176418 -6.0541322 -32.4919207
O -3.4466607 -6.006089 -33.4557384
C -6.0082382 -4.3507814 -32.1811528
H -6.3304691 -3.4699273 -31.624698
H -6.7175425 -5.152142 -31.9705056
O -6.0325689 -4.0692292 -33.5679786
H -5.899994 -3.1163384 -33.6518697
N -4.8741413 -7.1737827 -32.163381
H -5.4716506 -7.1487112 -31.3494907
C -4.8665038 -8.4037354 -32.9751168
H -3.8421984 -8.7622554 -33.024553

C -5.296619 -8.1459118 -34.4291164
O -4.6251069 -8.5946963 -35.3565651
C -5.7191938 -9.4992887 -32.2851475
H -5.2617698 -9.6962688 -31.3135578
C -7.1749464 -9.0643969 -32.0189038
H -7.7180113 -8.9252308 -32.9539572
H -7.2157387 -8.1483551 -31.4309694
H -7.6857446 -9.8419113 -31.4478146
C -5.7591062 -10.8389807 -33.0521625
H -6.3486961 -11.5529072 -32.4750014
H -6.2623358 -10.6958331 -34.0075932
C -4.3868889 -11.4732058 -33.3065014
H -3.86465 -11.6184485 -32.3614838
H -3.7929475 -10.8453952 -33.9693974
H -4.5241808 -12.4437534 -33.7840225
N -6.3197801 -7.3081212 -34.6459337
H -6.7867577 -6.9174776 -33.8407924
C -6.8108968 -6.9429317 -35.9822531
H -6.9295796 -7.8697976 -36.5438468
C -5.8010042 -6.0946841 -36.7703672
O -5.7534856 -6.2049544 -37.9993011
C -8.1929232 -6.2678168 -35.828368
H -8.8936588 -6.9935093 -35.4113768
H -8.1035006 -5.4462128 -35.1147961
C -8.795135 -5.6932521 -37.1238768
H -9.7559562 -5.2332397 -36.8894435
H -8.1411857 -4.8942535 -37.4703831
C -9.0162358 -6.7421972 -38.2346595
H -9.9843958 -7.2211919 -38.076221
H -8.2596339 -7.5233973 -38.1830747
C -8.9707821 -6.1540308 -39.6519625
H -9.7832162 -5.4324325 -39.7803459
H -9.1312743 -6.9749907 -40.3575952
N -7.6582374 -5.5172426 -39.9201159
H -7.5238449 -4.6979612 -39.3466649
H -7.5232324 -5.260878 -40.8933727
H -6.8930147 -6.1433565 -39.6825251
N -4.9831557 -5.2914209 -36.0868715
H -5.1114733 -5.2316311 -35.0833861
C -3.8621657 -4.5557308 -36.6755995
H -4.2125805 -3.9476157 -37.5101316
H -3.4303927 -3.8987074 -35.9202956
C -2.7652385 -5.4947951 -37.177779
O -2.4814048 -5.5085537 -38.373279
N -2.2677174 -6.3806718 -36.3070245
H -2.5945535 -6.342489 -35.3479413

C -1.2607822 -7.3958332 -36.6512525
H -0.3445124 -6.8925585 -36.961776
C -1.6939054 -8.2984128 -37.8176158
O -0.9311845 -8.4717807 -38.7673839
C -0.9587692 -8.2239316 -35.3945171
H -0.5126998 -7.5591821 -34.6545479
H -1.9003645 -8.5918536 -34.9852958
C -0.0205586 -9.403772 -35.5957312
C -0.476939 -10.7237244 -35.4080804
H -1.509788 -10.9114615 -35.1526382
C 0.4018223 -11.8087914 -35.5828742
H 0.0411055 -12.8215519 -35.4605813
C 1.7396798 -11.5814567 -35.9510346
H 2.4073452 -12.4177367 -36.1126058
C 2.197147 -10.2669613 -36.1448257
H 3.2167878 -10.0866761 -36.4561706
C 1.3158525 -9.1847377 -35.975059
H 1.6594669 -8.1812815 -36.1674815
N -2.9508894 -8.7602365 -37.8162996
H -3.5180262 -8.6013591 -36.9897401
C -3.5164419 -9.5705426 -38.9055008
H -2.8830213 -10.4489155 -39.0412821
C -3.4884375 -8.8191223 -40.2441416
O -3.0854456 -9.4032043 -41.2444796
C -4.9353343 -10.0606912 -38.5229828
H -5.4930771 -9.2156787 -38.1147458
C -5.7199256 -10.6008875 -39.7362459
H -5.1748012 -11.428586 -40.1934969
H -5.8704642 -9.8203191 -40.4808494
H -6.7059161 -10.950001 -39.4339102
C -4.8187313 -11.1576382 -37.4374046
H -4.3660146 -12.0503218 -37.8713733
H -4.1598068 -10.814504 -36.6407699
C -6.1553173 -11.5460725 -36.7923586
H -6.7955629 -12.0580817 -37.5094256
H -6.6616767 -10.6550823 -36.4198857
H -5.9674939 -12.2254155 -35.9610021
N -3.8323925 -7.5248185 -40.2879394
H -4.0656676 -7.0556824 -39.4218744
C -3.7691065 -6.745843 -41.5391746
H -4.3005425 -7.2994218 -42.3139513
C -2.3393446 -6.5519942 -42.0526116
O -2.1145754 -6.6247537 -43.2621331
C -4.4332741 -5.3718712 -41.3703427
H -3.8537605 -4.6109815 -41.8970195
H -4.4733346 -5.1006908 -40.3138927

O -5.7412933 -5.3987764 -41.9212897
H -5.6390313 -5.3843889 -42.8827614
N -1.3861787 -6.3126124 -41.1495095
H -1.6547347 -6.2826085 -40.1714205
C 0.0158965 -6.0425797 -41.481886
H 0.0577829 -5.3112942 -42.2894989
C 0.740751 -7.3012983 -41.9896076
O 1.4825599 -7.2298511 -42.9719905
C 0.6993255 -5.4537121 -40.2332656
H 0.5759614 -6.161702 -39.4119538
H 1.768021 -5.3583632 -40.4270006
C 0.1460188 -4.0790425 -39.7977291
H -0.9410754 -4.0968931 -39.8068985
C 0.5881962 -3.7787531 -38.3651539
H 1.6688825 -3.8436922 -38.2817429
H 0.1358835 -4.4987693 -37.6855946
H 0.2728934 -2.7773283 -38.0815491
C 0.5838749 -2.9467471 -40.7300837
H 1.6679734 -2.8669288 -40.751094
H 0.1714852 -2.0035095 -40.3724266
H 0.2150466 -3.1247907 -41.7383577
N 0.4550266 -8.4611916 -41.3890336
H -0.1379686 -8.4416052 -40.564342
C 0.9646168 -9.7595951 -41.8364496
H 2.0196674 -9.6433663 -42.0892642
C 0.2485841 -10.2596498 -43.1035624
O 0.8987943 -10.7616169 -44.0158858
C 0.8455888 -10.7511466 -40.6638834
H 1.3523783 -10.326376 -39.7955445
H -0.210257 -10.8640917 -40.4110157
C 1.4357562 -12.1477194 -40.9424942
H 0.9147976 -12.6038002 -41.7839374
C 2.9345315 -12.1074372 -41.2461209
H 3.4682867 -11.5756565 -40.4587843
H 3.1048157 -11.6059846 -42.1990644
H 3.3258448 -13.1209115 -41.3293987
C 1.228386 -13.036948 -39.7157139
H 1.7534838 -12.6171902 -38.8578697
H 1.6078131 -14.0384338 -39.9180822
H 0.1647686 -13.1057515 -39.4881301
N -1.0698244 -10.0583188 -43.2209953
H -1.5784126 -9.6705319 -42.4337335
C -1.8288138 -10.4543792 -44.4133578
H -1.6406428 -11.512455 -44.6026739
C -1.3632127 -9.6903915 -45.6534104
O -1.0336879 -10.3147171 -46.6612399

C -3.3368109 -10.2801862 -44.1613091
H -3.5384081 -9.2595103 -43.8397539
H -3.6148275 -10.9493552 -43.3461491
C -4.2354917 -10.6192243 -45.3629775
H -5.2387681 -10.8120393 -44.981554
H -3.8764807 -11.5469091 -45.8110439
S -4.390968 -9.3657168 -46.6781462
C -5.1799099 -8.0079527 -45.7729399
H -5.4383667 -7.2116522 -46.4701612
H -4.4931662 -7.6138861 -45.0242914
H -6.0870682 -8.3705426 -45.2881865
N -1.2836702 -8.3547994 -45.597801
H -1.5351508 -7.8696527 -44.7427603
C -0.9788531 -7.5879528 -46.8043397
H -1.457325 -8.1034868 -47.6360249
C 0.521404 -7.5756896 -47.1451478
O 0.8637811 -7.5102657 -48.3264741
C -1.6430715 -6.2032923 -46.748782
H -1.1355764 -5.5871202 -46.0089988
H -2.6802938 -6.3144597 -46.4385527
C -1.6394516 -5.5151375 -48.1094585
N -0.9709235 -4.389742 -48.2157069
H -0.4685428 -4.0199304 -47.4270163
H -0.9501478 -3.9346266 -49.1103779
O -2.2486604 -5.9590541 -49.0785727
N 1.416904 -7.7516498 -46.1678263
H 1.0980991 -7.8154289 -45.2078644
C 2.8375119 -7.9866019 -46.4363406
H 3.1969886 -7.2710589 -47.1758242
H 3.3911814 -7.8211393 -45.5225839
C 3.1279278 -9.4043741 -46.945777
O 3.8298141 -9.5493483 -47.9453507
N 2.447086 -10.4225332 -46.4101552
H 1.9315207 -10.2624063 -45.5490469
C 2.4854058 -11.7979805 -46.9139969
H 3.5152832 -12.1562071 -46.9087821
H 1.8931677 -12.4278665 -46.2504679
C 1.9239234 -11.9295401 -48.3328251
O 2.5932635 -12.4826701 -49.2022689
N 0.7929469 -11.2777152 -48.628854
H 0.267872 -10.8632915 -47.8641561
C 0.2683398 -11.1228987 -49.9938683
H 0.0573556 -12.1067264 -50.4128921
C 1.2826057 -10.4351154 -50.919821
O 1.4660857 -10.8427349 -52.0678355
C -1.03995 -10.3126457 -49.9511539

H -1.8060414 -10.9034882 -49.4476874
H -0.8743164 -9.4136616 -49.3588149
C -1.5543657 -9.8836418 -51.2950987
C -1.1187229 -8.7991881 -51.9758444
H -0.3563124 -8.1147227 -51.6133052
N -1.6791194 -8.7882528 -53.2363242
H -1.3736528 -8.1625098 -53.9676252
C -2.5438482 -9.8457334 -53.4186151
C -3.3411695 -10.2526051 -54.4992947
H -3.3476281 -9.6931163 -55.422582
C -4.1539985 -11.3894695 -54.358604
H -4.801853 -11.6994674 -55.1695271
C -4.1314936 -12.1172815 -53.1550939
H -4.7589197 -12.9925633 -53.044334
C -3.3101688 -11.7105559 -52.0842815
H -3.3046579 -12.2804389 -51.1652787
C -2.499787 -10.5592805 -52.1831049
N 1.96428 -9.3993704 -50.4253499
H 1.7809878 -9.1156849 -49.4709382
C 2.9636726 -8.654149 -51.2011413
H 2.5137098 -8.3680451 -52.1510126
C 4.1751778 -9.5252683 -51.5380323
O 4.584636 -9.5468988 -52.6991712
C 3.3562456 -7.3613529 -50.4714756
H 3.7076416 -7.5770001 -49.4645344
C 4.4220107 -6.5580398 -51.2039862
H 4.0784879 -6.3762013 -52.2179027
H 5.3645424 -7.1059384 -51.2128172
H 4.574357 -5.6062331 -50.6915057
O 2.2203371 -6.5288334 -50.4111686
H 1.6945377 -6.8265429 -49.6442254
N 4.6517677 -10.3483214 -50.5991615
H 4.2697926 -10.2894075 -49.6593257
C 5.6772822 -11.3580867 -50.8568759
H 6.5451977 -10.8700353 -51.3041686
C 5.1923812 -12.4341222 -51.8490049
O 5.8968642 -12.7354913 -52.8110289
C 6.1067911 -11.9647842 -49.5145566
H 5.2580115 -12.4435631 -49.0255728
H 6.499734 -11.1841373 -48.8617151
H 6.8863492 -12.7084471 -49.6781218
N 3.9534366 -12.9249302 -51.7086818
H 3.4267844 -12.6496469 -50.8836684
C 3.334672 -13.8997941 -52.6291921
H 3.9737587 -14.7791788 -52.6745089
C 3.2311635 -13.3581777 -54.0601941

O 3.501972 -14.0879971 -55.0132972
C 1.9611062 -14.3607064 -52.0920771
H 1.4005589 -13.4924383 -51.7508018
C 1.0987965 -15.094003 -53.1308964
H 1.6582642 -15.9270397 -53.5595551
H 0.8015677 -14.4086898 -53.9226587
H 0.1946828 -15.4764729 -52.6554527
C 2.1541222 -15.3188572 -50.9094302
H 2.6359165 -16.2380737 -51.2437664
H 1.18814 -15.559973 -50.4658729
H 2.7765934 -14.8574841 -50.1444956
N 2.9105554 -12.073782 -54.2549198
H 2.6628376 -11.510579 -53.4444772
C 2.8382215 -11.4848562 -55.603466
H 2.5477263 -12.2731829 -56.294507
C 4.1932573 -11.0523229 -56.1735234
O 4.3050569 -11.0014922 -57.3976465
C 1.7500749 -10.3972881 -55.6684607
H 1.973677 -9.6224873 -54.932472
H 1.7766916 -9.9369756 -56.657886
C 0.3181127 -10.9321047 -55.4242165
H 0.2277581 -11.265099 -54.390261
H -0.3748739 -10.1024204 -55.5677658
C -0.0987512 -12.0894509 -56.3560436
H 0.544746 -12.9544503 -56.17014
H 0.0153415 -11.7777132 -57.39766
C -1.5577073 -12.5187353 -56.1029819
H -2.2269318 -11.6802266 -56.3325773
H -1.6736438 -12.7561101 -55.0366384
N -1.920309 -13.7026939 -56.930816
H -1.8494992 -13.5031153 -57.9228767
H -2.903941 -13.9547134 -56.8042702
H -1.3493329 -14.5134468 -56.71273
N 5.239108 -10.8811623 -55.358782
H 5.0832237 -10.9063656 -54.3572045
C 6.6252368 -10.7490509 -55.8568326
H 6.6136699 -10.1725473 -56.7835268
C 7.23075 -12.111499 -56.2120809
O 7.8163342 -12.237301 -57.2830009
C 7.5248285 -9.9796415 -54.8686851
H 7.4639628 -10.4501565 -53.8877875
C 8.9905939 -9.944826 -55.3227672
H 9.0568977 -9.5997377 -56.3555774
H 9.4372889 -10.9357012 -55.2428839
H 9.5579733 -9.2635562 -54.6896495
C 7.0793244 -8.5186274 -54.7449023

H 7.2214826 -8.0117064 -55.6995769
H 7.6884329 -8.0243809 -53.9867521
H 6.0354079 -8.4688468 -54.4529373
N 7.0278598 -13.1595392 -55.4024952
H 6.5409429 -13.011564 -54.5230944
C 7.6271667 -14.4858677 -55.6559459
H 8.6634613 -14.3031979 -55.9438593
C 6.9906641 -15.2279986 -56.8419336
O 7.6716931 -15.9879257 -57.5225639
C 7.6575082 -15.3317962 -54.3638388
H 7.9124323 -14.6589608 -53.5431137
C 6.2996944 -15.9707711 -54.0400428
H 6.0816508 -16.7851927 -54.7321286
H 5.5262014 -15.2171777 -54.1221082
H 6.2972443 -16.3543042 -53.0201916
C 8.777694 -16.3911983 -54.4438692
H 8.5331507 -17.1303691 -55.2074872
H 9.7068405 -15.8989223 -54.7355244
C 9.0351373 -17.1189259 -53.1182629
H 8.173149 -17.7267722 -52.8436591
H 9.2387219 -16.3944539 -52.3289923
H 9.8999306 -17.7735446 -53.2299662
N 5.7387061 -14.9039989 -57.1868191
H 5.2127234 -14.3222172 -56.5485866
C 5.1062204 -15.3009733 -58.4534771
H 5.3874356 -16.3318692 -58.6680816
C 5.6030433 -14.4874707 -59.672003
O 5.0849158 -14.6599443 -60.773668
C 3.5735904 -15.2270815 -58.2905951
H 3.3168007 -14.2373792 -57.910768
H 3.1057086 -15.3358337 -59.269651
C 2.9775319 -16.3018904 -57.3585176
H 3.4899355 -16.2918303 -56.3990126
C 1.4957103 -15.9962792 -57.1210999
H 0.9610944 -15.9934025 -58.0710052
H 1.4152946 -15.0272253 -56.6348142
H 1.0717324 -16.755584 -56.4636815
C 3.0815974 -17.7101515 -57.947436
H 2.616133 -17.7444842 -58.9324024
H 2.5847047 -18.4241014 -57.2906702
H 4.1275589 -18.0041337 -58.0324987
N 6.5584693 -13.5695965 -59.4840437
H 6.9635998 -13.4805897 -58.559394
C 7.1419561 -12.7294521 -60.533019
H 8.0300305 -12.2428227 -60.1296168
H 7.4428545 -13.3604889 -61.3694312

C 6.2113295 -11.6372743 -61.0664694
O 6.4668133 -11.1007314 -62.1414297
N 5.0989407 -11.3521821 -60.3807085
H 4.9813281 -11.7997437 -59.4822739
C 4.011173 -10.5016823 -60.8819061
H 4.0372055 -10.488733 -61.9692609
C 4.1458654 -9.0360507 -60.4550618
O 4.0311999 -8.1487806 -61.2999311
C 2.6640571 -11.1114873 -60.4638078
H 1.8926477 -10.351034 -60.4842377
H 2.730934 -11.4831747 -59.4414438
C 2.2535011 -12.2657448 -61.3868724
H 1.4591474 -12.8339579 -60.902344
H 3.1069454 -12.932034 -61.5231857
S 1.6498686 -11.7327541 -63.0139356
C 1.6214996 -13.3280329 -63.8756168
H 1.2174771 -13.1904539 -64.8799789
H 0.9970689 -14.0358844 -63.3303468
H 2.6341481 -13.7267153 -63.9450518
N 4.4403578 -8.7747006 -59.1755103
H 4.5741277 -9.5745046 -58.565276
C 4.7721047 -7.4432695 -58.6461105
H 4.2329189 -6.6929209 -59.2287529
C 6.2628089 -7.1513591 -58.8255713
O 7.1132365 -8.0286716 -58.7190631
C 4.3124856 -7.3225627 -57.1776623
H 3.2218766 -7.3201566 -57.1538076
H 4.664482 -8.1940433 -56.6248092
C 4.8413101 -6.0586761 -56.4669405
H 5.9269408 -6.1201353 -56.3940316
H 4.5835999 -5.1738691 -57.0510368
C 4.2767258 -5.8952971 -55.0518431
H 4.2724669 -6.8599056 -54.5417714
H 3.2469937 -5.5465683 -55.1283597
C 5.0870934 -4.8828455 -54.2236432
H 4.4787623 -4.580427 -53.3658758
H 5.2958632 -3.991815 -54.8235343
N 6.3573003 -5.466491 -53.7188249
H 6.2395348 -6.3113772 -53.1868018
H 6.9070721 -4.8226841 -53.1439003
H 7.0461657 -5.6510471 -54.4521823
N 6.5831059 -5.8796803 -59.0166479
H 5.8235267 -5.2105063 -59.113986
C 7.9375657 -5.3227778 -59.0310328
H 8.6143732 -5.9856498 -58.4939189
C 7.9299697 -3.9753104 -58.3016906

O 6.8849011 -3.3299543 -58.2056804
C 8.3940536 -5.2114097 -60.498702
H 7.5989185 -4.7461053 -61.083704
C 9.6867695 -4.4393521 -60.7534294
H 10.461601 -4.7634189 -60.0624864
H 9.5092652 -3.3712309 -60.637248
H 10.0261984 -4.6212524 -61.7728531
O 8.6164189 -6.5095718 -60.9963761
H 8.1255677 -7.1366062 -60.4489059
N 9.0868513 -3.5097747 -57.8288001
H 9.9133854 -4.0836863 -57.9019445
C 9.2842826 -2.0970461 -57.484942
H 8.3344924 -1.5820776 -57.6283965
C 10.2397767 -1.4588863 -58.488381
O 11.1822076 -2.0878065 -58.9748673
C 9.6723067 -1.8988096 -55.9998239
H 9.4745394 -2.8358148 -55.4851521
C 11.1589555 -1.5434698 -55.8111287
H 11.3819425 -0.5661663 -56.2441685
H 11.7740098 -2.3086833 -56.2769605
H 11.4226248 -1.4985324 -54.7557414
C 8.7636256 -0.8305709 -55.3496209
H 8.9602225 0.1445593 -55.7991688
H 7.7226656 -1.0931424 -55.5430241
C 8.9349449 -0.7232145 -53.8274753
H 9.9006709 -0.2819557 -53.5826193
H 8.8562241 -1.7099889 -53.3708842
H 8.1532925 -0.0869728 -53.4164581
N 9.9517901 -0.2103244 -58.8094482
H 9.1682783 0.231535 -58.3373574
C 10.654951 0.6360141 -59.7608788
H 11.4393476 0.0802286 -60.2761536
C 11.3137825 1.7707113 -58.9834636
O 10.6744127 2.3280577 -58.0948429
C 9.5994166 1.0937026 -60.7929687
H 8.6089785 1.028997 -60.3467226
H 9.6158493 0.3827572 -61.6188321
C 9.6756245 2.4856417 -61.3678588
N 9.6560584 2.7683857 -62.7304314
C 9.6304442 4.1041051 -62.8320208
H 9.6677495 4.647484 -63.7669766
N 9.599696 4.6686446 -61.6145544
H 9.6754936 5.6576488 -61.4159251
C 9.6037052 3.6646771 -60.6808175
H 9.5692624 3.7963986 -59.6106405
N 12.5267095 2.1509732 -59.3842939

H 13.0058659 1.5797279 -60.0714761
C 13.1036285 3.4786336 -59.1419166
H 12.3791542 4.0801703 -58.6065819
C 13.4003518 4.137936 -60.4980748
O 13.9804095 3.4895145 -61.3751488
C 14.3486196 3.3947989 -58.2404563
H 14.0355429 3.0345998 -57.2602812
H 15.0443904 2.6671256 -58.661884
C 15.0998792 4.7290328 -58.0561606
H 15.5972644 4.9762276 -58.9978091
H 15.8861092 4.5829474 -57.3125827
C 14.2028056 5.9034645 -57.6227446
O 14.1451209 6.8963968 -58.3833069
O 13.5576779 5.8054743 -56.5593021
N 13.0453446 5.4126846 -60.6712948
H 12.6425479 5.9066621 -59.8759317
C 13.3632583 6.2062371 -61.8533931
H 14.1139036 5.6662568 -62.4167197
C 13.9496037 7.5757594 -61.5001384
O 13.3110616 8.3851175 -60.8299668
C 12.1357501 6.3519771 -62.752082
H 11.3121412 6.7960327 -62.1946084
H 11.8710934 5.3727228 -63.1411174
H 12.3807769 6.9994993 -63.5956822
N 15.109189 7.8745898 -62.0873356
H 15.4773862 7.2053572 -62.7563538
C 15.8432781 9.1316037 -61.9126432
H 15.2448618 9.8123746 -61.30726
C 16.0561624 9.7692783 -63.2824116
O 16.4188304 9.0839867 -64.2349318
C 17.1881392 8.8969502 -61.188147
H 17.8076121 8.2561194 -61.8169794
C 16.9764192 8.1765845 -59.8366834
H 16.3622953 8.7964619 -59.1838709
H 16.4432474 7.2414061 -60.0042166
C 17.9261826 10.2389898 -60.9959842
H 17.3004281 10.929227 -60.4298202
H 18.1645092 10.6855659 -61.9607414
H 18.8686102 10.0894265 -60.4738383
C 18.2749524 7.8099776 -59.110778
H 18.7708445 8.7035771 -58.7336403
H 18.9412565 7.2714474 -59.7855677
H 18.0370505 7.1684199 -58.2616609
N 15.8801458 11.0823937 -63.3805037
H 15.5331748 11.5872138 -62.5688742
C 15.9623305 11.817571 -64.6442913

H 16.7952455 11.4316431 -65.2351664
C 16.2306405 13.3158319 -64.4223685
O 16.001996 13.8407697 -63.3256233
C 14.6780471 11.5721577 -65.4405948
H 14.5614297 12.3592306 -66.1828294
H 14.8143697 10.6397859 -65.9771394
C 13.3981278 11.4233831 -64.5967226
H 13.2409175 12.3213791 -64.0054631
H 13.4843556 10.5777558 -63.912386
C 12.2010246 11.1445692 -65.486179
O 12.4220538 10.9836834 -66.7062945
O 11.0711677 11.0577901 -64.952999
N 16.7127389 14.0282412 -65.4533237
H 16.8577185 13.566903 -66.349382
C 16.9273486 15.4814323 -65.3898288
H 16.3871383 15.8476574 -64.5193038
C 16.3447654 16.2645858 -66.5817988
O 16.1558116 15.7419953 -67.6835179
C 18.4109936 15.8096423 -65.1287159
H 18.8261429 15.0930979 -64.4212154
H 18.4804463 16.7986531 -64.677849
C 19.2719235 15.817902 -66.3805842
O 19.9812442 14.8726451 -66.670495
N 19.2629312 16.8803734 -67.1552989
H 19.7115882 16.7659892 -68.0471681
H 18.6938739 17.6945154 -66.9492704
N 16.1171737 17.5560555 -66.3461034
H 16.3244087 17.9218972 -65.4216912
C 15.6409335 18.5213459 -67.3311437
H 15.1563779 17.9880668 -68.1515813
C 16.8207639 19.3019001 -67.9050657
O 17.814395 19.5196466 -67.2156087
C 14.6025547 19.470826 -66.7156745
H 15.0457816 19.9951369 -65.8663646
H 13.7532703 18.8943344 -66.3687152
C 14.1049765 20.4776596 -67.7549781
O 14.0497 21.6808959 -67.4357038
O 13.8939207 20.0301459 -68.9036003
N 16.6932574 19.7127438 -69.1587056
H 15.7361606 19.6535177 -69.5221682
C 17.6417525 20.4891035 -69.9664808
H 18.1709276 19.7718774 -70.5775385
C 18.7397635 21.2279062 -69.1911158
O 19.6163392 20.5303848 -68.6231238
O 18.7905354 22.4280729 -69.0989548
C 16.8603575 21.4153103 -70.9236658

H 16.3055296 22.1240006 -70.3041713
H 17.5800612 21.9809911 -71.516076
C 15.8643249 20.7213124 -71.8783228
H 15.1037594 20.1933291 -71.2956912
C 15.1458514 21.7861056 -72.7164382
H 15.8579213 22.3362843 -73.3312233
H 14.6219113 22.4788679 -72.0561909
H 14.4058411 21.3071409 -73.3609419
C 16.5435476 19.7199673 -72.8232848
H 17.329409 20.2094041 -73.398468
H 15.8002973 19.3095617 -73.5106352
H 16.9602655 18.8885803 -72.2557724

**Supplementary Data 3. Cartesian coordinates for TePeNMT homology model.**

N -2.1385667 21.7753992 -15.9844882
H -2.8881317 22.3451992 -15.612073
H -2.4187043 21.4412796 -16.898962
H -1.952445 20.9940613 -15.3721405
C -0.8983927 22.6008534 -16.1048954
H -1.0805973 23.3575681 -16.8709468
C 0.33849 21.7897622 -16.5659561
O 1.2553163 22.327476 -17.1874333
C -0.6053818 23.3116885 -14.7771474
H 0.3668547 23.80077 -14.8479421
H -0.5567763 22.5711725 -13.9778747
C -1.639947 24.3835703 -14.4160168
H -1.396285 24.7758699 -13.426708
H -2.6273748 23.9290305 -14.3587376
S -1.7285502 25.7740286 -15.5733039
C -2.9534829 26.7844932 -14.6928753
H -3.1342375 27.7045542 -15.2488035
H -2.5798462 27.0332981 -13.6984196
H -3.8891334 26.2323095 -14.5953778
N 0.3673335 20.4886197 -16.2657559
H -0.4109708 20.1080689 -15.7405706
C 1.3938537 19.5859753 -16.7771625
H 1.3580658 19.6103407 -17.8672226
H 2.3753087 19.9422459 -16.4680576
C 1.2496557 18.1385969 -16.3224611
O 0.7986647 17.8537127 -15.2154758
N 1.7628938 17.2390447 -17.1522057
H 2.1121366 17.5432004 -18.0501647
C 2.0288584 15.8382482 -16.8338954
H 1.9028599 15.7027836 -15.7604402
C 3.4957816 15.5076051 -17.1394012
O 4.119702 16.1812749 -17.9609391
C 0.9990926 14.9223526 -17.5032517
H 1.2572628 13.8928438 -17.2749979
H 0.0423459 15.1358366 -17.026009
C 0.8149397 15.0455348 -19.020348
H 1.6157817 14.5050359 -19.5288645
H 0.8699863 16.0954136 -19.3176114
C -0.5554566 14.4754363 -19.4180847
O -1.297864 15.2053148 -20.1119062
O -0.875195 13.3478418 -18.9747065
N 4.1061184 14.7016468 -16.2632362
H 3.5442127 14.1102083 -15.6715657
C 5.5629684 14.6122266 -16.0983634

H 6.0446756 14.8186955 -17.0600251
C 5.9695982 13.2129132 -15.6736854
O 5.3463653 12.639753 -14.786269
C 6.0594875 15.5900786 -15.0182406
H 5.5966651 15.3285324 -14.0641537
H 7.1389711 15.4636292 -14.915048
C 5.7688522 17.0627619 -15.3210483
H 6.199365 17.3109927 -16.2919241
H 4.6914859 17.2284646 -15.351451
C 6.3694499 17.9615862 -14.2310701
H 7.4516987 17.8220036 -14.192695
H 5.9514537 17.6743696 -13.2638997
C 6.0413128 19.4366002 -14.4861744
H 6.4275873 20.029751 -13.6516003
H 4.9522862 19.5499365 -14.5026758
N 6.6253332 19.9107507 -15.7656037
H 7.6331593 19.818804 -15.7546612
H 6.3843249 20.8792368 -15.933991
H 6.2683797 19.3550584 -16.5356419
N 7.106455 12.7593095 -16.1705568
H 7.4956956 13.2759729 -16.9574346
C 7.5639831 11.383838 -16.027934
H 6.7220861 10.7521143 -15.7410336
C 8.6256294 11.3080201 -14.9245199
O 9.6775993 11.9537076 -14.9983873
C 8.0743326 10.8706348 -17.3888252
H 8.9570019 11.4416081 -17.6791626
H 8.3787829 9.8302572 -17.2664366
C 7.0169986 10.9370419 -18.5172246
H 6.1405201 10.371064 -18.1914962
H 7.409044 10.4393792 -19.4038394
C 6.5974706 12.3719046 -18.896483
O 7.4785217 13.2681533 -18.9084406
O 5.3745041 12.6072441 -18.9936412
N 8.3366279 10.5700175 -13.8495991
H 7.428984 10.1185224 -13.7997984
C 9.2537472 10.4628666 -12.7165948
H 9.411715 11.4704627 -12.3299929
C 10.6243327 9.9036971 -13.153139
O 10.6986872 8.9385328 -13.9069108
C 8.6037136 9.6350831 -11.6036772
H 8.4204313 8.619799 -11.9600852
H 7.655759 10.0896643 -11.3114457
H 9.2647124 9.595986 -10.737355
N 11.703524 10.5367378 -12.6721267
H 11.5296608 11.3030954 -12.0418231

C 13.1142625 10.3142897 -13.0702562
H 13.6516325 11.146085 -12.6173586
C 13.4243279 10.5119185 -14.5675031
O 14.4681525 10.0599575 -15.0339048
C 13.7765064 9.0619071 -12.4395625
H 14.845682 9.122573 -12.6445486
C 13.6265139 9.0675566 -10.9118445
H 12.5803642 8.9423089 -10.6318339
H 14.0018322 10.0070241 -10.506304
H 14.2061243 8.2486758 -10.485005
C 13.2962742 7.7038875 -12.9652798
H 12.2890587 7.4911381 -12.6080123
H 13.9637118 6.9192992 -12.6112727
H 13.2950871 7.7094786 -14.0550769
N 12.605294 11.3349009 -15.2378421
H 11.7369915 11.5814565 -14.7805248
C 12.7314974 11.833814 -16.6111805
H 11.8724211 12.4921825 -16.7495073
C 12.6154246 10.7757957 -17.7237248
O 13.1625965 9.6728211 -17.644985
C 13.9716254 12.728998 -16.7542726
H 14.876663 12.1276992 -16.6566322
H 13.9650252 13.5017634 -15.9855672
H 13.9695782 13.2008787 -17.7373648
N 11.9102878 11.1457732 -18.7984009
H 11.4901366 12.0606529 -18.8311174
C 11.6113386 10.2161504 -19.8849453
H 11.4683386 9.2575284 -19.3922061
C 12.7697246 10.041093 -20.8749029
O 13.4841971 10.9883786 -21.2103005
C 10.2956217 10.5834444 -20.6018401
H 9.5642602 10.8737195 -19.8541547
H 10.4613373 11.4376453 -21.2598264
C 9.7172444 9.4047201 -21.4196476
H 10.350257 9.2350742 -22.2929309
H 8.7362979 9.7002982 -21.7986705
C 9.5899082 8.089336 -20.6183328
O 8.4815808 7.7830257 -20.1380566
O 10.6205886 7.3866881 -20.4498584
N 12.8901662 8.8305339 -21.4204041
H 12.2017952 8.1247666 -21.1470783
C 13.7998552 8.4972353 -22.5092701
H 14.1771403 9.4236569 -22.9434863
C 13.0399482 7.741936 -23.6088019
O 12.2666108 6.8259826 -23.3460136
C 14.9936922 7.7225667 -21.9245386

H 15.5173435 8.3745069 -21.2235556
H 14.6001376 6.8863099 -21.3570609
C 16.0029798 7.189281 -22.9591753
H 15.4878043 6.568367 -23.6890142
C 16.7306364 8.3210255 -23.6896215
H 17.2504768 8.9574053 -22.9726436
H 16.0235264 8.9250527 -24.2541046
H 17.4549132 7.90003 -24.3877077
C 17.0509741 6.3236455 -22.2584349
H 17.5894864 6.9110609 -21.5149428
H 17.7563171 5.9395483 -22.9960493
H 16.5630007 5.4766885 -21.778768
N 13.2962296 8.1160457 -24.8611726
H 13.9627613 8.8548464 -25.0049569
C 12.7297022 7.4720004 -26.0467334
H 11.6552424 7.6561114 -26.0668372
C 12.963355 5.9483119 -26.0268958
O 14.0825531 5.4964434 -25.7932219
C 13.3669937 8.1475173 -27.2671617
H 13.1403425 9.2139322 -27.2266732
H 14.4513269 8.049724 -27.187207
C 12.9656811 7.6189319 -28.6307211
C 12.1494424 8.3971049 -29.4761888
H 11.769383 9.3526089 -29.1454154
C 11.8565891 7.9520163 -30.7798108
H 11.2562024 8.555452 -31.4472005
C 12.3673248 6.7178848 -31.2374061
O 12.1160781 6.2857259 -32.4969706
H 11.5991475 6.9488647 -33.0185918
C 13.1697677 5.9340538 -30.3847744
H 13.5492799 4.9896701 -30.7459872
C 13.4827043 6.3907892 -29.091865
H 14.1273567 5.791667 -28.4621886
N 11.8951892 5.1788313 -26.2599184
H 11.0304658 5.6427895 -26.4800882
C 11.8426092 3.7036502 -26.2137578
H 10.7811114 3.4599108 -26.2631227
C 12.3180866 3.0493723 -24.893297
O 12.588105 1.8486882 -24.8229072
C 12.4769853 3.1028593 -27.4818833
H 13.5610504 3.0598399 -27.3602348
H 12.2582179 3.7523311 -28.3311342
C 11.9188234 1.7098089 -27.8129528
O 10.6734795 1.5608478 -27.7641107
O 12.7303627 0.8270589 -28.169355
N 12.3786583 3.8077385 -23.7892188

H 12.1567416 4.7960764 -23.8565606
C 12.7236866 3.2518266 -22.4780661
H 13.5786598 2.5902866 -22.6181944
C 11.5733328 2.4149249 -21.9209309
O 10.5345585 2.9233827 -21.5059758
C 13.1586237 4.3717878 -21.5322934
H 14.0067916 4.8753226 -21.9927235
H 12.3398627 5.0838206 -21.4074609
C 13.5708913 3.8411374 -20.1519667
H 12.6903613 3.4453154 -19.6452956
H 14.3110593 3.0473288 -20.2626016
C 14.1595742 4.9730826 -19.3074994
H 13.477531 5.8273082 -19.3386531
H 15.1255757 5.265841 -19.7198963
C 14.3246592 4.5251755 -17.8558798
H 14.9647772 3.6393135 -17.8237746
H 13.3343658 4.2459127 -17.4769636
N 14.894404 5.6206571 -17.034387
H 15.8026652 5.9022239 -17.3743342
H 14.9585333 5.3479973 -16.0632593
H 14.281514 6.4323719 -17.0835563
N 11.8145313 1.1129027 -21.796752
H 12.636759 0.7428888 -22.2469702
C 10.915173 0.206206 -21.075752
H 9.9261211 0.3116122 -21.5233934
C 10.7879049 0.6022152 -19.5986625
O 11.7565775 1.0128309 -18.9601126
C 11.3345322 -1.270057 -21.22834
H 10.6296722 -1.8783407 -20.6606041
C 11.2418011 -1.7098514 -22.6941182
H 11.9608367 -1.1650701 -23.3087444
H 10.2395993 -1.5174528 -23.0780593
H 11.4468521 -2.7773516 -22.7753339
C 12.7458793 -1.5768076 -20.7095885
H 13.4907476 -1.0077613 -21.2656222
H 12.9570051 -2.6395233 -20.8284199
H 12.8214824 -1.3296274 -19.6506744
N 9.6090573 0.3641778 -19.0190877
H 8.8636369 0.042936 -19.61778
C 9.3030641 0.4995601 -17.5765515
H 8.2498576 0.2378134 -17.4888618
C 9.3464409 1.9102799 -16.9589934
O 9.2888891 2.018324 -15.7386553
C 10.0341763 -0.545724 -16.7016046
H 9.6693134 -0.4676256 -15.6770594
C 9.7794235 -1.9797159 -17.169852

H 10.215899 -2.1450359 -18.1540049
H 8.7079161 -2.1740235 -17.2032576
H 10.239578 -2.6699839 -16.4624216
O 11.4299275 -0.3613229 -16.691399
H 11.6667592 0.1678767 -17.4727227
N 9.2779939 2.9909397 -17.7450031
H 9.3809 2.8980378 -18.7455964
C 8.8924366 4.3186524 -17.2226461
H 9.484174 4.5459203 -16.3361664
C 7.3950813 4.3363773 -16.8420819
O 6.54833 4.1563548 -17.7157283
C 9.1737252 5.4060425 -18.2724269
H 8.8412563 6.3707971 -17.8859208
H 8.6096819 5.1895781 -19.1816057
O 10.5517975 5.4957348 -18.5958262
H 10.5894134 6.1754387 -19.3178923
N 7.0367162 4.4791013 -15.5515566
H 7.7637049 4.4744686 -14.8485576
C 5.6335974 4.372326 -15.0850094
H 5.0070584 4.6318068 -15.9400818
C 5.0839267 5.3356751 -13.9496445
O 4.1802507 4.8921545 -13.2441249
C 5.3501324 2.8742188 -14.7954481
H 5.6601937 2.2734319 -15.6495461
H 4.2770818 2.7385852 -14.6680957
C 6.0481724 2.3356799 -13.5563096
O 6.9355157 2.9397106 -12.9802825
N 5.6839717 1.1536912 -13.1130988
H 6.1721809 0.8278604 -12.2984982
H 4.9434468 0.6468731 -13.5561149
N 5.4209234 6.6581869 -13.7954432
H 6.1313111 7.0377768 -14.4037847
C 4.6727308 7.6690857 -12.8730958
H 3.6722803 7.2716519 -12.7031805
H 5.1665915 7.6425688 -11.902207
C 4.4605241 9.2667223 -13.2399755
O 5.4811072 9.6663177 -13.812121
N 3.2536409 10.1109433 -12.9872607
H 2.6474233 9.6614132 -12.3206553
C 2.3771526 11.4343133 -13.6922496
H 3.1658733 12.0715482 -14.0792353
C 1.0505375 12.8198719 -12.978889
O 0.738086 12.1140384 -12.0071596
C 1.9540762 10.4352165 -14.9544058
H 0.9329988 10.1097298 -14.746017
C 2.7417697 9.1177858 -15.5124699

H 3.7821969 9.3932306 -15.6797871
H 2.6711718 8.3582525 -14.7352443
C 1.9181844 11.302737 -16.1856385
H 2.9290002 11.590086 -16.4899791
H 1.3394983 12.1952761 -15.9637221
H 1.4171187 10.8031861 -17.0150758
C 2.3461044 8.3062146 -16.7841532
H 1.2838958 8.043668 -16.7411571
H 2.9411495 7.3919643 -16.8207085
H 2.5328117 8.8854332 -17.695951
N 0.2333932 14.4003043 -13.1834479
H 0.7437184 14.7653437 -13.9704046
C -1.2123799 15.955379 -12.9597387
H -1.0028175 16.3292374 -13.960482
C -2.5312399 15.1695335 -13.1010222
H -2.3258053 14.1092915 -13.2484796
H -3.0618878 15.2689053 -12.1555772
C -3.4175086 15.6977339 -14.2585728
H -3.2432872 16.7657087 -14.3985759
C -3.0783778 14.9714564 -15.5675978
H -3.2811317 13.903087 -15.4809576
H -2.0269104 15.1037593 -15.8111044
H -3.6757972 15.3801559 -16.3812018
C -4.9110054 15.512951 -13.9555661
H -5.1400087 14.4586235 -13.8201664
H -5.5040244 15.9052164 -14.7807417
H -5.172629 16.0607214 -13.0540348
C -2.0596333 17.9254177 -12.3430272
O -1.3832877 18.2475852 -11.374691
N -3.271554 19.2249776 -12.5340521
H -3.8017849 18.9469697 -13.3433045
C -4.1853409 20.9390259 -11.9780666
H -4.1201797 20.8378689 -10.8941409
C -5.8142116 21.9798425 -12.0685304
O -6.4577917 21.990306 -13.1238488
C -3.3553264 22.2522131 -12.3368714
H -2.7508435 21.9810248 -13.199742
H -3.9431639 23.1195955 -12.6376299
C -2.3935577 22.6639575 -11.2141059
H -1.5875927 23.2551301 -11.6539897
H -1.9417954 21.7737149 -10.7710014
C -3.1027948 23.4921229 -10.1365626
O -4.1338848 23.0043489 -9.6189633
O -2.6283562 24.6174618 -9.8712537
N -6.4475046 23.0719258 -11.2698861
H -5.8681384 23.316696 -10.4654281

C -7.9445496 23.8116708 -11.2481781
H -8.5313975 23.1447577 -11.8786809
C -8.4505552 25.3183528 -11.7282815
O -7.9562339 26.3541144 -11.2608184
C -8.5827225 23.7319299 -9.8142739
H -8.3098143 22.7644516 -9.3910452
H -8.1237068 24.4929663 -9.1820639
C -10.1128287 23.8553926 -9.699581
H -10.4295997 24.8665053 -9.9608487
H -10.3884539 23.7049408 -8.6526486
C -10.8425833 22.8299823 -10.5754585
O -10.9583405 23.1023724 -11.793048
O -11.2574013 21.7859035 -10.0265513
N -9.602907 25.5151302 -12.4609812
H -10.1970189 24.6947314 -12.6038089
C -10.0093518 26.7968804 -13.139427
H -9.6033406 27.6153906 -12.5465371
C -11.5373224 27.0770783 -13.3129338
O -12.3430523 26.1660776 -13.4651874
C -9.3158021 26.7963794 -14.5218177
H -8.2652178 26.5407625 -14.3723088
H -9.7616841 25.9988297 -15.1187812
C -9.3656862 28.1006173 -15.3396433
H -10.3980501 28.3527717 -15.5746801
C -8.7151959 29.2767986 -14.6053596
H -7.6855865 29.0272619 -14.3458923
H -9.2616884 29.5103425 -13.6946444
H -8.721151 30.1580805 -15.2468748
C -8.6202634 27.902131 -16.6598573
H -7.5714266 27.6723762 -16.4704162
H -8.6904418 28.808201 -17.2612677
H -9.0736317 27.0808258 -17.214682
N -11.9412789 28.3572898 -13.4297045
H -11.2467902 29.0828588 -13.3712445
C -13.3321225 28.7589476 -13.708797
H -13.9922764 28.0787705 -13.1669539
C -13.6973559 28.668285 -15.20173
O -13.2212429 29.4599915 -16.0166546
C -13.5976231 30.1744778 -13.1694646
H -13.3339687 30.2032292 -12.1112647
H -12.9526566 30.8857408 -13.6884859
C -15.0439728 30.6091374 -13.3172145
C -16.0007793 30.2075446 -12.3654889
H -15.7026398 29.6095012 -11.5164327
C -17.3497155 30.5734271 -12.5229045
H -18.084102 30.2600018 -11.7942258

C -17.7459906 31.3401123 -13.632773
H -18.784204 31.6156339 -13.7569758
C -16.7930155 31.742311 -14.5848431
H -17.0984245 32.326439 -15.4416514
C -15.4440982 31.3771052 -14.4276294
H -14.7165289 31.6734974 -15.1713742
N -14.6326318 27.780819 -15.5396291
H -14.9521194 27.1150618 -14.8471413
C -15.2390964 27.6886577 -16.8666391
H -15.735075 28.6312197 -17.0994544
H -14.4624243 27.5134405 -17.6111349
C -16.2666536 26.5596064 -16.957317
O -16.3953891 25.7524785 -16.0401148
N -16.9764725 26.4878918 -18.0819378
H -16.8289854 27.176356 -18.8044725
C -17.9575033 25.4408391 -18.3831007
H -17.7899232 24.5827215 -17.7304426
C -17.7792012 24.9927546 -19.8397808
O -17.7959146 25.8188285 -20.750926
C -19.3742105 25.9740925 -18.1133224
H -19.4072767 26.3626889 -17.0943744
H -19.5911057 26.7896238 -18.8053619
C -20.4418095 24.8784784 -18.2588574
H -20.2170614 24.0779836 -17.5490798
H -20.3816661 24.4589508 -19.2655484
C -21.873747 25.3850101 -18.0153021
O -22.0329252 26.4946208 -17.4568844
O -22.8044396 24.6367793 -18.3888393
N -17.5576422 23.6953514 -20.0649298
H -17.6282623 23.0479037 -19.293315
C -17.3269048 23.1256682 -21.396853
H -17.6295227 23.8524687 -22.1539443
C -18.173276 21.8705255 -21.6133325
O -18.4492201 21.1344351 -20.6665456
C -15.8303326 22.833808 -21.5971231
H -15.5031065 22.1074362 -20.8521036
H -15.2664698 23.755845 -21.4523805
C -15.5235885 22.3038595 -22.9769314
N -16.0573017 22.8083599 -24.1656877
C -15.6125519 22.0028993 -25.140073
H -15.854249 22.1181749 -26.1888167
N -14.8449621 21.02673 -24.6277346
H -14.429788 20.2755673 -25.1579691
C -14.7816493 21.1953241 -23.2621255
H -14.2791179 20.561661 -22.5447457
N -18.5185024 21.5818415 -22.8701808

H -18.1432063 22.1738025 -23.6009505
C -19.2288161 20.3661674 -23.257331
H -19.9116933 20.1067755 -22.4457962
C -18.2431347 19.2015468 -23.4437471
O -17.5034608 19.1367934 -24.4310812
C -20.0753688 20.6536823 -24.5101802
H -20.7622502 21.4721288 -24.2876809
H -19.4111275 20.9769363 -25.3134978
C -20.8899068 19.4416545 -25.0031907
H -20.2061397 18.632868 -25.252551
C -21.8900833 18.9305006 -23.9653095
H -22.5664824 19.7301243 -23.6638583
H -21.3716057 18.543429 -23.0898099
H -22.4721546 18.1090986 -24.387707
C -21.6558409 19.8178908 -26.2704486
H -22.3670983 20.6156827 -26.0549008
H -22.1966695 18.9460329 -26.6408658
H -20.9593117 20.149503 -27.0399621
N -18.305586 18.2264973 -22.5363312
H -18.9848484 18.3084584 -21.7879638
C -17.4987295 17.0022879 -22.56464
H -16.4445225 17.2767669 -22.5352472
C -17.7136686 16.1851165 -23.8479637
O -16.7324037 15.7061853 -24.4089253
C -17.8183622 16.1676056 -21.3118248
H -17.6392762 16.7679502 -20.4189096
H -17.1541192 15.3031549 -21.279748
C -19.2469521 15.6902349 -21.2993199
N -20.3555822 16.5291505 -21.1770985
C -21.409785 15.8002387 -21.5780685
H -22.404937 16.1941265 -21.7444035
N -21.0189659 14.5690479 -21.940586
H -21.5461555 13.999855 -22.611493
C -19.6585309 14.4792161 -21.7690455
H -19.0246639 13.6887216 -22.1457658
N -18.9342631 16.1416122 -24.3967885
H -19.7139306 16.4659898 -23.8383305
C -19.2306073 15.4622993 -25.6698397
H -18.9191364 14.4191048 -25.5943322
C -18.4740459 16.0915514 -26.8531044
O -17.994577 15.3816032 -27.732199
C -20.7422481 15.4827005 -25.9605596
H -20.9140981 15.0341016 -26.9402223
H -21.0845436 16.5175317 -26.0055241
C -21.5715667 14.7083567 -24.9337949
O -21.1731487 13.5796127 -24.57134

O -22.5721805 15.2767049 -24.4473625
N -18.2496141 17.4112082 -26.8323886
H -18.5716361 17.9329869 -26.0288096
C -17.4531297 18.1047217 -27.8489587
H -17.7977212 17.8135617 -28.8424323
H -17.5879439 19.1789559 -27.7304308
C -15.956921 17.7942072 -27.7454374
O -15.2888172 17.6280449 -28.7627028
N -15.439025 17.6201509 -26.5247796
H -16.0609943 17.7043393 -25.7294654
C -14.0743246 17.1266898 -26.3031017
H -13.3826248 17.7038566 -26.9186169
C -13.9256189 15.6534948 -26.724003
O -12.9564171 15.2990751 -27.3953963
C -13.702554 17.3422062 -24.8291262
H -13.7609572 18.4063683 -24.6038623
H -14.431762 16.8391893 -24.1953113
C -12.3186415 16.8515113 -24.4543145
C -11.189853 17.6482056 -24.7279804
H -11.3040053 18.609059 -25.2098121
C -9.9051194 17.1911998 -24.3777829
H -9.0312386 17.792005 -24.5808524
C -9.7460546 15.9368061 -23.7535044
O -8.5019852 15.5094553 -23.4137942
H -8.5099345 14.6569228 -22.9779248
C -10.8775819 15.1370117 -23.4808996
H -10.7593813 14.1747477 -23.0085044
C -12.161335 15.5959257 -23.8337612
H -13.0275541 14.9793858 -23.6353157
N -14.9268359 14.8201123 -26.423841
H -15.7014839 15.176822 -25.8741997
C -14.9828394 13.4197593 -26.8408227
H -14.0769896 12.9181517 -26.4986893
C -15.0310294 13.2711907 -28.3685764
O -14.316041 12.4302505 -28.9112801
C -16.1878629 12.7409781 -26.1724631
H -16.0987507 12.8425083 -25.0898935
H -17.1015123 13.2506904 -26.4752072
C -16.3194695 11.2716038 -26.5208461
C -15.679925 10.2960364 -25.7306851
H -15.1296517 10.5903453 -24.8487413
C -15.7427155 8.9385754 -26.1018896
H -15.2470437 8.1817352 -25.5143806
C -16.4327316 8.5582085 -27.2723703
O -16.4601685 7.2539103 -27.6505408
H -16.8895214 7.1591939 -28.5014005

C -17.087672 9.5346431 -28.0532165
H -17.603424 9.2540987 -28.9595907
C -17.035941 10.887789 -27.6716777
H -17.5222426 11.6367727 -28.2851494
N -15.7781968 14.1209507 -29.0785084
H -16.4213335 14.735133 -28.58639
C -15.8302485 14.0991621 -30.5458371
H -16.023302 13.0723458 -30.857743
C -14.4935853 14.5210283 -31.1785911
O -13.967663 13.81329 -32.0333803
C -16.994066 14.9750367 -31.0424011
H -17.90762 14.6865268 -30.519727
H -16.7819786 16.0183928 -30.8031416
C -17.238316 14.8508303 -32.5555673
H -16.3290961 15.1177248 -33.0992785
H -18.0038016 15.5779021 -32.8375267
C -17.6986865 13.4395358 -32.9540066
O -18.9264795 13.2539991 -33.097649
O -16.8225467 12.5548429 -33.0888592
N -13.837982 15.5729289 -30.6701081
H -14.2892367 16.1201696 -29.9440593
C -12.4898729 15.9648231 -31.1351682
H -12.5346458 16.1337827 -32.2113024
C -11.4694612 14.8360737 -30.9194148
O -10.6875397 14.5295424 -31.8238719
C -12.0510367 17.2817301 -30.4618222
H -12.2135606 17.2075124 -29.385962
C -10.5745282 17.6261432 -30.7001145
H -10.360449 17.6415137 -31.7693089
H -9.9326989 16.8899406 -30.2164525
H -10.3471552 18.6032195 -30.2737645
C -12.8768666 18.454303 -31.009272
H -12.6844 18.5843449 -32.0745506
H -12.6143381 19.3701809 -30.4802161
H -13.9403605 18.2682826 -30.8668087
N -11.5418689 14.1369177 -29.7810758
H -12.2068198 14.4377171 -29.0741423
C -10.7457172 12.934077 -29.5229422
H -9.6880299 13.1653715 -29.6509979
H -10.9128238 12.6231638 -28.4919327
C -11.1015855 11.7534031 -30.4368387
O -10.2052046 11.0525872 -30.9091058
N -12.3806552 11.5703557 -30.7665667
H -13.0669852 12.1981408 -30.3596135
C -12.8830576 10.4904933 -31.6335773
H -12.4536585 9.5465298 -31.299618

C -12.4546029 10.6930136 -33.0856704
O -11.8379689 9.7967565 -33.662114
C -14.4115139 10.3664975 -31.5108397
H -14.8837716 11.3195435 -31.7430037
C -15.0075931 9.281508 -32.405127
H -14.497612 8.3332624 -32.2385487
H -14.9184434 9.5764776 -33.4497101
H -16.0681325 9.1738969 -32.1753859
O -14.7191285 9.9990169 -30.1858573
H -14.6611214 10.8041515 -29.6445041
N -12.6061399 11.9050074 -33.6278135
H -13.1277568 12.5991368 -33.0943308
C -12.1108057 12.2899864 -34.96069
H -12.5626445 11.6330904 -35.7039491
C -10.5924574 12.1086371 -35.0606404
O -10.1118234 11.5017037 -36.0194828
C -12.5253388 13.7419652 -35.2846723
H -12.2047179 14.3950538 -34.4716908
C -11.9014053 14.2583629 -36.5896139
H -12.1485647 13.587198 -37.4129321
H -10.8182307 14.3273054 -36.4922998
H -12.2846337 15.2548982 -36.8119629
C -14.0473735 13.8569901 -35.4397281
H -14.390893 13.2481522 -36.2760117
H -14.3288191 14.8958381 -35.6094773
H -14.5551193 13.5168148 -34.5364626
N -9.8327613 12.5183483 -34.0376294
H -10.2691124 13.0158676 -33.2677802
C -8.3866883 12.2947826 -33.9964652
H -7.94803 12.713808 -34.9032965
C -8.0253687 10.7951613 -33.964585
O -7.1750255 10.3535353 -34.7385592
C -7.808095 13.0498299 -32.7940855
H -8.2322251 12.6640796 -31.8665257
H -8.0425363 14.1119177 -32.8768195
H -6.7249933 12.9258191 -32.7693519
N -8.7383063 9.9928143 -33.1690346
H -9.4399586 10.4216383 -32.5757625
C -8.5261026 8.5373236 -33.0519002
H -7.476906 8.3564513 -32.8190178
C -8.836892 7.7959867 -34.3557814
O -8.0721417 6.9214099 -34.7582541
C -9.3671494 7.960609 -31.9009351
H -10.4270834 8.1175921 -32.1029293
C -9.1210937 6.4714297 -31.6602582
H -8.0569718 6.2924769 -31.5037895

H -9.4663804 5.8883818 -32.5134126
H -9.6722286 6.1524028 -30.7752524
O -9.0287859 8.6085484 -30.6970051
H -9.4300372 9.4946216 -30.711502
N -9.8765834 8.2071453 -35.0894348
H -10.4751493 8.9342993 -34.7066042
C -10.218649 7.6484065 -36.4087586
H -10.3078933 6.565189 -36.3149444
C -9.0961985 7.9031619 -37.4291875
O -8.8107229 7.0319327 -38.2457573
C -11.5883493 8.2058925 -36.8724934
H -11.5800876 9.2895272 -36.7418409
C -11.8649231 7.9080697 -38.3605028
H -11.8182208 6.8336763 -38.5433383
H -11.1330206 8.4106419 -38.9928817
H -12.8465962 8.2794081 -38.6492608
C -12.7194695 7.6055139 -36.002287
H -12.8725628 6.5606242 -36.275951
H -12.4291906 7.6247014 -34.9527788
C -14.0547499 8.3529914 -36.1158443
H -14.469557 8.2558855 -37.1180749
H -13.9111854 9.4081356 -35.8800284
H -14.765007 7.9268607 -35.4065953
N -8.3937191 9.0388911 -37.3367375
H -8.6846259 9.7176446 -36.6427142
C -7.265239 9.3812153 -38.2229573
H -7.5533014 9.1179589 -39.2408252
C -5.9497281 8.6286735 -37.9283472
O -4.9741865 8.7465544 -38.6840646
C -7.0390962 10.899473 -38.1899562
H -8.0014138 11.4136505 -38.2093944
H -6.4800146 11.1924024 -39.0771632
O -6.2990547 11.3175345 -37.05805
H -6.6658562 10.914714 -36.2473093
N -5.8900761 7.8921141 -36.8100548
H -6.7249579 7.8243468 -36.2397125
C -4.6565738 7.3302429 -36.256341
H -3.8362244 8.0007177 -36.5123651
C -4.2800623 5.9408626 -36.8074929
O -3.1352983 5.5205919 -36.6491747
C -4.7709993 7.3240925 -34.7269169
H -5.5236596 6.6026775 -34.4107831
H -5.0474193 8.3167635 -34.3677705
H -3.8122191 7.0434425 -34.2890576
N -5.1800992 5.2659515 -37.5310566
H -6.0659794 5.7270073 -37.7010793

C -4.9863126 3.9392569 -38.1512939
H -5.0038412 3.1815968 -37.3671395
C -3.6467069 3.7571432 -38.8886477
O -2.9731921 2.7386014 -38.7264792
C -6.1697208 3.6740353 -39.1048049
H -7.0402153 3.4328757 -38.4935022
H -5.9499646 2.7970377 -39.7150907
C -6.5637775 4.8197352 -40.0201111
N -5.7699764 5.9207787 -40.3641479
C -6.5674691 6.752385 -41.049963
H -6.277414 7.7318812 -41.4028678
N -7.7926363 6.2244398 -41.1826337
H -8.5854946 6.6964794 -41.5940588
C -7.8129371 5.0107498 -40.5332027
H -8.6720512 4.3747877 -40.3736965
N -3.1845526 4.7824287 -39.6094995
H -3.8397485 5.5490973 -39.7402001
C -1.9094941 4.7790629 -40.3503428
H -1.9058746 3.9210138 -41.0259016
C -0.6755178 4.6000095 -39.4590681
O 0.3500762 4.1523355 -39.9634635
C -1.8108003 6.0655976 -41.1818839
H -1.9802055 6.9149351 -40.5182176
H -0.8101594 6.155225 -41.6088882
C -2.8366963 6.0737128 -42.3295833
H -3.7805566 5.6467642 -41.9998432
H -2.4675716 5.4691415 -43.158521
C -3.090666 7.5062857 -42.7975055
H -2.21557 7.8834898 -43.328123
H -3.2604295 8.1357064 -41.9227019
N -4.2898586 7.6146451 -43.6353146
H -5.0985667 8.0752973 -43.2163844
C -4.3632322 7.5927761 -44.9465972
N -3.4118636 7.0991039 -45.680707
H -3.2172373 7.5904414 -46.5573116
H -2.6560279 6.644648 -45.2205878
N -5.4028833 8.1065841 -45.5234574
H -6.0523492 8.6180683 -44.9202338
H -5.4423454 8.1830156 -46.5185052
N -0.7546542 4.8641941 -38.1528525
H -1.6478001 5.1514024 -37.7665882
C 0.3441051 4.6164829 -37.2169392
H 1.2323201 5.1264964 -37.5885598
C 0.6996833 3.1210416 -37.0906775
O 1.8637562 2.7984179 -36.8587498
C -0.0133202 5.2297426 -35.858726

H -0.9017486 4.748926 -35.4486878
H -0.2065543 6.2971976 -35.9704315
H 0.8169899 5.0909097 -35.1647239
N -0.2423564 2.2117153 -37.3765593
H -1.1831707 2.5200405 -37.5942453
C 0.0441412 0.7786215 -37.4595675
H 0.5673747 0.4724402 -36.5515695
C 0.9552045 0.4341165 -38.6550826
O 1.7906982 -0.4590169 -38.5465271
C -1.2872208 0.0221727 -37.522223
H -1.8167315 0.2672544 -38.4447155
H -1.9079066 0.2944651 -36.668599
H -1.1024213 -1.0504495 -37.4916568
N 0.8781965 1.1977947 -39.7544587
H 0.2102942 1.9555281 -39.7611379
C 1.8112123 1.0873253 -40.8905783
H 2.0039892 0.0301488 -41.0804667
C 3.1532747 1.7347408 -40.5479506
O 4.1918602 1.1211358 -40.7786454
C 1.2242149 1.7004207 -42.1796402
H 1.0712918 2.7691655 -42.0362597
C 2.1680245 1.5163423 -43.3761676
H 2.3959256 0.458749 -43.5151264
H 3.0968456 2.0616945 -43.2111974
H 1.7047064 1.9050683 -44.2829471
C -0.1270984 1.0684292 -42.5415657
H -0.0134202 -0.0094525 -42.6618157
H -0.4995357 1.4948188 -43.4727939
H -0.8607535 1.2658962 -41.7604837
N 3.1497115 2.9075864 -39.8983128
H 2.2584563 3.3646896 -39.7477886
C 4.3829764 3.5769137 -39.4271644
H 5.0316617 3.7532182 -40.287047
C 5.1714679 2.6870662 -38.4572727
O 6.3914124 2.6466122 -38.5342007
C 4.0854947 4.9447377 -38.7734971
H 3.4492565 4.7868126 -37.9038288
C 5.3579813 5.667574 -38.3077099
H 6.0473034 5.7700222 -39.1479344
H 5.8520541 5.0988091 -37.5198768
H 5.1091859 6.6515776 -37.9116408
C 3.3657123 5.9094878 -39.7292705
H 4.0142422 6.146524 -40.5715909
H 3.1117469 6.8290265 -39.2030367
H 2.4524687 5.4645623 -40.1099666
N 4.5099699 1.8691594 -37.6261136

H 3.501018 1.9661057 -37.5679173
C 5.1768327 0.8861925 -36.7517641
H 5.9189905 1.4157616 -36.1524491
C 5.9655105 -0.1805465 -37.5278367
O 6.9655235 -0.6580613 -37.005066
C 4.1151826 0.2654933 -35.821984
H 3.6872668 1.0584642 -35.2070059
H 3.3115988 -0.1409395 -36.4366679
C 4.6026419 -0.8663175 -34.8987054
H 4.9701494 -1.6946528 -35.5041773
H 3.7425868 -1.2253198 -34.3369632
C 5.691348 -0.4487287 -33.8979481
H 5.305648 0.3329929 -33.2411631
H 6.5347315 -0.0299299 -34.4515579
N 6.1852786 -1.5988657 -33.1065998
H 7.0302672 -2.023504 -33.4565587
C 5.6931212 -2.0929878 -31.9796119
N 4.6137617 -1.6238221 -31.4121932
H 4.2496792 -2.0315972 -30.5644291
H 4.1500759 -0.8388203 -31.8268032
N 6.2826626 -3.0960025 -31.392016
H 7.158683 -3.4460252 -31.7437805
H 5.9220268 -3.4384356 -30.5210906
N 5.5130667 -0.5786026 -38.7196586
H 4.6989518 -0.1098716 -39.0946508
C 6.155844 -1.6215128 -39.5477068
H 6.7171868 -2.2989085 -38.9044692
C 7.1870424 -1.0471198 -40.5141252
O 8.1672071 -1.7115576 -40.8379074
C 5.0729423 -2.429844 -40.2970744
H 4.4719049 -1.7392024 -40.8919679
C 5.6505611 -3.4778044 -41.2668926
H 6.3652394 -4.1194898 -40.7503349
H 6.1577817 -2.9777053 -42.0922651
H 4.8537515 -4.0898952 -41.6888069
C 4.1272498 -3.1381766 -39.3001663
H 3.4521743 -2.3981917 -38.8745679
H 3.519856 -3.8567523 -39.8472324
C 4.8022508 -3.8790266 -38.1377464
H 5.1620025 -3.1674568 -37.3956499
H 5.6444629 -4.4611102 -38.5038849
H 4.0797422 -4.5405924 -37.6627072
N 6.9634562 0.1827065 -40.9660073
H 6.1059518 0.645179 -40.6903566
C 7.8918005 0.9019409 -41.8307362
H 8.3560989 0.1968688 -42.5199967

C 9.0541694 1.5083209 -41.0392823
O 10.1751688 1.4175308 -41.5113396
C 7.0910835 1.9372281 -42.6425295
H 6.4755174 2.5141194 -41.9494156
C 8.015055 2.9405889 -43.3353853
H 8.8170607 2.4130698 -43.8517664
H 8.4436482 3.6079035 -42.5859543
H 7.4537262 3.5413332 -44.0376991
C 6.1329575 1.2547909 -43.6511673
H 5.3388 0.7519175 -43.1009127
H 5.6559528 2.0240523 -44.2518133
C 6.7607953 0.2206656 -44.5975052
H 7.0017951 -0.6915304 -44.0477814
H 7.6692296 0.6123557 -45.0487245
H 6.0481605 -0.0291608 -45.3837918
N 8.798992 1.9819869 -39.8195457
H 7.8269675 1.9509433 -39.5448266
C 9.6149082 2.8267209 -38.9312898
H 9.2170332 2.5797365 -37.9459199
C 9.3250434 4.3409519 -39.0306445
O 8.5238253 4.7993794 -39.8542185
C 11.099113 2.4020543 -38.8279642
H 11.7082873 3.1408157 -39.3434734
H 11.2510327 1.4269978 -39.2904088
C 11.5965131 2.3038259 -37.3833269
O 11.4492724 3.3205405 -36.6703128
O 12.1249463 1.2330349 -37.0094698
N 9.8588958 5.1058401 -38.0754761
H 10.516748 4.6466534 -37.4396315
C 9.3325992 6.4041998 -37.6512809
H 8.3526531 6.2042132 -37.2199303
C 9.1395831 7.4234129 -38.7896545
O 10.0494829 7.7492656 -39.5515223
C 10.2024572 6.9760756 -36.5200309
H 10.3373002 6.2059069 -35.7593743
H 11.1814432 7.2466722 -36.9189787
C 9.5601829 8.2096529 -35.8657287
H 8.5933201 7.9247972 -35.443284
H 9.3792816 8.9717041 -36.6261681
C 10.4478246 8.8008571 -34.7647909
O 10.8020372 8.0456266 -33.8314376
O 10.7428137 10.012847 -34.8410345
N 7.9356494 8.0039627 -38.8202964
H 7.2747294 7.6949909 -38.1276181
C 7.4979327 9.066309 -39.7246913
H 6.4475177 9.2219085 -39.4784392

C 7.4607978 8.7250306 -41.2328113
O 7.2182063 9.6195186 -42.0501456
C 8.189382 10.3821855 -39.339387
H 9.2533861 10.3291555 -39.5703739
H 8.0772996 10.5585665 -38.2684137
H 7.7299772 11.2135753 -39.8729389
N 7.5456185 7.4423073 -41.6155564
H 7.7541 6.7354562 -40.9135248
C 7.344726 6.9801426 -42.997353
H 6.9802142 7.8289962 -43.5714683
C 6.2228111 5.9398072 -43.1423331
O 5.8136015 5.298277 -42.1811172
C 8.7040589 6.5417867 -43.5937623
H 9.3769214 6.2789443 -42.781539
H 8.5857406 5.6690598 -44.2357264
C 9.2911549 7.7100855 -44.425368
H 9.1907816 8.6410518 -43.8675988
C 10.7676023 7.5379856 -44.7641775
H 10.9094523 6.6899523 -45.4326336
H 11.347273 7.4053009 -43.8535032
H 11.1408564 8.4293545 -45.2677901
C 8.5546316 7.8484516 -45.7653518
H 8.6226347 6.9085588 -46.3141622
H 9.0244539 8.6267692 -46.3653165
H 7.5122898 8.121164 -45.6183874
N 5.6747873 5.8134553 -44.3622881
H 6.051826 6.3868802 -45.1032308
C 4.5891328 4.8649351 -44.6980872
H 4.5631062 4.1139725 -43.9051263
C 4.8447935 4.0459821 -45.973334
O 3.9017478 3.6498304 -46.6520897
C 3.2105171 5.5603428 -44.666108
H 2.4533252 4.7738203 -44.6831793
H 3.1054279 6.0864467 -43.7153432
C 2.9161759 6.5454592 -45.8179066
H 3.1261552 6.0697121 -46.7735315
H 3.5448542 7.4315362 -45.7241825
C 1.4308664 6.9484155 -45.7776237
H 1.2590212 7.5693356 -44.8966521
H 0.8212512 6.0482975 -45.6740764
N 0.981203 7.6811794 -46.9780385
H 0.7524881 8.6673509 -46.8923419
C 0.6228584 7.177358 -48.1443321
N 0.8186153 5.929421 -48.4685675
H 0.5752193 5.6132175 -49.3870961
H 1.388143 5.3577715 -47.8677134

N 0.0406873 7.949872 -49.0092621
H -0.2933387 8.855375 -48.6610897
H -0.2623347 7.6163466 -49.9005672
N 6.1190758 3.8578277 -46.3209588
H 6.8301691 4.1363819 -45.6635574
C 6.5958218 3.2428633 -47.5626859
H 5.830005 2.5719029 -47.9542416
C 7.8916732 2.4367347 -47.343168
O 8.5668267 2.64766 -46.3401455
C 6.8674228 4.3491416 -48.5851797
H 7.7970464 4.8464069 -48.304794
H 7.032437 3.8770175 -49.5533758
C 5.793543 5.4093771 -48.7466088
C 4.5694425 5.071977 -49.3430479
H 4.3617759 4.0434651 -49.5976387
C 3.6563468 6.0786253 -49.6922932
H 2.7837729 5.8310304 -50.2772817
C 3.9157208 7.4151039 -49.3467661
H 3.2349086 8.196869 -49.6514519
C 5.0976506 7.7468283 -48.6644446
H 5.3156042 8.7854837 -48.4512446
C 6.0470079 6.7471097 -48.3865945
H 6.9997683 7.0245668 -47.966154
N 8.2414517 1.5676536 -48.3001291
H 7.6597218 1.5224626 -49.118932
C 9.3979198 0.6578697 -48.2914649
H 9.1016447 -0.2646165 -47.791358
C 10.6602547 1.1969945 -47.5742744
O 11.1786285 2.245083 -47.9427162
C 9.7087369 0.3224341 -49.75895
H 9.967396 1.234773 -50.3010873
H 8.8483582 -0.1504811 -50.2328802
H 10.5608699 -0.3531544 -49.8204047
N 11.1058984 0.4683517 -46.5404187
H 10.6020477 -0.3756197 -46.3181145
C 12.3498292 0.6411544 -45.7620579
H 12.0842348 0.2321049 -44.78769
C 12.7677959 2.0582833 -45.3295639
O 13.9257997 2.2966503 -44.965395
C 13.488643 -0.2542563 -46.2966031
H 14.4106292 0.3125754 -46.3885813
H 13.233996 -0.5667301 -47.3062372
C 13.7404825 -1.5012609 -45.4212878
O 12.782663 -1.9446545 -44.7479203
O 14.8558588 -2.0616043 -45.4835954
N 11.7676098 2.9221305 -45.1175289

H 10.8663841 2.6705479 -45.5032949
C 11.8434355 4.1137479 -44.2565654
H 10.9702733 4.7131008 -44.4953178
C 13.0474841 5.0013958 -44.5690478
O 13.2566867 5.4146228 -45.7033659
C 11.6839069 3.7215239 -42.7654516
H 10.7042933 3.2772 -42.7433744
C 12.7172875 2.6995277 -42.243958
H 13.7411237 3.021697 -42.4103113
H 12.5627012 1.7344082 -42.7242798
H 12.5786212 2.5252549 -41.1832717
C 11.6078686 4.8907969 -41.7666777
H 12.5088229 4.963613 -41.1569353
H 10.7748965 4.7152793 -41.0967139
H 11.4403098 5.837781 -42.2673589
N 13.7997814 5.3551081 -43.5357305
H 13.5427582 4.9757027 -42.6392564
C 14.9722896 6.1999847 -43.5773004
H 14.7513394 7.1316738 -44.1038787
C 16.1325397 5.5277022 -44.3124657
O 16.971215 6.2121722 -44.8791594
C 15.3363167 6.5059358 -42.1167684
H 14.5993989 7.1982355 -41.7073855
H 15.2826306 5.5845827 -41.5328418
C 16.7149966 7.0988042 -41.9408267
C 16.9780411 8.3984896 -42.403626
H 16.1908624 8.9809401 -42.8654804
C 17.7536204 6.3218351 -41.3924162
H 17.5600729 5.3123153 -41.0611771
C 18.2759221 8.9240865 -42.299878
H 18.4802673 9.9218835 -42.6403416
C 19.0546992 6.8452967 -41.3078177
H 19.8555147 6.2433117 -40.90308
C 19.314826 8.149128 -41.7601691
H 20.3159865 8.5514257 -41.6963891
N 16.2058513 4.196207 -44.3268838
H 15.3924863 3.6449569 -44.0809636
C 17.3365519 3.5317345 -44.9880782
H 18.247824 4.0148928 -44.6328091
C 17.3362498 3.7162578 -46.5202865
O 18.411362 3.6889561 -47.1212283
C 17.4139431 2.0551039 -44.544089
H 17.2640812 2.0186564 -43.4640312
C 18.7685605 1.4097662 -44.8475106
H 18.7772956 0.3945679 -44.4540022
H 19.5699411 1.9849547 -44.3903993

H 18.9200973 1.3641078 -45.9264033
O 16.4573901 1.2286298 -45.1599023
H 15.5577683 1.5969757 -45.0914068
N 16.1818698 4.0690221 -47.1078517
H 15.3465728 4.1192239 -46.5422669
C 16.0387294 4.3659216 -48.539581
H 16.9320402 3.9392428 -48.9945931
C 16.0907108 5.8966401 -49.0803814
O 16.0794458 5.9767522 -50.2908172
C 14.8556147 3.4957892 -49.0773819
H 13.9798023 4.1366509 -49.1933825
H 14.5968292 2.7346661 -48.3378472
C 15.1275989 2.7525254 -50.4024108
O 14.1400315 2.2286328 -50.965854
O 16.307982 2.6299838 -50.8096799
N 16.2900343 7.1238414 -48.4203981
H 16.4497442 7.0131143 -47.4277553
C 16.1613041 8.688626 -48.8951202
H 15.5643498 8.6270493 -49.8051702
C 17.4543756 9.8556596 -49.3389581
O 18.4787587 9.2741032 -49.0634247
C 15.1839224 9.2297703 -47.7945255
H 14.6070166 8.4267525 -47.3302822
H 14.4664933 9.8869038 -48.2885934
C 15.9020718 10.0680588 -46.713811
O 16.3872989 11.1886415 -47.009515
O 15.9809798 9.6471257 -45.537687
N 17.6582507 11.2710311 -50.0187799
H 16.7432841 11.6473397 -50.215925
C 18.9196021 12.4999217 -50.4996121
H 19.4239745 12.7072473 -49.557024
C 19.8910586 11.6284874 -51.3214552
H 19.3040143 10.926469 -51.916006
H 20.4598406 12.2416738 -52.0211191
C 20.8996192 10.8577388 -50.4511839
H 20.4908512 10.6758339 -49.4595413
H 21.799341 11.4600084 -50.3269145
C 21.2793501 9.5187183 -51.0747255
O 22.2065281 9.4215954 -51.8603748
N 20.5592735 8.459898 -50.7803827
H 20.794535 7.5910496 -51.2218482
H 19.7724068 8.5434667 -50.1355936
C 19.092366 14.182846 -51.3913657
O 18.0971829 14.350783 -52.0856275
N 20.1085432 15.3856543 -51.5493382
H 20.9260687 15.212857 -50.986138

C 20.2948739 16.9461427 -52.3131094
H 20.2227503 16.7196393 -53.3770916
C 21.4078328 18.3179152 -52.2406162
O 22.5468109 18.183726 -51.7988277
C 19.0686253 17.7464106 -51.9697146
H 19.2338989 18.2728335 -51.0391949
H 18.2005998 17.0932304 -51.9204808
H 18.8667864 18.4802587 -52.7438545
N 21.1702238 19.6749261 -52.6514902
H 20.2345851 19.8379585 -52.9935248
C 22.0024176 21.0115075 -52.4479166
H 22.8683875 20.7431257 -51.8441611
C 21.2832516 22.1579803 -51.6588977
O 20.098186 22.3880363 -51.8856444
C 22.5113401 21.7661131 -53.7349204
H 21.6390627 22.0773934 -54.3111731
H 23.0283835 22.6774737 -53.4283394
C 23.4507222 21.0203275 -54.6889031
H 22.9115143 20.1707727 -55.1106251
H 23.7236569 21.6915994 -55.5045992
C 24.7266258 20.5101083 -54.0066815
H 24.4530278 19.8220364 -53.2042071
H 25.2791937 21.3544152 -53.5879837
C 25.5984771 19.7681775 -55.0268131
H 25.909443 20.4728439 -55.8051529
H 24.9888066 18.9920389 -55.5006992
N 26.7854235 19.1515691 -54.3842775
H 27.3653918 19.8580064 -53.9512649
H 27.3408378 18.6543916 -55.0695705
H 26.4940335 18.4923601 -53.6723818
N 22.0141776 22.9790716 -50.8776924
H 22.9848334 22.7561847 -50.7481297
C 21.4363151 24.0595014 -50.0406279
H 20.6651534 23.5880877 -49.4217957
C 20.6932301 25.1571828 -50.8434591
O 21.1123859 25.4928957 -51.9501055
C 22.4954675 24.6021902 -49.0576117
H 21.9681595 25.0697176 -48.2281703
H 23.0446883 23.7611733 -48.6298199
C 23.4967813 25.6136439 -49.6395768
H 24.1308012 25.1185841 -50.3761876
H 22.9616084 26.426131 -50.1294037
C 24.3600993 26.2058559 -48.5122502
H 25.0695733 25.4548231 -48.159759
H 23.7129745 26.4715583 -47.6719088
C 25.1007745 27.47991 -48.9414417

H 25.6487684 27.8643149 -48.0751497
H 24.3538345 28.231996 -49.2207631
N 26.033051 27.2429212 -50.0717632
H 26.729265 26.5557402 -49.8134378
H 26.5058074 28.1019389 -50.3270792
H 25.5216444 26.9054137 -50.8778729
N 19.6653497 25.7902966 -50.2502701
C 18.5260935 26.4199977 -50.9302971
H 17.683172 26.2611953 -50.25573
C 17.9666354 25.9230463 -52.2825018
O 17.2120042 26.6858309 -52.8855268
C 18.8245867 27.9163968 -50.8994079
H 19.6064095 28.1600803 -51.6210341
H 17.9311673 28.5198536 -51.0658878
C 19.9982801 26.7186724 -49.1631419
H 19.6497361 26.3469128 -48.204326
H 21.0706298 26.8855675 -49.103357
C 19.3411503 28.0678737 -49.4726293
H 18.5032263 28.2293829 -48.7937525
H 20.0551531 28.8885143 -49.395854
N 18.1444302 24.6585658 -52.7170806
H 18.7575415 24.0313991 -52.2081714
C 17.2497154 24.0944877 -53.7603078
H 17.1809009 24.8305569 -54.5644565
C 15.8108259 23.9536811 -53.2184155
O 15.6116559 23.7733122 -52.0168994
C 17.7905208 22.7718561 -54.367871
H 18.2164828 22.1394345 -53.5915213
H 16.9504542 22.2274091 -54.799801
C 18.8275157 22.9966047 -55.490398
H 19.7490875 23.3914824 -55.0618205
H 18.4292468 23.7618966 -56.1540239
C 19.1471149 21.7322105 -56.331149
H 18.3107319 21.0347568 -56.2604309
H 20.0375569 21.2460875 -55.9308658
N 19.3493069 22.0762304 -57.7577649
H 19.2235844 23.0639289 -58.0004847
C 19.3902273 21.3013719 -58.8329725
N 19.5688706 20.0105413 -58.789513
H 19.5868354 19.4947518 -59.6476506
H 19.6328429 19.5190873 -57.9186205
N 19.2411043 21.8219077 -60.01574
H 18.7989704 22.7503794 -60.07748
H 19.2658129 21.2691169 -60.8598066
N 14.821947 23.9635607 -54.1171727
H 15.0640462 24.1295175 -55.0795782

C 13.4070457 23.7053603 -53.8023649
H 13.2318405 23.8314889 -52.7303655
C 13.0148305 22.2690653 -54.1901151
O 13.2650034 21.8554559 -55.3268438
C 12.5042714 24.70102 -54.554355
H 11.4657565 24.4018961 -54.4176821
H 12.7363243 24.6603459 -55.6166903
C 12.5918951 26.14774 -54.1111584
O 13.1869008 26.5106661 -53.1113018
N 11.9857542 27.0281791 -54.8750885
H 12.0663734 27.990201 -54.6088372
H 11.474667 26.7304553 -55.7012412
N 12.3104354 21.5618859 -53.3039725
H 12.1319232 21.9882772 -52.3959269
C 11.7798987 20.2088839 -53.5218785
H 12.1354009 19.8236533 -54.4750295
C 10.2499869 20.2006078 -53.5304971
O 9.636071 20.6928764 -52.5837385
C 12.2939867 19.2814334 -52.4104324
H 12.0003572 19.7111831 -51.4550013
H 13.3811827 19.2399207 -52.459261
C 11.7331357 17.8531895 -52.451972
H 10.6473428 17.8812916 -52.5280063
H 11.9562163 17.3702826 -51.5054196
S 12.3827862 16.813984 -53.7779941
C 13.8931229 16.2105975 -52.9728887
H 14.4627188 15.602152 -53.6752431
H 13.6285852 15.6018071 -52.1075691
H 14.5036321 17.0516099 -52.6454799
N 9.6561252 19.476022 -54.4810838
H 10.2356146 19.1014098 -55.228018
C 8.2618632 19.0294284 -54.4110718
H 7.7432147 19.5950109 -53.6411374
C 8.2012683 17.5367186 -54.0391612
O 8.6091906 16.6916482 -54.8315145
C 7.5848191 19.3377223 -55.7563497
H 7.6533027 20.404727 -55.9472126
H 8.1407942 18.8235216 -56.5354205
C 6.1094319 18.9161937 -55.8471269
H 6.0223168 17.8621598 -55.5987058
C 5.2008171 19.7105303 -54.9111302
H 5.2842959 20.7773402 -55.1160026
H 5.4789999 19.5191252 -53.8783082
H 4.1706631 19.3882301 -55.0433692
C 5.5958432 19.1157678 -57.2735893
H 5.6369121 20.1715894 -57.5396105

H 4.5696247 18.7588489 -57.3505647
H 6.2159663 18.5490014 -57.9673902
N 7.650806 17.207511 -52.8705797
H 7.2809473 17.9542867 -52.2913677
C 7.3710433 15.8279812 -52.4464654
H 8.0060151 15.1508264 -53.0126345
C 5.9027388 15.4793458 -52.7611271
O 4.9709185 16.075372 -52.2065813
C 7.75986 15.6405981 -50.9667627
H 7.1203425 16.2573012 -50.3370099
H 8.7862314 15.9905422 -50.8370111
C 7.6824443 14.1863395 -50.4760692
O 6.8708591 13.4090086 -51.019936
O 8.4087115 13.8653462 -49.5055914
N 5.6864177 14.5711102 -53.7189983
H 6.4974395 14.0713565 -54.0777662
C 4.3447916 14.1842873 -54.1843736
H 3.6345714 14.9349283 -53.8406842
C 3.9351687 12.8544228 -53.5605881
O 4.3232699 11.7831128 -54.0192536
C 4.2336867 14.1575708 -55.7201635
H 4.8568936 13.3737071 -56.1278179
C 2.7974381 13.8629456 -56.1479142
H 2.1426664 14.6095088 -55.703318
H 2.4969621 12.8683791 -55.8186253
H 2.7229068 13.8922377 -57.2343704
C 4.6461988 15.4961607 -56.3443168
H 4.0765308 16.3124174 -55.897042
H 4.4772019 15.482627 -57.4198195
H 5.7094427 15.6583488 -56.1699027
N 3.0257182 12.9205146 -52.5909151
H 2.717312 13.8404901 -52.2952782
C 2.6511322 11.7941466 -51.7378099
H 1.5919382 11.8736081 -51.495824
H 2.826334 10.8515972 -52.2505211
C 3.4378837 11.764253 -50.4292475
O 3.7783718 10.6859284 -49.9559292
N 3.6725557 12.9262538 -49.8125931
H 3.3218807 13.7634739 -50.2589932
C 4.5639542 13.1282033 -48.6574106
H 5.5763021 12.8435635 -48.9560907
C 4.2394702 12.317265 -47.379758
O 4.9944892 12.3378076 -46.4012804
C 4.557135 14.6320349 -48.3562011
H 4.9178299 15.1702423 -49.2356233
H 5.2327209 14.8285762 -47.5267004

S 2.8894722 15.2074225 -47.9074698
H 2.7061254 14.4032963 -46.8479299
N 3.0995178 11.6227276 -47.3420058
H 2.5537956 11.6005599 -48.1895828
C 2.6973692 10.7695608 -46.2284903
H 1.7296098 10.3306215 -46.4382888
H 3.4252333 9.964224 -46.1301078
C 2.5964225 11.5335775 -44.9096616
O 2.0756425 12.6458024 -44.8617237
N 3.1475683 10.9405853 -43.8409104
H 3.5995258 10.050913 -43.9813798
C 3.2644505 11.5676352 -42.5110625
H 2.5363332 12.3721322 -42.4922009
C 4.6024855 12.3215117 -42.2989558
O 4.9330438 12.7035656 -41.1837733
C 2.7685976 10.6219858 -41.3876643
H 3.329892 9.6866489 -41.3966698
H 2.9110664 11.1133101 -40.4234859
C 1.2482459 10.3497499 -41.5881163
H 0.7919812 11.3134804 -41.7977461
H 1.1127473 9.7337692 -42.4784474
C 0.4227669 9.7161131 -40.4391496
H 0.7314226 10.1512193 -39.4901884
H 0.5960085 8.639396 -40.4025918
C -1.0758004 10.0217816 -40.7095582
H -1.41602 9.4220592 -41.5561837
H -1.1388727 11.0716698 -41.0127608
N -2.0107233 9.8625097 -39.5598994
H -2.0996365 8.9127489 -39.2334567
H -2.9406933 10.1949484 -39.8135329
H -1.7309402 10.4529809 -38.7839952
N 5.3030928 12.6456133 -43.3932679
H 4.9509657 12.3158047 -44.2839729
C 6.263606 13.7539353 -43.4685514
H 6.339384 14.0373157 -44.5168193
H 5.8664254 14.6025745 -42.9108326
C 7.6966603 13.5217271 -42.9844836
O 8.5229441 14.4189115 -43.1446022
N 8.052298 12.3272621 -42.5032093
H 7.3432049 11.6119669 -42.3818921
C 9.4128024 12.0200965 -42.040962
H 9.6646061 12.6654394 -41.1981862
H 9.4474178 10.9834761 -41.7123813
C 10.481707 12.1834795 -43.1192423
O 11.5222019 12.7773601 -42.8556136
N 10.1750316 11.7843984 -44.3589289

H 9.2846575 11.3332788 -44.4955374
C 11.0186339 12.0413754 -45.5425943
H 11.9980506 11.588156 -45.3790969
C 11.2701527 13.5389522 -45.7486745
O 12.4236188 13.9518113 -45.8260238
C 10.4258214 11.3832015 -46.8083101
H 10.5905208 10.3099959 -46.7269139
C 8.9312833 11.6033541 -47.0463739
H 8.7099762 12.6632397 -47.1442538
H 8.3393666 11.1827496 -46.2344072
H 8.6361123 11.1131843 -47.9758533
O 11.0895781 11.8306891 -47.9597574
H 10.484578 12.3844576 -48.4701324
N 10.2252136 14.371186 -45.663101
H 9.3074535 13.9722939 -45.5446904
C 10.2857366 15.8236971 -45.8547703
H 10.7235008 16.0265557 -46.8268567
C 11.1106349 16.537419 -44.7650356
O 11.8863183 17.4526382 -45.0535196
C 8.8657689 16.3970441 -45.8856242
H 8.8986116 17.4030429 -46.3069571
H 8.4867539 16.4648006 -44.8668808
S 7.7163401 15.3759861 -46.8510702
H 8.4533831 15.1086649 -47.9512639
N 10.9632892 16.0999518 -43.508704
H 10.2825801 15.3646959 -43.3383485
C 11.7666912 16.5933009 -42.3777083
H 11.7210304 17.682167 -42.3558897
C 13.2337723 16.2149077 -42.5646649
O 14.1041332 17.0794444 -42.4631432
C 11.2181531 16.0596538 -41.0385742
H 11.0788588 14.9803609 -41.1088303
C 12.1583111 16.3454275 -39.8608403
H 12.413162 17.4044263 -39.8365978
H 13.0713173 15.7583224 -39.9544092
H 11.6747709 16.0658247 -38.9240883
C 9.8619149 16.7101419 -40.7311641
H 9.9738266 17.7906836 -40.6408162
H 9.465485 16.313601 -39.7953368
H 9.147258 16.4904463 -41.5236473
N 13.5086787 14.9554922 -42.9189931
H 12.7374487 14.300883 -43.0121126
C 14.8616324 14.4600437 -43.1763215
H 15.4605874 14.5795315 -42.2714439
C 15.5717593 15.257738 -44.2626702
O 16.6984701 15.6980688 -44.0520232

C 14.8119003 12.9696725 -43.5454865
H 15.47017 12.778969 -44.3885111
H 13.8159203 12.6800994 -43.8734669
C 15.2350372 12.1081566 -42.3887418
N 16.5503927 11.9810507 -41.9354251
C 16.4753185 11.2196844 -40.8299497
H 17.3253826 10.9192111 -40.2306378
N 15.2029599 10.8627996 -40.5836198
H 14.890693 10.2906702 -39.8085624
C 14.4037633 11.4233475 -41.5538219
H 13.3257702 11.3738953 -41.6274226
N 14.9078435 15.5075468 -45.387301
H 13.989292 15.091668 -45.5152898
C 15.5112271 16.2350314 -46.5011762
H 16.4944499 15.8005517 -46.6810838
C 15.788866 17.702243 -46.1455768
O 16.9395239 18.1374501 -46.2012692
C 14.6897345 16.0179734 -47.7882799
H 15.1327128 16.6236197 -48.5603665
C 14.7908711 14.5234165 -48.182398
H 14.1962195 13.8987777 -47.5198436
H 15.8220997 14.1781548 -48.1262874
H 14.4372692 14.3716629 -49.2011021
C 13.2366266 16.4975189 -47.7320752
H 12.7646062 16.0195718 -46.8976035
H 13.2059957 17.5711979 -47.5585925
C 12.4407379 16.1837724 -49.0033466
H 12.2244151 15.1164961 -49.062049
H 13.0189562 16.48687 -49.8741816
H 11.4970485 16.7230964 -48.9888032
N 14.8187234 18.4245126 -45.5793613
H 13.9008259 18.0120409 -45.4473034
C 15.0314858 19.8172963 -45.1857533
H 15.4248899 20.3646793 -46.0422485
C 16.0654803 19.9815299 -44.052507
O 16.775709 20.9872676 -44.0197683
C 13.6739593 20.4215669 -44.8288207
H 13.2487701 19.8884107 -43.9775321
H 12.9997992 20.3346726 -45.682005
H 13.7950387 21.4777533 -44.588073
N 16.1945128 19.0031707 -43.1417661
H 15.5550477 18.2125874 -43.1960906
C 17.1426889 19.0652184 -42.0165262
H 17.2682905 20.115006 -41.7540704
C 18.5555244 18.5806105 -42.3633946
O 19.5022617 19.2022987 -41.8920977

C 16.5356863 18.3408313 -40.8014493
H 15.5546198 18.7709715 -40.5929376
H 16.3868088 17.2899072 -41.0560764
C 17.3783728 18.4041283 -39.5137768
H 18.2761985 17.8007418 -39.655826
H 16.803368 17.9420041 -38.7110996
C 17.8159365 19.8120997 -39.0658109
H 18.4907804 20.2290409 -39.816017
H 18.3932875 19.7217441 -38.1440225
N 16.6897875 20.7491832 -38.8609668
H 16.5665419 21.4502419 -39.5716769
C 15.8807065 20.8193544 -37.8169131
N 14.9853566 21.7589121 -37.7435133
H 14.3645694 21.8061651 -36.956886
H 14.8726153 22.4170928 -38.4946404
N 15.9472023 19.9843426 -36.8189475
H 16.6241288 19.2455779 -36.8502505
H 15.2949912 20.0477172 -36.0589826
N 18.7300829 17.5290883 -43.1764286
H 17.9068943 17.0488226 -43.5306661
C 20.0684893 17.0390531 -43.5746573
H 20.7065707 16.9291063 -42.6964889
C 20.7865705 18.0002679 -44.5366372
O 22.0134416 18.0053269 -44.5586972
C 19.9813346 15.699411 -44.3219547
H 19.304658 15.8747004 -45.1616404
H 20.9671976 15.5209637 -44.7569981
C 19.5628823 14.3578569 -43.6784448
H 20.3290906 14.0366348 -42.971993
H 18.6129878 14.4437749 -43.1562454
C 19.4543242 13.3380914 -44.844005
H 20.418487 13.3271226 -45.354375
H 18.7167883 13.694725 -45.5669937
C 19.1362141 11.8730891 -44.4980398
H 19.490443 11.6480878 -43.4892399
H 19.686472 11.2395421 -45.2001729
N 17.6955314 11.5486037 -44.6343123
H 17.1272324 12.0169392 -43.9401613
H 17.4806865 10.5531329 -44.5600195
H 17.3202713 11.7308229 -45.5705452
N 20.0478271 18.6624719 -45.4364741
H 19.046923 18.5093921 -45.436498
C 20.6358853 19.3309775 -46.6085273
H 21.6749389 19.5607893 -46.3684505
C 20.0541746 20.7091617 -46.9622581
O 20.3152965 21.2210728 -48.0515675

C 20.6868561 18.354292 -47.7957786
H 21.0473326 18.879063 -48.6814314
H 21.4566581 17.6177888 -47.5613333
C 19.4373287 17.602644 -48.1667651
C 19.394892 16.1940631 -48.1271936
H 20.196105 15.6434021 -47.6560693
C 18.3451897 15.5113149 -48.7728262
H 18.2923324 14.4372359 -48.7484014
C 17.3896775 16.2497781 -49.5044677
O 16.5031357 15.6745726 -50.3589183
H 16.8175994 14.8393012 -50.723428
C 17.3695605 17.6457992 -49.4114359
H 16.6398896 18.1806343 -49.9827419
C 18.3848271 18.32481 -48.7295873
H 18.4231516 19.3990316 -48.7319403
N 19.3517895 21.3539171 -46.024161
H 19.1651048 20.8775033 -45.1539639
C 19.0384741 22.7911583 -46.0750765
H 18.3539854 22.9771147 -45.2457682
C 18.2660689 23.2374827 -47.3416898
O 18.3498463 24.3844502 -47.7803859
C 20.3254224 23.5929801 -45.7682459
H 21.0190659 23.4837652 -46.6030665
H 20.8007875 23.1682607 -44.8823023
C 20.0831759 25.0814136 -45.500918
O 19.0655738 25.4065578 -44.8420731
O 20.8655128 25.9146396 -46.0124582
N 17.4548563 22.3556176 -47.9295338
H 17.4238769 21.424189 -47.5465261
C 16.4823397 22.7281121 -48.9710723
H 16.9464183 23.4477338 -49.6473765
C 15.24281 23.3977265 -48.3725054
O 14.9522023 23.2457856 -47.1836795
C 16.0422905 21.5162724 -49.8130752
H 15.2603537 21.8499514 -50.4920367
C 17.2011516 21.0558416 -50.7125561
H 18.1143336 20.9147098 -50.1404907
H 17.4026366 21.8357731 -51.4439333
H 16.943333 20.1436236 -51.2459316
C 15.398919 20.4182088 -48.9405732
H 16.1665696 19.8238888 -48.4571155
H 14.7955406 20.8598812 -48.151009
C 14.4493492 19.5350595 -49.7478417
H 14.8862927 19.2531794 -50.7036183
H 13.5268763 20.0795328 -49.9194258
H 14.2201392 18.6408149 -49.1834476

N 14.4360211 24.010578 -49.2375354
H 14.7402083 24.0446147 -50.2085445
C 12.9993631 24.158699 -49.0076405
H 12.8175786 24.2573771 -47.9397338
C 12.2662528 22.9046082 -49.5103378
O 12.6107608 22.3634176 -50.5604784
C 12.5234879 25.4517901 -49.6859483
H 13.0406288 26.283309 -49.2048236
H 12.830272 25.44303 -50.731898
C 11.0112581 25.7338392 -49.6079951
H 10.8575365 26.7986213 -49.7828324
H 10.658697 25.5126149 -48.6002629
C 10.1424015 24.9839405 -50.6247752
O 10.5934761 24.3664138 -51.5757024
N 8.8434225 24.9954737 -50.4307784
H 8.2650384 24.4828781 -51.0906918
H 8.4331033 25.5005896 -49.6701504
N 11.2359635 22.4621827 -48.7844512
H 11.0578115 22.9049529 -47.8891113
C 10.3429926 21.383246 -49.214284
H 10.5949631 21.0920705 -50.235015
C 8.8711568 21.8264889 -49.2272665
O 8.3637704 22.3460571 -48.2286926
C 10.5732244 20.1618264 -48.309774
H 10.3332883 20.4190762 -47.2777976
H 11.6227969 19.8672222 -48.3609254
S 9.5400098 18.7650291 -48.8429061
H 8.3402798 19.3259352 -48.6493615
N 8.1565084 21.4877271 -50.299685
H 8.6580875 21.1050509 -51.0997332
C 6.6915622 21.4723115 -50.3796701
H 6.2622286 21.8728319 -49.4665579
C 6.2201094 20.031677 -50.5303368
O 6.4399128 19.4158451 -51.5664035
C 6.1943466 22.3420754 -51.5434893
H 6.7074003 22.060723 -52.4640156
C 4.6826819 22.2560971 -51.7615391
H 4.1511206 22.4652032 -50.8329504
H 4.408085 21.2625488 -52.117419
H 4.3832472 22.9787312 -52.5207642
O 6.4597851 23.693421 -51.2481149
H 6.0035097 23.908674 -50.423727
N 5.5750189 19.4858299 -49.5003679
H 5.4489048 20.0421054 -48.6609263
C 4.8934822 18.1915638 -49.5707935
H 5.3805521 17.5583986 -50.3075649

H 4.9561912 17.6920084 -48.6044006
C 3.4178491 18.3381219 -49.9451647
O 2.7499778 19.2457229 -49.4508098
N 2.8759735 17.4192718 -50.7447628
H 3.489359 16.728848 -51.172119
C 1.4218424 17.2768095 -50.9304672
H 0.9352328 17.8212694 -50.1261785
C 0.9560668 15.8305307 -50.7697592
O 1.6406314 14.8966804 -51.1867693
C 0.9136146 17.8905378 -52.2565359
H -0.1448096 17.6360649 -52.3435319
C 1.6387051 17.3248981 -53.5004349
H 2.6843131 17.6363338 -53.4916327
H 1.6106378 16.2363657 -53.4623999
C 0.9961788 19.4249524 -52.1858548
H 2.0365676 19.7521869 -52.1926341
H 0.5247676 19.7783184 -51.266841
H 0.4642133 19.8755679 -53.0218273
C 1.0052803 17.7519525 -54.8316522
H 1.1088672 18.8260369 -54.9781292
H -0.0515642 17.481976 -54.8436029
H 1.510511 17.2397663 -55.65089
N -0.2477491 15.6495421 -50.2260329
H -0.7465903 16.4668413 -49.8863459
C -0.9656272 14.3704334 -50.212057
H -0.6470092 13.784003 -51.0743246
C -2.4736961 14.6046495 -50.3361702
O -2.9923013 15.6108318 -49.855645
C -0.6345655 13.5626088 -48.9469876
H -1.035592 14.063649 -48.0667914
H 0.4483293 13.4718768 -48.8486547
O -1.2039403 12.2647033 -49.0661658
H -0.8230701 11.6633302 -48.3915009
N -3.189996 13.6624306 -50.9545782
H -2.693636 12.8593069 -51.3085801
C -4.6570956 13.7038033 -51.1109514
H -4.9453094 14.6681594 -51.5324737
C -5.3659256 13.5807981 -49.7505522
O -6.4750307 14.0756237 -49.5707605
C -5.0838402 12.5810381 -52.0946599
H -4.5868338 11.656628 -51.7934535
C -6.60297 12.3174183 -52.0927149
H -7.1427278 13.2367823 -52.3255887
H -6.9278242 11.951214 -51.1188279
H -6.8636599 11.5560362 -52.8261546
C -4.6265523 12.9470809 -53.5247128

H -5.2699191 13.744181 -53.8956398
H -3.6030739 13.3215718 -53.5000957
C -4.6555746 11.788384 -54.5300713
H -5.6777838 11.458972 -54.7113472
H -4.0625365 10.9539342 -54.1543668
H -4.2321241 12.1260136 -55.4769672
N -4.7271601 12.913471 -48.788543
H -3.775143 12.6272275 -48.9723444
C -5.2999645 12.5558781 -47.4919011
H -6.3581933 12.3426238 -47.6238231
C -5.12327 13.6827605 -46.4548777
O -3.9915082 13.9538372 -46.0403971
C -4.6452439 11.2446981 -47.0565748
H -3.587913 11.2809028 -47.3174804
H -5.1004324 10.4159779 -47.6001787
O -4.7366918 11.0128571 -45.6723862
H -5.6758702 11.0025864 -45.3613888
N -6.2181431 14.2734505 -45.9265974
C -6.1448261 15.2535177 -44.8392723
H -5.5037262 16.0768951 -45.1469994
C -5.5994107 14.6593758 -43.5365247
O -4.990551 15.3639187 -42.7414758
C -7.5767455 15.7683423 -44.6357585
H -8.056633 15.2366621 -43.8112928
H -7.5919532 16.8425494 -44.4489233
C -7.6005661 14.1249832 -46.3585443
H -7.6892693 14.0050037 -47.4364082
H -8.051605 13.2737373 -45.845865
C -8.2932888 15.4153499 -45.9362787
H -8.111043 16.1900229 -46.680453
H -9.3633916 15.2698172 -45.7853592
N -5.798928 13.3603836 -43.3244477
H -6.3334307 12.8472974 -44.0199305
C -5.3819279 12.5909009 -42.1533687
H -5.5086175 13.2194111 -41.2710266
C -3.889537 12.1953296 -42.1819728
O -3.2267231 12.2061379 -41.1417984
C -6.3412377 11.3905832 -41.9922851
H -7.3113502 11.7703061 -41.6675729
H -5.9575394 10.7498656 -41.1971473
C -6.5663179 10.5388814 -43.2538189
O -6.908747 11.078591 -44.3329274
O -6.3508923 9.3107568 -43.1812859
N -3.3008267 11.9683216 -43.3619274
H -3.8866016 11.9470974 -44.1933319
C -1.8430178 11.9831623 -43.5618261

H -1.3723892 11.257173 -42.9150736
C -1.2618897 13.3502526 -43.1982805
O -0.4067809 13.4546671 -42.3135894
C -1.4776396 11.6187214 -45.0103698
H -2.1218696 12.1563712 -45.7031098
H -0.4492902 11.9202012 -45.2048059
C -1.5826308 10.1109484 -45.2530965
H -2.542401 9.7508069 -44.8866851
H -0.8029225 9.6052696 -44.6769365
C -1.4265517 9.7523301 -46.7342513
O -2.1887154 8.9014817 -47.2378949
O -0.4387212 10.1870761 -47.3704002
N -1.8238071 14.3981459 -43.7998905
H -2.5652821 14.2183388 -44.4716944
C -1.3629664 15.7743681 -43.644951
H -0.2966211 15.7971953 -43.8619811
C -1.5011144 16.2671664 -42.1931932
O -0.5669668 16.8774303 -41.6842395
C -2.0937697 16.6432986 -44.6956286
H -3.1616424 16.4331521 -44.6208048
C -1.6453572 16.3062192 -46.1415492
H -1.8060816 15.2459139 -46.3228022
H -2.2857106 16.8442621 -46.8391984
C -1.9169274 18.1412204 -44.4519846
H -0.8624135 18.3648704 -44.4776171
H -2.351247 18.4235725 -43.4927015
H -2.4173335 18.7036014 -45.2397571
C -0.1850993 16.6247081 -46.5047295
H 0.502714 16.0852862 -45.8550906
H 0.005957 17.6948585 -46.4390428
H -0.0012286 16.3148047 -47.5317343
N -2.5671139 15.9256739 -41.4689525
H -3.3267024 15.4447242 -41.9411177
C -2.7735512 16.3191514 -40.0675664
H -2.7786111 17.4067291 -40.0129945
C -1.6436206 15.8404344 -39.1348382
O -1.1884872 16.6049745 -38.2795479
C -4.1712491 15.8244989 -39.6523631
H -4.8964345 16.3669418 -40.2600162
H -4.276831 14.7663523 -39.8970248
C -4.5547229 16.0502673 -38.1793378
H -5.6347657 16.1922723 -38.1337433
H -4.0852274 16.9595111 -37.8033673
C -4.2106249 14.87867 -37.2602295
O -3.478791 13.9624486 -37.5900405
N -4.7524033 14.838826 -36.0651956

H -4.5608134 14.0108061 -35.5281391
H -5.3835633 15.5549238 -35.7602172
N -1.1004638 14.6424705 -39.3708469
H -1.5013084 14.0764096 -40.106404
C 0.0418472 14.1322373 -38.6116889
H -0.0436926 14.4501244 -37.5713118
C 1.3850099 14.6518156 -39.1239408
O 2.2660308 14.8908551 -38.3035319
C 0.0435491 12.6104591 -38.6527647
H 1.0465421 12.2488474 -38.4138007
H -0.2262749 12.3017169 -39.6598596
S -1.1117245 11.9445062 -37.4241918
H -2.0986728 12.845829 -37.6105998
N 1.5575739 14.8170369 -40.4417058
H 0.8054631 14.5690666 -41.0762493
C 2.7701594 15.422793 -40.987119
H 3.6444812 14.9045317 -40.5883586
C 2.8696392 16.8863013 -40.5636503
O 3.5847836 17.2487937 -39.6283336
C 2.7907183 15.3258836 -42.5238653
H 1.9156234 15.7985274 -42.9702768
H 2.7822393 14.2994419 -42.8465516
H 3.7036186 15.7788831 -42.9060338
N 2.1002966 17.7180647 -41.2791163
H 1.1970245 17.3057366 -41.4945638
C 2.6117307 18.5517644 -42.3861192
H 3.1341734 17.8723516 -43.0516016
C 1.3970127 19.0600589 -43.2242848
H 1.358215 18.4262558 -44.0974844
H 0.5018309 18.6692998 -42.7733033
C 1.1166488 20.5727843 -43.5444475
H 1.2915895 21.151315 -42.6635274
H 1.8411256 20.9728381 -44.2381746
C -0.3230066 20.9360951 -43.9914732
H -1.0227159 20.3720331 -43.3859128
H -0.4699022 20.6178342 -45.0179328
C -0.6963571 22.4298742 -43.8025919
H 0.1995555 22.9755746 -43.4898631
H -1.4260614 22.4979107 -42.9928553
N -1.2635441 23.0865246 -45.012343
H -0.5895041 23.0906335 -45.7804691
H -1.4992876 24.0597524 -44.8902245
H -2.0834365 22.6400856 -45.4285437
C 3.7642777 19.4606364 -41.9383207
O 4.6934606 18.9814363 -41.3095185
N 3.6311882 20.7771193 -41.9989438

H 2.8876535 21.1417414 -42.5604266
C 4.0788258 21.6000937 -40.8865894
H 5.1237588 21.3834285 -40.6673247
C 3.9999911 23.0820359 -41.3327063
H 3.0493725 23.2473995 -41.8409777
H 4.7944901 23.2717171 -42.0553756
C 4.1026739 24.1179475 -40.2399803
N 3.0918861 25.0331692 -39.9409579
C 3.4223691 25.5589047 -38.7527186
H 2.8010558 26.2448663 -38.192991
N 4.5752577 25.0358408 -38.3042514
H 4.8930887 25.1113105 -37.3429746
C 5.0364284 24.1390869 -39.2428601
H 5.8685618 23.4582498 -39.1352582
C 3.2420288 21.199852 -39.6385361
O 3.0797663 22.025223 -38.7500341
N 2.530232 20.0383135 -39.640792
H 2.865367 19.2416506 -40.1623869
C 1.4823966 19.8353562 -38.7041374
H 1.2481924 20.7922532 -38.2384218
C 2.0858318 19.0933861 -37.5361898
O 3.067777 19.6165891 -37.0050026
C 0.1413169 19.4928001 -39.3442083
H 0.1442248 18.4588372 -39.6817178
H -0.6141341 19.569777 -38.5611991
C -0.267998 20.4123699 -40.5175301
H 0.1183068 19.9509151 -41.3678312
C -1.7776922 20.4353344 -40.70414
H -2.2492346 20.8376265 -39.8066394
H -2.1313518 19.4171078 -40.8620015
H -2.0806715 21.041301 -41.5528518
C 0.1944014 21.8747087 -40.5369619
H 1.2344464 22.003532 -40.7787416
H 0.035205 22.3050513 -39.5470494
H -0.3843236 22.4742456 -41.2283432
N 1.4878269 18.020571 -37.0275607
H 0.680745 17.619781 -37.490007
C 1.8370793 17.567107 -35.682473
H 1.6480608 18.3923588 -34.994206
C 3.3255548 17.1909504 -35.5105984
O 3.975249 17.6990186 -34.5966676
C 0.897452 16.4265003 -35.2938252
H 1.0844826 15.5580817 -35.9233497
H -0.1416476 16.7397962 -35.4098183
H 1.0723946 16.150676 -34.2533891
N 3.895938 16.3935091 -36.4204664

H 3.3273487 16.0340792 -37.1822979
C 5.2891756 15.9425756 -36.3358269
H 5.4806631 15.6036171 -35.3165186
C 6.3276383 17.0397494 -36.6405866
O 7.4910534 16.9063308 -36.2670223
C 5.4618095 14.744717 -37.274927
H 5.2733635 15.0489023 -38.3063018
H 4.7617187 13.9547081 -37.0015401
H 6.47971 14.3605827 -37.1961688
N 5.9285687 18.1328717 -37.2982429
H 4.962632 18.1795496 -37.6016175
C 6.8314934 19.2418052 -37.635999
H 7.8429605 18.8505207 -37.6832907
C 6.844892 20.3829303 -36.6121256
O 7.5699517 21.3645417 -36.8076774
C 6.4883151 19.7654549 -39.0249614
H 5.4763035 20.1549587 -38.9918722
H 6.5452826 18.940506 -39.7349417
O 7.3909096 20.7802827 -39.4271918
H 7.5897174 21.3108912 -38.6363837
N 6.0301942 20.3258281 -35.5583977
H 5.4633655 19.4990513 -35.4131305
C 5.9102691 21.4436743 -34.6242259
H 5.8569889 22.3420848 -35.2375565
C 7.1639527 21.593333 -33.7492034
O 7.6308825 20.6298769 -33.149975
C 4.6035516 21.3149437 -33.8160339
H 3.9610644 20.5586151 -34.2705869
H 4.8261419 20.983383 -32.8007128
C 3.8146552 22.6339158 -33.7630863
H 4.3761121 23.361526 -33.1761685
H 2.8597389 22.454182 -33.2696363
C 3.5503843 23.2164719 -35.1512038
O 4.0029425 24.3035214 -35.4804174
N 2.9488639 22.4629638 -36.0411139
H 2.950617 22.7735293 -37.0102779
H 2.7918841 21.4817724 -35.8719266
N 7.7210904 22.8081187 -33.6914249
H 7.2987774 23.5567177 -34.2190106
C 8.9972279 23.0788989 -33.0131334
H 9.0701395 24.1482844 -32.8145984
H 9.0214978 22.5526099 -32.0586359
C 10.2493908 22.6746477 -33.805537
O 11.3286693 22.5814544 -33.2268775
N 10.1271567 22.3925806 -35.1093067
H 9.2142375 22.4390954 -35.5407744

C 11.2828804 22.1739636 -35.9765919
H 12.1470555 21.9182832 -35.3639204
C 11.6311631 23.4425166 -36.7703543
O 10.8468171 23.8739616 -37.6062429
C 11.0248787 21.0051895 -36.9419587
H 10.2282802 21.2982624 -37.627223
H 11.9359448 20.8776523 -37.5163873
C 10.6521745 19.6372191 -36.3484215
H 9.6886744 19.7031776 -35.8480046
C 10.554009 18.6256642 -37.4922608
H 11.5421073 18.4392663 -37.908598
H 9.9033941 19.0025286 -38.2786216
H 10.1402736 17.6890333 -37.1141196
C 11.700169 19.1190278 -35.363519
H 12.6761902 19.0615675 -35.8409176
H 11.4049478 18.1266817 -35.0192001
H 11.7434391 19.7788958 -34.4960605
N 12.8948299 23.8661966 -36.7091582
H 13.3352564 23.7065076 -35.812184
C 13.4155394 25.0179326 -37.4790076
H 12.6681205 25.8145446 -37.4282954
C 13.6732908 24.7387133 -38.9836209
O 14.3679382 25.4938641 -39.6582383
C 14.7076514 25.5493859 -36.8171534
H 15.5330047 24.8870948 -37.0856383
H 14.9254605 26.5347873 -37.2302733
C 14.686988 25.6625318 -35.2845744
H 15.5866248 26.1902592 -34.9596257
H 14.7320807 24.6561241 -34.8586402
C 13.4378158 26.3886491 -34.7758366
O 13.4280188 27.6368107 -34.819975
O 12.4900572 25.6633847 -34.4019596
N 13.2648524 23.5772005 -39.5105643
H 12.5354699 23.085907 -39.0068225
C 13.5570177 23.1795007 -40.8976494
H 14.5254876 23.6009869 -41.1761233
C 12.542468 23.7627457 -41.8860466
O 11.3445727 23.7642826 -41.6133993
C 13.6717175 21.6505914 -41.0511484
H 14.6737933 21.3409764 -40.7633607
H 13.5409867 21.3814689 -42.0977122
C 12.6807254 20.8409029 -40.234541
O 13.0819143 19.9571612 -39.4998137
N 11.4063889 21.1652132 -40.2593715
H 10.7663929 20.6171536 -39.7086994
H 11.0835354 21.9909526 -40.7547637

N 13.0133136 24.1123106 -43.0888082
H 14.003369 23.9861169 -43.252692
C 12.2559559 24.8063065 -44.14478
H 11.634147 25.5703974 -43.6725806
C 11.2900693 23.8636724 -44.8836361
O 11.4794884 23.5296958 -46.0528063
C 13.2402964 25.5207442 -45.0935189
H 13.7974318 24.7585126 -45.6299273
H 12.6747639 26.1075278 -45.8195301
C 14.2367 26.4457212 -44.363787
H 13.6782029 27.2076148 -43.8182901
H 14.8081668 25.8651489 -43.6389009
C 15.2530414 27.1445263 -45.2802328
H 15.9662655 27.6665972 -44.6388995
H 14.7410762 27.8844194 -45.8974409
C 16.0017683 26.1590706 -46.1895801
H 15.4383225 26.0365783 -47.1185131
H 16.0485386 25.1791219 -45.7028254
N 17.3833513 26.6013149 -46.4784672
H 17.4501689 27.5257558 -46.8646949
H 17.8394313 25.9286781 -47.0934372
H 17.9672793 26.5200239 -45.6378488
N 10.2586788 23.4106932 -44.1767312
H 10.1841574 23.7178249 -43.2124954
C 9.2046284 22.5279462 -44.6902692
H 9.4646982 22.2318761 -45.7066837
C 7.8664666 23.2561846 -44.773848
O 7.5173305 24.0732073 -43.9268036
C 9.0728533 21.2239852 -43.8778855
H 8.316634 20.6091786 -44.3642886
C 10.3766276 20.4230278 -43.8853184
H 11.1563743 20.9672999 -43.3545985
H 10.6901088 20.2389319 -44.9118237
H 10.2204614 19.4614051 -43.3955968
C 8.6444678 21.4437578 -42.4225151
H 9.3296493 22.1190396 -41.9159669
H 8.6253233 20.491972 -41.8909169
H 7.6444986 21.8751438 -42.3818422
N 7.0848025 22.9020958 -45.7835561
H 7.4542566 22.2468272 -46.4660422
C 5.688535 23.2901281 -45.9603944
H 5.2592379 23.5642908 -44.9971347
C 4.9368154 22.090787 -46.5162855
O 5.5175932 21.2799915 -47.2402207
C 5.572196 24.4861637 -46.9093264
H 6.1315988 25.3254024 -46.4927753

H 4.5253632 24.7775338 -47.0057321
O 6.0920421 24.166681 -48.1936715
H 6.9019288 23.6412406 -48.069159
N 3.6649475 21.9328174 -46.1572492
H 3.1924113 22.6509743 -45.6290061
C 2.8540294 20.891737 -46.7718369
H 3.2603949 20.7420939 -47.768523
C 1.4232672 21.3469321 -46.9993605
O 0.9655742 22.2461308 -46.2958749
C 2.8723023 19.5311039 -46.0727033
H 1.8998189 19.3593649 -45.6756641
H 2.831625 18.8431272 -46.9167144
C 4.0238134 18.9905664 -45.2169582
C 4.3071869 17.6253821 -45.3860567
H 3.6888653 17.0224533 -46.0309537
C 4.8919789 19.744078 -44.3979068
H 4.7530479 20.7996534 -44.2590226
C 5.3992881 17.0235059 -44.752768
H 5.5957283 15.9801922 -44.9285487
C 5.9619852 19.1264943 -43.7177794
H 6.5828412 19.700868 -43.0462872
C 6.2154413 17.761582 -43.8888187
H 7.0295482 17.2877361 -43.3583586
N 0.7049211 20.6843602 -47.9071269
H 1.1580329 19.9917536 -48.4999263
C -0.7426341 20.8387991 -48.0115702
H -1.0824204 20.9641876 -46.9898499
C -1.5361428 19.617803 -48.4852348
O -0.9924186 18.6595985 -49.0384563
C -1.11455 22.1121096 -48.8003637
H -1.7026334 21.8618073 -49.6844732
H -0.2187602 22.6483505 -49.119513
C -1.9431406 22.9948283 -47.8556361
O -2.8487189 22.4398679 -47.1824651
O -1.4945848 24.1167787 -47.540539
N -2.8489941 19.6723543 -48.2388324
H -3.2118883 20.5386921 -47.8451067
C -3.8093695 18.7281656 -48.8164363
H -3.3973647 17.7249245 -48.7310293
C -3.9734597 19.0752603 -50.2936677
O -4.3973474 20.175993 -50.6415666
C -5.1732061 18.7435119 -48.0992768
H -5.6730347 19.6950951 -48.2825975
C -6.0637773 17.6068255 -48.6193746
H -5.6052813 16.639463 -48.4108445
H -6.2151491 17.7031978 -49.6940242

H -7.0395154 17.6576898 -48.1394967
C -5.0359728 18.5600012 -46.5826229
H -4.4763312 17.6507279 -46.373113
H -6.0235974 18.4938287 -46.1264681
H -4.5200769 19.4177832 -46.1531065
N -3.6215795 18.1436875 -51.1741785
H -3.2879382 17.2507786 -50.8282856
C -3.6115106 18.3903012 -52.6077545
H -4.5005105 18.9612831 -52.879691
H -2.7342712 18.9845956 -52.864576
C -3.5764643 17.1062873 -53.4192378
O -2.760003 16.2192524 -53.1788795
N -4.4347018 17.0516582 -54.4326559
H -5.0732818 17.8204822 -54.5581563
C -4.3864658 16.05468 -55.4956002
H -4.0915978 15.0912157 -55.0883004
C -3.3390088 16.4662952 -56.5310674
O -3.4495113 17.5140222 -57.164842
C -5.7890218 15.9175663 -56.0942179
H -6.1264015 16.9003061 -56.4288134
H -6.4684305 15.5787826 -55.3090581
C -5.884318 14.9468145 -57.2730016
O -4.8383242 14.4918336 -57.7921059
O -7.0322555 14.7422459 -57.7062923
N -2.3237965 15.6294866 -56.7222922
H -2.3348741 14.7515165 -56.2290766
C -1.1959614 15.917138 -57.5968154
H -0.8042434 16.9026035 -57.3349828
C -1.5477935 15.9520276 -59.0940706
O -0.7986989 16.5414193 -59.8719076
C -0.1261968 14.8713849 -57.2991538
H -0.4332321 13.8964363 -57.6832249
H 0.0211865 14.8034629 -56.2217764
H 0.8044678 15.171372 -57.7763393
N -2.6894224 15.3899282 -59.5099429
H -3.3057519 14.9434825 -58.8268714
C -3.1680469 15.5081525 -60.8889374
H -2.2977069 15.4764396 -61.5426236
C -3.8714604 16.8578481 -61.1353749
O -3.8881519 17.3427345 -62.2717246
C -4.0589673 14.2899356 -61.2019821
H -3.5800509 13.3944349 -60.8022451
H -5.0094949 14.4145144 -60.6792547
C -4.3336983 14.054691 -62.7013104
H -4.8066073 14.9365606 -63.1323418
C -3.0607759 13.732287 -63.4923323

H -2.5395826 12.8897156 -63.037984
H -2.3987076 14.5952794 -63.5160859
H -3.3198034 13.4669686 -64.5171777
C -5.2831101 12.8667808 -62.868187
H -4.8269732 11.9632398 -62.4652122
H -5.5106869 12.7195113 -63.9246257
H -6.2135557 13.0676634 -62.3361046
N -4.3438173 17.5317856 -60.0767708
H -4.239743 17.0913676 -59.1689351
C -5.1392953 18.7670931 -60.1508346
H -5.3101043 19.014173 -61.1954728
C -4.4652452 20.0192241 -59.553186
O -4.7247452 21.1197762 -60.036646
C -6.5275032 18.4864972 -59.5562995
H -7.1041324 19.4099779 -59.5149897
H -6.4332717 18.0886791 -58.546576
C -7.2894454 17.5033969 -60.4354308
O -7.5563357 17.7986585 -61.5883171
N -7.5704357 16.2932746 -60.0008698
H -8.0536807 15.6856149 -60.632247
H -7.3300548 15.9419432 -59.0695682
N -3.5291447 19.8666245 -58.6129897
H -3.3778296 18.9328279 -58.2480178
C -2.7161973 20.948246 -58.0426512
H -3.3974747 21.6988151 -57.6398965
C -1.8381347 21.6244684 -59.108357
O -1.3603305 20.9679296 -60.0376461
C -1.8708616 20.3801509 -56.8857777
H -1.2155043 19.5960797 -57.2688558
H -2.5403692 19.9339514 -56.1488536
C -1.0113161 21.4298223 -56.1713507
H -0.2443703 21.7912918 -56.8567282
H -0.5059511 20.9491715 -55.3327881
S -1.9402459 22.8506813 -55.5394919
C -0.5701828 23.8292679 -54.8690719
H -0.9613234 24.74235 -54.4197375
H 0.1193026 24.0926016 -55.6715556
H -0.0410525 23.252321 -54.1098706
N -1.6015408 22.9369568 -58.9811233
H -1.9423219 23.4054305 -58.1486339
C -0.8192637 23.7390643 -59.9395053
H -0.2576772 23.064157 -60.5851844
C 0.2282464 24.5957342 -59.2270807
O 0.004776 25.0706294 -58.1183271
C -1.7509353 24.6002576 -60.822006
H -2.2485006 25.3361505 -60.1875853

H -1.1470927 25.1432575 -61.5512182
C -2.8355023 23.8041089 -61.576147
H -3.4692309 24.5095959 -62.1148217
H -3.4670243 23.293783 -60.85025
C -2.285634 22.7794283 -62.5838831
H -1.469829 22.2166229 -62.1362081
H -1.8727529 23.3160784 -63.440655
N -3.3480238 21.8600974 -63.0454371
H -4.1592275 22.2924263 -63.4494333
C -3.3712553 20.539867 -62.9367282
N -4.3632524 19.8552986 -63.436589
H -4.394121 18.8635073 -63.2050823
H -5.2093213 20.3214733 -63.7056537
N -2.4470361 19.8412767 -62.3456859
H -1.8072108 20.3142319 -61.7116381
H -2.5589807 18.8412342 -62.2636225
N 1.3477479 24.8147152 -59.9086339
H 1.3980397 24.4714973 -60.8561952
C 2.4801481 25.6382775 -59.4817666
H 2.171737 26.310206 -58.6796175
C 2.9540436 26.4820188 -60.6742369
O 2.5493109 26.2124622 -61.8048482
C 3.6101897 24.7291168 -58.9715134
H 3.9556652 24.0994097 -59.7921348
H 4.4519802 25.3525014 -58.6710504
C 3.2414508 23.8502424 -57.790578
C 3.2255769 24.3988016 -56.4940977
H 3.4578802 25.4439975 -56.3432923
C 2.944317 22.484486 -57.9805326
H 2.957988 22.0603603 -58.9749413
C 2.9118273 23.5871558 -55.3886793
H 2.893409 23.9989979 -54.390721
C 2.6376701 21.6672874 -56.8746023
H 2.4174114 20.6212776 -57.0198648
C 2.6196309 22.2218223 -55.5761608
O 2.3179465 21.4526177 -54.4997302
H 2.18435 20.5319836 -54.724826
N 3.8334992 27.4628261 -60.4509498
H 4.2082822 27.584594 -59.5219213
C 4.4939213 28.1620494 -61.5616058
H 3.7309422 28.5488971 -62.2384471
C 5.3882736 27.2020676 -62.3485732
O 6.0176283 26.3114741 -61.7616727
C 5.3217809 29.3477141 -61.0436929
H 6.091829 28.9980095 -60.3569351
H 4.6612574 30.029898 -60.5071673

O 5.9393037 30.0538829 -62.1032955
H 6.7243398 29.5613144 -62.4093461
N 5.5820288 27.4940815 -63.6317522
H 5.0106585 28.1958685 -64.0746187
C 6.7307802 26.9874338 -64.3760478
H 6.6366855 25.9085016 -64.4254029
C 8.028407 27.3558119 -63.6304401
O 8.1187891 28.4400932 -63.0417593
C 6.7479946 27.5450176 -65.8100117
H 7.719706 27.3149662 -66.2505881
H 6.6485008 28.6315395 -65.7731873
C 5.6672195 26.9697377 -66.7358325
O 4.772257 26.2412801 -66.2508656
O 5.7522914 27.2543315 -67.9504335
N 9.0046514 26.4438149 -63.5827792
H 8.8767 25.5757138 -64.0999271
C 10.2928452 26.6834173 -62.9168859
H 10.9628395 25.863104 -63.1596956
H 10.726691 27.5994604 -63.3176861
C 10.2581107 26.8168125 -61.3836782
O 11.1929088 27.3706979 -60.812188
N 9.2068629 26.3463782 -60.7000323
H 8.4713915 25.8971 -61.2280263
C 9.0731089 26.4374344 -59.2283582
H 9.1851045 27.4791121 -58.9258279
C 10.1069491 25.6136815 -58.4432941
O 10.4836519 25.994552 -57.3288833
C 7.6897912 25.9495096 -58.7796666
H 7.6527518 25.9341602 -57.6888603
H 7.5223822 24.9374057 -59.148062
O 6.6580577 26.7898418 -59.2502181
H 6.5249451 26.5892104 -60.1998395
N 10.5092653 24.4553591 -58.9775317
H 10.148396 24.2047745 -59.8925566
C 11.2703098 23.4359545 -58.2511685
H 11.53296 23.831252 -57.2725145
C 12.5826537 23.0539197 -58.9343996
O 12.7237566 23.0892687 -60.153064
C 10.4016771 22.1945978 -58.0069762
H 10.1678104 21.7295215 -58.9665107
H 10.9725225 21.4720044 -57.4234963
C 9.1174901 22.4918067 -57.2616053
C 9.1440574 22.7882535 -55.8861933
H 10.0770687 22.7663086 -55.3446341
C 7.8965917 22.5106125 -57.9559699
H 7.8822044 22.2996273 -59.0114969

C 7.9532069 23.1130515 -55.2127899
H 7.9771022 23.3415634 -54.1554797
C 6.7052395 22.8277978 -57.284616
H 5.7704859 22.841606 -57.8236995
C 6.7337942 23.135257 -55.9133376
H 5.8189192 23.3828901 -55.3936062
N 13.517583 22.5839715 -58.1188466
H 13.3072881 22.5350507 -57.1278463
C 14.8179288 22.0601079 -58.5444263
H 15.0314648 22.3351386 -59.5799209
C 14.8820061 20.5283609 -58.4571595
O 15.7122647 19.8905221 -59.1081171
C 15.861854 22.6970743 -57.6264017
H 15.4958654 22.6849266 -56.6000881
H 16.7664742 22.1004187 -57.6568063
C 16.1785143 24.1454572 -58.0345006
H 16.4935678 24.6931106 -57.143149
H 15.2907343 24.6425045 -58.4312527
C 17.305602 24.1476694 -59.067901
O 17.1190339 23.5958259 -60.1692975
O 18.44763 24.4781666 -58.6847995
N 14.0008482 19.9431131 -57.6415347
H 13.375073 20.5442361 -57.11928
C 13.8557196 18.5135042 -57.3961515
H 14.3000178 17.9505765 -58.2132188
C 12.3562485 18.1809218 -57.2916426
O 11.6055412 18.9392478 -56.6695756
C 14.5613807 18.1437631 -56.0764476
H 13.9703084 18.5856413 -55.2756477
H 14.5331341 17.0604963 -55.9537592
C 16.0075358 18.6382877 -55.8749913
H 16.0774433 19.7022195 -56.0961422
C 16.4272853 18.4705187 -54.4170553
H 16.4803628 17.4129 -54.1558347
H 15.7119752 18.972793 -53.7650525
H 17.3987076 18.9361346 -54.2814031
C 17.0207935 17.8886108 -56.7338335
H 16.9321508 16.814618 -56.5697651
H 18.0321975 18.2073195 -56.4892937
H 16.8337156 18.1120355 -57.7796173
N 11.9268476 17.0254245 -57.802589
H 12.5667476 16.4679935 -58.3580903
C 10.6134922 16.4408486 -57.4727742
H 10.200647 16.9817986 -56.6240596
C 10.8460628 14.9974708 -57.0338666
O 11.4968357 14.2196056 -57.7343886

C 9.584129 16.5453119 -58.623281
H 9.8891 15.8453796 -59.384544
C 9.5544193 17.9798466 -59.2141413
H 9.2019235 18.6905423 -58.4697116
H 10.5636784 18.2729503 -59.500348
C 8.1844474 16.0604285 -58.2049168
H 7.7911132 16.6907992 -57.4099126
H 8.2346799 15.0332518 -57.8434264
H 7.5072373 16.083636 -59.0554359
C 8.7221937 18.1222675 -60.4791129
H 7.6661354 18.1114382 -60.2270789
H 8.9737782 17.3097286 -61.1536164
H 8.9770545 19.0592215 -60.9698104
N 10.3564225 14.6574374 -55.8469035
H 9.7941865 15.3397182 -55.3446681
C 10.5610505 13.361721 -55.2088406
H 11.1527113 12.721799 -55.8623133
C 9.2265463 12.6740554 -54.9607565
O 8.2716464 13.3041168 -54.5053329
C 11.33476 13.5316808 -53.9000093
H 12.3374437 13.8914798 -54.1303004
H 10.8329836 14.2807638 -53.2847405
C 11.4560679 12.2414929 -53.107116
C 12.2584577 11.1893237 -53.5913009
H 12.8093437 11.3079028 -54.5115507
C 12.3285074 9.9712448 -52.892086
H 12.935959 9.1595666 -53.2719667
C 11.6002187 9.80203 -51.7022888
H 11.6597189 8.8679635 -51.1608939
C 10.7885537 10.8448729 -51.2227153
H 10.2173261 10.7141468 -50.3149552
C 10.7115946 12.0603749 -51.9250526
H 10.0647863 12.8469327 -51.5572318
N 9.1602094 11.3777451 -55.2613215
H 9.967209 10.9333375 -55.6892098
C 7.9328766 10.5948167 -55.1189933
H 7.301889 11.0701356 -54.3646456
C 8.255057 9.1831792 -54.6477118
O 9.0891373 8.509012 -55.2435789
C 7.1597415 10.5440188 -56.4471444
H 7.7114147 9.9222563 -57.1494066
C 5.8095002 9.8629732 -56.2113163
H 5.4896513 10.0212756 -55.1882349
H 5.9140703 8.7885214 -56.3956002
H 5.0585275 10.2781186 -56.8705113
C 7.0334598 11.9183157 -57.1168015

H 6.7794035 12.6694729 -56.3779092
H 6.2773547 11.9109892 -57.8926942
H 7.9869524 12.1928344 -57.5734035
N 7.5304832 8.6922801 -53.6430702
H 6.8760449 9.3036927 -53.1726615
C 7.6681163 7.3252057 -53.1358409
H 8.2068826 6.7406371 -53.8801015
C 6.2876119 6.6673452 -52.9862792
O 5.4097723 7.1751835 -52.2984167
C 8.5623854 7.3067255 -51.8731256
H 9.5579647 7.6240205 -52.1890165
C 8.094785 8.2976582 -50.7891014
H 7.0661867 8.0877915 -50.5071326
H 8.1462975 9.3195321 -51.1659953
H 8.7354529 8.2405992 -49.9112774
C 8.6896144 5.865025 -51.3446308
H 7.7397634 5.5693468 -50.9079072
H 8.8984866 5.198318 -52.1818034
C 9.804598 5.6643442 -50.3096542
H 9.5898272 6.2220642 -49.3977556
H 10.7584064 5.9929064 -50.7211156
H 9.8773923 4.6049462 -50.0589894
N 6.0735872 5.5791504 -53.7322323
H 6.8322139 5.3057652 -54.3480851
C 4.8587987 4.7465541 -53.7745885
H 4.9169297 4.1923917 -54.7089597
C 3.516118 5.5057891 -53.8614048
O 2.5191161 5.1118456 -53.2532807
C 4.8607276 3.7088065 -52.6438668
H 4.6254463 4.2273965 -51.7263386
H 4.0580549 2.9937707 -52.8329263
C 6.1573545 2.9237375 -52.4338794
H 5.9667513 2.1123083 -51.7318169
H 6.9089625 3.5794581 -51.9982039
C 6.7097361 2.3381834 -53.7201912
O 7.8204849 2.6367907 -54.124086
N 5.9596121 1.5408023 -54.4434662
H 6.3860314 1.1601171 -55.2659575
H 4.9443976 1.5731702 -54.3602218
N 3.4922299 6.609794 -54.6151327
H 4.3734201 6.9183661 -54.9925454
C 2.3102518 7.4619524 -54.7883182
H 1.523502 7.1422596 -54.1042785
C 1.7272705 7.3825057 -56.209906
O 0.5213259 7.2247492 -56.3878293
C 2.7044591 8.890572 -54.3960494

H 3.4554001 9.2689719 -55.0839756
H 3.1257726 8.8895343 -53.3873539
S 1.267988 9.9956678 -54.455393
H 1.9581013 11.1116626 -54.1764562
N 2.5785643 7.3956426 -57.2408166
H 3.5667076 7.4519611 -57.0465178
C 2.1412431 7.4998195 -58.6453044
H 1.3706166 8.2684934 -58.6961616
C 1.455654 6.224998 -59.1639819
O 0.6846533 6.2812467 -60.1185169
C 3.3271499 7.9979964 -59.498981
H 4.2271993 7.4604443 -59.1935633
C 3.5120954 9.5030321 -59.2234801
H 2.7433385 10.070109 -59.7525158
H 3.3888159 9.7067547 -58.1618482
C 3.1360484 7.7830679 -61.0060698
H 2.1735433 8.1823495 -61.3276595
H 3.2063335 6.7193164 -61.2171017
H 3.9295564 8.2650958 -61.5743967
C 4.8911054 10.0097662 -59.6504475
H 5.0099305 9.9628237 -60.7314375
H 5.6735898 9.4236341 -59.1685517
H 4.9797716 11.0455733 -59.3443017
N 1.5717582 5.1041732 -58.4564899
H 2.1827045 5.1206104 -57.6529454
C 0.7328439 3.9149034 -58.651388
H 0.8947388 3.5462889 -59.6639284
C -0.7823536 4.1812482 -58.5331897
O -1.5480582 3.5308124 -59.2486143
C 1.1982313 2.8047712 -57.6992863
H 0.5310809 1.9524817 -57.8022436
H 2.1807305 2.4791663 -58.0396875
C 1.2976814 3.1948038 -56.2119626
H 0.3012642 3.1688227 -55.7657597
H 1.6944627 4.205053 -56.0979221
C 2.2384522 2.2254211 -55.4939914
O 3.4720441 2.4194285 -55.590334
O 1.7426194 1.2099946 -54.9540701
N -1.2002262 5.2068174 -57.7761104
H -0.5129244 5.7123035 -57.2254629
C -2.598939 5.6605511 -57.6700557
H -3.2650489 4.7962077 -57.7183515
C -3.0249164 6.6303356 -58.7861373
O -4.212663 6.7435325 -59.0897442
C -2.8084363 6.3407486 -56.3074555
H -3.8525083 6.6461592 -56.2269157

H -2.1951503 7.2411331 -56.2540386
C -2.494111 5.4648105 -55.1222584
N -3.2748369 4.3995001 -54.6813414
C -1.402258 5.5724542 -54.3123075
H -0.6083186 6.2963122 -54.4092074
C -2.6439835 3.8919381 -53.61534
H -2.9894624 3.0311948 -53.0587202
N -1.5076655 4.5702295 -53.3754836
H -0.8044458 4.3204002 -52.6953078
N -2.0794667 7.3391896 -59.4136958
H -1.1158875 7.1556197 -59.1719981
C -2.3616997 8.3287098 -60.4667779
H -3.2276859 8.9173499 -60.1602128
C -2.7464924 7.5875773 -61.7548098
O -2.001621 6.7304906 -62.2333305
C -1.169923 9.3091347 -60.6337605
H -0.2399088 8.7535995 -60.5132491
C -1.1908147 10.461597 -59.5993055
H -0.3002541 11.0742561 -59.749503
H -2.0610367 11.0934907 -59.7853356
C -1.1248122 9.968303 -62.0256172
H -2.0519598 10.5058078 -62.2273489
H -0.9656117 9.221869 -62.8030621
H -0.2929621 10.6711498 -62.0761009
C -1.2165483 10.0459058 -58.1225757
H -0.4028053 9.3553923 -57.9105932
H -2.1699243 9.5788426 -57.8756608
H -1.0975652 10.9321865 -57.4986943
N -3.9191623 7.9002296 -62.3131057
H -4.4943561 8.5906527 -61.8567087
C -4.4811881 7.1647627 -63.4544377
H -4.2521094 6.1048541 -63.3300885
C -3.8496852 7.5951677 -64.7889643
O -3.2727482 6.7640869 -65.4867586
C -6.0132403 7.3088003 -63.4796484
H -6.2694267 8.3537053 -63.6622205
H -6.4039576 6.7184002 -64.3096062
C -6.7166646 6.8608593 -62.1861027
H -7.7887787 7.0203155 -62.3018223
H -6.3872767 7.4812947 -61.3521355
C -6.4970237 5.3889815 -61.8459751
O -6.9995508 4.4963847 -62.5114945
N -5.7572628 5.0902401 -60.8042215
H -5.833909 4.14822 -60.4307188
H -5.3482376 5.8097848 -60.2156199
N -3.8569681 8.8981386 -65.0946658

H -4.3202539 9.5439063 -64.4785248
C -3.1401507 9.4583446 -66.2477274
H -3.0411782 8.6833109 -67.0107969
C -1.7236527 9.8698317 -65.8282051
O -1.4618108 10.9868417 -65.364135
C -3.9305454 10.5951731 -66.9132344
H -4.0322571 11.4261952 -66.2131145
H -4.9304066 10.2295566 -67.1538309
C -3.2659619 11.0967388 -68.2087631
O -2.0957811 10.7292587 -68.4770125
O -3.9030893 11.919111 -68.8986723
N -0.803794 8.9111941 -65.9612056
H -1.116004 8.0117478 -66.302686
C 0.6148988 9.1220086 -65.6716635
H 0.701243 9.6227611 -64.7065084
C 1.2656105 10.0741342 -66.6689175
O 2.098196 10.8648421 -66.2441968
C 1.3529267 7.7757301 -65.5926566
H 1.1126767 7.1689474 -66.4679944
H 2.4297305 7.955249 -65.5762918
C 0.9574017 7.0366377 -64.3062496
H 1.2856483 7.6316669 -63.4540904
H -0.1280695 6.9419732 -64.2625537
C 1.5717334 5.633152 -64.2128231
H 2.6407791 5.6991469 -64.4166036
H 1.1128656 4.9710833 -64.9484462
C 1.3819517 5.0770151 -62.7971826
H 1.7874912 5.8138986 -62.1066302
H 1.9538723 4.1524789 -62.6816544
N -0.0351696 4.8430507 -62.4374898
H -0.6226182 5.656999 -62.6128357
H -0.1159741 4.6848988 -61.4413045
H -0.410129 4.0257107 -62.9061462
N 0.857613 10.1083443 -67.9375912
H 0.0306158 9.5949737 -68.2147783
C 1.4627276 11.0419099 -68.8908006
H 2.5495178 10.9536548 -68.8456466
C 1.0955327 12.4971915 -68.5945729
O 1.9570158 13.368256 -68.6695282
C 1.0240044 10.7936445 -70.3246517
H -0.0383867 11.0268319 -70.4208665
H 1.5987591 11.5133366 -70.9013528
C 1.2870139 9.4334093 -70.9591885
H 0.6207275 8.6875652 -70.5204886
H 2.321262 9.1555211 -70.7495981
C 1.0670261 9.5267931 -72.4818624

O 1.5837226 8.6346019 -73.1863309
O 0.449561 10.5278113 -72.9382914
N -0.1651418 12.7909911 -68.2489103
H -0.8538143 12.0316837 -68.2624408
C -0.597046 14.1400191 -67.8500139
H -0.3606733 14.8368522 -68.6493023
C 0.1788132 14.6128788 -66.6331437
O 0.6875716 15.7306213 -66.6664212
C -2.1126775 14.1466946 -67.5941196
H -2.6227997 14.0571914 -68.5530931
H -2.3576696 13.272807 -66.9910062
C -2.6896842 15.3612165 -66.8459386
H -3.7652708 15.2087699 -66.7430743
H -2.2667664 15.3935762 -65.8416272
C -2.4521123 16.7168604 -67.5216241
H -2.901966 16.7154243 -68.5155877
H -1.3820194 16.9074539 -67.6060141
C -3.0951271 17.8042082 -66.6541248
H -2.7560098 17.6658543 -65.6223159
H -4.1813943 17.6762267 -66.6795216
N -2.71647 19.159312 -67.1134731
H -1.700728 19.2804625 -67.0277463
H -3.1123092 19.8764223 -66.5258363
H -2.9676407 19.3212056 -68.075571
N 0.3204954 13.766832 -65.6119406
H -0.1301998 12.8600258 -65.6626692
C 1.1372004 14.0988161 -64.4489276
H 0.7681291 15.0311553 -64.0186157
C 2.6010132 14.3488103 -64.8440332
O 3.1203401 15.4231067 -64.5515805
C 1.0048208 13.0052398 -63.3833035
H -0.0274297 12.9715737 -63.0338871
H 1.2416141 12.0362933 -63.8251743
C 1.9196494 13.2544928 -62.2026704
C 1.5902525 14.2330469 -61.2449909
H 0.6507624 14.7645696 -61.3091331
C 2.5054578 14.5589645 -60.2283914
H 2.2645 15.3361693 -59.5185997
C 3.7477584 13.9046379 -60.163313
H 4.4610224 14.1635952 -59.3939693
C 4.075634 12.9324819 -61.1222156
H 5.0420339 12.4538226 -61.0965527
C 3.1634757 12.6011592 -62.1355357
H 3.4384687 11.8742898 -62.8856969
N 3.2219607 13.4431795 -65.6129092
H 2.7236004 12.5935768 -65.8627977

C 4.6285742 13.5613066 -66.0272132
H 5.2190656 13.6913037 -65.136371
C 4.8554892 14.8103973 -66.9071792
O 5.7538651 15.5982238 -66.6248052
C 5.1469566 12.2423806 -66.6375441
H 4.5264298 12.0049949 -67.503892
C 6.6112232 12.3569321 -67.0848273
H 7.294004 12.1712733 -66.2581363
H 6.8394987 13.3368053 -67.4929083
H 6.78462 11.6171695 -67.8603109
C 5.0988351 11.0728698 -65.6156212
H 5.8917595 11.1882859 -64.8758281
H 4.1709873 11.0890031 -65.0529447
C 5.2309691 9.6913749 -66.2729081
H 6.195366 9.5950969 -66.7695518
H 4.4343643 9.5515472 -67.0036455
H 5.1519131 8.9179254 -65.5089451
N 3.9636351 15.1292163 -67.8520424
H 3.2265652 14.4600239 -68.0547949
C 4.0218868 16.3871815 -68.6226211
H 5.0042765 16.4596209 -69.0901891
C 3.8797531 17.6361744 -67.751329
O 4.3853571 18.6900604 -68.1393617
C 2.9423549 16.3991231 -69.7126516
H 1.9845357 16.1193492 -69.2705717
H 2.8543825 17.4158555 -70.0989016
C 3.2701847 15.4763868 -70.8965743
H 3.2834725 14.434007 -70.5792135
H 4.2600221 15.7274747 -71.2810722
C 2.2462379 15.6651832 -72.0260354
H 2.5661878 15.0769262 -72.8877034
H 2.2537104 16.7118842 -72.3344896
N 0.873631 15.2917612 -71.613203
H 0.3736673 15.9475456 -71.0440993
C 0.2606047 14.1451865 -71.8617678
N 0.8324717 13.2109762 -72.562001
H 0.4215544 12.2750905 -72.7048818
H 1.7817286 13.3170273 -72.8573687
N -0.9394563 13.9204432 -71.4047869
H -1.4254517 14.5628763 -70.815964
H -1.3235025 12.9947997 -71.5411287
N 3.1468849 17.5726836 -66.640555
H 2.7147858 16.6893069 -66.3908726
C 3.0422072 18.6861717 -65.6891141
H 2.9536692 19.6115987 -66.2544648
C 4.3172776 18.8331359 -64.8542892

O 4.818539 19.9522103 -64.7316767
C 1.7891423 18.5422215 -64.8007602
H 1.0555546 17.9044931 -65.2924164
H 2.0612306 18.0606817 -63.8602092
C 1.1195113 19.8957002 -64.5028273
H 1.8473703 20.5638183 -64.0369374
H 0.3048766 19.7346467 -63.7929669
C 0.5491297 20.5310302 -65.7790337
O -0.2282403 19.8629453 -66.4986209
O 0.9457575 21.6510811 -66.1609826
N 4.94132 17.725144 -64.4234338
H 4.4998686 16.8199543 -64.5638902
C 6.2140599 17.8019982 -63.6909489
H 6.0524835 18.5815138 -62.9562327
C 7.3890477 18.3481159 -64.518059
O 8.2732855 18.9822581 -63.9445966
C 6.5604161 16.5360761 -62.8697259
H 7.6036421 16.612572 -62.5826921
C 5.687129 16.5745079 -61.5928064
H 4.6438896 16.3754152 -61.8451636
H 5.7360653 17.5498829 -61.1123979
H 6.0256541 15.83006 -60.8757306
C 6.3611885 15.1622226 -63.5084856
H 5.298497 14.9474787 -63.5301493
H 6.74844 15.1843201 -64.5161972
C 7.0610066 14.0140301 -62.7739185
H 6.6684855 13.8981371 -61.7667884
H 8.122155 14.2216184 -62.7084251
H 6.9110273 13.0840706 -63.3228865
N 7.30301 18.2991724 -65.852446
H 6.594574 17.6978391 -66.2541671
C 8.2430857 18.9945263 -66.7521705
H 9.256973 18.7089679 -66.4899572
C 8.1690656 20.51075 -66.5818082
O 9.1990222 21.1611148 -66.460308
C 7.977772 18.6399987 -68.229651
H 6.9892592 19.0220523 -68.484297
C 8.9845327 19.2914653 -69.1874602
H 10.0014391 18.9979181 -68.9194354
H 8.9126701 20.3778295 -69.1491886
H 8.7886603 18.966429 -70.2109033
C 7.96589 17.1408008 -68.5243393
H 8.943109 16.8146886 -68.8779901
H 7.2357508 16.9438971 -69.3046957
H 7.7171778 16.551408 -67.6508645
N 6.9615765 21.0893303 -66.532496

H 6.1488449 20.486938 -66.5022801
C 6.780615 22.5474602 -66.4135413
H 7.4330732 23.0541204 -67.1286945
C 7.2202667 23.0717325 -65.0569848
O 7.912679 24.0815246 -64.9821131
C 5.3159302 22.9327156 -66.6760503
H 4.6831159 22.049465 -66.5844177
H 4.9780527 23.653042 -65.9286919
C 5.1585947 23.5769347 -68.0589042
H 5.5905341 22.9257093 -68.8152424
H 5.7275276 24.5057287 -68.0837541
C 3.6905468 23.8966561 -68.3952165
H 3.6380314 24.2920585 -69.4108383
H 3.3606373 24.6917945 -67.7218206
N 2.7656957 22.7454938 -68.2527494
H 2.0283473 22.845749 -67.5579178
C 2.88091 21.5274568 -68.7454514
N 3.8693124 21.140862 -69.4991559
H 4.0949378 20.1565639 -69.4957941
H 4.5709032 21.8214323 -69.7161739
N 1.9903342 20.6334323 -68.4586807
H 1.2955849 20.8980357 -67.7519289
H 2.1477213 19.6774308 -68.6872359
N 6.8385406 22.3850171 -63.9793491
H 6.2678414 21.5594362 -64.1188139
C 7.0762139 22.8882716 -62.6154744
H 6.7536171 23.9302454 -62.5989197
C 8.5529194 22.901266 -62.209056
O 8.8936518 23.5075295 -61.1960446
C 6.2302789 22.1490035 -61.5659052
H 6.3002834 22.7158685 -60.6375336
C 4.7406745 22.1370212 -61.9312552
H 4.5608851 21.5072413 -62.7992222
H 4.4104188 23.1524728 -62.155835
H 4.1599278 21.7513496 -61.0951958
C 6.7187884 20.740197 -61.2639461
H 6.946144 20.2365441 -62.1944734
H 5.9697426 20.1820863 -60.7022476
H 7.6299397 20.7966649 -60.6732389
N 9.4330305 22.2598839 -62.9727844
H 9.1190088 21.8290885 -63.831774
C 10.8586508 22.2157185 -62.6987607
H 11.0205601 22.3896924 -61.6385775
C 11.6587618 23.2613323 -63.4943908
O 11.250341 23.686733 -64.571033
C 11.3162717 20.8126336 -63.009183

H 11.2222436 20.673128 -64.0870845
H 10.6948542 20.0962774 -62.4706286
H 12.34619 20.6925253 -62.698318
N 12.8048385 23.6814376 -62.9622398
H 13.0708624 23.3246349 -62.0518115
C 13.7495271 24.5578181 -63.6509041
H 13.1778 25.2862108 -64.2238724
C 14.6719354 23.7794981 -64.6223379
O 14.975169 22.6149417 -64.3663321
C 14.5534374 25.3133525 -62.584314
H 15.1349368 24.6050227 -61.9900518
H 13.8784704 25.8645982 -61.9278724
H 15.2402197 26.0126728 -63.0612602
N 15.2562672 24.4346856 -65.6428965
C 16.3457623 23.8565942 -66.4329207
H 15.9645131 23.0084207 -66.9972777
C 17.5084646 23.3942187 -65.5437926
O 17.9666931 24.1420647 -64.6815619
C 16.7604415 24.9539896 -67.41675
H 17.5396302 25.5804047 -66.9808329
H 17.0916402 24.5356973 -68.368371
C 15.4800201 25.7739039 -67.5748403
H 14.8102338 25.2746548 -68.2775496
H 15.6876092 26.7949285 -67.8961341
C 14.8755827 25.7311526 -66.1737705
H 13.7917232 25.8368128 -66.2352599
H 15.3069372 26.5190763 -65.555338
N 17.9761557 22.1537945 -65.7184936
H 17.5578649 21.5750566 -66.4370437
C 18.9691881 21.5330127 -64.828799
H 19.4528712 20.7173447 -65.3653614
H 19.7268063 22.271958 -64.5676816
C 18.3945524 20.9579177 -63.5231969
O 19.1431088 20.4234825 -62.6929978
N 17.0763422 20.9995032 -63.325819
H 16.4871733 21.5126475 -63.9742723
C 16.4200515 20.282351 -62.2438022
H 16.8992307 20.5608404 -61.3042546
C 16.5430015 18.7673145 -62.4295642
O 16.5979498 18.2344241 -63.5433115
C 14.9511917 20.6876016 -62.1594518
H 14.4759675 20.4721613 -63.1160417
H 14.876111 21.7519627 -61.935609
H 14.4412873 20.1255796 -61.3775577
N 16.4551063 18.0541021 -61.3125826
H 16.2855776 18.5690471 -60.4511779

C 16.3207142 16.6066863 -61.2672556
H 16.667364 16.1436285 -62.1925943
C 14.8430527 16.4876811 -61.1714432
O 14.2703098 16.4918358 -60.0760996
C 17.1025975 16.0055349 -60.0967373
H 16.7267382 16.4254048 -59.1674263
H 16.9332383 14.927969 -60.0696969
C 18.6151246 16.2789271 -60.2035818
H 18.7934009 17.3000794 -60.5397025
H 19.0498173 15.6006874 -60.9384452
C 19.3309473 16.1098317 -58.8681021
O 19.8209138 17.063498 -58.2767724
N 19.4280626 14.9016131 -58.3528361
H 19.9160454 14.8206799 -57.47876
H 19.0269415 14.1074667 -58.8221429
N 14.2900884 16.7928196 -62.3621341
H 14.9082487 16.9578584 -63.1449686
C 12.8660165 16.8921841 -62.5786515
H 12.50313 17.8129391 -62.118628
C 12.3945187 16.8195691 -64.0583579
H 11.5182164 16.1806944 -64.1205188
C 11.927267 18.2080647 -64.4477732
H 12.7140434 18.9289267 -64.2293846
H 11.0102521 18.4103486 -63.8975376
H 11.6871596 18.2879955 -65.5028421
C 13.3973882 16.3601632 -65.1198995
H 13.1242213 16.743578 -66.1014841
H 14.3632553 16.7589904 -64.8585091
C 13.4802412 14.8541873 -65.2635705
H 12.5137973 14.4524802 -65.5470749
H 13.7913259 14.4259101 -64.3196536
H 14.1809111 14.6061911 -66.0551717
C 12.2832975 15.764129 -61.7971147
O 11.2494559 16.0014477 -61.1925983
N 12.9953431 14.6160479 -61.6922139
H 13.8487719 14.4634771 -62.2140222
C 12.4409562 13.5648977 -60.9104669
H 11.9661348 14.0429408 -60.056893
C 13.3950247 12.5287825 -60.3251963
O 14.3898634 12.1321749 -60.9337501
C 11.3641755 12.9163869 -61.7760491
H 11.69877 11.9263678 -62.0871113
C 10.1448767 12.7675339 -60.8818847
H 9.7797381 13.7482455 -60.5753726
H 10.389334 12.200937 -59.985975
H 9.3515675 12.2659959 -61.4236928

C 10.8142608 13.647437 -63.0167973
H 10.2651247 14.5505195 -62.7809551
H 10.1246692 12.9691349 -63.5158578
H 11.6027534 13.8396563 -63.7244018
N 12.992268 12.0624941 -59.1407394
H 12.2290494 12.5696896 -58.7005972
C 13.3517678 10.7991302 -58.4875107
H 13.8691189 10.1454328 -59.1901295
C 12.022295 10.1409558 -58.0722694
O 11.2124517 10.7801555 -57.3905926
C 14.2732964 11.0619765 -57.2714117
H 13.7556098 11.7467959 -56.5983282
C 14.5539489 9.7552096 -56.5040559
H 15.098085 9.0547942 -57.1393494
H 13.6255968 9.2720965 -56.1977078
H 15.1326487 9.9473315 -55.6028041
C 15.5967541 11.7351388 -57.7104154
H 16.1322448 11.0671286 -58.3834904
H 15.3704105 12.6510699 -58.2557262
C 16.5299331 12.1281646 -56.5575202
H 16.9353557 11.2398739 -56.0737671
H 15.9862613 12.7290053 -55.8275692
H 17.3598219 12.7148318 -56.9523203
N 11.7580335 8.9046503 -58.5018071
H 12.4971308 8.4049395 -59.0006782
C 10.4983895 8.1867901 -58.2471085
H 9.9675434 8.6814518 -57.4396159
C 10.7986262 6.7492106 -57.8252037
O 11.102295 5.9443846 -58.7003625
C 9.5576584 8.1569015 -59.488233
H 10.0657489 7.6118162 -60.2850846
C 9.2280089 9.5527545 -60.0531933
H 8.7489912 10.1650888 -59.2892799
H 10.1684049 10.018405 -60.3410172
C 8.2538598 7.3944316 -59.1507915
H 7.707404 7.913199 -58.3643917
H 8.4738782 6.3822778 -58.8197286
H 7.6197814 7.2880813 -60.0276344
C 8.3419087 9.5094014 -61.311817
H 7.3388524 9.1658198 -61.0701747
H 8.7868381 8.8466241 -62.055071
H 8.2449684 10.4990386 -61.7479271
N 10.3778145 6.3647875 -56.6173089
H 10.0867171 7.0875804 -55.969359
C 10.0771669 4.965996 -56.2829504
H 10.6161371 4.3342597 -56.978693

C 8.5735844 4.707958 -56.4325046
O 7.7547345 5.5273144 -56.0055968
C 10.6004787 4.5910992 -54.8968489
H 10.1061673 5.1945444 -54.1366728
H 11.6735419 4.7889482 -54.8607758
O 10.3828632 3.2140938 -54.6418497
H 9.4472056 3.0366627 -54.4338089
N 8.1607001 3.6309068 -57.1055167
H 8.8769937 2.9804449 -57.4273788
C 6.7371884 3.300688 -57.3620578
H 6.1596131 3.5083897 -56.4621738
C 6.5491433 1.8064488 -57.6597888
O 7.4787363 1.1227106 -58.0834072
C 6.1822375 4.201798 -58.4902398
H 6.9381585 4.3099942 -59.2694911
C 4.8723056 3.766805 -59.1432156
H 4.1171477 3.6139869 -58.3752925
H 5.0235349 2.8460121 -59.7068042
H 4.5357146 4.5394803 -59.8340945
O 5.8902408 5.4700017 -57.9372644
H 6.6083844 5.6589023 -57.3062582
N 5.3715908 1.2506011 -57.3572631
H 4.6553821 1.8125038 -56.8983762
C 5.0533527 -0.1448405 -57.6414244
H 5.9175855 -0.7539901 -57.3692877
C 4.7686066 -0.3651727 -59.1348527
O 3.9382459 0.3149866 -59.7426248
C 3.8790949 -0.6020956 -56.7719286
H 2.9852171 -0.0491227 -57.0590648
H 4.0992285 -0.4156114 -55.7198245
H 3.7121333 -1.6687299 -56.9224242
N 5.384449 -1.4132231 -59.6710028
H 5.9615485 -1.978817 -59.0585404
C 5.274073 -1.8707675 -61.0481607
H 4.52063 -1.2819514 -61.5709594
C 4.819635 -3.3368143 -61.111532
O 4.71484 -4.0327033 -60.0985856
C 6.6290324 -1.6468094 -61.7413333
H 6.7247861 -2.2962304 -62.6122492
H 7.4480343 -1.8571985 -61.0513851
S 6.7161264 0.0744788 -62.2988248
H 5.7073454 -0.0156779 -63.1767116
N 4.5804398 -3.8163232 -62.3306187
H 4.6770828 -3.1691214 -63.1064769
C 4.346574 -5.2294842 -62.6551505
H 4.459447 -5.8264837 -61.7510928

C 5.4032849 -5.7429896 -63.6329606
O 6.1897972 -4.9572822 -64.1649364
C 2.9059161 -5.4309908 -63.1662228
H 2.7672556 -6.4849372 -63.4143421
H 2.2061259 -5.1920836 -62.3647088
C 2.5629917 -4.5759187 -64.3962885
H 3.3538419 -4.6507815 -65.1381943
H 2.4774119 -3.5317996 -64.1082129
C 1.239358 -5.0199314 -65.0200982
H 0.4409992 -4.8935195 -64.2851521
H 1.2911458 -6.0804497 -65.2787751
N 0.9258565 -4.2126199 -66.212061
H 0.4081314 -3.3590145 -66.0351975
C 1.2259418 -4.5210722 -67.4631425
N 1.9688723 -5.5468659 -67.7776155
H 2.1152903 -5.8721481 -68.7219343
H 2.3570204 -6.1389378 -67.0480665
N 0.7801215 -3.779182 -68.4391839
H 0.1596895 -3.0162654 -68.2328983
H 0.9754474 -4.0537471 -69.3841574
N 5.3899017 -7.0384504 -63.929528
H 4.7383203 -7.6344439 -63.4363782
C 6.0942102 -7.5596355 -65.1036889
H 7.0977374 -7.1300138 -65.1006469
C 5.4232967 -7.0925274 -66.4226341
O 4.3354509 -6.5106139 -66.4170726
C 6.2677109 -9.0838745 -64.9594063
H 6.6936781 -9.2945219 -63.9792089
H 6.9785677 -9.4402086 -65.7029615
C 4.9912565 -9.885612 -65.1365704
O 4.2025192 -9.6370408 -66.0222303
N 4.7808317 -10.9165378 -64.3629691
H 3.901636 -11.390675 -64.4666077
H 5.4341878 -11.1196392 -63.6117621
N 6.087026 -7.301176 -67.5654737
H 6.97492 -7.7768733 -67.5279906
C 5.5686416 -6.8960675 -68.8824775
H 5.3268371 -5.8354456 -68.847767
C 4.2717675 -7.636028 -69.2757134
O 3.3960169 -7.0361793 -69.9063768
C 6.6966829 -7.1057223 -69.9118019
H 7.5465829 -6.4850685 -69.622889
H 7.0144693 -8.1489414 -69.8602459
C 6.3428838 -6.7968035 -71.3816833
H 5.5274226 -7.4439008 -71.7033055
C 5.9359276 -5.3387014 -71.5982807

H 6.7312629 -4.6712647 -71.263949
H 5.0185384 -5.1252485 -71.0539862
H 5.7487945 -5.1681951 -72.6580208
C 7.5553389 -7.0841982 -72.2684394
H 8.3866897 -6.4341584 -71.9947723
H 7.2949337 -6.9136733 -73.313153
H 7.8578121 -8.1251123 -72.1530906
N 4.1334798 -8.8979656 -68.8636108
H 4.8250579 -9.2749777 -68.2335885
C 3.0282568 -9.7990911 -69.2066128
H 2.994302 -9.8630138 -70.2915907
C 1.6602333 -9.3119172 -68.6927965
O 1.5945623 -8.516802 -67.7502102
C 3.3072741 -11.2092038 -68.6610331
H 2.5364612 -11.8929918 -69.0166306
H 3.2739869 -11.2045757 -67.5732222
O 4.5717364 -11.6737482 -69.1032695
H 4.7435367 -12.5260357 -68.6915725
N 0.5433211 -9.7783737 -69.2905212
C -0.7985269 -9.4881579 -68.7920022
H -0.974344 -8.4156383 -68.878059
C -0.9754439 -9.9212677 -67.3331253
O -0.527133 -10.9988821 -66.9390965
C -1.7659872 -10.2366409 -69.7171588
H -1.976375 -11.2318292 -69.3196156
H -2.693753 -9.6827823 -69.8650191
C 0.4554123 -10.5666366 -70.5124142
H 1.1583618 -10.2258453 -71.2721388
H 0.6242898 -11.6193873 -70.280253
C -0.973233 -10.3730318 -71.0152853
H -1.0393598 -9.4455678 -71.586019
H -1.3115491 -11.2200922 -71.6130074
N -1.6717958 -9.0991233 -66.547554
H -1.9774122 -8.2113845 -66.9104634
C -1.9426282 -9.3712957 -65.1363939
H -0.9869589 -9.4906116 -64.6280691
C -2.7599268 -10.6578647 -64.9656591
O -3.8400503 -10.7898855 -65.5414905
C -2.701309 -8.1901542 -64.5041285
H -2.7718945 -8.3380187 -63.4261289
H -3.7101012 -8.1606276 -64.9151862
O -2.0772418 -6.945159 -64.7763668
H -2.4140318 -6.2669389 -64.1668781
N -2.292544 -11.5662814 -64.1043462
H -1.3627723 -11.44079 -63.7365719
C -3.0207712 -12.7921241 -63.7457168

H -3.2244875 -13.323451 -64.6781217
C -4.3966937 -12.4684447 -63.1343671
O -4.6230048 -11.3825377 -62.579092
C -2.1096583 -13.72106 -62.9038127
H -1.910148 -14.6234283 -63.4834728
H -1.1418491 -13.2284864 -62.8004951
C -2.5148567 -14.1252411 -61.4840284
H -2.7946617 -13.2267165 -60.9280213
H -1.6416138 -14.553518 -60.9864657
C -3.6505368 -15.1492933 -61.4302847
O -4.2452488 -15.22482 -60.332582
O -4.0275463 -15.7087837 -62.4791995
N -5.3275999 -13.4242844 -63.2208885
H -5.0081732 -14.3620026 -63.4746751
C -6.7373853 -13.2558538 -62.8708644
H -7.1275333 -12.4809128 -63.5271843
C -6.946757 -12.7679238 -61.4410492
O -7.9117529 -12.0322895 -61.2615722
C -7.4812086 -14.5788163 -63.1367407
H -7.2969487 -14.8900304 -64.1668939
H -7.0568127 -15.3459934 -62.4846586
C -8.9995942 -14.5392484 -62.9048429
H -9.3885239 -15.5543231 -62.9959171
H -9.1947796 -14.2219061 -61.8805407
C -9.7553918 -13.6445641 -63.9063237
H -10.1326223 -14.2766743 -64.7127237
H -9.0952164 -12.9031723 -64.3565096
C -10.9343519 -12.935928 -63.2296966
H -11.4774207 -13.6713244 -62.6279936
H -11.6147183 -12.563259 -64.0023007
N -10.4659423 -11.8111498 -62.3804404
H -9.7675482 -12.1340372 -61.710333
H -11.2343898 -11.3862407 -61.8810379
H -10.0067443 -11.1094506 -62.9439105
N -6.0816248 -13.0886981 -60.4718269
H -5.3969747 -13.8110332 -60.6825751
C -6.0968226 -12.5397581 -59.1017552
H -7.1172031 -12.2357542 -58.8791371
C -5.2212816 -11.2922417 -58.8815129
O -5.4720267 -10.5585664 -57.9262851
C -5.7549487 -13.6167123 -58.0640669
H -6.3952429 -14.4869167 -58.2181221
H -5.9482444 -13.2182487 -57.0671658
O -4.3992676 -14.0098058 -58.1432093
H -4.343102 -14.5866227 -58.960205
N -4.2784466 -10.9636136 -59.7796263

H -4.191226 -11.5356199 -60.6116894
C -3.5645761 -9.675878 -59.7459851
H -3.2982635 -9.4612025 -58.710529
C -4.4764878 -8.5391038 -60.2170946
O -4.6424323 -7.5619692 -59.492832
C -2.2637066 -9.7208699 -60.5782299
H -1.5860132 -10.4576694 -60.1459091
H -2.5060962 -10.0411425 -61.5913691
C -1.54592 -8.348949 -60.6594275
H -2.2252017 -7.6004366 -61.0640231
C -1.049715 -7.869789 -59.2926931
H -0.3925492 -8.6162577 -58.8481829
H -1.895953 -7.6778135 -58.6345231
H -0.5004847 -6.9352805 -59.4146506
C -0.3461376 -8.4080241 -61.5969519
H 0.3865714 -9.1103831 -61.226032
H 0.1108929 -7.4212003 -61.6794127
H -0.6523785 -8.7230191 -62.5882788
N -5.1169134 -8.689943 -61.3849318
H -4.9457027 -9.5459942 -61.90868
C -6.0412299 -7.6860189 -61.9416269
H -5.4551489 -6.7857532 -62.1437637
C -7.1240799 -7.231914 -60.9375162
O -7.2297424 -6.0346225 -60.7273625
C -6.5934143 -8.1928127 -63.2907024
H -5.7661369 -8.2380254 -63.9984981
H -6.9541082 -9.2147834 -63.16253
C -7.7329938 -7.3712305 -63.9310273
H -7.8971692 -7.7500898 -64.9408023
H -8.6442331 -7.5659483 -63.3636415
C -7.5575872 -5.8442729 -64.0094301
H -7.4980538 -5.4352241 -63.0077227
H -8.4615509 -5.4163366 -64.4482493
C -6.3614685 -5.3390559 -64.8205827
H -6.662531 -5.2804452 -65.8695286
H -5.5195257 -6.0309988 -64.7300999
N -5.9571078 -4.0034871 -64.3263948
H -6.7679802 -3.395136 -64.17414
H -5.2630821 -3.5587778 -64.9038722
H -5.4837897 -4.0957187 -63.4295262
N -7.871645 -8.0931999 -60.2239082
C -8.873936 -7.6616363 -59.2573345
H -9.5234761 -6.9139405 -59.7159941
C -8.2649088 -7.0320422 -57.9992117
O -8.9337415 -6.198365 -57.3998078
C -9.7099499 -8.8997193 -58.9176722

H -10.0869565 -8.8805224 -57.8938592
H -10.5345133 -8.9914458 -59.6264198
C -7.9351771 -9.5147877 -60.3439147
H -8.4774898 -9.7379089 -61.262144
H -6.9490279 -9.9651637 -60.3563997
C -8.720291 -10.0246495 -59.1344312
H -9.2281338 -10.9685741 -59.3297095
H -8.0659944 -10.10908 -58.2666461
N -7.021786 -7.3606245 -57.6043131
H -6.4998668 -8.0322734 -58.1535313
C -6.3056253 -6.6353359 -56.5348897
H -6.9728801 -6.5111878 -55.6824392
C -5.9322881 -5.2270172 -57.0036012
O -6.1602659 -4.2700824 -56.2681912
C -5.0687729 -7.4390979 -56.0860549
H -5.3895788 -8.424192 -55.7418211
H -4.4305065 -7.5948128 -56.9556649
C -4.1992388 -6.7757217 -54.993123
H -3.2115868 -7.2341931 -55.0481208
H -4.0668317 -5.7140503 -55.2046993
C -4.7169933 -6.9550987 -53.5545555
H -4.8133604 -8.0239603 -53.351868
H -5.6999526 -6.4874112 -53.4551922
C -3.7314371 -6.3211879 -52.5553124
H -3.9467867 -5.2535484 -52.4822222
H -2.7197203 -6.4281609 -52.9491867
N -3.7880118 -6.9492504 -51.2117734
H -4.7008678 -6.8646728 -50.7950431
H -3.1294465 -6.5095671 -50.5629086
H -3.5314121 -7.9249485 -51.2652042
N -5.4101965 -5.0928218 -58.2204721
H -5.2513331 -5.9347831 -58.7634563
C -5.1139113 -3.8079675 -58.8626221
H -4.4172266 -3.2441876 -58.2414818
C -6.3827766 -2.9486339 -58.9977917
O -6.424878 -1.8461333 -58.4601641
C -4.4306352 -4.0925634 -60.2147685
H -3.4889227 -4.612916 -60.0251308
H -5.0681263 -4.750292 -60.8032861
C -4.1402352 -2.8329003 -61.0402102
H -3.5223312 -2.1630568 -60.4391233
H -5.0881717 -2.3339607 -61.2560169
C -3.4537445 -3.1310734 -62.3813936
O -3.6879035 -4.2298107 -62.9487496
O -2.7668995 -2.224337 -62.9038172
N -7.4664462 -3.5002571 -59.5499567

H -7.3734285 -4.4240226 -59.9575726
C -8.7116426 -2.7639545 -59.7964255
H -8.4353106 -1.8095062 -60.2480485
C -9.4638687 -2.4089424 -58.5074308
O -9.9528278 -1.288221 -58.3755535
C -9.6118555 -3.5466591 -60.7821039
H -9.0607949 -4.3693492 -61.2330581
H -10.4668038 -3.9705644 -60.2532929
C -10.1111674 -2.6392604 -61.9230753
H -10.7214885 -1.8416841 -61.4932013
H -10.7558622 -3.2309231 -62.5768496
C -8.9667387 -2.0313695 -62.7636765
O -8.2371866 -2.7994259 -63.4391515
O -8.8066446 -0.7847794 -62.7441526
N -9.47042 -3.2962327 -57.4999447
H -9.0417097 -4.20374 -57.6524768
C -10.1010431 -3.0170332 -56.1973756
H -11.0288266 -2.4764864 -56.3973745
C -9.2769901 -2.0761537 -55.3142795
O -9.8498703 -1.4157787 -54.4554675
C -10.4660358 -4.334596 -55.4944689
H -11.0058283 -4.9560219 -56.2112972
H -9.5566922 -4.8544857 -55.1890845
C -11.3704398 -4.1053174 -54.2696322
H -10.7558442 -3.807233 -53.4188719
H -12.0692675 -3.2953097 -54.4855386
C -12.2007088 -5.3376816 -53.8874421
H -12.8839548 -5.5640901 -54.7082912
H -12.7938671 -5.0865768 -53.0059178
C -11.3301939 -6.5636396 -53.5898349
H -10.6293076 -6.3072624 -52.7892185
H -10.7480368 -6.8020651 -54.4859351
N -12.1634946 -7.726805 -53.1936491
H -12.701381 -7.5095402 -52.3633894
H -11.5843428 -8.534113 -52.9993255
H -12.8073944 -7.9650457 -53.9371857
N -7.9716141 -1.9444161 -55.5661454
H -7.5758423 -2.5040646 -56.3110106
C -7.1488087 -0.8533673 -55.0040867
H -7.6222074 -0.5067725 -54.0856447
C -7.1064176 0.3871534 -55.9041949
O -6.4532318 1.363562 -55.5514579
C -5.7327836 -1.3132419 -54.6059369
H -5.1704231 -0.4493004 -54.2512706
C -5.7686133 -2.3392788 -53.4714731
H -6.3200593 -3.2274988 -53.7777969

H -6.2540956 -1.8980821 -52.6008782
H -4.7497511 -2.6174525 -53.2046511
O -5.0121386 -1.9015539 -55.6648057
H -5.5081724 -2.690785 -55.9458878
N -7.8166632 0.3654646 -57.0417343
H -8.3462446 -0.4726061 -57.2427193
C -7.8502309 1.3899778 -58.0923619
H -8.3461036 0.901136 -58.9292334
C -6.4594769 1.7892235 -58.633805
O -6.3001095 2.8344891 -59.2640218
C -8.7526115 2.5574542 -57.6322694
H -9.6187161 2.1305136 -57.1235023
H -8.1999693 3.1478449 -56.8997247
C -9.2803135 3.5032468 -58.7320415
H -8.4579137 4.0436594 -59.1952806
C -10.0659324 2.771923 -59.8264709
H -10.8596408 2.1741863 -59.3761475
H -9.4021933 2.1159166 -60.3858728
H -10.5024011 3.4948423 -60.5147045
C -10.2132689 4.5376623 -58.0986059
H -11.0746174 4.0426197 -57.649767
H -10.5543428 5.2422816 -58.85601
H -9.6730952 5.0878871 -57.3275553
N -5.4426723 0.9434196 -58.4481881
H -5.6678236 0.0355238 -58.0548346
C -4.0798935 1.185912 -58.9357536
H -3.8150141 2.2211407 -58.7113507
C -4.0051982 1.032849 -60.4518383
O -4.7678273 0.27695 -61.0434649
C -3.0949649 0.2568287 -58.2130025
H -3.5138127 -0.7490093 -58.1422002
H -2.1678002 0.2159683 -58.784929
C -2.795052 0.8141 -56.8097441
H -2.3119699 1.7866437 -56.9208895
H -3.7301621 0.9656837 -56.2690902
C -1.8893985 -0.0773955 -55.9502131
H -2.4545609 -0.9480272 -55.6154601
H -1.5950424 0.5033683 -55.0736083
C -0.6439782 -0.5463608 -56.7108241
H -0.3018319 0.268945 -57.3493463
H -0.9215663 -1.3830817 -57.3566641
N 0.4467047 -0.9339892 -55.7916971
H 0.7465728 -0.112789 -55.2498367
H 1.2748543 -1.2105249 -56.2979961
H 0.1611309 -1.6486113 -55.1440047
N -3.0487572 1.7262357 -61.0774219

H -2.4915023 2.3509615 -60.5015827
C -2.7384924 1.6213578 -62.5167704
H -3.2764944 0.767828 -62.9341887
C -1.2710996 1.2723612 -62.7035448
O -0.4066791 2.1504201 -62.6343866
C -3.1835379 2.9095297 -63.2545982
H -3.4195988 3.6894009 -62.5300726
H -2.3764105 3.2816492 -63.8842477
C -4.4157079 2.6815394 -64.1627795
H -4.7723658 3.6569256 -64.4963477
H -4.1117891 2.1108096 -65.0407656
C -5.5713926 1.9356428 -63.4710507
H -5.670715 2.3288555 -62.460506
H -5.3424209 0.8725846 -63.4125411
C -6.9189723 2.0674559 -64.1837241
H -6.9312356 1.427326 -65.0694601
H -7.0421595 3.108668 -64.4916204
N -8.0097774 1.7033435 -63.2510456
H -7.9605382 0.719054 -62.960874
H -8.919042 1.7965361 -63.6756161
H -7.9574248 2.2846703 -62.4272009
N -0.9851324 -0.0152724 -62.8685472
H -1.7507671 -0.6921228 -62.9343551
C 0.3622951 -0.5630426 -62.7563147
H 0.9805693 0.166144 -62.2307181
C 0.9623679 -0.7635467 -64.1569443
O 0.4536773 -1.5034089 -65.0035394
C 0.3662204 -1.8384641 -61.8795023
H -0.0630003 -2.6610733 -62.4548948
C 1.819163 -2.1735927 -61.5191769
H 2.2062048 -1.4454116 -60.8056847
H 2.4410637 -2.152216 -62.4072739
H 1.8767011 -3.1699076 -61.08193
C -0.4305112 -1.6738873 -60.5577608
H 0.0351023 -0.9039866 -59.9410057
H -1.442414 -1.3440835 -60.7786126
C -0.5557201 -2.9669112 -59.7395202
H 0.4082644 -3.2467547 -59.3150915
H -0.9233073 -3.7726035 -60.3764866
H -1.265722 -2.8133331 -58.927048
N 2.0820627 -0.0851478 -64.4029156
H 2.4655832 0.475942 -63.6575331
C 2.9068083 -0.2770199 -65.5962123
H 2.3018658 -0.6952979 -66.4020926
C 4.0396154 -1.2643317 -65.292497
O 4.4307098 -1.4269569 -64.1332412

C 3.4495725 1.0814523 -66.061202
H 4.0958151 0.9525892 -66.9322466
H 4.0432311 1.5292122 -65.261646
S 2.0709612 2.1846743 -66.4876026
H 1.3003227 1.8921115 -65.4342484
N 4.6448883 -1.8495044 -66.3268338
H 4.3466771 -1.6176306 -67.261478
C 5.9932657 -2.3856146 -66.1595621
H 6.0491473 -2.8706147 -65.1886549
C 7.0088041 -1.2347704 -66.1249314
O 6.737833 -0.132612 -66.6086652
C 6.3073177 -3.4524623 -67.2163512
H 5.4957053 -4.1766962 -67.2517385
H 7.2191217 -3.9815768 -66.9375472
C 6.5230252 -2.8364934 -68.5773457
O 7.5678347 -2.280934 -68.8635157
N 5.5086591 -2.8222407 -69.403895
H 5.6807276 -2.3748843 -70.3032807
H 4.6772054 -3.3460722 -69.2191437
N 8.1801732 -1.5131329 -65.5618081
H 8.3470506 -2.4508453 -65.2320084
C 9.2452561 -0.5331003 -65.3748175
H 8.8589516 0.286887 -64.7647094
C 9.6957767 0.098359 -66.7072817
O 9.7789872 1.3222783 -66.8014925
C 10.3873618 -1.2029495 -64.5780293
H 11.2574185 -0.549599 -64.5671677
H 10.0590078 -1.2929866 -63.5414857
C 10.8177317 -2.5865022 -65.0490904
C 11.9181375 -2.7365552 -65.9169673
H 12.4819757 -1.8755661 -66.2390297
C 12.3053206 -4.0136478 -66.363435
H 13.1452796 -4.1187607 -67.0335377
C 11.597087 -5.1547648 -65.9319127
O 11.9592972 -6.3881262 -66.3699191
H 12.7274721 -6.3595775 -66.9421564
C 10.5129368 -5.0118722 -65.040923
H 9.9908157 -5.8930459 -64.6965091
C 10.1282528 -3.7322316 -64.5973171
H 9.308215 -3.6379075 -63.8983316
N 9.914116 -0.6953223 -67.7660937
H 9.7185 -1.6813804 -67.6780474
C 10.4258322 -0.1969978 -69.0526211
H 11.3531195 0.3334907 -68.8684677
C 9.4744436 0.8147866 -69.7068052
O 9.9050969 1.903903 -70.0918449

C 10.7555701 -1.3866897 -69.9704476
H 11.4799712 -2.0264374 -69.4643361
H 9.8468928 -1.968167 -70.1290859
C 11.319076 -0.9912917 -71.3499645
H 10.5886703 -0.3825456 -71.8825983
C 12.633967 -0.2142643 -71.2488822
H 13.3625122 -0.779567 -70.6682336
H 12.462991 0.7532524 -70.7772734
H 13.0319279 -0.0350601 -72.2476385
C 11.5735498 -2.2547509 -72.1719436
H 12.3160219 -2.8816896 -71.6783682
H 11.9344282 -1.9816748 -73.1636924
H 10.6437644 -2.8131246 -72.2827093
N 8.1721243 0.5219023 -69.7300654
H 7.8781855 -0.3931025 -69.4008782
C 7.1627271 1.4694778 -70.216607
H 7.465206 1.8258281 -71.2019721
C 7.0485684 2.7218684 -69.344085
O 6.7977963 3.7974389 -69.8845881
C 5.8037654 0.7721156 -70.3591403
H 5.0142317 1.5249066 -70.3314356
H 5.6424419 0.0910384 -69.5229777
C 5.6722935 0.0300481 -71.6647418
N 5.9532087 -1.3209948 -71.8725283
C 5.7499517 -1.545272 -73.178947
H 5.897551 -2.4993019 -73.6695688
N 5.3500598 -0.4176791 -73.790252
H 5.1471925 -0.3273254 -74.7758818
C 5.2980715 0.5887433 -72.851667
H 5.0341347 1.6258243 -73.0140616
N 7.3164396 2.6332608 -68.0360767
H 7.5599259 1.7314697 -67.6420576
C 7.2614763 3.7968527 -67.1509479
H 6.3620564 4.3393293 -67.4291629
C 8.4261241 4.7738874 -67.3880553
O 8.1695259 5.9696373 -67.5199989
C 7.0948289 3.3363918 -65.6891448
H 6.3288467 2.5601209 -65.6683109
H 8.0271337 2.8928056 -65.3390609
C 6.6681347 4.4549309 -64.7117129
H 7.5189438 5.109165 -64.5258907
C 5.4997377 5.3156355 -65.2074584
H 4.6757884 4.679795 -65.5306189
H 5.8301918 5.9439793 -66.0348046
H 5.1677863 5.9791257 -64.4099764
C 6.2145039 3.8387536 -63.3851094

H 5.3016056 3.2618588 -63.5260214
H 6.0445249 4.6237662 -62.6487135
H 6.9882306 3.1722436 -63.0041928
N 9.6480168 4.2850584 -67.6533181
H 9.7997559 3.2846604 -67.5630426
C 10.7262591 5.1507473 -68.1801541
H 10.8557976 5.9933419 -67.499146
C 10.3952471 5.6900104 -69.5631661
O 10.7418516 6.824877 -69.8713061
C 12.0605472 4.4240518 -68.3016877
H 12.1166467 3.8610945 -69.2350101
H 12.121121 3.7293563 -67.4803064
O 13.1491082 5.3247477 -68.1948499
H 13.269569 5.8510611 -69.0056855
N 9.6793356 4.9112594 -70.3797769
H 9.4739327 3.9631262 -70.084374
C 9.1529553 5.3651188 -71.6612905
H 9.9800387 5.6169965 -72.3253587
H 8.5721268 4.5597261 -72.1096179
C 8.2436502 6.5889815 -71.5206075
O 8.4964877 7.5881455 -72.1829673
N 7.271716 6.5802667 -70.6017955
H 7.1008963 5.7179054 -70.0934582
C 6.3753098 7.7248641 -70.3564864
H 5.913338 8.0175862 -71.2986071
C 7.1064366 8.967594 -69.8279953
O 6.7901523 10.0960769 -70.2162708
C 5.2706065 7.3106618 -69.3736671
H 5.7289851 6.9026424 -68.4718638
H 4.7203521 8.2063148 -69.0832355
C 4.2527546 6.3259864 -69.9274463
C 3.9569052 5.1377987 -69.2312677
H 4.470828 4.90886 -68.3126257
C 2.9868958 4.2463839 -69.7247382
H 2.7656118 3.3367646 -69.1875547
C 2.2932897 4.5473782 -70.9100298
H 1.5351606 3.8746796 -71.2849157
C 2.5749518 5.7375725 -71.6014384
H 2.0297721 5.995317 -72.5016579
C 3.5527959 6.6201892 -71.1146242
H 3.7383022 7.5375435 -71.6559592
N 8.1333138 8.7500029 -69.0098282
H 8.2811018 7.8069615 -68.6678179
C 9.07562 9.7716512 -68.5730746
H 8.5547353 10.5922679 -68.0912188
C 9.8605926 10.3853852 -69.7513712

O 9.7566054 11.5847949 -70.0065046
C 9.9597189 9.1076336 -67.5219712
H 10.8339721 9.7058895 -67.3759088
H 10.2895408 8.1279684 -67.8552867
S 9.0625811 8.9471298 -65.9503501
H 9.8351265 9.8474364 -65.3019252
N 10.4805271 9.5504548 -70.5898632
H 10.5030738 8.5688311 -70.3340113
C 11.2121939 9.9579073 -71.8004392
H 12.0155281 10.6371849 -71.5108146
C 10.3189662 10.6842638 -72.8220458
O 10.6706195 11.7496569 -73.3218699
C 11.8287769 8.7012708 -72.4308687
H 11.0787146 8.1829872 -73.0301747
H 12.1581597 8.0264929 -71.641054
O 12.9535704 9.016732 -73.2287619
H 13.2151123 8.2260534 -73.7115009
N 9.0917743 10.1931924 -73.0341901
H 8.8462209 9.3288767 -72.56349
C 8.0758771 10.7997598 -73.908273
H 8.5230395 10.9933282 -74.8827677
C 7.5601152 12.1572996 -73.4059811
O 6.9709603 12.9057875 -74.1813275
C 6.8980951 9.8177845 -74.0665242
H 6.5963806 9.4797408 -73.0746319
H 6.0469815 10.3468555 -74.4990079
C 7.2010356 8.596359 -74.9559661
H 8.1359891 8.1311667 -74.6495739
C 6.078985 7.5667695 -74.8172612
H 5.1265064 8.0041828 -75.1217947
H 6.002105 7.244648 -73.7790986
H 6.2918725 6.6974123 -75.4373065
C 7.3195826 8.9722824 -76.4354788
H 6.4050387 9.4644876 -76.7684964
H 7.4802548 8.0744735 -77.031481
H 8.1643764 9.6422728 -76.586479
N 7.8083199 12.5056056 -72.1428918
H 8.3267403 11.8644839 -71.5540116
C 7.4700463 13.8201725 -71.5844624
H 6.6249218 14.2248713 -72.140381
C 8.5965015 14.8537852 -71.7283394
O 8.4412966 15.9715581 -71.2569983
C 7.0096966 13.6702722 -70.1392581
H 6.7278152 14.646009 -69.7435907
H 7.8241777 13.2648047 -69.5378654
O 5.8818799 12.8084382 -70.0899695

H 6.1934362 11.8939971 -70.2199382
N 9.6527789 14.5194905 -72.4726668
H 9.6609405 13.583641 -72.8564542
C 10.8450282 15.3307806 -72.7823003
H 11.3215018 14.7681748 -73.5857787
C 11.9223128 15.3449434 -71.6844862
O 12.7208264 16.2703293 -71.5460031
C 10.4885248 16.6983789 -73.4123986
H 10.3052822 17.4216129 -72.6154718
H 9.5630553 16.5886969 -73.9810148
C 11.5483248 17.2610639 -74.3780375
O 12.3958398 16.4764423 -74.8659835
O 11.4371665 18.4657644 -74.7049311
N 11.9834183 14.26542 -70.8998863
H 11.310854 13.5258562 -71.0438426
C 13.0028759 14.0912496 -69.8714545
H 13.306885 15.0728525 -69.5156226
C 14.2829326 13.4339092 -70.3807713
O 14.2769208 12.3465484 -70.963321
C 12.4106606 13.3349922 -68.6923727
H 12.0504773 12.3665639 -69.0383334
H 13.215576 13.1470367 -67.9823596
C 11.3089208 14.0511588 -67.9372688
C 10.4143586 13.2895275 -67.1652494
H 10.4207411 12.2129036 -67.2452246
C 9.622269 13.9020251 -66.183757
H 9.0420531 13.2912381 -65.5123696
C 9.6896914 15.2939169 -66.0117228
O 9.0688287 15.8810816 -64.9659394
H 8.6517885 15.2197967 -64.4135997
C 10.476267 16.0688853 -66.8777947
H 10.5008894 17.1334103 -66.7732363
C 11.2869235 15.4568867 -67.8389113
H 11.9309688 16.084664 -68.4366982
N 15.413457 14.0666489 -70.0648519
H 15.3470006 14.8954175 -69.4865486
C 16.7466287 13.6072225 -70.4479379
H 16.6439033 13.0148765 -71.3578561
C 17.3845581 12.7089696 -69.3861672
O 17.0498046 12.785335 -68.2019845
C 17.6340751 14.8021464 -70.8021027
H 17.0888067 15.4656922 -71.4759625
H 18.5250258 14.4458703 -71.3186039
O 18.0387931 15.5263655 -69.6592734
H 17.3011331 15.5682376 -69.0191956
N 18.3085085 11.8526167 -69.839663

H 18.540374 11.8840546 -70.8182419
C 19.0346307 10.8940314 -69.0057114
H 19.4675616 10.1472293 -69.6713046
C 18.1121144 10.1118048 -68.0476025
O 18.3664568 10.0593789 -66.8463913
C 20.2060694 11.6112569 -68.3167241
H 19.8233584 12.3411857 -67.6012615
H 20.822286 12.1241624 -69.055329
H 20.8190484 10.8836954 -67.7848943
N 17.0082147 9.5547067 -68.5680509
H 16.8205954 9.6340163 -69.5538031
C 16.1258953 8.7216485 -67.7571872
H 16.0455573 9.2076642 -66.7871337
C 16.7726785 7.3640399 -67.4612841
O 17.037384 6.5613184 -68.3551552
C 14.6792575 8.6592675 -68.293561
H 14.2122908 9.6269136 -68.1101076
H 14.1203773 7.931987 -67.704949
C 14.479784 8.3334479 -69.7606206
O 14.1690558 7.2130685 -70.1375173
N 14.5042034 9.3331716 -70.6191546
H 14.1908225 9.1433714 -71.5603483
H 14.5384129 10.2922772 -70.2963576
N 17.0392665 7.1424113 -66.177116
H 16.830673 7.8928146 -65.5251381
C 17.5157217 5.8978433 -65.5954028
H 17.9177789 5.2616605 -66.384763
C 16.3357946 5.207524 -64.9288189
O 15.6159122 5.8225069 -64.1466295
C 18.6212937 6.1737034 -64.5672921
H 18.1711266 6.6156535 -63.6769974
H 19.0702786 5.2249355 -64.2716113
C 19.7093446 7.0913046 -65.0380035
C 19.7931824 8.4044157 -64.7311366
H 19.0720161 8.9404149 -64.124824
N 20.8504143 8.9783652 -65.4059943
H 20.9953366 9.9773354 -65.4322483
C 21.4912532 8.0630289 -66.2111008
C 22.5607112 8.1678796 -67.1126483
H 23.0547657 9.1163029 -67.2628505
C 22.976687 7.0234162 -67.8144749
H 23.7994497 7.0845023 -68.5131482
C 22.3164877 5.7977862 -67.6094385
H 22.6337933 4.9207785 -68.1562422
C 21.2368826 5.7069224 -66.707767
H 20.7248034 4.766601 -66.5741431

C 20.7952953 6.8365602 -65.9837824
N 16.1848563 3.912817 -65.176291
H 16.821799 3.469261 -65.8205527
C 15.0796393 3.1104436 -64.6775838
H 14.54308 3.6333506 -63.8833365
C 15.6247004 1.7985474 -64.1325112
O 16.0577958 0.9402323 -64.9020713
C 14.131694 2.8887033 -65.8487103
H 13.64995 3.8395378 -66.0623428
H 14.7289034 2.6128478 -66.7202805
C 13.0901348 1.8002141 -65.5654381
H 13.4487714 1.0489044 -64.8679001
C 11.8698551 2.4330043 -64.9143906
H 11.3898081 3.1481945 -65.575671
H 12.1699034 2.9421886 -63.9979327
H 11.1508884 1.6662489 -64.6483251
C 12.8433832 1.0456084 -66.8611406
H 12.3756177 1.6885177 -67.5980209
H 12.2143535 0.1981272 -66.6495824
H 13.7858171 0.6635186 -67.2549475
N 15.5022455 1.6123315 -62.8227831
H 15.0176951 2.3212374 -62.2749974
C 15.9819101 0.4156113 -62.1354336
H 16.5767406 -0.1688088 -62.8309248
C 14.7948239 -0.412656 -61.638592
O 14.0877462 0.0314062 -60.7298387
C 16.9228566 0.7713672 -60.9737509
H 16.3487296 1.1324943 -60.119959
C 17.7625409 -0.435787 -60.5611816
H 18.3692173 -0.7740933 -61.4018301
H 17.1180753 -1.2500821 -60.2345302
H 18.4224701 -0.1512396 -59.7425426
O 17.83844 1.7718527 -61.3585353
H 17.3314862 2.5714806 -61.5236629
N 14.5757734 -1.6381667 -62.1522123
C 13.7832369 -2.6243789 -61.4326093
H 12.8078798 -2.2156713 -61.1651937
C 14.5459745 -3.0172483 -60.1610187
O 15.6270337 -3.5983735 -60.2304277
C 13.5987399 -3.7952014 -62.405327
H 13.5382697 -4.7567764 -61.8952256
H 12.6985773 -3.6284725 -62.9974839
C 15.1494966 -2.2202696 -63.3610773
H 14.6833669 -1.7644417 -64.236083
H 16.2312348 -2.0961457 -63.4033807
C 14.823558 -3.7137458 -63.3158808

H 14.6132543 -4.1093829 -64.3096971
H 15.6551597 -4.2543635 -62.8618106
N 13.9662979 -2.746838 -58.9908767
H 13.0661721 -2.2754632 -59.0090904
C 14.5182136 -3.1102436 -57.6789229
H 15.4998097 -3.5591397 -57.8229977
C 13.7014467 -4.2044761 -56.9428997
O 13.6238671 -4.1604091 -55.7146881
C 14.714941 -1.81596 -56.8514598
H 13.7756341 -1.2643591 -56.8431172
H 14.9534945 -2.0737327 -55.819355
C 15.8229594 -0.873091 -57.3541501
H 15.6404761 -0.6146515 -58.3959849
C 15.8120489 0.4127376 -56.5303236
H 16.028931 0.1980959 -55.4826737
H 14.8309305 0.8833728 -56.593059
H 16.5554212 1.1109018 -56.9145674
C 17.2136708 -1.5018124 -57.2217831
H 17.3755269 -1.8389114 -56.1975999
H 17.9750578 -0.7631874 -57.470298
H 17.3151205 -2.3433035 -57.9050428
N 13.0948036 -5.214763 -57.6151661
C 12.2053006 -6.1721577 -56.9514439
H 11.3888186 -5.644905 -56.4617025
C 12.9249789 -7.0386965 -55.9075064
O 12.3195788 -7.4085346 -54.9111168
C 11.6162382 -7.0363963 -58.0682371
H 11.3764525 -8.0460584 -57.7312578
H 10.7270492 -6.5549761 -58.4702904
C 13.2765494 -5.6211828 -59.0043124
H 12.7053484 -4.9577645 -59.6524598
H 14.33053 -5.6213029 -59.2835882
C 12.7120005 -7.0340446 -59.1282169
H 12.3119336 -7.2235629 -60.1252091
H 13.4796364 -7.7662537 -58.8743209
N 14.2193922 -7.3309605 -56.0875591
H 14.6758111 -6.9951267 -56.9187628
C 15.0103271 -8.0990672 -55.112049
H 14.434692 -8.9910839 -54.8655987
C 15.2070223 -7.332927 -53.7964256
O 15.0766997 -7.9116657 -52.717904
C 16.3497863 -8.552893 -55.7413196
H 16.1067111 -9.1238207 -56.6393262
C 17.2457079 -7.3784555 -56.1790683
H 17.6104389 -6.833258 -55.3064419
H 16.7096291 -6.695959 -56.8363059

H 18.1109972 -7.7622999 -56.7213065
C 17.1758962 -9.4774166 -54.8192359
H 18.0864798 -9.7670527 -55.3455807
H 17.4716894 -8.935527 -53.9193684
C 16.4479887 -10.7630839 -54.4053729
H 16.0929498 -11.2919093 -55.2902479
H 15.605773 -10.5327737 -53.7532482
H 17.1389073 -11.4070984 -53.8607548
N 15.3935547 -6.0141343 -53.87863
H 15.4044782 -5.57695 -54.785056
C 15.4989416 -5.1573755 -52.7005961
H 16.1260958 -5.6549509 -51.9609007
C 14.1275042 -4.979588 -52.0493859
O 14.0041547 -5.2410058 -50.8625342
C 16.1529791 -3.8128398 -53.0612849
H 15.5453294 -3.3096024 -53.8136461
H 16.1749842 -3.1849001 -52.1697456
C 17.5888162 -3.9565456 -53.5969701
H 17.9604463 -2.9636573 -53.8583359
H 17.5825816 -4.5500643 -54.513358
C 18.5239075 -4.5941443 -52.560969
O 18.5957494 -5.8428049 -52.4662396
O 19.14147 -3.8633975 -51.7551244
N 13.0718701 -4.7526557 -52.834066
H 13.246795 -4.5814834 -53.8192455
C 11.6674641 -4.7573474 -52.3915481
H 11.5110988 -3.9424476 -51.6838844
C 11.2638044 -6.055375 -51.6705936
O 10.7000723 -5.9916155 -50.5815018
C 10.8225721 -4.4658085 -53.6373247
H 11.2703018 -4.9725609 -54.4847155
H 10.8980451 -3.3996945 -53.8162933
C 9.3400869 -4.8422281 -53.6032018
O 8.5960027 -4.4336681 -52.6834481
O 8.8625046 -5.4577929 -54.5871172
N 11.6917214 -7.2238873 -52.1577427
H 12.1378338 -7.2256039 -53.0701622
C 11.4803921 -8.5084706 -51.4804795
H 10.4192842 -8.5902366 -51.2439924
C 12.2581264 -8.616735 -50.1576347
O 11.7173718 -9.1125472 -49.1729519
C 11.8590517 -9.6523727 -52.4351888
H 12.9441046 -9.6885066 -52.5437256
H 11.4310558 -9.4628766 -53.4176041
C 11.3583046 -11.0199242 -51.9562242
H 11.7482812 -11.2164564 -50.9570363

H 11.7634607 -11.7769659 -52.6281464
S 9.5513028 -11.2041661 -51.9296248
C 9.4280423 -12.9497713 -51.463499
H 8.3802362 -13.2482151 -51.4521366
H 9.8585708 -13.0948903 -50.4720088
H 9.9653893 -13.5614913 -52.1881289
N 13.4992124 -8.1155487 -50.1013757
H 13.8751445 -7.7085546 -50.9531837
C 14.3462607 -8.0992723 -48.8917458
H 14.3261654 -9.0880359 -48.4302966
C 13.8414349 -7.109273 -47.8321094
O 13.8696885 -7.4088854 -46.6438106
C 15.7803139 -7.7771225 -49.3386629
H 16.0728206 -8.4953921 -50.1072518
H 15.78842 -6.7853287 -49.7877521
C 16.8146607 -7.8333564 -48.2019743
H 16.5459291 -7.1366515 -47.4067893
H 16.8280532 -8.8437172 -47.7908348
C 18.2187971 -7.4873246 -48.7193819
H 18.4184365 -8.0829343 -49.6128579
H 18.9536731 -7.7499724 -47.9570584
C 18.3487833 -5.992413 -49.0458139
H 18.4319706 -5.4217815 -48.1165061
H 17.4451803 -5.6561907 -49.5619324
N 19.5031294 -5.7293883 -49.9309239
H 20.3954323 -5.9966135 -49.5549232
H 19.5174901 -4.749167 -50.2346865
H 19.3464276 -6.1609146 -50.848426
N 13.3322727 -5.9630881 -48.2710839
H 13.3916351 -5.7966786 -49.2700395
C 12.6752334 -4.9259791 -47.4652083
H 13.3066394 -4.639705 -46.6204001
C 11.3514425 -5.4659331 -46.907204
O 11.0801258 -5.3692947 -45.708615
C 12.463281 -3.6936637 -48.3766892
H 12.0638674 -4.0566426 -49.3241162
C 11.3939495 -2.735481 -47.8251092
H 11.5699158 -2.5482408 -46.7698325
H 10.4027027 -3.1721763 -47.9439616
H 11.4167651 -1.795219 -48.3641339
C 13.8090055 -3.0011842 -48.7056933
H 14.1943818 -2.5034339 -47.8276267
H 14.5553025 -3.7476582 -48.9678682
C 13.7415405 -1.9940272 -49.8643963
H 13.1388901 -1.128049 -49.5957665
H 13.3172396 -2.4665957 -50.7493734

H 14.7489847 -1.6485579 -50.0979748
N 10.5569209 -6.1251239 -47.7575478
H 10.8058184 -6.1502801 -48.7417588
C 9.3406062 -6.8155937 -47.3499945
H 8.687436 -6.0980664 -46.8515223
C 9.6451199 -7.9320667 -46.3426887
O 8.9789309 -7.9857565 -45.3181334
C 8.6090851 -7.3439192 -48.5900398
H 9.2347282 -8.0646011 -49.1166765
H 8.3743305 -6.5167723 -49.2610325
H 7.6825766 -7.8315087 -48.286617
N 10.6967849 -8.7327052 -46.5459848
H 11.1877227 -8.6812436 -47.4322827
C 11.1410652 -9.7413254 -45.5766499
H 10.3095113 -10.4188421 -45.3765601
C 11.5417736 -9.1181235 -44.2316211
O 11.0027769 -9.5243077 -43.2065835
C 12.2879134 -10.5821685 -46.1574739
H 13.1383181 -9.9410307 -46.3900585
H 11.9539012 -11.054893 -47.0827182
C 12.7241101 -11.6616207 -45.1622198
O 13.5224159 -11.3337599 -44.252894
O 12.2018212 -12.793503 -45.265959
N 12.3829782 -8.0769071 -44.2118859
H 12.8033958 -7.7736289 -45.0842588
C 12.8418208 -7.4756884 -42.9545116
H 13.20788 -8.2795145 -42.315361
C 11.7105569 -6.7843906 -42.179033
O 11.5105007 -7.0762136 -40.995856
C 14.0107534 -6.5158331 -43.2086705
H 14.8199205 -7.0620265 -43.6953675
H 13.6867322 -5.7238326 -43.8863098
C 14.5352889 -5.889822 -41.9505213
C 15.3513898 -6.4920806 -41.0560262
H 15.7532765 -7.4912347 -41.1718495
N 15.5335855 -5.6654725 -39.9629168
H 16.0897329 -5.9158626 -39.160673
C 14.8148192 -4.49376 -40.0872147
C 14.6321954 -3.3866221 -39.242576
H 15.1032791 -3.3500841 -38.2735638
C 13.819259 -2.3236695 -39.676404
H 13.6551516 -1.4597462 -39.0434416
C 13.2231146 -2.3758491 -40.9490175
H 12.61592 -1.5480294 -41.2963996
C 13.413713 -3.4925454 -41.785488
H 12.9690569 -3.4997165 -42.7718436

C 14.1985453 -4.5894945 -41.3712467
N 10.9043925 -5.9527161 -42.8492866
H 11.1134384 -5.7630804 -43.8240366
C 9.7438538 -5.2921988 -42.2212211
H 10.0844229 -4.773692 -41.3238945
C 8.6828352 -6.2978319 -41.7797415
O 8.1475363 -6.1684389 -40.6765001
C 9.0796136 -4.2487583 -43.1304959
H 8.177974 -3.8895772 -42.6371761
C 9.9720763 -3.0467225 -43.4104666
H 10.8669468 -3.3593763 -43.9445606
H 10.2661708 -2.5763767 -42.4714854
H 9.4314004 -2.3171273 -44.0117579
O 8.7181591 -4.8066588 -44.3726664
H 9.5496823 -4.9479732 -44.8577988
N 8.4358032 -7.3581704 -42.5633335
H 8.8963046 -7.4314515 -43.466883
C 7.5723311 -8.4495715 -42.1268535
H 6.6350766 -8.0155664 -41.7795424
C 8.1700975 -9.1517994 -40.9247922
O 7.4758181 -9.174459 -39.9330543
C 7.2448461 -9.4620492 -43.2330206
H 8.1639771 -9.7685181 -43.7320877
H 6.8107539 -10.3548548 -42.779457
C 6.239774 -8.9211213 -44.2597927
H 6.2060887 -9.6067299 -45.1067549
H 6.5649357 -7.9487869 -44.6272582
C 4.8301142 -8.7907822 -43.6962677
O 4.0185217 -9.6936336 -43.7800733
N 4.4373901 -7.6329512 -43.2092787
H 3.4549186 -7.5555601 -42.9669857
H 5.0762198 -6.8618807 -43.140931
N 9.4273277 -9.6006942 -40.9284614
H 9.9686587 -9.4839231 -41.780828
C 10.0214635 -10.412134 -39.8567427
H 9.3084547 -11.1990181 -39.609992
C 10.253828 -9.6601678 -38.5291229
O 10.2250454 -10.2893742 -37.4666527
C 11.2944858 -11.0964365 -40.3945858
H 11.8618845 -11.5134649 -39.5643313
H 11.9244235 -10.3674132 -40.904203
C 10.9694015 -12.2521054 -41.3360884
O 10.0165345 -12.9890257 -41.1245311
N 11.7728694 -12.4937022 -42.343769
H 11.4995019 -13.1733068 -43.0362986
H 12.499172 -11.8467642 -42.6449554

N 10.2929531 -8.3240194 -38.537167
H 10.3624872 -7.8437779 -39.4304161
C 10.2075646 -7.5261957 -37.311539
H 10.9370343 -7.9083139 -36.5955343
C 8.8173012 -7.6078404 -36.6366541
O 8.7224682 -7.6976754 -35.4119872
C 10.5911494 -6.0827475 -37.6571958
H 9.8848154 -5.6694188 -38.3793169
H 11.5923603 -6.0574983 -38.0909011
H 10.5793001 -5.471428 -36.7549126
N 7.7281362 -7.648853 -37.4093658
H 7.8464977 -7.6620426 -38.4175533
C 6.3615845 -7.7065981 -36.8837774
H 6.2525433 -6.8732437 -36.1891368
C 5.9991926 -8.9928891 -36.0905116
O 5.3798625 -8.8602907 -35.0312186
C 5.3875801 -7.4605357 -38.0417105
H 5.317507 -8.3466364 -38.6717451
H 5.7360643 -6.6360806 -38.6607618
H 4.3964641 -7.2320804 -37.651936
N 6.3537059 -10.227742 -36.5201682
C 6.0921003 -11.4546801 -35.7992746
H 5.0595046 -11.4728627 -35.4552056
C 7.0141917 -11.5868196 -34.5825291
O 6.5442037 -12.022288 -33.5387212
C 6.2944393 -12.5878507 -36.8162142
H 6.5796654 -13.5239315 -36.3340225
H 5.3790469 -12.7278175 -37.3944718
C 6.9710996 -10.6108314 -37.7595143
H 6.2311127 -10.4906961 -38.55036
H 7.8621115 -10.0271148 -37.9027458
C 7.3852351 -12.0701274 -37.7139111
H 7.3632158 -12.5333561 -38.7012058
H 8.3603036 -12.1870356 -37.2408137
N 8.2600128 -11.0993172 -34.6473682
H 8.5974122 -10.7351963 -35.529587
C 9.1638845 -11.0324667 -33.493428
H 9.2559856 -12.0264496 -33.0531509
C 8.6246533 -10.1030979 -32.3918366
O 8.5898426 -10.4804776 -31.2234502
C 10.5463395 -10.5906214 -33.9929249
H 10.9400176 -11.3588206 -34.6604347
H 10.4349591 -9.6795107 -34.5821898
C 11.5645544 -10.3336599 -32.8982995
C 12.1676299 -11.4139774 -32.226674
H 11.9007571 -12.4282594 -32.4853161

C 13.115578 -11.1764532 -31.2147571
H 13.5750579 -12.007349 -30.6985902
C 13.464218 -9.8578058 -30.873536
H 14.194 -9.6750549 -30.0974929
C 12.8624226 -8.7769348 -31.541472
H 13.1324783 -7.7636193 -31.281157
C 11.9107974 -9.0133361 -32.5496745
H 11.4482941 -8.1777601 -33.0568941
N 8.0603085 -8.9495652 -32.7631566
H 8.1434311 -8.665565 -33.7337643
C 7.4168677 -8.0211198 -31.8271501
H 7.9214617 -8.1082216 -30.8649975
C 5.9315139 -8.326438 -31.5456683
O 5.2535744 -7.4801421 -30.961817
C 7.5941536 -6.574922 -32.324633
H 7.124499 -6.4941971 -33.3064549
H 7.0538484 -5.9068789 -31.6550215
C 9.0136829 -6.0386909 -32.3987931
C 9.4463363 -5.359517 -33.5561945
H 8.7912065 -5.2787379 -34.4129759
C 10.7433545 -4.8150995 -33.6203321
H 11.075635 -4.3075551 -34.5142454
C 11.6148391 -4.9453172 -32.5180674
O 12.8717441 -4.435554 -32.5709646
H 13.0345853 -3.9454995 -33.3800472
C 11.174962 -5.6029107 -31.3511048
H 11.8453005 -5.6829437 -30.507972
C 9.8769556 -6.144496 -31.2893049
H 9.5563376 -6.6474406 -30.3872537
N 5.3871766 -9.481238 -31.956426
H 6.0033601 -10.1538206 -32.4006634
C 3.9274016 -9.6951489 -32.0875758
H 3.5788743 -9.0167038 -32.8656313
C 3.101727 -9.3795924 -30.8315989
O 1.9946272 -8.8568 -30.9597617
C 3.6184426 -11.1199467 -32.5819641
H 4.1373467 -11.2775613 -33.5237867
C 4.0262775 -12.229422 -31.6076246
H 3.4924745 -12.148523 -30.6645446
H 5.0959923 -12.1726016 -31.405786
H 3.820187 -13.1996915 -32.0601955
O 2.2341245 -11.2757929 -32.8256162
H 2.1031661 -11.0808149 -33.7675107
N 3.6312656 -9.6287895 -29.6257628
H 4.5649006 -10.0097299 -29.5872965
C 2.9405287 -9.3177676 -28.3687039

H 1.9221669 -9.6969241 -28.445873
C 2.8276563 -7.8017421 -28.1507249
O 1.7167741 -7.2833488 -28.1555535
C 3.6362754 -10.0368499 -27.1973246
H 3.6353806 -11.1098669 -27.3948457
H 4.6745595 -9.7054275 -27.1474315
C 2.978396 -9.7839054 -25.8233671
H 2.9966347 -8.7178673 -25.6005677
C 1.5310448 -10.2812004 -25.7697653
H 1.4817257 -11.3377576 -26.0293787
H 0.9096754 -9.6938931 -26.4446131
H 1.1425322 -10.1385847 -24.7604909
C 3.7720642 -10.514862 -24.7416685
H 3.7563083 -11.5903824 -24.9153329
H 3.3368301 -10.2987848 -23.7649057
H 4.8026104 -10.1599394 -24.7388168
N 3.9616382 -7.0915666 -28.1298216
H 4.8305866 -7.5964343 -28.1841824
C 4.0327294 -5.6286734 -27.9990707
H 3.5475118 -5.3373421 -27.066531
C 3.3223149 -4.9052738 -29.1588863
O 2.7887892 -3.8112358 -29.0000078
C 5.522688 -5.2420034 -27.9073454
H 5.9417744 -5.7125461 -27.0159418
H 6.0387796 -5.6508386 -28.77724
C 5.8148372 -3.7290084 -27.839953
H 5.4302001 -3.2507583 -28.740477
C 5.2052008 -3.0369301 -26.6265279
H 5.5270028 -3.5308572 -25.7079944
H 4.1165442 -3.0902357 -26.6670599
H 5.5002705 -1.9890255 -26.5938106
C 7.3263404 -3.5054407 -27.8182893
H 7.7487717 -3.9367071 -26.9075049
H 7.5460048 -2.4366408 -27.8259105
H 7.7977514 -3.9704279 -28.681937
N 3.3240981 -5.4922525 -30.3565642
H 3.8200164 -6.3736162 -30.4481514
C 2.6217824 -4.9657189 -31.5266407
H 2.8835245 -3.9148142 -31.6417372
C 1.0943839 -5.029999 -31.3514762
O 0.4108227 -4.0473604 -31.6319395
C 3.127867 -5.7370606 -32.7582961
H 4.209855 -5.607543 -32.8206349
H 2.9292394 -6.7975108 -32.6056175
C 2.5111026 -5.3318787 -34.1077532
H 1.4532277 -5.5947615 -34.1197899

C 2.6686143 -3.8435188 -34.4097948
H 3.7203921 -3.5715168 -34.3494023
H 2.0853599 -3.2554452 -33.704076
H 2.2981698 -3.6320755 -35.4139741
C 3.226156 -6.095398 -35.2194507
H 4.2560045 -5.7514367 -35.3113525
H 2.7083162 -5.937219 -36.1650524
H 3.2404669 -7.1588567 -34.9842874
N 0.5617721 -6.1321914 -30.8100106
H 1.1854655 -6.8955181 -30.565335
C -0.8774119 -6.2932057 -30.5294087
H -1.4450707 -5.8133694 -31.3273322
C -1.3387857 -5.5989339 -29.2472027
O -2.4972833 -5.2107496 -29.1799714
C -1.2259879 -7.7857952 -30.5059816
H -0.5395652 -8.3161481 -29.8425807
H -2.2423577 -7.9080194 -30.1304602
C -1.1550235 -8.3653732 -31.9235204
H -0.1595575 -8.2283015 -32.3457716
H -1.8592111 -7.825137 -32.5503861
C -1.517416 -9.8499764 -31.9535904
H -1.7789138 -10.1101266 -32.9786297
H -2.3926915 -10.0247935 -31.3275004
N -0.3815104 -10.6888054 -31.5472774
H 0.4686067 -10.5891001 -32.0922565
C -0.2819976 -11.5284697 -30.5397689
N 0.7987728 -12.2399241 -30.4453776
H 0.9213052 -12.9191248 -29.7195572
H 1.4926744 -12.1243647 -31.1788985
N -1.2140364 -11.691179 -29.6413828
H -2.0524106 -11.1461476 -29.7056909
H -1.0837421 -12.3425586 -28.8909044
N -0.4359914 -5.4137744 -28.291648
H 0.4565919 -5.878876 -28.4106519
C -0.6121594 -4.6637386 -27.0405899
H -1.5484464 -4.9448234 -26.5574101
C -0.6419177 -3.1442925 -27.2699944
O -1.4719593 -2.454789 -26.6849815
C 0.5604857 -5.0831334 -26.1452238
H 0.4619065 -6.1502756 -25.9371324
H 1.4675594 -4.941078 -26.7285774
C 0.7664583 -4.3591921 -24.8161269
H -0.1072872 -4.5150072 -24.1788573
H 0.8875321 -3.2879945 -24.9917974
C 2.035639 -4.9226683 -24.1581527
O 1.8929473 -5.6917077 -23.1831504

O 3.1384206 -4.6239782 -24.6732221
N 0.1917402 -2.6286023 -28.1822229
H 0.9142711 -3.2259607 -28.5641261
C 0.1832659 -1.2150978 -28.5658876
H 0.0037262 -0.6137299 -27.6722889
C -0.9246664 -0.8558517 -29.5768898
O -1.3427934 0.2986789 -29.6289793
C 1.5712314 -0.8568396 -29.111109
H 1.7755088 -1.4407739 -30.0082925
H 2.3275326 -1.0683264 -28.3535512
H 1.5991124 0.2052893 -29.3567835
N -1.3690744 -1.8151607 -30.399589
H -0.9570034 -2.7323259 -30.3107115
C -2.4122034 -1.6197109 -31.4130676
H -2.8401389 -0.6237023 -31.2850044
C -3.5838278 -2.5891446 -31.1989809
O -4.4984085 -2.2893246 -30.4387788
C -1.8158924 -1.6685056 -32.833723
H -1.3145361 -2.624073 -32.987596
H -2.63415 -1.6141187 -33.553035
C -0.8409829 -0.5505884 -33.1512774
C -1.3096435 0.649862 -33.7150379
H -2.3647469 0.7833783 -33.9051926
C 0.531843 -0.7025696 -32.8802269
H 0.8864084 -1.6149005 -32.4260483
C -0.4110316 1.6879453 -34.01348
H -0.7806818 2.6123909 -34.4308109
C 1.4300197 0.3388147 -33.1757978
H 2.4760809 0.2347951 -32.9481857
C 0.9602995 1.5313944 -33.7512922
H 1.6466328 2.3356076 -33.9776088
N -3.6205628 -3.7132496 -31.9236746
H -2.8479066 -3.9034345 -32.5436994
C -4.6046085 -4.7861899 -31.7255437
H -4.6477881 -5.0219769 -30.6606922
C -4.2176418 -6.0541322 -32.4919207
O -3.4466607 -6.006089 -33.4557384
C -6.0082382 -4.3507814 -32.1811528
H -6.3304691 -3.4699273 -31.624698
H -6.7175425 -5.152142 -31.9705056
O -6.0325689 -4.0692292 -33.5679786
H -5.899994 -3.1163384 -33.6518697
N -4.8741413 -7.1737827 -32.163381
H -5.4716506 -7.1487112 -31.3494907
C -4.8665038 -8.4037354 -32.9751168
H -3.8421984 -8.7622554 -33.024553

C -5.296619 -8.1459118 -34.4291164
O -4.6251069 -8.5946963 -35.3565651
C -5.7191938 -9.4992887 -32.2851475
H -5.2617698 -9.6962688 -31.3135578
C -7.1749464 -9.0643969 -32.0189038
H -7.7180113 -8.9252308 -32.9539572
H -7.2157387 -8.1483551 -31.4309694
H -7.6857446 -9.8419113 -31.4478146
C -5.7591062 -10.8389807 -33.0521625
H -6.3486961 -11.5529072 -32.4750014
H -6.2623358 -10.6958331 -34.0075932
C -4.3868889 -11.4732058 -33.3065014
H -3.86465 -11.6184485 -32.3614838
H -3.7929475 -10.8453952 -33.9693974
H -4.5241808 -12.4437534 -33.7840225
N -6.3197801 -7.3081212 -34.6459337
H -6.7867577 -6.9174776 -33.8407924
C -6.8108968 -6.9429317 -35.9822531
H -6.9295796 -7.8697976 -36.5438468
C -5.8010042 -6.0946841 -36.7703672
O -5.7534856 -6.2049544 -37.9993011
C -8.1929232 -6.2678168 -35.828368
H -8.8936588 -6.9935093 -35.4113768
H -8.1035006 -5.4462128 -35.1147961
C -8.795135 -5.6932521 -37.1238768
H -9.7559562 -5.2332397 -36.8894435
H -8.1411857 -4.8942535 -37.4703831
C -9.0162358 -6.7421972 -38.2346595
H -9.9843958 -7.2211919 -38.076221
H -8.2596339 -7.5233973 -38.1830747
C -8.9707821 -6.1540308 -39.6519625
H -9.7832162 -5.4324325 -39.7803459
H -9.1312743 -6.9749907 -40.3575952
N -7.6582374 -5.5172426 -39.9201159
H -7.5238449 -4.6979612 -39.3466649
H -7.5232324 -5.260878 -40.8933727
H -6.8930147 -6.1433565 -39.6825251
N -4.9831557 -5.2914209 -36.0868715
H -5.1114733 -5.2316311 -35.0833861
C -3.8621657 -4.5557308 -36.6755995
H -4.2125805 -3.9476157 -37.5101316
H -3.4303927 -3.8987074 -35.9202956
C -2.7652385 -5.4947951 -37.177779
O -2.4814048 -5.5085537 -38.373279
N -2.2677174 -6.3806718 -36.3070245
H -2.5945535 -6.342489 -35.3479413

C -1.2607822 -7.3958332 -36.6512525
H -0.3445124 -6.8925585 -36.961776
C -1.6939054 -8.2984128 -37.8176158
O -0.9311845 -8.4717807 -38.7673839
C -0.9587692 -8.2239316 -35.3945171
H -0.5126998 -7.5591821 -34.6545479
H -1.9003645 -8.5918536 -34.9852958
C -0.0205586 -9.403772 -35.5957312
C -0.476939 -10.7237244 -35.4080804
H -1.509788 -10.9114615 -35.1526382
C 0.4018223 -11.8087914 -35.5828742
H 0.0411055 -12.8215519 -35.4605813
C 1.7396798 -11.5814567 -35.9510346
H 2.4073452 -12.4177367 -36.1126058
C 2.197147 -10.2669613 -36.1448257
H 3.2167878 -10.0866761 -36.4561706
C 1.3158525 -9.1847377 -35.975059
H 1.6594669 -8.1812815 -36.1674815
N -2.9508894 -8.7602365 -37.8162996
H -3.5180262 -8.6013591 -36.9897401
C -3.5164419 -9.5705426 -38.9055008
H -2.8830213 -10.4489155 -39.0412821
C -3.4884375 -8.8191223 -40.2441416
O -3.0854456 -9.4032043 -41.2444796
C -4.9353343 -10.0606912 -38.5229828
H -5.4930771 -9.2156787 -38.1147458
C -5.7199256 -10.6008875 -39.7362459
H -5.1748012 -11.428586 -40.1934969
H -5.8704642 -9.8203191 -40.4808494
H -6.7059161 -10.950001 -39.4339102
C -4.8187313 -11.1576382 -37.4374046
H -4.3660146 -12.0503218 -37.8713733
H -4.1598068 -10.814504 -36.6407699
C -6.1553173 -11.5460725 -36.7923586
H -6.7955629 -12.0580817 -37.5094256
H -6.6616767 -10.6550823 -36.4198857
H -5.9674939 -12.2254155 -35.9610021
N -3.8323925 -7.5248185 -40.2879394
H -4.0656676 -7.0556824 -39.4218744
C -3.7691065 -6.745843 -41.5391746
H -4.3005425 -7.2994218 -42.3139513
C -2.3393446 -6.5519942 -42.0526116
O -2.1145754 -6.6247537 -43.2621331
C -4.4332741 -5.3718712 -41.3703427
H -3.8537605 -4.6109815 -41.8970195
H -4.4733346 -5.1006908 -40.3138927

O -5.7412933 -5.3987764 -41.9212897
H -5.6390313 -5.3843889 -42.8827614
N -1.3861787 -6.3126124 -41.1495095
H -1.6547347 -6.2826085 -40.1714205
C 0.0158965 -6.0425797 -41.481886
H 0.0577829 -5.3112942 -42.2894989
C 0.740751 -7.3012983 -41.9896076
O 1.4825599 -7.2298511 -42.9719905
C 0.6993255 -5.4537121 -40.2332656
H 0.5759614 -6.161702 -39.4119538
H 1.768021 -5.3583632 -40.4270006
C 0.1460188 -4.0790425 -39.7977291
H -0.9410754 -4.0968931 -39.8068985
C 0.5881962 -3.7787531 -38.3651539
H 1.6688825 -3.8436922 -38.2817429
H 0.1358835 -4.4987693 -37.6855946
H 0.2728934 -2.7773283 -38.0815491
C 0.5838749 -2.9467471 -40.7300837
H 1.6679734 -2.8669288 -40.751094
H 0.1714852 -2.0035095 -40.3724266
H 0.2150466 -3.1247907 -41.7383577
N 0.4550266 -8.4611916 -41.3890336
H -0.1379686 -8.4416052 -40.564342
C 0.9646168 -9.7595951 -41.8364496
H 2.0196674 -9.6433663 -42.0892642
C 0.2485841 -10.2596498 -43.1035624
O 0.8987943 -10.7616169 -44.0158858
C 0.8455888 -10.7511466 -40.6638834
H 1.3523783 -10.326376 -39.7955445
H -0.210257 -10.8640917 -40.4110157
C 1.4357562 -12.1477194 -40.9424942
H 0.9147976 -12.6038002 -41.7839374
C 2.9345315 -12.1074372 -41.2461209
H 3.4682867 -11.5756565 -40.4587843
H 3.1048157 -11.6059846 -42.1990644
H 3.3258448 -13.1209115 -41.3293987
C 1.228386 -13.036948 -39.7157139
H 1.7534838 -12.6171902 -38.8578697
H 1.6078131 -14.0384338 -39.9180822
H 0.1647686 -13.1057515 -39.4881301
N -1.0698244 -10.0583188 -43.2209953
H -1.5784126 -9.6705319 -42.4337335
C -1.8288138 -10.4543792 -44.4133578
H -1.6406428 -11.512455 -44.6026739
C -1.3632127 -9.6903915 -45.6534104
O -1.0336879 -10.3147171 -46.6612399

C -3.3368109 -10.2801862 -44.1613091
H -3.5384081 -9.2595103 -43.8397539
H -3.6148275 -10.9493552 -43.3461491
C -4.2354917 -10.6192243 -45.3629775
H -5.2387681 -10.8120393 -44.981554
H -3.8764807 -11.5469091 -45.8110439
S -4.390968 -9.3657168 -46.6781462
C -5.1799099 -8.0079527 -45.7729399
H -5.4383667 -7.2116522 -46.4701612
H -4.4931662 -7.6138861 -45.0242914
H -6.0870682 -8.3705426 -45.2881865
N -1.2836702 -8.3547994 -45.597801
H -1.5351508 -7.8696527 -44.7427603
C -0.9788531 -7.5879528 -46.8043397
H -1.457325 -8.1034868 -47.6360249
C 0.521404 -7.5756896 -47.1451478
O 0.8637811 -7.5102657 -48.3264741
C -1.6430715 -6.2032923 -46.748782
H -1.1355764 -5.5871202 -46.0089988
H -2.6802938 -6.3144597 -46.4385527
C -1.6394516 -5.5151375 -48.1094585
N -0.9709235 -4.389742 -48.2157069
H -0.4685428 -4.0199304 -47.4270163
H -0.9501478 -3.9346266 -49.1103779
O -2.2486604 -5.9590541 -49.0785727
N 1.416904 -7.7516498 -46.1678263
H 1.0980991 -7.8154289 -45.2078644
C 2.8375119 -7.9866019 -46.4363406
H 3.1969886 -7.2710589 -47.1758242
H 3.3911814 -7.8211393 -45.5225839
C 3.1279278 -9.4043741 -46.945777
O 3.8298141 -9.5493483 -47.9453507
N 2.447086 -10.4225332 -46.4101552
H 1.9315207 -10.2624063 -45.5490469
C 2.4854058 -11.7979805 -46.9139969
H 3.5152832 -12.1562071 -46.9087821
H 1.8931677 -12.4278665 -46.2504679
C 1.9239234 -11.9295401 -48.3328251
O 2.5932635 -12.4826701 -49.2022689
N 0.7929469 -11.2777152 -48.628854
H 0.267872 -10.8632915 -47.8641561
C 0.2683398 -11.1228987 -49.9938683
H 0.0573556 -12.1067264 -50.4128921
C 1.2826057 -10.4351154 -50.919821
O 1.4660857 -10.8427349 -52.0678355
C -1.03995 -10.3126457 -49.9511539

H -1.8060414 -10.9034882 -49.4476874
H -0.8743164 -9.4136616 -49.3588149
C -1.5543657 -9.8836418 -51.2950987
C -1.1187229 -8.7991881 -51.9758444
H -0.3563124 -8.1147227 -51.6133052
N -1.6791194 -8.7882528 -53.2363242
H -1.3736528 -8.1625098 -53.9676252
C -2.5438482 -9.8457334 -53.4186151
C -3.3411695 -10.2526051 -54.4992947
H -3.3476281 -9.6931163 -55.422582
C -4.1539985 -11.3894695 -54.358604
H -4.801853 -11.6994674 -55.1695271
C -4.1314936 -12.1172815 -53.1550939
H -4.7589197 -12.9925633 -53.044334
C -3.3101688 -11.7105559 -52.0842815
H -3.3046579 -12.2804389 -51.1652787
C -2.499787 -10.5592805 -52.1831049
N 1.96428 -9.3993704 -50.4253499
H 1.7809878 -9.1156849 -49.4709382
C 2.9636726 -8.654149 -51.2011413
H 2.5137098 -8.3680451 -52.1510126
C 4.1751778 -9.5252683 -51.5380323
O 4.584636 -9.5468988 -52.6991712
C 3.3562456 -7.3613529 -50.4714756
H 3.7076416 -7.5770001 -49.4645344
C 4.4220107 -6.5580398 -51.2039862
H 4.0784879 -6.3762013 -52.2179027
H 5.3645424 -7.1059384 -51.2128172
H 4.574357 -5.6062331 -50.6915057
O 2.2203371 -6.5288334 -50.4111686
H 1.6945377 -6.8265429 -49.6442254
N 4.6517677 -10.3483214 -50.5991615
H 4.2697926 -10.2894075 -49.6593257
C 5.6772822 -11.3580867 -50.8568759
H 6.5451977 -10.8700353 -51.3041686
C 5.1923812 -12.4341222 -51.8490049
O 5.8968642 -12.7354913 -52.8110289
C 6.1067911 -11.9647842 -49.5145566
H 5.2580115 -12.4435631 -49.0255728
H 6.499734 -11.1841373 -48.8617151
H 6.8863492 -12.7084471 -49.6781218
N 3.9534366 -12.9249302 -51.7086818
H 3.4267844 -12.6496469 -50.8836684
C 3.334672 -13.8997941 -52.6291921
H 3.9737587 -14.7791788 -52.6745089
C 3.2311635 -13.3581777 -54.0601941

O 3.501972 -14.0879971 -55.0132972
C 1.9611062 -14.3607064 -52.0920771
H 1.4005589 -13.4924383 -51.7508018
C 1.0987965 -15.094003 -53.1308964
H 1.6582642 -15.9270397 -53.5595551
H 0.8015677 -14.4086898 -53.9226587
H 0.1946828 -15.4764729 -52.6554527
C 2.1541222 -15.3188572 -50.9094302
H 2.6359165 -16.2380737 -51.2437664
H 1.18814 -15.559973 -50.4658729
H 2.7765934 -14.8574841 -50.1444956
N 2.9105554 -12.073782 -54.2549198
H 2.6628376 -11.510579 -53.4444772
C 2.8382215 -11.4848562 -55.603466
H 2.5477263 -12.2731829 -56.294507
C 4.1932573 -11.0523229 -56.1735234
O 4.3050569 -11.0014922 -57.3976465
C 1.7500749 -10.3972881 -55.6684607
H 1.973677 -9.6224873 -54.932472
H 1.7766916 -9.9369756 -56.657886
C 0.3181127 -10.9321047 -55.4242165
H 0.2277581 -11.265099 -54.390261
H -0.3748739 -10.1024204 -55.5677658
C -0.0987512 -12.0894509 -56.3560436
H 0.544746 -12.9544503 -56.17014
H 0.0153415 -11.7777132 -57.39766
C -1.5577073 -12.5187353 -56.1029819
H -2.2269318 -11.6802266 -56.3325773
H -1.6736438 -12.7561101 -55.0366384
N -1.920309 -13.7026939 -56.930816
H -1.8494992 -13.5031153 -57.9228767
H -2.903941 -13.9547134 -56.8042702
H -1.3493329 -14.5134468 -56.71273
N 5.239108 -10.8811623 -55.358782
H 5.0832237 -10.9063656 -54.3572045
C 6.6252368 -10.7490509 -55.8568326
H 6.6136699 -10.1725473 -56.7835268
C 7.23075 -12.111499 -56.2120809
O 7.8163342 -12.237301 -57.2830009
C 7.5248285 -9.9796415 -54.8686851
H 7.4639628 -10.4501565 -53.8877875
C 8.9905939 -9.944826 -55.3227672
H 9.0568977 -9.5997377 -56.3555774
H 9.4372889 -10.9357012 -55.2428839
H 9.5579733 -9.2635562 -54.6896495
C 7.0793244 -8.5186274 -54.7449023

H 7.2214826 -8.0117064 -55.6995769
H 7.6884329 -8.0243809 -53.9867521
H 6.0354079 -8.4688468 -54.4529373
N 7.0278598 -13.1595392 -55.4024952
H 6.5409429 -13.011564 -54.5230944
C 7.6271667 -14.4858677 -55.6559459
H 8.6634613 -14.3031979 -55.9438593
C 6.9906641 -15.2279986 -56.8419336
O 7.6716931 -15.9879257 -57.5225639
C 7.6575082 -15.3317962 -54.3638388
H 7.9124323 -14.6589608 -53.5431137
C 6.2996944 -15.9707711 -54.0400428
H 6.0816508 -16.7851927 -54.7321286
H 5.5262014 -15.2171777 -54.1221082
H 6.2972443 -16.3543042 -53.0201916
C 8.777694 -16.3911983 -54.4438692
H 8.5331507 -17.1303691 -55.2074872
H 9.7068405 -15.8989223 -54.7355244
C 9.0351373 -17.1189259 -53.1182629
H 8.173149 -17.7267722 -52.8436591
H 9.2387219 -16.3944539 -52.3289923
H 9.8999306 -17.7735446 -53.2299662
N 5.7387061 -14.9039989 -57.1868191
H 5.2127234 -14.3222172 -56.5485866
C 5.1062204 -15.3009733 -58.4534771
H 5.3874356 -16.3318692 -58.6680816
C 5.6030433 -14.4874707 -59.672003
O 5.0849158 -14.6599443 -60.773668
C 3.5735904 -15.2270815 -58.2905951
H 3.3168007 -14.2373792 -57.910768
H 3.1057086 -15.3358337 -59.269651
C 2.9775319 -16.3018904 -57.3585176
H 3.4899355 -16.2918303 -56.3990126
C 1.4957103 -15.9962792 -57.1210999
H 0.9610944 -15.9934025 -58.0710052
H 1.4152946 -15.0272253 -56.6348142
H 1.0717324 -16.755584 -56.4636815
C 3.0815974 -17.7101515 -57.947436
H 2.616133 -17.7444842 -58.9324024
H 2.5847047 -18.4241014 -57.2906702
H 4.1275589 -18.0041337 -58.0324987
N 6.5584693 -13.5695965 -59.4840437
H 6.9635998 -13.4805897 -58.559394
C 7.1419561 -12.7294521 -60.533019
H 8.0300305 -12.2428227 -60.1296168
H 7.4428545 -13.3604889 -61.3694312

C 6.2113295 -11.6372743 -61.0664694
O 6.4668133 -11.1007314 -62.1414297
N 5.0989407 -11.3521821 -60.3807085
H 4.9813281 -11.7997437 -59.4822739
C 4.011173 -10.5016823 -60.8819061
H 4.0372055 -10.488733 -61.9692609
C 4.1458654 -9.0360507 -60.4550618
O 4.0311999 -8.1487806 -61.2999311
C 2.6640571 -11.1114873 -60.4638078
H 1.8926477 -10.351034 -60.4842377
H 2.730934 -11.4831747 -59.4414438
C 2.2535011 -12.2657448 -61.3868724
H 1.4591474 -12.8339579 -60.902344
H 3.1069454 -12.932034 -61.5231857
S 1.6498686 -11.7327541 -63.0139356
C 1.6214996 -13.3280329 -63.8756168
H 1.2174771 -13.1904539 -64.8799789
H 0.9970689 -14.0358844 -63.3303468
H 2.6341481 -13.7267153 -63.9450518
N 4.4403578 -8.7747006 -59.1755103
H 4.5741277 -9.5745046 -58.565276
C 4.7721047 -7.4432695 -58.6461105
H 4.2329189 -6.6929209 -59.2287529
C 6.2628089 -7.1513591 -58.8255713
O 7.1132365 -8.0286716 -58.7190631
C 4.3124856 -7.3225627 -57.1776623
H 3.2218766 -7.3201566 -57.1538076
H 4.664482 -8.1940433 -56.6248092
C 4.8413101 -6.0586761 -56.4669405
H 5.9269408 -6.1201353 -56.3940316
H 4.5835999 -5.1738691 -57.0510368
C 4.2767258 -5.8952971 -55.0518431
H 4.2724669 -6.8599056 -54.5417714
H 3.2469937 -5.5465683 -55.1283597
C 5.0870934 -4.8828455 -54.2236432
H 4.4787623 -4.580427 -53.3658758
H 5.2958632 -3.991815 -54.8235343
N 6.3573003 -5.466491 -53.7188249
H 6.2395348 -6.3113772 -53.1868018
H 6.9070721 -4.8226841 -53.1439003
H 7.0461657 -5.6510471 -54.4521823
N 6.5831059 -5.8796803 -59.0166479
H 5.8235267 -5.2105063 -59.113986
C 7.9375657 -5.3227778 -59.0310328
H 8.6143732 -5.9856498 -58.4939189
C 7.9299697 -3.9753104 -58.3016906

O 6.8849011 -3.3299543 -58.2056804
C 8.3940536 -5.2114097 -60.498702
H 7.5989185 -4.7461053 -61.083704
C 9.6867695 -4.4393521 -60.7534294
H 10.461601 -4.7634189 -60.0624864
H 9.5092652 -3.3712309 -60.637248
H 10.0261984 -4.6212524 -61.7728531
O 8.6164189 -6.5095718 -60.9963761
H 8.1255677 -7.1366062 -60.4489059
N 9.0868513 -3.5097747 -57.8288001
H 9.9133854 -4.0836863 -57.9019445
C 9.2842826 -2.0970461 -57.484942
H 8.3344924 -1.5820776 -57.6283965
C 10.2397767 -1.4588863 -58.488381
O 11.1822076 -2.0878065 -58.9748673
C 9.6723067 -1.8988096 -55.9998239
H 9.4745394 -2.8358148 -55.4851521
C 11.1589555 -1.5434698 -55.8111287
H 11.3819425 -0.5661663 -56.2441685
H 11.7740098 -2.3086833 -56.2769605
H 11.4226248 -1.4985324 -54.7557414
C 8.7636256 -0.8305709 -55.3496209
H 8.9602225 0.1445593 -55.7991688
H 7.7226656 -1.0931424 -55.5430241
C 8.9349449 -0.7232145 -53.8274753
H 9.9006709 -0.2819557 -53.5826193
H 8.8562241 -1.7099889 -53.3708842
H 8.1532925 -0.0869728 -53.4164581
N 9.9517901 -0.2103244 -58.8094482
H 9.1682783 0.231535 -58.3373574
C 10.654951 0.6360141 -59.7608788
H 11.4393476 0.0802286 -60.2761536
C 11.3137825 1.7707113 -58.9834636
O 10.6744127 2.3280577 -58.0948429
C 9.5994166 1.0937026 -60.7929687
H 8.6089785 1.028997 -60.3467226
H 9.6158493 0.3827572 -61.6188321
C 9.6756245 2.4856417 -61.3678588
N 9.6560584 2.7683857 -62.7304314
C 9.6304442 4.1041051 -62.8320208
H 9.6677495 4.647484 -63.7669766
N 9.599696 4.6686446 -61.6145544
H 9.6754936 5.6576488 -61.4159251
C 9.6037052 3.6646771 -60.6808175
H 9.5692624 3.7963986 -59.6106405
N 12.5267095 2.1509732 -59.3842939

H 13.0058659 1.5797279 -60.0714761
C 13.1036285 3.4786336 -59.1419166
H 12.3791542 4.0801703 -58.6065819
C 13.4003518 4.137936 -60.4980748
O 13.9804095 3.4895145 -61.3751488
C 14.3486196 3.3947989 -58.2404563
H 14.0355429 3.0345998 -57.2602812
H 15.0443904 2.6671256 -58.661884
C 15.0998792 4.7290328 -58.0561606
H 15.5972644 4.9762276 -58.9978091
H 15.8861092 4.5829474 -57.3125827
C 14.2028056 5.9034645 -57.6227446
O 14.1451209 6.8963968 -58.3833069
O 13.5576779 5.8054743 -56.5593021
N 13.0453446 5.4126846 -60.6712948
H 12.6425479 5.9066621 -59.8759317
C 13.3632583 6.2062371 -61.8533931
H 14.1139036 5.6662568 -62.4167197
C 13.9496037 7.5757594 -61.5001384
O 13.3110616 8.3851175 -60.8299668
C 12.1357501 6.3519771 -62.752082
H 11.3121412 6.7960327 -62.1946084
H 11.8710934 5.3727228 -63.1411174
H 12.3807769 6.9994993 -63.5956822
N 15.109189 7.8745898 -62.0873356
H 15.4773862 7.2053572 -62.7563538
C 15.8432781 9.1316037 -61.9126432
H 15.2448618 9.8123746 -61.30726
C 16.0561624 9.7692783 -63.2824116
O 16.4188304 9.0839867 -64.2349318
C 17.1881392 8.8969502 -61.188147
H 17.8076121 8.2561194 -61.8169794
C 16.9764192 8.1765845 -59.8366834
H 16.3622953 8.7964619 -59.1838709
H 16.4432474 7.2414061 -60.0042166
C 17.9261826 10.2389898 -60.9959842
H 17.3004281 10.929227 -60.4298202
H 18.1645092 10.6855659 -61.9607414
H 18.8686102 10.0894265 -60.4738383
C 18.2749524 7.8099776 -59.110778
H 18.7708445 8.7035771 -58.7336403
H 18.9412565 7.2714474 -59.7855677
H 18.0370505 7.1684199 -58.2616609
N 15.8801458 11.0823937 -63.3805037
H 15.5331748 11.5872138 -62.5688742
C 15.9623305 11.817571 -64.6442913

H 16.7952455 11.4316431 -65.2351664
C 16.2306405 13.3158319 -64.4223685
O 16.001996 13.8407697 -63.3256233
C 14.6780471 11.5721577 -65.4405948
H 14.5614297 12.3592306 -66.1828294
H 14.8143697 10.6397859 -65.9771394
C 13.3981278 11.4233831 -64.5967226
H 13.2409175 12.3213791 -64.0054631
H 13.4843556 10.5777558 -63.912386
C 12.2010246 11.1445692 -65.486179
O 12.4220538 10.9836834 -66.7062945
O 11.0711677 11.0577901 -64.952999
N 16.7127389 14.0282412 -65.4533237
H 16.8577185 13.566903 -66.349382
C 16.9273486 15.4814323 -65.3898288
H 16.3871383 15.8476574 -64.5193038
C 16.3447654 16.2645858 -66.5817988
O 16.1558116 15.7419953 -67.6835179
C 18.4109936 15.8096423 -65.1287159
H 18.8261429 15.0930979 -64.4212154
H 18.4804463 16.7986531 -64.677849
C 19.2719235 15.817902 -66.3805842
O 19.9812442 14.8726451 -66.670495
N 19.2629312 16.8803734 -67.1552989
H 19.7115882 16.7659892 -68.0471681
H 18.6938739 17.6945154 -66.9492704
N 16.1171737 17.5560555 -66.3461034
H 16.3244087 17.9218972 -65.4216912
C 15.6409335 18.5213459 -67.3311437
H 15.1563779 17.9880668 -68.1515813
C 16.8207639 19.3019001 -67.9050657
O 17.814395 19.5196466 -67.2156087
C 14.6025547 19.470826 -66.7156745
H 15.0457816 19.9951369 -65.8663646
H 13.7532703 18.8943344 -66.3687152
C 14.1049765 20.4776596 -67.7549781
O 14.0497 21.6808959 -67.4357038
O 13.8939207 20.0301459 -68.9036003
N 16.6932574 19.7127438 -69.1587056
H 15.7361606 19.6535177 -69.5221682
C 17.6417525 20.4891035 -69.9664808
H 18.1709276 19.7718774 -70.5775385
C 18.7397635 21.2279062 -69.1911158
O 19.6163392 20.5303848 -68.6231238
O 18.7905354 22.4280729 -69.0989548
C 16.8603575 21.4153103 -70.9236658

H 16.3055296 22.1240006 -70.3041713
H 17.5800612 21.9809911 -71.516076
C 15.8643249 20.7213124 -71.8783228
H 15.1037594 20.1933291 -71.2956912
C 15.1458514 21.7861056 -72.7164382
H 15.8579213 22.3362843 -73.3312233
H 14.6219113 22.4788679 -72.0561909
H 14.4058411 21.3071409 -73.3609419
C 16.5435476 19.7199673 -72.8232848
H 17.329409 20.2094041 -73.398468
H 15.8002973 19.3095617 -73.5106352
H 16.9602655 18.8885803 -72.2557724

**Supplementary Data 4. Cartesian coordinates for CrPeNMT homology model with bound SAM.**

N -2.1385667 21.7753992 -15.9844882
H -2.8881317 22.3451992 -15.612073
H -2.4187043 21.4412796 -16.898962
H -1.952445 20.9940613 -15.3721405
C -0.8983927 22.6008534 -16.1048954
H -1.0805973 23.3575681 -16.8709468
C 0.33849 21.7897622 -16.5659561
O 1.2553163 22.327476 -17.1874333
C -0.6053818 23.3116885 -14.7771474
H 0.3668547 23.80077 -14.8479421
H -0.5567763 22.5711725 -13.9778747
C -1.639947 24.3835703 -14.4160168
H -1.396285 24.7758699 -13.426708
H -2.6273748 23.9290305 -14.3587376
S -1.7285502 25.7740286 -15.5733039
C -2.9534829 26.7844932 -14.6928753
H -3.1342375 27.7045542 -15.2488035
H -2.5798462 27.0332981 -13.6984196
H -3.8891334 26.2323095 -14.5953778
N 0.3673335 20.4886197 -16.2657559
H -0.4109708 20.1080689 -15.7405706
C 1.3938537 19.5859753 -16.7771625
H 1.3580658 19.6103407 -17.8672226
H 2.3753087 19.9422459 -16.4680576
C 1.2496557 18.1385969 -16.3224611
O 0.7986647 17.8537127 -15.2154758
N 1.7628938 17.2390447 -17.1522057
H 2.1121366 17.5432004 -18.0501647
C 2.0288584 15.8382482 -16.8338954
H 1.9028599 15.7027836 -15.7604402
C 3.4957816 15.5076051 -17.1394012
O 4.119702 16.1812749 -17.9609391
C 0.9990926 14.9223526 -17.5032517
H 1.2572628 13.8928438 -17.2749979
H 0.0423459 15.1358366 -17.026009
C 0.8149397 15.0455348 -19.020348
H 1.6157817 14.5050359 -19.5288645
H 0.8699863 16.0954136 -19.3176114
C -0.5554566 14.4754363 -19.4180847
O -1.297864 15.2053148 -20.1119062
O -0.875195 13.3478418 -18.9747065
N 4.1061184 14.7016468 -16.2632362
H 3.5442127 14.1102083 -15.6715657

C 5.5629684 14.6122266 -16.0983634
H 6.0446756 14.8186955 -17.0600251
C 5.9695982 13.2129132 -15.6736854
O 5.3463653 12.639753 -14.786269
C 6.0594875 15.5900786 -15.0182406
H 5.5966651 15.3285324 -14.0641537
H 7.1389711 15.4636292 -14.915048
C 5.7688522 17.0627619 -15.3210483
H 6.199365 17.3109927 -16.2919241
H 4.6914859 17.2284646 -15.351451
C 6.3694499 17.9615862 -14.2310701
H 7.4516987 17.8220036 -14.192695
H 5.9514537 17.6743696 -13.2638997
C 6.0413128 19.4366002 -14.4861744
H 6.4275873 20.029751 -13.6516003
H 4.9522862 19.5499365 -14.5026758
N 6.6253332 19.9107507 -15.7656037
H 7.6331593 19.818804 -15.7546612
H 6.3843249 20.8792368 -15.933991
H 6.2683797 19.3550584 -16.5356419
N 7.106455 12.7593095 -16.1705568
H 7.4956956 13.2759729 -16.9574346
C 7.5639831 11.383838 -16.027934
H 6.7220861 10.7521143 -15.7410336
C 8.6256294 11.3080201 -14.9245199
O 9.6775993 11.9537076 -14.9983873
C 8.0743326 10.8706348 -17.3888252
H 8.9570019 11.4416081 -17.6791626
H 8.3787829 9.8302572 -17.2664366
C 7.0169986 10.9370419 -18.5172246
H 6.1405201 10.371064 -18.1914962
H 7.409044 10.4393792 -19.4038394
C 6.5974706 12.3719046 -18.896483
O 7.4785217 13.2681533 -18.9084406
O 5.3745041 12.6072441 -18.9936412
N 8.3366279 10.5700175 -13.8495991
H 7.428984 10.1185224 -13.7997984
C 9.2537472 10.4628666 -12.7165948
H 9.411715 11.4704627 -12.3299929
C 10.6243327 9.9036971 -13.153139
O 10.6986872 8.9385328 -13.9069108
C 8.6037136 9.6350831 -11.6036772
H 8.4204313 8.619799 -11.9600852
H 7.655759 10.0896643 -11.3114457
H 9.2647124 9.595986 -10.737355
N 11.703524 10.5367378 -12.6721267

H 11.5296608 11.3030954 -12.0418231
C 13.1142625 10.3142897 -13.0702562
H 13.6516325 11.146085 -12.6173586
C 13.4243279 10.5119185 -14.5675031
O 14.4681525 10.0599575 -15.0339048
C 13.7765064 9.0619071 -12.4395625
H 14.845682 9.122573 -12.6445486
C 13.6265139 9.0675566 -10.9118445
H 12.5803642 8.9423089 -10.6318339
H 14.0018322 10.0070241 -10.506304
H 14.2061243 8.2486758 -10.485005
C 13.2962742 7.7038875 -12.9652798
H 12.2890587 7.4911381 -12.6080123
H 13.9637118 6.9192992 -12.6112727
H 13.2950871 7.7094786 -14.0550769
N 12.605294 11.3349009 -15.2378421
H 11.7369915 11.5814565 -14.7805248
C 12.7314974 11.833814 -16.6111805
H 11.8724211 12.4921825 -16.7495073
C 12.6154246 10.7757957 -17.7237248
O 13.1625965 9.6728211 -17.644985
C 13.9716254 12.728998 -16.7542726
H 14.876663 12.1276992 -16.6566322
H 13.9650252 13.5017634 -15.9855672
H 13.9695782 13.2008787 -17.7373648
N 11.9102878 11.1457732 -18.7984009
H 11.4901366 12.0606529 -18.8311174
C 11.6113386 10.2161504 -19.8849453
H 11.4683386 9.2575284 -19.3922061
C 12.7697246 10.041093 -20.8749029
O 13.4841971 10.9883786 -21.2103005
C 10.2956217 10.5834444 -20.6018401
H 9.5642602 10.8737195 -19.8541547
H 10.4613373 11.4376453 -21.2598264
C 9.7172444 9.4047201 -21.4196476
H 10.350257 9.2350742 -22.2929309
H 8.7362979 9.7002982 -21.7986705
C 9.5899082 8.089336 -20.6183328
O 8.4815808 7.7830257 -20.1380566
O 10.6205886 7.3866881 -20.4498584
N 12.8901662 8.8305339 -21.4204041
H 12.2017952 8.1247666 -21.1470783
C 13.7998552 8.4972353 -22.5092701
H 14.1771403 9.4236569 -22.9434863
C 13.0399482 7.741936 -23.6088019
O 12.2666108 6.8259826 -23.3460136

C 14.9936922 7.7225667 -21.9245386
H 15.5173435 8.3745069 -21.2235556
H 14.6001376 6.8863099 -21.3570609
C 16.0029798 7.189281 -22.9591753
H 15.4878043 6.568367 -23.6890142
C 16.7306364 8.3210255 -23.6896215
H 17.2504768 8.9574053 -22.9726436
H 16.0235264 8.9250527 -24.2541046
H 17.4549132 7.90003 -24.3877077
C 17.0509741 6.3236455 -22.2584349
H 17.5894864 6.9110609 -21.5149428
H 17.7563171 5.9395483 -22.9960493
H 16.5630007 5.4766885 -21.778768
N 13.2962296 8.1160457 -24.8611726
H 13.9627613 8.8548464 -25.0049569
C 12.7297022 7.4720004 -26.0467334
H 11.6552424 7.6561114 -26.0668372
C 12.963355 5.9483119 -26.0268958
O 14.0825531 5.4964434 -25.7932219
C 13.3669937 8.1475173 -27.2671617
H 13.1403425 9.2139322 -27.2266732
H 14.4513269 8.049724 -27.187207
C 12.9656811 7.6189319 -28.6307211
C 12.1494424 8.3971049 -29.4761888
H 11.769383 9.3526089 -29.1454154
C 11.8565891 7.9520163 -30.7798108
H 11.2562024 8.555452 -31.4472005
C 12.3673248 6.7178848 -31.2374061
O 12.1160781 6.2857259 -32.4969706
H 11.5991475 6.9488647 -33.0185918
C 13.1697677 5.9340538 -30.3847744
H 13.5492799 4.9896701 -30.7459872
C 13.4827043 6.3907892 -29.091865
H 14.1273567 5.791667 -28.4621886
N 11.8951892 5.1788313 -26.2599184
H 11.0304658 5.6427895 -26.4800882
C 11.8426092 3.7036502 -26.2137578
H 10.7811114 3.4599108 -26.2631227
C 12.3180866 3.0493723 -24.893297
O 12.588105 1.8486882 -24.8229072
C 12.4769853 3.1028593 -27.4818833
H 13.5610504 3.0598399 -27.3602348
H 12.2582179 3.7523311 -28.3311342
C 11.9188234 1.7098089 -27.8129528
O 10.6734795 1.5608478 -27.7641107
O 12.7303627 0.8270589 -28.169355

N 12.3786583 3.8077385 -23.7892188
H 12.1567416 4.7960764 -23.8565606
C 12.7236866 3.2518266 -22.4780661
H 13.5786598 2.5902866 -22.6181944
C 11.5733328 2.4149249 -21.9209309
O 10.5345585 2.9233827 -21.5059758
C 13.1586237 4.3717878 -21.5322934
H 14.0067916 4.8753226 -21.9927235
H 12.3398627 5.0838206 -21.4074609
C 13.5708913 3.8411374 -20.1519667
H 12.6903613 3.4453154 -19.6452956
H 14.3110593 3.0473288 -20.2626016
C 14.1595742 4.9730826 -19.3074994
H 13.477531 5.8273082 -19.3386531
H 15.1255757 5.265841 -19.7198963
C 14.3246592 4.5251755 -17.8558798
H 14.9647772 3.6393135 -17.8237746
H 13.3343658 4.2459127 -17.4769636
N 14.894404 5.6206571 -17.034387
H 15.8026652 5.9022239 -17.3743342
H 14.9585333 5.3479973 -16.0632593
H 14.281514 6.4323719 -17.0835563
N 11.8145313 1.1129027 -21.796752
H 12.636759 0.7428888 -22.2469702
C 10.915173 0.206206 -21.075752
H 9.9261211 0.3116122 -21.5233934
C 10.7879049 0.6022152 -19.5986625
O 11.7565775 1.0128309 -18.9601126
C 11.3345322 -1.270057 -21.22834
H 10.6296722 -1.8783407 -20.6606041
C 11.2418011 -1.7098514 -22.6941182
H 11.9608367 -1.1650701 -23.3087444
H 10.2395993 -1.5174528 -23.0780593
H 11.4468521 -2.7773516 -22.7753339
C 12.7458793 -1.5768076 -20.7095885
H 13.4907476 -1.0077613 -21.2656222
H 12.9570051 -2.6395233 -20.8284199
H 12.8214824 -1.3296274 -19.6506744
N 9.6090573 0.3641778 -19.0190877
H 8.8636369 0.042936 -19.61778
C 9.3030641 0.4995601 -17.5765515
H 8.2498576 0.2378134 -17.4888618
C 9.3464409 1.9102799 -16.9589934
O 9.2888891 2.018324 -15.7386553
C 10.0341763 -0.545724 -16.7016046
H 9.6693134 -0.4676256 -15.6770594

C 9.7794235 -1.9797159 -17.169852
H 10.215899 -2.1450359 -18.1540049
H 8.7079161 -2.1740235 -17.2032576
H 10.239578 -2.6699839 -16.4624216
O 11.4299275 -0.3613229 -16.691399
H 11.6667592 0.1678767 -17.4727227
N 9.2779939 2.9909397 -17.7450031
H 9.3809 2.8980378 -18.7455964
C 8.8924366 4.3186524 -17.2226461
H 9.484174 4.5459203 -16.3361664
C 7.3950813 4.3363773 -16.8420819
O 6.54833 4.1563548 -17.7157283
C 9.1737252 5.4060425 -18.2724269
H 8.8412563 6.3707971 -17.8859208
H 8.6096819 5.1895781 -19.1816057
O 10.5517975 5.4957348 -18.5958262
H 10.5894134 6.1754387 -19.3178923
N 7.0367162 4.4791013 -15.5515566
H 7.7637049 4.4744686 -14.8485576
C 5.6335974 4.372326 -15.0850094
H 5.0070584 4.6318068 -15.9400818
C 5.0839267 5.3356751 -13.9496445
O 4.1802507 4.8921545 -13.2441249
C 5.3501324 2.8742188 -14.7954481
H 5.6601937 2.2734319 -15.6495461
H 4.2770818 2.7385852 -14.6680957
C 6.0481724 2.3356799 -13.5563096
O 6.9355157 2.9397106 -12.9802825
N 5.6839717 1.1536912 -13.1130988
H 6.1721809 0.8278604 -12.2984982
H 4.9434468 0.6468731 -13.5561149
N 5.4209234 6.6581869 -13.7954432
H 6.1313111 7.0377768 -14.4037847
C 4.6727308 7.6690857 -12.8730958
H 3.6722803 7.2716519 -12.7031805
H 5.1665915 7.6425688 -11.902207
C 4.4605241 9.2667223 -13.2399755
O 5.4811072 9.6663177 -13.812121
N 3.2536409 10.1109433 -12.9872607
H 2.6474233 9.6614132 -12.3206553
C 2.3771526 11.4343133 -13.6922496
H 3.1658733 12.0715482 -14.0792353
C 1.0505375 12.8198719 -12.978889
O 0.738086 12.1140384 -12.0071596
C 1.9540762 10.4352165 -14.9544058
H 0.9329988 10.1097298 -14.746017

C 2.7417697 9.1177858 -15.5124699
H 3.7821969 9.3932306 -15.6797871
H 2.6711718 8.3582525 -14.7352443
C 1.9181844 11.302737 -16.1856385
H 2.9290002 11.590086 -16.4899791
H 1.3394983 12.1952761 -15.9637221
H 1.4171187 10.8031861 -17.0150758
C 2.3461044 8.3062146 -16.7841532
H 1.2838958 8.043668 -16.7411571
H 2.9411495 7.3919643 -16.8207085
H 2.5328117 8.8854332 -17.695951
N 0.2333932 14.4003043 -13.1834479
H 0.7437184 14.7653437 -13.9704046
C -1.2123799 15.955379 -12.9597387
H -1.0028175 16.3292374 -13.960482
C -2.5312399 15.1695335 -13.1010222
H -2.3258053 14.1092915 -13.2484796
H -3.0618878 15.2689053 -12.1555772
C -3.4175086 15.6977339 -14.2585728
H -3.2432872 16.7657087 -14.3985759
C -3.0783778 14.9714564 -15.5675978
H -3.2811317 13.903087 -15.4809576
H -2.0269104 15.1037593 -15.8111044
H -3.6757972 15.3801559 -16.3812018
C -4.9110054 15.512951 -13.9555661
H -5.1400087 14.4586235 -13.8201664
H -5.5040244 15.9052164 -14.7807417
H -5.172629 16.0607214 -13.0540348
C -2.0596333 17.9254177 -12.3430272
O -1.3832877 18.2475852 -11.374691
N -3.271554 19.2249776 -12.5340521
H -3.8017849 18.9469697 -13.3433045
C -4.1853409 20.9390259 -11.9780666
H -4.1201797 20.8378689 -10.8941409
C -5.8142116 21.9798425 -12.0685304
O -6.4577917 21.990306 -13.1238488
C -3.3553264 22.2522131 -12.3368714
H -2.7508435 21.9810248 -13.199742
H -3.9431639 23.1195955 -12.6376299
C -2.3935577 22.6639575 -11.2141059
H -1.5875927 23.2551301 -11.6539897
H -1.9417954 21.7737149 -10.7710014
C -3.1027948 23.4921229 -10.1365626
O -4.1338848 23.0043489 -9.6189633
O -2.6283562 24.6174618 -9.8712537
N -6.4475046 23.0719258 -11.2698861

H -5.8681384 23.316696 -10.4654281
C -7.9445496 23.8116708 -11.2481781
H -8.5313975 23.1447577 -11.8786809
C -8.4505552 25.3183528 -11.7282815
O -7.9562339 26.3541144 -11.2608184
C -8.5827225 23.7319299 -9.8142739
H -8.3098143 22.7644516 -9.3910452
H -8.1237068 24.4929663 -9.1820639
C -10.1128287 23.8553926 -9.699581
H -10.4295997 24.8665053 -9.9608487
H -10.3884539 23.7049408 -8.6526486
C -10.8425833 22.8299823 -10.5754585
O -10.9583405 23.1023724 -11.793048
O -11.2574013 21.7859035 -10.0265513
N -9.602907 25.5151302 -12.4609812
H -10.1970189 24.6947314 -12.6038089
C -10.0093518 26.7968804 -13.139427
H -9.6033406 27.6153906 -12.5465371
C -11.5373224 27.0770783 -13.3129338
O -12.3430523 26.1660776 -13.4651874
C -9.3158021 26.7963794 -14.5218177
H -8.2652178 26.5407625 -14.3723088
H -9.7616841 25.9988297 -15.1187812
C -9.3656862 28.1006173 -15.3396433
H -10.3980501 28.3527717 -15.5746801
C -8.7151959 29.2767986 -14.6053596
H -7.6855865 29.0272619 -14.3458923
H -9.2616884 29.5103425 -13.6946444
H -8.721151 30.1580805 -15.2468748
C -8.6202634 27.902131 -16.6598573
H -7.5714266 27.6723762 -16.4704162
H -8.6904418 28.808201 -17.2612677
H -9.0736317 27.0808258 -17.214682
N -11.9412789 28.3572898 -13.4297045
H -11.2467902 29.0828588 -13.3712445
C -13.3321225 28.7589476 -13.708797
H -13.9922764 28.0787705 -13.1669539
C -13.6973559 28.668285 -15.20173
O -13.2212429 29.4599915 -16.0166546
C -13.5976231 30.1744778 -13.1694646
H -13.3339687 30.2032292 -12.1112647
H -12.9526566 30.8857408 -13.6884859
C -15.0439728 30.6091374 -13.3172145
C -16.0007793 30.2075446 -12.3654889
H -15.7026398 29.6095012 -11.5164327
C -17.3497155 30.5734271 -12.5229045

H -18.084102 30.2600018 -11.7942258
C -17.7459906 31.3401123 -13.632773
H -18.784204 31.6156339 -13.7569758
C -16.7930155 31.742311 -14.5848431
H -17.0984245 32.326439 -15.4416514
C -15.4440982 31.3771052 -14.4276294
H -14.7165289 31.6734974 -15.1713742
N -14.6326318 27.780819 -15.5396291
H -14.9521194 27.1150618 -14.8471413
C -15.2390964 27.6886577 -16.8666391
H -15.735075 28.6312197 -17.0994544
H -14.4624243 27.5134405 -17.6111349
C -16.2666536 26.5596064 -16.957317
O -16.3953891 25.7524785 -16.0401148
N -16.9764725 26.4878918 -18.0819378
H -16.8289854 27.176356 -18.8044725
C -17.9575033 25.4408391 -18.3831007
H -17.7899232 24.5827215 -17.7304426
C -17.7792012 24.9927546 -19.8397808
O -17.7959146 25.8188285 -20.750926
C -19.3742105 25.9740925 -18.1133224
H -19.4072767 26.3626889 -17.0943744
H -19.5911057 26.7896238 -18.8053619
C -20.4418095 24.8784784 -18.2588574
H -20.2170614 24.0779836 -17.5490798
H -20.3816661 24.4589508 -19.2655484
C -21.873747 25.3850101 -18.0153021
O -22.0329252 26.4946208 -17.4568844
O -22.8044396 24.6367793 -18.3888393
N -17.5576422 23.6953514 -20.0649298
H -17.6282623 23.0479037 -19.293315
C -17.3269048 23.1256682 -21.396853
H -17.6295227 23.8524687 -22.1539443
C -18.173276 21.8705255 -21.6133325
O -18.4492201 21.1344351 -20.6665456
C -15.8303326 22.833808 -21.5971231
H -15.5031065 22.1074362 -20.8521036
H -15.2664698 23.755845 -21.4523805
C -15.5235885 22.3038595 -22.9769314
N -16.0573017 22.8083599 -24.1656877
C -15.6125519 22.0028993 -25.140073
H -15.854249 22.1181749 -26.1888167
N -14.8449621 21.02673 -24.6277346
H -14.429788 20.2755673 -25.1579691
C -14.7816493 21.1953241 -23.2621255
H -14.2791179 20.561661 -22.5447457

N -18.5185024 21.5818415 -22.8701808
H -18.1432063 22.1738025 -23.6009505
C -19.2288161 20.3661674 -23.257331
H -19.9116933 20.1067755 -22.4457962
C -18.2431347 19.2015468 -23.4437471
O -17.5034608 19.1367934 -24.4310812
C -20.0753688 20.6536823 -24.5101802
H -20.7622502 21.4721288 -24.2876809
H -19.4111275 20.9769363 -25.3134978
C -20.8899068 19.4416545 -25.0031907
H -20.2061397 18.632868 -25.252551
C -21.8900833 18.9305006 -23.9653095
H -22.5664824 19.7301243 -23.6638583
H -21.3716057 18.543429 -23.0898099
H -22.4721546 18.1090986 -24.387707
C -21.6558409 19.8178908 -26.2704486
H -22.3670983 20.6156827 -26.0549008
H -22.1966695 18.9460329 -26.6408658
H -20.9593117 20.149503 -27.0399621
N -18.305586 18.2264973 -22.5363312
H -18.9848484 18.3084584 -21.7879638
C -17.4987295 17.0022879 -22.56464
H -16.4445225 17.2767669 -22.5352472
C -17.7136686 16.1851165 -23.8479637
O -16.7324037 15.7061853 -24.4089253
C -17.8183622 16.1676056 -21.3118248
H -17.6392762 16.7679502 -20.4189096
H -17.1541192 15.3031549 -21.279748
C -19.2469521 15.6902349 -21.2993199
N -20.3555822 16.5291505 -21.1770985
C -21.409785 15.8002387 -21.5780685
H -22.404937 16.1941265 -21.7444035
N -21.0189659 14.5690479 -21.940586
H -21.5461555 13.999855 -22.611493
C -19.6585309 14.4792161 -21.7690455
H -19.0246639 13.6887216 -22.1457658
N -18.9342631 16.1416122 -24.3967885
H -19.7139306 16.4659898 -23.8383305
C -19.2306073 15.4622993 -25.6698397
H -18.9191364 14.4191048 -25.5943322
C -18.4740459 16.0915514 -26.8531044
O -17.994577 15.3816032 -27.732199
C -20.7422481 15.4827005 -25.9605596
H -20.9140981 15.0341016 -26.9402223
H -21.0845436 16.5175317 -26.0055241
C -21.5715667 14.7083567 -24.9337949

O -21.1731487 13.5796127 -24.57134
O -22.5721805 15.2767049 -24.4473625
N -18.2496141 17.4112082 -26.8323886
H -18.5716361 17.9329869 -26.0288096
C -17.4531297 18.1047217 -27.8489587
H -17.7977212 17.8135617 -28.8424323
H -17.5879439 19.1789559 -27.7304308
C -15.956921 17.7942072 -27.7454374
O -15.2888172 17.6280449 -28.7627028
N -15.439025 17.6201509 -26.5247796
H -16.0609943 17.7043393 -25.7294654
C -14.0743246 17.1266898 -26.3031017
H -13.3826248 17.7038566 -26.9186169
C -13.9256189 15.6534948 -26.724003
O -12.9564171 15.2990751 -27.3953963
C -13.702554 17.3422062 -24.8291262
H -13.7609572 18.4063683 -24.6038623
H -14.431762 16.8391893 -24.1953113
C -12.3186415 16.8515113 -24.4543145
C -11.189853 17.6482056 -24.7279804
H -11.3040053 18.609059 -25.2098121
C -9.9051194 17.1911998 -24.3777829
H -9.0312386 17.792005 -24.5808524
C -9.7460546 15.9368061 -23.7535044
O -8.5019852 15.5094553 -23.4137942
H -8.5099345 14.6569228 -22.9779248
C -10.8775819 15.1370117 -23.4808996
H -10.7593813 14.1747477 -23.0085044
C -12.161335 15.5959257 -23.8337612
H -13.0275541 14.9793858 -23.6353157
N -14.9268359 14.8201123 -26.423841
H -15.7014839 15.176822 -25.8741997
C -14.9828394 13.4197593 -26.8408227
H -14.0769896 12.9181517 -26.4986893
C -15.0310294 13.2711907 -28.3685764
O -14.316041 12.4302505 -28.9112801
C -16.1878629 12.7409781 -26.1724631
H -16.0987507 12.8425083 -25.0898935
H -17.1015123 13.2506904 -26.4752072
C -16.3194695 11.2716038 -26.5208461
C -15.679925 10.2960364 -25.7306851
H -15.1296517 10.5903453 -24.8487413
C -15.7427155 8.9385754 -26.1018896
H -15.2470437 8.1817352 -25.5143806
C -16.4327316 8.5582085 -27.2723703
O -16.4601685 7.2539103 -27.6505408

H -16.8895214 7.1591939 -28.5014005
C -17.087672 9.5346431 -28.0532165
H -17.603424 9.2540987 -28.9595907
C -17.035941 10.887789 -27.6716777
H -17.5222426 11.6367727 -28.2851494
N -15.7781968 14.1209507 -29.0785084
H -16.4213335 14.735133 -28.58639
C -15.8302485 14.0991621 -30.5458371
H -16.023302 13.0723458 -30.857743
C -14.4935853 14.5210283 -31.1785911
O -13.967663 13.81329 -32.0333803
C -16.994066 14.9750367 -31.0424011
H -17.90762 14.6865268 -30.519727
H -16.7819786 16.0183928 -30.8031416
C -17.238316 14.8508303 -32.5555673
H -16.3290961 15.1177248 -33.0992785
H -18.0038016 15.5779021 -32.8375267
C -17.6986865 13.4395358 -32.9540066
O -18.9264795 13.2539991 -33.097649
O -16.8225467 12.5548429 -33.0888592
N -13.837982 15.5729289 -30.6701081
H -14.2892367 16.1201696 -29.9440593
C -12.4898729 15.9648231 -31.1351682
H -12.5346458 16.1337827 -32.2113024
C -11.4694612 14.8360737 -30.9194148
O -10.6875397 14.5295424 -31.8238719
C -12.0510367 17.2817301 -30.4618222
H -12.2135606 17.2075124 -29.385962
C -10.5745282 17.6261432 -30.7001145
H -10.360449 17.6415137 -31.7693089
H -9.9326989 16.8899406 -30.2164525
H -10.3471552 18.6032195 -30.2737645
C -12.8768666 18.454303 -31.009272
H -12.6844 18.5843449 -32.0745506
H -12.6143381 19.3701809 -30.4802161
H -13.9403605 18.2682826 -30.8668087
N -11.5418689 14.1369177 -29.7810758
H -12.2068198 14.4377171 -29.0741423
C -10.7457172 12.934077 -29.5229422
H -9.6880299 13.1653715 -29.6509979
H -10.9128238 12.6231638 -28.4919327
C -11.1015855 11.7534031 -30.4368387
O -10.2052046 11.0525872 -30.9091058
N -12.3806552 11.5703557 -30.7665667
H -13.0669852 12.1981408 -30.3596135
C -12.8830576 10.4904933 -31.6335773

H -12.4536585 9.5465298 -31.299618
C -12.4546029 10.6930136 -33.0856704
O -11.8379689 9.7967565 -33.662114
C -14.4115139 10.3664975 -31.5108397
H -14.8837716 11.3195435 -31.7430037
C -15.0075931 9.281508 -32.405127
H -14.497612 8.3332624 -32.2385487
H -14.9184434 9.5764776 -33.4497101
H -16.0681325 9.1738969 -32.1753859
O -14.7191285 9.9990169 -30.1858573
H -14.6611214 10.8041515 -29.6445041
N -12.6061399 11.9050074 -33.6278135
H -13.1277568 12.5991368 -33.0943308
C -12.1108057 12.2899864 -34.96069
H -12.5626445 11.6330904 -35.7039491
C -10.5924574 12.1086371 -35.0606404
O -10.1118234 11.5017037 -36.0194828
C -12.5253388 13.7419652 -35.2846723
H -12.2047179 14.3950538 -34.4716908
C -11.9014053 14.2583629 -36.5896139
H -12.1485647 13.587198 -37.4129321
H -10.8182307 14.3273054 -36.4922998
H -12.2846337 15.2548982 -36.8119629
C -14.0473735 13.8569901 -35.4397281
H -14.390893 13.2481522 -36.2760117
H -14.3288191 14.8958381 -35.6094773
H -14.5551193 13.5168148 -34.5364626
N -9.8327613 12.5183483 -34.0376294
H -10.2691124 13.0158676 -33.2677802
C -8.3866883 12.2947826 -33.9964652
H -7.94803 12.713808 -34.9032965
C -8.0253687 10.7951613 -33.964585
O -7.1750255 10.3535353 -34.7385592
C -7.808095 13.0498299 -32.7940855
H -8.2322251 12.6640796 -31.8665257
H -8.0425363 14.1119177 -32.8768195
H -6.7249933 12.9258191 -32.7693519
N -8.7383063 9.9928143 -33.1690346
H -9.4399586 10.4216383 -32.5757625
C -8.5261026 8.5373236 -33.0519002
H -7.476906 8.3564513 -32.8190178
C -8.836892 7.7959867 -34.3557814
O -8.0721417 6.9214099 -34.7582541
C -9.3671494 7.960609 -31.9009351
H -10.4270834 8.1175921 -32.1029293
C -9.1210937 6.4714297 -31.6602582

H -8.0569718 6.2924769 -31.5037895
H -9.4663804 5.8883818 -32.5134126
H -9.6722286 6.1524028 -30.7752524
O -9.0287859 8.6085484 -30.6970051
H -9.4300372 9.4946216 -30.711502
N -9.8765834 8.2071453 -35.0894348
H -10.4751493 8.9342993 -34.7066042
C -10.218649 7.6484065 -36.4087586
H -10.3078933 6.565189 -36.3149444
C -9.0961985 7.9031619 -37.4291875
O -8.8107229 7.0319327 -38.2457573
C -11.5883493 8.2058925 -36.8724934
H -11.5800876 9.2895272 -36.7418409
C -11.8649231 7.9080697 -38.3605028
H -11.8182208 6.8336763 -38.5433383
H -11.1330206 8.4106419 -38.9928817
H -12.8465962 8.2794081 -38.6492608
C -12.7194695 7.6055139 -36.002287
H -12.8725628 6.5606242 -36.275951
H -12.4291906 7.6247014 -34.9527788
C -14.0547499 8.3529914 -36.1158443
H -14.469557 8.2558855 -37.1180749
H -13.9111854 9.4081356 -35.8800284
H -14.765007 7.9268607 -35.4065953
N -8.3937191 9.0388911 -37.3367375
H -8.6846259 9.7176446 -36.6427142
C -7.265239 9.3812153 -38.2229573
H -7.5533014 9.1179589 -39.2408252
C -5.9497281 8.6286735 -37.9283472
O -4.9741865 8.7465544 -38.6840646
C -7.0390962 10.899473 -38.1899562
H -8.0014138 11.4136505 -38.2093944
H -6.4800146 11.1924024 -39.0771632
O -6.2990547 11.3175345 -37.05805
H -6.6658562 10.914714 -36.2473093
N -5.8900761 7.8921141 -36.8100548
H -6.7249579 7.8243468 -36.2397125
C -4.6565738 7.3302429 -36.256341
H -3.8362244 8.0007177 -36.5123651
C -4.2800623 5.9408626 -36.8074929
O -3.1352983 5.5205919 -36.6491747
C -4.7709993 7.3240925 -34.7269169
H -5.5236596 6.6026775 -34.4107831
H -5.0474193 8.3167635 -34.3677705
H -3.8122191 7.0434425 -34.2890576
N -5.1800992 5.2659515 -37.5310566

H -6.0659794 5.7270073 -37.7010793
C -4.9863126 3.9392569 -38.1512939
H -5.0038412 3.1815968 -37.3671395
C -3.6467069 3.7571432 -38.8886477
O -2.9731921 2.7386014 -38.7264792
C -6.1697208 3.6740353 -39.1048049
H -7.0402153 3.4328757 -38.4935022
H -5.9499646 2.7970377 -39.7150907
C -6.5637775 4.8197352 -40.0201111
N -5.7699764 5.9207787 -40.3641479
C -6.5674691 6.752385 -41.049963
H -6.277414 7.7318812 -41.4028678
N -7.7926363 6.2244398 -41.1826337
H -8.5854946 6.6964794 -41.5940588
C -7.8129371 5.0107498 -40.5332027
H -8.6720512 4.3747877 -40.3736965
N -3.1845526 4.7824287 -39.6094995
H -3.8397485 5.5490973 -39.7402001
C -1.9094941 4.7790629 -40.3503428
H -1.9058746 3.9210138 -41.0259016
C -0.6755178 4.6000095 -39.4590681
O 0.3500762 4.1523355 -39.9634635
C -1.8108003 6.0655976 -41.1818839
H -1.9802055 6.9149351 -40.5182176
H -0.8101594 6.155225 -41.6088882
C -2.8366963 6.0737128 -42.3295833
H -3.7805566 5.6467642 -41.9998432
H -2.4675716 5.4691415 -43.158521
C -3.090666 7.5062857 -42.7975055
H -2.21557 7.8834898 -43.328123
H -3.2604295 8.1357064 -41.9227019
N -4.2898586 7.6146451 -43.6353146
H -5.0985667 8.0752973 -43.2163844
C -4.3632322 7.5927761 -44.9465972
N -3.4118636 7.0991039 -45.680707
H -3.2172373 7.5904414 -46.5573116
H -2.6560279 6.644648 -45.2205878
N -5.4028833 8.1065841 -45.5234574
H -6.0523492 8.6180683 -44.9202338
H -5.4423454 8.1830156 -46.5185052
N -0.7546542 4.8641941 -38.1528525
H -1.6478001 5.1514024 -37.7665882
C 0.3441051 4.6164829 -37.2169392
H 1.2323201 5.1264964 -37.5885598
C 0.6996833 3.1210416 -37.0906775
O 1.8637562 2.7984179 -36.8587498

C -0.0133202 5.2297426 -35.858726
H -0.9017486 4.748926 -35.4486878
H -0.2065543 6.2971976 -35.9704315
H 0.8169899 5.0909097 -35.1647239
N -0.2423564 2.2117153 -37.3765593
H -1.1831707 2.5200405 -37.5942453
C 0.0441412 0.7786215 -37.4595675
H 0.5673747 0.4724402 -36.5515695
C 0.9552045 0.4341165 -38.6550826
O 1.7906982 -0.4590169 -38.5465271
C -1.2872208 0.0221727 -37.522223
H -1.8167315 0.2672544 -38.4447155
H -1.9079066 0.2944651 -36.668599
H -1.1024213 -1.0504495 -37.4916568
N 0.8781965 1.1977947 -39.7544587
H 0.2102942 1.9555281 -39.7611379
C 1.8112123 1.0873253 -40.8905783
H 2.0039892 0.0301488 -41.0804667
C 3.1532747 1.7347408 -40.5479506
O 4.1918602 1.1211358 -40.7786454
C 1.2242149 1.7004207 -42.1796402
H 1.0712918 2.7691655 -42.0362597
C 2.1680245 1.5163423 -43.3761676
H 2.3959256 0.458749 -43.5151264
H 3.0968456 2.0616945 -43.2111974
H 1.7047064 1.9050683 -44.2829471
C -0.1270984 1.0684292 -42.5415657
H -0.0134202 -0.0094525 -42.6618157
H -0.4995357 1.4948188 -43.4727939
H -0.8607535 1.2658962 -41.7604837
N 3.1497115 2.9075864 -39.8983128
H 2.2584563 3.3646896 -39.7477886
C 4.3829764 3.5769137 -39.4271644
H 5.0316617 3.7532182 -40.287047
C 5.1714679 2.6870662 -38.4572727
O 6.3914124 2.6466122 -38.5342007
C 4.0854947 4.9447377 -38.7734971
H 3.4492565 4.7868126 -37.9038288
C 5.3579813 5.667574 -38.3077099
H 6.0473034 5.7700222 -39.1479344
H 5.8520541 5.0988091 -37.5198768
H 5.1091859 6.6515776 -37.9116408
C 3.3657123 5.9094878 -39.7292705
H 4.0142422 6.146524 -40.5715909
H 3.1117469 6.8290265 -39.2030367
H 2.4524687 5.4645623 -40.1099666

N 4.5099699 1.8691594 -37.6261136
H 3.501018 1.9661057 -37.5679173
C 5.1768327 0.8861925 -36.7517641
H 5.9189905 1.4157616 -36.1524491
C 5.9655105 -0.1805465 -37.5278367
O 6.9655235 -0.6580613 -37.005066
C 4.1151826 0.2654933 -35.821984
H 3.6872668 1.0584642 -35.2070059
H 3.3115988 -0.1409395 -36.4366679
C 4.6026419 -0.8663175 -34.8987054
H 4.9701494 -1.6946528 -35.5041773
H 3.7425868 -1.2253198 -34.3369632
C 5.691348 -0.4487287 -33.8979481
H 5.305648 0.3329929 -33.2411631
H 6.5347315 -0.0299299 -34.4515579
N 6.1852786 -1.5988657 -33.1065998
H 7.0302672 -2.023504 -33.4565587
C 5.6931212 -2.0929878 -31.9796119
N 4.6137617 -1.6238221 -31.4121932
H 4.2496792 -2.0315972 -30.5644291
H 4.1500759 -0.8388203 -31.8268032
N 6.2826626 -3.0960025 -31.392016
H 7.158683 -3.4460252 -31.7437805
H 5.9220268 -3.4384356 -30.5210906
N 5.5130667 -0.5786026 -38.7196586
H 4.6989518 -0.1098716 -39.0946508
C 6.155844 -1.6215128 -39.5477068
H 6.7171868 -2.2989085 -38.9044692
C 7.1870424 -1.0471198 -40.5141252
O 8.1672071 -1.7115576 -40.8379074
C 5.0729423 -2.429844 -40.2970744
H 4.4719049 -1.7392024 -40.8919679
C 5.6505611 -3.4778044 -41.2668926
H 6.3652394 -4.1194898 -40.7503349
H 6.1577817 -2.9777053 -42.0922651
H 4.8537515 -4.0898952 -41.6888069
C 4.1272498 -3.1381766 -39.3001663
H 3.4521743 -2.3981917 -38.8745679
H 3.519856 -3.8567523 -39.8472324
C 4.8022508 -3.8790266 -38.1377464
H 5.1620025 -3.1674568 -37.3956499
H 5.6444629 -4.4611102 -38.5038849
H 4.0797422 -4.5405924 -37.6627072
N 6.9634562 0.1827065 -40.9660073
H 6.1059518 0.645179 -40.6903566
C 7.8918005 0.9019409 -41.8307362

H 8.3560989 0.1968688 -42.5199967
C 9.0541694 1.5083209 -41.0392823
O 10.1751688 1.4175308 -41.5113396
C 7.0910835 1.9372281 -42.6425295
H 6.4755174 2.5141194 -41.9494156
C 8.015055 2.9405889 -43.3353853
H 8.8170607 2.4130698 -43.8517664
H 8.4436482 3.6079035 -42.5859543
H 7.4537262 3.5413332 -44.0376991
C 6.1329575 1.2547909 -43.6511673
H 5.3388 0.7519175 -43.1009127
H 5.6559528 2.0240523 -44.2518133
C 6.7607953 0.2206656 -44.5975052
H 7.0017951 -0.6915304 -44.0477814
H 7.6692296 0.6123557 -45.0487245
H 6.0481605 -0.0291608 -45.3837918
N 8.798992 1.9819869 -39.8195457
H 7.8269675 1.9509433 -39.5448266
C 9.6149082 2.8267209 -38.9312898
H 9.2170332 2.5797365 -37.9459199
C 9.3250434 4.3409519 -39.0306445
O 8.5238253 4.7993794 -39.8542185
C 11.099113 2.4020543 -38.8279642
H 11.7082873 3.1408157 -39.3434734
H 11.2510327 1.4269978 -39.2904088
C 11.5965131 2.3038259 -37.3833269
O 11.4492724 3.3205405 -36.6703128
O 12.1249463 1.2330349 -37.0094698
N 9.8588958 5.1058401 -38.0754761
H 10.516748 4.6466534 -37.4396315
C 9.3325992 6.4041998 -37.6512809
H 8.3526531 6.2042132 -37.2199303
C 9.1395831 7.4234129 -38.7896545
O 10.0494829 7.7492656 -39.5515223
C 10.2024572 6.9760756 -36.5200309
H 10.3373002 6.2059069 -35.7593743
H 11.1814432 7.2466722 -36.9189787
C 9.5601829 8.2096529 -35.8657287
H 8.5933201 7.9247972 -35.443284
H 9.3792816 8.9717041 -36.6261681
C 10.4478246 8.8008571 -34.7647909
O 10.8020372 8.0456266 -33.8314376
O 10.7428137 10.012847 -34.8410345
N 7.9356494 8.0039627 -38.8202964
H 7.2747294 7.6949909 -38.1276181
C 7.4979327 9.066309 -39.7246913

H 6.4475177 9.2219085 -39.4784392
C 7.4607978 8.7250306 -41.2328113
O 7.2182063 9.6195186 -42.0501456
C 8.189382 10.3821855 -39.339387
H 9.2533861 10.3291555 -39.5703739
H 8.0772996 10.5585665 -38.2684137
H 7.7299772 11.2135753 -39.8729389
N 7.5456185 7.4423073 -41.6155564
H 7.7541 6.7354562 -40.9135248
C 7.344726 6.9801426 -42.997353
H 6.9802142 7.8289962 -43.5714683
C 6.2228111 5.9398072 -43.1423331
O 5.8136015 5.298277 -42.1811172
C 8.7040589 6.5417867 -43.5937623
H 9.3769214 6.2789443 -42.781539
H 8.5857406 5.6690598 -44.2357264
C 9.2911549 7.7100855 -44.425368
H 9.1907816 8.6410518 -43.8675988
C 10.7676023 7.5379856 -44.7641775
H 10.9094523 6.6899523 -45.4326336
H 11.347273 7.4053009 -43.8535032
H 11.1408564 8.4293545 -45.2677901
C 8.5546316 7.8484516 -45.7653518
H 8.6226347 6.9085588 -46.3141622
H 9.0244539 8.6267692 -46.3653165
H 7.5122898 8.121164 -45.6183874
N 5.6747873 5.8134553 -44.3622881
H 6.051826 6.3868802 -45.1032308
C 4.5891328 4.8649351 -44.6980872
H 4.5631062 4.1139725 -43.9051263
C 4.8447935 4.0459821 -45.973334
O 3.9017478 3.6498304 -46.6520897
C 3.2105171 5.5603428 -44.666108
H 2.4533252 4.7738203 -44.6831793
H 3.1054279 6.0864467 -43.7153432
C 2.9161759 6.5454592 -45.8179066
H 3.1261552 6.0697121 -46.7735315
H 3.5448542 7.4315362 -45.7241825
C 1.4308664 6.9484155 -45.7776237
H 1.2590212 7.5693356 -44.8966521
H 0.8212512 6.0482975 -45.6740764
N 0.981203 7.6811794 -46.9780385
H 0.7524881 8.6673509 -46.8923419
C 0.6228584 7.177358 -48.1443321
N 0.8186153 5.929421 -48.4685675
H 0.5752193 5.6132175 -49.3870961

H 1.388143 5.3577715 -47.8677134
N 0.0406873 7.949872 -49.0092621
H -0.2933387 8.855375 -48.6610897
H -0.2623347 7.6163466 -49.9005672
N 6.1190758 3.8578277 -46.3209588
H 6.8301691 4.1363819 -45.6635574
C 6.5958218 3.2428633 -47.5626859
H 5.830005 2.5719029 -47.9542416
C 7.8916732 2.4367347 -47.343168
O 8.5668267 2.64766 -46.3401455
C 6.8674228 4.3491416 -48.5851797
H 7.7970464 4.8464069 -48.304794
H 7.032437 3.8770175 -49.5533758
C 5.793543 5.4093771 -48.7466088
C 4.5694425 5.071977 -49.3430479
H 4.3617759 4.0434651 -49.5976387
C 3.6563468 6.0786253 -49.6922932
H 2.7837729 5.8310304 -50.2772817
C 3.9157208 7.4151039 -49.3467661
H 3.2349086 8.196869 -49.6514519
C 5.0976506 7.7468283 -48.6644446
H 5.3156042 8.7854837 -48.4512446
C 6.0470079 6.7471097 -48.3865945
H 6.9997683 7.0245668 -47.966154
N 8.2414517 1.5676536 -48.3001291
H 7.6597218 1.5224626 -49.118932
C 9.3979198 0.6578697 -48.2914649
H 9.1016447 -0.2646165 -47.791358
C 10.6602547 1.1969945 -47.5742744
O 11.1786285 2.245083 -47.9427162
C 9.7087369 0.3224341 -49.75895
H 9.967396 1.234773 -50.3010873
H 8.8483582 -0.1504811 -50.2328802
H 10.5608699 -0.3531544 -49.8204047
N 11.1058984 0.4683517 -46.5404187
H 10.6020477 -0.3756197 -46.3181145
C 12.3498292 0.6411544 -45.7620579
H 12.0842348 0.2321049 -44.78769
C 12.7677959 2.0582833 -45.3295639
O 13.9257997 2.2966503 -44.965395
C 13.488643 -0.2542563 -46.2966031
H 14.4106292 0.3125754 -46.3885813
H 13.233996 -0.5667301 -47.3062372
C 13.7404825 -1.5012609 -45.4212878
O 12.782663 -1.9446545 -44.7479203
O 14.8558588 -2.0616043 -45.4835954

N 11.7676098 2.9221305 -45.1175289
H 10.8663841 2.6705479 -45.5032949
C 11.8434355 4.1137479 -44.2565654
H 10.9702733 4.7131008 -44.4953178
C 13.0474841 5.0013958 -44.5690478
O 13.2566867 5.4146228 -45.7033659
C 11.6839069 3.7215239 -42.7654516
H 10.7042933 3.2772 -42.7433744
C 12.7172875 2.6995277 -42.243958
H 13.7411237 3.021697 -42.4103113
H 12.5627012 1.7344082 -42.7242798
H 12.5786212 2.5252549 -41.1832717
C 11.6078686 4.8907969 -41.7666777
H 12.5088229 4.963613 -41.1569353
H 10.7748965 4.7152793 -41.0967139
H 11.4403098 5.837781 -42.2673589
N 13.7997814 5.3551081 -43.5357305
H 13.5427582 4.9757027 -42.6392564
C 14.9722896 6.1999847 -43.5773004
H 14.7513394 7.1316738 -44.1038787
C 16.1325397 5.5277022 -44.3124657
O 16.971215 6.2121722 -44.8791594
C 15.3363167 6.5059358 -42.1167684
H 14.5993989 7.1982355 -41.7073855
H 15.2826306 5.5845827 -41.5328418
C 16.7149966 7.0988042 -41.9408267
C 16.9780411 8.3984896 -42.403626
H 16.1908624 8.9809401 -42.8654804
C 17.7536204 6.3218351 -41.3924162
H 17.5600729 5.3123153 -41.0611771
C 18.2759221 8.9240865 -42.299878
H 18.4802673 9.9218835 -42.6403416
C 19.0546992 6.8452967 -41.3078177
H 19.8555147 6.2433117 -40.90308
C 19.314826 8.149128 -41.7601691
H 20.3159865 8.5514257 -41.6963891
N 16.2058513 4.196207 -44.3268838
H 15.3924863 3.6449569 -44.0809636
C 17.3365519 3.5317345 -44.9880782
H 18.247824 4.0148928 -44.6328091
C 17.3362498 3.7162578 -46.5202865
O 18.411362 3.6889561 -47.1212283
C 17.4139431 2.0551039 -44.544089
H 17.2640812 2.0186564 -43.4640312
C 18.7685605 1.4097662 -44.8475106
H 18.7772956 0.3945679 -44.4540022

H 19.5699411 1.9849547 -44.3903993
H 18.9200973 1.3641078 -45.9264033
O 16.4573901 1.2286298 -45.1599023
H 15.5577683 1.5969757 -45.0914068
N 16.1818698 4.0690221 -47.1078517
H 15.3465728 4.1192239 -46.5422669
C 16.0387294 4.3659216 -48.539581
H 16.9320402 3.9392428 -48.9945931
C 16.0907108 5.8966401 -49.0803814
O 16.0794458 5.9767522 -50.2908172
C 14.8556147 3.4957892 -49.0773819
H 13.9798023 4.1366509 -49.1933825
H 14.5968292 2.7346661 -48.3378472
C 15.1275989 2.7525254 -50.4024108
O 14.1400315 2.2286328 -50.965854
O 16.307982 2.6299838 -50.8096799
N 16.2900343 7.1238414 -48.4203981
H 16.4497442 7.0131143 -47.4277553
C 16.1613041 8.688626 -48.8951202
H 15.5643498 8.6270493 -49.8051702
C 17.4543756 9.8556596 -49.3389581
O 18.4787587 9.2741032 -49.0634247
C 15.1839224 9.2297703 -47.7945255
H 14.6070166 8.4267525 -47.3302822
H 14.4664933 9.8869038 -48.2885934
C 15.9020718 10.0680588 -46.713811
O 16.3872989 11.1886415 -47.009515
O 15.9809798 9.6471257 -45.537687
N 17.6582507 11.2710311 -50.0187799
H 16.7432841 11.6473397 -50.215925
C 18.9196021 12.4999217 -50.4996121
H 19.4239745 12.7072473 -49.557024
C 19.8910586 11.6284874 -51.3214552
H 19.3040143 10.926469 -51.916006
H 20.4598406 12.2416738 -52.0211191
C 20.8996192 10.8577388 -50.4511839
H 20.4908512 10.6758339 -49.4595413
H 21.799341 11.4600084 -50.3269145
C 21.2793501 9.5187183 -51.0747255
O 22.2065281 9.4215954 -51.8603748
N 20.5592735 8.459898 -50.7803827
H 20.794535 7.5910496 -51.2218482
H 19.7724068 8.5434667 -50.1355936
C 19.092366 14.182846 -51.3913657
O 18.0971829 14.350783 -52.0856275
N 20.1085432 15.3856543 -51.5493382

H 20.9260687 15.212857 -50.986138
C 20.2948739 16.9461427 -52.3131094
H 20.2227503 16.7196393 -53.3770916
C 21.4078328 18.3179152 -52.2406162
O 22.5468109 18.183726 -51.7988277
C 19.0686253 17.7464106 -51.9697146
H 19.2338989 18.2728335 -51.0391949
H 18.2005998 17.0932304 -51.9204808
H 18.8667864 18.4802587 -52.7438545
N 21.1702238 19.6749261 -52.6514902
H 20.2345851 19.8379585 -52.9935248
C 22.0024176 21.0115075 -52.4479166
H 22.8683875 20.7431257 -51.8441611
C 21.2832516 22.1579803 -51.6588977
O 20.098186 22.3880363 -51.8856444
C 22.5113401 21.7661131 -53.7349204
H 21.6390627 22.0773934 -54.3111731
H 23.0283835 22.6774737 -53.4283394
C 23.4507222 21.0203275 -54.6889031
H 22.9115143 20.1707727 -55.1106251
H 23.7236569 21.6915994 -55.5045992
C 24.7266258 20.5101083 -54.0066815
H 24.4530278 19.8220364 -53.2042071
H 25.2791937 21.3544152 -53.5879837
C 25.5984771 19.7681775 -55.0268131
H 25.909443 20.4728439 -55.8051529
H 24.9888066 18.9920389 -55.5006992
N 26.7854235 19.1515691 -54.3842775
H 27.3653918 19.8580064 -53.9512649
H 27.3408378 18.6543916 -55.0695705
H 26.4940335 18.4923601 -53.6723818
N 22.0141776 22.9790716 -50.8776924
H 22.9848334 22.7561847 -50.7481297
C 21.4363151 24.0595014 -50.0406279
H 20.6651534 23.5880877 -49.4217957
C 20.6932301 25.1571828 -50.8434591
O 21.1123859 25.4928957 -51.9501055
C 22.4954675 24.6021902 -49.0576117
H 21.9681595 25.0697176 -48.2281703
H 23.0446883 23.7611733 -48.6298199
C 23.4967813 25.6136439 -49.6395768
H 24.1308012 25.1185841 -50.3761876
H 22.9616084 26.426131 -50.1294037
C 24.3600993 26.2058559 -48.5122502
H 25.0695733 25.4548231 -48.159759
H 23.7129745 26.4715583 -47.6719088

C 25.1007745 27.47991 -48.9414417
H 25.6487684 27.8643149 -48.0751497
H 24.3538345 28.231996 -49.2207631
N 26.033051 27.2429212 -50.0717632
H 26.729265 26.5557402 -49.8134378
H 26.5058074 28.1019389 -50.3270792
H 25.5216444 26.9054137 -50.8778729
N 19.6653497 25.7902966 -50.2502701
C 18.5260935 26.4199977 -50.9302971
H 17.683172 26.2611953 -50.25573
C 17.9666354 25.9230463 -52.2825018
O 17.2120042 26.6858309 -52.8855268
C 18.8245867 27.9163968 -50.8994079
H 19.6064095 28.1600803 -51.6210341
H 17.9311673 28.5198536 -51.0658878
C 19.9982801 26.7186724 -49.1631419
H 19.6497361 26.3469128 -48.204326
H 21.0706298 26.8855675 -49.103357
C 19.3411503 28.0678737 -49.4726293
H 18.5032263 28.2293829 -48.7937525
H 20.0551531 28.8885143 -49.395854
N 18.1444302 24.6585658 -52.7170806
H 18.7575415 24.0313991 -52.2081714
C 17.2497154 24.0944877 -53.7603078
H 17.1809009 24.8305569 -54.5644565
C 15.8108259 23.9536811 -53.2184155
O 15.6116559 23.7733122 -52.0168994
C 17.7905208 22.7718561 -54.367871
H 18.2164828 22.1394345 -53.5915213
H 16.9504542 22.2274091 -54.799801
C 18.8275157 22.9966047 -55.490398
H 19.7490875 23.3914824 -55.0618205
H 18.4292468 23.7618966 -56.1540239
C 19.1471149 21.7322105 -56.331149
H 18.3107319 21.0347568 -56.2604309
H 20.0375569 21.2460875 -55.9308658
N 19.3493069 22.0762304 -57.7577649
H 19.2235844 23.0639289 -58.0004847
C 19.3902273 21.3013719 -58.8329725
N 19.5688706 20.0105413 -58.789513
H 19.5868354 19.4947518 -59.6476506
H 19.6328429 19.5190873 -57.9186205
N 19.2411043 21.8219077 -60.01574
H 18.7989704 22.7503794 -60.07748
H 19.2658129 21.2691169 -60.8598066
N 14.821947 23.9635607 -54.1171727

H 15.0640462 24.1295175 -55.0795782
C 13.4070457 23.7053603 -53.8023649
H 13.2318405 23.8314889 -52.7303655
C 13.0148305 22.2690653 -54.1901151
O 13.2650034 21.8554559 -55.3268438
C 12.5042714 24.70102 -54.554355
H 11.4657565 24.4018961 -54.4176821
H 12.7363243 24.6603459 -55.6166903
C 12.5918951 26.14774 -54.1111584
O 13.1869008 26.5106661 -53.1113018
N 11.9857542 27.0281791 -54.8750885
H 12.0663734 27.990201 -54.6088372
H 11.474667 26.7304553 -55.7012412
N 12.3104354 21.5618859 -53.3039725
H 12.1319232 21.9882772 -52.3959269
C 11.7798987 20.2088839 -53.5218785
H 12.1354009 19.8236533 -54.4750295
C 10.2499869 20.2006078 -53.5304971
O 9.636071 20.6928764 -52.5837385
C 12.2939867 19.2814334 -52.4104324
H 12.0003572 19.7111831 -51.4550013
H 13.3811827 19.2399207 -52.459261
C 11.7331357 17.8531895 -52.451972
H 10.6473428 17.8812916 -52.5280063
H 11.9562163 17.3702826 -51.5054196
S 12.3827862 16.813984 -53.7779941
C 13.8931229 16.2105975 -52.9728887
H 14.4627188 15.602152 -53.6752431
H 13.6285852 15.6018071 -52.1075691
H 14.5036321 17.0516099 -52.6454799
N 9.6561252 19.476022 -54.4810838
H 10.2356146 19.1014098 -55.228018
C 8.2618632 19.0294284 -54.4110718
H 7.7432147 19.5950109 -53.6411374
C 8.2012683 17.5367186 -54.0391612
O 8.6091906 16.6916482 -54.8315145
C 7.5848191 19.3377223 -55.7563497
H 7.6533027 20.404727 -55.9472126
H 8.1407942 18.8235216 -56.5354205
C 6.1094319 18.9161937 -55.8471269
H 6.0223168 17.8621598 -55.5987058
C 5.2008171 19.7105303 -54.9111302
H 5.2842959 20.7773402 -55.1160026
H 5.4789999 19.5191252 -53.8783082
H 4.1706631 19.3882301 -55.0433692
C 5.5958432 19.1157678 -57.2735893

H 5.6369121 20.1715894 -57.5396105
H 4.5696247 18.7588489 -57.3505647
H 6.2159663 18.5490014 -57.9673902
N 7.650806 17.207511 -52.8705797
H 7.2809473 17.9542867 -52.2913677
C 7.3710433 15.8279812 -52.4464654
H 8.0060151 15.1508264 -53.0126345
C 5.9027388 15.4793458 -52.7611271
O 4.9709185 16.075372 -52.2065813
C 7.75986 15.6405981 -50.9667627
H 7.1203425 16.2573012 -50.3370099
H 8.7862314 15.9905422 -50.8370111
C 7.6824443 14.1863395 -50.4760692
O 6.8708591 13.4090086 -51.019936
O 8.4087115 13.8653462 -49.5055914
N 5.6864177 14.5711102 -53.7189983
H 6.4974395 14.0713565 -54.0777662
C 4.3447916 14.1842873 -54.1843736
H 3.6345714 14.9349283 -53.8406842
C 3.9351687 12.8544228 -53.5605881
O 4.3232699 11.7831128 -54.0192536
C 4.2336867 14.1575708 -55.7201635
H 4.8568936 13.3737071 -56.1278179
C 2.7974381 13.8629456 -56.1479142
H 2.1426664 14.6095088 -55.703318
H 2.4969621 12.8683791 -55.8186253
H 2.7229068 13.8922377 -57.2343704
C 4.6461988 15.4961607 -56.3443168
H 4.0765308 16.3124174 -55.897042
H 4.4772019 15.482627 -57.4198195
H 5.7094427 15.6583488 -56.1699027
N 3.0257182 12.9205146 -52.5909151
H 2.717312 13.8404901 -52.2952782
C 2.6511322 11.7941466 -51.7378099
H 1.5919382 11.8736081 -51.495824
H 2.826334 10.8515972 -52.2505211
C 3.4378837 11.764253 -50.4292475
O 3.7783718 10.6859284 -49.9559292
N 3.6725557 12.9262538 -49.8125931
H 3.3218807 13.7634739 -50.2589932
C 4.5639542 13.1282033 -48.6574106
H 5.5763021 12.8435635 -48.9560907
C 4.2394702 12.317265 -47.379758
O 4.9944892 12.3378076 -46.4012804
C 4.557135 14.6320349 -48.3562011
H 4.9178299 15.1702423 -49.2356233

H 5.2327209 14.8285762 -47.5267004
S 2.8894722 15.2074225 -47.9074698
H 2.7061254 14.4032963 -46.8479299
N 3.0995178 11.6227276 -47.3420058
H 2.5537956 11.6005599 -48.1895828
C 2.6973692 10.7695608 -46.2284903
H 1.7296098 10.3306215 -46.4382888
H 3.4252333 9.964224 -46.1301078
C 2.5964225 11.5335775 -44.9096616
O 2.0756425 12.6458024 -44.8617237
N 3.1475683 10.9405853 -43.8409104
H 3.5995258 10.050913 -43.9813798
C 3.2644505 11.5676352 -42.5110625
H 2.5363332 12.3721322 -42.4922009
C 4.6024855 12.3215117 -42.2989558
O 4.9330438 12.7035656 -41.1837733
C 2.7685976 10.6219858 -41.3876643
H 3.329892 9.6866489 -41.3966698
H 2.9110664 11.1133101 -40.4234859
C 1.2482459 10.3497499 -41.5881163
H 0.7919812 11.3134804 -41.7977461
H 1.1127473 9.7337692 -42.4784474
C 0.4227669 9.7161131 -40.4391496
H 0.7314226 10.1512193 -39.4901884
H 0.5960085 8.639396 -40.4025918
C -1.0758004 10.0217816 -40.7095582
H -1.41602 9.4220592 -41.5561837
H -1.1388727 11.0716698 -41.0127608
N -2.0107233 9.8625097 -39.5598994
H -2.0996365 8.9127489 -39.2334567
H -2.9406933 10.1949484 -39.8135329
H -1.7309402 10.4529809 -38.7839952
N 5.3030928 12.6456133 -43.3932679
H 4.9509657 12.3158047 -44.2839729
C 6.263606 13.7539353 -43.4685514
H 6.339384 14.0373157 -44.5168193
H 5.8664254 14.6025745 -42.9108326
C 7.6966603 13.5217271 -42.9844836
O 8.5229441 14.4189115 -43.1446022
N 8.052298 12.3272621 -42.5032093
H 7.3432049 11.6119669 -42.3818921
C 9.4128024 12.0200965 -42.040962
H 9.6646061 12.6654394 -41.1981862
H 9.4474178 10.9834761 -41.7123813
C 10.481707 12.1834795 -43.1192423
O 11.5222019 12.7773601 -42.8556136

N 10.1750316 11.7843984 -44.3589289
H 9.2846575 11.3332788 -44.4955374
C 11.0186339 12.0413754 -45.5425943
H 11.9980506 11.588156 -45.3790969
C 11.2701527 13.5389522 -45.7486745
O 12.4236188 13.9518113 -45.8260238
C 10.4258214 11.3832015 -46.8083101
H 10.5905208 10.3099959 -46.7269139
C 8.9312833 11.6033541 -47.0463739
H 8.7099762 12.6632397 -47.1442538
H 8.3393666 11.1827496 -46.2344072
H 8.6361123 11.1131843 -47.9758533
O 11.0895781 11.8306891 -47.9597574
H 10.484578 12.3844576 -48.4701324
N 10.2252136 14.371186 -45.663101
H 9.3074535 13.9722939 -45.5446904
C 10.2857366 15.8236971 -45.8547703
H 10.7235008 16.0265557 -46.8268567
C 11.1106349 16.537419 -44.7650356
O 11.8863183 17.4526382 -45.0535196
C 8.8657689 16.3970441 -45.8856242
H 8.8986116 17.4030429 -46.3069571
H 8.4867539 16.4648006 -44.8668808
S 7.7163401 15.3759861 -46.8510702
H 8.4533831 15.1086649 -47.9512639
N 10.9632892 16.0999518 -43.508704
H 10.2825801 15.3646959 -43.3383485
C 11.7666912 16.5933009 -42.3777083
H 11.7210304 17.682167 -42.3558897
C 13.2337723 16.2149077 -42.5646649
O 14.1041332 17.0794444 -42.4631432
C 11.2181531 16.0596538 -41.0385742
H 11.0788588 14.9803609 -41.1088303
C 12.1583111 16.3454275 -39.8608403
H 12.413162 17.4044263 -39.8365978
H 13.0713173 15.7583224 -39.9544092
H 11.6747709 16.0658247 -38.9240883
C 9.8619149 16.7101419 -40.7311641
H 9.9738266 17.7906836 -40.6408162
H 9.465485 16.313601 -39.7953368
H 9.147258 16.4904463 -41.5236473
N 13.5086787 14.9554922 -42.9189931
H 12.7374487 14.300883 -43.0121126
C 14.8616324 14.4600437 -43.1763215
H 15.4605874 14.5795315 -42.2714439
C 15.5717593 15.257738 -44.2626702

O 16.6984701 15.6980688 -44.0520232
C 14.8119003 12.9696725 -43.5454865
H 15.47017 12.778969 -44.3885111
H 13.8159203 12.6800994 -43.8734669
C 15.2350372 12.1081566 -42.3887418
N 16.5503927 11.9810507 -41.9354251
C 16.4753185 11.2196844 -40.8299497
H 17.3253826 10.9192111 -40.2306378
N 15.2029599 10.8627996 -40.5836198
H 14.890693 10.2906702 -39.8085624
C 14.4037633 11.4233475 -41.5538219
H 13.3257702 11.3738953 -41.6274226
N 14.9078435 15.5075468 -45.387301
H 13.989292 15.091668 -45.5152898
C 15.5112271 16.2350314 -46.5011762
H 16.4944499 15.8005517 -46.6810838
C 15.788866 17.702243 -46.1455768
O 16.9395239 18.1374501 -46.2012692
C 14.6897345 16.0179734 -47.7882799
H 15.1327128 16.6236197 -48.5603665
C 14.7908711 14.5234165 -48.182398
H 14.1962195 13.8987777 -47.5198436
H 15.8220997 14.1781548 -48.1262874
H 14.4372692 14.3716629 -49.2011021
C 13.2366266 16.4975189 -47.7320752
H 12.7646062 16.0195718 -46.8976035
H 13.2059957 17.5711979 -47.5585925
C 12.4407379 16.1837724 -49.0033466
H 12.2244151 15.1164961 -49.062049
H 13.0189562 16.48687 -49.8741816
H 11.4970485 16.7230964 -48.9888032
N 14.8187234 18.4245126 -45.5793613
H 13.9008259 18.0120409 -45.4473034
C 15.0314858 19.8172963 -45.1857533
H 15.4248899 20.3646793 -46.0422485
C 16.0654803 19.9815299 -44.052507
O 16.775709 20.9872676 -44.0197683
C 13.6739593 20.4215669 -44.8288207
H 13.2487701 19.8884107 -43.9775321
H 12.9997992 20.3346726 -45.682005
H 13.7950387 21.4777533 -44.588073
N 16.1945128 19.0031707 -43.1417661
H 15.5550477 18.2125874 -43.1960906
C 17.1426889 19.0652184 -42.0165262
H 17.2682905 20.115006 -41.7540704
C 18.5555244 18.5806105 -42.3633946

O 19.5022617 19.2022987 -41.8920977
C 16.5356863 18.3408313 -40.8014493
H 15.5546198 18.7709715 -40.5929376
H 16.3868088 17.2899072 -41.0560764
C 17.3783728 18.4041283 -39.5137768
H 18.2761985 17.8007418 -39.655826
H 16.803368 17.9420041 -38.7110996
C 17.8159365 19.8120997 -39.0658109
H 18.4907804 20.2290409 -39.816017
H 18.3932875 19.7217441 -38.1440225
N 16.6897875 20.7491832 -38.8609668
H 16.5665419 21.4502419 -39.5716769
C 15.8807065 20.8193544 -37.8169131
N 14.9853566 21.7589121 -37.7435133
H 14.3645694 21.8061651 -36.956886
H 14.8726153 22.4170928 -38.4946404
N 15.9472023 19.9843426 -36.8189475
H 16.6241288 19.2455779 -36.8502505
H 15.2949912 20.0477172 -36.0589826
N 18.7300829 17.5290883 -43.1764286
H 17.9068943 17.0488226 -43.5306661
C 20.0684893 17.0390531 -43.5746573
H 20.7065707 16.9291063 -42.6964889
C 20.7865705 18.0002679 -44.5366372
O 22.0134416 18.0053269 -44.5586972
C 19.9813346 15.699411 -44.3219547
H 19.304658 15.8747004 -45.1616404
H 20.9671976 15.5209637 -44.7569981
C 19.5628823 14.3578569 -43.6784448
H 20.3290906 14.0366348 -42.971993
H 18.6129878 14.4437749 -43.1562454
C 19.4543242 13.3380914 -44.844005
H 20.418487 13.3271226 -45.354375
H 18.7167883 13.694725 -45.5669937
C 19.1362141 11.8730891 -44.4980398
H 19.490443 11.6480878 -43.4892399
H 19.686472 11.2395421 -45.2001729
N 17.6955314 11.5486037 -44.6343123
H 17.1272324 12.0169392 -43.9401613
H 17.4806865 10.5531329 -44.5600195
H 17.3202713 11.7308229 -45.5705452
N 20.0478271 18.6624719 -45.4364741
H 19.046923 18.5093921 -45.436498
C 20.6358853 19.3309775 -46.6085273
H 21.6749389 19.5607893 -46.3684505
C 20.0541746 20.7091617 -46.9622581

O 20.3152965 21.2210728 -48.0515675
C 20.6868561 18.354292 -47.7957786
H 21.0473326 18.879063 -48.6814314
H 21.4566581 17.6177888 -47.5613333
C 19.4373287 17.602644 -48.1667651
C 19.394892 16.1940631 -48.1271936
H 20.196105 15.6434021 -47.6560693
C 18.3451897 15.5113149 -48.7728262
H 18.2923324 14.4372359 -48.7484014
C 17.3896775 16.2497781 -49.5044677
O 16.5031357 15.6745726 -50.3589183
H 16.8175994 14.8393012 -50.723428
C 17.3695605 17.6457992 -49.4114359
H 16.6398896 18.1806343 -49.9827419
C 18.3848271 18.32481 -48.7295873
H 18.4231516 19.3990316 -48.7319403
N 19.3517895 21.3539171 -46.024161
H 19.1651048 20.8775033 -45.1539639
C 19.0384741 22.7911583 -46.0750765
H 18.3539854 22.9771147 -45.2457682
C 18.2660689 23.2374827 -47.3416898
O 18.3498463 24.3844502 -47.7803859
C 20.3254224 23.5929801 -45.7682459
H 21.0190659 23.4837652 -46.6030665
H 20.8007875 23.1682607 -44.8823023
C 20.0831759 25.0814136 -45.500918
O 19.0655738 25.4065578 -44.8420731
O 20.8655128 25.9146396 -46.0124582
N 17.4548563 22.3556176 -47.9295338
H 17.4238769 21.424189 -47.5465261
C 16.4823397 22.7281121 -48.9710723
H 16.9464183 23.4477338 -49.6473765
C 15.24281 23.3977265 -48.3725054
O 14.9522023 23.2457856 -47.1836795
C 16.0422905 21.5162724 -49.8130752
H 15.2603537 21.8499514 -50.4920367
C 17.2011516 21.0558416 -50.7125561
H 18.1143336 20.9147098 -50.1404907
H 17.4026366 21.8357731 -51.4439333
H 16.943333 20.1436236 -51.2459316
C 15.398919 20.4182088 -48.9405732
H 16.1665696 19.8238888 -48.4571155
H 14.7955406 20.8598812 -48.151009
C 14.4493492 19.5350595 -49.7478417
H 14.8862927 19.2531794 -50.7036183
H 13.5268763 20.0795328 -49.9194258

H 14.2201392 18.6408149 -49.1834476
N 14.4360211 24.010578 -49.2375354
H 14.7402083 24.0446147 -50.2085445
C 12.9993631 24.158699 -49.0076405
H 12.8175786 24.2573771 -47.9397338
C 12.2662528 22.9046082 -49.5103378
O 12.6107608 22.3634176 -50.5604784
C 12.5234879 25.4517901 -49.6859483
H 13.0406288 26.283309 -49.2048236
H 12.830272 25.44303 -50.731898
C 11.0112581 25.7338392 -49.6079951
H 10.8575365 26.7986213 -49.7828324
H 10.658697 25.5126149 -48.6002629
C 10.1424015 24.9839405 -50.6247752
O 10.5934761 24.3664138 -51.5757024
N 8.8434225 24.9954737 -50.4307784
H 8.2650384 24.4828781 -51.0906918
H 8.4331033 25.5005896 -49.6701504
N 11.2359635 22.4621827 -48.7844512
H 11.0578115 22.9049529 -47.8891113
C 10.3429926 21.383246 -49.214284
H 10.5949631 21.0920705 -50.235015
C 8.8711568 21.8264889 -49.2272665
O 8.3637704 22.3460571 -48.2286926
C 10.5732244 20.1618264 -48.309774
H 10.3332883 20.4190762 -47.2777976
H 11.6227969 19.8672222 -48.3609254
S 9.5400098 18.7650291 -48.8429061
H 8.3402798 19.3259352 -48.6493615
N 8.1565084 21.4877271 -50.299685
H 8.6580875 21.1050509 -51.0997332
C 6.6915622 21.4723115 -50.3796701
H 6.2622286 21.8728319 -49.4665579
C 6.2201094 20.031677 -50.5303368
O 6.4399128 19.4158451 -51.5664035
C 6.1943466 22.3420754 -51.5434893
H 6.7074003 22.060723 -52.4640156
C 4.6826819 22.2560971 -51.7615391
H 4.1511206 22.4652032 -50.8329504
H 4.408085 21.2625488 -52.117419
H 4.3832472 22.9787312 -52.5207642
O 6.4597851 23.693421 -51.2481149
H 6.0035097 23.908674 -50.423727
N 5.5750189 19.4858299 -49.5003679
H 5.4489048 20.0421054 -48.6609263
C 4.8934822 18.1915638 -49.5707935

H 5.3805521 17.5583986 -50.3075649
H 4.9561912 17.6920084 -48.6044006
C 3.4178491 18.3381219 -49.9451647
O 2.7499778 19.2457229 -49.4508098
N 2.8759735 17.4192718 -50.7447628
H 3.489359 16.728848 -51.172119
C 1.4218424 17.2768095 -50.9304672
H 0.9352328 17.8212694 -50.1261785
C 0.9560668 15.8305307 -50.7697592
O 1.6406314 14.8966804 -51.1867693
C 0.9136146 17.8905378 -52.2565359
H -0.1448096 17.6360649 -52.3435319
C 1.6387051 17.3248981 -53.5004349
H 2.6843131 17.6363338 -53.4916327
H 1.6106378 16.2363657 -53.4623999
C 0.9961788 19.4249524 -52.1858548
H 2.0365676 19.7521869 -52.1926341
H 0.5247676 19.7783184 -51.266841
H 0.4642133 19.8755679 -53.0218273
C 1.0052803 17.7519525 -54.8316522
H 1.1088672 18.8260369 -54.9781292
H -0.0515642 17.481976 -54.8436029
H 1.510511 17.2397663 -55.65089
N -0.2477491 15.6495421 -50.2260329
H -0.7465903 16.4668413 -49.8863459
C -0.9656272 14.3704334 -50.212057
H -0.6470092 13.784003 -51.0743246
C -2.4736961 14.6046495 -50.3361702
O -2.9923013 15.6108318 -49.855645
C -0.6345655 13.5626088 -48.9469876
H -1.035592 14.063649 -48.0667914
H 0.4483293 13.4718768 -48.8486547
O -1.2039403 12.2647033 -49.0661658
H -0.8230701 11.6633302 -48.3915009
N -3.189996 13.6624306 -50.9545782
H -2.693636 12.8593069 -51.3085801
C -4.6570956 13.7038033 -51.1109514
H -4.9453094 14.6681594 -51.5324737
C -5.3659256 13.5807981 -49.7505522
O -6.4750307 14.0756237 -49.5707605
C -5.0838402 12.5810381 -52.0946599
H -4.5868338 11.656628 -51.7934535
C -6.60297 12.3174183 -52.0927149
H -7.1427278 13.2367823 -52.3255887
H -6.9278242 11.951214 -51.1188279
H -6.8636599 11.5560362 -52.8261546

C -4.6265523 12.9470809 -53.5247128
H -5.2699191 13.744181 -53.8956398
H -3.6030739 13.3215718 -53.5000957
C -4.6555746 11.788384 -54.5300713
H -5.6777838 11.458972 -54.7113472
H -4.0625365 10.9539342 -54.1543668
H -4.2321241 12.1260136 -55.4769672
N -4.7271601 12.913471 -48.788543
H -3.775143 12.6272275 -48.9723444
C -5.2999645 12.5558781 -47.4919011
H -6.3581933 12.3426238 -47.6238231
C -5.12327 13.6827605 -46.4548777
O -3.9915082 13.9538372 -46.0403971
C -4.6452439 11.2446981 -47.0565748
H -3.587913 11.2809028 -47.3174804
H -5.1004324 10.4159779 -47.6001787
O -4.7366918 11.0128571 -45.6723862
H -5.6758702 11.0025864 -45.3613888
N -6.2181431 14.2734505 -45.9265974
C -6.1448261 15.2535177 -44.8392723
H -5.5037262 16.0768951 -45.1469994
C -5.5994107 14.6593758 -43.5365247
O -4.990551 15.3639187 -42.7414758
C -7.5767455 15.7683423 -44.6357585
H -8.056633 15.2366621 -43.8112928
H -7.5919532 16.8425494 -44.4489233
C -7.6005661 14.1249832 -46.3585443
H -7.6892693 14.0050037 -47.4364082
H -8.051605 13.2737373 -45.845865
C -8.2932888 15.4153499 -45.9362787
H -8.111043 16.1900229 -46.680453
H -9.3633916 15.2698172 -45.7853592
N -5.798928 13.3603836 -43.3244477
H -6.3334307 12.8472974 -44.0199305
C -5.3819279 12.5909009 -42.1533687
H -5.5086175 13.2194111 -41.2710266
C -3.889537 12.1953296 -42.1819728
O -3.2267231 12.2061379 -41.1417984
C -6.3412377 11.3905832 -41.9922851
H -7.3113502 11.7703061 -41.6675729
H -5.9575394 10.7498656 -41.1971473
C -6.5663179 10.5388814 -43.2538189
O -6.908747 11.078591 -44.3329274
O -6.3508923 9.3107568 -43.1812859
N -3.3008267 11.9683216 -43.3619274
H -3.8866016 11.9470974 -44.1933319

C -1.8430178 11.9831623 -43.5618261
H -1.3723892 11.257173 -42.9150736
C -1.2618897 13.3502526 -43.1982805
O -0.4067809 13.4546671 -42.3135894
C -1.4776396 11.6187214 -45.0103698
H -2.1218696 12.1563712 -45.7031098
H -0.4492902 11.9202012 -45.2048059
C -1.5826308 10.1109484 -45.2530965
H -2.542401 9.7508069 -44.8866851
H -0.8029225 9.6052696 -44.6769365
C -1.4265517 9.7523301 -46.7342513
O -2.1887154 8.9014817 -47.2378949
O -0.4387212 10.1870761 -47.3704002
N -1.8238071 14.3981459 -43.7998905
H -2.5652821 14.2183388 -44.4716944
C -1.3629664 15.7743681 -43.644951
H -0.2966211 15.7971953 -43.8619811
C -1.5011144 16.2671664 -42.1931932
O -0.5669668 16.8774303 -41.6842395
C -2.0937697 16.6432986 -44.6956286
H -3.1616424 16.4331521 -44.6208048
C -1.6453572 16.3062192 -46.1415492
H -1.8060816 15.2459139 -46.3228022
H -2.2857106 16.8442621 -46.8391984
C -1.9169274 18.1412204 -44.4519846
H -0.8624135 18.3648704 -44.4776171
H -2.351247 18.4235725 -43.4927015
H -2.4173335 18.7036014 -45.2397571
C -0.1850993 16.6247081 -46.5047295
H 0.502714 16.0852862 -45.8550906
H 0.005957 17.6948585 -46.4390428
H -0.0012286 16.3148047 -47.5317343
N -2.5671139 15.9256739 -41.4689525
H -3.3267024 15.4447242 -41.9411177
C -2.7735512 16.3191514 -40.0675664
H -2.7786111 17.4067291 -40.0129945
C -1.6436206 15.8404344 -39.1348382
O -1.1884872 16.6049745 -38.2795479
C -4.1712491 15.8244989 -39.6523631
H -4.8964345 16.3669418 -40.2600162
H -4.276831 14.7663523 -39.8970248
C -4.5547229 16.0502673 -38.1793378
H -5.6347657 16.1922723 -38.1337433
H -4.0852274 16.9595111 -37.8033673
C -4.2106249 14.87867 -37.2602295
O -3.478791 13.9624486 -37.5900405

N -4.7524033 14.838826 -36.0651956
H -4.5608134 14.0108061 -35.5281391
H -5.3835633 15.5549238 -35.7602172
N -1.1004638 14.6424705 -39.3708469
H -1.5013084 14.0764096 -40.106404
C 0.0418472 14.1322373 -38.6116889
H -0.0436926 14.4501244 -37.5713118
C 1.3850099 14.6518156 -39.1239408
O 2.2660308 14.8908551 -38.3035319
C 0.0435491 12.6104591 -38.6527647
H 1.0465421 12.2488474 -38.4138007
H -0.2262749 12.3017169 -39.6598596
S -1.1117245 11.9445062 -37.4241918
H -2.0986728 12.845829 -37.6105998
N 1.5575739 14.8170369 -40.4417058
H 0.8054631 14.5690666 -41.0762493
C 2.7701594 15.422793 -40.987119
H 3.6444812 14.9045317 -40.5883586
C 2.8696392 16.8863013 -40.5636503
O 3.5847836 17.2487937 -39.6283336
C 2.7907183 15.3258836 -42.5238653
H 1.9156234 15.7985274 -42.9702768
H 2.7822393 14.2994419 -42.8465516
H 3.7036186 15.7788831 -42.9060338
N 2.1002966 17.7180647 -41.2791163
H 1.1970245 17.3057366 -41.4945638
C 2.6117307 18.5517644 -42.3861192
H 3.1341734 17.8723516 -43.0516016
C 1.3970127 19.0600589 -43.2242848
H 1.358215 18.4262558 -44.0974844
H 0.5018309 18.6692998 -42.7733033
C 1.1166488 20.5727843 -43.5444475
H 1.2915895 21.151315 -42.6635274
H 1.8411256 20.9728381 -44.2381746
C -0.3230066 20.9360951 -43.9914732
H -1.0227159 20.3720331 -43.3859128
H -0.4699022 20.6178342 -45.0179328
C -0.6963571 22.4298742 -43.8025919
H 0.1995555 22.9755746 -43.4898631
H -1.4260614 22.4979107 -42.9928553
N -1.2635441 23.0865246 -45.012343
H -0.5895041 23.0906335 -45.7804691
H -1.4992876 24.0597524 -44.8902245
H -2.0834365 22.6400856 -45.4285437
C 3.7642777 19.4606364 -41.9383207
O 4.6934606 18.9814363 -41.3095185

N 3.6311882 20.7771193 -41.9989438
H 2.8876535 21.1417414 -42.5604266
C 4.0788258 21.6000937 -40.8865894
H 5.1237588 21.3834285 -40.6673247
C 3.9999911 23.0820359 -41.3327063
H 3.0493725 23.2473995 -41.8409777
H 4.7944901 23.2717171 -42.0553756
C 4.1026739 24.1179475 -40.2399803
N 3.0918861 25.0331692 -39.9409579
C 3.4223691 25.5589047 -38.7527186
H 2.8010558 26.2448663 -38.192991
N 4.5752577 25.0358408 -38.3042514
H 4.8930887 25.1113105 -37.3429746
C 5.0364284 24.1390869 -39.2428601
H 5.8685618 23.4582498 -39.1352582
C 3.2420288 21.199852 -39.6385361
O 3.0797663 22.025223 -38.7500341
N 2.530232 20.0383135 -39.640792
H 2.865367 19.2416506 -40.1623869
C 1.4823966 19.8353562 -38.7041374
H 1.2481924 20.7922532 -38.2384218
C 2.0858318 19.0933861 -37.5361898
O 3.067777 19.6165891 -37.0050026
C 0.1413169 19.4928001 -39.3442083
H 0.1442248 18.4588372 -39.6817178
H -0.6141341 19.569777 -38.5611991
C -0.267998 20.4123699 -40.5175301
H 0.1183068 19.9509151 -41.3678312
C -1.7776922 20.4353344 -40.70414
H -2.2492346 20.8376265 -39.8066394
H -2.1313518 19.4171078 -40.8620015
H -2.0806715 21.041301 -41.5528518
C 0.1944014 21.8747087 -40.5369619
H 1.2344464 22.003532 -40.7787416
H 0.035205 22.3050513 -39.5470494
H -0.3843236 22.4742456 -41.2283432
N 1.4878269 18.020571 -37.0275607
H 0.680745 17.619781 -37.490007
C 1.8370793 17.567107 -35.682473
H 1.6480608 18.3923588 -34.994206
C 3.3255548 17.1909504 -35.5105984
O 3.975249 17.6990186 -34.5966676
C 0.897452 16.4265003 -35.2938252
H 1.0844826 15.5580817 -35.9233497
H -0.1416476 16.7397962 -35.4098183
H 1.0723946 16.150676 -34.2533891

N 3.895938 16.3935091 -36.4204664
H 3.3273487 16.0340792 -37.1822979
C 5.2891756 15.9425756 -36.3358269
H 5.4806631 15.6036171 -35.3165186
C 6.3276383 17.0397494 -36.6405866
O 7.4910534 16.9063308 -36.2670223
C 5.4618095 14.744717 -37.274927
H 5.2733635 15.0489023 -38.3063018
H 4.7617187 13.9547081 -37.0015401
H 6.47971 14.3605827 -37.1961688
N 5.9285687 18.1328717 -37.2982429
H 4.962632 18.1795496 -37.6016175
C 6.8314934 19.2418052 -37.635999
H 7.8429605 18.8505207 -37.6832907
C 6.844892 20.3829303 -36.6121256
O 7.5699517 21.3645417 -36.8076774
C 6.4883151 19.7654549 -39.0249614
H 5.4763035 20.1549587 -38.9918722
H 6.5452826 18.940506 -39.7349417
O 7.3909096 20.7802827 -39.4271918
H 7.5897174 21.3108912 -38.6363837
N 6.0301942 20.3258281 -35.5583977
H 5.4633655 19.4990513 -35.4131305
C 5.9102691 21.4436743 -34.6242259
H 5.8569889 22.3420848 -35.2375565
C 7.1639527 21.593333 -33.7492034
O 7.6308825 20.6298769 -33.149975
C 4.6035516 21.3149437 -33.8160339
H 3.9610644 20.5586151 -34.2705869
H 4.8261419 20.983383 -32.8007128
C 3.8146552 22.6339158 -33.7630863
H 4.3761121 23.361526 -33.1761685
H 2.8597389 22.454182 -33.2696363
C 3.5503843 23.2164719 -35.1512038
O 4.0029425 24.3035214 -35.4804174
N 2.9488639 22.4629638 -36.0411139
H 2.950617 22.7735293 -37.0102779
H 2.7918841 21.4817724 -35.8719266
N 7.7210904 22.8081187 -33.6914249
H 7.2987774 23.5567177 -34.2190106
C 8.9972279 23.0788989 -33.0131334
H 9.0701395 24.1482844 -32.8145984
H 9.0214978 22.5526099 -32.0586359
C 10.2493908 22.6746477 -33.805537
O 11.3286693 22.5814544 -33.2268775
N 10.1271567 22.3925806 -35.1093067

H 9.2142375 22.4390954 -35.5407744
C 11.2828804 22.1739636 -35.9765919
H 12.1470555 21.9182832 -35.3639204
C 11.6311631 23.4425166 -36.7703543
O 10.8468171 23.8739616 -37.6062429
C 11.0248787 21.0051895 -36.9419587
H 10.2282802 21.2982624 -37.627223
H 11.9359448 20.8776523 -37.5163873
C 10.6521745 19.6372191 -36.3484215
H 9.6886744 19.7031776 -35.8480046
C 10.554009 18.6256642 -37.4922608
H 11.5421073 18.4392663 -37.908598
H 9.9033941 19.0025286 -38.2786216
H 10.1402736 17.6890333 -37.1141196
C 11.700169 19.1190278 -35.363519
H 12.6761902 19.0615675 -35.8409176
H 11.4049478 18.1266817 -35.0192001
H 11.7434391 19.7788958 -34.4960605
N 12.8948299 23.8661966 -36.7091582
H 13.3352564 23.7065076 -35.812184
C 13.4155394 25.0179326 -37.4790076
H 12.6681205 25.8145446 -37.4282954
C 13.6732908 24.7387133 -38.9836209
O 14.3679382 25.4938641 -39.6582383
C 14.7076514 25.5493859 -36.8171534
H 15.5330047 24.8870948 -37.0856383
H 14.9254605 26.5347873 -37.2302733
C 14.686988 25.6625318 -35.2845744
H 15.5866248 26.1902592 -34.9596257
H 14.7320807 24.6561241 -34.8586402
C 13.4378158 26.3886491 -34.7758366
O 13.4280188 27.6368107 -34.819975
O 12.4900572 25.6633847 -34.4019596
N 13.2648524 23.5772005 -39.5105643
H 12.5354699 23.085907 -39.0068225
C 13.5570177 23.1795007 -40.8976494
H 14.5254876 23.6009869 -41.1761233
C 12.542468 23.7627457 -41.8860466
O 11.3445727 23.7642826 -41.6133993
C 13.6717175 21.6505914 -41.0511484
H 14.6737933 21.3409764 -40.7633607
H 13.5409867 21.3814689 -42.0977122
C 12.6807254 20.8409029 -40.234541
O 13.0819143 19.9571612 -39.4998137
N 11.4063889 21.1652132 -40.2593715
H 10.7663929 20.6171536 -39.7086994

H 11.0835354 21.9909526 -40.7547637
N 13.0133136 24.1123106 -43.0888082
H 14.003369 23.9861169 -43.252692
C 12.2559559 24.8063065 -44.14478
H 11.634147 25.5703974 -43.6725806
C 11.2900693 23.8636724 -44.8836361
O 11.4794884 23.5296958 -46.0528063
C 13.2402964 25.5207442 -45.0935189
H 13.7974318 24.7585126 -45.6299273
H 12.6747639 26.1075278 -45.8195301
C 14.2367 26.4457212 -44.363787
H 13.6782029 27.2076148 -43.8182901
H 14.8081668 25.8651489 -43.6389009
C 15.2530414 27.1445263 -45.2802328
H 15.9662655 27.6665972 -44.6388995
H 14.7410762 27.8844194 -45.8974409
C 16.0017683 26.1590706 -46.1895801
H 15.4383225 26.0365783 -47.1185131
H 16.0485386 25.1791219 -45.7028254
N 17.3833513 26.6013149 -46.4784672
H 17.4501689 27.5257558 -46.8646949
H 17.8394313 25.9286781 -47.0934372
H 17.9672793 26.5200239 -45.6378488
N 10.2586788 23.4106932 -44.1767312
H 10.1841574 23.7178249 -43.2124954
C 9.2046284 22.5279462 -44.6902692
H 9.4646982 22.2318761 -45.7066837
C 7.8664666 23.2561846 -44.773848
O 7.5173305 24.0732073 -43.9268036
C 9.0728533 21.2239852 -43.8778855
H 8.316634 20.6091786 -44.3642886
C 10.3766276 20.4230278 -43.8853184
H 11.1563743 20.9672999 -43.3545985
H 10.6901088 20.2389319 -44.9118237
H 10.2204614 19.4614051 -43.3955968
C 8.6444678 21.4437578 -42.4225151
H 9.3296493 22.1190396 -41.9159669
H 8.6253233 20.491972 -41.8909169
H 7.6444986 21.8751438 -42.3818422
N 7.0848025 22.9020958 -45.7835561
H 7.4542566 22.2468272 -46.4660422
C 5.688535 23.2901281 -45.9603944
H 5.2592379 23.5642908 -44.9971347
C 4.9368154 22.090787 -46.5162855
O 5.5175932 21.2799915 -47.2402207
C 5.572196 24.4861637 -46.9093264

H 6.1315988 25.3254024 -46.4927753
H 4.5253632 24.7775338 -47.0057321
O 6.0920421 24.166681 -48.1936715
H 6.9019288 23.6412406 -48.069159
N 3.6649475 21.9328174 -46.1572492
H 3.1924113 22.6509743 -45.6290061
C 2.8540294 20.891737 -46.7718369
H 3.2603949 20.7420939 -47.768523
C 1.4232672 21.3469321 -46.9993605
O 0.9655742 22.2461308 -46.2958749
C 2.8723023 19.5311039 -46.0727033
H 1.8998189 19.3593649 -45.6756641
H 2.831625 18.8431272 -46.9167144
C 4.0238134 18.9905664 -45.2169582
C 4.3071869 17.6253821 -45.3860567
H 3.6888653 17.0224533 -46.0309537
C 4.8919789 19.744078 -44.3979068
H 4.7530479 20.7996534 -44.2590226
C 5.3992881 17.0235059 -44.752768
H 5.5957283 15.9801922 -44.9285487
C 5.9619852 19.1264943 -43.7177794
H 6.5828412 19.700868 -43.0462872
C 6.2154413 17.761582 -43.8888187
H 7.0295482 17.2877361 -43.3583586
N 0.7049211 20.6843602 -47.9071269
H 1.1580329 19.9917536 -48.4999263
C -0.7426341 20.8387991 -48.0115702
H -1.0824204 20.9641876 -46.9898499
C -1.5361428 19.617803 -48.4852348
O -0.9924186 18.6595985 -49.0384563
C -1.11455 22.1121096 -48.8003637
H -1.7026334 21.8618073 -49.6844732
H -0.2187602 22.6483505 -49.119513
C -1.9431406 22.9948283 -47.8556361
O -2.8487189 22.4398679 -47.1824651
O -1.4945848 24.1167787 -47.540539
N -2.8489941 19.6723543 -48.2388324
H -3.2118883 20.5386921 -47.8451067
C -3.8093695 18.7281656 -48.8164363
H -3.3973647 17.7249245 -48.7310293
C -3.9734597 19.0752603 -50.2936677
O -4.3973474 20.175993 -50.6415666
C -5.1732061 18.7435119 -48.0992768
H -5.6730347 19.6950951 -48.2825975
C -6.0637773 17.6068255 -48.6193746
H -5.6052813 16.639463 -48.4108445

H -6.2151491 17.7031978 -49.6940242
H -7.0395154 17.6576898 -48.1394967
C -5.0359728 18.5600012 -46.5826229
H -4.4763312 17.6507279 -46.373113
H -6.0235974 18.4938287 -46.1264681
H -4.5200769 19.4177832 -46.1531065
N -3.6215795 18.1436875 -51.1741785
H -3.2879382 17.2507786 -50.8282856
C -3.6115106 18.3903012 -52.6077545
H -4.5005105 18.9612831 -52.879691
H -2.7342712 18.9845956 -52.864576
C -3.5764643 17.1062873 -53.4192378
O -2.760003 16.2192524 -53.1788795
N -4.4347018 17.0516582 -54.4326559
H -5.0732818 17.8204822 -54.5581563
C -4.3864658 16.05468 -55.4956002
H -4.0915978 15.0912157 -55.0883004
C -3.3390088 16.4662952 -56.5310674
O -3.4495113 17.5140222 -57.164842
C -5.7890218 15.9175663 -56.0942179
H -6.1264015 16.9003061 -56.4288134
H -6.4684305 15.5787826 -55.3090581
C -5.884318 14.9468145 -57.2730016
O -4.8383242 14.4918336 -57.7921059
O -7.0322555 14.7422459 -57.7062923
N -2.3237965 15.6294866 -56.7222922
H -2.3348741 14.7515165 -56.2290766
C -1.1959614 15.917138 -57.5968154
H -0.8042434 16.9026035 -57.3349828
C -1.5477935 15.9520276 -59.0940706
O -0.7986989 16.5414193 -59.8719076
C -0.1261968 14.8713849 -57.2991538
H -0.4332321 13.8964363 -57.6832249
H 0.0211865 14.8034629 -56.2217764
H 0.8044678 15.171372 -57.7763393
N -2.6894224 15.3899282 -59.5099429
H -3.3057519 14.9434825 -58.8268714
C -3.1680469 15.5081525 -60.8889374
H -2.2977069 15.4764396 -61.5426236
C -3.8714604 16.8578481 -61.1353749
O -3.8881519 17.3427345 -62.2717246
C -4.0589673 14.2899356 -61.2019821
H -3.5800509 13.3944349 -60.8022451
H -5.0094949 14.4145144 -60.6792547
C -4.3336983 14.054691 -62.7013104
H -4.8066073 14.9365606 -63.1323418

C -3.0607759 13.732287 -63.4923323
H -2.5395826 12.8897156 -63.037984
H -2.3987076 14.5952794 -63.5160859
H -3.3198034 13.4669686 -64.5171777
C -5.2831101 12.8667808 -62.868187
H -4.8269732 11.9632398 -62.4652122
H -5.5106869 12.7195113 -63.9246257
H -6.2135557 13.0676634 -62.3361046
N -4.3438173 17.5317856 -60.0767708
H -4.239743 17.0913676 -59.1689351
C -5.1392953 18.7670931 -60.1508346
H -5.3101043 19.014173 -61.1954728
C -4.4652452 20.0192241 -59.553186
O -4.7247452 21.1197762 -60.036646
C -6.5275032 18.4864972 -59.5562995
H -7.1041324 19.4099779 -59.5149897
H -6.4332717 18.0886791 -58.546576
C -7.2894454 17.5033969 -60.4354308
O -7.5563357 17.7986585 -61.5883171
N -7.5704357 16.2932746 -60.0008698
H -8.0536807 15.6856149 -60.632247
H -7.3300548 15.9419432 -59.0695682
N -3.5291447 19.8666245 -58.6129897
H -3.3778296 18.9328279 -58.2480178
C -2.7161973 20.948246 -58.0426512
H -3.3974747 21.6988151 -57.6398965
C -1.8381347 21.6244684 -59.108357
O -1.3603305 20.9679296 -60.0376461
C -1.8708616 20.3801509 -56.8857777
H -1.2155043 19.5960797 -57.2688558
H -2.5403692 19.9339514 -56.1488536
C -1.0113161 21.4298223 -56.1713507
H -0.2443703 21.7912918 -56.8567282
H -0.5059511 20.9491715 -55.3327881
S -1.9402459 22.8506813 -55.5394919
C -0.5701828 23.8292679 -54.8690719
H -0.9613234 24.74235 -54.4197375
H 0.1193026 24.0926016 -55.6715556
H -0.0410525 23.252321 -54.1098706
N -1.6015408 22.9369568 -58.9811233
H -1.9423219 23.4054305 -58.1486339
C -0.8192637 23.7390643 -59.9395053
H -0.2576772 23.064157 -60.5851844
C 0.2282464 24.5957342 -59.2270807
O 0.004776 25.0706294 -58.1183271
C -1.7509353 24.6002576 -60.822006

H -2.2485006 25.3361505 -60.1875853
H -1.1470927 25.1432575 -61.5512182
C -2.8355023 23.8041089 -61.576147
H -3.4692309 24.5095959 -62.1148217
H -3.4670243 23.293783 -60.85025
C -2.285634 22.7794283 -62.5838831
H -1.469829 22.2166229 -62.1362081
H -1.8727529 23.3160784 -63.440655
N -3.3480238 21.8600974 -63.0454371
H -4.1592275 22.2924263 -63.4494333
C -3.3712553 20.539867 -62.9367282
N -4.3632524 19.8552986 -63.436589
H -4.394121 18.8635073 -63.2050823
H -5.2093213 20.3214733 -63.7056537
N -2.4470361 19.8412767 -62.3456859
H -1.8072108 20.3142319 -61.7116381
H -2.5589807 18.8412342 -62.2636225
N 1.3477479 24.8147152 -59.9086339
H 1.3980397 24.4714973 -60.8561952
C 2.4801481 25.6382775 -59.4817666
H 2.171737 26.310206 -58.6796175
C 2.9540436 26.4820188 -60.6742369
O 2.5493109 26.2124622 -61.8048482
C 3.6101897 24.7291168 -58.9715134
H 3.9556652 24.0994097 -59.7921348
H 4.4519802 25.3525014 -58.6710504
C 3.2414508 23.8502424 -57.790578
C 3.2255769 24.3988016 -56.4940977
H 3.4578802 25.4439975 -56.3432923
C 2.944317 22.484486 -57.9805326
H 2.957988 22.0603603 -58.9749413
C 2.9118273 23.5871558 -55.3886793
H 2.893409 23.9989979 -54.390721
C 2.6376701 21.6672874 -56.8746023
H 2.4174114 20.6212776 -57.0198648
C 2.6196309 22.2218223 -55.5761608
O 2.3179465 21.4526177 -54.4997302
H 2.18435 20.5319836 -54.724826
N 3.8334992 27.4628261 -60.4509498
H 4.2082822 27.584594 -59.5219213
C 4.4939213 28.1620494 -61.5616058
H 3.7309422 28.5488971 -62.2384471
C 5.3882736 27.2020676 -62.3485732
O 6.0176283 26.3114741 -61.7616727
C 5.3217809 29.3477141 -61.0436929
H 6.091829 28.9980095 -60.3569351

H 4.6612574 30.029898 -60.5071673
O 5.9393037 30.0538829 -62.1032955
H 6.7243398 29.5613144 -62.4093461
N 5.5820288 27.4940815 -63.6317522
H 5.0106585 28.1958685 -64.0746187
C 6.7307802 26.9874338 -64.3760478
H 6.6366855 25.9085016 -64.4254029
C 8.028407 27.3558119 -63.6304401
O 8.1187891 28.4400932 -63.0417593
C 6.7479946 27.5450176 -65.8100117
H 7.719706 27.3149662 -66.2505881
H 6.6485008 28.6315395 -65.7731873
C 5.6672195 26.9697377 -66.7358325
O 4.772257 26.2412801 -66.2508656
O 5.7522914 27.2543315 -67.9504335
N 9.0046514 26.4438149 -63.5827792
H 8.8767 25.5757138 -64.0999271
C 10.2928452 26.6834173 -62.9168859
H 10.9628395 25.863104 -63.1596956
H 10.726691 27.5994604 -63.3176861
C 10.2581107 26.8168125 -61.3836782
O 11.1929088 27.3706979 -60.812188
N 9.2068629 26.3463782 -60.7000323
H 8.4713915 25.8971 -61.2280263
C 9.0731089 26.4374344 -59.2283582
H 9.1851045 27.4791121 -58.9258279
C 10.1069491 25.6136815 -58.4432941
O 10.4836519 25.994552 -57.3288833
C 7.6897912 25.9495096 -58.7796666
H 7.6527518 25.9341602 -57.6888603
H 7.5223822 24.9374057 -59.148062
O 6.6580577 26.7898418 -59.2502181
H 6.5249451 26.5892104 -60.1998395
N 10.5092653 24.4553591 -58.9775317
H 10.148396 24.2047745 -59.8925566
C 11.2703098 23.4359545 -58.2511685
H 11.53296 23.831252 -57.2725145
C 12.5826537 23.0539197 -58.9343996
O 12.7237566 23.0892687 -60.153064
C 10.4016771 22.1945978 -58.0069762
H 10.1678104 21.7295215 -58.9665107
H 10.9725225 21.4720044 -57.4234963
C 9.1174901 22.4918067 -57.2616053
C 9.1440574 22.7882535 -55.8861933
H 10.0770687 22.7663086 -55.3446341
C 7.8965917 22.5106125 -57.9559699

H 7.8822044 22.2996273 -59.0114969
C 7.9532069 23.1130515 -55.2127899
H 7.9771022 23.3415634 -54.1554797
C 6.7052395 22.8277978 -57.284616
H 5.7704859 22.841606 -57.8236995
C 6.7337942 23.135257 -55.9133376
H 5.8189192 23.3828901 -55.3936062
N 13.517583 22.5839715 -58.1188466
H 13.3072881 22.5350507 -57.1278463
C 14.8179288 22.0601079 -58.5444263
H 15.0314648 22.3351386 -59.5799209
C 14.8820061 20.5283609 -58.4571595
O 15.7122647 19.8905221 -59.1081171
C 15.861854 22.6970743 -57.6264017
H 15.4958654 22.6849266 -56.6000881
H 16.7664742 22.1004187 -57.6568063
C 16.1785143 24.1454572 -58.0345006
H 16.4935678 24.6931106 -57.143149
H 15.2907343 24.6425045 -58.4312527
C 17.305602 24.1476694 -59.067901
O 17.1190339 23.5958259 -60.1692975
O 18.44763 24.4781666 -58.6847995
N 14.0008482 19.9431131 -57.6415347
H 13.375073 20.5442361 -57.11928
C 13.8557196 18.5135042 -57.3961515
H 14.3000178 17.9505765 -58.2132188
C 12.3562485 18.1809218 -57.2916426
O 11.6055412 18.9392478 -56.6695756
C 14.5613807 18.1437631 -56.0764476
H 13.9703084 18.5856413 -55.2756477
H 14.5331341 17.0604963 -55.9537592
C 16.0075358 18.6382877 -55.8749913
H 16.0774433 19.7022195 -56.0961422
C 16.4272853 18.4705187 -54.4170553
H 16.4803628 17.4129 -54.1558347
H 15.7119752 18.972793 -53.7650525
H 17.3987076 18.9361346 -54.2814031
C 17.0207935 17.8886108 -56.7338335
H 16.9321508 16.814618 -56.5697651
H 18.0321975 18.2073195 -56.4892937
H 16.8337156 18.1120355 -57.7796173
N 11.9268476 17.0254245 -57.802589
H 12.5667476 16.4679935 -58.3580903
C 10.6134922 16.4408486 -57.4727742
H 10.200647 16.9817986 -56.6240596
C 10.8460628 14.9974708 -57.0338666

O 11.4968357 14.2196056 -57.7343886
C 9.584129 16.5453119 -58.623281
H 9.8891 15.8453796 -59.384544
C 9.5544193 17.9798466 -59.2141413
H 9.2019235 18.6905423 -58.4697116
H 10.5636784 18.2729503 -59.500348
C 8.1844474 16.0604285 -58.2049168
H 7.7911132 16.6907992 -57.4099126
H 8.2346799 15.0332518 -57.8434264
H 7.5072373 16.083636 -59.0554359
C 8.7221937 18.1222675 -60.4791129
H 7.6661354 18.1114382 -60.2270789
H 8.9737782 17.3097286 -61.1536164
H 8.9770545 19.0592215 -60.9698104
N 10.3564225 14.6574374 -55.8469035
H 9.7941865 15.3397182 -55.3446681
C 10.5610505 13.361721 -55.2088406
H 11.1527113 12.721799 -55.8623133
C 9.2265463 12.6740554 -54.9607565
O 8.2716464 13.3041168 -54.5053329
C 11.33476 13.5316808 -53.9000093
H 12.3374437 13.8914798 -54.1303004
H 10.8329836 14.2807638 -53.2847405
C 11.4560679 12.2414929 -53.107116
C 12.2584577 11.1893237 -53.5913009
H 12.8093437 11.3079028 -54.5115507
C 12.3285074 9.9712448 -52.892086
H 12.935959 9.1595666 -53.2719667
C 11.6002187 9.80203 -51.7022888
H 11.6597189 8.8679635 -51.1608939
C 10.7885537 10.8448729 -51.2227153
H 10.2173261 10.7141468 -50.3149552
C 10.7115946 12.0603749 -51.9250526
H 10.0647863 12.8469327 -51.5572318
N 9.1602094 11.3777451 -55.2613215
H 9.967209 10.9333375 -55.6892098
C 7.9328766 10.5948167 -55.1189933
H 7.301889 11.0701356 -54.3646456
C 8.255057 9.1831792 -54.6477118
O 9.0891373 8.509012 -55.2435789
C 7.1597415 10.5440188 -56.4471444
H 7.7114147 9.9222563 -57.1494066
C 5.8095002 9.8629732 -56.2113163
H 5.4896513 10.0212756 -55.1882349
H 5.9140703 8.7885214 -56.3956002
H 5.0585275 10.2781186 -56.8705113

C 7.0334598 11.9183157 -57.1168015
H 6.7794035 12.6694729 -56.3779092
H 6.2773547 11.9109892 -57.8926942
H 7.9869524 12.1928344 -57.5734035
N 7.5304832 8.6922801 -53.6430702
H 6.8760449 9.3036927 -53.1726615
C 7.6681163 7.3252057 -53.1358409
H 8.2068826 6.7406371 -53.8801015
C 6.2876119 6.6673452 -52.9862792
O 5.4097723 7.1751835 -52.2984167
C 8.5623854 7.3067255 -51.8731256
H 9.5579647 7.6240205 -52.1890165
C 8.094785 8.2976582 -50.7891014
H 7.0661867 8.0877915 -50.5071326
H 8.1462975 9.3195321 -51.1659953
H 8.7354529 8.2405992 -49.9112774
C 8.6896144 5.865025 -51.3446308
H 7.7397634 5.5693468 -50.9079072
H 8.8984866 5.198318 -52.1818034
C 9.804598 5.6643442 -50.3096542
H 9.5898272 6.2220642 -49.3977556
H 10.7584064 5.9929064 -50.7211156
H 9.8773923 4.6049462 -50.0589894
N 6.0735872 5.5791504 -53.7322323
H 6.8322139 5.3057652 -54.3480851
C 4.8587987 4.7465541 -53.7745885
H 4.9169297 4.1923917 -54.7089597
C 3.516118 5.5057891 -53.8614048
O 2.5191161 5.1118456 -53.2532807
C 4.8607276 3.7088065 -52.6438668
H 4.6254463 4.2273965 -51.7263386
H 4.0580549 2.9937707 -52.8329263
C 6.1573545 2.9237375 -52.4338794
H 5.9667513 2.1123083 -51.7318169
H 6.9089625 3.5794581 -51.9982039
C 6.7097361 2.3381834 -53.7201912
O 7.8204849 2.6367907 -54.124086
N 5.9596121 1.5408023 -54.4434662
H 6.3860314 1.1601171 -55.2659575
H 4.9443976 1.5731702 -54.3602218
N 3.4922299 6.609794 -54.6151327
H 4.3734201 6.9183661 -54.9925454
C 2.3102518 7.4619524 -54.7883182
H 1.523502 7.1422596 -54.1042785
C 1.7272705 7.3825057 -56.209906
O 0.5213259 7.2247492 -56.3878293

C 2.7044591 8.890572 -54.3960494
H 3.4554001 9.2689719 -55.0839756
H 3.1257726 8.8895343 -53.3873539
S 1.267988 9.9956678 -54.455393
H 1.9581013 11.1116626 -54.1764562
N 2.5785643 7.3956426 -57.2408166
H 3.5667076 7.4519611 -57.0465178
C 2.1412431 7.4998195 -58.6453044
H 1.3706166 8.2684934 -58.6961616
C 1.455654 6.224998 -59.1639819
O 0.6846533 6.2812467 -60.1185169
C 3.3271499 7.9979964 -59.498981
H 4.2271993 7.4604443 -59.1935633
C 3.5120954 9.5030321 -59.2234801
H 2.7433385 10.070109 -59.7525158
H 3.3888159 9.7067547 -58.1618482
C 3.1360484 7.7830679 -61.0060698
H 2.1735433 8.1823495 -61.3276595
H 3.2063335 6.7193164 -61.2171017
H 3.9295564 8.2650958 -61.5743967
C 4.8911054 10.0097662 -59.6504475
H 5.0099305 9.9628237 -60.7314375
H 5.6735898 9.4236341 -59.1685517
H 4.9797716 11.0455733 -59.3443017
N 1.5717582 5.1041732 -58.4564899
H 2.1827045 5.1206104 -57.6529454
C 0.7328439 3.9149034 -58.651388
H 0.8947388 3.5462889 -59.6639284
C -0.7823536 4.1812482 -58.5331897
O -1.5480582 3.5308124 -59.2486143
C 1.1982313 2.8047712 -57.6992863
H 0.5310809 1.9524817 -57.8022436
H 2.1807305 2.4791663 -58.0396875
C 1.2976814 3.1948038 -56.2119626
H 0.3012642 3.1688227 -55.7657597
H 1.6944627 4.205053 -56.0979221
C 2.2384522 2.2254211 -55.4939914
O 3.4720441 2.4194285 -55.590334
O 1.7426194 1.2099946 -54.9540701
N -1.2002262 5.2068174 -57.7761104
H -0.5129244 5.7123035 -57.2254629
C -2.598939 5.6605511 -57.6700557
H -3.2650489 4.7962077 -57.7183515
C -3.0249164 6.6303356 -58.7861373
O -4.212663 6.7435325 -59.0897442
C -2.8084363 6.3407486 -56.3074555

H -3.8525083 6.6461592 -56.2269157
H -2.1951503 7.2411331 -56.2540386
C -2.494111 5.4648105 -55.1222584
N -3.2748369 4.3995001 -54.6813414
C -1.402258 5.5724542 -54.3123075
H -0.6083186 6.2963122 -54.4092074
C -2.6439835 3.8919381 -53.61534
H -2.9894624 3.0311948 -53.0587202
N -1.5076655 4.5702295 -53.3754836
H -0.8044458 4.3204002 -52.6953078
N -2.0794667 7.3391896 -59.4136958
H -1.1158875 7.1556197 -59.1719981
C -2.3616997 8.3287098 -60.4667779
H -3.2276859 8.9173499 -60.1602128
C -2.7464924 7.5875773 -61.7548098
O -2.001621 6.7304906 -62.2333305
C -1.169923 9.3091347 -60.6337605
H -0.2399088 8.7535995 -60.5132491
C -1.1908147 10.461597 -59.5993055
H -0.3002541 11.0742561 -59.749503
H -2.0610367 11.0934907 -59.7853356
C -1.1248122 9.968303 -62.0256172
H -2.0519598 10.5058078 -62.2273489
H -0.9656117 9.221869 -62.8030621
H -0.2929621 10.6711498 -62.0761009
C -1.2165483 10.0459058 -58.1225757
H -0.4028053 9.3553923 -57.9105932
H -2.1699243 9.5788426 -57.8756608
H -1.0975652 10.9321865 -57.4986943
N -3.9191623 7.9002296 -62.3131057
H -4.4943561 8.5906527 -61.8567087
C -4.4811881 7.1647627 -63.4544377
H -4.2521094 6.1048541 -63.3300885
C -3.8496852 7.5951677 -64.7889643
O -3.2727482 6.7640869 -65.4867586
C -6.0132403 7.3088003 -63.4796484
H -6.2694267 8.3537053 -63.6622205
H -6.4039576 6.7184002 -64.3096062
C -6.7166646 6.8608593 -62.1861027
H -7.7887787 7.0203155 -62.3018223
H -6.3872767 7.4812947 -61.3521355
C -6.4970237 5.3889815 -61.8459751
O -6.9995508 4.4963847 -62.5114945
N -5.7572628 5.0902401 -60.8042215
H -5.833909 4.14822 -60.4307188
H -5.3482376 5.8097848 -60.2156199

N -3.8569681 8.8981386 -65.0946658
H -4.3202539 9.5439063 -64.4785248
C -3.1401507 9.4583446 -66.2477274
H -3.0411782 8.6833109 -67.0107969
C -1.7236527 9.8698317 -65.8282051
O -1.4618108 10.9868417 -65.364135
C -3.9305454 10.5951731 -66.9132344
H -4.0322571 11.4261952 -66.2131145
H -4.9304066 10.2295566 -67.1538309
C -3.2659619 11.0967388 -68.2087631
O -2.0957811 10.7292587 -68.4770125
O -3.9030893 11.919111 -68.8986723
N -0.803794 8.9111941 -65.9612056
H -1.116004 8.0117478 -66.302686
C 0.6148988 9.1220086 -65.6716635
H 0.701243 9.6227611 -64.7065084
C 1.2656105 10.0741342 -66.6689175
O 2.098196 10.8648421 -66.2441968
C 1.3529267 7.7757301 -65.5926566
H 1.1126767 7.1689474 -66.4679944
H 2.4297305 7.955249 -65.5762918
C 0.9574017 7.0366377 -64.3062496
H 1.2856483 7.6316669 -63.4540904
H -0.1280695 6.9419732 -64.2625537
C 1.5717334 5.633152 -64.2128231
H 2.6407791 5.6991469 -64.4166036
H 1.1128656 4.9710833 -64.9484462
C 1.3819517 5.0770151 -62.7971826
H 1.7874912 5.8138986 -62.1066302
H 1.9538723 4.1524789 -62.6816544
N -0.0351696 4.8430507 -62.4374898
H -0.6226182 5.656999 -62.6128357
H -0.1159741 4.6848988 -61.4413045
H -0.410129 4.0257107 -62.9061462
N 0.857613 10.1083443 -67.9375912
H 0.0306158 9.5949737 -68.2147783
C 1.4627276 11.0419099 -68.8908006
H 2.5495178 10.9536548 -68.8456466
C 1.0955327 12.4971915 -68.5945729
O 1.9570158 13.368256 -68.6695282
C 1.0240044 10.7936445 -70.3246517
H -0.0383867 11.0268319 -70.4208665
H 1.5987591 11.5133366 -70.9013528
C 1.2870139 9.4334093 -70.9591885
H 0.6207275 8.6875652 -70.5204886
H 2.321262 9.1555211 -70.7495981

C 1.0670261 9.5267931 -72.4818624
O 1.5837226 8.6346019 -73.1863309
O 0.449561 10.5278113 -72.9382914
N -0.1651418 12.7909911 -68.2489103
H -0.8538143 12.0316837 -68.2624408
C -0.597046 14.1400191 -67.8500139
H -0.3606733 14.8368522 -68.6493023
C 0.1788132 14.6128788 -66.6331437
O 0.6875716 15.7306213 -66.6664212
C -2.1126775 14.1466946 -67.5941196
H -2.6227997 14.0571914 -68.5530931
H -2.3576696 13.272807 -66.9910062
C -2.6896842 15.3612165 -66.8459386
H -3.7652708 15.2087699 -66.7430743
H -2.2667664 15.3935762 -65.8416272
C -2.4521123 16.7168604 -67.5216241
H -2.901966 16.7154243 -68.5155877
H -1.3820194 16.9074539 -67.6060141
C -3.0951271 17.8042082 -66.6541248
H -2.7560098 17.6658543 -65.6223159
H -4.1813943 17.6762267 -66.6795216
N -2.71647 19.159312 -67.1134731
H -1.700728 19.2804625 -67.0277463
H -3.1123092 19.8764223 -66.5258363
H -2.9676407 19.3212056 -68.075571
N 0.3204954 13.766832 -65.6119406
H -0.1301998 12.8600258 -65.6626692
C 1.1372004 14.0988161 -64.4489276
H 0.7681291 15.0311553 -64.0186157
C 2.6010132 14.3488103 -64.8440332
O 3.1203401 15.4231067 -64.5515805
C 1.0048208 13.0052398 -63.3833035
H -0.0274297 12.9715737 -63.0338871
H 1.2416141 12.0362933 -63.8251743
C 1.9196494 13.2544928 -62.2026704
C 1.5902525 14.2330469 -61.2449909
H 0.6507624 14.7645696 -61.3091331
C 2.5054578 14.5589645 -60.2283914
H 2.2645 15.3361693 -59.5185997
C 3.7477584 13.9046379 -60.163313
H 4.4610224 14.1635952 -59.3939693
C 4.075634 12.9324819 -61.1222156
H 5.0420339 12.4538226 -61.0965527
C 3.1634757 12.6011592 -62.1355357
H 3.4384687 11.8742898 -62.8856969
N 3.2219607 13.4431795 -65.6129092

H 2.7236004 12.5935768 -65.8627977
C 4.6285742 13.5613066 -66.0272132
H 5.2190656 13.6913037 -65.136371
C 4.8554892 14.8103973 -66.9071792
O 5.7538651 15.5982238 -66.6248052
C 5.1469566 12.2423806 -66.6375441
H 4.5264298 12.0049949 -67.503892
C 6.6112232 12.3569321 -67.0848273
H 7.294004 12.1712733 -66.2581363
H 6.8394987 13.3368053 -67.4929083
H 6.78462 11.6171695 -67.8603109
C 5.0988351 11.0728698 -65.6156212
H 5.8917595 11.1882859 -64.8758281
H 4.1709873 11.0890031 -65.0529447
C 5.2309691 9.6913749 -66.2729081
H 6.195366 9.5950969 -66.7695518
H 4.4343643 9.5515472 -67.0036455
H 5.1519131 8.9179254 -65.5089451
N 3.9636351 15.1292163 -67.8520424
H 3.2265652 14.4600239 -68.0547949
C 4.0218868 16.3871815 -68.6226211
H 5.0042765 16.4596209 -69.0901891
C 3.8797531 17.6361744 -67.751329
O 4.3853571 18.6900604 -68.1393617
C 2.9423549 16.3991231 -69.7126516
H 1.9845357 16.1193492 -69.2705717
H 2.8543825 17.4158555 -70.0989016
C 3.2701847 15.4763868 -70.8965743
H 3.2834725 14.434007 -70.5792135
H 4.2600221 15.7274747 -71.2810722
C 2.2462379 15.6651832 -72.0260354
H 2.5661878 15.0769262 -72.8877034
H 2.2537104 16.7118842 -72.3344896
N 0.873631 15.2917612 -71.613203
H 0.3736673 15.9475456 -71.0440993
C 0.2606047 14.1451865 -71.8617678
N 0.8324717 13.2109762 -72.562001
H 0.4215544 12.2750905 -72.7048818
H 1.7817286 13.3170273 -72.8573687
N -0.9394563 13.9204432 -71.4047869
H -1.4254517 14.5628763 -70.815964
H -1.3235025 12.9947997 -71.5411287
N 3.1468849 17.5726836 -66.640555
H 2.7147858 16.6893069 -66.3908726
C 3.0422072 18.6861717 -65.6891141
H 2.9536692 19.6115987 -66.2544648

C 4.3172776 18.8331359 -64.8542892
O 4.818539 19.9522103 -64.7316767
C 1.7891423 18.5422215 -64.8007602
H 1.0555546 17.9044931 -65.2924164
H 2.0612306 18.0606817 -63.8602092
C 1.1195113 19.8957002 -64.5028273
H 1.8473703 20.5638183 -64.0369374
H 0.3048766 19.7346467 -63.7929669
C 0.5491297 20.5310302 -65.7790337
O -0.2282403 19.8629453 -66.4986209
O 0.9457575 21.6510811 -66.1609826
N 4.94132 17.725144 -64.4234338
H 4.4998686 16.8199543 -64.5638902
C 6.2140599 17.8019982 -63.6909489
H 6.0524835 18.5815138 -62.9562327
C 7.3890477 18.3481159 -64.518059
O 8.2732855 18.9822581 -63.9445966
C 6.5604161 16.5360761 -62.8697259
H 7.6036421 16.612572 -62.5826921
C 5.687129 16.5745079 -61.5928064
H 4.6438896 16.3754152 -61.8451636
H 5.7360653 17.5498829 -61.1123979
H 6.0256541 15.83006 -60.8757306
C 6.3611885 15.1622226 -63.5084856
H 5.298497 14.9474787 -63.5301493
H 6.74844 15.1843201 -64.5161972
C 7.0610066 14.0140301 -62.7739185
H 6.6684855 13.8981371 -61.7667884
H 8.122155 14.2216184 -62.7084251
H 6.9110273 13.0840706 -63.3228865
N 7.30301 18.2991724 -65.852446
H 6.594574 17.6978391 -66.2541671
C 8.2430857 18.9945263 -66.7521705
H 9.256973 18.7089679 -66.4899572
C 8.1690656 20.51075 -66.5818082
O 9.1990222 21.1611148 -66.460308
C 7.977772 18.6399987 -68.229651
H 6.9892592 19.0220523 -68.484297
C 8.9845327 19.2914653 -69.1874602
H 10.0014391 18.9979181 -68.9194354
H 8.9126701 20.3778295 -69.1491886
H 8.7886603 18.966429 -70.2109033
C 7.96589 17.1408008 -68.5243393
H 8.943109 16.8146886 -68.8779901
H 7.2357508 16.9438971 -69.3046957
H 7.7171778 16.551408 -67.6508645

N 6.9615765 21.0893303 -66.532496
H 6.1488449 20.486938 -66.5022801
C 6.780615 22.5474602 -66.4135413
H 7.4330732 23.0541204 -67.1286945
C 7.2202667 23.0717325 -65.0569848
O 7.912679 24.0815246 -64.9821131
C 5.3159302 22.9327156 -66.6760503
H 4.6831159 22.049465 -66.5844177
H 4.9780527 23.653042 -65.9286919
C 5.1585947 23.5769347 -68.0589042
H 5.5905341 22.9257093 -68.8152424
H 5.7275276 24.5057287 -68.0837541
C 3.6905468 23.8966561 -68.3952165
H 3.6380314 24.2920585 -69.4108383
H 3.3606373 24.6917945 -67.7218206
N 2.7656957 22.7454938 -68.2527494
H 2.0283473 22.845749 -67.5579178
C 2.88091 21.5274568 -68.7454514
N 3.8693124 21.140862 -69.4991559
H 4.0949378 20.1565639 -69.4957941
H 4.5709032 21.8214323 -69.7161739
N 1.9903342 20.6334323 -68.4586807
H 1.2955849 20.8980357 -67.7519289
H 2.1477213 19.6774308 -68.6872359
N 6.8385406 22.3850171 -63.9793491
H 6.2678414 21.5594362 -64.1188139
C 7.0762139 22.8882716 -62.6154744
H 6.7536171 23.9302454 -62.5989197
C 8.5529194 22.901266 -62.209056
O 8.8936518 23.5075295 -61.1960446
C 6.2302789 22.1490035 -61.5659052
H 6.3002834 22.7158685 -60.6375336
C 4.7406745 22.1370212 -61.9312552
H 4.5608851 21.5072413 -62.7992222
H 4.4104188 23.1524728 -62.155835
H 4.1599278 21.7513496 -61.0951958
C 6.7187884 20.740197 -61.2639461
H 6.946144 20.2365441 -62.1944734
H 5.9697426 20.1820863 -60.7022476
H 7.6299397 20.7966649 -60.6732389
N 9.4330305 22.2598839 -62.9727844
H 9.1190088 21.8290885 -63.831774
C 10.8586508 22.2157185 -62.6987607
H 11.0205601 22.3896924 -61.6385775
C 11.6587618 23.2613323 -63.4943908
O 11.250341 23.686733 -64.571033

C 11.3162717 20.8126336 -63.009183
H 11.2222436 20.673128 -64.0870845
H 10.6948542 20.0962774 -62.4706286
H 12.34619 20.6925253 -62.698318
N 12.8048385 23.6814376 -62.9622398
H 13.0708624 23.3246349 -62.0518115
C 13.7495271 24.5578181 -63.6509041
H 13.1778 25.2862108 -64.2238724
C 14.6719354 23.7794981 -64.6223379
O 14.975169 22.6149417 -64.3663321
C 14.5534374 25.3133525 -62.584314
H 15.1349368 24.6050227 -61.9900518
H 13.8784704 25.8645982 -61.9278724
H 15.2402197 26.0126728 -63.0612602
N 15.2562672 24.4346856 -65.6428965
C 16.3457623 23.8565942 -66.4329207
H 15.9645131 23.0084207 -66.9972777
C 17.5084646 23.3942187 -65.5437926
O 17.9666931 24.1420647 -64.6815619
C 16.7604415 24.9539896 -67.41675
H 17.5396302 25.5804047 -66.9808329
H 17.0916402 24.5356973 -68.368371
C 15.4800201 25.7739039 -67.5748403
H 14.8102338 25.2746548 -68.2775496
H 15.6876092 26.7949285 -67.8961341
C 14.8755827 25.7311526 -66.1737705
H 13.7917232 25.8368128 -66.2352599
H 15.3069372 26.5190763 -65.555338
N 17.9761557 22.1537945 -65.7184936
H 17.5578649 21.5750566 -66.4370437
C 18.9691881 21.5330127 -64.828799
H 19.4528712 20.7173447 -65.3653614
H 19.7268063 22.271958 -64.5676816
C 18.3945524 20.9579177 -63.5231969
O 19.1431088 20.4234825 -62.6929978
N 17.0763422 20.9995032 -63.325819
H 16.4871733 21.5126475 -63.9742723
C 16.4200515 20.282351 -62.2438022
H 16.8992307 20.5608404 -61.3042546
C 16.5430015 18.7673145 -62.4295642
O 16.5979498 18.2344241 -63.5433115
C 14.9511917 20.6876016 -62.1594518
H 14.4759675 20.4721613 -63.1160417
H 14.876111 21.7519627 -61.935609
H 14.4412873 20.1255796 -61.3775577
N 16.4551063 18.0541021 -61.3125826

H 16.2855776 18.5690471 -60.4511779
C 16.3207142 16.6066863 -61.2672556
H 16.667364 16.1436285 -62.1925943
C 14.8430527 16.4876811 -61.1714432
O 14.2703098 16.4918358 -60.0760996
C 17.1025975 16.0055349 -60.0967373
H 16.7267382 16.4254048 -59.1674263
H 16.9332383 14.927969 -60.0696969
C 18.6151246 16.2789271 -60.2035818
H 18.7934009 17.3000794 -60.5397025
H 19.0498173 15.6006874 -60.9384452
C 19.3309473 16.1098317 -58.8681021
O 19.8209138 17.063498 -58.2767724
N 19.4280626 14.9016131 -58.3528361
H 19.9160454 14.8206799 -57.47876
H 19.0269415 14.1074667 -58.8221429
N 14.2900884 16.7928196 -62.3621341
H 14.9082487 16.9578584 -63.1449686
C 12.8660165 16.8921841 -62.5786515
H 12.50313 17.8129391 -62.118628
C 12.3945187 16.8195691 -64.0583579
H 11.5182164 16.1806944 -64.1205188
C 11.927267 18.2080647 -64.4477732
H 12.7140434 18.9289267 -64.2293846
H 11.0102521 18.4103486 -63.8975376
H 11.6871596 18.2879955 -65.5028421
C 13.3973882 16.3601632 -65.1198995
H 13.1242213 16.743578 -66.1014841
H 14.3632553 16.7589904 -64.8585091
C 13.4802412 14.8541873 -65.2635705
H 12.5137973 14.4524802 -65.5470749
H 13.7913259 14.4259101 -64.3196536
H 14.1809111 14.6061911 -66.0551717
C 12.2832975 15.764129 -61.7971147
O 11.2494559 16.0014477 -61.1925983
N 12.9953431 14.6160479 -61.6922139
H 13.8487719 14.4634771 -62.2140222
C 12.4409562 13.5648977 -60.9104669
H 11.9661348 14.0429408 -60.056893
C 13.3950247 12.5287825 -60.3251963
O 14.3898634 12.1321749 -60.9337501
C 11.3641755 12.9163869 -61.7760491
H 11.69877 11.9263678 -62.0871113
C 10.1448767 12.7675339 -60.8818847
H 9.7797381 13.7482455 -60.5753726
H 10.389334 12.200937 -59.985975

H 9.3515675 12.2659959 -61.4236928
C 10.8142608 13.647437 -63.0167973
H 10.2651247 14.5505195 -62.7809551
H 10.1246692 12.9691349 -63.5158578
H 11.6027534 13.8396563 -63.7244018
N 12.992268 12.0624941 -59.1407394
H 12.2290494 12.5696896 -58.7005972
C 13.3517678 10.7991302 -58.4875107
H 13.8691189 10.1454328 -59.1901295
C 12.022295 10.1409558 -58.0722694
O 11.2124517 10.7801555 -57.3905926
C 14.2732964 11.0619765 -57.2714117
H 13.7556098 11.7467959 -56.5983282
C 14.5539489 9.7552096 -56.5040559
H 15.098085 9.0547942 -57.1393494
H 13.6255968 9.2720965 -56.1977078
H 15.1326487 9.9473315 -55.6028041
C 15.5967541 11.7351388 -57.7104154
H 16.1322448 11.0671286 -58.3834904
H 15.3704105 12.6510699 -58.2557262
C 16.5299331 12.1281646 -56.5575202
H 16.9353557 11.2398739 -56.0737671
H 15.9862613 12.7290053 -55.8275692
H 17.3598219 12.7148318 -56.9523203
N 11.7580335 8.9046503 -58.5018071
H 12.4971308 8.4049395 -59.0006782
C 10.4983895 8.1867901 -58.2471085
H 9.9675434 8.6814518 -57.4396159
C 10.7986262 6.7492106 -57.8252037
O 11.102295 5.9443846 -58.7003625
C 9.5576584 8.1569015 -59.488233
H 10.0657489 7.6118162 -60.2850846
C 9.2280089 9.5527545 -60.0531933
H 8.7489912 10.1650888 -59.2892799
H 10.1684049 10.018405 -60.3410172
C 8.2538598 7.3944316 -59.1507915
H 7.707404 7.913199 -58.3643917
H 8.4738782 6.3822778 -58.8197286
H 7.6197814 7.2880813 -60.0276344
C 8.3419087 9.5094014 -61.311817
H 7.3388524 9.1658198 -61.0701747
H 8.7868381 8.8466241 -62.055071
H 8.2449684 10.4990386 -61.7479271
N 10.3778145 6.3647875 -56.6173089
H 10.0867171 7.0875804 -55.969359
C 10.0771669 4.965996 -56.2829504

H 10.6161371 4.3342597 -56.978693
C 8.5735844 4.707958 -56.4325046
O 7.7547345 5.5273144 -56.0055968
C 10.6004787 4.5910992 -54.8968489
H 10.1061673 5.1945444 -54.1366728
H 11.6735419 4.7889482 -54.8607758
O 10.3828632 3.2140938 -54.6418497
H 9.4472056 3.0366627 -54.4338089
N 8.1607001 3.6309068 -57.1055167
H 8.8769937 2.9804449 -57.4273788
C 6.7371884 3.300688 -57.3620578
H 6.1596131 3.5083897 -56.4621738
C 6.5491433 1.8064488 -57.6597888
O 7.4787363 1.1227106 -58.0834072
C 6.1822375 4.201798 -58.4902398
H 6.9381585 4.3099942 -59.2694911
C 4.8723056 3.766805 -59.1432156
H 4.1171477 3.6139869 -58.3752925
H 5.0235349 2.8460121 -59.7068042
H 4.5357146 4.5394803 -59.8340945
O 5.8902408 5.4700017 -57.9372644
H 6.6083844 5.6589023 -57.3062582
N 5.3715908 1.2506011 -57.3572631
H 4.6553821 1.8125038 -56.8983762
C 5.0533527 -0.1448405 -57.6414244
H 5.9175855 -0.7539901 -57.3692877
C 4.7686066 -0.3651727 -59.1348527
O 3.9382459 0.3149866 -59.7426248
C 3.8790949 -0.6020956 -56.7719286
H 2.9852171 -0.0491227 -57.0590648
H 4.0992285 -0.4156114 -55.7198245
H 3.7121333 -1.6687299 -56.9224242
N 5.384449 -1.4132231 -59.6710028
H 5.9615485 -1.978817 -59.0585404
C 5.274073 -1.8707675 -61.0481607
H 4.52063 -1.2819514 -61.5709594
C 4.819635 -3.3368143 -61.111532
O 4.71484 -4.0327033 -60.0985856
C 6.6290324 -1.6468094 -61.7413333
H 6.7247861 -2.2962304 -62.6122492
H 7.4480343 -1.8571985 -61.0513851
S 6.7161264 0.0744788 -62.2988248
H 5.7073454 -0.0156779 -63.1767116
N 4.5804398 -3.8163232 -62.3306187
H 4.6770828 -3.1691214 -63.1064769
C 4.346574 -5.2294842 -62.6551505

H 4.459447 -5.8264837 -61.7510928
C 5.4032849 -5.7429896 -63.6329606
O 6.1897972 -4.9572822 -64.1649364
C 2.9059161 -5.4309908 -63.1662228
H 2.7672556 -6.4849372 -63.4143421
H 2.2061259 -5.1920836 -62.3647088
C 2.5629917 -4.5759187 -64.3962885
H 3.3538419 -4.6507815 -65.1381943
H 2.4774119 -3.5317996 -64.1082129
C 1.239358 -5.0199314 -65.0200982
H 0.4409992 -4.8935195 -64.2851521
H 1.2911458 -6.0804497 -65.2787751
N 0.9258565 -4.2126199 -66.212061
H 0.4081314 -3.3590145 -66.0351975
C 1.2259418 -4.5210722 -67.4631425
N 1.9688723 -5.5468659 -67.7776155
H 2.1152903 -5.8721481 -68.7219343
H 2.3570204 -6.1389378 -67.0480665
N 0.7801215 -3.779182 -68.4391839
H 0.1596895 -3.0162654 -68.2328983
H 0.9754474 -4.0537471 -69.3841574
N 5.3899017 -7.0384504 -63.929528
H 4.7383203 -7.6344439 -63.4363782
C 6.0942102 -7.5596355 -65.1036889
H 7.0977374 -7.1300138 -65.1006469
C 5.4232967 -7.0925274 -66.4226341
O 4.3354509 -6.5106139 -66.4170726
C 6.2677109 -9.0838745 -64.9594063
H 6.6936781 -9.2945219 -63.9792089
H 6.9785677 -9.4402086 -65.7029615
C 4.9912565 -9.885612 -65.1365704
O 4.2025192 -9.6370408 -66.0222303
N 4.7808317 -10.9165378 -64.3629691
H 3.901636 -11.390675 -64.4666077
H 5.4341878 -11.1196392 -63.6117621
N 6.087026 -7.301176 -67.5654737
H 6.97492 -7.7768733 -67.5279906
C 5.5686416 -6.8960675 -68.8824775
H 5.3268371 -5.8354456 -68.847767
C 4.2717675 -7.636028 -69.2757134
O 3.3960169 -7.0361793 -69.9063768
C 6.6966829 -7.1057223 -69.9118019
H 7.5465829 -6.4850685 -69.622889
H 7.0144693 -8.1489414 -69.8602459
C 6.3428838 -6.7968035 -71.3816833
H 5.5274226 -7.4439008 -71.7033055

C 5.9359276 -5.3387014 -71.5982807
H 6.7312629 -4.6712647 -71.263949
H 5.0185384 -5.1252485 -71.0539862
H 5.7487945 -5.1681951 -72.6580208
C 7.5553389 -7.0841982 -72.2684394
H 8.3866897 -6.4341584 -71.9947723
H 7.2949337 -6.9136733 -73.313153
H 7.8578121 -8.1251123 -72.1530906
N 4.1334798 -8.8979656 -68.8636108
H 4.8250579 -9.2749777 -68.2335885
C 3.0282568 -9.7990911 -69.2066128
H 2.994302 -9.8630138 -70.2915907
C 1.6602333 -9.3119172 -68.6927965
O 1.5945623 -8.516802 -67.7502102
C 3.3072741 -11.2092038 -68.6610331
H 2.5364612 -11.8929918 -69.0166306
H 3.2739869 -11.2045757 -67.5732222
O 4.5717364 -11.6737482 -69.1032695
H 4.7435367 -12.5260357 -68.6915725
N 0.5433211 -9.7783737 -69.2905212
C -0.7985269 -9.4881579 -68.7920022
H -0.974344 -8.4156383 -68.878059
C -0.9754439 -9.9212677 -67.3331253
O -0.527133 -10.9988821 -66.9390965
C -1.7659872 -10.2366409 -69.7171588
H -1.976375 -11.2318292 -69.3196156
H -2.693753 -9.6827823 -69.8650191
C 0.4554123 -10.5666366 -70.5124142
H 1.1583618 -10.2258453 -71.2721388
H 0.6242898 -11.6193873 -70.280253
C -0.973233 -10.3730318 -71.0152853
H -1.0393598 -9.4455678 -71.586019
H -1.3115491 -11.2200922 -71.6130074
N -1.6717958 -9.0991233 -66.547554
H -1.9774122 -8.2113845 -66.9104634
C -1.9426282 -9.3712957 -65.1363939
H -0.9869589 -9.4906116 -64.6280691
C -2.7599268 -10.6578647 -64.9656591
O -3.8400503 -10.7898855 -65.5414905
C -2.701309 -8.1901542 -64.5041285
H -2.7718945 -8.3380187 -63.4261289
H -3.7101012 -8.1606276 -64.9151862
O -2.0772418 -6.945159 -64.7763668
H -2.4140318 -6.2669389 -64.1668781
N -2.292544 -11.5662814 -64.1043462
H -1.3627723 -11.44079 -63.7365719

C -3.0207712 -12.7921241 -63.7457168
H -3.2244875 -13.323451 -64.6781217
C -4.3966937 -12.4684447 -63.1343671
O -4.6230048 -11.3825377 -62.579092
C -2.1096583 -13.72106 -62.9038127
H -1.910148 -14.6234283 -63.4834728
H -1.1418491 -13.2284864 -62.8004951
C -2.5148567 -14.1252411 -61.4840284
H -2.7946617 -13.2267165 -60.9280213
H -1.6416138 -14.553518 -60.9864657
C -3.6505368 -15.1492933 -61.4302847
O -4.2452488 -15.22482 -60.332582
O -4.0275463 -15.7087837 -62.4791995
N -5.3275999 -13.4242844 -63.2208885
H -5.0081732 -14.3620026 -63.4746751
C -6.7373853 -13.2558538 -62.8708644
H -7.1275333 -12.4809128 -63.5271843
C -6.946757 -12.7679238 -61.4410492
O -7.9117529 -12.0322895 -61.2615722
C -7.4812086 -14.5788163 -63.1367407
H -7.2969487 -14.8900304 -64.1668939
H -7.0568127 -15.3459934 -62.4846586
C -8.9995942 -14.5392484 -62.9048429
H -9.3885239 -15.5543231 -62.9959171
H -9.1947796 -14.2219061 -61.8805407
C -9.7553918 -13.6445641 -63.9063237
H -10.1326223 -14.2766743 -64.7127237
H -9.0952164 -12.9031723 -64.3565096
C -10.9343519 -12.935928 -63.2296966
H -11.4774207 -13.6713244 -62.6279936
H -11.6147183 -12.563259 -64.0023007
N -10.4659423 -11.8111498 -62.3804404
H -9.7675482 -12.1340372 -61.710333
H -11.2343898 -11.3862407 -61.8810379
H -10.0067443 -11.1094506 -62.9439105
N -6.0816248 -13.0886981 -60.4718269
H -5.3969747 -13.8110332 -60.6825751
C -6.0968226 -12.5397581 -59.1017552
H -7.1172031 -12.2357542 -58.8791371
C -5.2212816 -11.2922417 -58.8815129
O -5.4720267 -10.5585664 -57.9262851
C -5.7549487 -13.6167123 -58.0640669
H -6.3952429 -14.4869167 -58.2181221
H -5.9482444 -13.2182487 -57.0671658
O -4.3992676 -14.0098058 -58.1432093
H -4.343102 -14.5866227 -58.960205

N -4.2784466 -10.9636136 -59.7796263
H -4.191226 -11.5356199 -60.6116894
C -3.5645761 -9.675878 -59.7459851
H -3.2982635 -9.4612025 -58.710529
C -4.4764878 -8.5391038 -60.2170946
O -4.6424323 -7.5619692 -59.492832
C -2.2637066 -9.7208699 -60.5782299
H -1.5860132 -10.4576694 -60.1459091
H -2.5060962 -10.0411425 -61.5913691
C -1.54592 -8.348949 -60.6594275
H -2.2252017 -7.6004366 -61.0640231
C -1.049715 -7.869789 -59.2926931
H -0.3925492 -8.6162577 -58.8481829
H -1.895953 -7.6778135 -58.6345231
H -0.5004847 -6.9352805 -59.4146506
C -0.3461376 -8.4080241 -61.5969519
H 0.3865714 -9.1103831 -61.226032
H 0.1108929 -7.4212003 -61.6794127
H -0.6523785 -8.7230191 -62.5882788
N -5.1169134 -8.689943 -61.3849318
H -4.9457027 -9.5459942 -61.90868
C -6.0412299 -7.6860189 -61.9416269
H -5.4551489 -6.7857532 -62.1437637
C -7.1240799 -7.231914 -60.9375162
O -7.2297424 -6.0346225 -60.7273625
C -6.5934143 -8.1928127 -63.2907024
H -5.7661369 -8.2380254 -63.9984981
H -6.9541082 -9.2147834 -63.16253
C -7.7329938 -7.3712305 -63.9310273
H -7.8971692 -7.7500898 -64.9408023
H -8.6442331 -7.5659483 -63.3636415
C -7.5575872 -5.8442729 -64.0094301
H -7.4980538 -5.4352241 -63.0077227
H -8.4615509 -5.4163366 -64.4482493
C -6.3614685 -5.3390559 -64.8205827
H -6.662531 -5.2804452 -65.8695286
H -5.5195257 -6.0309988 -64.7300999
N -5.9571078 -4.0034871 -64.3263948
H -6.7679802 -3.395136 -64.17414
H -5.2630821 -3.5587778 -64.9038722
H -5.4837897 -4.0957187 -63.4295262
N -7.871645 -8.0931999 -60.2239082
C -8.873936 -7.6616363 -59.2573345
H -9.5234761 -6.9139405 -59.7159941
C -8.2649088 -7.0320422 -57.9992117
O -8.9337415 -6.198365 -57.3998078

C -9.7099499 -8.8997193 -58.9176722
H -10.0869565 -8.8805224 -57.8938592
H -10.5345133 -8.9914458 -59.6264198
C -7.9351771 -9.5147877 -60.3439147
H -8.4774898 -9.7379089 -61.262144
H -6.9490279 -9.9651637 -60.3563997
C -8.720291 -10.0246495 -59.1344312
H -9.2281338 -10.9685741 -59.3297095
H -8.0659944 -10.10908 -58.2666461
N -7.021786 -7.3606245 -57.6043131
H -6.4998668 -8.0322734 -58.1535313
C -6.3056253 -6.6353359 -56.5348897
H -6.9728801 -6.5111878 -55.6824392
C -5.9322881 -5.2270172 -57.0036012
O -6.1602659 -4.2700824 -56.2681912
C -5.0687729 -7.4390979 -56.0860549
H -5.3895788 -8.424192 -55.7418211
H -4.4305065 -7.5948128 -56.9556649
C -4.1992388 -6.7757217 -54.993123
H -3.2115868 -7.2341931 -55.0481208
H -4.0668317 -5.7140503 -55.2046993
C -4.7169933 -6.9550987 -53.5545555
H -4.8133604 -8.0239603 -53.351868
H -5.6999526 -6.4874112 -53.4551922
C -3.7314371 -6.3211879 -52.5553124
H -3.9467867 -5.2535484 -52.4822222
H -2.7197203 -6.4281609 -52.9491867
N -3.7880118 -6.9492504 -51.2117734
H -4.7008678 -6.8646728 -50.7950431
H -3.1294465 -6.5095671 -50.5629086
H -3.5314121 -7.9249485 -51.2652042
N -5.4101965 -5.0928218 -58.2204721
H -5.2513331 -5.9347831 -58.7634563
C -5.1139113 -3.8079675 -58.8626221
H -4.4172266 -3.2441876 -58.2414818
C -6.3827766 -2.9486339 -58.9977917
O -6.424878 -1.8461333 -58.4601641
C -4.4306352 -4.0925634 -60.2147685
H -3.4889227 -4.612916 -60.0251308
H -5.0681263 -4.750292 -60.8032861
C -4.1402352 -2.8329003 -61.0402102
H -3.5223312 -2.1630568 -60.4391233
H -5.0881717 -2.3339607 -61.2560169
C -3.4537445 -3.1310734 -62.3813936
O -3.6879035 -4.2298107 -62.9487496
O -2.7668995 -2.224337 -62.9038172

N -7.4664462 -3.5002571 -59.5499567
H -7.3734285 -4.4240226 -59.9575726
C -8.7116426 -2.7639545 -59.7964255
H -8.4353106 -1.8095062 -60.2480485
C -9.4638687 -2.4089424 -58.5074308
O -9.9528278 -1.288221 -58.3755535
C -9.6118555 -3.5466591 -60.7821039
H -9.0607949 -4.3693492 -61.2330581
H -10.4668038 -3.9705644 -60.2532929
C -10.1111674 -2.6392604 -61.9230753
H -10.7214885 -1.8416841 -61.4932013
H -10.7558622 -3.2309231 -62.5768496
C -8.9667387 -2.0313695 -62.7636765
O -8.2371866 -2.7994259 -63.4391515
O -8.8066446 -0.7847794 -62.7441526
N -9.47042 -3.2962327 -57.4999447
H -9.0417097 -4.20374 -57.6524768
C -10.1010431 -3.0170332 -56.1973756
H -11.0288266 -2.4764864 -56.3973745
C -9.2769901 -2.0761537 -55.3142795
O -9.8498703 -1.4157787 -54.4554675
C -10.4660358 -4.334596 -55.4944689
H -11.0058283 -4.9560219 -56.2112972
H -9.5566922 -4.8544857 -55.1890845
C -11.3704398 -4.1053174 -54.2696322
H -10.7558442 -3.807233 -53.4188719
H -12.0692675 -3.2953097 -54.4855386
C -12.2007088 -5.3376816 -53.8874421
H -12.8839548 -5.5640901 -54.7082912
H -12.7938671 -5.0865768 -53.0059178
C -11.3301939 -6.5636396 -53.5898349
H -10.6293076 -6.3072624 -52.7892185
H -10.7480368 -6.8020651 -54.4859351
N -12.1634946 -7.726805 -53.1936491
H -12.701381 -7.5095402 -52.3633894
H -11.5843428 -8.534113 -52.9993255
H -12.8073944 -7.9650457 -53.9371857
N -7.9716141 -1.9444161 -55.5661454
H -7.5758423 -2.5040646 -56.3110106
C -7.1488087 -0.8533673 -55.0040867
H -7.6222074 -0.5067725 -54.0856447
C -7.1064176 0.3871534 -55.9041949
O -6.4532318 1.363562 -55.5514579
C -5.7327836 -1.3132419 -54.6059369
H -5.1704231 -0.4493004 -54.2512706
C -5.7686133 -2.3392788 -53.4714731

H -6.3200593 -3.2274988 -53.7777969
H -6.2540956 -1.8980821 -52.6008782
H -4.7497511 -2.6174525 -53.2046511
O -5.0121386 -1.9015539 -55.6648057
H -5.5081724 -2.690785 -55.9458878
N -7.8166632 0.3654646 -57.0417343
H -8.3462446 -0.4726061 -57.2427193
C -7.8502309 1.3899778 -58.0923619
H -8.3461036 0.901136 -58.9292334
C -6.4594769 1.7892235 -58.633805
O -6.3001095 2.8344891 -59.2640218
C -8.7526115 2.5574542 -57.6322694
H -9.6187161 2.1305136 -57.1235023
H -8.1999693 3.1478449 -56.8997247
C -9.2803135 3.5032468 -58.7320415
H -8.4579137 4.0436594 -59.1952806
C -10.0659324 2.771923 -59.8264709
H -10.8596408 2.1741863 -59.3761475
H -9.4021933 2.1159166 -60.3858728
H -10.5024011 3.4948423 -60.5147045
C -10.2132689 4.5376623 -58.0986059
H -11.0746174 4.0426197 -57.649767
H -10.5543428 5.2422816 -58.85601
H -9.6730952 5.0878871 -57.3275553
N -5.4426723 0.9434196 -58.4481881
H -5.6678236 0.0355238 -58.0548346
C -4.0798935 1.185912 -58.9357536
H -3.8150141 2.2211407 -58.7113507
C -4.0051982 1.032849 -60.4518383
O -4.7678273 0.27695 -61.0434649
C -3.0949649 0.2568287 -58.2130025
H -3.5138127 -0.7490093 -58.1422002
H -2.1678002 0.2159683 -58.784929
C -2.795052 0.8141 -56.8097441
H -2.3119699 1.7866437 -56.9208895
H -3.7301621 0.9656837 -56.2690902
C -1.8893985 -0.0773955 -55.9502131
H -2.4545609 -0.9480272 -55.6154601
H -1.5950424 0.5033683 -55.0736083
C -0.6439782 -0.5463608 -56.7108241
H -0.3018319 0.268945 -57.3493463
H -0.9215663 -1.3830817 -57.3566641
N 0.4467047 -0.9339892 -55.7916971
H 0.7465728 -0.112789 -55.2498367
H 1.2748543 -1.2105249 -56.2979961
H 0.1611309 -1.6486113 -55.1440047

N -3.0487572 1.7262357 -61.0774219
H -2.4915023 2.3509615 -60.5015827
C -2.7384924 1.6213578 -62.5167704
H -3.2764944 0.767828 -62.9341887
C -1.2710996 1.2723612 -62.7035448
O -0.4066791 2.1504201 -62.6343866
C -3.1835379 2.9095297 -63.2545982
H -3.4195988 3.6894009 -62.5300726
H -2.3764105 3.2816492 -63.8842477
C -4.4157079 2.6815394 -64.1627795
H -4.7723658 3.6569256 -64.4963477
H -4.1117891 2.1108096 -65.0407656
C -5.5713926 1.9356428 -63.4710507
H -5.670715 2.3288555 -62.460506
H -5.3424209 0.8725846 -63.4125411
C -6.9189723 2.0674559 -64.1837241
H -6.9312356 1.427326 -65.0694601
H -7.0421595 3.108668 -64.4916204
N -8.0097774 1.7033435 -63.2510456
H -7.9605382 0.719054 -62.960874
H -8.919042 1.7965361 -63.6756161
H -7.9574248 2.2846703 -62.4272009
N -0.9851324 -0.0152724 -62.8685472
H -1.7507671 -0.6921228 -62.9343551
C 0.3622951 -0.5630426 -62.7563147
H 0.9805693 0.166144 -62.2307181
C 0.9623679 -0.7635467 -64.1569443
O 0.4536773 -1.5034089 -65.0035394
C 0.3662204 -1.8384641 -61.8795023
H -0.0630003 -2.6610733 -62.4548948
C 1.819163 -2.1735927 -61.5191769
H 2.2062048 -1.4454116 -60.8056847
H 2.4410637 -2.152216 -62.4072739
H 1.8767011 -3.1699076 -61.08193
C -0.4305112 -1.6738873 -60.5577608
H 0.0351023 -0.9039866 -59.9410057
H -1.442414 -1.3440835 -60.7786126
C -0.5557201 -2.9669112 -59.7395202
H 0.4082644 -3.2467547 -59.3150915
H -0.9233073 -3.7726035 -60.3764866
H -1.265722 -2.8133331 -58.927048
N 2.0820627 -0.0851478 -64.4029156
H 2.4655832 0.475942 -63.6575331
C 2.9068083 -0.2770199 -65.5962123
H 2.3018658 -0.6952979 -66.4020926
C 4.0396154 -1.2643317 -65.292497

O 4.4307098 -1.4269569 -64.1332412
C 3.4495725 1.0814523 -66.061202
H 4.0958151 0.9525892 -66.9322466
H 4.0432311 1.5292122 -65.261646
S 2.0709612 2.1846743 -66.4876026
H 1.3003227 1.8921115 -65.4342484
N 4.6448883 -1.8495044 -66.3268338
H 4.3466771 -1.6176306 -67.261478
C 5.9932657 -2.3856146 -66.1595621
H 6.0491473 -2.8706147 -65.1886549
C 7.0088041 -1.2347704 -66.1249314
O 6.737833 -0.132612 -66.6086652
C 6.3073177 -3.4524623 -67.2163512
H 5.4957053 -4.1766962 -67.2517385
H 7.2191217 -3.9815768 -66.9375472
C 6.5230252 -2.8364934 -68.5773457
O 7.5678347 -2.280934 -68.8635157
N 5.5086591 -2.8222407 -69.403895
H 5.6807276 -2.3748843 -70.3032807
H 4.6772054 -3.3460722 -69.2191437
N 8.1801732 -1.5131329 -65.5618081
H 8.3470506 -2.4508453 -65.2320084
C 9.2452561 -0.5331003 -65.3748175
H 8.8589516 0.286887 -64.7647094
C 9.6957767 0.098359 -66.7072817
O 9.7789872 1.3222783 -66.8014925
C 10.3873618 -1.2029495 -64.5780293
H 11.2574185 -0.549599 -64.5671677
H 10.0590078 -1.2929866 -63.5414857
C 10.8177317 -2.5865022 -65.0490904
C 11.9181375 -2.7365552 -65.9169673
H 12.4819757 -1.8755661 -66.2390297
C 12.3053206 -4.0136478 -66.363435
H 13.1452796 -4.1187607 -67.0335377
C 11.597087 -5.1547648 -65.9319127
O 11.9592972 -6.3881262 -66.3699191
H 12.7274721 -6.3595775 -66.9421564
C 10.5129368 -5.0118722 -65.040923
H 9.9908157 -5.8930459 -64.6965091
C 10.1282528 -3.7322316 -64.5973171
H 9.308215 -3.6379075 -63.8983316
N 9.914116 -0.6953223 -67.7660937
H 9.7185 -1.6813804 -67.6780474
C 10.4258322 -0.1969978 -69.0526211
H 11.3531195 0.3334907 -68.8684677
C 9.4744436 0.8147866 -69.7068052

O 9.9050969 1.903903 -70.0918449
C 10.7555701 -1.3866897 -69.9704476
H 11.4799712 -2.0264374 -69.4643361
H 9.8468928 -1.968167 -70.1290859
C 11.319076 -0.9912917 -71.3499645
H 10.5886703 -0.3825456 -71.8825983
C 12.633967 -0.2142643 -71.2488822
H 13.3625122 -0.779567 -70.6682336
H 12.462991 0.7532524 -70.7772734
H 13.0319279 -0.0350601 -72.2476385
C 11.5735498 -2.2547509 -72.1719436
H 12.3160219 -2.8816896 -71.6783682
H 11.9344282 -1.9816748 -73.1636924
H 10.6437644 -2.8131246 -72.2827093
N 8.1721243 0.5219023 -69.7300654
H 7.8781855 -0.3931025 -69.4008782
C 7.1627271 1.4694778 -70.216607
H 7.465206 1.8258281 -71.2019721
C 7.0485684 2.7218684 -69.344085
O 6.7977963 3.7974389 -69.8845881
C 5.8037654 0.7721156 -70.3591403
H 5.0142317 1.5249066 -70.3314356
H 5.6424419 0.0910384 -69.5229777
C 5.6722935 0.0300481 -71.6647418
N 5.9532087 -1.3209948 -71.8725283
C 5.7499517 -1.545272 -73.178947
H 5.897551 -2.4993019 -73.6695688
N 5.3500598 -0.4176791 -73.790252
H 5.1471925 -0.3273254 -74.7758818
C 5.2980715 0.5887433 -72.851667
H 5.0341347 1.6258243 -73.0140616
N 7.3164396 2.6332608 -68.0360767
H 7.5599259 1.7314697 -67.6420576
C 7.2614763 3.7968527 -67.1509479
H 6.3620564 4.3393293 -67.4291629
C 8.4261241 4.7738874 -67.3880553
O 8.1695259 5.9696373 -67.5199989
C 7.0948289 3.3363918 -65.6891448
H 6.3288467 2.5601209 -65.6683109
H 8.0271337 2.8928056 -65.3390609
C 6.6681347 4.4549309 -64.7117129
H 7.5189438 5.109165 -64.5258907
C 5.4997377 5.3156355 -65.2074584
H 4.6757884 4.679795 -65.5306189
H 5.8301918 5.9439793 -66.0348046
H 5.1677863 5.9791257 -64.4099764

C 6.2145039 3.8387536 -63.3851094
H 5.3016056 3.2618588 -63.5260214
H 6.0445249 4.6237662 -62.6487135
H 6.9882306 3.1722436 -63.0041928
N 9.6480168 4.2850584 -67.6533181
H 9.7997559 3.2846604 -67.5630426
C 10.7262591 5.1507473 -68.1801541
H 10.8557976 5.9933419 -67.499146
C 10.3952471 5.6900104 -69.5631661
O 10.7418516 6.824877 -69.8713061
C 12.0605472 4.4240518 -68.3016877
H 12.1166467 3.8610945 -69.2350101
H 12.121121 3.7293563 -67.4803064
O 13.1491082 5.3247477 -68.1948499
H 13.269569 5.8510611 -69.0056855
N 9.6793356 4.9112594 -70.3797769
H 9.4739327 3.9631262 -70.084374
C 9.1529553 5.3651188 -71.6612905
H 9.9800387 5.6169965 -72.3253587
H 8.5721268 4.5597261 -72.1096179
C 8.2436502 6.5889815 -71.5206075
O 8.4964877 7.5881455 -72.1829673
N 7.271716 6.5802667 -70.6017955
H 7.1008963 5.7179054 -70.0934582
C 6.3753098 7.7248641 -70.3564864
H 5.913338 8.0175862 -71.2986071
C 7.1064366 8.967594 -69.8279953
O 6.7901523 10.0960769 -70.2162708
C 5.2706065 7.3106618 -69.3736671
H 5.7289851 6.9026424 -68.4718638
H 4.7203521 8.2063148 -69.0832355
C 4.2527546 6.3259864 -69.9274463
C 3.9569052 5.1377987 -69.2312677
H 4.470828 4.90886 -68.3126257
C 2.9868958 4.2463839 -69.7247382
H 2.7656118 3.3367646 -69.1875547
C 2.2932897 4.5473782 -70.9100298
H 1.5351606 3.8746796 -71.2849157
C 2.5749518 5.7375725 -71.6014384
H 2.0297721 5.995317 -72.5016579
C 3.5527959 6.6201892 -71.1146242
H 3.7383022 7.5375435 -71.6559592
N 8.1333138 8.7500029 -69.0098282
H 8.2811018 7.8069615 -68.6678179
C 9.07562 9.7716512 -68.5730746
H 8.5547353 10.5922679 -68.0912188

C 9.8605926 10.3853852 -69.7513712
O 9.7566054 11.5847949 -70.0065046
C 9.9597189 9.1076336 -67.5219712
H 10.8339721 9.7058895 -67.3759088
H 10.2895408 8.1279684 -67.8552867
S 9.0625811 8.9471298 -65.9503501
H 9.8351265 9.8474364 -65.3019252
N 10.4805271 9.5504548 -70.5898632
H 10.5030738 8.5688311 -70.3340113
C 11.2121939 9.9579073 -71.8004392
H 12.0155281 10.6371849 -71.5108146
C 10.3189662 10.6842638 -72.8220458
O 10.6706195 11.7496569 -73.3218699
C 11.8287769 8.7012708 -72.4308687
H 11.0787146 8.1829872 -73.0301747
H 12.1581597 8.0264929 -71.641054
O 12.9535704 9.016732 -73.2287619
H 13.2151123 8.2260534 -73.7115009
N 9.0917743 10.1931924 -73.0341901
H 8.8462209 9.3288767 -72.56349
C 8.0758771 10.7997598 -73.908273
H 8.5230395 10.9933282 -74.8827677
C 7.5601152 12.1572996 -73.4059811
O 6.9709603 12.9057875 -74.1813275
C 6.8980951 9.8177845 -74.0665242
H 6.5963806 9.4797408 -73.0746319
H 6.0469815 10.3468555 -74.4990079
C 7.2010356 8.596359 -74.9559661
H 8.1359891 8.1311667 -74.6495739
C 6.078985 7.5667695 -74.8172612
H 5.1265064 8.0041828 -75.1217947
H 6.002105 7.244648 -73.7790986
H 6.2918725 6.6974123 -75.4373065
C 7.3195826 8.9722824 -76.4354788
H 6.4050387 9.4644876 -76.7684964
H 7.4802548 8.0744735 -77.031481
H 8.1643764 9.6422728 -76.586479
N 7.8083199 12.5056056 -72.1428918
H 8.3267403 11.8644839 -71.5540116
C 7.4700463 13.8201725 -71.5844624
H 6.6249218 14.2248713 -72.140381
C 8.5965015 14.8537852 -71.7283394
O 8.4412966 15.9715581 -71.2569983
C 7.0096966 13.6702722 -70.1392581
H 6.7278152 14.646009 -69.7435907
H 7.8241777 13.2648047 -69.5378654

O 5.8818799 12.8084382 -70.0899695
H 6.1934362 11.8939971 -70.2199382
N 9.6527789 14.5194905 -72.4726668
H 9.6609405 13.583641 -72.8564542
C 10.8450282 15.3307806 -72.7823003
H 11.3215018 14.7681748 -73.5857787
C 11.9223128 15.3449434 -71.6844862
O 12.7208264 16.2703293 -71.5460031
C 10.4885248 16.6983789 -73.4123986
H 10.3052822 17.4216129 -72.6154718
H 9.5630553 16.5886969 -73.9810148
C 11.5483248 17.2610639 -74.3780375
O 12.3958398 16.4764423 -74.8659835
O 11.4371665 18.4657644 -74.7049311
N 11.9834183 14.26542 -70.8998863
H 11.310854 13.5258562 -71.0438426
C 13.0028759 14.0912496 -69.8714545
H 13.306885 15.0728525 -69.5156226
C 14.2829326 13.4339092 -70.3807713
O 14.2769208 12.3465484 -70.963321
C 12.4106606 13.3349922 -68.6923727
H 12.0504773 12.3665639 -69.0383334
H 13.215576 13.1470367 -67.9823596
C 11.3089208 14.0511588 -67.9372688
C 10.4143586 13.2895275 -67.1652494
H 10.4207411 12.2129036 -67.2452246
C 9.622269 13.9020251 -66.183757
H 9.0420531 13.2912381 -65.5123696
C 9.6896914 15.2939169 -66.0117228
O 9.0688287 15.8810816 -64.9659394
H 8.6517885 15.2197967 -64.4135997
C 10.476267 16.0688853 -66.8777947
H 10.5008894 17.1334103 -66.7732363
C 11.2869235 15.4568867 -67.8389113
H 11.9309688 16.084664 -68.4366982
N 15.413457 14.0666489 -70.0648519
H 15.3470006 14.8954175 -69.4865486
C 16.7466287 13.6072225 -70.4479379
H 16.6439033 13.0148765 -71.3578561
C 17.3845581 12.7089696 -69.3861672
O 17.0498046 12.785335 -68.2019845
C 17.6340751 14.8021464 -70.8021027
H 17.0888067 15.4656922 -71.4759625
H 18.5250258 14.4458703 -71.3186039
O 18.0387931 15.5263655 -69.6592734
H 17.3011331 15.5682376 -69.0191956

N 18.3085085 11.8526167 -69.839663
H 18.540374 11.8840546 -70.8182419
C 19.0346307 10.8940314 -69.0057114
H 19.4675616 10.1472293 -69.6713046
C 18.1121144 10.1118048 -68.0476025
O 18.3664568 10.0593789 -66.8463913
C 20.2060694 11.6112569 -68.3167241
H 19.8233584 12.3411857 -67.6012615
H 20.822286 12.1241624 -69.055329
H 20.8190484 10.8836954 -67.7848943
N 17.0082147 9.5547067 -68.5680509
H 16.8205954 9.6340163 -69.5538031
C 16.1258953 8.7216485 -67.7571872
H 16.0455573 9.2076642 -66.7871337
C 16.7726785 7.3640399 -67.4612841
O 17.037384 6.5613184 -68.3551552
C 14.6792575 8.6592675 -68.293561
H 14.2122908 9.6269136 -68.1101076
H 14.1203773 7.931987 -67.704949
C 14.479784 8.3334479 -69.7606206
O 14.1690558 7.2130685 -70.1375173
N 14.5042034 9.3331716 -70.6191546
H 14.1908225 9.1433714 -71.5603483
H 14.5384129 10.2922772 -70.2963576
N 17.0392665 7.1424113 -66.177116
H 16.830673 7.8928146 -65.5251381
C 17.5157217 5.8978433 -65.5954028
H 17.9177789 5.2616605 -66.384763
C 16.3357946 5.207524 -64.9288189
O 15.6159122 5.8225069 -64.1466295
C 18.6212937 6.1737034 -64.5672921
H 18.1711266 6.6156535 -63.6769974
H 19.0702786 5.2249355 -64.2716113
C 19.7093446 7.0913046 -65.0380035
C 19.7931824 8.4044157 -64.7311366
H 19.0720161 8.9404149 -64.124824
N 20.8504143 8.9783652 -65.4059943
H 20.9953366 9.9773354 -65.4322483
C 21.4912532 8.0630289 -66.2111008
C 22.5607112 8.1678796 -67.1126483
H 23.0547657 9.1163029 -67.2628505
C 22.976687 7.0234162 -67.8144749
H 23.7994497 7.0845023 -68.5131482
C 22.3164877 5.7977862 -67.6094385
H 22.6337933 4.9207785 -68.1562422
C 21.2368826 5.7069224 -66.707767

H 20.7248034 4.766601 -66.5741431
C 20.7952953 6.8365602 -65.9837824
N 16.1848563 3.912817 -65.176291
H 16.821799 3.469261 -65.8205527
C 15.0796393 3.1104436 -64.6775838
H 14.54308 3.6333506 -63.8833365
C 15.6247004 1.7985474 -64.1325112
O 16.0577958 0.9402323 -64.9020713
C 14.131694 2.8887033 -65.8487103
H 13.64995 3.8395378 -66.0623428
H 14.7289034 2.6128478 -66.7202805
C 13.0901348 1.8002141 -65.5654381
H 13.4487714 1.0489044 -64.8679001
C 11.8698551 2.4330043 -64.9143906
H 11.3898081 3.1481945 -65.575671
H 12.1699034 2.9421886 -63.9979327
H 11.1508884 1.6662489 -64.6483251
C 12.8433832 1.0456084 -66.8611406
H 12.3756177 1.6885177 -67.5980209
H 12.2143535 0.1981272 -66.6495824
H 13.7858171 0.6635186 -67.2549475
N 15.5022455 1.6123315 -62.8227831
H 15.0176951 2.3212374 -62.2749974
C 15.9819101 0.4156113 -62.1354336
H 16.5767406 -0.1688088 -62.8309248
C 14.7948239 -0.412656 -61.638592
O 14.0877462 0.0314062 -60.7298387
C 16.9228566 0.7713672 -60.9737509
H 16.3487296 1.1324943 -60.119959
C 17.7625409 -0.435787 -60.5611816
H 18.3692173 -0.7740933 -61.4018301
H 17.1180753 -1.2500821 -60.2345302
H 18.4224701 -0.1512396 -59.7425426
O 17.83844 1.7718527 -61.3585353
H 17.3314862 2.5714806 -61.5236629
N 14.5757734 -1.6381667 -62.1522123
C 13.7832369 -2.6243789 -61.4326093
H 12.8078798 -2.2156713 -61.1651937
C 14.5459745 -3.0172483 -60.1610187
O 15.6270337 -3.5983735 -60.2304277
C 13.5987399 -3.7952014 -62.405327
H 13.5382697 -4.7567764 -61.8952256
H 12.6985773 -3.6284725 -62.9974839
C 15.1494966 -2.2202696 -63.3610773
H 14.6833669 -1.7644417 -64.236083
H 16.2312348 -2.0961457 -63.4033807

C 14.823558 -3.7137458 -63.3158808
H 14.6132543 -4.1093829 -64.3096971
H 15.6551597 -4.2543635 -62.8618106
N 13.9662979 -2.746838 -58.9908767
H 13.0661721 -2.2754632 -59.0090904
C 14.5182136 -3.1102436 -57.6789229
H 15.4998097 -3.5591397 -57.8229977
C 13.7014467 -4.2044761 -56.9428997
O 13.6238671 -4.1604091 -55.7146881
C 14.714941 -1.81596 -56.8514598
H 13.7756341 -1.2643591 -56.8431172
H 14.9534945 -2.0737327 -55.819355
C 15.8229594 -0.873091 -57.3541501
H 15.6404761 -0.6146515 -58.3959849
C 15.8120489 0.4127376 -56.5303236
H 16.028931 0.1980959 -55.4826737
H 14.8309305 0.8833728 -56.593059
H 16.5554212 1.1109018 -56.9145674
C 17.2136708 -1.5018124 -57.2217831
H 17.3755269 -1.8389114 -56.1975999
H 17.9750578 -0.7631874 -57.470298
H 17.3151205 -2.3433035 -57.9050428
N 13.0948036 -5.214763 -57.6151661
C 12.2053006 -6.1721577 -56.9514439
H 11.3888186 -5.644905 -56.4617025
C 12.9249789 -7.0386965 -55.9075064
O 12.3195788 -7.4085346 -54.9111168
C 11.6162382 -7.0363963 -58.0682371
H 11.3764525 -8.0460584 -57.7312578
H 10.7270492 -6.5549761 -58.4702904
C 13.2765494 -5.6211828 -59.0043124
H 12.7053484 -4.9577645 -59.6524598
H 14.33053 -5.6213029 -59.2835882
C 12.7120005 -7.0340446 -59.1282169
H 12.3119336 -7.2235629 -60.1252091
H 13.4796364 -7.7662537 -58.8743209
N 14.2193922 -7.3309605 -56.0875591
H 14.6758111 -6.9951267 -56.9187628
C 15.0103271 -8.0990672 -55.112049
H 14.434692 -8.9910839 -54.8655987
C 15.2070223 -7.332927 -53.7964256
O 15.0766997 -7.9116657 -52.717904
C 16.3497863 -8.552893 -55.7413196
H 16.1067111 -9.1238207 -56.6393262
C 17.2457079 -7.3784555 -56.1790683
H 17.6104389 -6.833258 -55.3064419

H 16.7096291 -6.695959 -56.8363059
H 18.1109972 -7.7622999 -56.7213065
C 17.1758962 -9.4774166 -54.8192359
H 18.0864798 -9.7670527 -55.3455807
H 17.4716894 -8.935527 -53.9193684
C 16.4479887 -10.7630839 -54.4053729
H 16.0929498 -11.2919093 -55.2902479
H 15.605773 -10.5327737 -53.7532482
H 17.1389073 -11.4070984 -53.8607548
N 15.3935547 -6.0141343 -53.87863
H 15.4044782 -5.57695 -54.785056
C 15.4989416 -5.1573755 -52.7005961
H 16.1260958 -5.6549509 -51.9609007
C 14.1275042 -4.979588 -52.0493859
O 14.0041547 -5.2410058 -50.8625342
C 16.1529791 -3.8128398 -53.0612849
H 15.5453294 -3.3096024 -53.8136461
H 16.1749842 -3.1849001 -52.1697456
C 17.5888162 -3.9565456 -53.5969701
H 17.9604463 -2.9636573 -53.8583359
H 17.5825816 -4.5500643 -54.513358
C 18.5239075 -4.5941443 -52.560969
O 18.5957494 -5.8428049 -52.4662396
O 19.14147 -3.8633975 -51.7551244
N 13.0718701 -4.7526557 -52.834066
H 13.246795 -4.5814834 -53.8192455
C 11.6674641 -4.7573474 -52.3915481
H 11.5110988 -3.9424476 -51.6838844
C 11.2638044 -6.055375 -51.6705936
O 10.7000723 -5.9916155 -50.5815018
C 10.8225721 -4.4658085 -53.6373247
H 11.2703018 -4.9725609 -54.4847155
H 10.8980451 -3.3996945 -53.8162933
C 9.3400869 -4.8422281 -53.6032018
O 8.5960027 -4.4336681 -52.6834481
O 8.8625046 -5.4577929 -54.5871172
N 11.6917214 -7.2238873 -52.1577427
H 12.1378338 -7.2256039 -53.0701622
C 11.4803921 -8.5084706 -51.4804795
H 10.4192842 -8.5902366 -51.2439924
C 12.2581264 -8.616735 -50.1576347
O 11.7173718 -9.1125472 -49.1729519
C 11.8590517 -9.6523727 -52.4351888
H 12.9441046 -9.6885066 -52.5437256
H 11.4310558 -9.4628766 -53.4176041
C 11.3583046 -11.0199242 -51.9562242

H 11.7482812 -11.2164564 -50.9570363
H 11.7634607 -11.7769659 -52.6281464
S 9.5513028 -11.2041661 -51.9296248
C 9.4280423 -12.9497713 -51.463499
H 8.3802362 -13.2482151 -51.4521366
H 9.8585708 -13.0948903 -50.4720088
H 9.9653893 -13.5614913 -52.1881289
N 13.4992124 -8.1155487 -50.1013757
H 13.8751445 -7.7085546 -50.9531837
C 14.3462607 -8.0992723 -48.8917458
H 14.3261654 -9.0880359 -48.4302966
C 13.8414349 -7.109273 -47.8321094
O 13.8696885 -7.4088854 -46.6438106
C 15.7803139 -7.7771225 -49.3386629
H 16.0728206 -8.4953921 -50.1072518
H 15.78842 -6.7853287 -49.7877521
C 16.8146607 -7.8333564 -48.2019743
H 16.5459291 -7.1366515 -47.4067893
H 16.8280532 -8.8437172 -47.7908348
C 18.2187971 -7.4873246 -48.7193819
H 18.4184365 -8.0829343 -49.6128579
H 18.9536731 -7.7499724 -47.9570584
C 18.3487833 -5.992413 -49.0458139
H 18.4319706 -5.4217815 -48.1165061
H 17.4451803 -5.6561907 -49.5619324
N 19.5031294 -5.7293883 -49.9309239
H 20.3954323 -5.9966135 -49.5549232
H 19.5174901 -4.749167 -50.2346865
H 19.3464276 -6.1609146 -50.848426
N 13.3322727 -5.9630881 -48.2710839
H 13.3916351 -5.7966786 -49.2700395
C 12.6752334 -4.9259791 -47.4652083
H 13.3066394 -4.639705 -46.6204001
C 11.3514425 -5.4659331 -46.907204
O 11.0801258 -5.3692947 -45.708615
C 12.463281 -3.6936637 -48.3766892
H 12.0638674 -4.0566426 -49.3241162
C 11.3939495 -2.735481 -47.8251092
H 11.5699158 -2.5482408 -46.7698325
H 10.4027027 -3.1721763 -47.9439616
H 11.4167651 -1.795219 -48.3641339
C 13.8090055 -3.0011842 -48.7056933
H 14.1943818 -2.5034339 -47.8276267
H 14.5553025 -3.7476582 -48.9678682
C 13.7415405 -1.9940272 -49.8643963
H 13.1388901 -1.128049 -49.5957665

H 13.3172396 -2.4665957 -50.7493734
H 14.7489847 -1.6485579 -50.0979748
N 10.5569209 -6.1251239 -47.7575478
H 10.8058184 -6.1502801 -48.7417588
C 9.3406062 -6.8155937 -47.3499945
H 8.687436 -6.0980664 -46.8515223
C 9.6451199 -7.9320667 -46.3426887
O 8.9789309 -7.9857565 -45.3181334
C 8.6090851 -7.3439192 -48.5900398
H 9.2347282 -8.0646011 -49.1166765
H 8.3743305 -6.5167723 -49.2610325
H 7.6825766 -7.8315087 -48.286617
N 10.6967849 -8.7327052 -46.5459848
H 11.1877227 -8.6812436 -47.4322827
C 11.1410652 -9.7413254 -45.5766499
H 10.3095113 -10.4188421 -45.3765601
C 11.5417736 -9.1181235 -44.2316211
O 11.0027769 -9.5243077 -43.2065835
C 12.2879134 -10.5821685 -46.1574739
H 13.1383181 -9.9410307 -46.3900585
H 11.9539012 -11.054893 -47.0827182
C 12.7241101 -11.6616207 -45.1622198
O 13.5224159 -11.3337599 -44.252894
O 12.2018212 -12.793503 -45.265959
N 12.3829782 -8.0769071 -44.2118859
H 12.8033958 -7.7736289 -45.0842588
C 12.8418208 -7.4756884 -42.9545116
H 13.20788 -8.2795145 -42.315361
C 11.7105569 -6.7843906 -42.179033
O 11.5105007 -7.0762136 -40.995856
C 14.0107534 -6.5158331 -43.2086705
H 14.8199205 -7.0620265 -43.6953675
H 13.6867322 -5.7238326 -43.8863098
C 14.5352889 -5.889822 -41.9505213
C 15.3513898 -6.4920806 -41.0560262
H 15.7532765 -7.4912347 -41.1718495
N 15.5335855 -5.6654725 -39.9629168
H 16.0897329 -5.9158626 -39.160673
C 14.8148192 -4.49376 -40.0872147
C 14.6321954 -3.3866221 -39.242576
H 15.1032791 -3.3500841 -38.2735638
C 13.819259 -2.3236695 -39.676404
H 13.6551516 -1.4597462 -39.0434416
C 13.2231146 -2.3758491 -40.9490175
H 12.61592 -1.5480294 -41.2963996
C 13.413713 -3.4925454 -41.785488

H 12.9690569 -3.4997165 -42.7718436
C 14.1985453 -4.5894945 -41.3712467
N 10.9043925 -5.9527161 -42.8492866
H 11.1134384 -5.7630804 -43.8240366
C 9.7438538 -5.2921988 -42.2212211
H 10.0844229 -4.773692 -41.3238945
C 8.6828352 -6.2978319 -41.7797415
O 8.1475363 -6.1684389 -40.6765001
C 9.0796136 -4.2487583 -43.1304959
H 8.177974 -3.8895772 -42.6371761
C 9.9720763 -3.0467225 -43.4104666
H 10.8669468 -3.3593763 -43.9445606
H 10.2661708 -2.5763767 -42.4714854
H 9.4314004 -2.3171273 -44.0117579
O 8.7181591 -4.8066588 -44.3726664
H 9.5496823 -4.9479732 -44.8577988
N 8.4358032 -7.3581704 -42.5633335
H 8.8963046 -7.4314515 -43.466883
C 7.5723311 -8.4495715 -42.1268535
H 6.6350766 -8.0155664 -41.7795424
C 8.1700975 -9.1517994 -40.9247922
O 7.4758181 -9.174459 -39.9330543
C 7.2448461 -9.4620492 -43.2330206
H 8.1639771 -9.7685181 -43.7320877
H 6.8107539 -10.3548548 -42.779457
C 6.239774 -8.9211213 -44.2597927
H 6.2060887 -9.6067299 -45.1067549
H 6.5649357 -7.9487869 -44.6272582
C 4.8301142 -8.7907822 -43.6962677
O 4.0185217 -9.6936336 -43.7800733
N 4.4373901 -7.6329512 -43.2092787
H 3.4549186 -7.5555601 -42.9669857
H 5.0762198 -6.8618807 -43.140931
N 9.4273277 -9.6006942 -40.9284614
H 9.9686587 -9.4839231 -41.780828
C 10.0214635 -10.412134 -39.8567427
H 9.3084547 -11.1990181 -39.609992
C 10.253828 -9.6601678 -38.5291229
O 10.2250454 -10.2893742 -37.4666527
C 11.2944858 -11.0964365 -40.3945858
H 11.8618845 -11.5134649 -39.5643313
H 11.9244235 -10.3674132 -40.904203
C 10.9694015 -12.2521054 -41.3360884
O 10.0165345 -12.9890257 -41.1245311
N 11.7728694 -12.4937022 -42.343769
H 11.4995019 -13.1733068 -43.0362986

H 12.499172 -11.8467642 -42.6449554
N 10.2929531 -8.3240194 -38.537167
H 10.3624872 -7.8437779 -39.4304161
C 10.2075646 -7.5261957 -37.311539
H 10.9370343 -7.9083139 -36.5955343
C 8.8173012 -7.6078404 -36.6366541
O 8.7224682 -7.6976754 -35.4119872
C 10.5911494 -6.0827475 -37.6571958
H 9.8848154 -5.6694188 -38.3793169
H 11.5923603 -6.0574983 -38.0909011
H 10.5793001 -5.471428 -36.7549126
N 7.7281362 -7.648853 -37.4093658
H 7.8464977 -7.6620426 -38.4175533
C 6.3615845 -7.7065981 -36.8837774
H 6.2525433 -6.8732437 -36.1891368
C 5.9991926 -8.9928891 -36.0905116
O 5.3798625 -8.8602907 -35.0312186
C 5.3875801 -7.4605357 -38.0417105
H 5.317507 -8.3466364 -38.6717451
H 5.7360643 -6.6360806 -38.6607618
H 4.3964641 -7.2320804 -37.651936
N 6.3537059 -10.227742 -36.5201682
C 6.0921003 -11.4546801 -35.7992746
H 5.0595046 -11.4728627 -35.4552056
C 7.0141917 -11.5868196 -34.5825291
O 6.5442037 -12.022288 -33.5387212
C 6.2944393 -12.5878507 -36.8162142
H 6.5796654 -13.5239315 -36.3340225
H 5.3790469 -12.7278175 -37.3944718
C 6.9710996 -10.6108314 -37.7595143
H 6.2311127 -10.4906961 -38.55036
H 7.8621115 -10.0271148 -37.9027458
C 7.3852351 -12.0701274 -37.7139111
H 7.3632158 -12.5333561 -38.7012058
H 8.3603036 -12.1870356 -37.2408137
N 8.2600128 -11.0993172 -34.6473682
H 8.5974122 -10.7351963 -35.529587
C 9.1638845 -11.0324667 -33.493428
H 9.2559856 -12.0264496 -33.0531509
C 8.6246533 -10.1030979 -32.3918366
O 8.5898426 -10.4804776 -31.2234502
C 10.5463395 -10.5906214 -33.9929249
H 10.9400176 -11.3588206 -34.6604347
H 10.4349591 -9.6795107 -34.5821898
C 11.5645544 -10.3336599 -32.8982995
C 12.1676299 -11.4139774 -32.226674

H 11.9007571 -12.4282594 -32.4853161
C 13.115578 -11.1764532 -31.2147571
H 13.5750579 -12.007349 -30.6985902
C 13.464218 -9.8578058 -30.873536
H 14.194 -9.6750549 -30.0974929
C 12.8624226 -8.7769348 -31.541472
H 13.1324783 -7.7636193 -31.281157
C 11.9107974 -9.0133361 -32.5496745
H 11.4482941 -8.1777601 -33.0568941
N 8.0603085 -8.9495652 -32.7631566
H 8.1434311 -8.665565 -33.7337643
C 7.4168677 -8.0211198 -31.8271501
H 7.9214617 -8.1082216 -30.8649975
C 5.9315139 -8.326438 -31.5456683
O 5.2535744 -7.4801421 -30.961817
C 7.5941536 -6.574922 -32.324633
H 7.124499 -6.4941971 -33.3064549
H 7.0538484 -5.9068789 -31.6550215
C 9.0136829 -6.0386909 -32.3987931
C 9.4463363 -5.359517 -33.5561945
H 8.7912065 -5.2787379 -34.4129759
C 10.7433545 -4.8150995 -33.6203321
H 11.075635 -4.3075551 -34.5142454
C 11.6148391 -4.9453172 -32.5180674
O 12.8717441 -4.435554 -32.5709646
H 13.0345853 -3.9454995 -33.3800472
C 11.174962 -5.6029107 -31.3511048
H 11.8453005 -5.6829437 -30.507972
C 9.8769556 -6.144496 -31.2893049
H 9.5563376 -6.6474406 -30.3872537
N 5.3871766 -9.481238 -31.956426
H 6.0033601 -10.1538206 -32.4006634
C 3.9274016 -9.6951489 -32.0875758
H 3.5788743 -9.0167038 -32.8656313
C 3.101727 -9.3795924 -30.8315989
O 1.9946272 -8.8568 -30.9597617
C 3.6184426 -11.1199467 -32.5819641
H 4.1373467 -11.2775613 -33.5237867
C 4.0262775 -12.229422 -31.6076246
H 3.4924745 -12.148523 -30.6645446
H 5.0959923 -12.1726016 -31.405786
H 3.820187 -13.1996915 -32.0601955
O 2.2341245 -11.2757929 -32.8256162
H 2.1031661 -11.0808149 -33.7675107
N 3.6312656 -9.6287895 -29.6257628
H 4.5649006 -10.0097299 -29.5872965

C 2.9405287 -9.3177676 -28.3687039
H 1.9221669 -9.6969241 -28.445873
C 2.8276563 -7.8017421 -28.1507249
O 1.7167741 -7.2833488 -28.1555535
C 3.6362754 -10.0368499 -27.1973246
H 3.6353806 -11.1098669 -27.3948457
H 4.6745595 -9.7054275 -27.1474315
C 2.978396 -9.7839054 -25.8233671
H 2.9966347 -8.7178673 -25.6005677
C 1.5310448 -10.2812004 -25.7697653
H 1.4817257 -11.3377576 -26.0293787
H 0.9096754 -9.6938931 -26.4446131
H 1.1425322 -10.1385847 -24.7604909
C 3.7720642 -10.514862 -24.7416685
H 3.7563083 -11.5903824 -24.9153329
H 3.3368301 -10.2987848 -23.7649057
H 4.8026104 -10.1599394 -24.7388168
N 3.9616382 -7.0915666 -28.1298216
H 4.8305866 -7.5964343 -28.1841824
C 4.0327294 -5.6286734 -27.9990707
H 3.5475118 -5.3373421 -27.066531
C 3.3223149 -4.9052738 -29.1588863
O 2.7887892 -3.8112358 -29.0000078
C 5.522688 -5.2420034 -27.9073454
H 5.9417744 -5.7125461 -27.0159418
H 6.0387796 -5.6508386 -28.77724
C 5.8148372 -3.7290084 -27.839953
H 5.4302001 -3.2507583 -28.740477
C 5.2052008 -3.0369301 -26.6265279
H 5.5270028 -3.5308572 -25.7079944
H 4.1165442 -3.0902357 -26.6670599
H 5.5002705 -1.9890255 -26.5938106
C 7.3263404 -3.5054407 -27.8182893
H 7.7487717 -3.9367071 -26.9075049
H 7.5460048 -2.4366408 -27.8259105
H 7.7977514 -3.9704279 -28.681937
N 3.3240981 -5.4922525 -30.3565642
H 3.8200164 -6.3736162 -30.4481514
C 2.6217824 -4.9657189 -31.5266407
H 2.8835245 -3.9148142 -31.6417372
C 1.0943839 -5.029999 -31.3514762
O 0.4108227 -4.0473604 -31.6319395
C 3.127867 -5.7370606 -32.7582961
H 4.209855 -5.607543 -32.8206349
H 2.9292394 -6.7975108 -32.6056175
C 2.5111026 -5.3318787 -34.1077532

H 1.4532277 -5.5947615 -34.1197899
C 2.6686143 -3.8435188 -34.4097948
H 3.7203921 -3.5715168 -34.3494023
H 2.0853599 -3.2554452 -33.704076
H 2.2981698 -3.6320755 -35.4139741
C 3.226156 -6.095398 -35.2194507
H 4.2560045 -5.7514367 -35.3113525
H 2.7083162 -5.937219 -36.1650524
H 3.2404669 -7.1588567 -34.9842874
N 0.5617721 -6.1321914 -30.8100106
H 1.1854655 -6.8955181 -30.565335
C -0.8774119 -6.2932057 -30.5294087
H -1.4450707 -5.8133694 -31.3273322
C -1.3387857 -5.5989339 -29.2472027
O -2.4972833 -5.2107496 -29.1799714
C -1.2259879 -7.7857952 -30.5059816
H -0.5395652 -8.3161481 -29.8425807
H -2.2423577 -7.9080194 -30.1304602
C -1.1550235 -8.3653732 -31.9235204
H -0.1595575 -8.2283015 -32.3457716
H -1.8592111 -7.825137 -32.5503861
C -1.517416 -9.8499764 -31.9535904
H -1.7789138 -10.1101266 -32.9786297
H -2.3926915 -10.0247935 -31.3275004
N -0.3815104 -10.6888054 -31.5472774
H 0.4686067 -10.5891001 -32.0922565
C -0.2819976 -11.5284697 -30.5397689
N 0.7987728 -12.2399241 -30.4453776
H 0.9213052 -12.9191248 -29.7195572
H 1.4926744 -12.1243647 -31.1788985
N -1.2140364 -11.691179 -29.6413828
H -2.0524106 -11.1461476 -29.7056909
H -1.0837421 -12.3425586 -28.8909044
N -0.4359914 -5.4137744 -28.291648
H 0.4565919 -5.878876 -28.4106519
C -0.6121594 -4.6637386 -27.0405899
H -1.5484464 -4.9448234 -26.5574101
C -0.6419177 -3.1442925 -27.2699944
O -1.4719593 -2.454789 -26.6849815
C 0.5604857 -5.0831334 -26.1452238
H 0.4619065 -6.1502756 -25.9371324
H 1.4675594 -4.941078 -26.7285774
C 0.7664583 -4.3591921 -24.8161269
H -0.1072872 -4.5150072 -24.1788573
H 0.8875321 -3.2879945 -24.9917974
C 2.035639 -4.9226683 -24.1581527

O 1.8929473 -5.6917077 -23.1831504
O 3.1384206 -4.6239782 -24.6732221
N 0.1917402 -2.6286023 -28.1822229
H 0.9142711 -3.2259607 -28.5641261
C 0.1832659 -1.2150978 -28.5658876
H 0.0037262 -0.6137299 -27.6722889
C -0.9246664 -0.8558517 -29.5768898
O -1.3427934 0.2986789 -29.6289793
C 1.5712314 -0.8568396 -29.111109
H 1.7755088 -1.4407739 -30.0082925
H 2.3275326 -1.0683264 -28.3535512
H 1.5991124 0.2052893 -29.3567835
N -1.3690744 -1.8151607 -30.399589
H -0.9570034 -2.7323259 -30.3107115
C -2.4122034 -1.6197109 -31.4130676
H -2.8401389 -0.6237023 -31.2850044
C -3.5838278 -2.5891446 -31.1989809
O -4.4984085 -2.2893246 -30.4387788
C -1.8158924 -1.6685056 -32.833723
H -1.3145361 -2.624073 -32.987596
H -2.63415 -1.6141187 -33.553035
C -0.8409829 -0.5505884 -33.1512774
C -1.3096435 0.649862 -33.7150379
H -2.3647469 0.7833783 -33.9051926
C 0.531843 -0.7025696 -32.8802269
H 0.8864084 -1.6149005 -32.4260483
C -0.4110316 1.6879453 -34.01348
H -0.7806818 2.6123909 -34.4308109
C 1.4300197 0.3388147 -33.1757978
H 2.4760809 0.2347951 -32.9481857
C 0.9602995 1.5313944 -33.7512922
H 1.6466328 2.3356076 -33.9776088
N -3.6205628 -3.7132496 -31.9236746
H -2.8479066 -3.9034345 -32.5436994
C -4.6046085 -4.7861899 -31.7255437
H -4.6477881 -5.0219769 -30.6606922
C -4.2176418 -6.0541322 -32.4919207
O -3.4466607 -6.006089 -33.4557384
C -6.0082382 -4.3507814 -32.1811528
H -6.3304691 -3.4699273 -31.624698
H -6.7175425 -5.152142 -31.9705056
O -6.0325689 -4.0692292 -33.5679786
H -5.899994 -3.1163384 -33.6518697
N -4.8741413 -7.1737827 -32.163381
H -5.4716506 -7.1487112 -31.3494907
C -4.8665038 -8.4037354 -32.9751168

H -3.8421984 -8.7622554 -33.024553
C -5.296619 -8.1459118 -34.4291164
O -4.6251069 -8.5946963 -35.3565651
C -5.7191938 -9.4992887 -32.2851475
H -5.2617698 -9.6962688 -31.3135578
C -7.1749464 -9.0643969 -32.0189038
H -7.7180113 -8.9252308 -32.9539572
H -7.2157387 -8.1483551 -31.4309694
H -7.6857446 -9.8419113 -31.4478146
C -5.7591062 -10.8389807 -33.0521625
H -6.3486961 -11.5529072 -32.4750014
H -6.2623358 -10.6958331 -34.0075932
C -4.3868889 -11.4732058 -33.3065014
H -3.86465 -11.6184485 -32.3614838
H -3.7929475 -10.8453952 -33.9693974
H -4.5241808 -12.4437534 -33.7840225
N -6.3197801 -7.3081212 -34.6459337
H -6.7867577 -6.9174776 -33.8407924
C -6.8108968 -6.9429317 -35.9822531
H -6.9295796 -7.8697976 -36.5438468
C -5.8010042 -6.0946841 -36.7703672
O -5.7534856 -6.2049544 -37.9993011
C -8.1929232 -6.2678168 -35.828368
H -8.8936588 -6.9935093 -35.4113768
H -8.1035006 -5.4462128 -35.1147961
C -8.795135 -5.6932521 -37.1238768
H -9.7559562 -5.2332397 -36.8894435
H -8.1411857 -4.8942535 -37.4703831
C -9.0162358 -6.7421972 -38.2346595
H -9.9843958 -7.2211919 -38.076221
H -8.2596339 -7.5233973 -38.1830747
C -8.9707821 -6.1540308 -39.6519625
H -9.7832162 -5.4324325 -39.7803459
H -9.1312743 -6.9749907 -40.3575952
N -7.6582374 -5.5172426 -39.9201159
H -7.5238449 -4.6979612 -39.3466649
H -7.5232324 -5.260878 -40.8933727
H -6.8930147 -6.1433565 -39.6825251
N -4.9831557 -5.2914209 -36.0868715
H -5.1114733 -5.2316311 -35.0833861
C -3.8621657 -4.5557308 -36.6755995
H -4.2125805 -3.9476157 -37.5101316
H -3.4303927 -3.8987074 -35.9202956
C -2.7652385 -5.4947951 -37.177779
O -2.4814048 -5.5085537 -38.373279
N -2.2677174 -6.3806718 -36.3070245

H -2.5945535 -6.342489 -35.3479413
C -1.2607822 -7.3958332 -36.6512525
H -0.3445124 -6.8925585 -36.961776
C -1.6939054 -8.2984128 -37.8176158
O -0.9311845 -8.4717807 -38.7673839
C -0.9587692 -8.2239316 -35.3945171
H -0.5126998 -7.5591821 -34.6545479
H -1.9003645 -8.5918536 -34.9852958
C -0.0205586 -9.403772 -35.5957312
C -0.476939 -10.7237244 -35.4080804
H -1.509788 -10.9114615 -35.1526382
C 0.4018223 -11.8087914 -35.5828742
H 0.0411055 -12.8215519 -35.4605813
C 1.7396798 -11.5814567 -35.9510346
H 2.4073452 -12.4177367 -36.1126058
C 2.197147 -10.2669613 -36.1448257
H 3.2167878 -10.0866761 -36.4561706
C 1.3158525 -9.1847377 -35.975059
H 1.6594669 -8.1812815 -36.1674815
N -2.9508894 -8.7602365 -37.8162996
H -3.5180262 -8.6013591 -36.9897401
C -3.5164419 -9.5705426 -38.9055008
H -2.8830213 -10.4489155 -39.0412821
C -3.4884375 -8.8191223 -40.2441416
O -3.0854456 -9.4032043 -41.2444796
C -4.9353343 -10.0606912 -38.5229828
H -5.4930771 -9.2156787 -38.1147458
C -5.7199256 -10.6008875 -39.7362459
H -5.1748012 -11.428586 -40.1934969
H -5.8704642 -9.8203191 -40.4808494
H -6.7059161 -10.950001 -39.4339102
C -4.8187313 -11.1576382 -37.4374046
H -4.3660146 -12.0503218 -37.8713733
H -4.1598068 -10.814504 -36.6407699
C -6.1553173 -11.5460725 -36.7923586
H -6.7955629 -12.0580817 -37.5094256
H -6.6616767 -10.6550823 -36.4198857
H -5.9674939 -12.2254155 -35.9610021
N -3.8323925 -7.5248185 -40.2879394
H -4.0656676 -7.0556824 -39.4218744
C -3.7691065 -6.745843 -41.5391746
H -4.3005425 -7.2994218 -42.3139513
C -2.3393446 -6.5519942 -42.0526116
O -2.1145754 -6.6247537 -43.2621331
C -4.4332741 -5.3718712 -41.3703427
H -3.8537605 -4.6109815 -41.8970195

H -4.4733346 -5.1006908 -40.3138927
O -5.7412933 -5.3987764 -41.9212897
H -5.6390313 -5.3843889 -42.8827614
N -1.3861787 -6.3126124 -41.1495095
H -1.6547347 -6.2826085 -40.1714205
C 0.0158965 -6.0425797 -41.481886
H 0.0577829 -5.3112942 -42.2894989
C 0.740751 -7.3012983 -41.9896076
O 1.4825599 -7.2298511 -42.9719905
C 0.6993255 -5.4537121 -40.2332656
H 0.5759614 -6.161702 -39.4119538
H 1.768021 -5.3583632 -40.4270006
C 0.1460188 -4.0790425 -39.7977291
H -0.9410754 -4.0968931 -39.8068985
C 0.5881962 -3.7787531 -38.3651539
H 1.6688825 -3.8436922 -38.2817429
H 0.1358835 -4.4987693 -37.6855946
H 0.2728934 -2.7773283 -38.0815491
C 0.5838749 -2.9467471 -40.7300837
H 1.6679734 -2.8669288 -40.751094
H 0.1714852 -2.0035095 -40.3724266
H 0.2150466 -3.1247907 -41.7383577
N 0.4550266 -8.4611916 -41.3890336
H -0.1379686 -8.4416052 -40.564342
C 0.9646168 -9.7595951 -41.8364496
H 2.0196674 -9.6433663 -42.0892642
C 0.2485841 -10.2596498 -43.1035624
O 0.8987943 -10.7616169 -44.0158858
C 0.8455888 -10.7511466 -40.6638834
H 1.3523783 -10.326376 -39.7955445
H -0.210257 -10.8640917 -40.4110157
C 1.4357562 -12.1477194 -40.9424942
H 0.9147976 -12.6038002 -41.7839374
C 2.9345315 -12.1074372 -41.2461209
H 3.4682867 -11.5756565 -40.4587843
H 3.1048157 -11.6059846 -42.1990644
H 3.3258448 -13.1209115 -41.3293987
C 1.228386 -13.036948 -39.7157139
H 1.7534838 -12.6171902 -38.8578697
H 1.6078131 -14.0384338 -39.9180822
H 0.1647686 -13.1057515 -39.4881301
N -1.0698244 -10.0583188 -43.2209953
H -1.5784126 -9.6705319 -42.4337335
C -1.8288138 -10.4543792 -44.4133578
H -1.6406428 -11.512455 -44.6026739
C -1.3632127 -9.6903915 -45.6534104

O -1.0336879 -10.3147171 -46.6612399
C -3.3368109 -10.2801862 -44.1613091
H -3.5384081 -9.2595103 -43.8397539
H -3.6148275 -10.9493552 -43.3461491
C -4.2354917 -10.6192243 -45.3629775
H -5.2387681 -10.8120393 -44.981554
H -3.8764807 -11.5469091 -45.8110439
S -4.390968 -9.3657168 -46.6781462
C -5.1799099 -8.0079527 -45.7729399
H -5.4383667 -7.2116522 -46.4701612
H -4.4931662 -7.6138861 -45.0242914
H -6.0870682 -8.3705426 -45.2881865
N -1.2836702 -8.3547994 -45.597801
H -1.5351508 -7.8696527 -44.7427603
C -0.9788531 -7.5879528 -46.8043397
H -1.457325 -8.1034868 -47.6360249
C 0.521404 -7.5756896 -47.1451478
O 0.8637811 -7.5102657 -48.3264741
C -1.6430715 -6.2032923 -46.748782
H -1.1355764 -5.5871202 -46.0089988
H -2.6802938 -6.3144597 -46.4385527
C -1.6394516 -5.5151375 -48.1094585
N -0.9709235 -4.389742 -48.2157069
H -0.4685428 -4.0199304 -47.4270163
H -0.9501478 -3.9346266 -49.1103779
O -2.2486604 -5.9590541 -49.0785727
N 1.416904 -7.7516498 -46.1678263
H 1.0980991 -7.8154289 -45.2078644
C 2.8375119 -7.9866019 -46.4363406
H 3.1969886 -7.2710589 -47.1758242
H 3.3911814 -7.8211393 -45.5225839
C 3.1279278 -9.4043741 -46.945777
O 3.8298141 -9.5493483 -47.9453507
N 2.447086 -10.4225332 -46.4101552
H 1.9315207 -10.2624063 -45.5490469
C 2.4854058 -11.7979805 -46.9139969
H 3.5152832 -12.1562071 -46.9087821
H 1.8931677 -12.4278665 -46.2504679
C 1.9239234 -11.9295401 -48.3328251
O 2.5932635 -12.4826701 -49.2022689
N 0.7929469 -11.2777152 -48.628854
H 0.267872 -10.8632915 -47.8641561
C 0.2683398 -11.1228987 -49.9938683
H 0.0573556 -12.1067264 -50.4128921
C 1.2826057 -10.4351154 -50.919821
O 1.4660857 -10.8427349 -52.0678355

C -1.03995 -10.3126457 -49.9511539
H -1.8060414 -10.9034882 -49.4476874
H -0.8743164 -9.4136616 -49.3588149
C -1.5543657 -9.8836418 -51.2950987
C -1.1187229 -8.7991881 -51.9758444
H -0.3563124 -8.1147227 -51.6133052
N -1.6791194 -8.7882528 -53.2363242
H -1.3736528 -8.1625098 -53.9676252
C -2.5438482 -9.8457334 -53.4186151
C -3.3411695 -10.2526051 -54.4992947
H -3.3476281 -9.6931163 -55.422582
C -4.1539985 -11.3894695 -54.358604
H -4.801853 -11.6994674 -55.1695271
C -4.1314936 -12.1172815 -53.1550939
H -4.7589197 -12.9925633 -53.044334
C -3.3101688 -11.7105559 -52.0842815
H -3.3046579 -12.2804389 -51.1652787
C -2.499787 -10.5592805 -52.1831049
N 1.96428 -9.3993704 -50.4253499
H 1.7809878 -9.1156849 -49.4709382
C 2.9636726 -8.654149 -51.2011413
H 2.5137098 -8.3680451 -52.1510126
C 4.1751778 -9.5252683 -51.5380323
O 4.584636 -9.5468988 -52.6991712
C 3.3562456 -7.3613529 -50.4714756
H 3.7076416 -7.5770001 -49.4645344
C 4.4220107 -6.5580398 -51.2039862
H 4.0784879 -6.3762013 -52.2179027
H 5.3645424 -7.1059384 -51.2128172
H 4.574357 -5.6062331 -50.6915057
O 2.2203371 -6.5288334 -50.4111686
H 1.6945377 -6.8265429 -49.6442254
N 4.6517677 -10.3483214 -50.5991615
H 4.2697926 -10.2894075 -49.6593257
C 5.6772822 -11.3580867 -50.8568759
H 6.5451977 -10.8700353 -51.3041686
C 5.1923812 -12.4341222 -51.8490049
O 5.8968642 -12.7354913 -52.8110289
C 6.1067911 -11.9647842 -49.5145566
H 5.2580115 -12.4435631 -49.0255728
H 6.499734 -11.1841373 -48.8617151
H 6.8863492 -12.7084471 -49.6781218
N 3.9534366 -12.9249302 -51.7086818
H 3.4267844 -12.6496469 -50.8836684
C 3.334672 -13.8997941 -52.6291921
H 3.9737587 -14.7791788 -52.6745089

C 3.2311635 -13.3581777 -54.0601941
O 3.501972 -14.0879971 -55.0132972
C 1.9611062 -14.3607064 -52.0920771
H 1.4005589 -13.4924383 -51.7508018
C 1.0987965 -15.094003 -53.1308964
H 1.6582642 -15.9270397 -53.5595551
H 0.8015677 -14.4086898 -53.9226587
H 0.1946828 -15.4764729 -52.6554527
C 2.1541222 -15.3188572 -50.9094302
H 2.6359165 -16.2380737 -51.2437664
H 1.18814 -15.559973 -50.4658729
H 2.7765934 -14.8574841 -50.1444956
N 2.9105554 -12.073782 -54.2549198
H 2.6628376 -11.510579 -53.4444772
C 2.8382215 -11.4848562 -55.603466
H 2.5477263 -12.2731829 -56.294507
C 4.1932573 -11.0523229 -56.1735234
O 4.3050569 -11.0014922 -57.3976465
C 1.7500749 -10.3972881 -55.6684607
H 1.973677 -9.6224873 -54.932472
H 1.7766916 -9.9369756 -56.657886
C 0.3181127 -10.9321047 -55.4242165
H 0.2277581 -11.265099 -54.390261
H -0.3748739 -10.1024204 -55.5677658
C -0.0987512 -12.0894509 -56.3560436
H 0.544746 -12.9544503 -56.17014
H 0.0153415 -11.7777132 -57.39766
C -1.5577073 -12.5187353 -56.1029819
H -2.2269318 -11.6802266 -56.3325773
H -1.6736438 -12.7561101 -55.0366384
N -1.920309 -13.7026939 -56.930816
H -1.8494992 -13.5031153 -57.9228767
H -2.903941 -13.9547134 -56.8042702
H -1.3493329 -14.5134468 -56.71273
N 5.239108 -10.8811623 -55.358782
H 5.0832237 -10.9063656 -54.3572045
C 6.6252368 -10.7490509 -55.8568326
H 6.6136699 -10.1725473 -56.7835268
C 7.23075 -12.111499 -56.2120809
O 7.8163342 -12.237301 -57.2830009
C 7.5248285 -9.9796415 -54.8686851
H 7.4639628 -10.4501565 -53.8877875
C 8.9905939 -9.944826 -55.3227672
H 9.0568977 -9.5997377 -56.3555774
H 9.4372889 -10.9357012 -55.2428839
H 9.5579733 -9.2635562 -54.6896495

C 7.0793244 -8.5186274 -54.7449023
H 7.2214826 -8.0117064 -55.6995769
H 7.6884329 -8.0243809 -53.9867521
H 6.0354079 -8.4688468 -54.4529373
N 7.0278598 -13.1595392 -55.4024952
H 6.5409429 -13.011564 -54.5230944
C 7.6271667 -14.4858677 -55.6559459
H 8.6634613 -14.3031979 -55.9438593
C 6.9906641 -15.2279986 -56.8419336
O 7.6716931 -15.9879257 -57.5225639
C 7.6575082 -15.3317962 -54.3638388
H 7.9124323 -14.6589608 -53.5431137
C 6.2996944 -15.9707711 -54.0400428
H 6.0816508 -16.7851927 -54.7321286
H 5.5262014 -15.2171777 -54.1221082
H 6.2972443 -16.3543042 -53.0201916
C 8.777694 -16.3911983 -54.4438692
H 8.5331507 -17.1303691 -55.2074872
H 9.7068405 -15.8989223 -54.7355244
C 9.0351373 -17.1189259 -53.1182629
H 8.173149 -17.7267722 -52.8436591
H 9.2387219 -16.3944539 -52.3289923
H 9.8999306 -17.7735446 -53.2299662
N 5.7387061 -14.9039989 -57.1868191
H 5.2127234 -14.3222172 -56.5485866
C 5.1062204 -15.3009733 -58.4534771
H 5.3874356 -16.3318692 -58.6680816
C 5.6030433 -14.4874707 -59.672003
O 5.0849158 -14.6599443 -60.773668
C 3.5735904 -15.2270815 -58.2905951
H 3.3168007 -14.2373792 -57.910768
H 3.1057086 -15.3358337 -59.269651
C 2.9775319 -16.3018904 -57.3585176
H 3.4899355 -16.2918303 -56.3990126
C 1.4957103 -15.9962792 -57.1210999
H 0.9610944 -15.9934025 -58.0710052
H 1.4152946 -15.0272253 -56.6348142
H 1.0717324 -16.755584 -56.4636815
C 3.0815974 -17.7101515 -57.947436
H 2.616133 -17.7444842 -58.9324024
H 2.5847047 -18.4241014 -57.2906702
H 4.1275589 -18.0041337 -58.0324987
N 6.5584693 -13.5695965 -59.4840437
H 6.9635998 -13.4805897 -58.559394
C 7.1419561 -12.7294521 -60.533019
H 8.0300305 -12.2428227 -60.1296168

H 7.4428545 -13.3604889 -61.3694312
C 6.2113295 -11.6372743 -61.0664694
O 6.4668133 -11.1007314 -62.1414297
N 5.0989407 -11.3521821 -60.3807085
H 4.9813281 -11.7997437 -59.4822739
C 4.011173 -10.5016823 -60.8819061
H 4.0372055 -10.488733 -61.9692609
C 4.1458654 -9.0360507 -60.4550618
O 4.0311999 -8.1487806 -61.2999311
C 2.6640571 -11.1114873 -60.4638078
H 1.8926477 -10.351034 -60.4842377
H 2.730934 -11.4831747 -59.4414438
C 2.2535011 -12.2657448 -61.3868724
H 1.4591474 -12.8339579 -60.902344
H 3.1069454 -12.932034 -61.5231857
S 1.6498686 -11.7327541 -63.0139356
C 1.6214996 -13.3280329 -63.8756168
H 1.2174771 -13.1904539 -64.8799789
H 0.9970689 -14.0358844 -63.3303468
H 2.6341481 -13.7267153 -63.9450518
N 4.4403578 -8.7747006 -59.1755103
H 4.5741277 -9.5745046 -58.565276
C 4.7721047 -7.4432695 -58.6461105
H 4.2329189 -6.6929209 -59.2287529
C 6.2628089 -7.1513591 -58.8255713
O 7.1132365 -8.0286716 -58.7190631
C 4.3124856 -7.3225627 -57.1776623
H 3.2218766 -7.3201566 -57.1538076
H 4.664482 -8.1940433 -56.6248092
C 4.8413101 -6.0586761 -56.4669405
H 5.9269408 -6.1201353 -56.3940316
H 4.5835999 -5.1738691 -57.0510368
C 4.2767258 -5.8952971 -55.0518431
H 4.2724669 -6.8599056 -54.5417714
H 3.2469937 -5.5465683 -55.1283597
C 5.0870934 -4.8828455 -54.2236432
H 4.4787623 -4.580427 -53.3658758
H 5.2958632 -3.991815 -54.8235343
N 6.3573003 -5.466491 -53.7188249
H 6.2395348 -6.3113772 -53.1868018
H 6.9070721 -4.8226841 -53.1439003
H 7.0461657 -5.6510471 -54.4521823
N 6.5831059 -5.8796803 -59.0166479
H 5.8235267 -5.2105063 -59.113986
C 7.9375657 -5.3227778 -59.0310328
H 8.6143732 -5.9856498 -58.4939189

C 7.9299697 -3.9753104 -58.3016906
O 6.8849011 -3.3299543 -58.2056804
C 8.3940536 -5.2114097 -60.498702
H 7.5989185 -4.7461053 -61.083704
C 9.6867695 -4.4393521 -60.7534294
H 10.461601 -4.7634189 -60.0624864
H 9.5092652 -3.3712309 -60.637248
H 10.0261984 -4.6212524 -61.7728531
O 8.6164189 -6.5095718 -60.9963761
H 8.1255677 -7.1366062 -60.4489059
N 9.0868513 -3.5097747 -57.8288001
H 9.9133854 -4.0836863 -57.9019445
C 9.2842826 -2.0970461 -57.484942
H 8.3344924 -1.5820776 -57.6283965
C 10.2397767 -1.4588863 -58.488381
O 11.1822076 -2.0878065 -58.9748673
C 9.6723067 -1.8988096 -55.9998239
H 9.4745394 -2.8358148 -55.4851521
C 11.1589555 -1.5434698 -55.8111287
H 11.3819425 -0.5661663 -56.2441685
H 11.7740098 -2.3086833 -56.2769605
H 11.4226248 -1.4985324 -54.7557414
C 8.7636256 -0.8305709 -55.3496209
H 8.9602225 0.1445593 -55.7991688
H 7.7226656 -1.0931424 -55.5430241
C 8.9349449 -0.7232145 -53.8274753
H 9.9006709 -0.2819557 -53.5826193
H 8.8562241 -1.7099889 -53.3708842
H 8.1532925 -0.0869728 -53.4164581
N 9.9517901 -0.2103244 -58.8094482
H 9.1682783 0.231535 -58.3373574
C 10.654951 0.6360141 -59.7608788
H 11.4393476 0.0802286 -60.2761536
C 11.3137825 1.7707113 -58.9834636
O 10.6744127 2.3280577 -58.0948429
C 9.5994166 1.0937026 -60.7929687
H 8.6089785 1.028997 -60.3467226
H 9.6158493 0.3827572 -61.6188321
C 9.6756245 2.4856417 -61.3678588
N 9.6560584 2.7683857 -62.7304314
C 9.6304442 4.1041051 -62.8320208
H 9.6677495 4.647484 -63.7669766
N 9.599696 4.6686446 -61.6145544
H 9.6754936 5.6576488 -61.4159251
C 9.6037052 3.6646771 -60.6808175
H 9.5692624 3.7963986 -59.6106405

N 12.5267095 2.1509732 -59.3842939
H 13.0058659 1.5797279 -60.0714761
C 13.1036285 3.4786336 -59.1419166
H 12.3791542 4.0801703 -58.6065819
C 13.4003518 4.137936 -60.4980748
O 13.9804095 3.4895145 -61.3751488
C 14.3486196 3.3947989 -58.2404563
H 14.0355429 3.0345998 -57.2602812
H 15.0443904 2.6671256 -58.661884
C 15.0998792 4.7290328 -58.0561606
H 15.5972644 4.9762276 -58.9978091
H 15.8861092 4.5829474 -57.3125827
C 14.2028056 5.9034645 -57.6227446
O 14.1451209 6.8963968 -58.3833069
O 13.5576779 5.8054743 -56.5593021
N 13.0453446 5.4126846 -60.6712948
H 12.6425479 5.9066621 -59.8759317
C 13.3632583 6.2062371 -61.8533931
H 14.1139036 5.6662568 -62.4167197
C 13.9496037 7.5757594 -61.5001384
O 13.3110616 8.3851175 -60.8299668
C 12.1357501 6.3519771 -62.752082
H 11.3121412 6.7960327 -62.1946084
H 11.8710934 5.3727228 -63.1411174
H 12.3807769 6.9994993 -63.5956822
N 15.109189 7.8745898 -62.0873356
H 15.4773862 7.2053572 -62.7563538
C 15.8432781 9.1316037 -61.9126432
H 15.2448618 9.8123746 -61.30726
C 16.0561624 9.7692783 -63.2824116
O 16.4188304 9.0839867 -64.2349318
C 17.1881392 8.8969502 -61.188147
H 17.8076121 8.2561194 -61.8169794
C 16.9764192 8.1765845 -59.8366834
H 16.3622953 8.7964619 -59.1838709
H 16.4432474 7.2414061 -60.0042166
C 17.9261826 10.2389898 -60.9959842
H 17.3004281 10.929227 -60.4298202
H 18.1645092 10.6855659 -61.9607414
H 18.8686102 10.0894265 -60.4738383
C 18.2749524 7.8099776 -59.110778
H 18.7708445 8.7035771 -58.7336403
H 18.9412565 7.2714474 -59.7855677
H 18.0370505 7.1684199 -58.2616609
N 15.8801458 11.0823937 -63.3805037
H 15.5331748 11.5872138 -62.5688742

C 15.9623305 11.817571 -64.6442913
H 16.7952455 11.4316431 -65.2351664
C 16.2306405 13.3158319 -64.4223685
O 16.001996 13.8407697 -63.3256233
C 14.6780471 11.5721577 -65.4405948
H 14.5614297 12.3592306 -66.1828294
H 14.8143697 10.6397859 -65.9771394
C 13.3981278 11.4233831 -64.5967226
H 13.2409175 12.3213791 -64.0054631
H 13.4843556 10.5777558 -63.912386
C 12.2010246 11.1445692 -65.486179
O 12.4220538 10.9836834 -66.7062945
O 11.0711677 11.0577901 -64.952999
N 16.7127389 14.0282412 -65.4533237
H 16.8577185 13.566903 -66.349382
C 16.9273486 15.4814323 -65.3898288
H 16.3871383 15.8476574 -64.5193038
C 16.3447654 16.2645858 -66.5817988
O 16.1558116 15.7419953 -67.6835179
C 18.4109936 15.8096423 -65.1287159
H 18.8261429 15.0930979 -64.4212154
H 18.4804463 16.7986531 -64.677849
C 19.2719235 15.817902 -66.3805842
O 19.9812442 14.8726451 -66.670495
N 19.2629312 16.8803734 -67.1552989
H 19.7115882 16.7659892 -68.0471681
H 18.6938739 17.6945154 -66.9492704
N 16.1171737 17.5560555 -66.3461034
H 16.3244087 17.9218972 -65.4216912
C 15.6409335 18.5213459 -67.3311437
H 15.1563779 17.9880668 -68.1515813
C 16.8207639 19.3019001 -67.9050657
O 17.814395 19.5196466 -67.2156087
C 14.6025547 19.470826 -66.7156745
H 15.0457816 19.9951369 -65.8663646
H 13.7532703 18.8943344 -66.3687152
C 14.1049765 20.4776596 -67.7549781
O 14.0497 21.6808959 -67.4357038
O 13.8939207 20.0301459 -68.9036003
N 16.6932574 19.7127438 -69.1587056
H 15.7361606 19.6535177 -69.5221682
C 17.6417525 20.4891035 -69.9664808
H 18.1709276 19.7718774 -70.5775385
C 18.7397635 21.2279062 -69.1911158
O 19.6163392 20.5303848 -68.6231238
O 18.7905354 22.4280729 -69.0989548

C 16.8603575 21.4153103 -70.9236658
H 16.3055296 22.1240006 -70.3041713
H 17.5800612 21.9809911 -71.516076
C 15.8643249 20.7213124 -71.8783228
H 15.1037594 20.1933291 -71.2956912
C 15.1458514 21.7861056 -72.7164382
H 15.8579213 22.3362843 -73.3312233
H 14.6219113 22.4788679 -72.0561909
H 14.4058411 21.3071409 -73.3609419
C 16.5435476 19.7199673 -72.8232848
H 17.329409 20.2094041 -73.398468
H 15.8002973 19.3095617 -73.5106352
H 16.9602655 18.8885803 -72.2557724

**Supplementary Data 5. Cartesian coordinates for TePeNMT homology model with bound SAM.**

N -2.1385667 21.7753992 -15.9844882
H -2.8881317 22.3451992 -15.612073
H -2.4187043 21.4412796 -16.898962
H -1.952445 20.9940613 -15.3721405
C -0.8983927 22.6008534 -16.1048954
H -1.0805973 23.3575681 -16.8709468
C 0.33849 21.7897622 -16.5659561
O 1.2553163 22.327476 -17.1874333
C -0.6053818 23.3116885 -14.7771474
H 0.3668547 23.80077 -14.8479421
H -0.5567763 22.5711725 -13.9778747
C -1.639947 24.3835703 -14.4160168
H -1.396285 24.7758699 -13.426708
H -2.6273748 23.9290305 -14.3587376
S -1.7285502 25.7740286 -15.5733039
C -2.9534829 26.7844932 -14.6928753
H -3.1342375 27.7045542 -15.2488035
H -2.5798462 27.0332981 -13.6984196
H -3.8891334 26.2323095 -14.5953778
N 0.3673335 20.4886197 -16.2657559
H -0.4109708 20.1080689 -15.7405706
C 1.3938537 19.5859753 -16.7771625
H 1.3580658 19.6103407 -17.8672226
H 2.3753087 19.9422459 -16.4680576
C 1.2496557 18.1385969 -16.3224611
O 0.7986647 17.8537127 -15.2154758
N 1.7628938 17.2390447 -17.1522057
H 2.1121366 17.5432004 -18.0501647
C 2.0288584 15.8382482 -16.8338954
H 1.9028599 15.7027836 -15.7604402
C 3.4957816 15.5076051 -17.1394012
O 4.119702 16.1812749 -17.9609391
C 0.9990926 14.9223526 -17.5032517
H 1.2572628 13.8928438 -17.2749979
H 0.0423459 15.1358366 -17.026009
C 0.8149397 15.0455348 -19.020348
H 1.6157817 14.5050359 -19.5288645
H 0.8699863 16.0954136 -19.3176114
C -0.5554566 14.4754363 -19.4180847
O -1.297864 15.2053148 -20.1119062
O -0.875195 13.3478418 -18.9747065
N 4.1061184 14.7016468 -16.2632362
H 3.5442127 14.1102083 -15.6715657

C 5.5629684 14.6122266 -16.0983634
H 6.0446756 14.8186955 -17.0600251
C 5.9695982 13.2129132 -15.6736854
O 5.3463653 12.639753 -14.786269
C 6.0594875 15.5900786 -15.0182406
H 5.5966651 15.3285324 -14.0641537
H 7.1389711 15.4636292 -14.915048
C 5.7688522 17.0627619 -15.3210483
H 6.199365 17.3109927 -16.2919241
H 4.6914859 17.2284646 -15.351451
C 6.3694499 17.9615862 -14.2310701
H 7.4516987 17.8220036 -14.192695
H 5.9514537 17.6743696 -13.2638997
C 6.0413128 19.4366002 -14.4861744
H 6.4275873 20.029751 -13.6516003
H 4.9522862 19.5499365 -14.5026758
N 6.6253332 19.9107507 -15.7656037
H 7.6331593 19.818804 -15.7546612
H 6.3843249 20.8792368 -15.933991
H 6.2683797 19.3550584 -16.5356419
N 7.106455 12.7593095 -16.1705568
H 7.4956956 13.2759729 -16.9574346
C 7.5639831 11.383838 -16.027934
H 6.7220861 10.7521143 -15.7410336
C 8.6256294 11.3080201 -14.9245199
O 9.6775993 11.9537076 -14.9983873
C 8.0743326 10.8706348 -17.3888252
H 8.9570019 11.4416081 -17.6791626
H 8.3787829 9.8302572 -17.2664366
C 7.0169986 10.9370419 -18.5172246
H 6.1405201 10.371064 -18.1914962
H 7.409044 10.4393792 -19.4038394
C 6.5974706 12.3719046 -18.896483
O 7.4785217 13.2681533 -18.9084406
O 5.3745041 12.6072441 -18.9936412
N 8.3366279 10.5700175 -13.8495991
H 7.428984 10.1185224 -13.7997984
C 9.2537472 10.4628666 -12.7165948
H 9.411715 11.4704627 -12.3299929
C 10.6243327 9.9036971 -13.153139
O 10.6986872 8.9385328 -13.9069108
C 8.6037136 9.6350831 -11.6036772
H 8.4204313 8.619799 -11.9600852
H 7.655759 10.0896643 -11.3114457
H 9.2647124 9.595986 -10.737355
N 11.703524 10.5367378 -12.6721267

H 11.5296608 11.3030954 -12.0418231
C 13.1142625 10.3142897 -13.0702562
H 13.6516325 11.146085 -12.6173586
C 13.4243279 10.5119185 -14.5675031
O 14.4681525 10.0599575 -15.0339048
C 13.7765064 9.0619071 -12.4395625
H 14.845682 9.122573 -12.6445486
C 13.6265139 9.0675566 -10.9118445
H 12.5803642 8.9423089 -10.6318339
H 14.0018322 10.0070241 -10.506304
H 14.2061243 8.2486758 -10.485005
C 13.2962742 7.7038875 -12.9652798
H 12.2890587 7.4911381 -12.6080123
H 13.9637118 6.9192992 -12.6112727
H 13.2950871 7.7094786 -14.0550769
N 12.605294 11.3349009 -15.2378421
H 11.7369915 11.5814565 -14.7805248
C 12.7314974 11.833814 -16.6111805
H 11.8724211 12.4921825 -16.7495073
C 12.6154246 10.7757957 -17.7237248
O 13.1625965 9.6728211 -17.644985
C 13.9716254 12.728998 -16.7542726
H 14.876663 12.1276992 -16.6566322
H 13.9650252 13.5017634 -15.9855672
H 13.9695782 13.2008787 -17.7373648
N 11.9102878 11.1457732 -18.7984009
H 11.4901366 12.0606529 -18.8311174
C 11.6113386 10.2161504 -19.8849453
H 11.4683386 9.2575284 -19.3922061
C 12.7697246 10.041093 -20.8749029
O 13.4841971 10.9883786 -21.2103005
C 10.2956217 10.5834444 -20.6018401
H 9.5642602 10.8737195 -19.8541547
H 10.4613373 11.4376453 -21.2598264
C 9.7172444 9.4047201 -21.4196476
H 10.350257 9.2350742 -22.2929309
H 8.7362979 9.7002982 -21.7986705
C 9.5899082 8.089336 -20.6183328
O 8.4815808 7.7830257 -20.1380566
O 10.6205886 7.3866881 -20.4498584
N 12.8901662 8.8305339 -21.4204041
H 12.2017952 8.1247666 -21.1470783
C 13.7998552 8.4972353 -22.5092701
H 14.1771403 9.4236569 -22.9434863
C 13.0399482 7.741936 -23.6088019
O 12.2666108 6.8259826 -23.3460136

C 14.9936922 7.7225667 -21.9245386
H 15.5173435 8.3745069 -21.2235556
H 14.6001376 6.8863099 -21.3570609
C 16.0029798 7.189281 -22.9591753
H 15.4878043 6.568367 -23.6890142
C 16.7306364 8.3210255 -23.6896215
H 17.2504768 8.9574053 -22.9726436
H 16.0235264 8.9250527 -24.2541046
H 17.4549132 7.90003 -24.3877077
C 17.0509741 6.3236455 -22.2584349
H 17.5894864 6.9110609 -21.5149428
H 17.7563171 5.9395483 -22.9960493
H 16.5630007 5.4766885 -21.778768
N 13.2962296 8.1160457 -24.8611726
H 13.9627613 8.8548464 -25.0049569
C 12.7297022 7.4720004 -26.0467334
H 11.6552424 7.6561114 -26.0668372
C 12.963355 5.9483119 -26.0268958
O 14.0825531 5.4964434 -25.7932219
C 13.3669937 8.1475173 -27.2671617
H 13.1403425 9.2139322 -27.2266732
H 14.4513269 8.049724 -27.187207
C 12.9656811 7.6189319 -28.6307211
C 12.1494424 8.3971049 -29.4761888
H 11.769383 9.3526089 -29.1454154
C 11.8565891 7.9520163 -30.7798108
H 11.2562024 8.555452 -31.4472005
C 12.3673248 6.7178848 -31.2374061
O 12.1160781 6.2857259 -32.4969706
H 11.5991475 6.9488647 -33.0185918
C 13.1697677 5.9340538 -30.3847744
H 13.5492799 4.9896701 -30.7459872
C 13.4827043 6.3907892 -29.091865
H 14.1273567 5.791667 -28.4621886
N 11.8951892 5.1788313 -26.2599184
H 11.0304658 5.6427895 -26.4800882
C 11.8426092 3.7036502 -26.2137578
H 10.7811114 3.4599108 -26.2631227
C 12.3180866 3.0493723 -24.893297
O 12.588105 1.8486882 -24.8229072
C 12.4769853 3.1028593 -27.4818833
H 13.5610504 3.0598399 -27.3602348
H 12.2582179 3.7523311 -28.3311342
C 11.9188234 1.7098089 -27.8129528
O 10.6734795 1.5608478 -27.7641107
O 12.7303627 0.8270589 -28.169355

N 12.3786583 3.8077385 -23.7892188
H 12.1567416 4.7960764 -23.8565606
C 12.7236866 3.2518266 -22.4780661
H 13.5786598 2.5902866 -22.6181944
C 11.5733328 2.4149249 -21.9209309
O 10.5345585 2.9233827 -21.5059758
C 13.1586237 4.3717878 -21.5322934
H 14.0067916 4.8753226 -21.9927235
H 12.3398627 5.0838206 -21.4074609
C 13.5708913 3.8411374 -20.1519667
H 12.6903613 3.4453154 -19.6452956
H 14.3110593 3.0473288 -20.2626016
C 14.1595742 4.9730826 -19.3074994
H 13.477531 5.8273082 -19.3386531
H 15.1255757 5.265841 -19.7198963
C 14.3246592 4.5251755 -17.8558798
H 14.9647772 3.6393135 -17.8237746
H 13.3343658 4.2459127 -17.4769636
N 14.894404 5.6206571 -17.034387
H 15.8026652 5.9022239 -17.3743342
H 14.9585333 5.3479973 -16.0632593
H 14.281514 6.4323719 -17.0835563
N 11.8145313 1.1129027 -21.796752
H 12.636759 0.7428888 -22.2469702
C 10.915173 0.206206 -21.075752
H 9.9261211 0.3116122 -21.5233934
C 10.7879049 0.6022152 -19.5986625
O 11.7565775 1.0128309 -18.9601126
C 11.3345322 -1.270057 -21.22834
H 10.6296722 -1.8783407 -20.6606041
C 11.2418011 -1.7098514 -22.6941182
H 11.9608367 -1.1650701 -23.3087444
H 10.2395993 -1.5174528 -23.0780593
H 11.4468521 -2.7773516 -22.7753339
C 12.7458793 -1.5768076 -20.7095885
H 13.4907476 -1.0077613 -21.2656222
H 12.9570051 -2.6395233 -20.8284199
H 12.8214824 -1.3296274 -19.6506744
N 9.6090573 0.3641778 -19.0190877
H 8.8636369 0.042936 -19.61778
C 9.3030641 0.4995601 -17.5765515
H 8.2498576 0.2378134 -17.4888618
C 9.3464409 1.9102799 -16.9589934
O 9.2888891 2.018324 -15.7386553
C 10.0341763 -0.545724 -16.7016046
H 9.6693134 -0.4676256 -15.6770594

C 9.7794235 -1.9797159 -17.169852
H 10.215899 -2.1450359 -18.1540049
H 8.7079161 -2.1740235 -17.2032576
H 10.239578 -2.6699839 -16.4624216
O 11.4299275 -0.3613229 -16.691399
H 11.6667592 0.1678767 -17.4727227
N 9.2779939 2.9909397 -17.7450031
H 9.3809 2.8980378 -18.7455964
C 8.8924366 4.3186524 -17.2226461
H 9.484174 4.5459203 -16.3361664
C 7.3950813 4.3363773 -16.8420819
O 6.54833 4.1563548 -17.7157283
C 9.1737252 5.4060425 -18.2724269
H 8.8412563 6.3707971 -17.8859208
H 8.6096819 5.1895781 -19.1816057
O 10.5517975 5.4957348 -18.5958262
H 10.5894134 6.1754387 -19.3178923
N 7.0367162 4.4791013 -15.5515566
H 7.7637049 4.4744686 -14.8485576
C 5.6335974 4.372326 -15.0850094
H 5.0070584 4.6318068 -15.9400818
C 5.0839267 5.3356751 -13.9496445
O 4.1802507 4.8921545 -13.2441249
C 5.3501324 2.8742188 -14.7954481
H 5.6601937 2.2734319 -15.6495461
H 4.2770818 2.7385852 -14.6680957
C 6.0481724 2.3356799 -13.5563096
O 6.9355157 2.9397106 -12.9802825
N 5.6839717 1.1536912 -13.1130988
H 6.1721809 0.8278604 -12.2984982
H 4.9434468 0.6468731 -13.5561149
N 5.4209234 6.6581869 -13.7954432
H 6.1313111 7.0377768 -14.4037847
C 4.6727308 7.6690857 -12.8730958
H 3.6722803 7.2716519 -12.7031805
H 5.1665915 7.6425688 -11.902207
C 4.4605241 9.2667223 -13.2399755
O 5.4811072 9.6663177 -13.812121
N 3.2536409 10.1109433 -12.9872607
H 2.6474233 9.6614132 -12.3206553
C 2.3771526 11.4343133 -13.6922496
H 3.1658733 12.0715482 -14.0792353
C 1.0505375 12.8198719 -12.978889
O 0.738086 12.1140384 -12.0071596
C 1.9540762 10.4352165 -14.9544058
H 0.9329988 10.1097298 -14.746017

C 2.7417697 9.1177858 -15.5124699
H 3.7821969 9.3932306 -15.6797871
H 2.6711718 8.3582525 -14.7352443
C 1.9181844 11.302737 -16.1856385
H 2.9290002 11.590086 -16.4899791
H 1.3394983 12.1952761 -15.9637221
H 1.4171187 10.8031861 -17.0150758
C 2.3461044 8.3062146 -16.7841532
H 1.2838958 8.043668 -16.7411571
H 2.9411495 7.3919643 -16.8207085
H 2.5328117 8.8854332 -17.695951
N 0.2333932 14.4003043 -13.1834479
H 0.7437184 14.7653437 -13.9704046
C -1.2123799 15.955379 -12.9597387
H -1.0028175 16.3292374 -13.960482
C -2.5312399 15.1695335 -13.1010222
H -2.3258053 14.1092915 -13.2484796
H -3.0618878 15.2689053 -12.1555772
C -3.4175086 15.6977339 -14.2585728
H -3.2432872 16.7657087 -14.3985759
C -3.0783778 14.9714564 -15.5675978
H -3.2811317 13.903087 -15.4809576
H -2.0269104 15.1037593 -15.8111044
H -3.6757972 15.3801559 -16.3812018
C -4.9110054 15.512951 -13.9555661
H -5.1400087 14.4586235 -13.8201664
H -5.5040244 15.9052164 -14.7807417
H -5.172629 16.0607214 -13.0540348
C -2.0596333 17.9254177 -12.3430272
O -1.3832877 18.2475852 -11.374691
N -3.271554 19.2249776 -12.5340521
H -3.8017849 18.9469697 -13.3433045
C -4.1853409 20.9390259 -11.9780666
H -4.1201797 20.8378689 -10.8941409
C -5.8142116 21.9798425 -12.0685304
O -6.4577917 21.990306 -13.1238488
C -3.3553264 22.2522131 -12.3368714
H -2.7508435 21.9810248 -13.199742
H -3.9431639 23.1195955 -12.6376299
C -2.3935577 22.6639575 -11.2141059
H -1.5875927 23.2551301 -11.6539897
H -1.9417954 21.7737149 -10.7710014
C -3.1027948 23.4921229 -10.1365626
O -4.1338848 23.0043489 -9.6189633
O -2.6283562 24.6174618 -9.8712537
N -6.4475046 23.0719258 -11.2698861

H -5.8681384 23.316696 -10.4654281
C -7.9445496 23.8116708 -11.2481781
H -8.5313975 23.1447577 -11.8786809
C -8.4505552 25.3183528 -11.7282815
O -7.9562339 26.3541144 -11.2608184
C -8.5827225 23.7319299 -9.8142739
H -8.3098143 22.7644516 -9.3910452
H -8.1237068 24.4929663 -9.1820639
C -10.1128287 23.8553926 -9.699581
H -10.4295997 24.8665053 -9.9608487
H -10.3884539 23.7049408 -8.6526486
C -10.8425833 22.8299823 -10.5754585
O -10.9583405 23.1023724 -11.793048
O -11.2574013 21.7859035 -10.0265513
N -9.602907 25.5151302 -12.4609812
H -10.1970189 24.6947314 -12.6038089
C -10.0093518 26.7968804 -13.139427
H -9.6033406 27.6153906 -12.5465371
C -11.5373224 27.0770783 -13.3129338
O -12.3430523 26.1660776 -13.4651874
C -9.3158021 26.7963794 -14.5218177
H -8.2652178 26.5407625 -14.3723088
H -9.7616841 25.9988297 -15.1187812
C -9.3656862 28.1006173 -15.3396433
H -10.3980501 28.3527717 -15.5746801
C -8.7151959 29.2767986 -14.6053596
H -7.6855865 29.0272619 -14.3458923
H -9.2616884 29.5103425 -13.6946444
H -8.721151 30.1580805 -15.2468748
C -8.6202634 27.902131 -16.6598573
H -7.5714266 27.6723762 -16.4704162
H -8.6904418 28.808201 -17.2612677
H -9.0736317 27.0808258 -17.214682
N -11.9412789 28.3572898 -13.4297045
H -11.2467902 29.0828588 -13.3712445
C -13.3321225 28.7589476 -13.708797
H -13.9922764 28.0787705 -13.1669539
C -13.6973559 28.668285 -15.20173
O -13.2212429 29.4599915 -16.0166546
C -13.5976231 30.1744778 -13.1694646
H -13.3339687 30.2032292 -12.1112647
H -12.9526566 30.8857408 -13.6884859
C -15.0439728 30.6091374 -13.3172145
C -16.0007793 30.2075446 -12.3654889
H -15.7026398 29.6095012 -11.5164327
C -17.3497155 30.5734271 -12.5229045

H -18.084102 30.2600018 -11.7942258
C -17.7459906 31.3401123 -13.632773
H -18.784204 31.6156339 -13.7569758
C -16.7930155 31.742311 -14.5848431
H -17.0984245 32.326439 -15.4416514
C -15.4440982 31.3771052 -14.4276294
H -14.7165289 31.6734974 -15.1713742
N -14.6326318 27.780819 -15.5396291
H -14.9521194 27.1150618 -14.8471413
C -15.2390964 27.6886577 -16.8666391
H -15.735075 28.6312197 -17.0994544
H -14.4624243 27.5134405 -17.6111349
C -16.2666536 26.5596064 -16.957317
O -16.3953891 25.7524785 -16.0401148
N -16.9764725 26.4878918 -18.0819378
H -16.8289854 27.176356 -18.8044725
C -17.9575033 25.4408391 -18.3831007
H -17.7899232 24.5827215 -17.7304426
C -17.7792012 24.9927546 -19.8397808
O -17.7959146 25.8188285 -20.750926
C -19.3742105 25.9740925 -18.1133224
H -19.4072767 26.3626889 -17.0943744
H -19.5911057 26.7896238 -18.8053619
C -20.4418095 24.8784784 -18.2588574
H -20.2170614 24.0779836 -17.5490798
H -20.3816661 24.4589508 -19.2655484
C -21.873747 25.3850101 -18.0153021
O -22.0329252 26.4946208 -17.4568844
O -22.8044396 24.6367793 -18.3888393
N -17.5576422 23.6953514 -20.0649298
H -17.6282623 23.0479037 -19.293315
C -17.3269048 23.1256682 -21.396853
H -17.6295227 23.8524687 -22.1539443
C -18.173276 21.8705255 -21.6133325
O -18.4492201 21.1344351 -20.6665456
C -15.8303326 22.833808 -21.5971231
H -15.5031065 22.1074362 -20.8521036
H -15.2664698 23.755845 -21.4523805
C -15.5235885 22.3038595 -22.9769314
N -16.0573017 22.8083599 -24.1656877
C -15.6125519 22.0028993 -25.140073
H -15.854249 22.1181749 -26.1888167
N -14.8449621 21.02673 -24.6277346
H -14.429788 20.2755673 -25.1579691
C -14.7816493 21.1953241 -23.2621255
H -14.2791179 20.561661 -22.5447457

N -18.5185024 21.5818415 -22.8701808
H -18.1432063 22.1738025 -23.6009505
C -19.2288161 20.3661674 -23.257331
H -19.9116933 20.1067755 -22.4457962
C -18.2431347 19.2015468 -23.4437471
O -17.5034608 19.1367934 -24.4310812
C -20.0753688 20.6536823 -24.5101802
H -20.7622502 21.4721288 -24.2876809
H -19.4111275 20.9769363 -25.3134978
C -20.8899068 19.4416545 -25.0031907
H -20.2061397 18.632868 -25.252551
C -21.8900833 18.9305006 -23.9653095
H -22.5664824 19.7301243 -23.6638583
H -21.3716057 18.543429 -23.0898099
H -22.4721546 18.1090986 -24.387707
C -21.6558409 19.8178908 -26.2704486
H -22.3670983 20.6156827 -26.0549008
H -22.1966695 18.9460329 -26.6408658
H -20.9593117 20.149503 -27.0399621
N -18.305586 18.2264973 -22.5363312
H -18.9848484 18.3084584 -21.7879638
C -17.4987295 17.0022879 -22.56464
H -16.4445225 17.2767669 -22.5352472
C -17.7136686 16.1851165 -23.8479637
O -16.7324037 15.7061853 -24.4089253
C -17.8183622 16.1676056 -21.3118248
H -17.6392762 16.7679502 -20.4189096
H -17.1541192 15.3031549 -21.279748
C -19.2469521 15.6902349 -21.2993199
N -20.3555822 16.5291505 -21.1770985
C -21.409785 15.8002387 -21.5780685
H -22.404937 16.1941265 -21.7444035
N -21.0189659 14.5690479 -21.940586
H -21.5461555 13.999855 -22.611493
C -19.6585309 14.4792161 -21.7690455
H -19.0246639 13.6887216 -22.1457658
N -18.9342631 16.1416122 -24.3967885
H -19.7139306 16.4659898 -23.8383305
C -19.2306073 15.4622993 -25.6698397
H -18.9191364 14.4191048 -25.5943322
C -18.4740459 16.0915514 -26.8531044
O -17.994577 15.3816032 -27.732199
C -20.7422481 15.4827005 -25.9605596
H -20.9140981 15.0341016 -26.9402223
H -21.0845436 16.5175317 -26.0055241
C -21.5715667 14.7083567 -24.9337949

O -21.1731487 13.5796127 -24.57134
O -22.5721805 15.2767049 -24.4473625
N -18.2496141 17.4112082 -26.8323886
H -18.5716361 17.9329869 -26.0288096
C -17.4531297 18.1047217 -27.8489587
H -17.7977212 17.8135617 -28.8424323
H -17.5879439 19.1789559 -27.7304308
C -15.956921 17.7942072 -27.7454374
O -15.2888172 17.6280449 -28.7627028
N -15.439025 17.6201509 -26.5247796
H -16.0609943 17.7043393 -25.7294654
C -14.0743246 17.1266898 -26.3031017
H -13.3826248 17.7038566 -26.9186169
C -13.9256189 15.6534948 -26.724003
O -12.9564171 15.2990751 -27.3953963
C -13.702554 17.3422062 -24.8291262
H -13.7609572 18.4063683 -24.6038623
H -14.431762 16.8391893 -24.1953113
C -12.3186415 16.8515113 -24.4543145
C -11.189853 17.6482056 -24.7279804
H -11.3040053 18.609059 -25.2098121
C -9.9051194 17.1911998 -24.3777829
H -9.0312386 17.792005 -24.5808524
C -9.7460546 15.9368061 -23.7535044
O -8.5019852 15.5094553 -23.4137942
H -8.5099345 14.6569228 -22.9779248
C -10.8775819 15.1370117 -23.4808996
H -10.7593813 14.1747477 -23.0085044
C -12.161335 15.5959257 -23.8337612
H -13.0275541 14.9793858 -23.6353157
N -14.9268359 14.8201123 -26.423841
H -15.7014839 15.176822 -25.8741997
C -14.9828394 13.4197593 -26.8408227
H -14.0769896 12.9181517 -26.4986893
C -15.0310294 13.2711907 -28.3685764
O -14.316041 12.4302505 -28.9112801
C -16.1878629 12.7409781 -26.1724631
H -16.0987507 12.8425083 -25.0898935
H -17.1015123 13.2506904 -26.4752072
C -16.3194695 11.2716038 -26.5208461
C -15.679925 10.2960364 -25.7306851
H -15.1296517 10.5903453 -24.8487413
C -15.7427155 8.9385754 -26.1018896
H -15.2470437 8.1817352 -25.5143806
C -16.4327316 8.5582085 -27.2723703
O -16.4601685 7.2539103 -27.6505408

H -16.8895214 7.1591939 -28.5014005
C -17.087672 9.5346431 -28.0532165
H -17.603424 9.2540987 -28.9595907
C -17.035941 10.887789 -27.6716777
H -17.5222426 11.6367727 -28.2851494
N -15.7781968 14.1209507 -29.0785084
H -16.4213335 14.735133 -28.58639
C -15.8302485 14.0991621 -30.5458371
H -16.023302 13.0723458 -30.857743
C -14.4935853 14.5210283 -31.1785911
O -13.967663 13.81329 -32.0333803
C -16.994066 14.9750367 -31.0424011
H -17.90762 14.6865268 -30.519727
H -16.7819786 16.0183928 -30.8031416
C -17.238316 14.8508303 -32.5555673
H -16.3290961 15.1177248 -33.0992785
H -18.0038016 15.5779021 -32.8375267
C -17.6986865 13.4395358 -32.9540066
O -18.9264795 13.2539991 -33.097649
O -16.8225467 12.5548429 -33.0888592
N -13.837982 15.5729289 -30.6701081
H -14.2892367 16.1201696 -29.9440593
C -12.4898729 15.9648231 -31.1351682
H -12.5346458 16.1337827 -32.2113024
C -11.4694612 14.8360737 -30.9194148
O -10.6875397 14.5295424 -31.8238719
C -12.0510367 17.2817301 -30.4618222
H -12.2135606 17.2075124 -29.385962
C -10.5745282 17.6261432 -30.7001145
H -10.360449 17.6415137 -31.7693089
H -9.9326989 16.8899406 -30.2164525
H -10.3471552 18.6032195 -30.2737645
C -12.8768666 18.454303 -31.009272
H -12.6844 18.5843449 -32.0745506
H -12.6143381 19.3701809 -30.4802161
H -13.9403605 18.2682826 -30.8668087
N -11.5418689 14.1369177 -29.7810758
H -12.2068198 14.4377171 -29.0741423
C -10.7457172 12.934077 -29.5229422
H -9.6880299 13.1653715 -29.6509979
H -10.9128238 12.6231638 -28.4919327
C -11.1015855 11.7534031 -30.4368387
O -10.2052046 11.0525872 -30.9091058
N -12.3806552 11.5703557 -30.7665667
H -13.0669852 12.1981408 -30.3596135
C -12.8830576 10.4904933 -31.6335773

H -12.4536585 9.5465298 -31.299618
C -12.4546029 10.6930136 -33.0856704
O -11.8379689 9.7967565 -33.662114
C -14.4115139 10.3664975 -31.5108397
H -14.8837716 11.3195435 -31.7430037
C -15.0075931 9.281508 -32.405127
H -14.497612 8.3332624 -32.2385487
H -14.9184434 9.5764776 -33.4497101
H -16.0681325 9.1738969 -32.1753859
O -14.7191285 9.9990169 -30.1858573
H -14.6611214 10.8041515 -29.6445041
N -12.6061399 11.9050074 -33.6278135
H -13.1277568 12.5991368 -33.0943308
C -12.1108057 12.2899864 -34.96069
H -12.5626445 11.6330904 -35.7039491
C -10.5924574 12.1086371 -35.0606404
O -10.1118234 11.5017037 -36.0194828
C -12.5253388 13.7419652 -35.2846723
H -12.2047179 14.3950538 -34.4716908
C -11.9014053 14.2583629 -36.5896139
H -12.1485647 13.587198 -37.4129321
H -10.8182307 14.3273054 -36.4922998
H -12.2846337 15.2548982 -36.8119629
C -14.0473735 13.8569901 -35.4397281
H -14.390893 13.2481522 -36.2760117
H -14.3288191 14.8958381 -35.6094773
H -14.5551193 13.5168148 -34.5364626
N -9.8327613 12.5183483 -34.0376294
H -10.2691124 13.0158676 -33.2677802
C -8.3866883 12.2947826 -33.9964652
H -7.94803 12.713808 -34.9032965
C -8.0253687 10.7951613 -33.964585
O -7.1750255 10.3535353 -34.7385592
C -7.808095 13.0498299 -32.7940855
H -8.2322251 12.6640796 -31.8665257
H -8.0425363 14.1119177 -32.8768195
H -6.7249933 12.9258191 -32.7693519
N -8.7383063 9.9928143 -33.1690346
H -9.4399586 10.4216383 -32.5757625
C -8.5261026 8.5373236 -33.0519002
H -7.476906 8.3564513 -32.8190178
C -8.836892 7.7959867 -34.3557814
O -8.0721417 6.9214099 -34.7582541
C -9.3671494 7.960609 -31.9009351
H -10.4270834 8.1175921 -32.1029293
C -9.1210937 6.4714297 -31.6602582

H -8.0569718 6.2924769 -31.5037895
H -9.4663804 5.8883818 -32.5134126
H -9.6722286 6.1524028 -30.7752524
O -9.0287859 8.6085484 -30.6970051
H -9.4300372 9.4946216 -30.711502
N -9.8765834 8.2071453 -35.0894348
H -10.4751493 8.9342993 -34.7066042
C -10.218649 7.6484065 -36.4087586
H -10.3078933 6.565189 -36.3149444
C -9.0961985 7.9031619 -37.4291875
O -8.8107229 7.0319327 -38.2457573
C -11.5883493 8.2058925 -36.8724934
H -11.5800876 9.2895272 -36.7418409
C -11.8649231 7.9080697 -38.3605028
H -11.8182208 6.8336763 -38.5433383
H -11.1330206 8.4106419 -38.9928817
H -12.8465962 8.2794081 -38.6492608
C -12.7194695 7.6055139 -36.002287
H -12.8725628 6.5606242 -36.275951
H -12.4291906 7.6247014 -34.9527788
C -14.0547499 8.3529914 -36.1158443
H -14.469557 8.2558855 -37.1180749
H -13.9111854 9.4081356 -35.8800284
H -14.765007 7.9268607 -35.4065953
N -8.3937191 9.0388911 -37.3367375
H -8.6846259 9.7176446 -36.6427142
C -7.265239 9.3812153 -38.2229573
H -7.5533014 9.1179589 -39.2408252
C -5.9497281 8.6286735 -37.9283472
O -4.9741865 8.7465544 -38.6840646
C -7.0390962 10.899473 -38.1899562
H -8.0014138 11.4136505 -38.2093944
H -6.4800146 11.1924024 -39.0771632
O -6.2990547 11.3175345 -37.05805
H -6.6658562 10.914714 -36.2473093
N -5.8900761 7.8921141 -36.8100548
H -6.7249579 7.8243468 -36.2397125
C -4.6565738 7.3302429 -36.256341
H -3.8362244 8.0007177 -36.5123651
C -4.2800623 5.9408626 -36.8074929
O -3.1352983 5.5205919 -36.6491747
C -4.7709993 7.3240925 -34.7269169
H -5.5236596 6.6026775 -34.4107831
H -5.0474193 8.3167635 -34.3677705
H -3.8122191 7.0434425 -34.2890576
N -5.1800992 5.2659515 -37.5310566

H -6.0659794 5.7270073 -37.7010793
C -4.9863126 3.9392569 -38.1512939
H -5.0038412 3.1815968 -37.3671395
C -3.6467069 3.7571432 -38.8886477
O -2.9731921 2.7386014 -38.7264792
C -6.1697208 3.6740353 -39.1048049
H -7.0402153 3.4328757 -38.4935022
H -5.9499646 2.7970377 -39.7150907
C -6.5637775 4.8197352 -40.0201111
N -5.7699764 5.9207787 -40.3641479
C -6.5674691 6.752385 -41.049963
H -6.277414 7.7318812 -41.4028678
N -7.7926363 6.2244398 -41.1826337
H -8.5854946 6.6964794 -41.5940588
C -7.8129371 5.0107498 -40.5332027
H -8.6720512 4.3747877 -40.3736965
N -3.1845526 4.7824287 -39.6094995
H -3.8397485 5.5490973 -39.7402001
C -1.9094941 4.7790629 -40.3503428
H -1.9058746 3.9210138 -41.0259016
C -0.6755178 4.6000095 -39.4590681
O 0.3500762 4.1523355 -39.9634635
C -1.8108003 6.0655976 -41.1818839
H -1.9802055 6.9149351 -40.5182176
H -0.8101594 6.155225 -41.6088882
C -2.8366963 6.0737128 -42.3295833
H -3.7805566 5.6467642 -41.9998432
H -2.4675716 5.4691415 -43.158521
C -3.090666 7.5062857 -42.7975055
H -2.21557 7.8834898 -43.328123
H -3.2604295 8.1357064 -41.9227019
N -4.2898586 7.6146451 -43.6353146
H -5.0985667 8.0752973 -43.2163844
C -4.3632322 7.5927761 -44.9465972
N -3.4118636 7.0991039 -45.680707
H -3.2172373 7.5904414 -46.5573116
H -2.6560279 6.644648 -45.2205878
N -5.4028833 8.1065841 -45.5234574
H -6.0523492 8.6180683 -44.9202338
H -5.4423454 8.1830156 -46.5185052
N -0.7546542 4.8641941 -38.1528525
H -1.6478001 5.1514024 -37.7665882
C 0.3441051 4.6164829 -37.2169392
H 1.2323201 5.1264964 -37.5885598
C 0.6996833 3.1210416 -37.0906775
O 1.8637562 2.7984179 -36.8587498

C -0.0133202 5.2297426 -35.858726
H -0.9017486 4.748926 -35.4486878
H -0.2065543 6.2971976 -35.9704315
H 0.8169899 5.0909097 -35.1647239
N -0.2423564 2.2117153 -37.3765593
H -1.1831707 2.5200405 -37.5942453
C 0.0441412 0.7786215 -37.4595675
H 0.5673747 0.4724402 -36.5515695
C 0.9552045 0.4341165 -38.6550826
O 1.7906982 -0.4590169 -38.5465271
C -1.2872208 0.0221727 -37.522223
H -1.8167315 0.2672544 -38.4447155
H -1.9079066 0.2944651 -36.668599
H -1.1024213 -1.0504495 -37.4916568
N 0.8781965 1.1977947 -39.7544587
H 0.2102942 1.9555281 -39.7611379
C 1.8112123 1.0873253 -40.8905783
H 2.0039892 0.0301488 -41.0804667
C 3.1532747 1.7347408 -40.5479506
O 4.1918602 1.1211358 -40.7786454
C 1.2242149 1.7004207 -42.1796402
H 1.0712918 2.7691655 -42.0362597
C 2.1680245 1.5163423 -43.3761676
H 2.3959256 0.458749 -43.5151264
H 3.0968456 2.0616945 -43.2111974
H 1.7047064 1.9050683 -44.2829471
C -0.1270984 1.0684292 -42.5415657
H -0.0134202 -0.0094525 -42.6618157
H -0.4995357 1.4948188 -43.4727939
H -0.8607535 1.2658962 -41.7604837
N 3.1497115 2.9075864 -39.8983128
H 2.2584563 3.3646896 -39.7477886
C 4.3829764 3.5769137 -39.4271644
H 5.0316617 3.7532182 -40.287047
C 5.1714679 2.6870662 -38.4572727
O 6.3914124 2.6466122 -38.5342007
C 4.0854947 4.9447377 -38.7734971
H 3.4492565 4.7868126 -37.9038288
C 5.3579813 5.667574 -38.3077099
H 6.0473034 5.7700222 -39.1479344
H 5.8520541 5.0988091 -37.5198768
H 5.1091859 6.6515776 -37.9116408
C 3.3657123 5.9094878 -39.7292705
H 4.0142422 6.146524 -40.5715909
H 3.1117469 6.8290265 -39.2030367
H 2.4524687 5.4645623 -40.1099666

N 4.5099699 1.8691594 -37.6261136
H 3.501018 1.9661057 -37.5679173
C 5.1768327 0.8861925 -36.7517641
H 5.9189905 1.4157616 -36.1524491
C 5.9655105 -0.1805465 -37.5278367
O 6.9655235 -0.6580613 -37.005066
C 4.1151826 0.2654933 -35.821984
H 3.6872668 1.0584642 -35.2070059
H 3.3115988 -0.1409395 -36.4366679
C 4.6026419 -0.8663175 -34.8987054
H 4.9701494 -1.6946528 -35.5041773
H 3.7425868 -1.2253198 -34.3369632
C 5.691348 -0.4487287 -33.8979481
H 5.305648 0.3329929 -33.2411631
H 6.5347315 -0.0299299 -34.4515579
N 6.1852786 -1.5988657 -33.1065998
H 7.0302672 -2.023504 -33.4565587
C 5.6931212 -2.0929878 -31.9796119
N 4.6137617 -1.6238221 -31.4121932
H 4.2496792 -2.0315972 -30.5644291
H 4.1500759 -0.8388203 -31.8268032
N 6.2826626 -3.0960025 -31.392016
H 7.158683 -3.4460252 -31.7437805
H 5.9220268 -3.4384356 -30.5210906
N 5.5130667 -0.5786026 -38.7196586
H 4.6989518 -0.1098716 -39.0946508
C 6.155844 -1.6215128 -39.5477068
H 6.7171868 -2.2989085 -38.9044692
C 7.1870424 -1.0471198 -40.5141252
O 8.1672071 -1.7115576 -40.8379074
C 5.0729423 -2.429844 -40.2970744
H 4.4719049 -1.7392024 -40.8919679
C 5.6505611 -3.4778044 -41.2668926
H 6.3652394 -4.1194898 -40.7503349
H 6.1577817 -2.9777053 -42.0922651
H 4.8537515 -4.0898952 -41.6888069
C 4.1272498 -3.1381766 -39.3001663
H 3.4521743 -2.3981917 -38.8745679
H 3.519856 -3.8567523 -39.8472324
C 4.8022508 -3.8790266 -38.1377464
H 5.1620025 -3.1674568 -37.3956499
H 5.6444629 -4.4611102 -38.5038849
H 4.0797422 -4.5405924 -37.6627072
N 6.9634562 0.1827065 -40.9660073
H 6.1059518 0.645179 -40.6903566
C 7.8918005 0.9019409 -41.8307362

H 8.3560989 0.1968688 -42.5199967
C 9.0541694 1.5083209 -41.0392823
O 10.1751688 1.4175308 -41.5113396
C 7.0910835 1.9372281 -42.6425295
H 6.4755174 2.5141194 -41.9494156
C 8.015055 2.9405889 -43.3353853
H 8.8170607 2.4130698 -43.8517664
H 8.4436482 3.6079035 -42.5859543
H 7.4537262 3.5413332 -44.0376991
C 6.1329575 1.2547909 -43.6511673
H 5.3388 0.7519175 -43.1009127
H 5.6559528 2.0240523 -44.2518133
C 6.7607953 0.2206656 -44.5975052
H 7.0017951 -0.6915304 -44.0477814
H 7.6692296 0.6123557 -45.0487245
H 6.0481605 -0.0291608 -45.3837918
N 8.798992 1.9819869 -39.8195457
H 7.8269675 1.9509433 -39.5448266
C 9.6149082 2.8267209 -38.9312898
H 9.2170332 2.5797365 -37.9459199
C 9.3250434 4.3409519 -39.0306445
O 8.5238253 4.7993794 -39.8542185
C 11.099113 2.4020543 -38.8279642
H 11.7082873 3.1408157 -39.3434734
H 11.2510327 1.4269978 -39.2904088
C 11.5965131 2.3038259 -37.3833269
O 11.4492724 3.3205405 -36.6703128
O 12.1249463 1.2330349 -37.0094698
N 9.8588958 5.1058401 -38.0754761
H 10.516748 4.6466534 -37.4396315
C 9.3325992 6.4041998 -37.6512809
H 8.3526531 6.2042132 -37.2199303
C 9.1395831 7.4234129 -38.7896545
O 10.0494829 7.7492656 -39.5515223
C 10.2024572 6.9760756 -36.5200309
H 10.3373002 6.2059069 -35.7593743
H 11.1814432 7.2466722 -36.9189787
C 9.5601829 8.2096529 -35.8657287
H 8.5933201 7.9247972 -35.443284
H 9.3792816 8.9717041 -36.6261681
C 10.4478246 8.8008571 -34.7647909
O 10.8020372 8.0456266 -33.8314376
O 10.7428137 10.012847 -34.8410345
N 7.9356494 8.0039627 -38.8202964
H 7.2747294 7.6949909 -38.1276181
C 7.4979327 9.066309 -39.7246913

H 6.4475177 9.2219085 -39.4784392
C 7.4607978 8.7250306 -41.2328113
O 7.2182063 9.6195186 -42.0501456
C 8.189382 10.3821855 -39.339387
H 9.2533861 10.3291555 -39.5703739
H 8.0772996 10.5585665 -38.2684137
H 7.7299772 11.2135753 -39.8729389
N 7.5456185 7.4423073 -41.6155564
H 7.7541 6.7354562 -40.9135248
C 7.344726 6.9801426 -42.997353
H 6.9802142 7.8289962 -43.5714683
C 6.2228111 5.9398072 -43.1423331
O 5.8136015 5.298277 -42.1811172
C 8.7040589 6.5417867 -43.5937623
H 9.3769214 6.2789443 -42.781539
H 8.5857406 5.6690598 -44.2357264
C 9.2911549 7.7100855 -44.425368
H 9.1907816 8.6410518 -43.8675988
C 10.7676023 7.5379856 -44.7641775
H 10.9094523 6.6899523 -45.4326336
H 11.347273 7.4053009 -43.8535032
H 11.1408564 8.4293545 -45.2677901
C 8.5546316 7.8484516 -45.7653518
H 8.6226347 6.9085588 -46.3141622
H 9.0244539 8.6267692 -46.3653165
H 7.5122898 8.121164 -45.6183874
N 5.6747873 5.8134553 -44.3622881
H 6.051826 6.3868802 -45.1032308
C 4.5891328 4.8649351 -44.6980872
H 4.5631062 4.1139725 -43.9051263
C 4.8447935 4.0459821 -45.973334
O 3.9017478 3.6498304 -46.6520897
C 3.2105171 5.5603428 -44.666108
H 2.4533252 4.7738203 -44.6831793
H 3.1054279 6.0864467 -43.7153432
C 2.9161759 6.5454592 -45.8179066
H 3.1261552 6.0697121 -46.7735315
H 3.5448542 7.4315362 -45.7241825
C 1.4308664 6.9484155 -45.7776237
H 1.2590212 7.5693356 -44.8966521
H 0.8212512 6.0482975 -45.6740764
N 0.981203 7.6811794 -46.9780385
H 0.7524881 8.6673509 -46.8923419
C 0.6228584 7.177358 -48.1443321
N 0.8186153 5.929421 -48.4685675
H 0.5752193 5.6132175 -49.3870961

H 1.388143 5.3577715 -47.8677134
N 0.0406873 7.949872 -49.0092621
H -0.2933387 8.855375 -48.6610897
H -0.2623347 7.6163466 -49.9005672
N 6.1190758 3.8578277 -46.3209588
H 6.8301691 4.1363819 -45.6635574
C 6.5958218 3.2428633 -47.5626859
H 5.830005 2.5719029 -47.9542416
C 7.8916732 2.4367347 -47.343168
O 8.5668267 2.64766 -46.3401455
C 6.8674228 4.3491416 -48.5851797
H 7.7970464 4.8464069 -48.304794
H 7.032437 3.8770175 -49.5533758
C 5.793543 5.4093771 -48.7466088
C 4.5694425 5.071977 -49.3430479
H 4.3617759 4.0434651 -49.5976387
C 3.6563468 6.0786253 -49.6922932
H 2.7837729 5.8310304 -50.2772817
C 3.9157208 7.4151039 -49.3467661
H 3.2349086 8.196869 -49.6514519
C 5.0976506 7.7468283 -48.6644446
H 5.3156042 8.7854837 -48.4512446
C 6.0470079 6.7471097 -48.3865945
H 6.9997683 7.0245668 -47.966154
N 8.2414517 1.5676536 -48.3001291
H 7.6597218 1.5224626 -49.118932
C 9.3979198 0.6578697 -48.2914649
H 9.1016447 -0.2646165 -47.791358
C 10.6602547 1.1969945 -47.5742744
O 11.1786285 2.245083 -47.9427162
C 9.7087369 0.3224341 -49.75895
H 9.967396 1.234773 -50.3010873
H 8.8483582 -0.1504811 -50.2328802
H 10.5608699 -0.3531544 -49.8204047
N 11.1058984 0.4683517 -46.5404187
H 10.6020477 -0.3756197 -46.3181145
C 12.3498292 0.6411544 -45.7620579
H 12.0842348 0.2321049 -44.78769
C 12.7677959 2.0582833 -45.3295639
O 13.9257997 2.2966503 -44.965395
C 13.488643 -0.2542563 -46.2966031
H 14.4106292 0.3125754 -46.3885813
H 13.233996 -0.5667301 -47.3062372
C 13.7404825 -1.5012609 -45.4212878
O 12.782663 -1.9446545 -44.7479203
O 14.8558588 -2.0616043 -45.4835954

N 11.7676098 2.9221305 -45.1175289
H 10.8663841 2.6705479 -45.5032949
C 11.8434355 4.1137479 -44.2565654
H 10.9702733 4.7131008 -44.4953178
C 13.0474841 5.0013958 -44.5690478
O 13.2566867 5.4146228 -45.7033659
C 11.6839069 3.7215239 -42.7654516
H 10.7042933 3.2772 -42.7433744
C 12.7172875 2.6995277 -42.243958
H 13.7411237 3.021697 -42.4103113
H 12.5627012 1.7344082 -42.7242798
H 12.5786212 2.5252549 -41.1832717
C 11.6078686 4.8907969 -41.7666777
H 12.5088229 4.963613 -41.1569353
H 10.7748965 4.7152793 -41.0967139
H 11.4403098 5.837781 -42.2673589
N 13.7997814 5.3551081 -43.5357305
H 13.5427582 4.9757027 -42.6392564
C 14.9722896 6.1999847 -43.5773004
H 14.7513394 7.1316738 -44.1038787
C 16.1325397 5.5277022 -44.3124657
O 16.971215 6.2121722 -44.8791594
C 15.3363167 6.5059358 -42.1167684
H 14.5993989 7.1982355 -41.7073855
H 15.2826306 5.5845827 -41.5328418
C 16.7149966 7.0988042 -41.9408267
C 16.9780411 8.3984896 -42.403626
H 16.1908624 8.9809401 -42.8654804
C 17.7536204 6.3218351 -41.3924162
H 17.5600729 5.3123153 -41.0611771
C 18.2759221 8.9240865 -42.299878
H 18.4802673 9.9218835 -42.6403416
C 19.0546992 6.8452967 -41.3078177
H 19.8555147 6.2433117 -40.90308
C 19.314826 8.149128 -41.7601691
H 20.3159865 8.5514257 -41.6963891
N 16.2058513 4.196207 -44.3268838
H 15.3924863 3.6449569 -44.0809636
C 17.3365519 3.5317345 -44.9880782
H 18.247824 4.0148928 -44.6328091
C 17.3362498 3.7162578 -46.5202865
O 18.411362 3.6889561 -47.1212283
C 17.4139431 2.0551039 -44.544089
H 17.2640812 2.0186564 -43.4640312
C 18.7685605 1.4097662 -44.8475106
H 18.7772956 0.3945679 -44.4540022

H 19.5699411 1.9849547 -44.3903993
H 18.9200973 1.3641078 -45.9264033
O 16.4573901 1.2286298 -45.1599023
H 15.5577683 1.5969757 -45.0914068
N 16.1818698 4.0690221 -47.1078517
H 15.3465728 4.1192239 -46.5422669
C 16.0387294 4.3659216 -48.539581
H 16.9320402 3.9392428 -48.9945931
C 16.0907108 5.8966401 -49.0803814
O 16.0794458 5.9767522 -50.2908172
C 14.8556147 3.4957892 -49.0773819
H 13.9798023 4.1366509 -49.1933825
H 14.5968292 2.7346661 -48.3378472
C 15.1275989 2.7525254 -50.4024108
O 14.1400315 2.2286328 -50.965854
O 16.307982 2.6299838 -50.8096799
N 16.2900343 7.1238414 -48.4203981
H 16.4497442 7.0131143 -47.4277553
C 16.1613041 8.688626 -48.8951202
H 15.5643498 8.6270493 -49.8051702
C 17.4543756 9.8556596 -49.3389581
O 18.4787587 9.2741032 -49.0634247
C 15.1839224 9.2297703 -47.7945255
H 14.6070166 8.4267525 -47.3302822
H 14.4664933 9.8869038 -48.2885934
C 15.9020718 10.0680588 -46.713811
O 16.3872989 11.1886415 -47.009515
O 15.9809798 9.6471257 -45.537687
N 17.6582507 11.2710311 -50.0187799
H 16.7432841 11.6473397 -50.215925
C 18.9196021 12.4999217 -50.4996121
H 19.4239745 12.7072473 -49.557024
C 19.8910586 11.6284874 -51.3214552
H 19.3040143 10.926469 -51.916006
H 20.4598406 12.2416738 -52.0211191
C 20.8996192 10.8577388 -50.4511839
H 20.4908512 10.6758339 -49.4595413
H 21.799341 11.4600084 -50.3269145
C 21.2793501 9.5187183 -51.0747255
O 22.2065281 9.4215954 -51.8603748
N 20.5592735 8.459898 -50.7803827
H 20.794535 7.5910496 -51.2218482
H 19.7724068 8.5434667 -50.1355936
C 19.092366 14.182846 -51.3913657
O 18.0971829 14.350783 -52.0856275
N 20.1085432 15.3856543 -51.5493382

H 20.9260687 15.212857 -50.986138
C 20.2948739 16.9461427 -52.3131094
H 20.2227503 16.7196393 -53.3770916
C 21.4078328 18.3179152 -52.2406162
O 22.5468109 18.183726 -51.7988277
C 19.0686253 17.7464106 -51.9697146
H 19.2338989 18.2728335 -51.0391949
H 18.2005998 17.0932304 -51.9204808
H 18.8667864 18.4802587 -52.7438545
N 21.1702238 19.6749261 -52.6514902
H 20.2345851 19.8379585 -52.9935248
C 22.0024176 21.0115075 -52.4479166
H 22.8683875 20.7431257 -51.8441611
C 21.2832516 22.1579803 -51.6588977
O 20.098186 22.3880363 -51.8856444
C 22.5113401 21.7661131 -53.7349204
H 21.6390627 22.0773934 -54.3111731
H 23.0283835 22.6774737 -53.4283394
C 23.4507222 21.0203275 -54.6889031
H 22.9115143 20.1707727 -55.1106251
H 23.7236569 21.6915994 -55.5045992
C 24.7266258 20.5101083 -54.0066815
H 24.4530278 19.8220364 -53.2042071
H 25.2791937 21.3544152 -53.5879837
C 25.5984771 19.7681775 -55.0268131
H 25.909443 20.4728439 -55.8051529
H 24.9888066 18.9920389 -55.5006992
N 26.7854235 19.1515691 -54.3842775
H 27.3653918 19.8580064 -53.9512649
H 27.3408378 18.6543916 -55.0695705
H 26.4940335 18.4923601 -53.6723818
N 22.0141776 22.9790716 -50.8776924
H 22.9848334 22.7561847 -50.7481297
C 21.4363151 24.0595014 -50.0406279
H 20.6651534 23.5880877 -49.4217957
C 20.6932301 25.1571828 -50.8434591
O 21.1123859 25.4928957 -51.9501055
C 22.4954675 24.6021902 -49.0576117
H 21.9681595 25.0697176 -48.2281703
H 23.0446883 23.7611733 -48.6298199
C 23.4967813 25.6136439 -49.6395768
H 24.1308012 25.1185841 -50.3761876
H 22.9616084 26.426131 -50.1294037
C 24.3600993 26.2058559 -48.5122502
H 25.0695733 25.4548231 -48.159759
H 23.7129745 26.4715583 -47.6719088

C 25.1007745 27.47991 -48.9414417
H 25.6487684 27.8643149 -48.0751497
H 24.3538345 28.231996 -49.2207631
N 26.033051 27.2429212 -50.0717632
H 26.729265 26.5557402 -49.8134378
H 26.5058074 28.1019389 -50.3270792
H 25.5216444 26.9054137 -50.8778729
N 19.6653497 25.7902966 -50.2502701
C 18.5260935 26.4199977 -50.9302971
H 17.683172 26.2611953 -50.25573
C 17.9666354 25.9230463 -52.2825018
O 17.2120042 26.6858309 -52.8855268
C 18.8245867 27.9163968 -50.8994079
H 19.6064095 28.1600803 -51.6210341
H 17.9311673 28.5198536 -51.0658878
C 19.9982801 26.7186724 -49.1631419
H 19.6497361 26.3469128 -48.204326
H 21.0706298 26.8855675 -49.103357
C 19.3411503 28.0678737 -49.4726293
H 18.5032263 28.2293829 -48.7937525
H 20.0551531 28.8885143 -49.395854
N 18.1444302 24.6585658 -52.7170806
H 18.7575415 24.0313991 -52.2081714
C 17.2497154 24.0944877 -53.7603078
H 17.1809009 24.8305569 -54.5644565
C 15.8108259 23.9536811 -53.2184155
O 15.6116559 23.7733122 -52.0168994
C 17.7905208 22.7718561 -54.367871
H 18.2164828 22.1394345 -53.5915213
H 16.9504542 22.2274091 -54.799801
C 18.8275157 22.9966047 -55.490398
H 19.7490875 23.3914824 -55.0618205
H 18.4292468 23.7618966 -56.1540239
C 19.1471149 21.7322105 -56.331149
H 18.3107319 21.0347568 -56.2604309
H 20.0375569 21.2460875 -55.9308658
N 19.3493069 22.0762304 -57.7577649
H 19.2235844 23.0639289 -58.0004847
C 19.3902273 21.3013719 -58.8329725
N 19.5688706 20.0105413 -58.789513
H 19.5868354 19.4947518 -59.6476506
H 19.6328429 19.5190873 -57.9186205
N 19.2411043 21.8219077 -60.01574
H 18.7989704 22.7503794 -60.07748
H 19.2658129 21.2691169 -60.8598066
N 14.821947 23.9635607 -54.1171727

H 15.0640462 24.1295175 -55.0795782
C 13.4070457 23.7053603 -53.8023649
H 13.2318405 23.8314889 -52.7303655
C 13.0148305 22.2690653 -54.1901151
O 13.2650034 21.8554559 -55.3268438
C 12.5042714 24.70102 -54.554355
H 11.4657565 24.4018961 -54.4176821
H 12.7363243 24.6603459 -55.6166903
C 12.5918951 26.14774 -54.1111584
O 13.1869008 26.5106661 -53.1113018
N 11.9857542 27.0281791 -54.8750885
H 12.0663734 27.990201 -54.6088372
H 11.474667 26.7304553 -55.7012412
N 12.3104354 21.5618859 -53.3039725
H 12.1319232 21.9882772 -52.3959269
C 11.7798987 20.2088839 -53.5218785
H 12.1354009 19.8236533 -54.4750295
C 10.2499869 20.2006078 -53.5304971
O 9.636071 20.6928764 -52.5837385
C 12.2939867 19.2814334 -52.4104324
H 12.0003572 19.7111831 -51.4550013
H 13.3811827 19.2399207 -52.459261
C 11.7331357 17.8531895 -52.451972
H 10.6473428 17.8812916 -52.5280063
H 11.9562163 17.3702826 -51.5054196
S 12.3827862 16.813984 -53.7779941
C 13.8931229 16.2105975 -52.9728887
H 14.4627188 15.602152 -53.6752431
H 13.6285852 15.6018071 -52.1075691
H 14.5036321 17.0516099 -52.6454799
N 9.6561252 19.476022 -54.4810838
H 10.2356146 19.1014098 -55.228018
C 8.2618632 19.0294284 -54.4110718
H 7.7432147 19.5950109 -53.6411374
C 8.2012683 17.5367186 -54.0391612
O 8.6091906 16.6916482 -54.8315145
C 7.5848191 19.3377223 -55.7563497
H 7.6533027 20.404727 -55.9472126
H 8.1407942 18.8235216 -56.5354205
C 6.1094319 18.9161937 -55.8471269
H 6.0223168 17.8621598 -55.5987058
C 5.2008171 19.7105303 -54.9111302
H 5.2842959 20.7773402 -55.1160026
H 5.4789999 19.5191252 -53.8783082
H 4.1706631 19.3882301 -55.0433692
C 5.5958432 19.1157678 -57.2735893

H 5.6369121 20.1715894 -57.5396105
H 4.5696247 18.7588489 -57.3505647
H 6.2159663 18.5490014 -57.9673902
N 7.650806 17.207511 -52.8705797
H 7.2809473 17.9542867 -52.2913677
C 7.3710433 15.8279812 -52.4464654
H 8.0060151 15.1508264 -53.0126345
C 5.9027388 15.4793458 -52.7611271
O 4.9709185 16.075372 -52.2065813
C 7.75986 15.6405981 -50.9667627
H 7.1203425 16.2573012 -50.3370099
H 8.7862314 15.9905422 -50.8370111
C 7.6824443 14.1863395 -50.4760692
O 6.8708591 13.4090086 -51.019936
O 8.4087115 13.8653462 -49.5055914
N 5.6864177 14.5711102 -53.7189983
H 6.4974395 14.0713565 -54.0777662
C 4.3447916 14.1842873 -54.1843736
H 3.6345714 14.9349283 -53.8406842
C 3.9351687 12.8544228 -53.5605881
O 4.3232699 11.7831128 -54.0192536
C 4.2336867 14.1575708 -55.7201635
H 4.8568936 13.3737071 -56.1278179
C 2.7974381 13.8629456 -56.1479142
H 2.1426664 14.6095088 -55.703318
H 2.4969621 12.8683791 -55.8186253
H 2.7229068 13.8922377 -57.2343704
C 4.6461988 15.4961607 -56.3443168
H 4.0765308 16.3124174 -55.897042
H 4.4772019 15.482627 -57.4198195
H 5.7094427 15.6583488 -56.1699027
N 3.0257182 12.9205146 -52.5909151
H 2.717312 13.8404901 -52.2952782
C 2.6511322 11.7941466 -51.7378099
H 1.5919382 11.8736081 -51.495824
H 2.826334 10.8515972 -52.2505211
C 3.4378837 11.764253 -50.4292475
O 3.7783718 10.6859284 -49.9559292
N 3.6725557 12.9262538 -49.8125931
H 3.3218807 13.7634739 -50.2589932
C 4.5639542 13.1282033 -48.6574106
H 5.5763021 12.8435635 -48.9560907
C 4.2394702 12.317265 -47.379758
O 4.9944892 12.3378076 -46.4012804
C 4.557135 14.6320349 -48.3562011
H 4.9178299 15.1702423 -49.2356233

H 5.2327209 14.8285762 -47.5267004
S 2.8894722 15.2074225 -47.9074698
H 2.7061254 14.4032963 -46.8479299
N 3.0995178 11.6227276 -47.3420058
H 2.5537956 11.6005599 -48.1895828
C 2.6973692 10.7695608 -46.2284903
H 1.7296098 10.3306215 -46.4382888
H 3.4252333 9.964224 -46.1301078
C 2.5964225 11.5335775 -44.9096616
O 2.0756425 12.6458024 -44.8617237
N 3.1475683 10.9405853 -43.8409104
H 3.5995258 10.050913 -43.9813798
C 3.2644505 11.5676352 -42.5110625
H 2.5363332 12.3721322 -42.4922009
C 4.6024855 12.3215117 -42.2989558
O 4.9330438 12.7035656 -41.1837733
C 2.7685976 10.6219858 -41.3876643
H 3.329892 9.6866489 -41.3966698
H 2.9110664 11.1133101 -40.4234859
C 1.2482459 10.3497499 -41.5881163
H 0.7919812 11.3134804 -41.7977461
H 1.1127473 9.7337692 -42.4784474
C 0.4227669 9.7161131 -40.4391496
H 0.7314226 10.1512193 -39.4901884
H 0.5960085 8.639396 -40.4025918
C -1.0758004 10.0217816 -40.7095582
H -1.41602 9.4220592 -41.5561837
H -1.1388727 11.0716698 -41.0127608
N -2.0107233 9.8625097 -39.5598994
H -2.0996365 8.9127489 -39.2334567
H -2.9406933 10.1949484 -39.8135329
H -1.7309402 10.4529809 -38.7839952
N 5.3030928 12.6456133 -43.3932679
H 4.9509657 12.3158047 -44.2839729
C 6.263606 13.7539353 -43.4685514
H 6.339384 14.0373157 -44.5168193
H 5.8664254 14.6025745 -42.9108326
C 7.6966603 13.5217271 -42.9844836
O 8.5229441 14.4189115 -43.1446022
N 8.052298 12.3272621 -42.5032093
H 7.3432049 11.6119669 -42.3818921
C 9.4128024 12.0200965 -42.040962
H 9.6646061 12.6654394 -41.1981862
H 9.4474178 10.9834761 -41.7123813
C 10.481707 12.1834795 -43.1192423
O 11.5222019 12.7773601 -42.8556136

N 10.1750316 11.7843984 -44.3589289
H 9.2846575 11.3332788 -44.4955374
C 11.0186339 12.0413754 -45.5425943
H 11.9980506 11.588156 -45.3790969
C 11.2701527 13.5389522 -45.7486745
O 12.4236188 13.9518113 -45.8260238
C 10.4258214 11.3832015 -46.8083101
H 10.5905208 10.3099959 -46.7269139
C 8.9312833 11.6033541 -47.0463739
H 8.7099762 12.6632397 -47.1442538
H 8.3393666 11.1827496 -46.2344072
H 8.6361123 11.1131843 -47.9758533
O 11.0895781 11.8306891 -47.9597574
H 10.484578 12.3844576 -48.4701324
N 10.2252136 14.371186 -45.663101
H 9.3074535 13.9722939 -45.5446904
C 10.2857366 15.8236971 -45.8547703
H 10.7235008 16.0265557 -46.8268567
C 11.1106349 16.537419 -44.7650356
O 11.8863183 17.4526382 -45.0535196
C 8.8657689 16.3970441 -45.8856242
H 8.8986116 17.4030429 -46.3069571
H 8.4867539 16.4648006 -44.8668808
S 7.7163401 15.3759861 -46.8510702
H 8.4533831 15.1086649 -47.9512639
N 10.9632892 16.0999518 -43.508704
H 10.2825801 15.3646959 -43.3383485
C 11.7666912 16.5933009 -42.3777083
H 11.7210304 17.682167 -42.3558897
C 13.2337723 16.2149077 -42.5646649
O 14.1041332 17.0794444 -42.4631432
C 11.2181531 16.0596538 -41.0385742
H 11.0788588 14.9803609 -41.1088303
C 12.1583111 16.3454275 -39.8608403
H 12.413162 17.4044263 -39.8365978
H 13.0713173 15.7583224 -39.9544092
H 11.6747709 16.0658247 -38.9240883
C 9.8619149 16.7101419 -40.7311641
H 9.9738266 17.7906836 -40.6408162
H 9.465485 16.313601 -39.7953368
H 9.147258 16.4904463 -41.5236473
N 13.5086787 14.9554922 -42.9189931
H 12.7374487 14.300883 -43.0121126
C 14.8616324 14.4600437 -43.1763215
H 15.4605874 14.5795315 -42.2714439
C 15.5717593 15.257738 -44.2626702

O 16.6984701 15.6980688 -44.0520232
C 14.8119003 12.9696725 -43.5454865
H 15.47017 12.778969 -44.3885111
H 13.8159203 12.6800994 -43.8734669
C 15.2350372 12.1081566 -42.3887418
N 16.5503927 11.9810507 -41.9354251
C 16.4753185 11.2196844 -40.8299497
H 17.3253826 10.9192111 -40.2306378
N 15.2029599 10.8627996 -40.5836198
H 14.890693 10.2906702 -39.8085624
C 14.4037633 11.4233475 -41.5538219
H 13.3257702 11.3738953 -41.6274226
N 14.9078435 15.5075468 -45.387301
H 13.989292 15.091668 -45.5152898
C 15.5112271 16.2350314 -46.5011762
H 16.4944499 15.8005517 -46.6810838
C 15.788866 17.702243 -46.1455768
O 16.9395239 18.1374501 -46.2012692
C 14.6897345 16.0179734 -47.7882799
H 15.1327128 16.6236197 -48.5603665
C 14.7908711 14.5234165 -48.182398
H 14.1962195 13.8987777 -47.5198436
H 15.8220997 14.1781548 -48.1262874
H 14.4372692 14.3716629 -49.2011021
C 13.2366266 16.4975189 -47.7320752
H 12.7646062 16.0195718 -46.8976035
H 13.2059957 17.5711979 -47.5585925
C 12.4407379 16.1837724 -49.0033466
H 12.2244151 15.1164961 -49.062049
H 13.0189562 16.48687 -49.8741816
H 11.4970485 16.7230964 -48.9888032
N 14.8187234 18.4245126 -45.5793613
H 13.9008259 18.0120409 -45.4473034
C 15.0314858 19.8172963 -45.1857533
H 15.4248899 20.3646793 -46.0422485
C 16.0654803 19.9815299 -44.052507
O 16.775709 20.9872676 -44.0197683
C 13.6739593 20.4215669 -44.8288207
H 13.2487701 19.8884107 -43.9775321
H 12.9997992 20.3346726 -45.682005
H 13.7950387 21.4777533 -44.588073
N 16.1945128 19.0031707 -43.1417661
H 15.5550477 18.2125874 -43.1960906
C 17.1426889 19.0652184 -42.0165262
H 17.2682905 20.115006 -41.7540704
C 18.5555244 18.5806105 -42.3633946

O 19.5022617 19.2022987 -41.8920977
C 16.5356863 18.3408313 -40.8014493
H 15.5546198 18.7709715 -40.5929376
H 16.3868088 17.2899072 -41.0560764
C 17.3783728 18.4041283 -39.5137768
H 18.2761985 17.8007418 -39.655826
H 16.803368 17.9420041 -38.7110996
C 17.8159365 19.8120997 -39.0658109
H 18.4907804 20.2290409 -39.816017
H 18.3932875 19.7217441 -38.1440225
N 16.6897875 20.7491832 -38.8609668
H 16.5665419 21.4502419 -39.5716769
C 15.8807065 20.8193544 -37.8169131
N 14.9853566 21.7589121 -37.7435133
H 14.3645694 21.8061651 -36.956886
H 14.8726153 22.4170928 -38.4946404
N 15.9472023 19.9843426 -36.8189475
H 16.6241288 19.2455779 -36.8502505
H 15.2949912 20.0477172 -36.0589826
N 18.7300829 17.5290883 -43.1764286
H 17.9068943 17.0488226 -43.5306661
C 20.0684893 17.0390531 -43.5746573
H 20.7065707 16.9291063 -42.6964889
C 20.7865705 18.0002679 -44.5366372
O 22.0134416 18.0053269 -44.5586972
C 19.9813346 15.699411 -44.3219547
H 19.304658 15.8747004 -45.1616404
H 20.9671976 15.5209637 -44.7569981
C 19.5628823 14.3578569 -43.6784448
H 20.3290906 14.0366348 -42.971993
H 18.6129878 14.4437749 -43.1562454
C 19.4543242 13.3380914 -44.844005
H 20.418487 13.3271226 -45.354375
H 18.7167883 13.694725 -45.5669937
C 19.1362141 11.8730891 -44.4980398
H 19.490443 11.6480878 -43.4892399
H 19.686472 11.2395421 -45.2001729
N 17.6955314 11.5486037 -44.6343123
H 17.1272324 12.0169392 -43.9401613
H 17.4806865 10.5531329 -44.5600195
H 17.3202713 11.7308229 -45.5705452
N 20.0478271 18.6624719 -45.4364741
H 19.046923 18.5093921 -45.436498
C 20.6358853 19.3309775 -46.6085273
H 21.6749389 19.5607893 -46.3684505
C 20.0541746 20.7091617 -46.9622581

O 20.3152965 21.2210728 -48.0515675
C 20.6868561 18.354292 -47.7957786
H 21.0473326 18.879063 -48.6814314
H 21.4566581 17.6177888 -47.5613333
C 19.4373287 17.602644 -48.1667651
C 19.394892 16.1940631 -48.1271936
H 20.196105 15.6434021 -47.6560693
C 18.3451897 15.5113149 -48.7728262
H 18.2923324 14.4372359 -48.7484014
C 17.3896775 16.2497781 -49.5044677
O 16.5031357 15.6745726 -50.3589183
H 16.8175994 14.8393012 -50.723428
C 17.3695605 17.6457992 -49.4114359
H 16.6398896 18.1806343 -49.9827419
C 18.3848271 18.32481 -48.7295873
H 18.4231516 19.3990316 -48.7319403
N 19.3517895 21.3539171 -46.024161
H 19.1651048 20.8775033 -45.1539639
C 19.0384741 22.7911583 -46.0750765
H 18.3539854 22.9771147 -45.2457682
C 18.2660689 23.2374827 -47.3416898
O 18.3498463 24.3844502 -47.7803859
C 20.3254224 23.5929801 -45.7682459
H 21.0190659 23.4837652 -46.6030665
H 20.8007875 23.1682607 -44.8823023
C 20.0831759 25.0814136 -45.500918
O 19.0655738 25.4065578 -44.8420731
O 20.8655128 25.9146396 -46.0124582
N 17.4548563 22.3556176 -47.9295338
H 17.4238769 21.424189 -47.5465261
C 16.4823397 22.7281121 -48.9710723
H 16.9464183 23.4477338 -49.6473765
C 15.24281 23.3977265 -48.3725054
O 14.9522023 23.2457856 -47.1836795
C 16.0422905 21.5162724 -49.8130752
H 15.2603537 21.8499514 -50.4920367
C 17.2011516 21.0558416 -50.7125561
H 18.1143336 20.9147098 -50.1404907
H 17.4026366 21.8357731 -51.4439333
H 16.943333 20.1436236 -51.2459316
C 15.398919 20.4182088 -48.9405732
H 16.1665696 19.8238888 -48.4571155
H 14.7955406 20.8598812 -48.151009
C 14.4493492 19.5350595 -49.7478417
H 14.8862927 19.2531794 -50.7036183
H 13.5268763 20.0795328 -49.9194258

H 14.2201392 18.6408149 -49.1834476
N 14.4360211 24.010578 -49.2375354
H 14.7402083 24.0446147 -50.2085445
C 12.9993631 24.158699 -49.0076405
H 12.8175786 24.2573771 -47.9397338
C 12.2662528 22.9046082 -49.5103378
O 12.6107608 22.3634176 -50.5604784
C 12.5234879 25.4517901 -49.6859483
H 13.0406288 26.283309 -49.2048236
H 12.830272 25.44303 -50.731898
C 11.0112581 25.7338392 -49.6079951
H 10.8575365 26.7986213 -49.7828324
H 10.658697 25.5126149 -48.6002629
C 10.1424015 24.9839405 -50.6247752
O 10.5934761 24.3664138 -51.5757024
N 8.8434225 24.9954737 -50.4307784
H 8.2650384 24.4828781 -51.0906918
H 8.4331033 25.5005896 -49.6701504
N 11.2359635 22.4621827 -48.7844512
H 11.0578115 22.9049529 -47.8891113
C 10.3429926 21.383246 -49.214284
H 10.5949631 21.0920705 -50.235015
C 8.8711568 21.8264889 -49.2272665
O 8.3637704 22.3460571 -48.2286926
C 10.5732244 20.1618264 -48.309774
H 10.3332883 20.4190762 -47.2777976
H 11.6227969 19.8672222 -48.3609254
S 9.5400098 18.7650291 -48.8429061
H 8.3402798 19.3259352 -48.6493615
N 8.1565084 21.4877271 -50.299685
H 8.6580875 21.1050509 -51.0997332
C 6.6915622 21.4723115 -50.3796701
H 6.2622286 21.8728319 -49.4665579
C 6.2201094 20.031677 -50.5303368
O 6.4399128 19.4158451 -51.5664035
C 6.1943466 22.3420754 -51.5434893
H 6.7074003 22.060723 -52.4640156
C 4.6826819 22.2560971 -51.7615391
H 4.1511206 22.4652032 -50.8329504
H 4.408085 21.2625488 -52.117419
H 4.3832472 22.9787312 -52.5207642
O 6.4597851 23.693421 -51.2481149
H 6.0035097 23.908674 -50.423727
N 5.5750189 19.4858299 -49.5003679
H 5.4489048 20.0421054 -48.6609263
C 4.8934822 18.1915638 -49.5707935

H 5.3805521 17.5583986 -50.3075649
H 4.9561912 17.6920084 -48.6044006
C 3.4178491 18.3381219 -49.9451647
O 2.7499778 19.2457229 -49.4508098
N 2.8759735 17.4192718 -50.7447628
H 3.489359 16.728848 -51.172119
C 1.4218424 17.2768095 -50.9304672
H 0.9352328 17.8212694 -50.1261785
C 0.9560668 15.8305307 -50.7697592
O 1.6406314 14.8966804 -51.1867693
C 0.9136146 17.8905378 -52.2565359
H -0.1448096 17.6360649 -52.3435319
C 1.6387051 17.3248981 -53.5004349
H 2.6843131 17.6363338 -53.4916327
H 1.6106378 16.2363657 -53.4623999
C 0.9961788 19.4249524 -52.1858548
H 2.0365676 19.7521869 -52.1926341
H 0.5247676 19.7783184 -51.266841
H 0.4642133 19.8755679 -53.0218273
C 1.0052803 17.7519525 -54.8316522
H 1.1088672 18.8260369 -54.9781292
H -0.0515642 17.481976 -54.8436029
H 1.510511 17.2397663 -55.65089
N -0.2477491 15.6495421 -50.2260329
H -0.7465903 16.4668413 -49.8863459
C -0.9656272 14.3704334 -50.212057
H -0.6470092 13.784003 -51.0743246
C -2.4736961 14.6046495 -50.3361702
O -2.9923013 15.6108318 -49.855645
C -0.6345655 13.5626088 -48.9469876
H -1.035592 14.063649 -48.0667914
H 0.4483293 13.4718768 -48.8486547
O -1.2039403 12.2647033 -49.0661658
H -0.8230701 11.6633302 -48.3915009
N -3.189996 13.6624306 -50.9545782
H -2.693636 12.8593069 -51.3085801
C -4.6570956 13.7038033 -51.1109514
H -4.9453094 14.6681594 -51.5324737
C -5.3659256 13.5807981 -49.7505522
O -6.4750307 14.0756237 -49.5707605
C -5.0838402 12.5810381 -52.0946599
H -4.5868338 11.656628 -51.7934535
C -6.60297 12.3174183 -52.0927149
H -7.1427278 13.2367823 -52.3255887
H -6.9278242 11.951214 -51.1188279
H -6.8636599 11.5560362 -52.8261546

C -4.6265523 12.9470809 -53.5247128
H -5.2699191 13.744181 -53.8956398
H -3.6030739 13.3215718 -53.5000957
C -4.6555746 11.788384 -54.5300713
H -5.6777838 11.458972 -54.7113472
H -4.0625365 10.9539342 -54.1543668
H -4.2321241 12.1260136 -55.4769672
N -4.7271601 12.913471 -48.788543
H -3.775143 12.6272275 -48.9723444
C -5.2999645 12.5558781 -47.4919011
H -6.3581933 12.3426238 -47.6238231
C -5.12327 13.6827605 -46.4548777
O -3.9915082 13.9538372 -46.0403971
C -4.6452439 11.2446981 -47.0565748
H -3.587913 11.2809028 -47.3174804
H -5.1004324 10.4159779 -47.6001787
O -4.7366918 11.0128571 -45.6723862
H -5.6758702 11.0025864 -45.3613888
N -6.2181431 14.2734505 -45.9265974
C -6.1448261 15.2535177 -44.8392723
H -5.5037262 16.0768951 -45.1469994
C -5.5994107 14.6593758 -43.5365247
O -4.990551 15.3639187 -42.7414758
C -7.5767455 15.7683423 -44.6357585
H -8.056633 15.2366621 -43.8112928
H -7.5919532 16.8425494 -44.4489233
C -7.6005661 14.1249832 -46.3585443
H -7.6892693 14.0050037 -47.4364082
H -8.051605 13.2737373 -45.845865
C -8.2932888 15.4153499 -45.9362787
H -8.111043 16.1900229 -46.680453
H -9.3633916 15.2698172 -45.7853592
N -5.798928 13.3603836 -43.3244477
H -6.3334307 12.8472974 -44.0199305
C -5.3819279 12.5909009 -42.1533687
H -5.5086175 13.2194111 -41.2710266
C -3.889537 12.1953296 -42.1819728
O -3.2267231 12.2061379 -41.1417984
C -6.3412377 11.3905832 -41.9922851
H -7.3113502 11.7703061 -41.6675729
H -5.9575394 10.7498656 -41.1971473
C -6.5663179 10.5388814 -43.2538189
O -6.908747 11.078591 -44.3329274
O -6.3508923 9.3107568 -43.1812859
N -3.3008267 11.9683216 -43.3619274
H -3.8866016 11.9470974 -44.1933319

C -1.8430178 11.9831623 -43.5618261
H -1.3723892 11.257173 -42.9150736
C -1.2618897 13.3502526 -43.1982805
O -0.4067809 13.4546671 -42.3135894
C -1.4776396 11.6187214 -45.0103698
H -2.1218696 12.1563712 -45.7031098
H -0.4492902 11.9202012 -45.2048059
C -1.5826308 10.1109484 -45.2530965
H -2.542401 9.7508069 -44.8866851
H -0.8029225 9.6052696 -44.6769365
C -1.4265517 9.7523301 -46.7342513
O -2.1887154 8.9014817 -47.2378949
O -0.4387212 10.1870761 -47.3704002
N -1.8238071 14.3981459 -43.7998905
H -2.5652821 14.2183388 -44.4716944
C -1.3629664 15.7743681 -43.644951
H -0.2966211 15.7971953 -43.8619811
C -1.5011144 16.2671664 -42.1931932
O -0.5669668 16.8774303 -41.6842395
C -2.0937697 16.6432986 -44.6956286
H -3.1616424 16.4331521 -44.6208048
C -1.6453572 16.3062192 -46.1415492
H -1.8060816 15.2459139 -46.3228022
H -2.2857106 16.8442621 -46.8391984
C -1.9169274 18.1412204 -44.4519846
H -0.8624135 18.3648704 -44.4776171
H -2.351247 18.4235725 -43.4927015
H -2.4173335 18.7036014 -45.2397571
C -0.1850993 16.6247081 -46.5047295
H 0.502714 16.0852862 -45.8550906
H 0.005957 17.6948585 -46.4390428
H -0.0012286 16.3148047 -47.5317343
N -2.5671139 15.9256739 -41.4689525
H -3.3267024 15.4447242 -41.9411177
C -2.7735512 16.3191514 -40.0675664
H -2.7786111 17.4067291 -40.0129945
C -1.6436206 15.8404344 -39.1348382
O -1.1884872 16.6049745 -38.2795479
C -4.1712491 15.8244989 -39.6523631
H -4.8964345 16.3669418 -40.2600162
H -4.276831 14.7663523 -39.8970248
C -4.5547229 16.0502673 -38.1793378
H -5.6347657 16.1922723 -38.1337433
H -4.0852274 16.9595111 -37.8033673
C -4.2106249 14.87867 -37.2602295
O -3.478791 13.9624486 -37.5900405

N -4.7524033 14.838826 -36.0651956
H -4.5608134 14.0108061 -35.5281391
H -5.3835633 15.5549238 -35.7602172
N -1.1004638 14.6424705 -39.3708469
H -1.5013084 14.0764096 -40.106404
C 0.0418472 14.1322373 -38.6116889
H -0.0436926 14.4501244 -37.5713118
C 1.3850099 14.6518156 -39.1239408
O 2.2660308 14.8908551 -38.3035319
C 0.0435491 12.6104591 -38.6527647
H 1.0465421 12.2488474 -38.4138007
H -0.2262749 12.3017169 -39.6598596
S -1.1117245 11.9445062 -37.4241918
H -2.0986728 12.845829 -37.6105998
N 1.5575739 14.8170369 -40.4417058
H 0.8054631 14.5690666 -41.0762493
C 2.7701594 15.422793 -40.987119
H 3.6444812 14.9045317 -40.5883586
C 2.8696392 16.8863013 -40.5636503
O 3.5847836 17.2487937 -39.6283336
C 2.7907183 15.3258836 -42.5238653
H 1.9156234 15.7985274 -42.9702768
H 2.7822393 14.2994419 -42.8465516
H 3.7036186 15.7788831 -42.9060338
N 2.1002966 17.7180647 -41.2791163
H 1.1970245 17.3057366 -41.4945638
C 2.6117307 18.5517644 -42.3861192
H 3.1341734 17.8723516 -43.0516016
C 1.3970127 19.0600589 -43.2242848
H 1.358215 18.4262558 -44.0974844
H 0.5018309 18.6692998 -42.7733033
C 1.1166488 20.5727843 -43.5444475
H 1.2915895 21.151315 -42.6635274
H 1.8411256 20.9728381 -44.2381746
C -0.3230066 20.9360951 -43.9914732
H -1.0227159 20.3720331 -43.3859128
H -0.4699022 20.6178342 -45.0179328
C -0.6963571 22.4298742 -43.8025919
H 0.1995555 22.9755746 -43.4898631
H -1.4260614 22.4979107 -42.9928553
N -1.2635441 23.0865246 -45.012343
H -0.5895041 23.0906335 -45.7804691
H -1.4992876 24.0597524 -44.8902245
H -2.0834365 22.6400856 -45.4285437
C 3.7642777 19.4606364 -41.9383207
O 4.6934606 18.9814363 -41.3095185

N 3.6311882 20.7771193 -41.9989438
H 2.8876535 21.1417414 -42.5604266
C 4.0788258 21.6000937 -40.8865894
H 5.1237588 21.3834285 -40.6673247
C 3.9999911 23.0820359 -41.3327063
H 3.0493725 23.2473995 -41.8409777
H 4.7944901 23.2717171 -42.0553756
C 4.1026739 24.1179475 -40.2399803
N 3.0918861 25.0331692 -39.9409579
C 3.4223691 25.5589047 -38.7527186
H 2.8010558 26.2448663 -38.192991
N 4.5752577 25.0358408 -38.3042514
H 4.8930887 25.1113105 -37.3429746
C 5.0364284 24.1390869 -39.2428601
H 5.8685618 23.4582498 -39.1352582
C 3.2420288 21.199852 -39.6385361
O 3.0797663 22.025223 -38.7500341
N 2.530232 20.0383135 -39.640792
H 2.865367 19.2416506 -40.1623869
C 1.4823966 19.8353562 -38.7041374
H 1.2481924 20.7922532 -38.2384218
C 2.0858318 19.0933861 -37.5361898
O 3.067777 19.6165891 -37.0050026
C 0.1413169 19.4928001 -39.3442083
H 0.1442248 18.4588372 -39.6817178
H -0.6141341 19.569777 -38.5611991
C -0.267998 20.4123699 -40.5175301
H 0.1183068 19.9509151 -41.3678312
C -1.7776922 20.4353344 -40.70414
H -2.2492346 20.8376265 -39.8066394
H -2.1313518 19.4171078 -40.8620015
H -2.0806715 21.041301 -41.5528518
C 0.1944014 21.8747087 -40.5369619
H 1.2344464 22.003532 -40.7787416
H 0.035205 22.3050513 -39.5470494
H -0.3843236 22.4742456 -41.2283432
N 1.4878269 18.020571 -37.0275607
H 0.680745 17.619781 -37.490007
C 1.8370793 17.567107 -35.682473
H 1.6480608 18.3923588 -34.994206
C 3.3255548 17.1909504 -35.5105984
O 3.975249 17.6990186 -34.5966676
C 0.897452 16.4265003 -35.2938252
H 1.0844826 15.5580817 -35.9233497
H -0.1416476 16.7397962 -35.4098183
H 1.0723946 16.150676 -34.2533891

N 3.895938 16.3935091 -36.4204664
H 3.3273487 16.0340792 -37.1822979
C 5.2891756 15.9425756 -36.3358269
H 5.4806631 15.6036171 -35.3165186
C 6.3276383 17.0397494 -36.6405866
O 7.4910534 16.9063308 -36.2670223
C 5.4618095 14.744717 -37.274927
H 5.2733635 15.0489023 -38.3063018
H 4.7617187 13.9547081 -37.0015401
H 6.47971 14.3605827 -37.1961688
N 5.9285687 18.1328717 -37.2982429
H 4.962632 18.1795496 -37.6016175
C 6.8314934 19.2418052 -37.635999
H 7.8429605 18.8505207 -37.6832907
C 6.844892 20.3829303 -36.6121256
O 7.5699517 21.3645417 -36.8076774
C 6.4883151 19.7654549 -39.0249614
H 5.4763035 20.1549587 -38.9918722
H 6.5452826 18.940506 -39.7349417
O 7.3909096 20.7802827 -39.4271918
H 7.5897174 21.3108912 -38.6363837
N 6.0301942 20.3258281 -35.5583977
H 5.4633655 19.4990513 -35.4131305
C 5.9102691 21.4436743 -34.6242259
H 5.8569889 22.3420848 -35.2375565
C 7.1639527 21.593333 -33.7492034
O 7.6308825 20.6298769 -33.149975
C 4.6035516 21.3149437 -33.8160339
H 3.9610644 20.5586151 -34.2705869
H 4.8261419 20.983383 -32.8007128
C 3.8146552 22.6339158 -33.7630863
H 4.3761121 23.361526 -33.1761685
H 2.8597389 22.454182 -33.2696363
C 3.5503843 23.2164719 -35.1512038
O 4.0029425 24.3035214 -35.4804174
N 2.9488639 22.4629638 -36.0411139
H 2.950617 22.7735293 -37.0102779
H 2.7918841 21.4817724 -35.8719266
N 7.7210904 22.8081187 -33.6914249
H 7.2987774 23.5567177 -34.2190106
C 8.9972279 23.0788989 -33.0131334
H 9.0701395 24.1482844 -32.8145984
H 9.0214978 22.5526099 -32.0586359
C 10.2493908 22.6746477 -33.805537
O 11.3286693 22.5814544 -33.2268775
N 10.1271567 22.3925806 -35.1093067

H 9.2142375 22.4390954 -35.5407744
C 11.2828804 22.1739636 -35.9765919
H 12.1470555 21.9182832 -35.3639204
C 11.6311631 23.4425166 -36.7703543
O 10.8468171 23.8739616 -37.6062429
C 11.0248787 21.0051895 -36.9419587
H 10.2282802 21.2982624 -37.627223
H 11.9359448 20.8776523 -37.5163873
C 10.6521745 19.6372191 -36.3484215
H 9.6886744 19.7031776 -35.8480046
C 10.554009 18.6256642 -37.4922608
H 11.5421073 18.4392663 -37.908598
H 9.9033941 19.0025286 -38.2786216
H 10.1402736 17.6890333 -37.1141196
C 11.700169 19.1190278 -35.363519
H 12.6761902 19.0615675 -35.8409176
H 11.4049478 18.1266817 -35.0192001
H 11.7434391 19.7788958 -34.4960605
N 12.8948299 23.8661966 -36.7091582
H 13.3352564 23.7065076 -35.812184
C 13.4155394 25.0179326 -37.4790076
H 12.6681205 25.8145446 -37.4282954
C 13.6732908 24.7387133 -38.9836209
O 14.3679382 25.4938641 -39.6582383
C 14.7076514 25.5493859 -36.8171534
H 15.5330047 24.8870948 -37.0856383
H 14.9254605 26.5347873 -37.2302733
C 14.686988 25.6625318 -35.2845744
H 15.5866248 26.1902592 -34.9596257
H 14.7320807 24.6561241 -34.8586402
C 13.4378158 26.3886491 -34.7758366
O 13.4280188 27.6368107 -34.819975
O 12.4900572 25.6633847 -34.4019596
N 13.2648524 23.5772005 -39.5105643
H 12.5354699 23.085907 -39.0068225
C 13.5570177 23.1795007 -40.8976494
H 14.5254876 23.6009869 -41.1761233
C 12.542468 23.7627457 -41.8860466
O 11.3445727 23.7642826 -41.6133993
C 13.6717175 21.6505914 -41.0511484
H 14.6737933 21.3409764 -40.7633607
H 13.5409867 21.3814689 -42.0977122
C 12.6807254 20.8409029 -40.234541
O 13.0819143 19.9571612 -39.4998137
N 11.4063889 21.1652132 -40.2593715
H 10.7663929 20.6171536 -39.7086994

H 11.0835354 21.9909526 -40.7547637
N 13.0133136 24.1123106 -43.0888082
H 14.003369 23.9861169 -43.252692
C 12.2559559 24.8063065 -44.14478
H 11.634147 25.5703974 -43.6725806
C 11.2900693 23.8636724 -44.8836361
O 11.4794884 23.5296958 -46.0528063
C 13.2402964 25.5207442 -45.0935189
H 13.7974318 24.7585126 -45.6299273
H 12.6747639 26.1075278 -45.8195301
C 14.2367 26.4457212 -44.363787
H 13.6782029 27.2076148 -43.8182901
H 14.8081668 25.8651489 -43.6389009
C 15.2530414 27.1445263 -45.2802328
H 15.9662655 27.6665972 -44.6388995
H 14.7410762 27.8844194 -45.8974409
C 16.0017683 26.1590706 -46.1895801
H 15.4383225 26.0365783 -47.1185131
H 16.0485386 25.1791219 -45.7028254
N 17.3833513 26.6013149 -46.4784672
H 17.4501689 27.5257558 -46.8646949
H 17.8394313 25.9286781 -47.0934372
H 17.9672793 26.5200239 -45.6378488
N 10.2586788 23.4106932 -44.1767312
H 10.1841574 23.7178249 -43.2124954
C 9.2046284 22.5279462 -44.6902692
H 9.4646982 22.2318761 -45.7066837
C 7.8664666 23.2561846 -44.773848
O 7.5173305 24.0732073 -43.9268036
C 9.0728533 21.2239852 -43.8778855
H 8.316634 20.6091786 -44.3642886
C 10.3766276 20.4230278 -43.8853184
H 11.1563743 20.9672999 -43.3545985
H 10.6901088 20.2389319 -44.9118237
H 10.2204614 19.4614051 -43.3955968
C 8.6444678 21.4437578 -42.4225151
H 9.3296493 22.1190396 -41.9159669
H 8.6253233 20.491972 -41.8909169
H 7.6444986 21.8751438 -42.3818422
N 7.0848025 22.9020958 -45.7835561
H 7.4542566 22.2468272 -46.4660422
C 5.688535 23.2901281 -45.9603944
H 5.2592379 23.5642908 -44.9971347
C 4.9368154 22.090787 -46.5162855
O 5.5175932 21.2799915 -47.2402207
C 5.572196 24.4861637 -46.9093264

H 6.1315988 25.3254024 -46.4927753
H 4.5253632 24.7775338 -47.0057321
O 6.0920421 24.166681 -48.1936715
H 6.9019288 23.6412406 -48.069159
N 3.6649475 21.9328174 -46.1572492
H 3.1924113 22.6509743 -45.6290061
C 2.8540294 20.891737 -46.7718369
H 3.2603949 20.7420939 -47.768523
C 1.4232672 21.3469321 -46.9993605
O 0.9655742 22.2461308 -46.2958749
C 2.8723023 19.5311039 -46.0727033
H 1.8998189 19.3593649 -45.6756641
H 2.831625 18.8431272 -46.9167144
C 4.0238134 18.9905664 -45.2169582
C 4.3071869 17.6253821 -45.3860567
H 3.6888653 17.0224533 -46.0309537
C 4.8919789 19.744078 -44.3979068
H 4.7530479 20.7996534 -44.2590226
C 5.3992881 17.0235059 -44.752768
H 5.5957283 15.9801922 -44.9285487
C 5.9619852 19.1264943 -43.7177794
H 6.5828412 19.700868 -43.0462872
C 6.2154413 17.761582 -43.8888187
H 7.0295482 17.2877361 -43.3583586
N 0.7049211 20.6843602 -47.9071269
H 1.1580329 19.9917536 -48.4999263
C -0.7426341 20.8387991 -48.0115702
H -1.0824204 20.9641876 -46.9898499
C -1.5361428 19.617803 -48.4852348
O -0.9924186 18.6595985 -49.0384563
C -1.11455 22.1121096 -48.8003637
H -1.7026334 21.8618073 -49.6844732
H -0.2187602 22.6483505 -49.119513
C -1.9431406 22.9948283 -47.8556361
O -2.8487189 22.4398679 -47.1824651
O -1.4945848 24.1167787 -47.540539
N -2.8489941 19.6723543 -48.2388324
H -3.2118883 20.5386921 -47.8451067
C -3.8093695 18.7281656 -48.8164363
H -3.3973647 17.7249245 -48.7310293
C -3.9734597 19.0752603 -50.2936677
O -4.3973474 20.175993 -50.6415666
C -5.1732061 18.7435119 -48.0992768
H -5.6730347 19.6950951 -48.2825975
C -6.0637773 17.6068255 -48.6193746
H -5.6052813 16.639463 -48.4108445

H -6.2151491 17.7031978 -49.6940242
H -7.0395154 17.6576898 -48.1394967
C -5.0359728 18.5600012 -46.5826229
H -4.4763312 17.6507279 -46.373113
H -6.0235974 18.4938287 -46.1264681
H -4.5200769 19.4177832 -46.1531065
N -3.6215795 18.1436875 -51.1741785
H -3.2879382 17.2507786 -50.8282856
C -3.6115106 18.3903012 -52.6077545
H -4.5005105 18.9612831 -52.879691
H -2.7342712 18.9845956 -52.864576
C -3.5764643 17.1062873 -53.4192378
O -2.760003 16.2192524 -53.1788795
N -4.4347018 17.0516582 -54.4326559
H -5.0732818 17.8204822 -54.5581563
C -4.3864658 16.05468 -55.4956002
H -4.0915978 15.0912157 -55.0883004
C -3.3390088 16.4662952 -56.5310674
O -3.4495113 17.5140222 -57.164842
C -5.7890218 15.9175663 -56.0942179
H -6.1264015 16.9003061 -56.4288134
H -6.4684305 15.5787826 -55.3090581
C -5.884318 14.9468145 -57.2730016
O -4.8383242 14.4918336 -57.7921059
O -7.0322555 14.7422459 -57.7062923
N -2.3237965 15.6294866 -56.7222922
H -2.3348741 14.7515165 -56.2290766
C -1.1959614 15.917138 -57.5968154
H -0.8042434 16.9026035 -57.3349828
C -1.5477935 15.9520276 -59.0940706
O -0.7986989 16.5414193 -59.8719076
C -0.1261968 14.8713849 -57.2991538
H -0.4332321 13.8964363 -57.6832249
H 0.0211865 14.8034629 -56.2217764
H 0.8044678 15.171372 -57.7763393
N -2.6894224 15.3899282 -59.5099429
H -3.3057519 14.9434825 -58.8268714
C -3.1680469 15.5081525 -60.8889374
H -2.2977069 15.4764396 -61.5426236
C -3.8714604 16.8578481 -61.1353749
O -3.8881519 17.3427345 -62.2717246
C -4.0589673 14.2899356 -61.2019821
H -3.5800509 13.3944349 -60.8022451
H -5.0094949 14.4145144 -60.6792547
C -4.3336983 14.054691 -62.7013104
H -4.8066073 14.9365606 -63.1323418

C -3.0607759 13.732287 -63.4923323
H -2.5395826 12.8897156 -63.037984
H -2.3987076 14.5952794 -63.5160859
H -3.3198034 13.4669686 -64.5171777
C -5.2831101 12.8667808 -62.868187
H -4.8269732 11.9632398 -62.4652122
H -5.5106869 12.7195113 -63.9246257
H -6.2135557 13.0676634 -62.3361046
N -4.3438173 17.5317856 -60.0767708
H -4.239743 17.0913676 -59.1689351
C -5.1392953 18.7670931 -60.1508346
H -5.3101043 19.014173 -61.1954728
C -4.4652452 20.0192241 -59.553186
O -4.7247452 21.1197762 -60.036646
C -6.5275032 18.4864972 -59.5562995
H -7.1041324 19.4099779 -59.5149897
H -6.4332717 18.0886791 -58.546576
C -7.2894454 17.5033969 -60.4354308
O -7.5563357 17.7986585 -61.5883171
N -7.5704357 16.2932746 -60.0008698
H -8.0536807 15.6856149 -60.632247
H -7.3300548 15.9419432 -59.0695682
N -3.5291447 19.8666245 -58.6129897
H -3.3778296 18.9328279 -58.2480178
C -2.7161973 20.948246 -58.0426512
H -3.3974747 21.6988151 -57.6398965
C -1.8381347 21.6244684 -59.108357
O -1.3603305 20.9679296 -60.0376461
C -1.8708616 20.3801509 -56.8857777
H -1.2155043 19.5960797 -57.2688558
H -2.5403692 19.9339514 -56.1488536
C -1.0113161 21.4298223 -56.1713507
H -0.2443703 21.7912918 -56.8567282
H -0.5059511 20.9491715 -55.3327881
S -1.9402459 22.8506813 -55.5394919
C -0.5701828 23.8292679 -54.8690719
H -0.9613234 24.74235 -54.4197375
H 0.1193026 24.0926016 -55.6715556
H -0.0410525 23.252321 -54.1098706
N -1.6015408 22.9369568 -58.9811233
H -1.9423219 23.4054305 -58.1486339
C -0.8192637 23.7390643 -59.9395053
H -0.2576772 23.064157 -60.5851844
C 0.2282464 24.5957342 -59.2270807
O 0.004776 25.0706294 -58.1183271
C -1.7509353 24.6002576 -60.822006

H -2.2485006 25.3361505 -60.1875853
H -1.1470927 25.1432575 -61.5512182
C -2.8355023 23.8041089 -61.576147
H -3.4692309 24.5095959 -62.1148217
H -3.4670243 23.293783 -60.85025
C -2.285634 22.7794283 -62.5838831
H -1.469829 22.2166229 -62.1362081
H -1.8727529 23.3160784 -63.440655
N -3.3480238 21.8600974 -63.0454371
H -4.1592275 22.2924263 -63.4494333
C -3.3712553 20.539867 -62.9367282
N -4.3632524 19.8552986 -63.436589
H -4.394121 18.8635073 -63.2050823
H -5.2093213 20.3214733 -63.7056537
N -2.4470361 19.8412767 -62.3456859
H -1.8072108 20.3142319 -61.7116381
H -2.5589807 18.8412342 -62.2636225
N 1.3477479 24.8147152 -59.9086339
H 1.3980397 24.4714973 -60.8561952
C 2.4801481 25.6382775 -59.4817666
H 2.171737 26.310206 -58.6796175
C 2.9540436 26.4820188 -60.6742369
O 2.5493109 26.2124622 -61.8048482
C 3.6101897 24.7291168 -58.9715134
H 3.9556652 24.0994097 -59.7921348
H 4.4519802 25.3525014 -58.6710504
C 3.2414508 23.8502424 -57.790578
C 3.2255769 24.3988016 -56.4940977
H 3.4578802 25.4439975 -56.3432923
C 2.944317 22.484486 -57.9805326
H 2.957988 22.0603603 -58.9749413
C 2.9118273 23.5871558 -55.3886793
H 2.893409 23.9989979 -54.390721
C 2.6376701 21.6672874 -56.8746023
H 2.4174114 20.6212776 -57.0198648
C 2.6196309 22.2218223 -55.5761608
O 2.3179465 21.4526177 -54.4997302
H 2.18435 20.5319836 -54.724826
N 3.8334992 27.4628261 -60.4509498
H 4.2082822 27.584594 -59.5219213
C 4.4939213 28.1620494 -61.5616058
H 3.7309422 28.5488971 -62.2384471
C 5.3882736 27.2020676 -62.3485732
O 6.0176283 26.3114741 -61.7616727
C 5.3217809 29.3477141 -61.0436929
H 6.091829 28.9980095 -60.3569351

H 4.6612574 30.029898 -60.5071673
O 5.9393037 30.0538829 -62.1032955
H 6.7243398 29.5613144 -62.4093461
N 5.5820288 27.4940815 -63.6317522
H 5.0106585 28.1958685 -64.0746187
C 6.7307802 26.9874338 -64.3760478
H 6.6366855 25.9085016 -64.4254029
C 8.028407 27.3558119 -63.6304401
O 8.1187891 28.4400932 -63.0417593
C 6.7479946 27.5450176 -65.8100117
H 7.719706 27.3149662 -66.2505881
H 6.6485008 28.6315395 -65.7731873
C 5.6672195 26.9697377 -66.7358325
O 4.772257 26.2412801 -66.2508656
O 5.7522914 27.2543315 -67.9504335
N 9.0046514 26.4438149 -63.5827792
H 8.8767 25.5757138 -64.0999271
C 10.2928452 26.6834173 -62.9168859
H 10.9628395 25.863104 -63.1596956
H 10.726691 27.5994604 -63.3176861
C 10.2581107 26.8168125 -61.3836782
O 11.1929088 27.3706979 -60.812188
N 9.2068629 26.3463782 -60.7000323
H 8.4713915 25.8971 -61.2280263
C 9.0731089 26.4374344 -59.2283582
H 9.1851045 27.4791121 -58.9258279
C 10.1069491 25.6136815 -58.4432941
O 10.4836519 25.994552 -57.3288833
C 7.6897912 25.9495096 -58.7796666
H 7.6527518 25.9341602 -57.6888603
H 7.5223822 24.9374057 -59.148062
O 6.6580577 26.7898418 -59.2502181
H 6.5249451 26.5892104 -60.1998395
N 10.5092653 24.4553591 -58.9775317
H 10.148396 24.2047745 -59.8925566
C 11.2703098 23.4359545 -58.2511685
H 11.53296 23.831252 -57.2725145
C 12.5826537 23.0539197 -58.9343996
O 12.7237566 23.0892687 -60.153064
C 10.4016771 22.1945978 -58.0069762
H 10.1678104 21.7295215 -58.9665107
H 10.9725225 21.4720044 -57.4234963
C 9.1174901 22.4918067 -57.2616053
C 9.1440574 22.7882535 -55.8861933
H 10.0770687 22.7663086 -55.3446341
C 7.8965917 22.5106125 -57.9559699

H 7.8822044 22.2996273 -59.0114969
C 7.9532069 23.1130515 -55.2127899
H 7.9771022 23.3415634 -54.1554797
C 6.7052395 22.8277978 -57.284616
H 5.7704859 22.841606 -57.8236995
C 6.7337942 23.135257 -55.9133376
H 5.8189192 23.3828901 -55.3936062
N 13.517583 22.5839715 -58.1188466
H 13.3072881 22.5350507 -57.1278463
C 14.8179288 22.0601079 -58.5444263
H 15.0314648 22.3351386 -59.5799209
C 14.8820061 20.5283609 -58.4571595
O 15.7122647 19.8905221 -59.1081171
C 15.861854 22.6970743 -57.6264017
H 15.4958654 22.6849266 -56.6000881
H 16.7664742 22.1004187 -57.6568063
C 16.1785143 24.1454572 -58.0345006
H 16.4935678 24.6931106 -57.143149
H 15.2907343 24.6425045 -58.4312527
C 17.305602 24.1476694 -59.067901
O 17.1190339 23.5958259 -60.1692975
O 18.44763 24.4781666 -58.6847995
N 14.0008482 19.9431131 -57.6415347
H 13.375073 20.5442361 -57.11928
C 13.8557196 18.5135042 -57.3961515
H 14.3000178 17.9505765 -58.2132188
C 12.3562485 18.1809218 -57.2916426
O 11.6055412 18.9392478 -56.6695756
C 14.5613807 18.1437631 -56.0764476
H 13.9703084 18.5856413 -55.2756477
H 14.5331341 17.0604963 -55.9537592
C 16.0075358 18.6382877 -55.8749913
H 16.0774433 19.7022195 -56.0961422
C 16.4272853 18.4705187 -54.4170553
H 16.4803628 17.4129 -54.1558347
H 15.7119752 18.972793 -53.7650525
H 17.3987076 18.9361346 -54.2814031
C 17.0207935 17.8886108 -56.7338335
H 16.9321508 16.814618 -56.5697651
H 18.0321975 18.2073195 -56.4892937
H 16.8337156 18.1120355 -57.7796173
N 11.9268476 17.0254245 -57.802589
H 12.5667476 16.4679935 -58.3580903
C 10.6134922 16.4408486 -57.4727742
H 10.200647 16.9817986 -56.6240596
C 10.8460628 14.9974708 -57.0338666

O 11.4968357 14.2196056 -57.7343886
C 9.584129 16.5453119 -58.623281
H 9.8891 15.8453796 -59.384544
C 9.5544193 17.9798466 -59.2141413
H 9.2019235 18.6905423 -58.4697116
H 10.5636784 18.2729503 -59.500348
C 8.1844474 16.0604285 -58.2049168
H 7.7911132 16.6907992 -57.4099126
H 8.2346799 15.0332518 -57.8434264
H 7.5072373 16.083636 -59.0554359
C 8.7221937 18.1222675 -60.4791129
H 7.6661354 18.1114382 -60.2270789
H 8.9737782 17.3097286 -61.1536164
H 8.9770545 19.0592215 -60.9698104
N 10.3564225 14.6574374 -55.8469035
H 9.7941865 15.3397182 -55.3446681
C 10.5610505 13.361721 -55.2088406
H 11.1527113 12.721799 -55.8623133
C 9.2265463 12.6740554 -54.9607565
O 8.2716464 13.3041168 -54.5053329
C 11.33476 13.5316808 -53.9000093
H 12.3374437 13.8914798 -54.1303004
H 10.8329836 14.2807638 -53.2847405
C 11.4560679 12.2414929 -53.107116
C 12.2584577 11.1893237 -53.5913009
H 12.8093437 11.3079028 -54.5115507
C 12.3285074 9.9712448 -52.892086
H 12.935959 9.1595666 -53.2719667
C 11.6002187 9.80203 -51.7022888
H 11.6597189 8.8679635 -51.1608939
C 10.7885537 10.8448729 -51.2227153
H 10.2173261 10.7141468 -50.3149552
C 10.7115946 12.0603749 -51.9250526
H 10.0647863 12.8469327 -51.5572318
N 9.1602094 11.3777451 -55.2613215
H 9.967209 10.9333375 -55.6892098
C 7.9328766 10.5948167 -55.1189933
H 7.301889 11.0701356 -54.3646456
C 8.255057 9.1831792 -54.6477118
O 9.0891373 8.509012 -55.2435789
C 7.1597415 10.5440188 -56.4471444
H 7.7114147 9.9222563 -57.1494066
C 5.8095002 9.8629732 -56.2113163
H 5.4896513 10.0212756 -55.1882349
H 5.9140703 8.7885214 -56.3956002
H 5.0585275 10.2781186 -56.8705113

C 7.0334598 11.9183157 -57.1168015
H 6.7794035 12.6694729 -56.3779092
H 6.2773547 11.9109892 -57.8926942
H 7.9869524 12.1928344 -57.5734035
N 7.5304832 8.6922801 -53.6430702
H 6.8760449 9.3036927 -53.1726615
C 7.6681163 7.3252057 -53.1358409
H 8.2068826 6.7406371 -53.8801015
C 6.2876119 6.6673452 -52.9862792
O 5.4097723 7.1751835 -52.2984167
C 8.5623854 7.3067255 -51.8731256
H 9.5579647 7.6240205 -52.1890165
C 8.094785 8.2976582 -50.7891014
H 7.0661867 8.0877915 -50.5071326
H 8.1462975 9.3195321 -51.1659953
H 8.7354529 8.2405992 -49.9112774
C 8.6896144 5.865025 -51.3446308
H 7.7397634 5.5693468 -50.9079072
H 8.8984866 5.198318 -52.1818034
C 9.804598 5.6643442 -50.3096542
H 9.5898272 6.2220642 -49.3977556
H 10.7584064 5.9929064 -50.7211156
H 9.8773923 4.6049462 -50.0589894
N 6.0735872 5.5791504 -53.7322323
H 6.8322139 5.3057652 -54.3480851
C 4.8587987 4.7465541 -53.7745885
H 4.9169297 4.1923917 -54.7089597
C 3.516118 5.5057891 -53.8614048
O 2.5191161 5.1118456 -53.2532807
C 4.8607276 3.7088065 -52.6438668
H 4.6254463 4.2273965 -51.7263386
H 4.0580549 2.9937707 -52.8329263
C 6.1573545 2.9237375 -52.4338794
H 5.9667513 2.1123083 -51.7318169
H 6.9089625 3.5794581 -51.9982039
C 6.7097361 2.3381834 -53.7201912
O 7.8204849 2.6367907 -54.124086
N 5.9596121 1.5408023 -54.4434662
H 6.3860314 1.1601171 -55.2659575
H 4.9443976 1.5731702 -54.3602218
N 3.4922299 6.609794 -54.6151327
H 4.3734201 6.9183661 -54.9925454
C 2.3102518 7.4619524 -54.7883182
H 1.523502 7.1422596 -54.1042785
C 1.7272705 7.3825057 -56.209906
O 0.5213259 7.2247492 -56.3878293

C 2.7044591 8.890572 -54.3960494
H 3.4554001 9.2689719 -55.0839756
H 3.1257726 8.8895343 -53.3873539
S 1.267988 9.9956678 -54.455393
H 1.9581013 11.1116626 -54.1764562
N 2.5785643 7.3956426 -57.2408166
H 3.5667076 7.4519611 -57.0465178
C 2.1412431 7.4998195 -58.6453044
H 1.3706166 8.2684934 -58.6961616
C 1.455654 6.224998 -59.1639819
O 0.6846533 6.2812467 -60.1185169
C 3.3271499 7.9979964 -59.498981
H 4.2271993 7.4604443 -59.1935633
C 3.5120954 9.5030321 -59.2234801
H 2.7433385 10.070109 -59.7525158
H 3.3888159 9.7067547 -58.1618482
C 3.1360484 7.7830679 -61.0060698
H 2.1735433 8.1823495 -61.3276595
H 3.2063335 6.7193164 -61.2171017
H 3.9295564 8.2650958 -61.5743967
C 4.8911054 10.0097662 -59.6504475
H 5.0099305 9.9628237 -60.7314375
H 5.6735898 9.4236341 -59.1685517
H 4.9797716 11.0455733 -59.3443017
N 1.5717582 5.1041732 -58.4564899
H 2.1827045 5.1206104 -57.6529454
C 0.7328439 3.9149034 -58.651388
H 0.8947388 3.5462889 -59.6639284
C -0.7823536 4.1812482 -58.5331897
O -1.5480582 3.5308124 -59.2486143
C 1.1982313 2.8047712 -57.6992863
H 0.5310809 1.9524817 -57.8022436
H 2.1807305 2.4791663 -58.0396875
C 1.2976814 3.1948038 -56.2119626
H 0.3012642 3.1688227 -55.7657597
H 1.6944627 4.205053 -56.0979221
C 2.2384522 2.2254211 -55.4939914
O 3.4720441 2.4194285 -55.590334
O 1.7426194 1.2099946 -54.9540701
N -1.2002262 5.2068174 -57.7761104
H -0.5129244 5.7123035 -57.2254629
C -2.598939 5.6605511 -57.6700557
H -3.2650489 4.7962077 -57.7183515
C -3.0249164 6.6303356 -58.7861373
O -4.212663 6.7435325 -59.0897442
C -2.8084363 6.3407486 -56.3074555

H -3.8525083 6.6461592 -56.2269157
H -2.1951503 7.2411331 -56.2540386
C -2.494111 5.4648105 -55.1222584
N -3.2748369 4.3995001 -54.6813414
C -1.402258 5.5724542 -54.3123075
H -0.6083186 6.2963122 -54.4092074
C -2.6439835 3.8919381 -53.61534
H -2.9894624 3.0311948 -53.0587202
N -1.5076655 4.5702295 -53.3754836
H -0.8044458 4.3204002 -52.6953078
N -2.0794667 7.3391896 -59.4136958
H -1.1158875 7.1556197 -59.1719981
C -2.3616997 8.3287098 -60.4667779
H -3.2276859 8.9173499 -60.1602128
C -2.7464924 7.5875773 -61.7548098
O -2.001621 6.7304906 -62.2333305
C -1.169923 9.3091347 -60.6337605
H -0.2399088 8.7535995 -60.5132491
C -1.1908147 10.461597 -59.5993055
H -0.3002541 11.0742561 -59.749503
H -2.0610367 11.0934907 -59.7853356
C -1.1248122 9.968303 -62.0256172
H -2.0519598 10.5058078 -62.2273489
H -0.9656117 9.221869 -62.8030621
H -0.2929621 10.6711498 -62.0761009
C -1.2165483 10.0459058 -58.1225757
H -0.4028053 9.3553923 -57.9105932
H -2.1699243 9.5788426 -57.8756608
H -1.0975652 10.9321865 -57.4986943
N -3.9191623 7.9002296 -62.3131057
H -4.4943561 8.5906527 -61.8567087
C -4.4811881 7.1647627 -63.4544377
H -4.2521094 6.1048541 -63.3300885
C -3.8496852 7.5951677 -64.7889643
O -3.2727482 6.7640869 -65.4867586
C -6.0132403 7.3088003 -63.4796484
H -6.2694267 8.3537053 -63.6622205
H -6.4039576 6.7184002 -64.3096062
C -6.7166646 6.8608593 -62.1861027
H -7.7887787 7.0203155 -62.3018223
H -6.3872767 7.4812947 -61.3521355
C -6.4970237 5.3889815 -61.8459751
O -6.9995508 4.4963847 -62.5114945
N -5.7572628 5.0902401 -60.8042215
H -5.833909 4.14822 -60.4307188
H -5.3482376 5.8097848 -60.2156199

N -3.8569681 8.8981386 -65.0946658
H -4.3202539 9.5439063 -64.4785248
C -3.1401507 9.4583446 -66.2477274
H -3.0411782 8.6833109 -67.0107969
C -1.7236527 9.8698317 -65.8282051
O -1.4618108 10.9868417 -65.364135
C -3.9305454 10.5951731 -66.9132344
H -4.0322571 11.4261952 -66.2131145
H -4.9304066 10.2295566 -67.1538309
C -3.2659619 11.0967388 -68.2087631
O -2.0957811 10.7292587 -68.4770125
O -3.9030893 11.919111 -68.8986723
N -0.803794 8.9111941 -65.9612056
H -1.116004 8.0117478 -66.302686
C 0.6148988 9.1220086 -65.6716635
H 0.701243 9.6227611 -64.7065084
C 1.2656105 10.0741342 -66.6689175
O 2.098196 10.8648421 -66.2441968
C 1.3529267 7.7757301 -65.5926566
H 1.1126767 7.1689474 -66.4679944
H 2.4297305 7.955249 -65.5762918
C 0.9574017 7.0366377 -64.3062496
H 1.2856483 7.6316669 -63.4540904
H -0.1280695 6.9419732 -64.2625537
C 1.5717334 5.633152 -64.2128231
H 2.6407791 5.6991469 -64.4166036
H 1.1128656 4.9710833 -64.9484462
C 1.3819517 5.0770151 -62.7971826
H 1.7874912 5.8138986 -62.1066302
H 1.9538723 4.1524789 -62.6816544
N -0.0351696 4.8430507 -62.4374898
H -0.6226182 5.656999 -62.6128357
H -0.1159741 4.6848988 -61.4413045
H -0.410129 4.0257107 -62.9061462
N 0.857613 10.1083443 -67.9375912
H 0.0306158 9.5949737 -68.2147783
C 1.4627276 11.0419099 -68.8908006
H 2.5495178 10.9536548 -68.8456466
C 1.0955327 12.4971915 -68.5945729
O 1.9570158 13.368256 -68.6695282
C 1.0240044 10.7936445 -70.3246517
H -0.0383867 11.0268319 -70.4208665
H 1.5987591 11.5133366 -70.9013528
C 1.2870139 9.4334093 -70.9591885
H 0.6207275 8.6875652 -70.5204886
H 2.321262 9.1555211 -70.7495981

C 1.0670261 9.5267931 -72.4818624
O 1.5837226 8.6346019 -73.1863309
O 0.449561 10.5278113 -72.9382914
N -0.1651418 12.7909911 -68.2489103
H -0.8538143 12.0316837 -68.2624408
C -0.597046 14.1400191 -67.8500139
H -0.3606733 14.8368522 -68.6493023
C 0.1788132 14.6128788 -66.6331437
O 0.6875716 15.7306213 -66.6664212
C -2.1126775 14.1466946 -67.5941196
H -2.6227997 14.0571914 -68.5530931
H -2.3576696 13.272807 -66.9910062
C -2.6896842 15.3612165 -66.8459386
H -3.7652708 15.2087699 -66.7430743
H -2.2667664 15.3935762 -65.8416272
C -2.4521123 16.7168604 -67.5216241
H -2.901966 16.7154243 -68.5155877
H -1.3820194 16.9074539 -67.6060141
C -3.0951271 17.8042082 -66.6541248
H -2.7560098 17.6658543 -65.6223159
H -4.1813943 17.6762267 -66.6795216
N -2.71647 19.159312 -67.1134731
H -1.700728 19.2804625 -67.0277463
H -3.1123092 19.8764223 -66.5258363
H -2.9676407 19.3212056 -68.075571
N 0.3204954 13.766832 -65.6119406
H -0.1301998 12.8600258 -65.6626692
C 1.1372004 14.0988161 -64.4489276
H 0.7681291 15.0311553 -64.0186157
C 2.6010132 14.3488103 -64.8440332
O 3.1203401 15.4231067 -64.5515805
C 1.0048208 13.0052398 -63.3833035
H -0.0274297 12.9715737 -63.0338871
H 1.2416141 12.0362933 -63.8251743
C 1.9196494 13.2544928 -62.2026704
C 1.5902525 14.2330469 -61.2449909
H 0.6507624 14.7645696 -61.3091331
C 2.5054578 14.5589645 -60.2283914
H 2.2645 15.3361693 -59.5185997
C 3.7477584 13.9046379 -60.163313
H 4.4610224 14.1635952 -59.3939693
C 4.075634 12.9324819 -61.1222156
H 5.0420339 12.4538226 -61.0965527
C 3.1634757 12.6011592 -62.1355357
H 3.4384687 11.8742898 -62.8856969
N 3.2219607 13.4431795 -65.6129092

H 2.7236004 12.5935768 -65.8627977
C 4.6285742 13.5613066 -66.0272132
H 5.2190656 13.6913037 -65.136371
C 4.8554892 14.8103973 -66.9071792
O 5.7538651 15.5982238 -66.6248052
C 5.1469566 12.2423806 -66.6375441
H 4.5264298 12.0049949 -67.503892
C 6.6112232 12.3569321 -67.0848273
H 7.294004 12.1712733 -66.2581363
H 6.8394987 13.3368053 -67.4929083
H 6.78462 11.6171695 -67.8603109
C 5.0988351 11.0728698 -65.6156212
H 5.8917595 11.1882859 -64.8758281
H 4.1709873 11.0890031 -65.0529447
C 5.2309691 9.6913749 -66.2729081
H 6.195366 9.5950969 -66.7695518
H 4.4343643 9.5515472 -67.0036455
H 5.1519131 8.9179254 -65.5089451
N 3.9636351 15.1292163 -67.8520424
H 3.2265652 14.4600239 -68.0547949
C 4.0218868 16.3871815 -68.6226211
H 5.0042765 16.4596209 -69.0901891
C 3.8797531 17.6361744 -67.751329
O 4.3853571 18.6900604 -68.1393617
C 2.9423549 16.3991231 -69.7126516
H 1.9845357 16.1193492 -69.2705717
H 2.8543825 17.4158555 -70.0989016
C 3.2701847 15.4763868 -70.8965743
H 3.2834725 14.434007 -70.5792135
H 4.2600221 15.7274747 -71.2810722
C 2.2462379 15.6651832 -72.0260354
H 2.5661878 15.0769262 -72.8877034
H 2.2537104 16.7118842 -72.3344896
N 0.873631 15.2917612 -71.613203
H 0.3736673 15.9475456 -71.0440993
C 0.2606047 14.1451865 -71.8617678
N 0.8324717 13.2109762 -72.562001
H 0.4215544 12.2750905 -72.7048818
H 1.7817286 13.3170273 -72.8573687
N -0.9394563 13.9204432 -71.4047869
H -1.4254517 14.5628763 -70.815964
H -1.3235025 12.9947997 -71.5411287
N 3.1468849 17.5726836 -66.640555
H 2.7147858 16.6893069 -66.3908726
C 3.0422072 18.6861717 -65.6891141
H 2.9536692 19.6115987 -66.2544648

C 4.3172776 18.8331359 -64.8542892
O 4.818539 19.9522103 -64.7316767
C 1.7891423 18.5422215 -64.8007602
H 1.0555546 17.9044931 -65.2924164
H 2.0612306 18.0606817 -63.8602092
C 1.1195113 19.8957002 -64.5028273
H 1.8473703 20.5638183 -64.0369374
H 0.3048766 19.7346467 -63.7929669
C 0.5491297 20.5310302 -65.7790337
O -0.2282403 19.8629453 -66.4986209
O 0.9457575 21.6510811 -66.1609826
N 4.94132 17.725144 -64.4234338
H 4.4998686 16.8199543 -64.5638902
C 6.2140599 17.8019982 -63.6909489
H 6.0524835 18.5815138 -62.9562327
C 7.3890477 18.3481159 -64.518059
O 8.2732855 18.9822581 -63.9445966
C 6.5604161 16.5360761 -62.8697259
H 7.6036421 16.612572 -62.5826921
C 5.687129 16.5745079 -61.5928064
H 4.6438896 16.3754152 -61.8451636
H 5.7360653 17.5498829 -61.1123979
H 6.0256541 15.83006 -60.8757306
C 6.3611885 15.1622226 -63.5084856
H 5.298497 14.9474787 -63.5301493
H 6.74844 15.1843201 -64.5161972
C 7.0610066 14.0140301 -62.7739185
H 6.6684855 13.8981371 -61.7667884
H 8.122155 14.2216184 -62.7084251
H 6.9110273 13.0840706 -63.3228865
N 7.30301 18.2991724 -65.852446
H 6.594574 17.6978391 -66.2541671
C 8.2430857 18.9945263 -66.7521705
H 9.256973 18.7089679 -66.4899572
C 8.1690656 20.51075 -66.5818082
O 9.1990222 21.1611148 -66.460308
C 7.977772 18.6399987 -68.229651
H 6.9892592 19.0220523 -68.484297
C 8.9845327 19.2914653 -69.1874602
H 10.0014391 18.9979181 -68.9194354
H 8.9126701 20.3778295 -69.1491886
H 8.7886603 18.966429 -70.2109033
C 7.96589 17.1408008 -68.5243393
H 8.943109 16.8146886 -68.8779901
H 7.2357508 16.9438971 -69.3046957
H 7.7171778 16.551408 -67.6508645

N 6.9615765 21.0893303 -66.532496
H 6.1488449 20.486938 -66.5022801
C 6.780615 22.5474602 -66.4135413
H 7.4330732 23.0541204 -67.1286945
C 7.2202667 23.0717325 -65.0569848
O 7.912679 24.0815246 -64.9821131
C 5.3159302 22.9327156 -66.6760503
H 4.6831159 22.049465 -66.5844177
H 4.9780527 23.653042 -65.9286919
C 5.1585947 23.5769347 -68.0589042
H 5.5905341 22.9257093 -68.8152424
H 5.7275276 24.5057287 -68.0837541
C 3.6905468 23.8966561 -68.3952165
H 3.6380314 24.2920585 -69.4108383
H 3.3606373 24.6917945 -67.7218206
N 2.7656957 22.7454938 -68.2527494
H 2.0283473 22.845749 -67.5579178
C 2.88091 21.5274568 -68.7454514
N 3.8693124 21.140862 -69.4991559
H 4.0949378 20.1565639 -69.4957941
H 4.5709032 21.8214323 -69.7161739
N 1.9903342 20.6334323 -68.4586807
H 1.2955849 20.8980357 -67.7519289
H 2.1477213 19.6774308 -68.6872359
N 6.8385406 22.3850171 -63.9793491
H 6.2678414 21.5594362 -64.1188139
C 7.0762139 22.8882716 -62.6154744
H 6.7536171 23.9302454 -62.5989197
C 8.5529194 22.901266 -62.209056
O 8.8936518 23.5075295 -61.1960446
C 6.2302789 22.1490035 -61.5659052
H 6.3002834 22.7158685 -60.6375336
C 4.7406745 22.1370212 -61.9312552
H 4.5608851 21.5072413 -62.7992222
H 4.4104188 23.1524728 -62.155835
H 4.1599278 21.7513496 -61.0951958
C 6.7187884 20.740197 -61.2639461
H 6.946144 20.2365441 -62.1944734
H 5.9697426 20.1820863 -60.7022476
H 7.6299397 20.7966649 -60.6732389
N 9.4330305 22.2598839 -62.9727844
H 9.1190088 21.8290885 -63.831774
C 10.8586508 22.2157185 -62.6987607
H 11.0205601 22.3896924 -61.6385775
C 11.6587618 23.2613323 -63.4943908
O 11.250341 23.686733 -64.571033

C 11.3162717 20.8126336 -63.009183
H 11.2222436 20.673128 -64.0870845
H 10.6948542 20.0962774 -62.4706286
H 12.34619 20.6925253 -62.698318
N 12.8048385 23.6814376 -62.9622398
H 13.0708624 23.3246349 -62.0518115
C 13.7495271 24.5578181 -63.6509041
H 13.1778 25.2862108 -64.2238724
C 14.6719354 23.7794981 -64.6223379
O 14.975169 22.6149417 -64.3663321
C 14.5534374 25.3133525 -62.584314
H 15.1349368 24.6050227 -61.9900518
H 13.8784704 25.8645982 -61.9278724
H 15.2402197 26.0126728 -63.0612602
N 15.2562672 24.4346856 -65.6428965
C 16.3457623 23.8565942 -66.4329207
H 15.9645131 23.0084207 -66.9972777
C 17.5084646 23.3942187 -65.5437926
O 17.9666931 24.1420647 -64.6815619
C 16.7604415 24.9539896 -67.41675
H 17.5396302 25.5804047 -66.9808329
H 17.0916402 24.5356973 -68.368371
C 15.4800201 25.7739039 -67.5748403
H 14.8102338 25.2746548 -68.2775496
H 15.6876092 26.7949285 -67.8961341
C 14.8755827 25.7311526 -66.1737705
H 13.7917232 25.8368128 -66.2352599
H 15.3069372 26.5190763 -65.555338
N 17.9761557 22.1537945 -65.7184936
H 17.5578649 21.5750566 -66.4370437
C 18.9691881 21.5330127 -64.828799
H 19.4528712 20.7173447 -65.3653614
H 19.7268063 22.271958 -64.5676816
C 18.3945524 20.9579177 -63.5231969
O 19.1431088 20.4234825 -62.6929978
N 17.0763422 20.9995032 -63.325819
H 16.4871733 21.5126475 -63.9742723
C 16.4200515 20.282351 -62.2438022
H 16.8992307 20.5608404 -61.3042546
C 16.5430015 18.7673145 -62.4295642
O 16.5979498 18.2344241 -63.5433115
C 14.9511917 20.6876016 -62.1594518
H 14.4759675 20.4721613 -63.1160417
H 14.876111 21.7519627 -61.935609
H 14.4412873 20.1255796 -61.3775577
N 16.4551063 18.0541021 -61.3125826

H 16.2855776 18.5690471 -60.4511779
C 16.3207142 16.6066863 -61.2672556
H 16.667364 16.1436285 -62.1925943
C 14.8430527 16.4876811 -61.1714432
O 14.2703098 16.4918358 -60.0760996
C 17.1025975 16.0055349 -60.0967373
H 16.7267382 16.4254048 -59.1674263
H 16.9332383 14.927969 -60.0696969
C 18.6151246 16.2789271 -60.2035818
H 18.7934009 17.3000794 -60.5397025
H 19.0498173 15.6006874 -60.9384452
C 19.3309473 16.1098317 -58.8681021
O 19.8209138 17.063498 -58.2767724
N 19.4280626 14.9016131 -58.3528361
H 19.9160454 14.8206799 -57.47876
H 19.0269415 14.1074667 -58.8221429
N 14.2900884 16.7928196 -62.3621341
H 14.9082487 16.9578584 -63.1449686
C 12.8660165 16.8921841 -62.5786515
H 12.50313 17.8129391 -62.118628
C 12.3945187 16.8195691 -64.0583579
H 11.5182164 16.1806944 -64.1205188
C 11.927267 18.2080647 -64.4477732
H 12.7140434 18.9289267 -64.2293846
H 11.0102521 18.4103486 -63.8975376
H 11.6871596 18.2879955 -65.5028421
C 13.3973882 16.3601632 -65.1198995
H 13.1242213 16.743578 -66.1014841
H 14.3632553 16.7589904 -64.8585091
C 13.4802412 14.8541873 -65.2635705
H 12.5137973 14.4524802 -65.5470749
H 13.7913259 14.4259101 -64.3196536
H 14.1809111 14.6061911 -66.0551717
C 12.2832975 15.764129 -61.7971147
O 11.2494559 16.0014477 -61.1925983
N 12.9953431 14.6160479 -61.6922139
H 13.8487719 14.4634771 -62.2140222
C 12.4409562 13.5648977 -60.9104669
H 11.9661348 14.0429408 -60.056893
C 13.3950247 12.5287825 -60.3251963
O 14.3898634 12.1321749 -60.9337501
C 11.3641755 12.9163869 -61.7760491
H 11.69877 11.9263678 -62.0871113
C 10.1448767 12.7675339 -60.8818847
H 9.7797381 13.7482455 -60.5753726
H 10.389334 12.200937 -59.985975

H 9.3515675 12.2659959 -61.4236928
C 10.8142608 13.647437 -63.0167973
H 10.2651247 14.5505195 -62.7809551
H 10.1246692 12.9691349 -63.5158578
H 11.6027534 13.8396563 -63.7244018
N 12.992268 12.0624941 -59.1407394
H 12.2290494 12.5696896 -58.7005972
C 13.3517678 10.7991302 -58.4875107
H 13.8691189 10.1454328 -59.1901295
C 12.022295 10.1409558 -58.0722694
O 11.2124517 10.7801555 -57.3905926
C 14.2732964 11.0619765 -57.2714117
H 13.7556098 11.7467959 -56.5983282
C 14.5539489 9.7552096 -56.5040559
H 15.098085 9.0547942 -57.1393494
H 13.6255968 9.2720965 -56.1977078
H 15.1326487 9.9473315 -55.6028041
C 15.5967541 11.7351388 -57.7104154
H 16.1322448 11.0671286 -58.3834904
H 15.3704105 12.6510699 -58.2557262
C 16.5299331 12.1281646 -56.5575202
H 16.9353557 11.2398739 -56.0737671
H 15.9862613 12.7290053 -55.8275692
H 17.3598219 12.7148318 -56.9523203
N 11.7580335 8.9046503 -58.5018071
H 12.4971308 8.4049395 -59.0006782
C 10.4983895 8.1867901 -58.2471085
H 9.9675434 8.6814518 -57.4396159
C 10.7986262 6.7492106 -57.8252037
O 11.102295 5.9443846 -58.7003625
C 9.5576584 8.1569015 -59.488233
H 10.0657489 7.6118162 -60.2850846
C 9.2280089 9.5527545 -60.0531933
H 8.7489912 10.1650888 -59.2892799
H 10.1684049 10.018405 -60.3410172
C 8.2538598 7.3944316 -59.1507915
H 7.707404 7.913199 -58.3643917
H 8.4738782 6.3822778 -58.8197286
H 7.6197814 7.2880813 -60.0276344
C 8.3419087 9.5094014 -61.311817
H 7.3388524 9.1658198 -61.0701747
H 8.7868381 8.8466241 -62.055071
H 8.2449684 10.4990386 -61.7479271
N 10.3778145 6.3647875 -56.6173089
H 10.0867171 7.0875804 -55.969359
C 10.0771669 4.965996 -56.2829504

H 10.6161371 4.3342597 -56.978693
C 8.5735844 4.707958 -56.4325046
O 7.7547345 5.5273144 -56.0055968
C 10.6004787 4.5910992 -54.8968489
H 10.1061673 5.1945444 -54.1366728
H 11.6735419 4.7889482 -54.8607758
O 10.3828632 3.2140938 -54.6418497
H 9.4472056 3.0366627 -54.4338089
N 8.1607001 3.6309068 -57.1055167
H 8.8769937 2.9804449 -57.4273788
C 6.7371884 3.300688 -57.3620578
H 6.1596131 3.5083897 -56.4621738
C 6.5491433 1.8064488 -57.6597888
O 7.4787363 1.1227106 -58.0834072
C 6.1822375 4.201798 -58.4902398
H 6.9381585 4.3099942 -59.2694911
C 4.8723056 3.766805 -59.1432156
H 4.1171477 3.6139869 -58.3752925
H 5.0235349 2.8460121 -59.7068042
H 4.5357146 4.5394803 -59.8340945
O 5.8902408 5.4700017 -57.9372644
H 6.6083844 5.6589023 -57.3062582
N 5.3715908 1.2506011 -57.3572631
H 4.6553821 1.8125038 -56.8983762
C 5.0533527 -0.1448405 -57.6414244
H 5.9175855 -0.7539901 -57.3692877
C 4.7686066 -0.3651727 -59.1348527
O 3.9382459 0.3149866 -59.7426248
C 3.8790949 -0.6020956 -56.7719286
H 2.9852171 -0.0491227 -57.0590648
H 4.0992285 -0.4156114 -55.7198245
H 3.7121333 -1.6687299 -56.9224242
N 5.384449 -1.4132231 -59.6710028
H 5.9615485 -1.978817 -59.0585404
C 5.274073 -1.8707675 -61.0481607
H 4.52063 -1.2819514 -61.5709594
C 4.819635 -3.3368143 -61.111532
O 4.71484 -4.0327033 -60.0985856
C 6.6290324 -1.6468094 -61.7413333
H 6.7247861 -2.2962304 -62.6122492
H 7.4480343 -1.8571985 -61.0513851
S 6.7161264 0.0744788 -62.2988248
H 5.7073454 -0.0156779 -63.1767116
N 4.5804398 -3.8163232 -62.3306187
H 4.6770828 -3.1691214 -63.1064769
C 4.346574 -5.2294842 -62.6551505

H 4.459447 -5.8264837 -61.7510928
C 5.4032849 -5.7429896 -63.6329606
O 6.1897972 -4.9572822 -64.1649364
C 2.9059161 -5.4309908 -63.1662228
H 2.7672556 -6.4849372 -63.4143421
H 2.2061259 -5.1920836 -62.3647088
C 2.5629917 -4.5759187 -64.3962885
H 3.3538419 -4.6507815 -65.1381943
H 2.4774119 -3.5317996 -64.1082129
C 1.239358 -5.0199314 -65.0200982
H 0.4409992 -4.8935195 -64.2851521
H 1.2911458 -6.0804497 -65.2787751
N 0.9258565 -4.2126199 -66.212061
H 0.4081314 -3.3590145 -66.0351975
C 1.2259418 -4.5210722 -67.4631425
N 1.9688723 -5.5468659 -67.7776155
H 2.1152903 -5.8721481 -68.7219343
H 2.3570204 -6.1389378 -67.0480665
N 0.7801215 -3.779182 -68.4391839
H 0.1596895 -3.0162654 -68.2328983
H 0.9754474 -4.0537471 -69.3841574
N 5.3899017 -7.0384504 -63.929528
H 4.7383203 -7.6344439 -63.4363782
C 6.0942102 -7.5596355 -65.1036889
H 7.0977374 -7.1300138 -65.1006469
C 5.4232967 -7.0925274 -66.4226341
O 4.3354509 -6.5106139 -66.4170726
C 6.2677109 -9.0838745 -64.9594063
H 6.6936781 -9.2945219 -63.9792089
H 6.9785677 -9.4402086 -65.7029615
C 4.9912565 -9.885612 -65.1365704
O 4.2025192 -9.6370408 -66.0222303
N 4.7808317 -10.9165378 -64.3629691
H 3.901636 -11.390675 -64.4666077
H 5.4341878 -11.1196392 -63.6117621
N 6.087026 -7.301176 -67.5654737
H 6.97492 -7.7768733 -67.5279906
C 5.5686416 -6.8960675 -68.8824775
H 5.3268371 -5.8354456 -68.847767
C 4.2717675 -7.636028 -69.2757134
O 3.3960169 -7.0361793 -69.9063768
C 6.6966829 -7.1057223 -69.9118019
H 7.5465829 -6.4850685 -69.622889
H 7.0144693 -8.1489414 -69.8602459
C 6.3428838 -6.7968035 -71.3816833
H 5.5274226 -7.4439008 -71.7033055

C 5.9359276 -5.3387014 -71.5982807
H 6.7312629 -4.6712647 -71.263949
H 5.0185384 -5.1252485 -71.0539862
H 5.7487945 -5.1681951 -72.6580208
C 7.5553389 -7.0841982 -72.2684394
H 8.3866897 -6.4341584 -71.9947723
H 7.2949337 -6.9136733 -73.313153
H 7.8578121 -8.1251123 -72.1530906
N 4.1334798 -8.8979656 -68.8636108
H 4.8250579 -9.2749777 -68.2335885
C 3.0282568 -9.7990911 -69.2066128
H 2.994302 -9.8630138 -70.2915907
C 1.6602333 -9.3119172 -68.6927965
O 1.5945623 -8.516802 -67.7502102
C 3.3072741 -11.2092038 -68.6610331
H 2.5364612 -11.8929918 -69.0166306
H 3.2739869 -11.2045757 -67.5732222
O 4.5717364 -11.6737482 -69.1032695
H 4.7435367 -12.5260357 -68.6915725
N 0.5433211 -9.7783737 -69.2905212
C -0.7985269 -9.4881579 -68.7920022
H -0.974344 -8.4156383 -68.878059
C -0.9754439 -9.9212677 -67.3331253
O -0.527133 -10.9988821 -66.9390965
C -1.7659872 -10.2366409 -69.7171588
H -1.976375 -11.2318292 -69.3196156
H -2.693753 -9.6827823 -69.8650191
C 0.4554123 -10.5666366 -70.5124142
H 1.1583618 -10.2258453 -71.2721388
H 0.6242898 -11.6193873 -70.280253
C -0.973233 -10.3730318 -71.0152853
H -1.0393598 -9.4455678 -71.586019
H -1.3115491 -11.2200922 -71.6130074
N -1.6717958 -9.0991233 -66.547554
H -1.9774122 -8.2113845 -66.9104634
C -1.9426282 -9.3712957 -65.1363939
H -0.9869589 -9.4906116 -64.6280691
C -2.7599268 -10.6578647 -64.9656591
O -3.8400503 -10.7898855 -65.5414905
C -2.701309 -8.1901542 -64.5041285
H -2.7718945 -8.3380187 -63.4261289
H -3.7101012 -8.1606276 -64.9151862
O -2.0772418 -6.945159 -64.7763668
H -2.4140318 -6.2669389 -64.1668781
N -2.292544 -11.5662814 -64.1043462
H -1.3627723 -11.44079 -63.7365719

C -3.0207712 -12.7921241 -63.7457168
H -3.2244875 -13.323451 -64.6781217
C -4.3966937 -12.4684447 -63.1343671
O -4.6230048 -11.3825377 -62.579092
C -2.1096583 -13.72106 -62.9038127
H -1.910148 -14.6234283 -63.4834728
H -1.1418491 -13.2284864 -62.8004951
C -2.5148567 -14.1252411 -61.4840284
H -2.7946617 -13.2267165 -60.9280213
H -1.6416138 -14.553518 -60.9864657
C -3.6505368 -15.1492933 -61.4302847
O -4.2452488 -15.22482 -60.332582
O -4.0275463 -15.7087837 -62.4791995
N -5.3275999 -13.4242844 -63.2208885
H -5.0081732 -14.3620026 -63.4746751
C -6.7373853 -13.2558538 -62.8708644
H -7.1275333 -12.4809128 -63.5271843
C -6.946757 -12.7679238 -61.4410492
O -7.9117529 -12.0322895 -61.2615722
C -7.4812086 -14.5788163 -63.1367407
H -7.2969487 -14.8900304 -64.1668939
H -7.0568127 -15.3459934 -62.4846586
C -8.9995942 -14.5392484 -62.9048429
H -9.3885239 -15.5543231 -62.9959171
H -9.1947796 -14.2219061 -61.8805407
C -9.7553918 -13.6445641 -63.9063237
H -10.1326223 -14.2766743 -64.7127237
H -9.0952164 -12.9031723 -64.3565096
C -10.9343519 -12.935928 -63.2296966
H -11.4774207 -13.6713244 -62.6279936
H -11.6147183 -12.563259 -64.0023007
N -10.4659423 -11.8111498 -62.3804404
H -9.7675482 -12.1340372 -61.710333
H -11.2343898 -11.3862407 -61.8810379
H -10.0067443 -11.1094506 -62.9439105
N -6.0816248 -13.0886981 -60.4718269
H -5.3969747 -13.8110332 -60.6825751
C -6.0968226 -12.5397581 -59.1017552
H -7.1172031 -12.2357542 -58.8791371
C -5.2212816 -11.2922417 -58.8815129
O -5.4720267 -10.5585664 -57.9262851
C -5.7549487 -13.6167123 -58.0640669
H -6.3952429 -14.4869167 -58.2181221
H -5.9482444 -13.2182487 -57.0671658
O -4.3992676 -14.0098058 -58.1432093
H -4.343102 -14.5866227 -58.960205

N -4.2784466 -10.9636136 -59.7796263
H -4.191226 -11.5356199 -60.6116894
C -3.5645761 -9.675878 -59.7459851
H -3.2982635 -9.4612025 -58.710529
C -4.4764878 -8.5391038 -60.2170946
O -4.6424323 -7.5619692 -59.492832
C -2.2637066 -9.7208699 -60.5782299
H -1.5860132 -10.4576694 -60.1459091
H -2.5060962 -10.0411425 -61.5913691
C -1.54592 -8.348949 -60.6594275
H -2.2252017 -7.6004366 -61.0640231
C -1.049715 -7.869789 -59.2926931
H -0.3925492 -8.6162577 -58.8481829
H -1.895953 -7.6778135 -58.6345231
H -0.5004847 -6.9352805 -59.4146506
C -0.3461376 -8.4080241 -61.5969519
H 0.3865714 -9.1103831 -61.226032
H 0.1108929 -7.4212003 -61.6794127
H -0.6523785 -8.7230191 -62.5882788
N -5.1169134 -8.689943 -61.3849318
H -4.9457027 -9.5459942 -61.90868
C -6.0412299 -7.6860189 -61.9416269
H -5.4551489 -6.7857532 -62.1437637
C -7.1240799 -7.231914 -60.9375162
O -7.2297424 -6.0346225 -60.7273625
C -6.5934143 -8.1928127 -63.2907024
H -5.7661369 -8.2380254 -63.9984981
H -6.9541082 -9.2147834 -63.16253
C -7.7329938 -7.3712305 -63.9310273
H -7.8971692 -7.7500898 -64.9408023
H -8.6442331 -7.5659483 -63.3636415
C -7.5575872 -5.8442729 -64.0094301
H -7.4980538 -5.4352241 -63.0077227
H -8.4615509 -5.4163366 -64.4482493
C -6.3614685 -5.3390559 -64.8205827
H -6.662531 -5.2804452 -65.8695286
H -5.5195257 -6.0309988 -64.7300999
N -5.9571078 -4.0034871 -64.3263948
H -6.7679802 -3.395136 -64.17414
H -5.2630821 -3.5587778 -64.9038722
H -5.4837897 -4.0957187 -63.4295262
N -7.871645 -8.0931999 -60.2239082
C -8.873936 -7.6616363 -59.2573345
H -9.5234761 -6.9139405 -59.7159941
C -8.2649088 -7.0320422 -57.9992117
O -8.9337415 -6.198365 -57.3998078

C -9.7099499 -8.8997193 -58.9176722
H -10.0869565 -8.8805224 -57.8938592
H -10.5345133 -8.9914458 -59.6264198
C -7.9351771 -9.5147877 -60.3439147
H -8.4774898 -9.7379089 -61.262144
H -6.9490279 -9.9651637 -60.3563997
C -8.720291 -10.0246495 -59.1344312
H -9.2281338 -10.9685741 -59.3297095
H -8.0659944 -10.10908 -58.2666461
N -7.021786 -7.3606245 -57.6043131
H -6.4998668 -8.0322734 -58.1535313
C -6.3056253 -6.6353359 -56.5348897
H -6.9728801 -6.5111878 -55.6824392
C -5.9322881 -5.2270172 -57.0036012
O -6.1602659 -4.2700824 -56.2681912
C -5.0687729 -7.4390979 -56.0860549
H -5.3895788 -8.424192 -55.7418211
H -4.4305065 -7.5948128 -56.9556649
C -4.1992388 -6.7757217 -54.993123
H -3.2115868 -7.2341931 -55.0481208
H -4.0668317 -5.7140503 -55.2046993
C -4.7169933 -6.9550987 -53.5545555
H -4.8133604 -8.0239603 -53.351868
H -5.6999526 -6.4874112 -53.4551922
C -3.7314371 -6.3211879 -52.5553124
H -3.9467867 -5.2535484 -52.4822222
H -2.7197203 -6.4281609 -52.9491867
N -3.7880118 -6.9492504 -51.2117734
H -4.7008678 -6.8646728 -50.7950431
H -3.1294465 -6.5095671 -50.5629086
H -3.5314121 -7.9249485 -51.2652042
N -5.4101965 -5.0928218 -58.2204721
H -5.2513331 -5.9347831 -58.7634563
C -5.1139113 -3.8079675 -58.8626221
H -4.4172266 -3.2441876 -58.2414818
C -6.3827766 -2.9486339 -58.9977917
O -6.424878 -1.8461333 -58.4601641
C -4.4306352 -4.0925634 -60.2147685
H -3.4889227 -4.612916 -60.0251308
H -5.0681263 -4.750292 -60.8032861
C -4.1402352 -2.8329003 -61.0402102
H -3.5223312 -2.1630568 -60.4391233
H -5.0881717 -2.3339607 -61.2560169
C -3.4537445 -3.1310734 -62.3813936
O -3.6879035 -4.2298107 -62.9487496
O -2.7668995 -2.224337 -62.9038172

N -7.4664462 -3.5002571 -59.5499567
H -7.3734285 -4.4240226 -59.9575726
C -8.7116426 -2.7639545 -59.7964255
H -8.4353106 -1.8095062 -60.2480485
C -9.4638687 -2.4089424 -58.5074308
O -9.9528278 -1.288221 -58.3755535
C -9.6118555 -3.5466591 -60.7821039
H -9.0607949 -4.3693492 -61.2330581
H -10.4668038 -3.9705644 -60.2532929
C -10.1111674 -2.6392604 -61.9230753
H -10.7214885 -1.8416841 -61.4932013
H -10.7558622 -3.2309231 -62.5768496
C -8.9667387 -2.0313695 -62.7636765
O -8.2371866 -2.7994259 -63.4391515
O -8.8066446 -0.7847794 -62.7441526
N -9.47042 -3.2962327 -57.4999447
H -9.0417097 -4.20374 -57.6524768
C -10.1010431 -3.0170332 -56.1973756
H -11.0288266 -2.4764864 -56.3973745
C -9.2769901 -2.0761537 -55.3142795
O -9.8498703 -1.4157787 -54.4554675
C -10.4660358 -4.334596 -55.4944689
H -11.0058283 -4.9560219 -56.2112972
H -9.5566922 -4.8544857 -55.1890845
C -11.3704398 -4.1053174 -54.2696322
H -10.7558442 -3.807233 -53.4188719
H -12.0692675 -3.2953097 -54.4855386
C -12.2007088 -5.3376816 -53.8874421
H -12.8839548 -5.5640901 -54.7082912
H -12.7938671 -5.0865768 -53.0059178
C -11.3301939 -6.5636396 -53.5898349
H -10.6293076 -6.3072624 -52.7892185
H -10.7480368 -6.8020651 -54.4859351
N -12.1634946 -7.726805 -53.1936491
H -12.701381 -7.5095402 -52.3633894
H -11.5843428 -8.534113 -52.9993255
H -12.8073944 -7.9650457 -53.9371857
N -7.9716141 -1.9444161 -55.5661454
H -7.5758423 -2.5040646 -56.3110106
C -7.1488087 -0.8533673 -55.0040867
H -7.6222074 -0.5067725 -54.0856447
C -7.1064176 0.3871534 -55.9041949
O -6.4532318 1.363562 -55.5514579
C -5.7327836 -1.3132419 -54.6059369
H -5.1704231 -0.4493004 -54.2512706
C -5.7686133 -2.3392788 -53.4714731

H -6.3200593 -3.2274988 -53.7777969
H -6.2540956 -1.8980821 -52.6008782
H -4.7497511 -2.6174525 -53.2046511
O -5.0121386 -1.9015539 -55.6648057
H -5.5081724 -2.690785 -55.9458878
N -7.8166632 0.3654646 -57.0417343
H -8.3462446 -0.4726061 -57.2427193
C -7.8502309 1.3899778 -58.0923619
H -8.3461036 0.901136 -58.9292334
C -6.4594769 1.7892235 -58.633805
O -6.3001095 2.8344891 -59.2640218
C -8.7526115 2.5574542 -57.6322694
H -9.6187161 2.1305136 -57.1235023
H -8.1999693 3.1478449 -56.8997247
C -9.2803135 3.5032468 -58.7320415
H -8.4579137 4.0436594 -59.1952806
C -10.0659324 2.771923 -59.8264709
H -10.8596408 2.1741863 -59.3761475
H -9.4021933 2.1159166 -60.3858728
H -10.5024011 3.4948423 -60.5147045
C -10.2132689 4.5376623 -58.0986059
H -11.0746174 4.0426197 -57.649767
H -10.5543428 5.2422816 -58.85601
H -9.6730952 5.0878871 -57.3275553
N -5.4426723 0.9434196 -58.4481881
H -5.6678236 0.0355238 -58.0548346
C -4.0798935 1.185912 -58.9357536
H -3.8150141 2.2211407 -58.7113507
C -4.0051982 1.032849 -60.4518383
O -4.7678273 0.27695 -61.0434649
C -3.0949649 0.2568287 -58.2130025
H -3.5138127 -0.7490093 -58.1422002
H -2.1678002 0.2159683 -58.784929
C -2.795052 0.8141 -56.8097441
H -2.3119699 1.7866437 -56.9208895
H -3.7301621 0.9656837 -56.2690902
C -1.8893985 -0.0773955 -55.9502131
H -2.4545609 -0.9480272 -55.6154601
H -1.5950424 0.5033683 -55.0736083
C -0.6439782 -0.5463608 -56.7108241
H -0.3018319 0.268945 -57.3493463
H -0.9215663 -1.3830817 -57.3566641
N 0.4467047 -0.9339892 -55.7916971
H 0.7465728 -0.112789 -55.2498367
H 1.2748543 -1.2105249 -56.2979961
H 0.1611309 -1.6486113 -55.1440047

N -3.0487572 1.7262357 -61.0774219
H -2.4915023 2.3509615 -60.5015827
C -2.7384924 1.6213578 -62.5167704
H -3.2764944 0.767828 -62.9341887
C -1.2710996 1.2723612 -62.7035448
O -0.4066791 2.1504201 -62.6343866
C -3.1835379 2.9095297 -63.2545982
H -3.4195988 3.6894009 -62.5300726
H -2.3764105 3.2816492 -63.8842477
C -4.4157079 2.6815394 -64.1627795
H -4.7723658 3.6569256 -64.4963477
H -4.1117891 2.1108096 -65.0407656
C -5.5713926 1.9356428 -63.4710507
H -5.670715 2.3288555 -62.460506
H -5.3424209 0.8725846 -63.4125411
C -6.9189723 2.0674559 -64.1837241
H -6.9312356 1.427326 -65.0694601
H -7.0421595 3.108668 -64.4916204
N -8.0097774 1.7033435 -63.2510456
H -7.9605382 0.719054 -62.960874
H -8.919042 1.7965361 -63.6756161
H -7.9574248 2.2846703 -62.4272009
N -0.9851324 -0.0152724 -62.8685472
H -1.7507671 -0.6921228 -62.9343551
C 0.3622951 -0.5630426 -62.7563147
H 0.9805693 0.166144 -62.2307181
C 0.9623679 -0.7635467 -64.1569443
O 0.4536773 -1.5034089 -65.0035394
C 0.3662204 -1.8384641 -61.8795023
H -0.0630003 -2.6610733 -62.4548948
C 1.819163 -2.1735927 -61.5191769
H 2.2062048 -1.4454116 -60.8056847
H 2.4410637 -2.152216 -62.4072739
H 1.8767011 -3.1699076 -61.08193
C -0.4305112 -1.6738873 -60.5577608
H 0.0351023 -0.9039866 -59.9410057
H -1.442414 -1.3440835 -60.7786126
C -0.5557201 -2.9669112 -59.7395202
H 0.4082644 -3.2467547 -59.3150915
H -0.9233073 -3.7726035 -60.3764866
H -1.265722 -2.8133331 -58.927048
N 2.0820627 -0.0851478 -64.4029156
H 2.4655832 0.475942 -63.6575331
C 2.9068083 -0.2770199 -65.5962123
H 2.3018658 -0.6952979 -66.4020926
C 4.0396154 -1.2643317 -65.292497

O 4.4307098 -1.4269569 -64.1332412
C 3.4495725 1.0814523 -66.061202
H 4.0958151 0.9525892 -66.9322466
H 4.0432311 1.5292122 -65.261646
S 2.0709612 2.1846743 -66.4876026
H 1.3003227 1.8921115 -65.4342484
N 4.6448883 -1.8495044 -66.3268338
H 4.3466771 -1.6176306 -67.261478
C 5.9932657 -2.3856146 -66.1595621
H 6.0491473 -2.8706147 -65.1886549
C 7.0088041 -1.2347704 -66.1249314
O 6.737833 -0.132612 -66.6086652
C 6.3073177 -3.4524623 -67.2163512
H 5.4957053 -4.1766962 -67.2517385
H 7.2191217 -3.9815768 -66.9375472
C 6.5230252 -2.8364934 -68.5773457
O 7.5678347 -2.280934 -68.8635157
N 5.5086591 -2.8222407 -69.403895
H 5.6807276 -2.3748843 -70.3032807
H 4.6772054 -3.3460722 -69.2191437
N 8.1801732 -1.5131329 -65.5618081
H 8.3470506 -2.4508453 -65.2320084
C 9.2452561 -0.5331003 -65.3748175
H 8.8589516 0.286887 -64.7647094
C 9.6957767 0.098359 -66.7072817
O 9.7789872 1.3222783 -66.8014925
C 10.3873618 -1.2029495 -64.5780293
H 11.2574185 -0.549599 -64.5671677
H 10.0590078 -1.2929866 -63.5414857
C 10.8177317 -2.5865022 -65.0490904
C 11.9181375 -2.7365552 -65.9169673
H 12.4819757 -1.8755661 -66.2390297
C 12.3053206 -4.0136478 -66.363435
H 13.1452796 -4.1187607 -67.0335377
C 11.597087 -5.1547648 -65.9319127
O 11.9592972 -6.3881262 -66.3699191
H 12.7274721 -6.3595775 -66.9421564
C 10.5129368 -5.0118722 -65.040923
H 9.9908157 -5.8930459 -64.6965091
C 10.1282528 -3.7322316 -64.5973171
H 9.308215 -3.6379075 -63.8983316
N 9.914116 -0.6953223 -67.7660937
H 9.7185 -1.6813804 -67.6780474
C 10.4258322 -0.1969978 -69.0526211
H 11.3531195 0.3334907 -68.8684677
C 9.4744436 0.8147866 -69.7068052

O 9.9050969 1.903903 -70.0918449
C 10.7555701 -1.3866897 -69.9704476
H 11.4799712 -2.0264374 -69.4643361
H 9.8468928 -1.968167 -70.1290859
C 11.319076 -0.9912917 -71.3499645
H 10.5886703 -0.3825456 -71.8825983
C 12.633967 -0.2142643 -71.2488822
H 13.3625122 -0.779567 -70.6682336
H 12.462991 0.7532524 -70.7772734
H 13.0319279 -0.0350601 -72.2476385
C 11.5735498 -2.2547509 -72.1719436
H 12.3160219 -2.8816896 -71.6783682
H 11.9344282 -1.9816748 -73.1636924
H 10.6437644 -2.8131246 -72.2827093
N 8.1721243 0.5219023 -69.7300654
H 7.8781855 -0.3931025 -69.4008782
C 7.1627271 1.4694778 -70.216607
H 7.465206 1.8258281 -71.2019721
C 7.0485684 2.7218684 -69.344085
O 6.7977963 3.7974389 -69.8845881
C 5.8037654 0.7721156 -70.3591403
H 5.0142317 1.5249066 -70.3314356
H 5.6424419 0.0910384 -69.5229777
C 5.6722935 0.0300481 -71.6647418
N 5.9532087 -1.3209948 -71.8725283
C 5.7499517 -1.545272 -73.178947
H 5.897551 -2.4993019 -73.6695688
N 5.3500598 -0.4176791 -73.790252
H 5.1471925 -0.3273254 -74.7758818
C 5.2980715 0.5887433 -72.851667
H 5.0341347 1.6258243 -73.0140616
N 7.3164396 2.6332608 -68.0360767
H 7.5599259 1.7314697 -67.6420576
C 7.2614763 3.7968527 -67.1509479
H 6.3620564 4.3393293 -67.4291629
C 8.4261241 4.7738874 -67.3880553
O 8.1695259 5.9696373 -67.5199989
C 7.0948289 3.3363918 -65.6891448
H 6.3288467 2.5601209 -65.6683109
H 8.0271337 2.8928056 -65.3390609
C 6.6681347 4.4549309 -64.7117129
H 7.5189438 5.109165 -64.5258907
C 5.4997377 5.3156355 -65.2074584
H 4.6757884 4.679795 -65.5306189
H 5.8301918 5.9439793 -66.0348046
H 5.1677863 5.9791257 -64.4099764

C 6.2145039 3.8387536 -63.3851094
H 5.3016056 3.2618588 -63.5260214
H 6.0445249 4.6237662 -62.6487135
H 6.9882306 3.1722436 -63.0041928
N 9.6480168 4.2850584 -67.6533181
H 9.7997559 3.2846604 -67.5630426
C 10.7262591 5.1507473 -68.1801541
H 10.8557976 5.9933419 -67.499146
C 10.3952471 5.6900104 -69.5631661
O 10.7418516 6.824877 -69.8713061
C 12.0605472 4.4240518 -68.3016877
H 12.1166467 3.8610945 -69.2350101
H 12.121121 3.7293563 -67.4803064
O 13.1491082 5.3247477 -68.1948499
H 13.269569 5.8510611 -69.0056855
N 9.6793356 4.9112594 -70.3797769
H 9.4739327 3.9631262 -70.084374
C 9.1529553 5.3651188 -71.6612905
H 9.9800387 5.6169965 -72.3253587
H 8.5721268 4.5597261 -72.1096179
C 8.2436502 6.5889815 -71.5206075
O 8.4964877 7.5881455 -72.1829673
N 7.271716 6.5802667 -70.6017955
H 7.1008963 5.7179054 -70.0934582
C 6.3753098 7.7248641 -70.3564864
H 5.913338 8.0175862 -71.2986071
C 7.1064366 8.967594 -69.8279953
O 6.7901523 10.0960769 -70.2162708
C 5.2706065 7.3106618 -69.3736671
H 5.7289851 6.9026424 -68.4718638
H 4.7203521 8.2063148 -69.0832355
C 4.2527546 6.3259864 -69.9274463
C 3.9569052 5.1377987 -69.2312677
H 4.470828 4.90886 -68.3126257
C 2.9868958 4.2463839 -69.7247382
H 2.7656118 3.3367646 -69.1875547
C 2.2932897 4.5473782 -70.9100298
H 1.5351606 3.8746796 -71.2849157
C 2.5749518 5.7375725 -71.6014384
H 2.0297721 5.995317 -72.5016579
C 3.5527959 6.6201892 -71.1146242
H 3.7383022 7.5375435 -71.6559592
N 8.1333138 8.7500029 -69.0098282
H 8.2811018 7.8069615 -68.6678179
C 9.07562 9.7716512 -68.5730746
H 8.5547353 10.5922679 -68.0912188

C 9.8605926 10.3853852 -69.7513712
O 9.7566054 11.5847949 -70.0065046
C 9.9597189 9.1076336 -67.5219712
H 10.8339721 9.7058895 -67.3759088
H 10.2895408 8.1279684 -67.8552867
S 9.0625811 8.9471298 -65.9503501
H 9.8351265 9.8474364 -65.3019252
N 10.4805271 9.5504548 -70.5898632
H 10.5030738 8.5688311 -70.3340113
C 11.2121939 9.9579073 -71.8004392
H 12.0155281 10.6371849 -71.5108146
C 10.3189662 10.6842638 -72.8220458
O 10.6706195 11.7496569 -73.3218699
C 11.8287769 8.7012708 -72.4308687
H 11.0787146 8.1829872 -73.0301747
H 12.1581597 8.0264929 -71.641054
O 12.9535704 9.016732 -73.2287619
H 13.2151123 8.2260534 -73.7115009
N 9.0917743 10.1931924 -73.0341901
H 8.8462209 9.3288767 -72.56349
C 8.0758771 10.7997598 -73.908273
H 8.5230395 10.9933282 -74.8827677
C 7.5601152 12.1572996 -73.4059811
O 6.9709603 12.9057875 -74.1813275
C 6.8980951 9.8177845 -74.0665242
H 6.5963806 9.4797408 -73.0746319
H 6.0469815 10.3468555 -74.4990079
C 7.2010356 8.596359 -74.9559661
H 8.1359891 8.1311667 -74.6495739
C 6.078985 7.5667695 -74.8172612
H 5.1265064 8.0041828 -75.1217947
H 6.002105 7.244648 -73.7790986
H 6.2918725 6.6974123 -75.4373065
C 7.3195826 8.9722824 -76.4354788
H 6.4050387 9.4644876 -76.7684964
H 7.4802548 8.0744735 -77.031481
H 8.1643764 9.6422728 -76.586479
N 7.8083199 12.5056056 -72.1428918
H 8.3267403 11.8644839 -71.5540116
C 7.4700463 13.8201725 -71.5844624
H 6.6249218 14.2248713 -72.140381
C 8.5965015 14.8537852 -71.7283394
O 8.4412966 15.9715581 -71.2569983
C 7.0096966 13.6702722 -70.1392581
H 6.7278152 14.646009 -69.7435907
H 7.8241777 13.2648047 -69.5378654

O 5.8818799 12.8084382 -70.0899695
H 6.1934362 11.8939971 -70.2199382
N 9.6527789 14.5194905 -72.4726668
H 9.6609405 13.583641 -72.8564542
C 10.8450282 15.3307806 -72.7823003
H 11.3215018 14.7681748 -73.5857787
C 11.9223128 15.3449434 -71.6844862
O 12.7208264 16.2703293 -71.5460031
C 10.4885248 16.6983789 -73.4123986
H 10.3052822 17.4216129 -72.6154718
H 9.5630553 16.5886969 -73.9810148
C 11.5483248 17.2610639 -74.3780375
O 12.3958398 16.4764423 -74.8659835
O 11.4371665 18.4657644 -74.7049311
N 11.9834183 14.26542 -70.8998863
H 11.310854 13.5258562 -71.0438426
C 13.0028759 14.0912496 -69.8714545
H 13.306885 15.0728525 -69.5156226
C 14.2829326 13.4339092 -70.3807713
O 14.2769208 12.3465484 -70.963321
C 12.4106606 13.3349922 -68.6923727
H 12.0504773 12.3665639 -69.0383334
H 13.215576 13.1470367 -67.9823596
C 11.3089208 14.0511588 -67.9372688
C 10.4143586 13.2895275 -67.1652494
H 10.4207411 12.2129036 -67.2452246
C 9.622269 13.9020251 -66.183757
H 9.0420531 13.2912381 -65.5123696
C 9.6896914 15.2939169 -66.0117228
O 9.0688287 15.8810816 -64.9659394
H 8.6517885 15.2197967 -64.4135997
C 10.476267 16.0688853 -66.8777947
H 10.5008894 17.1334103 -66.7732363
C 11.2869235 15.4568867 -67.8389113
H 11.9309688 16.084664 -68.4366982
N 15.413457 14.0666489 -70.0648519
H 15.3470006 14.8954175 -69.4865486
C 16.7466287 13.6072225 -70.4479379
H 16.6439033 13.0148765 -71.3578561
C 17.3845581 12.7089696 -69.3861672
O 17.0498046 12.785335 -68.2019845
C 17.6340751 14.8021464 -70.8021027
H 17.0888067 15.4656922 -71.4759625
H 18.5250258 14.4458703 -71.3186039
O 18.0387931 15.5263655 -69.6592734
H 17.3011331 15.5682376 -69.0191956

N 18.3085085 11.8526167 -69.839663
H 18.540374 11.8840546 -70.8182419
C 19.0346307 10.8940314 -69.0057114
H 19.4675616 10.1472293 -69.6713046
C 18.1121144 10.1118048 -68.0476025
O 18.3664568 10.0593789 -66.8463913
C 20.2060694 11.6112569 -68.3167241
H 19.8233584 12.3411857 -67.6012615
H 20.822286 12.1241624 -69.055329
H 20.8190484 10.8836954 -67.7848943
N 17.0082147 9.5547067 -68.5680509
H 16.8205954 9.6340163 -69.5538031
C 16.1258953 8.7216485 -67.7571872
H 16.0455573 9.2076642 -66.7871337
C 16.7726785 7.3640399 -67.4612841
O 17.037384 6.5613184 -68.3551552
C 14.6792575 8.6592675 -68.293561
H 14.2122908 9.6269136 -68.1101076
H 14.1203773 7.931987 -67.704949
C 14.479784 8.3334479 -69.7606206
O 14.1690558 7.2130685 -70.1375173
N 14.5042034 9.3331716 -70.6191546
H 14.1908225 9.1433714 -71.5603483
H 14.5384129 10.2922772 -70.2963576
N 17.0392665 7.1424113 -66.177116
H 16.830673 7.8928146 -65.5251381
C 17.5157217 5.8978433 -65.5954028
H 17.9177789 5.2616605 -66.384763
C 16.3357946 5.207524 -64.9288189
O 15.6159122 5.8225069 -64.1466295
C 18.6212937 6.1737034 -64.5672921
H 18.1711266 6.6156535 -63.6769974
H 19.0702786 5.2249355 -64.2716113
C 19.7093446 7.0913046 -65.0380035
C 19.7931824 8.4044157 -64.7311366
H 19.0720161 8.9404149 -64.124824
N 20.8504143 8.9783652 -65.4059943
H 20.9953366 9.9773354 -65.4322483
C 21.4912532 8.0630289 -66.2111008
C 22.5607112 8.1678796 -67.1126483
H 23.0547657 9.1163029 -67.2628505
C 22.976687 7.0234162 -67.8144749
H 23.7994497 7.0845023 -68.5131482
C 22.3164877 5.7977862 -67.6094385
H 22.6337933 4.9207785 -68.1562422
C 21.2368826 5.7069224 -66.707767

H 20.7248034 4.766601 -66.5741431
C 20.7952953 6.8365602 -65.9837824
N 16.1848563 3.912817 -65.176291
H 16.821799 3.469261 -65.8205527
C 15.0796393 3.1104436 -64.6775838
H 14.54308 3.6333506 -63.8833365
C 15.6247004 1.7985474 -64.1325112
O 16.0577958 0.9402323 -64.9020713
C 14.131694 2.8887033 -65.8487103
H 13.64995 3.8395378 -66.0623428
H 14.7289034 2.6128478 -66.7202805
C 13.0901348 1.8002141 -65.5654381
H 13.4487714 1.0489044 -64.8679001
C 11.8698551 2.4330043 -64.9143906
H 11.3898081 3.1481945 -65.575671
H 12.1699034 2.9421886 -63.9979327
H 11.1508884 1.6662489 -64.6483251
C 12.8433832 1.0456084 -66.8611406
H 12.3756177 1.6885177 -67.5980209
H 12.2143535 0.1981272 -66.6495824
H 13.7858171 0.6635186 -67.2549475
N 15.5022455 1.6123315 -62.8227831
H 15.0176951 2.3212374 -62.2749974
C 15.9819101 0.4156113 -62.1354336
H 16.5767406 -0.1688088 -62.8309248
C 14.7948239 -0.412656 -61.638592
O 14.0877462 0.0314062 -60.7298387
C 16.9228566 0.7713672 -60.9737509
H 16.3487296 1.1324943 -60.119959
C 17.7625409 -0.435787 -60.5611816
H 18.3692173 -0.7740933 -61.4018301
H 17.1180753 -1.2500821 -60.2345302
H 18.4224701 -0.1512396 -59.7425426
O 17.83844 1.7718527 -61.3585353
H 17.3314862 2.5714806 -61.5236629
N 14.5757734 -1.6381667 -62.1522123
C 13.7832369 -2.6243789 -61.4326093
H 12.8078798 -2.2156713 -61.1651937
C 14.5459745 -3.0172483 -60.1610187
O 15.6270337 -3.5983735 -60.2304277
C 13.5987399 -3.7952014 -62.405327
H 13.5382697 -4.7567764 -61.8952256
H 12.6985773 -3.6284725 -62.9974839
C 15.1494966 -2.2202696 -63.3610773
H 14.6833669 -1.7644417 -64.236083
H 16.2312348 -2.0961457 -63.4033807

C 14.823558 -3.7137458 -63.3158808
H 14.6132543 -4.1093829 -64.3096971
H 15.6551597 -4.2543635 -62.8618106
N 13.9662979 -2.746838 -58.9908767
H 13.0661721 -2.2754632 -59.0090904
C 14.5182136 -3.1102436 -57.6789229
H 15.4998097 -3.5591397 -57.8229977
C 13.7014467 -4.2044761 -56.9428997
O 13.6238671 -4.1604091 -55.7146881
C 14.714941 -1.81596 -56.8514598
H 13.7756341 -1.2643591 -56.8431172
H 14.9534945 -2.0737327 -55.819355
C 15.8229594 -0.873091 -57.3541501
H 15.6404761 -0.6146515 -58.3959849
C 15.8120489 0.4127376 -56.5303236
H 16.028931 0.1980959 -55.4826737
H 14.8309305 0.8833728 -56.593059
H 16.5554212 1.1109018 -56.9145674
C 17.2136708 -1.5018124 -57.2217831
H 17.3755269 -1.8389114 -56.1975999
H 17.9750578 -0.7631874 -57.470298
H 17.3151205 -2.3433035 -57.9050428
N 13.0948036 -5.214763 -57.6151661
C 12.2053006 -6.1721577 -56.9514439
H 11.3888186 -5.644905 -56.4617025
C 12.9249789 -7.0386965 -55.9075064
O 12.3195788 -7.4085346 -54.9111168
C 11.6162382 -7.0363963 -58.0682371
H 11.3764525 -8.0460584 -57.7312578
H 10.7270492 -6.5549761 -58.4702904
C 13.2765494 -5.6211828 -59.0043124
H 12.7053484 -4.9577645 -59.6524598
H 14.33053 -5.6213029 -59.2835882
C 12.7120005 -7.0340446 -59.1282169
H 12.3119336 -7.2235629 -60.1252091
H 13.4796364 -7.7662537 -58.8743209
N 14.2193922 -7.3309605 -56.0875591
H 14.6758111 -6.9951267 -56.9187628
C 15.0103271 -8.0990672 -55.112049
H 14.434692 -8.9910839 -54.8655987
C 15.2070223 -7.332927 -53.7964256
O 15.0766997 -7.9116657 -52.717904
C 16.3497863 -8.552893 -55.7413196
H 16.1067111 -9.1238207 -56.6393262
C 17.2457079 -7.3784555 -56.1790683
H 17.6104389 -6.833258 -55.3064419

H 16.7096291 -6.695959 -56.8363059
H 18.1109972 -7.7622999 -56.7213065
C 17.1758962 -9.4774166 -54.8192359
H 18.0864798 -9.7670527 -55.3455807
H 17.4716894 -8.935527 -53.9193684
C 16.4479887 -10.7630839 -54.4053729
H 16.0929498 -11.2919093 -55.2902479
H 15.605773 -10.5327737 -53.7532482
H 17.1389073 -11.4070984 -53.8607548
N 15.3935547 -6.0141343 -53.87863
H 15.4044782 -5.57695 -54.785056
C 15.4989416 -5.1573755 -52.7005961
H 16.1260958 -5.6549509 -51.9609007
C 14.1275042 -4.979588 -52.0493859
O 14.0041547 -5.2410058 -50.8625342
C 16.1529791 -3.8128398 -53.0612849
H 15.5453294 -3.3096024 -53.8136461
H 16.1749842 -3.1849001 -52.1697456
C 17.5888162 -3.9565456 -53.5969701
H 17.9604463 -2.9636573 -53.8583359
H 17.5825816 -4.5500643 -54.513358
C 18.5239075 -4.5941443 -52.560969
O 18.5957494 -5.8428049 -52.4662396
O 19.14147 -3.8633975 -51.7551244
N 13.0718701 -4.7526557 -52.834066
H 13.246795 -4.5814834 -53.8192455
C 11.6674641 -4.7573474 -52.3915481
H 11.5110988 -3.9424476 -51.6838844
C 11.2638044 -6.055375 -51.6705936
O 10.7000723 -5.9916155 -50.5815018
C 10.8225721 -4.4658085 -53.6373247
H 11.2703018 -4.9725609 -54.4847155
H 10.8980451 -3.3996945 -53.8162933
C 9.3400869 -4.8422281 -53.6032018
O 8.5960027 -4.4336681 -52.6834481
O 8.8625046 -5.4577929 -54.5871172
N 11.6917214 -7.2238873 -52.1577427
H 12.1378338 -7.2256039 -53.0701622
C 11.4803921 -8.5084706 -51.4804795
H 10.4192842 -8.5902366 -51.2439924
C 12.2581264 -8.616735 -50.1576347
O 11.7173718 -9.1125472 -49.1729519
C 11.8590517 -9.6523727 -52.4351888
H 12.9441046 -9.6885066 -52.5437256
H 11.4310558 -9.4628766 -53.4176041
C 11.3583046 -11.0199242 -51.9562242

H 11.7482812 -11.2164564 -50.9570363
H 11.7634607 -11.7769659 -52.6281464
S 9.5513028 -11.2041661 -51.9296248
C 9.4280423 -12.9497713 -51.463499
H 8.3802362 -13.2482151 -51.4521366
H 9.8585708 -13.0948903 -50.4720088
H 9.9653893 -13.5614913 -52.1881289
N 13.4992124 -8.1155487 -50.1013757
H 13.8751445 -7.7085546 -50.9531837
C 14.3462607 -8.0992723 -48.8917458
H 14.3261654 -9.0880359 -48.4302966
C 13.8414349 -7.109273 -47.8321094
O 13.8696885 -7.4088854 -46.6438106
C 15.7803139 -7.7771225 -49.3386629
H 16.0728206 -8.4953921 -50.1072518
H 15.78842 -6.7853287 -49.7877521
C 16.8146607 -7.8333564 -48.2019743
H 16.5459291 -7.1366515 -47.4067893
H 16.8280532 -8.8437172 -47.7908348
C 18.2187971 -7.4873246 -48.7193819
H 18.4184365 -8.0829343 -49.6128579
H 18.9536731 -7.7499724 -47.9570584
C 18.3487833 -5.992413 -49.0458139
H 18.4319706 -5.4217815 -48.1165061
H 17.4451803 -5.6561907 -49.5619324
N 19.5031294 -5.7293883 -49.9309239
H 20.3954323 -5.9966135 -49.5549232
H 19.5174901 -4.749167 -50.2346865
H 19.3464276 -6.1609146 -50.848426
N 13.3322727 -5.9630881 -48.2710839
H 13.3916351 -5.7966786 -49.2700395
C 12.6752334 -4.9259791 -47.4652083
H 13.3066394 -4.639705 -46.6204001
C 11.3514425 -5.4659331 -46.907204
O 11.0801258 -5.3692947 -45.708615
C 12.463281 -3.6936637 -48.3766892
H 12.0638674 -4.0566426 -49.3241162
C 11.3939495 -2.735481 -47.8251092
H 11.5699158 -2.5482408 -46.7698325
H 10.4027027 -3.1721763 -47.9439616
H 11.4167651 -1.795219 -48.3641339
C 13.8090055 -3.0011842 -48.7056933
H 14.1943818 -2.5034339 -47.8276267
H 14.5553025 -3.7476582 -48.9678682
C 13.7415405 -1.9940272 -49.8643963
H 13.1388901 -1.128049 -49.5957665

H 13.3172396 -2.4665957 -50.7493734
H 14.7489847 -1.6485579 -50.0979748
N 10.5569209 -6.1251239 -47.7575478
H 10.8058184 -6.1502801 -48.7417588
C 9.3406062 -6.8155937 -47.3499945
H 8.687436 -6.0980664 -46.8515223
C 9.6451199 -7.9320667 -46.3426887
O 8.9789309 -7.9857565 -45.3181334
C 8.6090851 -7.3439192 -48.5900398
H 9.2347282 -8.0646011 -49.1166765
H 8.3743305 -6.5167723 -49.2610325
H 7.6825766 -7.8315087 -48.286617
N 10.6967849 -8.7327052 -46.5459848
H 11.1877227 -8.6812436 -47.4322827
C 11.1410652 -9.7413254 -45.5766499
H 10.3095113 -10.4188421 -45.3765601
C 11.5417736 -9.1181235 -44.2316211
O 11.0027769 -9.5243077 -43.2065835
C 12.2879134 -10.5821685 -46.1574739
H 13.1383181 -9.9410307 -46.3900585
H 11.9539012 -11.054893 -47.0827182
C 12.7241101 -11.6616207 -45.1622198
O 13.5224159 -11.3337599 -44.252894
O 12.2018212 -12.793503 -45.265959
N 12.3829782 -8.0769071 -44.2118859
H 12.8033958 -7.7736289 -45.0842588
C 12.8418208 -7.4756884 -42.9545116
H 13.20788 -8.2795145 -42.315361
C 11.7105569 -6.7843906 -42.179033
O 11.5105007 -7.0762136 -40.995856
C 14.0107534 -6.5158331 -43.2086705
H 14.8199205 -7.0620265 -43.6953675
H 13.6867322 -5.7238326 -43.8863098
C 14.5352889 -5.889822 -41.9505213
C 15.3513898 -6.4920806 -41.0560262
H 15.7532765 -7.4912347 -41.1718495
N 15.5335855 -5.6654725 -39.9629168
H 16.0897329 -5.9158626 -39.160673
C 14.8148192 -4.49376 -40.0872147
C 14.6321954 -3.3866221 -39.242576
H 15.1032791 -3.3500841 -38.2735638
C 13.819259 -2.3236695 -39.676404
H 13.6551516 -1.4597462 -39.0434416
C 13.2231146 -2.3758491 -40.9490175
H 12.61592 -1.5480294 -41.2963996
C 13.413713 -3.4925454 -41.785488

H 12.9690569 -3.4997165 -42.7718436
C 14.1985453 -4.5894945 -41.3712467
N 10.9043925 -5.9527161 -42.8492866
H 11.1134384 -5.7630804 -43.8240366
C 9.7438538 -5.2921988 -42.2212211
H 10.0844229 -4.773692 -41.3238945
C 8.6828352 -6.2978319 -41.7797415
O 8.1475363 -6.1684389 -40.6765001
C 9.0796136 -4.2487583 -43.1304959
H 8.177974 -3.8895772 -42.6371761
C 9.9720763 -3.0467225 -43.4104666
H 10.8669468 -3.3593763 -43.9445606
H 10.2661708 -2.5763767 -42.4714854
H 9.4314004 -2.3171273 -44.0117579
O 8.7181591 -4.8066588 -44.3726664
H 9.5496823 -4.9479732 -44.8577988
N 8.4358032 -7.3581704 -42.5633335
H 8.8963046 -7.4314515 -43.466883
C 7.5723311 -8.4495715 -42.1268535
H 6.6350766 -8.0155664 -41.7795424
C 8.1700975 -9.1517994 -40.9247922
O 7.4758181 -9.174459 -39.9330543
C 7.2448461 -9.4620492 -43.2330206
H 8.1639771 -9.7685181 -43.7320877
H 6.8107539 -10.3548548 -42.779457
C 6.239774 -8.9211213 -44.2597927
H 6.2060887 -9.6067299 -45.1067549
H 6.5649357 -7.9487869 -44.6272582
C 4.8301142 -8.7907822 -43.6962677
O 4.0185217 -9.6936336 -43.7800733
N 4.4373901 -7.6329512 -43.2092787
H 3.4549186 -7.5555601 -42.9669857
H 5.0762198 -6.8618807 -43.140931
N 9.4273277 -9.6006942 -40.9284614
H 9.9686587 -9.4839231 -41.780828
C 10.0214635 -10.412134 -39.8567427
H 9.3084547 -11.1990181 -39.609992
C 10.253828 -9.6601678 -38.5291229
O 10.2250454 -10.2893742 -37.4666527
C 11.2944858 -11.0964365 -40.3945858
H 11.8618845 -11.5134649 -39.5643313
H 11.9244235 -10.3674132 -40.904203
C 10.9694015 -12.2521054 -41.3360884
O 10.0165345 -12.9890257 -41.1245311
N 11.7728694 -12.4937022 -42.343769
H 11.4995019 -13.1733068 -43.0362986

H 12.499172 -11.8467642 -42.6449554
N 10.2929531 -8.3240194 -38.537167
H 10.3624872 -7.8437779 -39.4304161
C 10.2075646 -7.5261957 -37.311539
H 10.9370343 -7.9083139 -36.5955343
C 8.8173012 -7.6078404 -36.6366541
O 8.7224682 -7.6976754 -35.4119872
C 10.5911494 -6.0827475 -37.6571958
H 9.8848154 -5.6694188 -38.3793169
H 11.5923603 -6.0574983 -38.0909011
H 10.5793001 -5.471428 -36.7549126
N 7.7281362 -7.648853 -37.4093658
H 7.8464977 -7.6620426 -38.4175533
C 6.3615845 -7.7065981 -36.8837774
H 6.2525433 -6.8732437 -36.1891368
C 5.9991926 -8.9928891 -36.0905116
O 5.3798625 -8.8602907 -35.0312186
C 5.3875801 -7.4605357 -38.0417105
H 5.317507 -8.3466364 -38.6717451
H 5.7360643 -6.6360806 -38.6607618
H 4.3964641 -7.2320804 -37.651936
N 6.3537059 -10.227742 -36.5201682
C 6.0921003 -11.4546801 -35.7992746
H 5.0595046 -11.4728627 -35.4552056
C 7.0141917 -11.5868196 -34.5825291
O 6.5442037 -12.022288 -33.5387212
C 6.2944393 -12.5878507 -36.8162142
H 6.5796654 -13.5239315 -36.3340225
H 5.3790469 -12.7278175 -37.3944718
C 6.9710996 -10.6108314 -37.7595143
H 6.2311127 -10.4906961 -38.55036
H 7.8621115 -10.0271148 -37.9027458
C 7.3852351 -12.0701274 -37.7139111
H 7.3632158 -12.5333561 -38.7012058
H 8.3603036 -12.1870356 -37.2408137
N 8.2600128 -11.0993172 -34.6473682
H 8.5974122 -10.7351963 -35.529587
C 9.1638845 -11.0324667 -33.493428
H 9.2559856 -12.0264496 -33.0531509
C 8.6246533 -10.1030979 -32.3918366
O 8.5898426 -10.4804776 -31.2234502
C 10.5463395 -10.5906214 -33.9929249
H 10.9400176 -11.3588206 -34.6604347
H 10.4349591 -9.6795107 -34.5821898
C 11.5645544 -10.3336599 -32.8982995
C 12.1676299 -11.4139774 -32.226674

H 11.9007571 -12.4282594 -32.4853161
C 13.115578 -11.1764532 -31.2147571
H 13.5750579 -12.007349 -30.6985902
C 13.464218 -9.8578058 -30.873536
H 14.194 -9.6750549 -30.0974929
C 12.8624226 -8.7769348 -31.541472
H 13.1324783 -7.7636193 -31.281157
C 11.9107974 -9.0133361 -32.5496745
H 11.4482941 -8.1777601 -33.0568941
N 8.0603085 -8.9495652 -32.7631566
H 8.1434311 -8.665565 -33.7337643
C 7.4168677 -8.0211198 -31.8271501
H 7.9214617 -8.1082216 -30.8649975
C 5.9315139 -8.326438 -31.5456683
O 5.2535744 -7.4801421 -30.961817
C 7.5941536 -6.574922 -32.324633
H 7.124499 -6.4941971 -33.3064549
H 7.0538484 -5.9068789 -31.6550215
C 9.0136829 -6.0386909 -32.3987931
C 9.4463363 -5.359517 -33.5561945
H 8.7912065 -5.2787379 -34.4129759
C 10.7433545 -4.8150995 -33.6203321
H 11.075635 -4.3075551 -34.5142454
C 11.6148391 -4.9453172 -32.5180674
O 12.8717441 -4.435554 -32.5709646
H 13.0345853 -3.9454995 -33.3800472
C 11.174962 -5.6029107 -31.3511048
H 11.8453005 -5.6829437 -30.507972
C 9.8769556 -6.144496 -31.2893049
H 9.5563376 -6.6474406 -30.3872537
N 5.3871766 -9.481238 -31.956426
H 6.0033601 -10.1538206 -32.4006634
C 3.9274016 -9.6951489 -32.0875758
H 3.5788743 -9.0167038 -32.8656313
C 3.101727 -9.3795924 -30.8315989
O 1.9946272 -8.8568 -30.9597617
C 3.6184426 -11.1199467 -32.5819641
H 4.1373467 -11.2775613 -33.5237867
C 4.0262775 -12.229422 -31.6076246
H 3.4924745 -12.148523 -30.6645446
H 5.0959923 -12.1726016 -31.405786
H 3.820187 -13.1996915 -32.0601955
O 2.2341245 -11.2757929 -32.8256162
H 2.1031661 -11.0808149 -33.7675107
N 3.6312656 -9.6287895 -29.6257628
H 4.5649006 -10.0097299 -29.5872965

C 2.9405287 -9.3177676 -28.3687039
H 1.9221669 -9.6969241 -28.445873
C 2.8276563 -7.8017421 -28.1507249
O 1.7167741 -7.2833488 -28.1555535
C 3.6362754 -10.0368499 -27.1973246
H 3.6353806 -11.1098669 -27.3948457
H 4.6745595 -9.7054275 -27.1474315
C 2.978396 -9.7839054 -25.8233671
H 2.9966347 -8.7178673 -25.6005677
C 1.5310448 -10.2812004 -25.7697653
H 1.4817257 -11.3377576 -26.0293787
H 0.9096754 -9.6938931 -26.4446131
H 1.1425322 -10.1385847 -24.7604909
C 3.7720642 -10.514862 -24.7416685
H 3.7563083 -11.5903824 -24.9153329
H 3.3368301 -10.2987848 -23.7649057
H 4.8026104 -10.1599394 -24.7388168
N 3.9616382 -7.0915666 -28.1298216
H 4.8305866 -7.5964343 -28.1841824
C 4.0327294 -5.6286734 -27.9990707
H 3.5475118 -5.3373421 -27.066531
C 3.3223149 -4.9052738 -29.1588863
O 2.7887892 -3.8112358 -29.0000078
C 5.522688 -5.2420034 -27.9073454
H 5.9417744 -5.7125461 -27.0159418
H 6.0387796 -5.6508386 -28.77724
C 5.8148372 -3.7290084 -27.839953
H 5.4302001 -3.2507583 -28.740477
C 5.2052008 -3.0369301 -26.6265279
H 5.5270028 -3.5308572 -25.7079944
H 4.1165442 -3.0902357 -26.6670599
H 5.5002705 -1.9890255 -26.5938106
C 7.3263404 -3.5054407 -27.8182893
H 7.7487717 -3.9367071 -26.9075049
H 7.5460048 -2.4366408 -27.8259105
H 7.7977514 -3.9704279 -28.681937
N 3.3240981 -5.4922525 -30.3565642
H 3.8200164 -6.3736162 -30.4481514
C 2.6217824 -4.9657189 -31.5266407
H 2.8835245 -3.9148142 -31.6417372
C 1.0943839 -5.029999 -31.3514762
O 0.4108227 -4.0473604 -31.6319395
C 3.127867 -5.7370606 -32.7582961
H 4.209855 -5.607543 -32.8206349
H 2.9292394 -6.7975108 -32.6056175
C 2.5111026 -5.3318787 -34.1077532

H 1.4532277 -5.5947615 -34.1197899
C 2.6686143 -3.8435188 -34.4097948
H 3.7203921 -3.5715168 -34.3494023
H 2.0853599 -3.2554452 -33.704076
H 2.2981698 -3.6320755 -35.4139741
C 3.226156 -6.095398 -35.2194507
H 4.2560045 -5.7514367 -35.3113525
H 2.7083162 -5.937219 -36.1650524
H 3.2404669 -7.1588567 -34.9842874
N 0.5617721 -6.1321914 -30.8100106
H 1.1854655 -6.8955181 -30.565335
C -0.8774119 -6.2932057 -30.5294087
H -1.4450707 -5.8133694 -31.3273322
C -1.3387857 -5.5989339 -29.2472027
O -2.4972833 -5.2107496 -29.1799714
C -1.2259879 -7.7857952 -30.5059816
H -0.5395652 -8.3161481 -29.8425807
H -2.2423577 -7.9080194 -30.1304602
C -1.1550235 -8.3653732 -31.9235204
H -0.1595575 -8.2283015 -32.3457716
H -1.8592111 -7.825137 -32.5503861
C -1.517416 -9.8499764 -31.9535904
H -1.7789138 -10.1101266 -32.9786297
H -2.3926915 -10.0247935 -31.3275004
N -0.3815104 -10.6888054 -31.5472774
H 0.4686067 -10.5891001 -32.0922565
C -0.2819976 -11.5284697 -30.5397689
N 0.7987728 -12.2399241 -30.4453776
H 0.9213052 -12.9191248 -29.7195572
H 1.4926744 -12.1243647 -31.1788985
N -1.2140364 -11.691179 -29.6413828
H -2.0524106 -11.1461476 -29.7056909
H -1.0837421 -12.3425586 -28.8909044
N -0.4359914 -5.4137744 -28.291648
H 0.4565919 -5.878876 -28.4106519
C -0.6121594 -4.6637386 -27.0405899
H -1.5484464 -4.9448234 -26.5574101
C -0.6419177 -3.1442925 -27.2699944
O -1.4719593 -2.454789 -26.6849815
C 0.5604857 -5.0831334 -26.1452238
H 0.4619065 -6.1502756 -25.9371324
H 1.4675594 -4.941078 -26.7285774
C 0.7664583 -4.3591921 -24.8161269
H -0.1072872 -4.5150072 -24.1788573
H 0.8875321 -3.2879945 -24.9917974
C 2.035639 -4.9226683 -24.1581527

O 1.8929473 -5.6917077 -23.1831504
O 3.1384206 -4.6239782 -24.6732221
N 0.1917402 -2.6286023 -28.1822229
H 0.9142711 -3.2259607 -28.5641261
C 0.1832659 -1.2150978 -28.5658876
H 0.0037262 -0.6137299 -27.6722889
C -0.9246664 -0.8558517 -29.5768898
O -1.3427934 0.2986789 -29.6289793
C 1.5712314 -0.8568396 -29.111109
H 1.7755088 -1.4407739 -30.0082925
H 2.3275326 -1.0683264 -28.3535512
H 1.5991124 0.2052893 -29.3567835
N -1.3690744 -1.8151607 -30.399589
H -0.9570034 -2.7323259 -30.3107115
C -2.4122034 -1.6197109 -31.4130676
H -2.8401389 -0.6237023 -31.2850044
C -3.5838278 -2.5891446 -31.1989809
O -4.4984085 -2.2893246 -30.4387788
C -1.8158924 -1.6685056 -32.833723
H -1.3145361 -2.624073 -32.987596
H -2.63415 -1.6141187 -33.553035
C -0.8409829 -0.5505884 -33.1512774
C -1.3096435 0.649862 -33.7150379
H -2.3647469 0.7833783 -33.9051926
C 0.531843 -0.7025696 -32.8802269
H 0.8864084 -1.6149005 -32.4260483
C -0.4110316 1.6879453 -34.01348
H -0.7806818 2.6123909 -34.4308109
C 1.4300197 0.3388147 -33.1757978
H 2.4760809 0.2347951 -32.9481857
C 0.9602995 1.5313944 -33.7512922
H 1.6466328 2.3356076 -33.9776088
N -3.6205628 -3.7132496 -31.9236746
H -2.8479066 -3.9034345 -32.5436994
C -4.6046085 -4.7861899 -31.7255437
H -4.6477881 -5.0219769 -30.6606922
C -4.2176418 -6.0541322 -32.4919207
O -3.4466607 -6.006089 -33.4557384
C -6.0082382 -4.3507814 -32.1811528
H -6.3304691 -3.4699273 -31.624698
H -6.7175425 -5.152142 -31.9705056
O -6.0325689 -4.0692292 -33.5679786
H -5.899994 -3.1163384 -33.6518697
N -4.8741413 -7.1737827 -32.163381
H -5.4716506 -7.1487112 -31.3494907
C -4.8665038 -8.4037354 -32.9751168

H -3.8421984 -8.7622554 -33.024553
C -5.296619 -8.1459118 -34.4291164
O -4.6251069 -8.5946963 -35.3565651
C -5.7191938 -9.4992887 -32.2851475
H -5.2617698 -9.6962688 -31.3135578
C -7.1749464 -9.0643969 -32.0189038
H -7.7180113 -8.9252308 -32.9539572
H -7.2157387 -8.1483551 -31.4309694
H -7.6857446 -9.8419113 -31.4478146
C -5.7591062 -10.8389807 -33.0521625
H -6.3486961 -11.5529072 -32.4750014
H -6.2623358 -10.6958331 -34.0075932
C -4.3868889 -11.4732058 -33.3065014
H -3.86465 -11.6184485 -32.3614838
H -3.7929475 -10.8453952 -33.9693974
H -4.5241808 -12.4437534 -33.7840225
N -6.3197801 -7.3081212 -34.6459337
H -6.7867577 -6.9174776 -33.8407924
C -6.8108968 -6.9429317 -35.9822531
H -6.9295796 -7.8697976 -36.5438468
C -5.8010042 -6.0946841 -36.7703672
O -5.7534856 -6.2049544 -37.9993011
C -8.1929232 -6.2678168 -35.828368
H -8.8936588 -6.9935093 -35.4113768
H -8.1035006 -5.4462128 -35.1147961
C -8.795135 -5.6932521 -37.1238768
H -9.7559562 -5.2332397 -36.8894435
H -8.1411857 -4.8942535 -37.4703831
C -9.0162358 -6.7421972 -38.2346595
H -9.9843958 -7.2211919 -38.076221
H -8.2596339 -7.5233973 -38.1830747
C -8.9707821 -6.1540308 -39.6519625
H -9.7832162 -5.4324325 -39.7803459
H -9.1312743 -6.9749907 -40.3575952
N -7.6582374 -5.5172426 -39.9201159
H -7.5238449 -4.6979612 -39.3466649
H -7.5232324 -5.260878 -40.8933727
H -6.8930147 -6.1433565 -39.6825251
N -4.9831557 -5.2914209 -36.0868715
H -5.1114733 -5.2316311 -35.0833861
C -3.8621657 -4.5557308 -36.6755995
H -4.2125805 -3.9476157 -37.5101316
H -3.4303927 -3.8987074 -35.9202956
C -2.7652385 -5.4947951 -37.177779
O -2.4814048 -5.5085537 -38.373279
N -2.2677174 -6.3806718 -36.3070245

H -2.5945535 -6.342489 -35.3479413
C -1.2607822 -7.3958332 -36.6512525
H -0.3445124 -6.8925585 -36.961776
C -1.6939054 -8.2984128 -37.8176158
O -0.9311845 -8.4717807 -38.7673839
C -0.9587692 -8.2239316 -35.3945171
H -0.5126998 -7.5591821 -34.6545479
H -1.9003645 -8.5918536 -34.9852958
C -0.0205586 -9.403772 -35.5957312
C -0.476939 -10.7237244 -35.4080804
H -1.509788 -10.9114615 -35.1526382
C 0.4018223 -11.8087914 -35.5828742
H 0.0411055 -12.8215519 -35.4605813
C 1.7396798 -11.5814567 -35.9510346
H 2.4073452 -12.4177367 -36.1126058
C 2.197147 -10.2669613 -36.1448257
H 3.2167878 -10.0866761 -36.4561706
C 1.3158525 -9.1847377 -35.975059
H 1.6594669 -8.1812815 -36.1674815
N -2.9508894 -8.7602365 -37.8162996
H -3.5180262 -8.6013591 -36.9897401
C -3.5164419 -9.5705426 -38.9055008
H -2.8830213 -10.4489155 -39.0412821
C -3.4884375 -8.8191223 -40.2441416
O -3.0854456 -9.4032043 -41.2444796
C -4.9353343 -10.0606912 -38.5229828
H -5.4930771 -9.2156787 -38.1147458
C -5.7199256 -10.6008875 -39.7362459
H -5.1748012 -11.428586 -40.1934969
H -5.8704642 -9.8203191 -40.4808494
H -6.7059161 -10.950001 -39.4339102
C -4.8187313 -11.1576382 -37.4374046
H -4.3660146 -12.0503218 -37.8713733
H -4.1598068 -10.814504 -36.6407699
C -6.1553173 -11.5460725 -36.7923586
H -6.7955629 -12.0580817 -37.5094256
H -6.6616767 -10.6550823 -36.4198857
H -5.9674939 -12.2254155 -35.9610021
N -3.8323925 -7.5248185 -40.2879394
H -4.0656676 -7.0556824 -39.4218744
C -3.7691065 -6.745843 -41.5391746
H -4.3005425 -7.2994218 -42.3139513
C -2.3393446 -6.5519942 -42.0526116
O -2.1145754 -6.6247537 -43.2621331
C -4.4332741 -5.3718712 -41.3703427
H -3.8537605 -4.6109815 -41.8970195

H -4.4733346 -5.1006908 -40.3138927
O -5.7412933 -5.3987764 -41.9212897
H -5.6390313 -5.3843889 -42.8827614
N -1.3861787 -6.3126124 -41.1495095
H -1.6547347 -6.2826085 -40.1714205
C 0.0158965 -6.0425797 -41.481886
H 0.0577829 -5.3112942 -42.2894989
C 0.740751 -7.3012983 -41.9896076
O 1.4825599 -7.2298511 -42.9719905
C 0.6993255 -5.4537121 -40.2332656
H 0.5759614 -6.161702 -39.4119538
H 1.768021 -5.3583632 -40.4270006
C 0.1460188 -4.0790425 -39.7977291
H -0.9410754 -4.0968931 -39.8068985
C 0.5881962 -3.7787531 -38.3651539
H 1.6688825 -3.8436922 -38.2817429
H 0.1358835 -4.4987693 -37.6855946
H 0.2728934 -2.7773283 -38.0815491
C 0.5838749 -2.9467471 -40.7300837
H 1.6679734 -2.8669288 -40.751094
H 0.1714852 -2.0035095 -40.3724266
H 0.2150466 -3.1247907 -41.7383577
N 0.4550266 -8.4611916 -41.3890336
H -0.1379686 -8.4416052 -40.564342
C 0.9646168 -9.7595951 -41.8364496
H 2.0196674 -9.6433663 -42.0892642
C 0.2485841 -10.2596498 -43.1035624
O 0.8987943 -10.7616169 -44.0158858
C 0.8455888 -10.7511466 -40.6638834
H 1.3523783 -10.326376 -39.7955445
H -0.210257 -10.8640917 -40.4110157
C 1.4357562 -12.1477194 -40.9424942
H 0.9147976 -12.6038002 -41.7839374
C 2.9345315 -12.1074372 -41.2461209
H 3.4682867 -11.5756565 -40.4587843
H 3.1048157 -11.6059846 -42.1990644
H 3.3258448 -13.1209115 -41.3293987
C 1.228386 -13.036948 -39.7157139
H 1.7534838 -12.6171902 -38.8578697
H 1.6078131 -14.0384338 -39.9180822
H 0.1647686 -13.1057515 -39.4881301
N -1.0698244 -10.0583188 -43.2209953
H -1.5784126 -9.6705319 -42.4337335
C -1.8288138 -10.4543792 -44.4133578
H -1.6406428 -11.512455 -44.6026739
C -1.3632127 -9.6903915 -45.6534104

O -1.0336879 -10.3147171 -46.6612399
C -3.3368109 -10.2801862 -44.1613091
H -3.5384081 -9.2595103 -43.8397539
H -3.6148275 -10.9493552 -43.3461491
C -4.2354917 -10.6192243 -45.3629775
H -5.2387681 -10.8120393 -44.981554
H -3.8764807 -11.5469091 -45.8110439
S -4.390968 -9.3657168 -46.6781462
C -5.1799099 -8.0079527 -45.7729399
H -5.4383667 -7.2116522 -46.4701612
H -4.4931662 -7.6138861 -45.0242914
H -6.0870682 -8.3705426 -45.2881865
N -1.2836702 -8.3547994 -45.597801
H -1.5351508 -7.8696527 -44.7427603
C -0.9788531 -7.5879528 -46.8043397
H -1.457325 -8.1034868 -47.6360249
C 0.521404 -7.5756896 -47.1451478
O 0.8637811 -7.5102657 -48.3264741
C -1.6430715 -6.2032923 -46.748782
H -1.1355764 -5.5871202 -46.0089988
H -2.6802938 -6.3144597 -46.4385527
C -1.6394516 -5.5151375 -48.1094585
N -0.9709235 -4.389742 -48.2157069
H -0.4685428 -4.0199304 -47.4270163
H -0.9501478 -3.9346266 -49.1103779
O -2.2486604 -5.9590541 -49.0785727
N 1.416904 -7.7516498 -46.1678263
H 1.0980991 -7.8154289 -45.2078644
C 2.8375119 -7.9866019 -46.4363406
H 3.1969886 -7.2710589 -47.1758242
H 3.3911814 -7.8211393 -45.5225839
C 3.1279278 -9.4043741 -46.945777
O 3.8298141 -9.5493483 -47.9453507
N 2.447086 -10.4225332 -46.4101552
H 1.9315207 -10.2624063 -45.5490469
C 2.4854058 -11.7979805 -46.9139969
H 3.5152832 -12.1562071 -46.9087821
H 1.8931677 -12.4278665 -46.2504679
C 1.9239234 -11.9295401 -48.3328251
O 2.5932635 -12.4826701 -49.2022689
N 0.7929469 -11.2777152 -48.628854
H 0.267872 -10.8632915 -47.8641561
C 0.2683398 -11.1228987 -49.9938683
H 0.0573556 -12.1067264 -50.4128921
C 1.2826057 -10.4351154 -50.919821
O 1.4660857 -10.8427349 -52.0678355

C -1.03995 -10.3126457 -49.9511539
H -1.8060414 -10.9034882 -49.4476874
H -0.8743164 -9.4136616 -49.3588149
C -1.5543657 -9.8836418 -51.2950987
C -1.1187229 -8.7991881 -51.9758444
H -0.3563124 -8.1147227 -51.6133052
N -1.6791194 -8.7882528 -53.2363242
H -1.3736528 -8.1625098 -53.9676252
C -2.5438482 -9.8457334 -53.4186151
C -3.3411695 -10.2526051 -54.4992947
H -3.3476281 -9.6931163 -55.422582
C -4.1539985 -11.3894695 -54.358604
H -4.801853 -11.6994674 -55.1695271
C -4.1314936 -12.1172815 -53.1550939
H -4.7589197 -12.9925633 -53.044334
C -3.3101688 -11.7105559 -52.0842815
H -3.3046579 -12.2804389 -51.1652787
C -2.499787 -10.5592805 -52.1831049
N 1.96428 -9.3993704 -50.4253499
H 1.7809878 -9.1156849 -49.4709382
C 2.9636726 -8.654149 -51.2011413
H 2.5137098 -8.3680451 -52.1510126
C 4.1751778 -9.5252683 -51.5380323
O 4.584636 -9.5468988 -52.6991712
C 3.3562456 -7.3613529 -50.4714756
H 3.7076416 -7.5770001 -49.4645344
C 4.4220107 -6.5580398 -51.2039862
H 4.0784879 -6.3762013 -52.2179027
H 5.3645424 -7.1059384 -51.2128172
H 4.574357 -5.6062331 -50.6915057
O 2.2203371 -6.5288334 -50.4111686
H 1.6945377 -6.8265429 -49.6442254
N 4.6517677 -10.3483214 -50.5991615
H 4.2697926 -10.2894075 -49.6593257
C 5.6772822 -11.3580867 -50.8568759
H 6.5451977 -10.8700353 -51.3041686
C 5.1923812 -12.4341222 -51.8490049
O 5.8968642 -12.7354913 -52.8110289
C 6.1067911 -11.9647842 -49.5145566
H 5.2580115 -12.4435631 -49.0255728
H 6.499734 -11.1841373 -48.8617151
H 6.8863492 -12.7084471 -49.6781218
N 3.9534366 -12.9249302 -51.7086818
H 3.4267844 -12.6496469 -50.8836684
C 3.334672 -13.8997941 -52.6291921
H 3.9737587 -14.7791788 -52.6745089

C 3.2311635 -13.3581777 -54.0601941
O 3.501972 -14.0879971 -55.0132972
C 1.9611062 -14.3607064 -52.0920771
H 1.4005589 -13.4924383 -51.7508018
C 1.0987965 -15.094003 -53.1308964
H 1.6582642 -15.9270397 -53.5595551
H 0.8015677 -14.4086898 -53.9226587
H 0.1946828 -15.4764729 -52.6554527
C 2.1541222 -15.3188572 -50.9094302
H 2.6359165 -16.2380737 -51.2437664
H 1.18814 -15.559973 -50.4658729
H 2.7765934 -14.8574841 -50.1444956
N 2.9105554 -12.073782 -54.2549198
H 2.6628376 -11.510579 -53.4444772
C 2.8382215 -11.4848562 -55.603466
H 2.5477263 -12.2731829 -56.294507
C 4.1932573 -11.0523229 -56.1735234
O 4.3050569 -11.0014922 -57.3976465
C 1.7500749 -10.3972881 -55.6684607
H 1.973677 -9.6224873 -54.932472
H 1.7766916 -9.9369756 -56.657886
C 0.3181127 -10.9321047 -55.4242165
H 0.2277581 -11.265099 -54.390261
H -0.3748739 -10.1024204 -55.5677658
C -0.0987512 -12.0894509 -56.3560436
H 0.544746 -12.9544503 -56.17014
H 0.0153415 -11.7777132 -57.39766
C -1.5577073 -12.5187353 -56.1029819
H -2.2269318 -11.6802266 -56.3325773
H -1.6736438 -12.7561101 -55.0366384
N -1.920309 -13.7026939 -56.930816
H -1.8494992 -13.5031153 -57.9228767
H -2.903941 -13.9547134 -56.8042702
H -1.3493329 -14.5134468 -56.71273
N 5.239108 -10.8811623 -55.358782
H 5.0832237 -10.9063656 -54.3572045
C 6.6252368 -10.7490509 -55.8568326
H 6.6136699 -10.1725473 -56.7835268
C 7.23075 -12.111499 -56.2120809
O 7.8163342 -12.237301 -57.2830009
C 7.5248285 -9.9796415 -54.8686851
H 7.4639628 -10.4501565 -53.8877875
C 8.9905939 -9.944826 -55.3227672
H 9.0568977 -9.5997377 -56.3555774
H 9.4372889 -10.9357012 -55.2428839
H 9.5579733 -9.2635562 -54.6896495

C 7.0793244 -8.5186274 -54.7449023
H 7.2214826 -8.0117064 -55.6995769
H 7.6884329 -8.0243809 -53.9867521
H 6.0354079 -8.4688468 -54.4529373
N 7.0278598 -13.1595392 -55.4024952
H 6.5409429 -13.011564 -54.5230944
C 7.6271667 -14.4858677 -55.6559459
H 8.6634613 -14.3031979 -55.9438593
C 6.9906641 -15.2279986 -56.8419336
O 7.6716931 -15.9879257 -57.5225639
C 7.6575082 -15.3317962 -54.3638388
H 7.9124323 -14.6589608 -53.5431137
C 6.2996944 -15.9707711 -54.0400428
H 6.0816508 -16.7851927 -54.7321286
H 5.5262014 -15.2171777 -54.1221082
H 6.2972443 -16.3543042 -53.0201916
C 8.777694 -16.3911983 -54.4438692
H 8.5331507 -17.1303691 -55.2074872
H 9.7068405 -15.8989223 -54.7355244
C 9.0351373 -17.1189259 -53.1182629
H 8.173149 -17.7267722 -52.8436591
H 9.2387219 -16.3944539 -52.3289923
H 9.8999306 -17.7735446 -53.2299662
N 5.7387061 -14.9039989 -57.1868191
H 5.2127234 -14.3222172 -56.5485866
C 5.1062204 -15.3009733 -58.4534771
H 5.3874356 -16.3318692 -58.6680816
C 5.6030433 -14.4874707 -59.672003
O 5.0849158 -14.6599443 -60.773668
C 3.5735904 -15.2270815 -58.2905951
H 3.3168007 -14.2373792 -57.910768
H 3.1057086 -15.3358337 -59.269651
C 2.9775319 -16.3018904 -57.3585176
H 3.4899355 -16.2918303 -56.3990126
C 1.4957103 -15.9962792 -57.1210999
H 0.9610944 -15.9934025 -58.0710052
H 1.4152946 -15.0272253 -56.6348142
H 1.0717324 -16.755584 -56.4636815
C 3.0815974 -17.7101515 -57.947436
H 2.616133 -17.7444842 -58.9324024
H 2.5847047 -18.4241014 -57.2906702
H 4.1275589 -18.0041337 -58.0324987
N 6.5584693 -13.5695965 -59.4840437
H 6.9635998 -13.4805897 -58.559394
C 7.1419561 -12.7294521 -60.533019
H 8.0300305 -12.2428227 -60.1296168

H 7.4428545 -13.3604889 -61.3694312
C 6.2113295 -11.6372743 -61.0664694
O 6.4668133 -11.1007314 -62.1414297
N 5.0989407 -11.3521821 -60.3807085
H 4.9813281 -11.7997437 -59.4822739
C 4.011173 -10.5016823 -60.8819061
H 4.0372055 -10.488733 -61.9692609
C 4.1458654 -9.0360507 -60.4550618
O 4.0311999 -8.1487806 -61.2999311
C 2.6640571 -11.1114873 -60.4638078
H 1.8926477 -10.351034 -60.4842377
H 2.730934 -11.4831747 -59.4414438
C 2.2535011 -12.2657448 -61.3868724
H 1.4591474 -12.8339579 -60.902344
H 3.1069454 -12.932034 -61.5231857
S 1.6498686 -11.7327541 -63.0139356
C 1.6214996 -13.3280329 -63.8756168
H 1.2174771 -13.1904539 -64.8799789
H 0.9970689 -14.0358844 -63.3303468
H 2.6341481 -13.7267153 -63.9450518
N 4.4403578 -8.7747006 -59.1755103
H 4.5741277 -9.5745046 -58.565276
C 4.7721047 -7.4432695 -58.6461105
H 4.2329189 -6.6929209 -59.2287529
C 6.2628089 -7.1513591 -58.8255713
O 7.1132365 -8.0286716 -58.7190631
C 4.3124856 -7.3225627 -57.1776623
H 3.2218766 -7.3201566 -57.1538076
H 4.664482 -8.1940433 -56.6248092
C 4.8413101 -6.0586761 -56.4669405
H 5.9269408 -6.1201353 -56.3940316
H 4.5835999 -5.1738691 -57.0510368
C 4.2767258 -5.8952971 -55.0518431
H 4.2724669 -6.8599056 -54.5417714
H 3.2469937 -5.5465683 -55.1283597
C 5.0870934 -4.8828455 -54.2236432
H 4.4787623 -4.580427 -53.3658758
H 5.2958632 -3.991815 -54.8235343
N 6.3573003 -5.466491 -53.7188249
H 6.2395348 -6.3113772 -53.1868018
H 6.9070721 -4.8226841 -53.1439003
H 7.0461657 -5.6510471 -54.4521823
N 6.5831059 -5.8796803 -59.0166479
H 5.8235267 -5.2105063 -59.113986
C 7.9375657 -5.3227778 -59.0310328
H 8.6143732 -5.9856498 -58.4939189

C 7.9299697 -3.9753104 -58.3016906
O 6.8849011 -3.3299543 -58.2056804
C 8.3940536 -5.2114097 -60.498702
H 7.5989185 -4.7461053 -61.083704
C 9.6867695 -4.4393521 -60.7534294
H 10.461601 -4.7634189 -60.0624864
H 9.5092652 -3.3712309 -60.637248
H 10.0261984 -4.6212524 -61.7728531
O 8.6164189 -6.5095718 -60.9963761
H 8.1255677 -7.1366062 -60.4489059
N 9.0868513 -3.5097747 -57.8288001
H 9.9133854 -4.0836863 -57.9019445
C 9.2842826 -2.0970461 -57.484942
H 8.3344924 -1.5820776 -57.6283965
C 10.2397767 -1.4588863 -58.488381
O 11.1822076 -2.0878065 -58.9748673
C 9.6723067 -1.8988096 -55.9998239
H 9.4745394 -2.8358148 -55.4851521
C 11.1589555 -1.5434698 -55.8111287
H 11.3819425 -0.5661663 -56.2441685
H 11.7740098 -2.3086833 -56.2769605
H 11.4226248 -1.4985324 -54.7557414
C 8.7636256 -0.8305709 -55.3496209
H 8.9602225 0.1445593 -55.7991688
H 7.7226656 -1.0931424 -55.5430241
C 8.9349449 -0.7232145 -53.8274753
H 9.9006709 -0.2819557 -53.5826193
H 8.8562241 -1.7099889 -53.3708842
H 8.1532925 -0.0869728 -53.4164581
N 9.9517901 -0.2103244 -58.8094482
H 9.1682783 0.231535 -58.3373574
C 10.654951 0.6360141 -59.7608788
H 11.4393476 0.0802286 -60.2761536
C 11.3137825 1.7707113 -58.9834636
O 10.6744127 2.3280577 -58.0948429
C 9.5994166 1.0937026 -60.7929687
H 8.6089785 1.028997 -60.3467226
H 9.6158493 0.3827572 -61.6188321
C 9.6756245 2.4856417 -61.3678588
N 9.6560584 2.7683857 -62.7304314
C 9.6304442 4.1041051 -62.8320208
H 9.6677495 4.647484 -63.7669766
N 9.599696 4.6686446 -61.6145544
H 9.6754936 5.6576488 -61.4159251
C 9.6037052 3.6646771 -60.6808175
H 9.5692624 3.7963986 -59.6106405

N 12.5267095 2.1509732 -59.3842939
H 13.0058659 1.5797279 -60.0714761
C 13.1036285 3.4786336 -59.1419166
H 12.3791542 4.0801703 -58.6065819
C 13.4003518 4.137936 -60.4980748
O 13.9804095 3.4895145 -61.3751488
C 14.3486196 3.3947989 -58.2404563
H 14.0355429 3.0345998 -57.2602812
H 15.0443904 2.6671256 -58.661884
C 15.0998792 4.7290328 -58.0561606
H 15.5972644 4.9762276 -58.9978091
H 15.8861092 4.5829474 -57.3125827
C 14.2028056 5.9034645 -57.6227446
O 14.1451209 6.8963968 -58.3833069
O 13.5576779 5.8054743 -56.5593021
N 13.0453446 5.4126846 -60.6712948
H 12.6425479 5.9066621 -59.8759317
C 13.3632583 6.2062371 -61.8533931
H 14.1139036 5.6662568 -62.4167197
C 13.9496037 7.5757594 -61.5001384
O 13.3110616 8.3851175 -60.8299668
C 12.1357501 6.3519771 -62.752082
H 11.3121412 6.7960327 -62.1946084
H 11.8710934 5.3727228 -63.1411174
H 12.3807769 6.9994993 -63.5956822
N 15.109189 7.8745898 -62.0873356
H 15.4773862 7.2053572 -62.7563538
C 15.8432781 9.1316037 -61.9126432
H 15.2448618 9.8123746 -61.30726
C 16.0561624 9.7692783 -63.2824116
O 16.4188304 9.0839867 -64.2349318
C 17.1881392 8.8969502 -61.188147
H 17.8076121 8.2561194 -61.8169794
C 16.9764192 8.1765845 -59.8366834
H 16.3622953 8.7964619 -59.1838709
H 16.4432474 7.2414061 -60.0042166
C 17.9261826 10.2389898 -60.9959842
H 17.3004281 10.929227 -60.4298202
H 18.1645092 10.6855659 -61.9607414
H 18.8686102 10.0894265 -60.4738383
C 18.2749524 7.8099776 -59.110778
H 18.7708445 8.7035771 -58.7336403
H 18.9412565 7.2714474 -59.7855677
H 18.0370505 7.1684199 -58.2616609
N 15.8801458 11.0823937 -63.3805037
H 15.5331748 11.5872138 -62.5688742

C 15.9623305 11.817571 -64.6442913
H 16.7952455 11.4316431 -65.2351664
C 16.2306405 13.3158319 -64.4223685
O 16.001996 13.8407697 -63.3256233
C 14.6780471 11.5721577 -65.4405948
H 14.5614297 12.3592306 -66.1828294
H 14.8143697 10.6397859 -65.9771394
C 13.3981278 11.4233831 -64.5967226
H 13.2409175 12.3213791 -64.0054631
H 13.4843556 10.5777558 -63.912386
C 12.2010246 11.1445692 -65.486179
O 12.4220538 10.9836834 -66.7062945
O 11.0711677 11.0577901 -64.952999
N 16.7127389 14.0282412 -65.4533237
H 16.8577185 13.566903 -66.349382
C 16.9273486 15.4814323 -65.3898288
H 16.3871383 15.8476574 -64.5193038
C 16.3447654 16.2645858 -66.5817988
O 16.1558116 15.7419953 -67.6835179
C 18.4109936 15.8096423 -65.1287159
H 18.8261429 15.0930979 -64.4212154
H 18.4804463 16.7986531 -64.677849
C 19.2719235 15.817902 -66.3805842
O 19.9812442 14.8726451 -66.670495
N 19.2629312 16.8803734 -67.1552989
H 19.7115882 16.7659892 -68.0471681
H 18.6938739 17.6945154 -66.9492704
N 16.1171737 17.5560555 -66.3461034
H 16.3244087 17.9218972 -65.4216912
C 15.6409335 18.5213459 -67.3311437
H 15.1563779 17.9880668 -68.1515813
C 16.8207639 19.3019001 -67.9050657
O 17.814395 19.5196466 -67.2156087
C 14.6025547 19.470826 -66.7156745
H 15.0457816 19.9951369 -65.8663646
H 13.7532703 18.8943344 -66.3687152
C 14.1049765 20.4776596 -67.7549781
O 14.0497 21.6808959 -67.4357038
O 13.8939207 20.0301459 -68.9036003
N 16.6932574 19.7127438 -69.1587056
H 15.7361606 19.6535177 -69.5221682
C 17.6417525 20.4891035 -69.9664808
H 18.1709276 19.7718774 -70.5775385
C 18.7397635 21.2279062 -69.1911158
O 19.6163392 20.5303848 -68.6231238
O 18.7905354 22.4280729 -69.0989548

C 16.8603575 21.4153103 -70.9236658
H 16.3055296 22.1240006 -70.3041713
H 17.5800612 21.9809911 -71.516076
C 15.8643249 20.7213124 -71.8783228
H 15.1037594 20.1933291 -71.2956912
C 15.1458514 21.7861056 -72.7164382
H 15.8579213 22.3362843 -73.3312233
H 14.6219113 22.4788679 -72.0561909
H 14.4058411 21.3071409 -73.3609419
C 16.5435476 19.7199673 -72.8232848
H 17.329409 20.2094041 -73.398468
H 15.8002973 19.3095617 -73.5106352
H 16.9602655 18.8885803 -72.2557724

**Supplementary Data 6. Cartesian coordinates for perivine docked into CrPeNMT+SAM.**

N -2.1385667 21.7753992 -15.9844882
H -2.8881317 22.3451992 -15.612073
H -2.4187043 21.4412796 -16.898962
H -1.952445 20.9940613 -15.3721405
C -0.8983927 22.6008534 -16.1048954
H -1.0805973 23.3575681 -16.8709468
C 0.33849 21.7897622 -16.5659561
O 1.2553163 22.327476 -17.1874333
C -0.6053818 23.3116885 -14.7771474
H 0.3668547 23.80077 -14.8479421
H -0.5567763 22.5711725 -13.9778747
C -1.639947 24.3835703 -14.4160168
H -1.396285 24.7758699 -13.426708
H -2.6273748 23.9290305 -14.3587376
S -1.7285502 25.7740286 -15.5733039
C -2.9534829 26.7844932 -14.6928753
H -3.1342375 27.7045542 -15.2488035
H -2.5798462 27.0332981 -13.6984196
H -3.8891334 26.2323095 -14.5953778
N 0.3673335 20.4886197 -16.2657559
H -0.4109708 20.1080689 -15.7405706
C 1.3938537 19.5859753 -16.7771625
H 1.3580658 19.6103407 -17.8672226
H 2.3753087 19.9422459 -16.4680576
C 1.2496557 18.1385969 -16.3224611
O 0.7986647 17.8537127 -15.2154758
N 1.7628938 17.2390447 -17.1522057
H 2.1121366 17.5432004 -18.0501647
C 2.0288584 15.8382482 -16.8338954
H 1.9028599 15.7027836 -15.7604402
C 3.4957816 15.5076051 -17.1394012
O 4.119702 16.1812749 -17.9609391
C 0.9990926 14.9223526 -17.5032517
H 1.2572628 13.8928438 -17.2749979
H 0.0423459 15.1358366 -17.026009
C 0.8149397 15.0455348 -19.020348
H 1.6157817 14.5050359 -19.5288645
H 0.8699863 16.0954136 -19.3176114
C -0.5554566 14.4754363 -19.4180847
O -1.297864 15.2053148 -20.1119062
O -0.875195 13.3478418 -18.9747065
N 4.1061184 14.7016468 -16.2632362
H 3.5442127 14.1102083 -15.6715657
C 5.5629684 14.6122266 -16.0983634

H 6.0446756 14.8186955 -17.0600251
C 5.9695982 13.2129132 -15.6736854
O 5.3463653 12.639753 -14.786269
C 6.0594875 15.5900786 -15.0182406
H 5.5966651 15.3285324 -14.0641537
H 7.1389711 15.4636292 -14.915048
C 5.7688522 17.0627619 -15.3210483
H 6.199365 17.3109927 -16.2919241
H 4.6914859 17.2284646 -15.351451
C 6.3694499 17.9615862 -14.2310701
H 7.4516987 17.8220036 -14.192695
H 5.9514537 17.6743696 -13.2638997
C 6.0413128 19.4366002 -14.4861744
H 6.4275873 20.029751 -13.6516003
H 4.9522862 19.5499365 -14.5026758
N 6.6253332 19.9107507 -15.7656037
H 7.6331593 19.818804 -15.7546612
H 6.3843249 20.8792368 -15.933991
H 6.2683797 19.3550584 -16.5356419
N 7.106455 12.7593095 -16.1705568
H 7.4956956 13.2759729 -16.9574346
C 7.5639831 11.383838 -16.027934
H 6.7220861 10.7521143 -15.7410336
C 8.6256294 11.3080201 -14.9245199
O 9.6775993 11.9537076 -14.9983873
C 8.0743326 10.8706348 -17.3888252
H 8.9570019 11.4416081 -17.6791626
H 8.3787829 9.8302572 -17.2664366
C 7.0169986 10.9370419 -18.5172246
H 6.1405201 10.371064 -18.1914962
H 7.409044 10.4393792 -19.4038394
C 6.5974706 12.3719046 -18.896483
O 7.4785217 13.2681533 -18.9084406
O 5.3745041 12.6072441 -18.9936412
N 8.3366279 10.5700175 -13.8495991
H 7.428984 10.1185224 -13.7997984
C 9.2537472 10.4628666 -12.7165948
H 9.411715 11.4704627 -12.3299929
C 10.6243327 9.9036971 -13.153139
O 10.6986872 8.9385328 -13.9069108
C 8.6037136 9.6350831 -11.6036772
H 8.4204313 8.619799 -11.9600852
H 7.655759 10.0896643 -11.3114457
H 9.2647124 9.595986 -10.737355
N 11.703524 10.5367378 -12.6721267
H 11.5296608 11.3030954 -12.0418231

C 13.1142625 10.3142897 -13.0702562
H 13.6516325 11.146085 -12.6173586
C 13.4243279 10.5119185 -14.5675031
O 14.4681525 10.0599575 -15.0339048
C 13.7765064 9.0619071 -12.4395625
H 14.845682 9.122573 -12.6445486
C 13.6265139 9.0675566 -10.9118445
H 12.5803642 8.9423089 -10.6318339
H 14.0018322 10.0070241 -10.506304
H 14.2061243 8.2486758 -10.485005
C 13.2962742 7.7038875 -12.9652798
H 12.2890587 7.4911381 -12.6080123
H 13.9637118 6.9192992 -12.6112727
H 13.2950871 7.7094786 -14.0550769
N 12.605294 11.3349009 -15.2378421
H 11.7369915 11.5814565 -14.7805248
C 12.7314974 11.833814 -16.6111805
H 11.8724211 12.4921825 -16.7495073
C 12.6154246 10.7757957 -17.7237248
O 13.1625965 9.6728211 -17.644985
C 13.9716254 12.728998 -16.7542726
H 14.876663 12.1276992 -16.6566322
H 13.9650252 13.5017634 -15.9855672
H 13.9695782 13.2008787 -17.7373648
N 11.9102878 11.1457732 -18.7984009
H 11.4901366 12.0606529 -18.8311174
C 11.6113386 10.2161504 -19.8849453
H 11.4683386 9.2575284 -19.3922061
C 12.7697246 10.041093 -20.8749029
O 13.4841971 10.9883786 -21.2103005
C 10.2956217 10.5834444 -20.6018401
H 9.5642602 10.8737195 -19.8541547
H 10.4613373 11.4376453 -21.2598264
C 9.7172444 9.4047201 -21.4196476
H 10.350257 9.2350742 -22.2929309
H 8.7362979 9.7002982 -21.7986705
C 9.5899082 8.089336 -20.6183328
O 8.4815808 7.7830257 -20.1380566
O 10.6205886 7.3866881 -20.4498584
N 12.8901662 8.8305339 -21.4204041
H 12.2017952 8.1247666 -21.1470783
C 13.7998552 8.4972353 -22.5092701
H 14.1771403 9.4236569 -22.9434863
C 13.0399482 7.741936 -23.6088019
O 12.2666108 6.8259826 -23.3460136
C 14.9936922 7.7225667 -21.9245386

H 15.5173435 8.3745069 -21.2235556
H 14.6001376 6.8863099 -21.3570609
C 16.0029798 7.189281 -22.9591753
H 15.4878043 6.568367 -23.6890142
C 16.7306364 8.3210255 -23.6896215
H 17.2504768 8.9574053 -22.9726436
H 16.0235264 8.9250527 -24.2541046
H 17.4549132 7.90003 -24.3877077
C 17.0509741 6.3236455 -22.2584349
H 17.5894864 6.9110609 -21.5149428
H 17.7563171 5.9395483 -22.9960493
H 16.5630007 5.4766885 -21.778768
N 13.2962296 8.1160457 -24.8611726
H 13.9627613 8.8548464 -25.0049569
C 12.7297022 7.4720004 -26.0467334
H 11.6552424 7.6561114 -26.0668372
C 12.963355 5.9483119 -26.0268958
O 14.0825531 5.4964434 -25.7932219
C 13.3669937 8.1475173 -27.2671617
H 13.1403425 9.2139322 -27.2266732
H 14.4513269 8.049724 -27.187207
C 12.9656811 7.6189319 -28.6307211
C 12.1494424 8.3971049 -29.4761888
H 11.769383 9.3526089 -29.1454154
C 11.8565891 7.9520163 -30.7798108
H 11.2562024 8.555452 -31.4472005
C 12.3673248 6.7178848 -31.2374061
O 12.1160781 6.2857259 -32.4969706
H 11.5991475 6.9488647 -33.0185918
C 13.1697677 5.9340538 -30.3847744
H 13.5492799 4.9896701 -30.7459872
C 13.4827043 6.3907892 -29.091865
H 14.1273567 5.791667 -28.4621886
N 11.8951892 5.1788313 -26.2599184
H 11.0304658 5.6427895 -26.4800882
C 11.8426092 3.7036502 -26.2137578
H 10.7811114 3.4599108 -26.2631227
C 12.3180866 3.0493723 -24.893297
O 12.588105 1.8486882 -24.8229072
C 12.4769853 3.1028593 -27.4818833
H 13.5610504 3.0598399 -27.3602348
H 12.2582179 3.7523311 -28.3311342
C 11.9188234 1.7098089 -27.8129528
O 10.6734795 1.5608478 -27.7641107
O 12.7303627 0.8270589 -28.169355
N 12.3786583 3.8077385 -23.7892188

H 12.1567416 4.7960764 -23.8565606
C 12.7236866 3.2518266 -22.4780661
H 13.5786598 2.5902866 -22.6181944
C 11.5733328 2.4149249 -21.9209309
O 10.5345585 2.9233827 -21.5059758
C 13.1586237 4.3717878 -21.5322934
H 14.0067916 4.8753226 -21.9927235
H 12.3398627 5.0838206 -21.4074609
C 13.5708913 3.8411374 -20.1519667
H 12.6903613 3.4453154 -19.6452956
H 14.3110593 3.0473288 -20.2626016
C 14.1595742 4.9730826 -19.3074994
H 13.477531 5.8273082 -19.3386531
H 15.1255757 5.265841 -19.7198963
C 14.3246592 4.5251755 -17.8558798
H 14.9647772 3.6393135 -17.8237746
H 13.3343658 4.2459127 -17.4769636
N 14.894404 5.6206571 -17.034387
H 15.8026652 5.9022239 -17.3743342
H 14.9585333 5.3479973 -16.0632593
H 14.281514 6.4323719 -17.0835563
N 11.8145313 1.1129027 -21.796752
H 12.636759 0.7428888 -22.2469702
C 10.915173 0.206206 -21.075752
H 9.9261211 0.3116122 -21.5233934
C 10.7879049 0.6022152 -19.5986625
O 11.7565775 1.0128309 -18.9601126
C 11.3345322 -1.270057 -21.22834
H 10.6296722 -1.8783407 -20.6606041
C 11.2418011 -1.7098514 -22.6941182
H 11.9608367 -1.1650701 -23.3087444
H 10.2395993 -1.5174528 -23.0780593
H 11.4468521 -2.7773516 -22.7753339
C 12.7458793 -1.5768076 -20.7095885
H 13.4907476 -1.0077613 -21.2656222
H 12.9570051 -2.6395233 -20.8284199
H 12.8214824 -1.3296274 -19.6506744
N 9.6090573 0.3641778 -19.0190877
H 8.8636369 0.042936 -19.61778
C 9.3030641 0.4995601 -17.5765515
H 8.2498576 0.2378134 -17.4888618
C 9.3464409 1.9102799 -16.9589934
O 9.2888891 2.018324 -15.7386553
C 10.0341763 -0.545724 -16.7016046
H 9.6693134 -0.4676256 -15.6770594
C 9.7794235 -1.9797159 -17.169852

H 10.215899 -2.1450359 -18.1540049
H 8.7079161 -2.1740235 -17.2032576
H 10.239578 -2.6699839 -16.4624216
O 11.4299275 -0.3613229 -16.691399
H 11.6667592 0.1678767 -17.4727227
N 9.2779939 2.9909397 -17.7450031
H 9.3809 2.8980378 -18.7455964
C 8.8924366 4.3186524 -17.2226461
H 9.484174 4.5459203 -16.3361664
C 7.3950813 4.3363773 -16.8420819
O 6.54833 4.1563548 -17.7157283
C 9.1737252 5.4060425 -18.2724269
H 8.8412563 6.3707971 -17.8859208
H 8.6096819 5.1895781 -19.1816057
O 10.5517975 5.4957348 -18.5958262
H 10.5894134 6.1754387 -19.3178923
N 7.0367162 4.4791013 -15.5515566
H 7.7637049 4.4744686 -14.8485576
C 5.6335974 4.372326 -15.0850094
H 5.0070584 4.6318068 -15.9400818
C 5.0839267 5.3356751 -13.9496445
O 4.1802507 4.8921545 -13.2441249
C 5.3501324 2.8742188 -14.7954481
H 5.6601937 2.2734319 -15.6495461
H 4.2770818 2.7385852 -14.6680957
C 6.0481724 2.3356799 -13.5563096
O 6.9355157 2.9397106 -12.9802825
N 5.6839717 1.1536912 -13.1130988
H 6.1721809 0.8278604 -12.2984982
H 4.9434468 0.6468731 -13.5561149
N 5.4209234 6.6581869 -13.7954432
H 6.1313111 7.0377768 -14.4037847
C 4.6727308 7.6690857 -12.8730958
H 3.6722803 7.2716519 -12.7031805
H 5.1665915 7.6425688 -11.902207
C 4.4605241 9.2667223 -13.2399755
O 5.4811072 9.6663177 -13.812121
N 3.2536409 10.1109433 -12.9872607
H 2.6474233 9.6614132 -12.3206553
C 2.3771526 11.4343133 -13.6922496
H 3.1658733 12.0715482 -14.0792353
C 1.0505375 12.8198719 -12.978889
O 0.738086 12.1140384 -12.0071596
C 1.9540762 10.4352165 -14.9544058
H 0.9329988 10.1097298 -14.746017
C 2.7417697 9.1177858 -15.5124699

H 3.7821969 9.3932306 -15.6797871
H 2.6711718 8.3582525 -14.7352443
C 1.9181844 11.302737 -16.1856385
H 2.9290002 11.590086 -16.4899791
H 1.3394983 12.1952761 -15.9637221
H 1.4171187 10.8031861 -17.0150758
C 2.3461044 8.3062146 -16.7841532
H 1.2838958 8.043668 -16.7411571
H 2.9411495 7.3919643 -16.8207085
H 2.5328117 8.8854332 -17.695951
N 0.2333932 14.4003043 -13.1834479
H 0.7437184 14.7653437 -13.9704046
C -1.2123799 15.955379 -12.9597387
H -1.0028175 16.3292374 -13.960482
C -2.5312399 15.1695335 -13.1010222
H -2.3258053 14.1092915 -13.2484796
H -3.0618878 15.2689053 -12.1555772
C -3.4175086 15.6977339 -14.2585728
H -3.2432872 16.7657087 -14.3985759
C -3.0783778 14.9714564 -15.5675978
H -3.2811317 13.903087 -15.4809576
H -2.0269104 15.1037593 -15.8111044
H -3.6757972 15.3801559 -16.3812018
C -4.9110054 15.512951 -13.9555661
H -5.1400087 14.4586235 -13.8201664
H -5.5040244 15.9052164 -14.7807417
H -5.172629 16.0607214 -13.0540348
C -2.0596333 17.9254177 -12.3430272
O -1.3832877 18.2475852 -11.374691
N -3.271554 19.2249776 -12.5340521
H -3.8017849 18.9469697 -13.3433045
C -4.1853409 20.9390259 -11.9780666
H -4.1201797 20.8378689 -10.8941409
C -5.8142116 21.9798425 -12.0685304
O -6.4577917 21.990306 -13.1238488
C -3.3553264 22.2522131 -12.3368714
H -2.7508435 21.9810248 -13.199742
H -3.9431639 23.1195955 -12.6376299
C -2.3935577 22.6639575 -11.2141059
H -1.5875927 23.2551301 -11.6539897
H -1.9417954 21.7737149 -10.7710014
C -3.1027948 23.4921229 -10.1365626
O -4.1338848 23.0043489 -9.6189633
O -2.6283562 24.6174618 -9.8712537
N -6.4475046 23.0719258 -11.2698861
H -5.8681384 23.316696 -10.4654281

C -7.9445496 23.8116708 -11.2481781
H -8.5313975 23.1447577 -11.8786809
C -8.4505552 25.3183528 -11.7282815
O -7.9562339 26.3541144 -11.2608184
C -8.5827225 23.7319299 -9.8142739
H -8.3098143 22.7644516 -9.3910452
H -8.1237068 24.4929663 -9.1820639
C -10.1128287 23.8553926 -9.699581
H -10.4295997 24.8665053 -9.9608487
H -10.3884539 23.7049408 -8.6526486
C -10.8425833 22.8299823 -10.5754585
O -10.9583405 23.1023724 -11.793048
O -11.2574013 21.7859035 -10.0265513
N -9.602907 25.5151302 -12.4609812
H -10.1970189 24.6947314 -12.6038089
C -10.0093518 26.7968804 -13.139427
H -9.6033406 27.6153906 -12.5465371
C -11.5373224 27.0770783 -13.3129338
O -12.3430523 26.1660776 -13.4651874
C -9.3158021 26.7963794 -14.5218177
H -8.2652178 26.5407625 -14.3723088
H -9.7616841 25.9988297 -15.1187812
C -9.3656862 28.1006173 -15.3396433
H -10.3980501 28.3527717 -15.5746801
C -8.7151959 29.2767986 -14.6053596
H -7.6855865 29.0272619 -14.3458923
H -9.2616884 29.5103425 -13.6946444
H -8.721151 30.1580805 -15.2468748
C -8.6202634 27.902131 -16.6598573
H -7.5714266 27.6723762 -16.4704162
H -8.6904418 28.808201 -17.2612677
H -9.0736317 27.0808258 -17.214682
N -11.9412789 28.3572898 -13.4297045
H -11.2467902 29.0828588 -13.3712445
C -13.3321225 28.7589476 -13.708797
H -13.9922764 28.0787705 -13.1669539
C -13.6973559 28.668285 -15.20173
O -13.2212429 29.4599915 -16.0166546
C -13.5976231 30.1744778 -13.1694646
H -13.3339687 30.2032292 -12.1112647
H -12.9526566 30.8857408 -13.6884859
C -15.0439728 30.6091374 -13.3172145
C -16.0007793 30.2075446 -12.3654889
H -15.7026398 29.6095012 -11.5164327
C -17.3497155 30.5734271 -12.5229045
H -18.084102 30.2600018 -11.7942258

C -17.7459906 31.3401123 -13.632773
H -18.784204 31.6156339 -13.7569758
C -16.7930155 31.742311 -14.5848431
H -17.0984245 32.326439 -15.4416514
C -15.4440982 31.3771052 -14.4276294
H -14.7165289 31.6734974 -15.1713742
N -14.6326318 27.780819 -15.5396291
H -14.9521194 27.1150618 -14.8471413
C -15.2390964 27.6886577 -16.8666391
H -15.735075 28.6312197 -17.0994544
H -14.4624243 27.5134405 -17.6111349
C -16.2666536 26.5596064 -16.957317
O -16.3953891 25.7524785 -16.0401148
N -16.9764725 26.4878918 -18.0819378
H -16.8289854 27.176356 -18.8044725
C -17.9575033 25.4408391 -18.3831007
H -17.7899232 24.5827215 -17.7304426
C -17.7792012 24.9927546 -19.8397808
O -17.7959146 25.8188285 -20.750926
C -19.3742105 25.9740925 -18.1133224
H -19.4072767 26.3626889 -17.0943744
H -19.5911057 26.7896238 -18.8053619
C -20.4418095 24.8784784 -18.2588574
H -20.2170614 24.0779836 -17.5490798
H -20.3816661 24.4589508 -19.2655484
C -21.873747 25.3850101 -18.0153021
O -22.0329252 26.4946208 -17.4568844
O -22.8044396 24.6367793 -18.3888393
N -17.5576422 23.6953514 -20.0649298
H -17.6282623 23.0479037 -19.293315
C -17.3269048 23.1256682 -21.396853
H -17.6295227 23.8524687 -22.1539443
C -18.173276 21.8705255 -21.6133325
O -18.4492201 21.1344351 -20.6665456
C -15.8303326 22.833808 -21.5971231
H -15.5031065 22.1074362 -20.8521036
H -15.2664698 23.755845 -21.4523805
C -15.5235885 22.3038595 -22.9769314
N -16.0573017 22.8083599 -24.1656877
C -15.6125519 22.0028993 -25.140073
H -15.854249 22.1181749 -26.1888167
N -14.8449621 21.02673 -24.6277346
H -14.429788 20.2755673 -25.1579691
C -14.7816493 21.1953241 -23.2621255
H -14.2791179 20.561661 -22.5447457
N -18.5185024 21.5818415 -22.8701808

H -18.1432063 22.1738025 -23.6009505
C -19.2288161 20.3661674 -23.257331
H -19.9116933 20.1067755 -22.4457962
C -18.2431347 19.2015468 -23.4437471
O -17.5034608 19.1367934 -24.4310812
C -20.0753688 20.6536823 -24.5101802
H -20.7622502 21.4721288 -24.2876809
H -19.4111275 20.9769363 -25.3134978
C -20.8899068 19.4416545 -25.0031907
H -20.2061397 18.632868 -25.252551
C -21.8900833 18.9305006 -23.9653095
H -22.5664824 19.7301243 -23.6638583
H -21.3716057 18.543429 -23.0898099
H -22.4721546 18.1090986 -24.387707
C -21.6558409 19.8178908 -26.2704486
H -22.3670983 20.6156827 -26.0549008
H -22.1966695 18.9460329 -26.6408658
H -20.9593117 20.149503 -27.0399621
N -18.305586 18.2264973 -22.5363312
H -18.9848484 18.3084584 -21.7879638
C -17.4987295 17.0022879 -22.56464
H -16.4445225 17.2767669 -22.5352472
C -17.7136686 16.1851165 -23.8479637
O -16.7324037 15.7061853 -24.4089253
C -17.8183622 16.1676056 -21.3118248
H -17.6392762 16.7679502 -20.4189096
H -17.1541192 15.3031549 -21.279748
C -19.2469521 15.6902349 -21.2993199
N -20.3555822 16.5291505 -21.1770985
C -21.409785 15.8002387 -21.5780685
H -22.404937 16.1941265 -21.7444035
N -21.0189659 14.5690479 -21.940586
H -21.5461555 13.999855 -22.611493
C -19.6585309 14.4792161 -21.7690455
H -19.0246639 13.6887216 -22.1457658
N -18.9342631 16.1416122 -24.3967885
H -19.7139306 16.4659898 -23.8383305
C -19.2306073 15.4622993 -25.6698397
H -18.9191364 14.4191048 -25.5943322
C -18.4740459 16.0915514 -26.8531044
O -17.994577 15.3816032 -27.732199
C -20.7422481 15.4827005 -25.9605596
H -20.9140981 15.0341016 -26.9402223
H -21.0845436 16.5175317 -26.0055241
C -21.5715667 14.7083567 -24.9337949
O -21.1731487 13.5796127 -24.57134

O -22.5721805 15.2767049 -24.4473625
N -18.2496141 17.4112082 -26.8323886
H -18.5716361 17.9329869 -26.0288096
C -17.4531297 18.1047217 -27.8489587
H -17.7977212 17.8135617 -28.8424323
H -17.5879439 19.1789559 -27.7304308
C -15.956921 17.7942072 -27.7454374
O -15.2888172 17.6280449 -28.7627028
N -15.439025 17.6201509 -26.5247796
H -16.0609943 17.7043393 -25.7294654
C -14.0743246 17.1266898 -26.3031017
H -13.3826248 17.7038566 -26.9186169
C -13.9256189 15.6534948 -26.724003
O -12.9564171 15.2990751 -27.3953963
C -13.702554 17.3422062 -24.8291262
H -13.7609572 18.4063683 -24.6038623
H -14.431762 16.8391893 -24.1953113
C -12.3186415 16.8515113 -24.4543145
C -11.189853 17.6482056 -24.7279804
H -11.3040053 18.609059 -25.2098121
C -9.9051194 17.1911998 -24.3777829
H -9.0312386 17.792005 -24.5808524
C -9.7460546 15.9368061 -23.7535044
O -8.5019852 15.5094553 -23.4137942
H -8.5099345 14.6569228 -22.9779248
C -10.8775819 15.1370117 -23.4808996
H -10.7593813 14.1747477 -23.0085044
C -12.161335 15.5959257 -23.8337612
H -13.0275541 14.9793858 -23.6353157
N -14.9268359 14.8201123 -26.423841
H -15.7014839 15.176822 -25.8741997
C -14.9828394 13.4197593 -26.8408227
H -14.0769896 12.9181517 -26.4986893
C -15.0310294 13.2711907 -28.3685764
O -14.316041 12.4302505 -28.9112801
C -16.1878629 12.7409781 -26.1724631
H -16.0987507 12.8425083 -25.0898935
H -17.1015123 13.2506904 -26.4752072
C -16.3194695 11.2716038 -26.5208461
C -15.679925 10.2960364 -25.7306851
H -15.1296517 10.5903453 -24.8487413
C -15.7427155 8.9385754 -26.1018896
H -15.2470437 8.1817352 -25.5143806
C -16.4327316 8.5582085 -27.2723703
O -16.4601685 7.2539103 -27.6505408
H -16.8895214 7.1591939 -28.5014005

C -17.087672 9.5346431 -28.0532165
H -17.603424 9.2540987 -28.9595907
C -17.035941 10.887789 -27.6716777
H -17.5222426 11.6367727 -28.2851494
N -15.7781968 14.1209507 -29.0785084
H -16.4213335 14.735133 -28.58639
C -15.8302485 14.0991621 -30.5458371
H -16.023302 13.0723458 -30.857743
C -14.4935853 14.5210283 -31.1785911
O -13.967663 13.81329 -32.0333803
C -16.994066 14.9750367 -31.0424011
H -17.90762 14.6865268 -30.519727
H -16.7819786 16.0183928 -30.8031416
C -17.238316 14.8508303 -32.5555673
H -16.3290961 15.1177248 -33.0992785
H -18.0038016 15.5779021 -32.8375267
C -17.6986865 13.4395358 -32.9540066
O -18.9264795 13.2539991 -33.097649
O -16.8225467 12.5548429 -33.0888592
N -13.837982 15.5729289 -30.6701081
H -14.2892367 16.1201696 -29.9440593
C -12.4898729 15.9648231 -31.1351682
H -12.5346458 16.1337827 -32.2113024
C -11.4694612 14.8360737 -30.9194148
O -10.6875397 14.5295424 -31.8238719
C -12.0510367 17.2817301 -30.4618222
H -12.2135606 17.2075124 -29.385962
C -10.5745282 17.6261432 -30.7001145
H -10.360449 17.6415137 -31.7693089
H -9.9326989 16.8899406 -30.2164525
H -10.3471552 18.6032195 -30.2737645
C -12.8768666 18.454303 -31.009272
H -12.6844 18.5843449 -32.0745506
H -12.6143381 19.3701809 -30.4802161
H -13.9403605 18.2682826 -30.8668087
N -11.5418689 14.1369177 -29.7810758
H -12.2068198 14.4377171 -29.0741423
C -10.7457172 12.934077 -29.5229422
H -9.6880299 13.1653715 -29.6509979
H -10.9128238 12.6231638 -28.4919327
C -11.1015855 11.7534031 -30.4368387
O -10.2052046 11.0525872 -30.9091058
N -12.3806552 11.5703557 -30.7665667
H -13.0669852 12.1981408 -30.3596135
C -12.8830576 10.4904933 -31.6335773
H -12.4536585 9.5465298 -31.299618

C -12.4546029 10.6930136 -33.0856704
O -11.8379689 9.7967565 -33.662114
C -14.4115139 10.3664975 -31.5108397
H -14.8837716 11.3195435 -31.7430037
C -15.0075931 9.281508 -32.405127
H -14.497612 8.3332624 -32.2385487
H -14.9184434 9.5764776 -33.4497101
H -16.0681325 9.1738969 -32.1753859
O -14.7191285 9.9990169 -30.1858573
H -14.6611214 10.8041515 -29.6445041
N -12.6061399 11.9050074 -33.6278135
H -13.1277568 12.5991368 -33.0943308
C -12.1108057 12.2899864 -34.96069
H -12.5626445 11.6330904 -35.7039491
C -10.5924574 12.1086371 -35.0606404
O -10.1118234 11.5017037 -36.0194828
C -12.5253388 13.7419652 -35.2846723
H -12.2047179 14.3950538 -34.4716908
C -11.9014053 14.2583629 -36.5896139
H -12.1485647 13.587198 -37.4129321
H -10.8182307 14.3273054 -36.4922998
H -12.2846337 15.2548982 -36.8119629
C -14.0473735 13.8569901 -35.4397281
H -14.390893 13.2481522 -36.2760117
H -14.3288191 14.8958381 -35.6094773
H -14.5551193 13.5168148 -34.5364626
N -9.8327613 12.5183483 -34.0376294
H -10.2691124 13.0158676 -33.2677802
C -8.3866883 12.2947826 -33.9964652
H -7.94803 12.713808 -34.9032965
C -8.0253687 10.7951613 -33.964585
O -7.1750255 10.3535353 -34.7385592
C -7.808095 13.0498299 -32.7940855
H -8.2322251 12.6640796 -31.8665257
H -8.0425363 14.1119177 -32.8768195
H -6.7249933 12.9258191 -32.7693519
N -8.7383063 9.9928143 -33.1690346
H -9.4399586 10.4216383 -32.5757625
C -8.5261026 8.5373236 -33.0519002
H -7.476906 8.3564513 -32.8190178
C -8.836892 7.7959867 -34.3557814
O -8.0721417 6.9214099 -34.7582541
C -9.3671494 7.960609 -31.9009351
H -10.4270834 8.1175921 -32.1029293
C -9.1210937 6.4714297 -31.6602582
H -8.0569718 6.2924769 -31.5037895

H -9.4663804 5.8883818 -32.5134126
H -9.6722286 6.1524028 -30.7752524
O -9.0287859 8.6085484 -30.6970051
H -9.4300372 9.4946216 -30.711502
N -9.8765834 8.2071453 -35.0894348
H -10.4751493 8.9342993 -34.7066042
C -10.218649 7.6484065 -36.4087586
H -10.3078933 6.565189 -36.3149444
C -9.0961985 7.9031619 -37.4291875
O -8.8107229 7.0319327 -38.2457573
C -11.5883493 8.2058925 -36.8724934
H -11.5800876 9.2895272 -36.7418409
C -11.8649231 7.9080697 -38.3605028
H -11.8182208 6.8336763 -38.5433383
H -11.1330206 8.4106419 -38.9928817
H -12.8465962 8.2794081 -38.6492608
C -12.7194695 7.6055139 -36.002287
H -12.8725628 6.5606242 -36.275951
H -12.4291906 7.6247014 -34.9527788
C -14.0547499 8.3529914 -36.1158443
H -14.469557 8.2558855 -37.1180749
H -13.9111854 9.4081356 -35.8800284
H -14.765007 7.9268607 -35.4065953
N -8.3937191 9.0388911 -37.3367375
H -8.6846259 9.7176446 -36.6427142
C -7.265239 9.3812153 -38.2229573
H -7.5533014 9.1179589 -39.2408252
C -5.9497281 8.6286735 -37.9283472
O -4.9741865 8.7465544 -38.6840646
C -7.0390962 10.899473 -38.1899562
H -8.0014138 11.4136505 -38.2093944
H -6.4800146 11.1924024 -39.0771632
O -6.2990547 11.3175345 -37.05805
H -6.6658562 10.914714 -36.2473093
N -5.8900761 7.8921141 -36.8100548
H -6.7249579 7.8243468 -36.2397125
C -4.6565738 7.3302429 -36.256341
H -3.8362244 8.0007177 -36.5123651
C -4.2800623 5.9408626 -36.8074929
O -3.1352983 5.5205919 -36.6491747
C -4.7709993 7.3240925 -34.7269169
H -5.5236596 6.6026775 -34.4107831
H -5.0474193 8.3167635 -34.3677705
H -3.8122191 7.0434425 -34.2890576
N -5.1800992 5.2659515 -37.5310566
H -6.0659794 5.7270073 -37.7010793

C -4.9863126 3.9392569 -38.1512939
H -5.0038412 3.1815968 -37.3671395
C -3.6467069 3.7571432 -38.8886477
O -2.9731921 2.7386014 -38.7264792
C -6.1697208 3.6740353 -39.1048049
H -7.0402153 3.4328757 -38.4935022
H -5.9499646 2.7970377 -39.7150907
C -6.5637775 4.8197352 -40.0201111
N -5.7699764 5.9207787 -40.3641479
C -6.5674691 6.752385 -41.049963
H -6.277414 7.7318812 -41.4028678
N -7.7926363 6.2244398 -41.1826337
H -8.5854946 6.6964794 -41.5940588
C -7.8129371 5.0107498 -40.5332027
H -8.6720512 4.3747877 -40.3736965
N -3.1845526 4.7824287 -39.6094995
H -3.8397485 5.5490973 -39.7402001
C -1.9094941 4.7790629 -40.3503428
H -1.9058746 3.9210138 -41.0259016
C -0.6755178 4.6000095 -39.4590681
O 0.3500762 4.1523355 -39.9634635
C -1.8108003 6.0655976 -41.1818839
H -1.9802055 6.9149351 -40.5182176
H -0.8101594 6.155225 -41.6088882
C -2.8366963 6.0737128 -42.3295833
H -3.7805566 5.6467642 -41.9998432
H -2.4675716 5.4691415 -43.158521
C -3.090666 7.5062857 -42.7975055
H -2.21557 7.8834898 -43.328123
H -3.2604295 8.1357064 -41.9227019
N -4.2898586 7.6146451 -43.6353146
H -5.0985667 8.0752973 -43.2163844
C -4.3632322 7.5927761 -44.9465972
N -3.4118636 7.0991039 -45.680707
H -3.2172373 7.5904414 -46.5573116
H -2.6560279 6.644648 -45.2205878
N -5.4028833 8.1065841 -45.5234574
H -6.0523492 8.6180683 -44.9202338
H -5.4423454 8.1830156 -46.5185052
N -0.7546542 4.8641941 -38.1528525
H -1.6478001 5.1514024 -37.7665882
C 0.3441051 4.6164829 -37.2169392
H 1.2323201 5.1264964 -37.5885598
C 0.6996833 3.1210416 -37.0906775
O 1.8637562 2.7984179 -36.8587498
C -0.0133202 5.2297426 -35.858726

H -0.9017486 4.748926 -35.4486878
H -0.2065543 6.2971976 -35.9704315
H 0.8169899 5.0909097 -35.1647239
N -0.2423564 2.2117153 -37.3765593
H -1.1831707 2.5200405 -37.5942453
C 0.0441412 0.7786215 -37.4595675
H 0.5673747 0.4724402 -36.5515695
C 0.9552045 0.4341165 -38.6550826
O 1.7906982 -0.4590169 -38.5465271
C -1.2872208 0.0221727 -37.522223
H -1.8167315 0.2672544 -38.4447155
H -1.9079066 0.2944651 -36.668599
H -1.1024213 -1.0504495 -37.4916568
N 0.8781965 1.1977947 -39.7544587
H 0.2102942 1.9555281 -39.7611379
C 1.8112123 1.0873253 -40.8905783
H 2.0039892 0.0301488 -41.0804667
C 3.1532747 1.7347408 -40.5479506
O 4.1918602 1.1211358 -40.7786454
C 1.2242149 1.7004207 -42.1796402
H 1.0712918 2.7691655 -42.0362597
C 2.1680245 1.5163423 -43.3761676
H 2.3959256 0.458749 -43.5151264
H 3.0968456 2.0616945 -43.2111974
H 1.7047064 1.9050683 -44.2829471
C -0.1270984 1.0684292 -42.5415657
H -0.0134202 -0.0094525 -42.6618157
H -0.4995357 1.4948188 -43.4727939
H -0.8607535 1.2658962 -41.7604837
N 3.1497115 2.9075864 -39.8983128
H 2.2584563 3.3646896 -39.7477886
C 4.3829764 3.5769137 -39.4271644
H 5.0316617 3.7532182 -40.287047
C 5.1714679 2.6870662 -38.4572727
O 6.3914124 2.6466122 -38.5342007
C 4.0854947 4.9447377 -38.7734971
H 3.4492565 4.7868126 -37.9038288
C 5.3579813 5.667574 -38.3077099
H 6.0473034 5.7700222 -39.1479344
H 5.8520541 5.0988091 -37.5198768
H 5.1091859 6.6515776 -37.9116408
C 3.3657123 5.9094878 -39.7292705
H 4.0142422 6.146524 -40.5715909
H 3.1117469 6.8290265 -39.2030367
H 2.4524687 5.4645623 -40.1099666
N 4.5099699 1.8691594 -37.6261136

H 3.501018 1.9661057 -37.5679173
C 5.1768327 0.8861925 -36.7517641
H 5.9189905 1.4157616 -36.1524491
C 5.9655105 -0.1805465 -37.5278367
O 6.9655235 -0.6580613 -37.005066
C 4.1151826 0.2654933 -35.821984
H 3.6872668 1.0584642 -35.2070059
H 3.3115988 -0.1409395 -36.4366679
C 4.6026419 -0.8663175 -34.8987054
H 4.9701494 -1.6946528 -35.5041773
H 3.7425868 -1.2253198 -34.3369632
C 5.691348 -0.4487287 -33.8979481
H 5.305648 0.3329929 -33.2411631
H 6.5347315 -0.0299299 -34.4515579
N 6.1852786 -1.5988657 -33.1065998
H 7.0302672 -2.023504 -33.4565587
C 5.6931212 -2.0929878 -31.9796119
N 4.6137617 -1.6238221 -31.4121932
H 4.2496792 -2.0315972 -30.5644291
H 4.1500759 -0.8388203 -31.8268032
N 6.2826626 -3.0960025 -31.392016
H 7.158683 -3.4460252 -31.7437805
H 5.9220268 -3.4384356 -30.5210906
N 5.5130667 -0.5786026 -38.7196586
H 4.6989518 -0.1098716 -39.0946508
C 6.155844 -1.6215128 -39.5477068
H 6.7171868 -2.2989085 -38.9044692
C 7.1870424 -1.0471198 -40.5141252
O 8.1672071 -1.7115576 -40.8379074
C 5.0729423 -2.429844 -40.2970744
H 4.4719049 -1.7392024 -40.8919679
C 5.6505611 -3.4778044 -41.2668926
H 6.3652394 -4.1194898 -40.7503349
H 6.1577817 -2.9777053 -42.0922651
H 4.8537515 -4.0898952 -41.6888069
C 4.1272498 -3.1381766 -39.3001663
H 3.4521743 -2.3981917 -38.8745679
H 3.519856 -3.8567523 -39.8472324
C 4.8022508 -3.8790266 -38.1377464
H 5.1620025 -3.1674568 -37.3956499
H 5.6444629 -4.4611102 -38.5038849
H 4.0797422 -4.5405924 -37.6627072
N 6.9634562 0.1827065 -40.9660073
H 6.1059518 0.645179 -40.6903566
C 7.8918005 0.9019409 -41.8307362
H 8.3560989 0.1968688 -42.5199967

C 9.0541694 1.5083209 -41.0392823
O 10.1751688 1.4175308 -41.5113396
C 7.0910835 1.9372281 -42.6425295
H 6.4755174 2.5141194 -41.9494156
C 8.015055 2.9405889 -43.3353853
H 8.8170607 2.4130698 -43.8517664
H 8.4436482 3.6079035 -42.5859543
H 7.4537262 3.5413332 -44.0376991
C 6.1329575 1.2547909 -43.6511673
H 5.3388 0.7519175 -43.1009127
H 5.6559528 2.0240523 -44.2518133
C 6.7607953 0.2206656 -44.5975052
H 7.0017951 -0.6915304 -44.0477814
H 7.6692296 0.6123557 -45.0487245
H 6.0481605 -0.0291608 -45.3837918
N 8.798992 1.9819869 -39.8195457
H 7.8269675 1.9509433 -39.5448266
C 9.6149082 2.8267209 -38.9312898
H 9.2170332 2.5797365 -37.9459199
C 9.3250434 4.3409519 -39.0306445
O 8.5238253 4.7993794 -39.8542185
C 11.099113 2.4020543 -38.8279642
H 11.7082873 3.1408157 -39.3434734
H 11.2510327 1.4269978 -39.2904088
C 11.5965131 2.3038259 -37.3833269
O 11.4492724 3.3205405 -36.6703128
O 12.1249463 1.2330349 -37.0094698
N 9.8588958 5.1058401 -38.0754761
H 10.516748 4.6466534 -37.4396315
C 9.3325992 6.4041998 -37.6512809
H 8.3526531 6.2042132 -37.2199303
C 9.1395831 7.4234129 -38.7896545
O 10.0494829 7.7492656 -39.5515223
C 10.2024572 6.9760756 -36.5200309
H 10.3373002 6.2059069 -35.7593743
H 11.1814432 7.2466722 -36.9189787
C 9.5601829 8.2096529 -35.8657287
H 8.5933201 7.9247972 -35.443284
H 9.3792816 8.9717041 -36.6261681
C 10.4478246 8.8008571 -34.7647909
O 10.8020372 8.0456266 -33.8314376
O 10.7428137 10.012847 -34.8410345
N 7.9356494 8.0039627 -38.8202964
H 7.2747294 7.6949909 -38.1276181
C 7.4979327 9.066309 -39.7246913
H 6.4475177 9.2219085 -39.4784392

C 7.4607978 8.7250306 -41.2328113
O 7.2182063 9.6195186 -42.0501456
C 8.189382 10.3821855 -39.339387
H 9.2533861 10.3291555 -39.5703739
H 8.0772996 10.5585665 -38.2684137
H 7.7299772 11.2135753 -39.8729389
N 7.5456185 7.4423073 -41.6155564
H 7.7541 6.7354562 -40.9135248
C 7.344726 6.9801426 -42.997353
H 6.9802142 7.8289962 -43.5714683
C 6.2228111 5.9398072 -43.1423331
O 5.8136015 5.298277 -42.1811172
C 8.7040589 6.5417867 -43.5937623
H 9.3769214 6.2789443 -42.781539
H 8.5857406 5.6690598 -44.2357264
C 9.2911549 7.7100855 -44.425368
H 9.1907816 8.6410518 -43.8675988
C 10.7676023 7.5379856 -44.7641775
H 10.9094523 6.6899523 -45.4326336
H 11.347273 7.4053009 -43.8535032
H 11.1408564 8.4293545 -45.2677901
C 8.5546316 7.8484516 -45.7653518
H 8.6226347 6.9085588 -46.3141622
H 9.0244539 8.6267692 -46.3653165
H 7.5122898 8.121164 -45.6183874
N 5.6747873 5.8134553 -44.3622881
H 6.051826 6.3868802 -45.1032308
C 4.5891328 4.8649351 -44.6980872
H 4.5631062 4.1139725 -43.9051263
C 4.8447935 4.0459821 -45.973334
O 3.9017478 3.6498304 -46.6520897
C 3.2105171 5.5603428 -44.666108
H 2.4533252 4.7738203 -44.6831793
H 3.1054279 6.0864467 -43.7153432
C 2.9161759 6.5454592 -45.8179066
H 3.1261552 6.0697121 -46.7735315
H 3.5448542 7.4315362 -45.7241825
C 1.4308664 6.9484155 -45.7776237
H 1.2590212 7.5693356 -44.8966521
H 0.8212512 6.0482975 -45.6740764
N 0.981203 7.6811794 -46.9780385
H 0.7524881 8.6673509 -46.8923419
C 0.6228584 7.177358 -48.1443321
N 0.8186153 5.929421 -48.4685675
H 0.5752193 5.6132175 -49.3870961
H 1.388143 5.3577715 -47.8677134

N 0.0406873 7.949872 -49.0092621
H -0.2933387 8.855375 -48.6610897
H -0.2623347 7.6163466 -49.9005672
N 6.1190758 3.8578277 -46.3209588
H 6.8301691 4.1363819 -45.6635574
C 6.5958218 3.2428633 -47.5626859
H 5.830005 2.5719029 -47.9542416
C 7.8916732 2.4367347 -47.343168
O 8.5668267 2.64766 -46.3401455
C 6.8674228 4.3491416 -48.5851797
H 7.7970464 4.8464069 -48.304794
H 7.032437 3.8770175 -49.5533758
C 5.793543 5.4093771 -48.7466088
C 4.5694425 5.071977 -49.3430479
H 4.3617759 4.0434651 -49.5976387
C 3.6563468 6.0786253 -49.6922932
H 2.7837729 5.8310304 -50.2772817
C 3.9157208 7.4151039 -49.3467661
H 3.2349086 8.196869 -49.6514519
C 5.0976506 7.7468283 -48.6644446
H 5.3156042 8.7854837 -48.4512446
C 6.0470079 6.7471097 -48.3865945
H 6.9997683 7.0245668 -47.966154
N 8.2414517 1.5676536 -48.3001291
H 7.6597218 1.5224626 -49.118932
C 9.3979198 0.6578697 -48.2914649
H 9.1016447 -0.2646165 -47.791358
C 10.6602547 1.1969945 -47.5742744
O 11.1786285 2.245083 -47.9427162
C 9.7087369 0.3224341 -49.75895
H 9.967396 1.234773 -50.3010873
H 8.8483582 -0.1504811 -50.2328802
H 10.5608699 -0.3531544 -49.8204047
N 11.1058984 0.4683517 -46.5404187
H 10.6020477 -0.3756197 -46.3181145
C 12.3498292 0.6411544 -45.7620579
H 12.0842348 0.2321049 -44.78769
C 12.7677959 2.0582833 -45.3295639
O 13.9257997 2.2966503 -44.965395
C 13.488643 -0.2542563 -46.2966031
H 14.4106292 0.3125754 -46.3885813
H 13.233996 -0.5667301 -47.3062372
C 13.7404825 -1.5012609 -45.4212878
O 12.782663 -1.9446545 -44.7479203
O 14.8558588 -2.0616043 -45.4835954
N 11.7676098 2.9221305 -45.1175289

H 10.8663841 2.6705479 -45.5032949
C 11.8434355 4.1137479 -44.2565654
H 10.9702733 4.7131008 -44.4953178
C 13.0474841 5.0013958 -44.5690478
O 13.2566867 5.4146228 -45.7033659
C 11.6839069 3.7215239 -42.7654516
H 10.7042933 3.2772 -42.7433744
C 12.7172875 2.6995277 -42.243958
H 13.7411237 3.021697 -42.4103113
H 12.5627012 1.7344082 -42.7242798
H 12.5786212 2.5252549 -41.1832717
C 11.6078686 4.8907969 -41.7666777
H 12.5088229 4.963613 -41.1569353
H 10.7748965 4.7152793 -41.0967139
H 11.4403098 5.837781 -42.2673589
N 13.7997814 5.3551081 -43.5357305
H 13.5427582 4.9757027 -42.6392564
C 14.9722896 6.1999847 -43.5773004
H 14.7513394 7.1316738 -44.1038787
C 16.1325397 5.5277022 -44.3124657
O 16.971215 6.2121722 -44.8791594
C 15.3363167 6.5059358 -42.1167684
H 14.5993989 7.1982355 -41.7073855
H 15.2826306 5.5845827 -41.5328418
C 16.7149966 7.0988042 -41.9408267
C 16.9780411 8.3984896 -42.403626
H 16.1908624 8.9809401 -42.8654804
C 17.7536204 6.3218351 -41.3924162
H 17.5600729 5.3123153 -41.0611771
C 18.2759221 8.9240865 -42.299878
H 18.4802673 9.9218835 -42.6403416
C 19.0546992 6.8452967 -41.3078177
H 19.8555147 6.2433117 -40.90308
C 19.314826 8.149128 -41.7601691
H 20.3159865 8.5514257 -41.6963891
N 16.2058513 4.196207 -44.3268838
H 15.3924863 3.6449569 -44.0809636
C 17.3365519 3.5317345 -44.9880782
H 18.247824 4.0148928 -44.6328091
C 17.3362498 3.7162578 -46.5202865
O 18.411362 3.6889561 -47.1212283
C 17.4139431 2.0551039 -44.544089
H 17.2640812 2.0186564 -43.4640312
C 18.7685605 1.4097662 -44.8475106
H 18.7772956 0.3945679 -44.4540022
H 19.5699411 1.9849547 -44.3903993

H 18.9200973 1.3641078 -45.9264033
O 16.4573901 1.2286298 -45.1599023
H 15.5577683 1.5969757 -45.0914068
N 16.1818698 4.0690221 -47.1078517
H 15.3465728 4.1192239 -46.5422669
C 16.0387294 4.3659216 -48.539581
H 16.9320402 3.9392428 -48.9945931
C 16.0907108 5.8966401 -49.0803814
O 16.0794458 5.9767522 -50.2908172
C 14.8556147 3.4957892 -49.0773819
H 13.9798023 4.1366509 -49.1933825
H 14.5968292 2.7346661 -48.3378472
C 15.1275989 2.7525254 -50.4024108
O 14.1400315 2.2286328 -50.965854
O 16.307982 2.6299838 -50.8096799
N 16.2900343 7.1238414 -48.4203981
H 16.4497442 7.0131143 -47.4277553
C 16.1613041 8.688626 -48.8951202
H 15.5643498 8.6270493 -49.8051702
C 17.4543756 9.8556596 -49.3389581
O 18.4787587 9.2741032 -49.0634247
C 15.1839224 9.2297703 -47.7945255
H 14.6070166 8.4267525 -47.3302822
H 14.4664933 9.8869038 -48.2885934
C 15.9020718 10.0680588 -46.713811
O 16.3872989 11.1886415 -47.009515
O 15.9809798 9.6471257 -45.537687
N 17.6582507 11.2710311 -50.0187799
H 16.7432841 11.6473397 -50.215925
C 18.9196021 12.4999217 -50.4996121
H 19.4239745 12.7072473 -49.557024
C 19.8910586 11.6284874 -51.3214552
H 19.3040143 10.926469 -51.916006
H 20.4598406 12.2416738 -52.0211191
C 20.8996192 10.8577388 -50.4511839
H 20.4908512 10.6758339 -49.4595413
H 21.799341 11.4600084 -50.3269145
C 21.2793501 9.5187183 -51.0747255
O 22.2065281 9.4215954 -51.8603748
N 20.5592735 8.459898 -50.7803827
H 20.794535 7.5910496 -51.2218482
H 19.7724068 8.5434667 -50.1355936
C 19.092366 14.182846 -51.3913657
O 18.0971829 14.350783 -52.0856275
N 20.1085432 15.3856543 -51.5493382
H 20.9260687 15.212857 -50.986138

C 20.2948739 16.9461427 -52.3131094
H 20.2227503 16.7196393 -53.3770916
C 21.4078328 18.3179152 -52.2406162
O 22.5468109 18.183726 -51.7988277
C 19.0686253 17.7464106 -51.9697146
H 19.2338989 18.2728335 -51.0391949
H 18.2005998 17.0932304 -51.9204808
H 18.8667864 18.4802587 -52.7438545
N 21.1702238 19.6749261 -52.6514902
H 20.2345851 19.8379585 -52.9935248
C 22.0024176 21.0115075 -52.4479166
H 22.8683875 20.7431257 -51.8441611
C 21.2832516 22.1579803 -51.6588977
O 20.098186 22.3880363 -51.8856444
C 22.5113401 21.7661131 -53.7349204
H 21.6390627 22.0773934 -54.3111731
H 23.0283835 22.6774737 -53.4283394
C 23.4507222 21.0203275 -54.6889031
H 22.9115143 20.1707727 -55.1106251
H 23.7236569 21.6915994 -55.5045992
C 24.7266258 20.5101083 -54.0066815
H 24.4530278 19.8220364 -53.2042071
H 25.2791937 21.3544152 -53.5879837
C 25.5984771 19.7681775 -55.0268131
H 25.909443 20.4728439 -55.8051529
H 24.9888066 18.9920389 -55.5006992
N 26.7854235 19.1515691 -54.3842775
H 27.3653918 19.8580064 -53.9512649
H 27.3408378 18.6543916 -55.0695705
H 26.4940335 18.4923601 -53.6723818
N 22.0141776 22.9790716 -50.8776924
H 22.9848334 22.7561847 -50.7481297
C 21.4363151 24.0595014 -50.0406279
H 20.6651534 23.5880877 -49.4217957
C 20.6932301 25.1571828 -50.8434591
O 21.1123859 25.4928957 -51.9501055
C 22.4954675 24.6021902 -49.0576117
H 21.9681595 25.0697176 -48.2281703
H 23.0446883 23.7611733 -48.6298199
C 23.4967813 25.6136439 -49.6395768
H 24.1308012 25.1185841 -50.3761876
H 22.9616084 26.426131 -50.1294037
C 24.3600993 26.2058559 -48.5122502
H 25.0695733 25.4548231 -48.159759
H 23.7129745 26.4715583 -47.6719088
C 25.1007745 27.47991 -48.9414417

H 25.6487684 27.8643149 -48.0751497
H 24.3538345 28.231996 -49.2207631
N 26.033051 27.2429212 -50.0717632
H 26.729265 26.5557402 -49.8134378
H 26.5058074 28.1019389 -50.3270792
H 25.5216444 26.9054137 -50.8778729
N 19.6653497 25.7902966 -50.2502701
C 18.5260935 26.4199977 -50.9302971
H 17.683172 26.2611953 -50.25573
C 17.9666354 25.9230463 -52.2825018
O 17.2120042 26.6858309 -52.8855268
C 18.8245867 27.9163968 -50.8994079
H 19.6064095 28.1600803 -51.6210341
H 17.9311673 28.5198536 -51.0658878
C 19.9982801 26.7186724 -49.1631419
H 19.6497361 26.3469128 -48.204326
H 21.0706298 26.8855675 -49.103357
C 19.3411503 28.0678737 -49.4726293
H 18.5032263 28.2293829 -48.7937525
H 20.0551531 28.8885143 -49.395854
N 18.1444302 24.6585658 -52.7170806
H 18.7575415 24.0313991 -52.2081714
C 17.2497154 24.0944877 -53.7603078
H 17.1809009 24.8305569 -54.5644565
C 15.8108259 23.9536811 -53.2184155
O 15.6116559 23.7733122 -52.0168994
C 17.7905208 22.7718561 -54.367871
H 18.2164828 22.1394345 -53.5915213
H 16.9504542 22.2274091 -54.799801
C 18.8275157 22.9966047 -55.490398
H 19.7490875 23.3914824 -55.0618205
H 18.4292468 23.7618966 -56.1540239
C 19.1471149 21.7322105 -56.331149
H 18.3107319 21.0347568 -56.2604309
H 20.0375569 21.2460875 -55.9308658
N 19.3493069 22.0762304 -57.7577649
H 19.2235844 23.0639289 -58.0004847
C 19.3902273 21.3013719 -58.8329725
N 19.5688706 20.0105413 -58.789513
H 19.5868354 19.4947518 -59.6476506
H 19.6328429 19.5190873 -57.9186205
N 19.2411043 21.8219077 -60.01574
H 18.7989704 22.7503794 -60.07748
H 19.2658129 21.2691169 -60.8598066
N 14.821947 23.9635607 -54.1171727
H 15.0640462 24.1295175 -55.0795782

C 13.4070457 23.7053603 -53.8023649
H 13.2318405 23.8314889 -52.7303655
C 13.0148305 22.2690653 -54.1901151
O 13.2650034 21.8554559 -55.3268438
C 12.5042714 24.70102 -54.554355
H 11.4657565 24.4018961 -54.4176821
H 12.7363243 24.6603459 -55.6166903
C 12.5918951 26.14774 -54.1111584
O 13.1869008 26.5106661 -53.1113018
N 11.9857542 27.0281791 -54.8750885
H 12.0663734 27.990201 -54.6088372
H 11.474667 26.7304553 -55.7012412
N 12.3104354 21.5618859 -53.3039725
H 12.1319232 21.9882772 -52.3959269
C 11.7798987 20.2088839 -53.5218785
H 12.1354009 19.8236533 -54.4750295
C 10.2499869 20.2006078 -53.5304971
O 9.636071 20.6928764 -52.5837385
C 12.2939867 19.2814334 -52.4104324
H 12.0003572 19.7111831 -51.4550013
H 13.3811827 19.2399207 -52.459261
C 11.7331357 17.8531895 -52.451972
H 10.6473428 17.8812916 -52.5280063
H 11.9562163 17.3702826 -51.5054196
S 12.3827862 16.813984 -53.7779941
C 13.8931229 16.2105975 -52.9728887
H 14.4627188 15.602152 -53.6752431
H 13.6285852 15.6018071 -52.1075691
H 14.5036321 17.0516099 -52.6454799
N 9.6561252 19.476022 -54.4810838
H 10.2356146 19.1014098 -55.228018
C 8.2618632 19.0294284 -54.4110718
H 7.7432147 19.5950109 -53.6411374
C 8.2012683 17.5367186 -54.0391612
O 8.6091906 16.6916482 -54.8315145
C 7.5848191 19.3377223 -55.7563497
H 7.6533027 20.404727 -55.9472126
H 8.1407942 18.8235216 -56.5354205
C 6.1094319 18.9161937 -55.8471269
H 6.0223168 17.8621598 -55.5987058
C 5.2008171 19.7105303 -54.9111302
H 5.2842959 20.7773402 -55.1160026
H 5.4789999 19.5191252 -53.8783082
H 4.1706631 19.3882301 -55.0433692
C 5.5958432 19.1157678 -57.2735893
H 5.6369121 20.1715894 -57.5396105

H 4.5696247 18.7588489 -57.3505647
H 6.2159663 18.5490014 -57.9673902
N 7.650806 17.207511 -52.8705797
H 7.2809473 17.9542867 -52.2913677
C 7.3710433 15.8279812 -52.4464654
H 8.0060151 15.1508264 -53.0126345
C 5.9027388 15.4793458 -52.7611271
O 4.9709185 16.075372 -52.2065813
C 7.75986 15.6405981 -50.9667627
H 7.1203425 16.2573012 -50.3370099
H 8.7862314 15.9905422 -50.8370111
C 7.6824443 14.1863395 -50.4760692
O 6.8708591 13.4090086 -51.019936
O 8.4087115 13.8653462 -49.5055914
N 5.6864177 14.5711102 -53.7189983
H 6.4974395 14.0713565 -54.0777662
C 4.3447916 14.1842873 -54.1843736
H 3.6345714 14.9349283 -53.8406842
C 3.9351687 12.8544228 -53.5605881
O 4.3232699 11.7831128 -54.0192536
C 4.2336867 14.1575708 -55.7201635
H 4.8568936 13.3737071 -56.1278179
C 2.7974381 13.8629456 -56.1479142
H 2.1426664 14.6095088 -55.703318
H 2.4969621 12.8683791 -55.8186253
H 2.7229068 13.8922377 -57.2343704
C 4.6461988 15.4961607 -56.3443168
H 4.0765308 16.3124174 -55.897042
H 4.4772019 15.482627 -57.4198195
H 5.7094427 15.6583488 -56.1699027
N 3.0257182 12.9205146 -52.5909151
H 2.717312 13.8404901 -52.2952782
C 2.6511322 11.7941466 -51.7378099
H 1.5919382 11.8736081 -51.495824
H 2.826334 10.8515972 -52.2505211
C 3.4378837 11.764253 -50.4292475
O 3.7783718 10.6859284 -49.9559292
N 3.6725557 12.9262538 -49.8125931
H 3.3218807 13.7634739 -50.2589932
C 4.5639542 13.1282033 -48.6574106
H 5.5763021 12.8435635 -48.9560907
C 4.2394702 12.317265 -47.379758
O 4.9944892 12.3378076 -46.4012804
C 4.557135 14.6320349 -48.3562011
H 4.9178299 15.1702423 -49.2356233
H 5.2327209 14.8285762 -47.5267004

S 2.8894722 15.2074225 -47.9074698
H 2.7061254 14.4032963 -46.8479299
N 3.0995178 11.6227276 -47.3420058
H 2.5537956 11.6005599 -48.1895828
C 2.6973692 10.7695608 -46.2284903
H 1.7296098 10.3306215 -46.4382888
H 3.4252333 9.964224 -46.1301078
C 2.5964225 11.5335775 -44.9096616
O 2.0756425 12.6458024 -44.8617237
N 3.1475683 10.9405853 -43.8409104
H 3.5995258 10.050913 -43.9813798
C 3.2644505 11.5676352 -42.5110625
H 2.5363332 12.3721322 -42.4922009
C 4.6024855 12.3215117 -42.2989558
O 4.9330438 12.7035656 -41.1837733
C 2.7685976 10.6219858 -41.3876643
H 3.329892 9.6866489 -41.3966698
H 2.9110664 11.1133101 -40.4234859
C 1.2482459 10.3497499 -41.5881163
H 0.7919812 11.3134804 -41.7977461
H 1.1127473 9.7337692 -42.4784474
C 0.4227669 9.7161131 -40.4391496
H 0.7314226 10.1512193 -39.4901884
H 0.5960085 8.639396 -40.4025918
C -1.0758004 10.0217816 -40.7095582
H -1.41602 9.4220592 -41.5561837
H -1.1388727 11.0716698 -41.0127608
N -2.0107233 9.8625097 -39.5598994
H -2.0996365 8.9127489 -39.2334567
H -2.9406933 10.1949484 -39.8135329
H -1.7309402 10.4529809 -38.7839952
N 5.3030928 12.6456133 -43.3932679
H 4.9509657 12.3158047 -44.2839729
C 6.263606 13.7539353 -43.4685514
H 6.339384 14.0373157 -44.5168193
H 5.8664254 14.6025745 -42.9108326
C 7.6966603 13.5217271 -42.9844836
O 8.5229441 14.4189115 -43.1446022
N 8.052298 12.3272621 -42.5032093
H 7.3432049 11.6119669 -42.3818921
C 9.4128024 12.0200965 -42.040962
H 9.6646061 12.6654394 -41.1981862
H 9.4474178 10.9834761 -41.7123813
C 10.481707 12.1834795 -43.1192423
O 11.5222019 12.7773601 -42.8556136
N 10.1750316 11.7843984 -44.3589289

H 9.2846575 11.3332788 -44.4955374
C 11.0186339 12.0413754 -45.5425943
H 11.9980506 11.588156 -45.3790969
C 11.2701527 13.5389522 -45.7486745
O 12.4236188 13.9518113 -45.8260238
C 10.4258214 11.3832015 -46.8083101
H 10.5905208 10.3099959 -46.7269139
C 8.9312833 11.6033541 -47.0463739
H 8.7099762 12.6632397 -47.1442538
H 8.3393666 11.1827496 -46.2344072
H 8.6361123 11.1131843 -47.9758533
O 11.0895781 11.8306891 -47.9597574
H 10.484578 12.3844576 -48.4701324
N 10.2252136 14.371186 -45.663101
H 9.3074535 13.9722939 -45.5446904
C 10.2857366 15.8236971 -45.8547703
H 10.7235008 16.0265557 -46.8268567
C 11.1106349 16.537419 -44.7650356
O 11.8863183 17.4526382 -45.0535196
C 8.8657689 16.3970441 -45.8856242
H 8.8986116 17.4030429 -46.3069571
H 8.4867539 16.4648006 -44.8668808
S 7.7163401 15.3759861 -46.8510702
H 8.4533831 15.1086649 -47.9512639
N 10.9632892 16.0999518 -43.508704
H 10.2825801 15.3646959 -43.3383485
C 11.7666912 16.5933009 -42.3777083
H 11.7210304 17.682167 -42.3558897
C 13.2337723 16.2149077 -42.5646649
O 14.1041332 17.0794444 -42.4631432
C 11.2181531 16.0596538 -41.0385742
H 11.0788588 14.9803609 -41.1088303
C 12.1583111 16.3454275 -39.8608403
H 12.413162 17.4044263 -39.8365978
H 13.0713173 15.7583224 -39.9544092
H 11.6747709 16.0658247 -38.9240883
C 9.8619149 16.7101419 -40.7311641
H 9.9738266 17.7906836 -40.6408162
H 9.465485 16.313601 -39.7953368
H 9.147258 16.4904463 -41.5236473
N 13.5086787 14.9554922 -42.9189931
H 12.7374487 14.300883 -43.0121126
C 14.8616324 14.4600437 -43.1763215
H 15.4605874 14.5795315 -42.2714439
C 15.5717593 15.257738 -44.2626702
O 16.6984701 15.6980688 -44.0520232

C 14.8119003 12.9696725 -43.5454865
H 15.47017 12.778969 -44.3885111
H 13.8159203 12.6800994 -43.8734669
C 15.2350372 12.1081566 -42.3887418
N 16.5503927 11.9810507 -41.9354251
C 16.4753185 11.2196844 -40.8299497
H 17.3253826 10.9192111 -40.2306378
N 15.2029599 10.8627996 -40.5836198
H 14.890693 10.2906702 -39.8085624
C 14.4037633 11.4233475 -41.5538219
H 13.3257702 11.3738953 -41.6274226
N 14.9078435 15.5075468 -45.387301
H 13.989292 15.091668 -45.5152898
C 15.5112271 16.2350314 -46.5011762
H 16.4944499 15.8005517 -46.6810838
C 15.788866 17.702243 -46.1455768
O 16.9395239 18.1374501 -46.2012692
C 14.6897345 16.0179734 -47.7882799
H 15.1327128 16.6236197 -48.5603665
C 14.7908711 14.5234165 -48.182398
H 14.1962195 13.8987777 -47.5198436
H 15.8220997 14.1781548 -48.1262874
H 14.4372692 14.3716629 -49.2011021
C 13.2366266 16.4975189 -47.7320752
H 12.7646062 16.0195718 -46.8976035
H 13.2059957 17.5711979 -47.5585925
C 12.4407379 16.1837724 -49.0033466
H 12.2244151 15.1164961 -49.062049
H 13.0189562 16.48687 -49.8741816
H 11.4970485 16.7230964 -48.9888032
N 14.8187234 18.4245126 -45.5793613
H 13.9008259 18.0120409 -45.4473034
C 15.0314858 19.8172963 -45.1857533
H 15.4248899 20.3646793 -46.0422485
C 16.0654803 19.9815299 -44.052507
O 16.775709 20.9872676 -44.0197683
C 13.6739593 20.4215669 -44.8288207
H 13.2487701 19.8884107 -43.9775321
H 12.9997992 20.3346726 -45.682005
H 13.7950387 21.4777533 -44.588073
N 16.1945128 19.0031707 -43.1417661
H 15.5550477 18.2125874 -43.1960906
C 17.1426889 19.0652184 -42.0165262
H 17.2682905 20.115006 -41.7540704
C 18.5555244 18.5806105 -42.3633946
O 19.5022617 19.2022987 -41.8920977

C 16.5356863 18.3408313 -40.8014493
H 15.5546198 18.7709715 -40.5929376
H 16.3868088 17.2899072 -41.0560764
C 17.3783728 18.4041283 -39.5137768
H 18.2761985 17.8007418 -39.655826
H 16.803368 17.9420041 -38.7110996
C 17.8159365 19.8120997 -39.0658109
H 18.4907804 20.2290409 -39.816017
H 18.3932875 19.7217441 -38.1440225
N 16.6897875 20.7491832 -38.8609668
H 16.5665419 21.4502419 -39.5716769
C 15.8807065 20.8193544 -37.8169131
N 14.9853566 21.7589121 -37.7435133
H 14.3645694 21.8061651 -36.956886
H 14.8726153 22.4170928 -38.4946404
N 15.9472023 19.9843426 -36.8189475
H 16.6241288 19.2455779 -36.8502505
H 15.2949912 20.0477172 -36.0589826
N 18.7300829 17.5290883 -43.1764286
H 17.9068943 17.0488226 -43.5306661
C 20.0684893 17.0390531 -43.5746573
H 20.7065707 16.9291063 -42.6964889
C 20.7865705 18.0002679 -44.5366372
O 22.0134416 18.0053269 -44.5586972
C 19.9813346 15.699411 -44.3219547
H 19.304658 15.8747004 -45.1616404
H 20.9671976 15.5209637 -44.7569981
C 19.5628823 14.3578569 -43.6784448
H 20.3290906 14.0366348 -42.971993
H 18.6129878 14.4437749 -43.1562454
C 19.4543242 13.3380914 -44.844005
H 20.418487 13.3271226 -45.354375
H 18.7167883 13.694725 -45.5669937
C 19.1362141 11.8730891 -44.4980398
H 19.490443 11.6480878 -43.4892399
H 19.686472 11.2395421 -45.2001729
N 17.6955314 11.5486037 -44.6343123
H 17.1272324 12.0169392 -43.9401613
H 17.4806865 10.5531329 -44.5600195
H 17.3202713 11.7308229 -45.5705452
N 20.0478271 18.6624719 -45.4364741
H 19.046923 18.5093921 -45.436498
C 20.6358853 19.3309775 -46.6085273
H 21.6749389 19.5607893 -46.3684505
C 20.0541746 20.7091617 -46.9622581
O 20.3152965 21.2210728 -48.0515675

C 20.6868561 18.354292 -47.7957786
H 21.0473326 18.879063 -48.6814314
H 21.4566581 17.6177888 -47.5613333
C 19.4373287 17.602644 -48.1667651
C 19.394892 16.1940631 -48.1271936
H 20.196105 15.6434021 -47.6560693
C 18.3451897 15.5113149 -48.7728262
H 18.2923324 14.4372359 -48.7484014
C 17.3896775 16.2497781 -49.5044677
O 16.5031357 15.6745726 -50.3589183
H 16.8175994 14.8393012 -50.723428
C 17.3695605 17.6457992 -49.4114359
H 16.6398896 18.1806343 -49.9827419
C 18.3848271 18.32481 -48.7295873
H 18.4231516 19.3990316 -48.7319403
N 19.3517895 21.3539171 -46.024161
H 19.1651048 20.8775033 -45.1539639
C 19.0384741 22.7911583 -46.0750765
H 18.3539854 22.9771147 -45.2457682
C 18.2660689 23.2374827 -47.3416898
O 18.3498463 24.3844502 -47.7803859
C 20.3254224 23.5929801 -45.7682459
H 21.0190659 23.4837652 -46.6030665
H 20.8007875 23.1682607 -44.8823023
C 20.0831759 25.0814136 -45.500918
O 19.0655738 25.4065578 -44.8420731
O 20.8655128 25.9146396 -46.0124582
N 17.4548563 22.3556176 -47.9295338
H 17.4238769 21.424189 -47.5465261
C 16.4823397 22.7281121 -48.9710723
H 16.9464183 23.4477338 -49.6473765
C 15.24281 23.3977265 -48.3725054
O 14.9522023 23.2457856 -47.1836795
C 16.0422905 21.5162724 -49.8130752
H 15.2603537 21.8499514 -50.4920367
C 17.2011516 21.0558416 -50.7125561
H 18.1143336 20.9147098 -50.1404907
H 17.4026366 21.8357731 -51.4439333
H 16.943333 20.1436236 -51.2459316
C 15.398919 20.4182088 -48.9405732
H 16.1665696 19.8238888 -48.4571155
H 14.7955406 20.8598812 -48.151009
C 14.4493492 19.5350595 -49.7478417
H 14.8862927 19.2531794 -50.7036183
H 13.5268763 20.0795328 -49.9194258
H 14.2201392 18.6408149 -49.1834476

N 14.4360211 24.010578 -49.2375354
H 14.7402083 24.0446147 -50.2085445
C 12.9993631 24.158699 -49.0076405
H 12.8175786 24.2573771 -47.9397338
C 12.2662528 22.9046082 -49.5103378
O 12.6107608 22.3634176 -50.5604784
C 12.5234879 25.4517901 -49.6859483
H 13.0406288 26.283309 -49.2048236
H 12.830272 25.44303 -50.731898
C 11.0112581 25.7338392 -49.6079951
H 10.8575365 26.7986213 -49.7828324
H 10.658697 25.5126149 -48.6002629
C 10.1424015 24.9839405 -50.6247752
O 10.5934761 24.3664138 -51.5757024
N 8.8434225 24.9954737 -50.4307784
H 8.2650384 24.4828781 -51.0906918
H 8.4331033 25.5005896 -49.6701504
N 11.2359635 22.4621827 -48.7844512
H 11.0578115 22.9049529 -47.8891113
C 10.3429926 21.383246 -49.214284
H 10.5949631 21.0920705 -50.235015
C 8.8711568 21.8264889 -49.2272665
O 8.3637704 22.3460571 -48.2286926
C 10.5732244 20.1618264 -48.309774
H 10.3332883 20.4190762 -47.2777976
H 11.6227969 19.8672222 -48.3609254
S 9.5400098 18.7650291 -48.8429061
H 8.3402798 19.3259352 -48.6493615
N 8.1565084 21.4877271 -50.299685
H 8.6580875 21.1050509 -51.0997332
C 6.6915622 21.4723115 -50.3796701
H 6.2622286 21.8728319 -49.4665579
C 6.2201094 20.031677 -50.5303368
O 6.4399128 19.4158451 -51.5664035
C 6.1943466 22.3420754 -51.5434893
H 6.7074003 22.060723 -52.4640156
C 4.6826819 22.2560971 -51.7615391
H 4.1511206 22.4652032 -50.8329504
H 4.408085 21.2625488 -52.117419
H 4.3832472 22.9787312 -52.5207642
O 6.4597851 23.693421 -51.2481149
H 6.0035097 23.908674 -50.423727
N 5.5750189 19.4858299 -49.5003679
H 5.4489048 20.0421054 -48.6609263
C 4.8934822 18.1915638 -49.5707935
H 5.3805521 17.5583986 -50.3075649

H 4.9561912 17.6920084 -48.6044006
C 3.4178491 18.3381219 -49.9451647
O 2.7499778 19.2457229 -49.4508098
N 2.8759735 17.4192718 -50.7447628
H 3.489359 16.728848 -51.172119
C 1.4218424 17.2768095 -50.9304672
H 0.9352328 17.8212694 -50.1261785
C 0.9560668 15.8305307 -50.7697592
O 1.6406314 14.8966804 -51.1867693
C 0.9136146 17.8905378 -52.2565359
H -0.1448096 17.6360649 -52.3435319
C 1.6387051 17.3248981 -53.5004349
H 2.6843131 17.6363338 -53.4916327
H 1.6106378 16.2363657 -53.4623999
C 0.9961788 19.4249524 -52.1858548
H 2.0365676 19.7521869 -52.1926341
H 0.5247676 19.7783184 -51.266841
H 0.4642133 19.8755679 -53.0218273
C 1.0052803 17.7519525 -54.8316522
H 1.1088672 18.8260369 -54.9781292
H -0.0515642 17.481976 -54.8436029
H 1.510511 17.2397663 -55.65089
N -0.2477491 15.6495421 -50.2260329
H -0.7465903 16.4668413 -49.8863459
C -0.9656272 14.3704334 -50.212057
H -0.6470092 13.784003 -51.0743246
C -2.4736961 14.6046495 -50.3361702
O -2.9923013 15.6108318 -49.855645
C -0.6345655 13.5626088 -48.9469876
H -1.035592 14.063649 -48.0667914
H 0.4483293 13.4718768 -48.8486547
O -1.2039403 12.2647033 -49.0661658
H -0.8230701 11.6633302 -48.3915009
N -3.189996 13.6624306 -50.9545782
H -2.693636 12.8593069 -51.3085801
C -4.6570956 13.7038033 -51.1109514
H -4.9453094 14.6681594 -51.5324737
C -5.3659256 13.5807981 -49.7505522
O -6.4750307 14.0756237 -49.5707605
C -5.0838402 12.5810381 -52.0946599
H -4.5868338 11.656628 -51.7934535
C -6.60297 12.3174183 -52.0927149
H -7.1427278 13.2367823 -52.3255887
H -6.9278242 11.951214 -51.1188279
H -6.8636599 11.5560362 -52.8261546
C -4.6265523 12.9470809 -53.5247128

H -5.2699191 13.744181 -53.8956398
H -3.6030739 13.3215718 -53.5000957
C -4.6555746 11.788384 -54.5300713
H -5.6777838 11.458972 -54.7113472
H -4.0625365 10.9539342 -54.1543668
H -4.2321241 12.1260136 -55.4769672
N -4.7271601 12.913471 -48.788543
H -3.775143 12.6272275 -48.9723444
C -5.2999645 12.5558781 -47.4919011
H -6.3581933 12.3426238 -47.6238231
C -5.12327 13.6827605 -46.4548777
O -3.9915082 13.9538372 -46.0403971
C -4.6452439 11.2446981 -47.0565748
H -3.587913 11.2809028 -47.3174804
H -5.1004324 10.4159779 -47.6001787
O -4.7366918 11.0128571 -45.6723862
H -5.6758702 11.0025864 -45.3613888
N -6.2181431 14.2734505 -45.9265974
C -6.1448261 15.2535177 -44.8392723
H -5.5037262 16.0768951 -45.1469994
C -5.5994107 14.6593758 -43.5365247
O -4.990551 15.3639187 -42.7414758
C -7.5767455 15.7683423 -44.6357585
H -8.056633 15.2366621 -43.8112928
H -7.5919532 16.8425494 -44.4489233
C -7.6005661 14.1249832 -46.3585443
H -7.6892693 14.0050037 -47.4364082
H -8.051605 13.2737373 -45.845865
C -8.2932888 15.4153499 -45.9362787
H -8.111043 16.1900229 -46.680453
H -9.3633916 15.2698172 -45.7853592
N -5.798928 13.3603836 -43.3244477
H -6.3334307 12.8472974 -44.0199305
C -5.3819279 12.5909009 -42.1533687
H -5.5086175 13.2194111 -41.2710266
C -3.889537 12.1953296 -42.1819728
O -3.2267231 12.2061379 -41.1417984
C -6.3412377 11.3905832 -41.9922851
H -7.3113502 11.7703061 -41.6675729
H -5.9575394 10.7498656 -41.1971473
C -6.5663179 10.5388814 -43.2538189
O -6.908747 11.078591 -44.3329274
O -6.3508923 9.3107568 -43.1812859
N -3.3008267 11.9683216 -43.3619274
H -3.8866016 11.9470974 -44.1933319
C -1.8430178 11.9831623 -43.5618261

H -1.3723892 11.257173 -42.9150736
C -1.2618897 13.3502526 -43.1982805
O -0.4067809 13.4546671 -42.3135894
C -1.4776396 11.6187214 -45.0103698
H -2.1218696 12.1563712 -45.7031098
H -0.4492902 11.9202012 -45.2048059
C -1.5826308 10.1109484 -45.2530965
H -2.542401 9.7508069 -44.8866851
H -0.8029225 9.6052696 -44.6769365
C -1.4265517 9.7523301 -46.7342513
O -2.1887154 8.9014817 -47.2378949
O -0.4387212 10.1870761 -47.3704002
N -1.8238071 14.3981459 -43.7998905
H -2.5652821 14.2183388 -44.4716944
C -1.3629664 15.7743681 -43.644951
H -0.2966211 15.7971953 -43.8619811
C -1.5011144 16.2671664 -42.1931932
O -0.5669668 16.8774303 -41.6842395
C -2.0937697 16.6432986 -44.6956286
H -3.1616424 16.4331521 -44.6208048
C -1.6453572 16.3062192 -46.1415492
H -1.8060816 15.2459139 -46.3228022
H -2.2857106 16.8442621 -46.8391984
C -1.9169274 18.1412204 -44.4519846
H -0.8624135 18.3648704 -44.4776171
H -2.351247 18.4235725 -43.4927015
H -2.4173335 18.7036014 -45.2397571
C -0.1850993 16.6247081 -46.5047295
H 0.502714 16.0852862 -45.8550906
H 0.005957 17.6948585 -46.4390428
H -0.0012286 16.3148047 -47.5317343
N -2.5671139 15.9256739 -41.4689525
H -3.3267024 15.4447242 -41.9411177
C -2.7735512 16.3191514 -40.0675664
H -2.7786111 17.4067291 -40.0129945
C -1.6436206 15.8404344 -39.1348382
O -1.1884872 16.6049745 -38.2795479
C -4.1712491 15.8244989 -39.6523631
H -4.8964345 16.3669418 -40.2600162
H -4.276831 14.7663523 -39.8970248
C -4.5547229 16.0502673 -38.1793378
H -5.6347657 16.1922723 -38.1337433
H -4.0852274 16.9595111 -37.8033673
C -4.2106249 14.87867 -37.2602295
O -3.478791 13.9624486 -37.5900405
N -4.7524033 14.838826 -36.0651956

H -4.5608134 14.0108061 -35.5281391
H -5.3835633 15.5549238 -35.7602172
N -1.1004638 14.6424705 -39.3708469
H -1.5013084 14.0764096 -40.106404
C 0.0418472 14.1322373 -38.6116889
H -0.0436926 14.4501244 -37.5713118
C 1.3850099 14.6518156 -39.1239408
O 2.2660308 14.8908551 -38.3035319
C 0.0435491 12.6104591 -38.6527647
H 1.0465421 12.2488474 -38.4138007
H -0.2262749 12.3017169 -39.6598596
S -1.1117245 11.9445062 -37.4241918
H -2.0986728 12.845829 -37.6105998
N 1.5575739 14.8170369 -40.4417058
H 0.8054631 14.5690666 -41.0762493
C 2.7701594 15.422793 -40.987119
H 3.6444812 14.9045317 -40.5883586
C 2.8696392 16.8863013 -40.5636503
O 3.5847836 17.2487937 -39.6283336
C 2.7907183 15.3258836 -42.5238653
H 1.9156234 15.7985274 -42.9702768
H 2.7822393 14.2994419 -42.8465516
H 3.7036186 15.7788831 -42.9060338
N 2.1002966 17.7180647 -41.2791163
H 1.1970245 17.3057366 -41.4945638
C 2.6117307 18.5517644 -42.3861192
H 3.1341734 17.8723516 -43.0516016
C 1.3970127 19.0600589 -43.2242848
H 1.358215 18.4262558 -44.0974844
H 0.5018309 18.6692998 -42.7733033
C 1.1166488 20.5727843 -43.5444475
H 1.2915895 21.151315 -42.6635274
H 1.8411256 20.9728381 -44.2381746
C -0.3230066 20.9360951 -43.9914732
H -1.0227159 20.3720331 -43.3859128
H -0.4699022 20.6178342 -45.0179328
C -0.6963571 22.4298742 -43.8025919
H 0.1995555 22.9755746 -43.4898631
H -1.4260614 22.4979107 -42.9928553
N -1.2635441 23.0865246 -45.012343
H -0.5895041 23.0906335 -45.7804691
H -1.4992876 24.0597524 -44.8902245
H -2.0834365 22.6400856 -45.4285437
C 3.7642777 19.4606364 -41.9383207
O 4.6934606 18.9814363 -41.3095185
N 3.6311882 20.7771193 -41.9989438

H 2.8876535 21.1417414 -42.5604266
C 4.0788258 21.6000937 -40.8865894
H 5.1237588 21.3834285 -40.6673247
C 3.9999911 23.0820359 -41.3327063
H 3.0493725 23.2473995 -41.8409777
H 4.7944901 23.2717171 -42.0553756
C 4.1026739 24.1179475 -40.2399803
N 3.0918861 25.0331692 -39.9409579
C 3.4223691 25.5589047 -38.7527186
H 2.8010558 26.2448663 -38.192991
N 4.5752577 25.0358408 -38.3042514
H 4.8930887 25.1113105 -37.3429746
C 5.0364284 24.1390869 -39.2428601
H 5.8685618 23.4582498 -39.1352582
C 3.2420288 21.199852 -39.6385361
O 3.0797663 22.025223 -38.7500341
N 2.530232 20.0383135 -39.640792
H 2.865367 19.2416506 -40.1623869
C 1.4823966 19.8353562 -38.7041374
H 1.2481924 20.7922532 -38.2384218
C 2.0858318 19.0933861 -37.5361898
O 3.067777 19.6165891 -37.0050026
C 0.1413169 19.4928001 -39.3442083
H 0.1442248 18.4588372 -39.6817178
H -0.6141341 19.569777 -38.5611991
C -0.267998 20.4123699 -40.5175301
H 0.1183068 19.9509151 -41.3678312
C -1.7776922 20.4353344 -40.70414
H -2.2492346 20.8376265 -39.8066394
H -2.1313518 19.4171078 -40.8620015
H -2.0806715 21.041301 -41.5528518
C 0.1944014 21.8747087 -40.5369619
H 1.2344464 22.003532 -40.7787416
H 0.035205 22.3050513 -39.5470494
H -0.3843236 22.4742456 -41.2283432
N 1.4878269 18.020571 -37.0275607
H 0.680745 17.619781 -37.490007
C 1.8370793 17.567107 -35.682473
H 1.6480608 18.3923588 -34.994206
C 3.3255548 17.1909504 -35.5105984
O 3.975249 17.6990186 -34.5966676
C 0.897452 16.4265003 -35.2938252
H 1.0844826 15.5580817 -35.9233497
H -0.1416476 16.7397962 -35.4098183
H 1.0723946 16.150676 -34.2533891
N 3.895938 16.3935091 -36.4204664

H 3.3273487 16.0340792 -37.1822979
C 5.2891756 15.9425756 -36.3358269
H 5.4806631 15.6036171 -35.3165186
C 6.3276383 17.0397494 -36.6405866
O 7.4910534 16.9063308 -36.2670223
C 5.4618095 14.744717 -37.274927
H 5.2733635 15.0489023 -38.3063018
H 4.7617187 13.9547081 -37.0015401
H 6.47971 14.3605827 -37.1961688
N 5.9285687 18.1328717 -37.2982429
H 4.962632 18.1795496 -37.6016175
C 6.8314934 19.2418052 -37.635999
H 7.8429605 18.8505207 -37.6832907
C 6.844892 20.3829303 -36.6121256
O 7.5699517 21.3645417 -36.8076774
C 6.4883151 19.7654549 -39.0249614
H 5.4763035 20.1549587 -38.9918722
H 6.5452826 18.940506 -39.7349417
O 7.3909096 20.7802827 -39.4271918
H 7.5897174 21.3108912 -38.6363837
N 6.0301942 20.3258281 -35.5583977
H 5.4633655 19.4990513 -35.4131305
C 5.9102691 21.4436743 -34.6242259
H 5.8569889 22.3420848 -35.2375565
C 7.1639527 21.593333 -33.7492034
O 7.6308825 20.6298769 -33.149975
C 4.6035516 21.3149437 -33.8160339
H 3.9610644 20.5586151 -34.2705869
H 4.8261419 20.983383 -32.8007128
C 3.8146552 22.6339158 -33.7630863
H 4.3761121 23.361526 -33.1761685
H 2.8597389 22.454182 -33.2696363
C 3.5503843 23.2164719 -35.1512038
O 4.0029425 24.3035214 -35.4804174
N 2.9488639 22.4629638 -36.0411139
H 2.950617 22.7735293 -37.0102779
H 2.7918841 21.4817724 -35.8719266
N 7.7210904 22.8081187 -33.6914249
H 7.2987774 23.5567177 -34.2190106
C 8.9972279 23.0788989 -33.0131334
H 9.0701395 24.1482844 -32.8145984
H 9.0214978 22.5526099 -32.0586359
C 10.2493908 22.6746477 -33.805537
O 11.3286693 22.5814544 -33.2268775
N 10.1271567 22.3925806 -35.1093067
H 9.2142375 22.4390954 -35.5407744

C 11.2828804 22.1739636 -35.9765919
H 12.1470555 21.9182832 -35.3639204
C 11.6311631 23.4425166 -36.7703543
O 10.8468171 23.8739616 -37.6062429
C 11.0248787 21.0051895 -36.9419587
H 10.2282802 21.2982624 -37.627223
H 11.9359448 20.8776523 -37.5163873
C 10.6521745 19.6372191 -36.3484215
H 9.6886744 19.7031776 -35.8480046
C 10.554009 18.6256642 -37.4922608
H 11.5421073 18.4392663 -37.908598
H 9.9033941 19.0025286 -38.2786216
H 10.1402736 17.6890333 -37.1141196
C 11.700169 19.1190278 -35.363519
H 12.6761902 19.0615675 -35.8409176
H 11.4049478 18.1266817 -35.0192001
H 11.7434391 19.7788958 -34.4960605
N 12.8948299 23.8661966 -36.7091582
H 13.3352564 23.7065076 -35.812184
C 13.4155394 25.0179326 -37.4790076
H 12.6681205 25.8145446 -37.4282954
C 13.6732908 24.7387133 -38.9836209
O 14.3679382 25.4938641 -39.6582383
C 14.7076514 25.5493859 -36.8171534
H 15.5330047 24.8870948 -37.0856383
H 14.9254605 26.5347873 -37.2302733
C 14.686988 25.6625318 -35.2845744
H 15.5866248 26.1902592 -34.9596257
H 14.7320807 24.6561241 -34.8586402
C 13.4378158 26.3886491 -34.7758366
O 13.4280188 27.6368107 -34.819975
O 12.4900572 25.6633847 -34.4019596
N 13.2648524 23.5772005 -39.5105643
H 12.5354699 23.085907 -39.0068225
C 13.5570177 23.1795007 -40.8976494
H 14.5254876 23.6009869 -41.1761233
C 12.542468 23.7627457 -41.8860466
O 11.3445727 23.7642826 -41.6133993
C 13.6717175 21.6505914 -41.0511484
H 14.6737933 21.3409764 -40.7633607
H 13.5409867 21.3814689 -42.0977122
C 12.6807254 20.8409029 -40.234541
O 13.0819143 19.9571612 -39.4998137
N 11.4063889 21.1652132 -40.2593715
H 10.7663929 20.6171536 -39.7086994
H 11.0835354 21.9909525 -40.7547637

N 13.0133136 24.1123106 -43.0888082
H 14.003369 23.9861169 -43.252692
C 12.2559559 24.8063065 -44.14478
H 11.634147 25.5703974 -43.6725806
C 11.2900693 23.8636724 -44.8836361
O 11.4794884 23.5296958 -46.0528063
C 13.2402964 25.5207442 -45.0935189
H 13.7974318 24.7585126 -45.6299273
H 12.6747639 26.1075278 -45.8195301
C 14.2367 26.4457212 -44.363787
H 13.6782029 27.2076148 -43.8182901
H 14.8081668 25.8651489 -43.6389009
C 15.2530414 27.1445263 -45.2802328
H 15.9662655 27.6665972 -44.6388995
H 14.7410762 27.8844194 -45.8974409
C 16.0017683 26.1590706 -46.1895801
H 15.4383225 26.0365783 -47.1185131
H 16.0485386 25.1791219 -45.7028254
N 17.3833513 26.6013149 -46.4784672
H 17.4501689 27.5257558 -46.8646949
H 17.8394313 25.9286781 -47.0934372
H 17.9672793 26.5200239 -45.6378488
N 10.2586788 23.4106932 -44.1767312
H 10.1841574 23.7178249 -43.2124954
C 9.2046284 22.5279462 -44.6902692
H 9.4646982 22.2318761 -45.7066837
C 7.8664666 23.2561846 -44.773848
O 7.5173305 24.0732073 -43.9268036
C 9.0728533 21.2239852 -43.8778855
H 8.316634 20.6091786 -44.3642886
C 10.3766276 20.4230278 -43.8853184
H 11.1563743 20.9672999 -43.3545985
H 10.6901088 20.2389319 -44.9118237
H 10.2204614 19.4614051 -43.3955968
C 8.6444678 21.4437578 -42.4225151
H 9.3296493 22.1190396 -41.9159669
H 8.6253233 20.491972 -41.8909169
H 7.6444986 21.8751438 -42.3818422
N 7.0848025 22.9020958 -45.7835561
H 7.4542566 22.2468272 -46.4660422
C 5.688535 23.2901281 -45.9603944
H 5.2592379 23.5642908 -44.9971347
C 4.9368154 22.090787 -46.5162855
O 5.5175932 21.2799915 -47.2402207
C 5.572196 24.4861637 -46.9093264
H 6.1315988 25.3254024 -46.4927753

H 4.5253632 24.7775338 -47.0057321
O 6.0920421 24.166681 -48.1936715
H 6.9019288 23.6412406 -48.069159
N 3.6649475 21.9328174 -46.1572492
H 3.1924113 22.6509743 -45.6290061
C 2.8540294 20.891737 -46.7718369
H 3.2603949 20.7420939 -47.768523
C 1.4232672 21.3469321 -46.9993605
O 0.9655742 22.2461308 -46.2958749
C 2.8723023 19.5311039 -46.0727033
H 1.8998189 19.3593649 -45.6756641
H 2.831625 18.8431272 -46.9167144
C 4.0238134 18.9905664 -45.2169582
C 4.3071869 17.6253821 -45.3860567
H 3.6888653 17.0224533 -46.0309537
C 4.8919789 19.744078 -44.3979068
H 4.7530479 20.7996534 -44.2590226
C 5.3992881 17.0235059 -44.752768
H 5.5957283 15.9801922 -44.9285487
C 5.9619852 19.1264943 -43.7177794
H 6.5828412 19.700868 -43.0462872
C 6.2154413 17.761582 -43.8888187
H 7.0295482 17.2877361 -43.3583586
N 0.7049211 20.6843602 -47.9071269
H 1.1580329 19.9917536 -48.4999263
C -0.7426341 20.8387991 -48.0115702
H -1.0824204 20.9641876 -46.9898499
C -1.5361428 19.617803 -48.4852348
O -0.9924186 18.6595985 -49.0384563
C -1.11455 22.1121096 -48.8003637
H -1.7026334 21.8618073 -49.6844732
H -0.2187602 22.6483505 -49.119513
C -1.9431406 22.9948283 -47.8556361
O -2.8487189 22.4398679 -47.1824651
O -1.4945848 24.1167787 -47.540539
N -2.8489941 19.6723543 -48.2388324
H -3.2118883 20.5386921 -47.8451067
C -3.8093695 18.7281656 -48.8164363
H -3.3973647 17.7249245 -48.7310293
C -3.9734597 19.0752603 -50.2936677
O -4.3973474 20.175993 -50.6415666
C -5.1732061 18.7435119 -48.0992768
H -5.6730347 19.6950951 -48.2825975
C -6.0637773 17.6068255 -48.6193746
H -5.6052813 16.639463 -48.4108445
H -6.2151491 17.7031978 -49.6940242

H -7.0395154 17.6576898 -48.1394967
C -5.0359728 18.5600012 -46.5826229
H -4.4763312 17.6507279 -46.373113
H -6.0235974 18.4938287 -46.1264681
H -4.5200769 19.4177832 -46.1531065
N -3.6215795 18.1436875 -51.1741785
H -3.2879382 17.2507786 -50.8282856
C -3.6115106 18.3903012 -52.6077545
H -4.5005105 18.9612831 -52.879691
H -2.7342712 18.9845956 -52.864576
C -3.5764643 17.1062873 -53.4192378
O -2.760003 16.2192524 -53.1788795
N -4.4347018 17.0516582 -54.4326559
H -5.0732818 17.8204822 -54.5581563
C -4.3864658 16.05468 -55.4956002
H -4.0915978 15.0912157 -55.0883004
C -3.3390088 16.4662952 -56.5310674
O -3.4495113 17.5140222 -57.164842
C -5.7890218 15.9175663 -56.0942179
H -6.1264015 16.9003061 -56.4288134
H -6.4684305 15.5787826 -55.3090581
C -5.884318 14.9468145 -57.2730016
O -4.8383242 14.4918336 -57.7921059
O -7.0322555 14.7422459 -57.7062923
N -2.3237965 15.6294866 -56.7222922
H -2.3348741 14.7515165 -56.2290766
C -1.1959614 15.917138 -57.5968154
H -0.8042434 16.9026035 -57.3349828
C -1.5477935 15.9520276 -59.0940706
O -0.7986989 16.5414193 -59.8719076
C -0.1261968 14.8713849 -57.2991538
H -0.4332321 13.8964363 -57.6832249
H 0.0211865 14.8034629 -56.2217764
H 0.8044678 15.171372 -57.7763393
N -2.6894224 15.3899282 -59.5099429
H -3.3057519 14.9434825 -58.8268714
C -3.1680469 15.5081525 -60.8889374
H -2.2977069 15.4764396 -61.5426236
C -3.8714604 16.8578481 -61.1353749
O -3.8881519 17.3427345 -62.2717246
C -4.0589673 14.2899356 -61.2019821
H -3.5800509 13.3944349 -60.8022451
H -5.0094949 14.4145144 -60.6792547
C -4.3336983 14.054691 -62.7013104
H -4.8066073 14.9365606 -63.1323418
C -3.0607759 13.732287 -63.4923323

H -2.5395826 12.8897156 -63.037984
H -2.3987076 14.5952794 -63.5160859
H -3.3198034 13.4669686 -64.5171777
C -5.2831101 12.8667808 -62.868187
H -4.8269732 11.9632398 -62.4652122
H -5.5106869 12.7195113 -63.9246257
H -6.2135557 13.0676634 -62.3361046
N -4.3438173 17.5317856 -60.0767708
H -4.239743 17.0913676 -59.1689351
C -5.1392953 18.7670931 -60.1508346
H -5.3101043 19.014173 -61.1954728
C -4.4652452 20.0192241 -59.553186
O -4.7247452 21.1197762 -60.036646
C -6.5275032 18.4864972 -59.5562995
H -7.1041324 19.4099779 -59.5149897
H -6.4332717 18.0886791 -58.546576
C -7.2894454 17.5033969 -60.4354308
O -7.5563357 17.7986585 -61.5883171
N -7.5704357 16.2932746 -60.0008698
H -8.0536807 15.6856149 -60.632247
H -7.3300548 15.9419432 -59.0695682
N -3.5291447 19.8666245 -58.6129897
H -3.3778296 18.9328279 -58.2480178
C -2.7161973 20.948246 -58.0426512
H -3.3974747 21.6988151 -57.6398965
C -1.8381347 21.6244684 -59.108357
O -1.3603305 20.9679296 -60.0376461
C -1.8708616 20.3801509 -56.8857777
H -1.2155043 19.5960797 -57.2688558
H -2.5403692 19.9339514 -56.1488536
C -1.0113161 21.4298223 -56.1713507
H -0.2443703 21.7912918 -56.8567282
H -0.5059511 20.9491715 -55.3327881
S -1.9402459 22.8506813 -55.5394919
C -0.5701828 23.8292679 -54.8690719
H -0.9613234 24.74235 -54.4197375
H 0.1193026 24.0926016 -55.6715556
H -0.0410525 23.252321 -54.1098706
N -1.6015408 22.9369568 -58.9811233
H -1.9423219 23.4054305 -58.1486339
C -0.8192637 23.7390643 -59.9395053
H -0.2576772 23.064157 -60.5851844
C 0.2282464 24.5957342 -59.2270807
O 0.004776 25.0706294 -58.1183271
C -1.7509353 24.6002576 -60.822006
H -2.2485006 25.3361505 -60.1875853

H -1.1470927 25.1432575 -61.5512182
C -2.8355023 23.8041089 -61.576147
H -3.4692309 24.5095959 -62.1148217
H -3.4670243 23.293783 -60.85025
C -2.285634 22.7794283 -62.5838831
H -1.469829 22.2166229 -62.1362081
H -1.8727529 23.3160784 -63.440655
N -3.3480238 21.8600974 -63.0454371
H -4.1592275 22.2924263 -63.4494333
C -3.3712553 20.539867 -62.9367282
N -4.3632524 19.8552986 -63.436589
H -4.394121 18.8635073 -63.2050823
H -5.2093213 20.3214733 -63.7056537
N -2.4470361 19.8412767 -62.3456859
H -1.8072108 20.3142319 -61.7116381
H -2.5589807 18.8412342 -62.2636225
N 1.3477479 24.8147152 -59.9086339
H 1.3980397 24.4714973 -60.8561952
C 2.4801481 25.6382775 -59.4817666
H 2.171737 26.310206 -58.6796175
C 2.9540436 26.4820188 -60.6742369
O 2.5493109 26.2124622 -61.8048482
C 3.6101897 24.7291168 -58.9715134
H 3.9556652 24.0994097 -59.7921348
H 4.4519802 25.3525014 -58.6710504
C 3.2414508 23.8502424 -57.790578
C 3.2255769 24.3988016 -56.4940977
H 3.4578802 25.4439975 -56.3432923
C 2.944317 22.484486 -57.9805326
H 2.957988 22.0603603 -58.9749413
C 2.9118273 23.5871558 -55.3886793
H 2.893409 23.9989979 -54.390721
C 2.6376701 21.6672874 -56.8746023
H 2.4174114 20.6212776 -57.0198648
C 2.6196309 22.2218223 -55.5761608
O 2.3179465 21.4526177 -54.4997302
H 2.18435 20.5319836 -54.724826
N 3.8334992 27.4628261 -60.4509498
H 4.2082822 27.584594 -59.5219213
C 4.4939213 28.1620494 -61.5616058
H 3.7309422 28.5488971 -62.2384471
C 5.3882736 27.2020676 -62.3485732
O 6.0176283 26.3114741 -61.7616727
C 5.3217809 29.3477141 -61.0436929
H 6.091829 28.9980095 -60.3569351
H 4.6612574 30.029898 -60.5071673

O 5.9393037 30.0538829 -62.1032955
H 6.7243398 29.5613144 -62.4093461
N 5.5820288 27.4940815 -63.6317522
H 5.0106585 28.1958685 -64.0746187
C 6.7307802 26.9874338 -64.3760478
H 6.6366855 25.9085016 -64.4254029
C 8.028407 27.3558119 -63.6304401
O 8.1187891 28.4400932 -63.0417593
C 6.7479946 27.5450176 -65.8100117
H 7.719706 27.3149662 -66.2505881
H 6.6485008 28.6315395 -65.7731873
C 5.6672195 26.9697377 -66.7358325
O 4.772257 26.2412801 -66.2508656
O 5.7522914 27.2543315 -67.9504335
N 9.0046514 26.4438149 -63.5827792
H 8.8767 25.5757138 -64.0999271
C 10.2928452 26.6834173 -62.9168859
H 10.9628395 25.863104 -63.1596956
H 10.726691 27.5994604 -63.3176861
C 10.2581107 26.8168125 -61.3836782
O 11.1929088 27.3706979 -60.812188
N 9.2068629 26.3463782 -60.7000323
H 8.4713915 25.8971 -61.2280263
C 9.0731089 26.4374344 -59.2283582
H 9.1851045 27.4791121 -58.9258279
C 10.1069491 25.6136815 -58.4432941
O 10.4836519 25.994552 -57.3288833
C 7.6897912 25.9495096 -58.7796666
H 7.6527518 25.9341602 -57.6888603
H 7.5223822 24.9374057 -59.148062
O 6.6580577 26.7898418 -59.2502181
H 6.5249451 26.5892104 -60.1998395
N 10.5092653 24.4553591 -58.9775317
H 10.148396 24.2047745 -59.8925566
C 11.2703098 23.4359545 -58.2511685
H 11.53296 23.831252 -57.2725145
C 12.5826537 23.0539197 -58.9343996
O 12.7237566 23.0892687 -60.153064
C 10.4016771 22.1945978 -58.0069762
H 10.1678104 21.7295215 -58.9665107
H 10.9725225 21.4720044 -57.4234963
C 9.1174901 22.4918067 -57.2616053
C 9.1440574 22.7882535 -55.8861933
H 10.0770687 22.7663086 -55.3446341
C 7.8965917 22.5106125 -57.9559699
H 7.8822044 22.2996273 -59.0114969

C 7.9532069 23.1130515 -55.2127899
H 7.9771022 23.3415634 -54.1554797
C 6.7052395 22.8277978 -57.284616
H 5.7704859 22.841606 -57.8236995
C 6.7337942 23.135257 -55.9133376
H 5.8189192 23.3828901 -55.3936062
N 13.517583 22.5839715 -58.1188466
H 13.3072881 22.5350507 -57.1278463
C 14.8179288 22.0601079 -58.5444263
H 15.0314648 22.3351386 -59.5799209
C 14.8820061 20.5283609 -58.4571595
O 15.7122647 19.8905221 -59.1081171
C 15.861854 22.6970743 -57.6264017
H 15.4958654 22.6849266 -56.6000881
H 16.7664742 22.1004187 -57.6568063
C 16.1785143 24.1454572 -58.0345006
H 16.4935678 24.6931106 -57.143149
H 15.2907343 24.6425045 -58.4312527
C 17.305602 24.1476694 -59.067901
O 17.1190339 23.5958259 -60.1692975
O 18.44763 24.4781666 -58.6847995
N 14.0008482 19.9431131 -57.6415347
H 13.375073 20.5442361 -57.11928
C 13.8557196 18.5135042 -57.3961515
H 14.3000178 17.9505765 -58.2132188
C 12.3562485 18.1809218 -57.2916426
O 11.6055412 18.9392478 -56.6695756
C 14.5613807 18.1437631 -56.0764476
H 13.9703084 18.5856413 -55.2756477
H 14.5331341 17.0604963 -55.9537592
C 16.0075358 18.6382877 -55.8749913
H 16.0774433 19.7022195 -56.0961422
C 16.4272853 18.4705187 -54.4170553
H 16.4803628 17.4129 -54.1558347
H 15.7119752 18.972793 -53.7650525
H 17.3987076 18.9361346 -54.2814031
C 17.0207935 17.8886108 -56.7338335
H 16.9321508 16.814618 -56.5697651
H 18.0321975 18.2073195 -56.4892937
H 16.8337156 18.1120355 -57.7796173
N 11.9268476 17.0254245 -57.802589
H 12.5667476 16.4679935 -58.3580903
C 10.6134922 16.4408486 -57.4727742
H 10.200647 16.9817986 -56.6240596
C 10.8460628 14.9974708 -57.0338666
O 11.4968357 14.2196056 -57.7343886

C 9.584129 16.5453119 -58.623281
H 9.8891 15.8453796 -59.384544
C 9.5544193 17.9798466 -59.2141413
H 9.2019235 18.6905423 -58.4697116
H 10.5636784 18.2729503 -59.500348
C 8.1844474 16.0604285 -58.2049168
H 7.7911132 16.6907992 -57.4099126
H 8.2346799 15.0332518 -57.8434264
H 7.5072373 16.083636 -59.0554359
C 8.7221937 18.1222675 -60.4791129
H 7.6661354 18.1114382 -60.2270789
H 8.9737782 17.3097286 -61.1536164
H 8.9770545 19.0592215 -60.9698104
N 10.3564225 14.6574374 -55.8469035
H 9.7941865 15.3397182 -55.3446681
C 10.5610505 13.361721 -55.2088406
H 11.1527113 12.721799 -55.8623133
C 9.2265463 12.6740554 -54.9607565
O 8.2716464 13.3041168 -54.5053329
C 11.33476 13.5316808 -53.9000093
H 12.3374437 13.8914798 -54.1303004
H 10.8329836 14.2807638 -53.2847405
C 11.4560679 12.2414929 -53.107116
C 12.2584577 11.1893237 -53.5913009
H 12.8093437 11.3079028 -54.5115507
C 12.3285074 9.9712448 -52.892086
H 12.935959 9.1595666 -53.2719667
C 11.6002187 9.80203 -51.7022888
H 11.6597189 8.8679635 -51.1608939
C 10.7885537 10.8448729 -51.2227153
H 10.2173261 10.7141468 -50.3149552
C 10.7115946 12.0603749 -51.9250526
H 10.0647863 12.8469327 -51.5572318
N 9.1602094 11.3777451 -55.2613215
H 9.967209 10.9333375 -55.6892098
C 7.9328766 10.5948167 -55.1189933
H 7.301889 11.0701356 -54.3646456
C 8.255057 9.1831792 -54.6477118
O 9.0891373 8.509012 -55.2435789
C 7.1597415 10.5440188 -56.4471444
H 7.7114147 9.9222563 -57.1494066
C 5.8095002 9.8629732 -56.2113163
H 5.4896513 10.0212756 -55.1882349
H 5.9140703 8.7885214 -56.3956002
H 5.0585275 10.2781186 -56.8705113
C 7.0334598 11.9183157 -57.1168015

H 6.7794035 12.6694729 -56.3779092
H 6.2773547 11.9109892 -57.8926942
H 7.9869524 12.1928344 -57.5734035
N 7.5304832 8.6922801 -53.6430702
H 6.8760449 9.3036927 -53.1726615
C 7.6681163 7.3252057 -53.1358409
H 8.2068826 6.7406371 -53.8801015
C 6.2876119 6.6673452 -52.9862792
O 5.4097723 7.1751835 -52.2984167
C 8.5623854 7.3067255 -51.8731256
H 9.5579647 7.6240205 -52.1890165
C 8.094785 8.2976582 -50.7891014
H 7.0661867 8.0877915 -50.5071326
H 8.1462975 9.3195321 -51.1659953
H 8.7354529 8.2405992 -49.9112774
C 8.6896144 5.865025 -51.3446308
H 7.7397634 5.5693468 -50.9079072
H 8.8984866 5.198318 -52.1818034
C 9.804598 5.6643442 -50.3096542
H 9.5898272 6.2220642 -49.3977556
H 10.7584064 5.9929064 -50.7211156
H 9.8773923 4.6049462 -50.0589894
N 6.0735872 5.5791504 -53.7322323
H 6.8322139 5.3057652 -54.3480851
C 4.8587987 4.7465541 -53.7745885
H 4.9169297 4.1923917 -54.7089597
C 3.516118 5.5057891 -53.8614048
O 2.5191161 5.1118456 -53.2532807
C 4.8607276 3.7088065 -52.6438668
H 4.6254463 4.2273965 -51.7263386
H 4.0580549 2.9937707 -52.8329263
C 6.1573545 2.9237375 -52.4338794
H 5.9667513 2.1123083 -51.7318169
H 6.9089625 3.5794581 -51.9982039
C 6.7097361 2.3381834 -53.7201912
O 7.8204849 2.6367907 -54.124086
N 5.9596121 1.5408023 -54.4434662
H 6.3860314 1.1601171 -55.2659575
H 4.9443976 1.5731702 -54.3602218
N 3.4922299 6.609794 -54.6151327
H 4.3734201 6.9183661 -54.9925454
C 2.3102518 7.4619524 -54.7883182
H 1.523502 7.1422596 -54.1042785
C 1.7272705 7.3825057 -56.209906
O 0.5213259 7.2247492 -56.3878293
C 2.7044591 8.890572 -54.3960494

H 3.4554001 9.2689719 -55.0839756
H 3.1257726 8.8895343 -53.3873539
S 1.267988 9.9956678 -54.455393
H 1.9581013 11.1116626 -54.1764562
N 2.5785643 7.3956426 -57.2408166
H 3.5667076 7.4519611 -57.0465178
C 2.1412431 7.4998195 -58.6453044
H 1.3706166 8.2684934 -58.6961616
C 1.455654 6.224998 -59.1639819
O 0.6846533 6.2812467 -60.1185169
C 3.3271499 7.9979964 -59.498981
H 4.2271993 7.4604443 -59.1935633
C 3.5120954 9.5030321 -59.2234801
H 2.7433385 10.070109 -59.7525158
H 3.3888159 9.7067547 -58.1618482
C 3.1360484 7.7830679 -61.0060698
H 2.1735433 8.1823495 -61.3276595
H 3.2063335 6.7193164 -61.2171017
H 3.9295564 8.2650958 -61.5743967
C 4.8911054 10.0097662 -59.6504475
H 5.0099305 9.9628237 -60.7314375
H 5.6735898 9.4236341 -59.1685517
H 4.9797716 11.0455733 -59.3443017
N 1.5717582 5.1041732 -58.4564899
H 2.1827045 5.1206104 -57.6529454
C 0.7328439 3.9149034 -58.651388
H 0.8947388 3.5462889 -59.6639284
C -0.7823536 4.1812482 -58.5331897
O -1.5480582 3.5308124 -59.2486143
C 1.1982313 2.8047712 -57.6992863
H 0.5310809 1.9524817 -57.8022436
H 2.1807305 2.4791663 -58.0396875
C 1.2976814 3.1948038 -56.2119626
H 0.3012642 3.1688227 -55.7657597
H 1.6944627 4.205053 -56.0979221
C 2.2384522 2.2254211 -55.4939914
O 3.4720441 2.4194285 -55.590334
O 1.7426194 1.2099946 -54.9540701
N -1.2002262 5.2068174 -57.7761104
H -0.5129244 5.7123035 -57.2254629
C -2.598939 5.6605511 -57.6700557
H -3.2650489 4.7962077 -57.7183515
C -3.0249164 6.6303356 -58.7861373
O -4.212663 6.7435325 -59.0897442
C -2.8084363 6.3407486 -56.3074555
H -3.8525083 6.6461592 -56.2269157

H -2.1951503 7.2411331 -56.2540386
C -2.494111 5.4648105 -55.1222584
N -3.2748369 4.3995001 -54.6813414
C -1.402258 5.5724542 -54.3123075
H -0.6083186 6.2963122 -54.4092074
C -2.6439835 3.8919381 -53.61534
H -2.9894624 3.0311948 -53.0587202
N -1.5076655 4.5702295 -53.3754836
H -0.8044458 4.3204002 -52.6953078
N -2.0794667 7.3391896 -59.4136958
H -1.1158875 7.1556197 -59.1719981
C -2.3616997 8.3287098 -60.4667779
H -3.2276859 8.9173499 -60.1602128
C -2.7464924 7.5875773 -61.7548098
O -2.001621 6.7304906 -62.2333305
C -1.169923 9.3091347 -60.6337605
H -0.2399088 8.7535995 -60.5132491
C -1.1908147 10.461597 -59.5993055
H -0.3002541 11.0742561 -59.749503
H -2.0610367 11.0934907 -59.7853356
C -1.1248122 9.968303 -62.0256172
H -2.0519598 10.5058078 -62.2273489
H -0.9656117 9.221869 -62.8030621
H -0.2929621 10.6711498 -62.0761009
C -1.2165483 10.0459058 -58.1225757
H -0.4028053 9.3553923 -57.9105932
H -2.1699243 9.5788426 -57.8756608
H -1.0975652 10.9321865 -57.4986943
N -3.9191623 7.9002296 -62.3131057
H -4.4943561 8.5906527 -61.8567087
C -4.4811881 7.1647627 -63.4544377
H -4.2521094 6.1048541 -63.3300885
C -3.8496852 7.5951677 -64.7889643
O -3.2727482 6.7640869 -65.4867586
C -6.0132403 7.3088003 -63.4796484
H -6.2694267 8.3537053 -63.6622205
H -6.4039576 6.7184002 -64.3096062
C -6.7166646 6.8608593 -62.1861027
H -7.7887787 7.0203155 -62.3018223
H -6.3872767 7.4812947 -61.3521355
C -6.4970237 5.3889815 -61.8459751
O -6.9995508 4.4963847 -62.5114945
N -5.7572628 5.0902401 -60.8042215
H -5.833909 4.14822 -60.4307188
H -5.3482376 5.8097848 -60.2156199
N -3.8569681 8.8981386 -65.0946658

H -4.3202539 9.5439063 -64.4785248
C -3.1401507 9.4583446 -66.2477274
H -3.0411782 8.6833109 -67.0107969
C -1.7236527 9.8698317 -65.8282051
O -1.4618108 10.9868417 -65.364135
C -3.9305454 10.5951731 -66.9132344
H -4.0322571 11.4261952 -66.2131145
H -4.9304066 10.2295566 -67.1538309
C -3.2659619 11.0967388 -68.2087631
O -2.0957811 10.7292587 -68.4770125
O -3.9030893 11.919111 -68.8986723
N -0.803794 8.9111941 -65.9612056
H -1.116004 8.0117478 -66.302686
C 0.6148988 9.1220086 -65.6716635
H 0.701243 9.6227611 -64.7065084
C 1.2656105 10.0741342 -66.6689175
O 2.098196 10.8648421 -66.2441968
C 1.3529267 7.7757301 -65.5926566
H 1.1126767 7.1689474 -66.4679944
H 2.4297305 7.955249 -65.5762918
C 0.9574017 7.0366377 -64.3062496
H 1.2856483 7.6316669 -63.4540904
H -0.1280695 6.9419732 -64.2625537
C 1.5717334 5.633152 -64.2128231
H 2.6407791 5.6991469 -64.4166036
H 1.1128656 4.9710833 -64.9484462
C 1.3819517 5.0770151 -62.7971826
H 1.7874912 5.8138986 -62.1066302
H 1.9538723 4.1524789 -62.6816544
N -0.0351696 4.8430507 -62.4374898
H -0.6226182 5.656999 -62.6128357
H -0.1159741 4.6848988 -61.4413045
H -0.410129 4.0257107 -62.9061462
N 0.857613 10.1083443 -67.9375912
H 0.0306158 9.5949737 -68.2147783
C 1.4627276 11.0419099 -68.8908006
H 2.5495178 10.9536548 -68.8456466
C 1.0955327 12.4971915 -68.5945729
O 1.9570158 13.368256 -68.6695282
C 1.0240044 10.7936445 -70.3246517
H -0.0383867 11.0268319 -70.4208665
H 1.5987591 11.5133366 -70.9013528
C 1.2870139 9.4334093 -70.9591885
H 0.6207275 8.6875652 -70.5204886
H 2.321262 9.1555211 -70.7495981
C 1.0670261 9.5267931 -72.4818624

O 1.5837226 8.6346019 -73.1863309
O 0.449561 10.5278113 -72.9382914
N -0.1651418 12.7909911 -68.2489103
H -0.8538143 12.0316837 -68.2624408
C -0.597046 14.1400191 -67.8500139
H -0.3606733 14.8368522 -68.6493023
C 0.1788132 14.6128788 -66.6331437
O 0.6875716 15.7306213 -66.6664212
C -2.1126775 14.1466946 -67.5941196
H -2.6227997 14.0571914 -68.5530931
H -2.3576696 13.272807 -66.9910062
C -2.6896842 15.3612165 -66.8459386
H -3.7652708 15.2087699 -66.7430743
H -2.2667664 15.3935762 -65.8416272
C -2.4521123 16.7168604 -67.5216241
H -2.901966 16.7154243 -68.5155877
H -1.3820194 16.9074539 -67.6060141
C -3.0951271 17.8042082 -66.6541248
H -2.7560098 17.6658543 -65.6223159
H -4.1813943 17.6762267 -66.6795216
N -2.71647 19.159312 -67.1134731
H -1.700728 19.2804625 -67.0277463
H -3.1123092 19.8764223 -66.5258363
H -2.9676407 19.3212056 -68.075571
N 0.3204954 13.766832 -65.6119406
H -0.1301998 12.8600258 -65.6626692
C 1.1372004 14.0988161 -64.4489276
H 0.7681291 15.0311553 -64.0186157
C 2.6010132 14.3488103 -64.8440332
O 3.1203401 15.4231067 -64.5515805
C 1.0048208 13.0052398 -63.3833035
H -0.0274297 12.9715737 -63.0338871
H 1.2416141 12.0362933 -63.8251743
C 1.9196494 13.2544928 -62.2026704
C 1.5902525 14.2330469 -61.2449909
H 0.6507624 14.7645696 -61.3091331
C 2.5054578 14.5589645 -60.2283914
H 2.2645 15.3361693 -59.5185997
C 3.7477584 13.9046379 -60.163313
H 4.4610224 14.1635952 -59.3939693
C 4.075634 12.9324819 -61.1222156
H 5.0420339 12.4538226 -61.0965527
C 3.1634757 12.6011592 -62.1355357
H 3.4384687 11.8742898 -62.8856969
N 3.2219607 13.4431795 -65.6129092
H 2.7236004 12.5935768 -65.8627977

C 4.6285742 13.5613066 -66.0272132
H 5.2190656 13.6913037 -65.136371
C 4.8554892 14.8103973 -66.9071792
O 5.7538651 15.5982238 -66.6248052
C 5.1469566 12.2423806 -66.6375441
H 4.5264298 12.0049949 -67.503892
C 6.6112232 12.3569321 -67.0848273
H 7.294004 12.1712733 -66.2581363
H 6.8394987 13.3368053 -67.4929083
H 6.78462 11.6171695 -67.8603109
C 5.0988351 11.0728698 -65.6156212
H 5.8917595 11.1882859 -64.8758281
H 4.1709873 11.0890031 -65.0529447
C 5.2309691 9.6913749 -66.2729081
H 6.195366 9.5950969 -66.7695518
H 4.4343643 9.5515472 -67.0036455
H 5.1519131 8.9179254 -65.5089451
N 3.9636351 15.1292163 -67.8520424
H 3.2265652 14.4600239 -68.0547949
C 4.0218868 16.3871815 -68.6226211
H 5.0042765 16.4596209 -69.0901891
C 3.8797531 17.6361744 -67.751329
O 4.3853571 18.6900604 -68.1393617
C 2.9423549 16.3991231 -69.7126516
H 1.9845357 16.1193492 -69.2705717
H 2.8543825 17.4158555 -70.0989016
C 3.2701847 15.4763868 -70.8965743
H 3.2834725 14.434007 -70.5792135
H 4.2600221 15.7274747 -71.2810722
C 2.2462379 15.6651832 -72.0260354
H 2.5661878 15.0769262 -72.8877034
H 2.2537104 16.7118842 -72.3344896
N 0.873631 15.2917612 -71.613203
H 0.3736673 15.9475456 -71.0440993
C 0.2606047 14.1451865 -71.8617678
N 0.8324717 13.2109762 -72.562001
H 0.4215544 12.2750905 -72.7048818
H 1.7817286 13.3170273 -72.8573687
N -0.9394563 13.9204432 -71.4047869
H -1.4254517 14.5628763 -70.815964
H -1.3235025 12.9947997 -71.5411287
N 3.1468849 17.5726836 -66.640555
H 2.7147858 16.6893069 -66.3908726
C 3.0422072 18.6861717 -65.6891141
H 2.9536692 19.6115987 -66.2544648
C 4.3172776 18.8331359 -64.8542892

O 4.818539 19.9522103 -64.7316767
C 1.7891423 18.5422215 -64.8007602
H 1.0555546 17.9044931 -65.2924164
H 2.0612306 18.0606817 -63.8602092
C 1.1195113 19.8957002 -64.5028273
H 1.8473703 20.5638183 -64.0369374
H 0.3048766 19.7346467 -63.7929669
C 0.5491297 20.5310302 -65.7790337
O -0.2282403 19.8629453 -66.4986209
O 0.9457575 21.6510811 -66.1609826
N 4.94132 17.725144 -64.4234338
H 4.4998686 16.8199543 -64.5638902
C 6.2140599 17.8019982 -63.6909489
H 6.0524835 18.5815138 -62.9562327
C 7.3890477 18.3481159 -64.518059
O 8.2732855 18.9822581 -63.9445966
C 6.5604161 16.5360761 -62.8697259
H 7.6036421 16.612572 -62.5826921
C 5.687129 16.5745079 -61.5928064
H 4.6438896 16.3754152 -61.8451636
H 5.7360653 17.5498829 -61.1123979
H 6.0256541 15.83006 -60.8757306
C 6.3611885 15.1622226 -63.5084856
H 5.298497 14.9474787 -63.5301493
H 6.74844 15.1843201 -64.5161972
C 7.0610066 14.0140301 -62.7739185
H 6.6684855 13.8981371 -61.7667884
H 8.122155 14.2216184 -62.7084251
H 6.9110273 13.0840706 -63.3228865
N 7.30301 18.2991724 -65.852446
H 6.594574 17.6978391 -66.2541671
C 8.2430857 18.9945263 -66.7521705
H 9.256973 18.7089679 -66.4899572
C 8.1690656 20.51075 -66.5818082
O 9.1990222 21.1611148 -66.460308
C 7.977772 18.6399987 -68.229651
H 6.9892592 19.0220523 -68.484297
C 8.9845327 19.2914653 -69.1874602
H 10.0014391 18.9979181 -68.9194354
H 8.9126701 20.3778295 -69.1491886
H 8.7886603 18.966429 -70.2109033
C 7.96589 17.1408008 -68.5243393
H 8.943109 16.8146886 -68.8779901
H 7.2357508 16.9438971 -69.3046957
H 7.7171778 16.551408 -67.6508645
N 6.9615765 21.0893303 -66.532496

H 6.1488449 20.486938 -66.5022801
C 6.780615 22.5474602 -66.4135413
H 7.4330732 23.0541204 -67.1286945
C 7.2202667 23.0717325 -65.0569848
O 7.912679 24.0815246 -64.9821131
C 5.3159302 22.9327156 -66.6760503
H 4.6831159 22.049465 -66.5844177
H 4.9780527 23.653042 -65.9286919
C 5.1585947 23.5769347 -68.0589042
H 5.5905341 22.9257093 -68.8152424
H 5.7275276 24.5057287 -68.0837541
C 3.6905468 23.8966561 -68.3952165
H 3.6380314 24.2920585 -69.4108383
H 3.3606373 24.6917945 -67.7218206
N 2.7656957 22.7454938 -68.2527494
H 2.0283473 22.845749 -67.5579178
C 2.88091 21.5274568 -68.7454514
N 3.8693124 21.140862 -69.4991559
H 4.0949378 20.1565639 -69.4957941
H 4.5709032 21.8214323 -69.7161739
N 1.9903342 20.6334323 -68.4586807
H 1.2955849 20.8980357 -67.7519289
H 2.1477213 19.6774308 -68.6872359
N 6.8385406 22.3850171 -63.9793491
H 6.2678414 21.5594362 -64.1188139
C 7.0762139 22.8882716 -62.6154744
H 6.7536171 23.9302454 -62.5989197
C 8.5529194 22.901266 -62.209056
O 8.8936518 23.5075295 -61.1960446
C 6.2302789 22.1490035 -61.5659052
H 6.3002834 22.7158685 -60.6375336
C 4.7406745 22.1370212 -61.9312552
H 4.5608851 21.5072413 -62.7992222
H 4.4104188 23.1524728 -62.155835
H 4.1599278 21.7513496 -61.0951958
C 6.7187884 20.740197 -61.2639461
H 6.946144 20.2365441 -62.1944734
H 5.9697426 20.1820863 -60.7022476
H 7.6299397 20.7966649 -60.6732389
N 9.4330305 22.2598839 -62.9727844
H 9.1190088 21.8290885 -63.831774
C 10.8586508 22.2157185 -62.6987607
H 11.0205601 22.3896924 -61.6385775
C 11.6587618 23.2613323 -63.4943908
O 11.250341 23.686733 -64.571033
C 11.3162717 20.8126336 -63.009183

H 11.2222436 20.673128 -64.0870845
H 10.6948542 20.0962774 -62.4706286
H 12.34619 20.6925253 -62.698318
N 12.8048385 23.6814376 -62.9622398
H 13.0708624 23.3246349 -62.0518115
C 13.7495271 24.5578181 -63.6509041
H 13.1778 25.2862108 -64.2238724
C 14.6719354 23.7794981 -64.6223379
O 14.975169 22.6149417 -64.3663321
C 14.5534374 25.3133525 -62.584314
H 15.1349368 24.6050227 -61.9900518
H 13.8784704 25.8645982 -61.9278724
H 15.2402197 26.0126728 -63.0612602
N 15.2562672 24.4346856 -65.6428965
C 16.3457623 23.8565942 -66.4329207
H 15.9645131 23.0084207 -66.9972777
C 17.5084646 23.3942187 -65.5437926
O 17.9666931 24.1420647 -64.6815619
C 16.7604415 24.9539896 -67.41675
H 17.5396302 25.5804047 -66.9808329
H 17.0916402 24.5356973 -68.368371
C 15.4800201 25.7739039 -67.5748403
H 14.8102338 25.2746548 -68.2775496
H 15.6876092 26.7949285 -67.8961341
C 14.8755827 25.7311526 -66.1737705
H 13.7917232 25.8368128 -66.2352599
H 15.3069372 26.5190763 -65.555338
N 17.9761557 22.1537945 -65.7184936
H 17.5578649 21.5750566 -66.4370437
C 18.9691881 21.5330127 -64.828799
H 19.4528712 20.7173447 -65.3653614
H 19.7268063 22.271958 -64.5676816
C 18.3945524 20.9579177 -63.5231969
O 19.1431088 20.4234825 -62.6929978
N 17.0763422 20.9995032 -63.325819
H 16.4871733 21.5126475 -63.9742723
C 16.4200515 20.282351 -62.2438022
H 16.8992307 20.5608404 -61.3042546
C 16.5430015 18.7673145 -62.4295642
O 16.5979498 18.2344241 -63.5433115
C 14.9511917 20.6876016 -62.1594518
H 14.4759675 20.4721613 -63.1160417
H 14.876111 21.7519627 -61.935609
H 14.4412873 20.1255796 -61.3775577
N 16.4551063 18.0541021 -61.3125826
H 16.2855776 18.5690471 -60.4511779

C 16.3207142 16.6066863 -61.2672556
H 16.667364 16.1436285 -62.1925943
C 14.8430527 16.4876811 -61.1714432
O 14.2703098 16.4918358 -60.0760996
C 17.1025975 16.0055349 -60.0967373
H 16.7267382 16.4254048 -59.1674263
H 16.9332383 14.927969 -60.0696969
C 18.6151246 16.2789271 -60.2035818
H 18.7934009 17.3000794 -60.5397025
H 19.0498173 15.6006874 -60.9384452
C 19.3309473 16.1098317 -58.8681021
O 19.8209138 17.063498 -58.2767724
N 19.4280626 14.9016131 -58.3528361
H 19.9160454 14.8206799 -57.47876
H 19.0269415 14.1074667 -58.8221429
N 14.2900884 16.7928196 -62.3621341
H 14.9082487 16.9578584 -63.1449686
C 12.8660165 16.8921841 -62.5786515
H 12.50313 17.8129391 -62.118628
C 12.3945187 16.8195691 -64.0583579
H 11.5182164 16.1806944 -64.1205188
C 11.927267 18.2080647 -64.4477732
H 12.7140434 18.9289267 -64.2293846
H 11.0102521 18.4103486 -63.8975376
H 11.6871596 18.2879955 -65.5028421
C 13.3973882 16.3601632 -65.1198995
H 13.1242213 16.743578 -66.1014841
H 14.3632553 16.7589904 -64.8585091
C 13.4802412 14.8541873 -65.2635705
H 12.5137973 14.4524802 -65.5470749
H 13.7913259 14.4259101 -64.3196536
H 14.1809111 14.6061911 -66.0551717
C 12.2832975 15.764129 -61.7971147
O 11.2494559 16.0014477 -61.1925983
N 12.9953431 14.6160479 -61.6922139
H 13.8487719 14.4634771 -62.2140222
C 12.4409562 13.5648977 -60.9104669
H 11.9661348 14.0429408 -60.056893
C 13.3950247 12.5287825 -60.3251963
O 14.3898634 12.1321749 -60.9337501
C 11.3641755 12.9163869 -61.7760491
H 11.69877 11.9263678 -62.0871113
C 10.1448767 12.7675339 -60.8818847
H 9.7797381 13.7482455 -60.5753726
H 10.389334 12.200937 -59.985975
H 9.3515675 12.2659959 -61.4236928

C 10.8142608 13.647437 -63.0167973
H 10.2651247 14.5505195 -62.7809551
H 10.1246692 12.9691349 -63.5158578
H 11.6027534 13.8396563 -63.7244018
N 12.992268 12.0624941 -59.1407394
H 12.2290494 12.5696896 -58.7005972
C 13.3517678 10.7991302 -58.4875107
H 13.8691189 10.1454328 -59.1901295
C 12.022295 10.1409558 -58.0722694
O 11.2124517 10.7801555 -57.3905926
C 14.2732964 11.0619765 -57.2714117
H 13.7556098 11.7467959 -56.5983282
C 14.5539489 9.7552096 -56.5040559
H 15.098085 9.0547942 -57.1393494
H 13.6255968 9.2720965 -56.1977078
H 15.1326487 9.9473315 -55.6028041
C 15.5967541 11.7351388 -57.7104154
H 16.1322448 11.0671286 -58.3834904
H 15.3704105 12.6510699 -58.2557262
C 16.5299331 12.1281646 -56.5575202
H 16.9353557 11.2398739 -56.0737671
H 15.9862613 12.7290053 -55.8275692
H 17.3598219 12.7148318 -56.9523203
N 11.7580335 8.9046503 -58.5018071
H 12.4971308 8.4049395 -59.0006782
C 10.4983895 8.1867901 -58.2471085
H 9.9675434 8.6814518 -57.4396159
C 10.7986262 6.7492106 -57.8252037
O 11.102295 5.9443846 -58.7003625
C 9.5576584 8.1569015 -59.488233
H 10.0657489 7.6118162 -60.2850846
C 9.2280089 9.5527545 -60.0531933
H 8.7489912 10.1650888 -59.2892799
H 10.1684049 10.018405 -60.3410172
C 8.2538598 7.3944316 -59.1507915
H 7.707404 7.913199 -58.3643917
H 8.4738782 6.3822778 -58.8197286
H 7.6197814 7.2880813 -60.0276344
C 8.3419087 9.5094014 -61.311817
H 7.3388524 9.1658198 -61.0701747
H 8.7868381 8.8466241 -62.055071
H 8.2449684 10.4990386 -61.7479271
N 10.3778145 6.3647875 -56.6173089
H 10.0867171 7.0875804 -55.969359
C 10.0771669 4.965996 -56.2829504
H 10.6161371 4.3342597 -56.978693

C 8.5735844 4.707958 -56.4325046
O 7.7547345 5.5273144 -56.0055968
C 10.6004787 4.5910992 -54.8968489
H 10.1061673 5.1945444 -54.1366728
H 11.6735419 4.7889482 -54.8607758
O 10.3828632 3.2140938 -54.6418497
H 9.4472056 3.0366627 -54.4338089
N 8.1607001 3.6309068 -57.1055167
H 8.8769937 2.9804449 -57.4273788
C 6.7371884 3.300688 -57.3620578
H 6.1596131 3.5083897 -56.4621738
C 6.5491433 1.8064488 -57.6597888
O 7.4787363 1.1227106 -58.0834072
C 6.1822375 4.201798 -58.4902398
H 6.9381585 4.3099942 -59.2694911
C 4.8723056 3.766805 -59.1432156
H 4.1171477 3.6139869 -58.3752925
H 5.0235349 2.8460121 -59.7068042
H 4.5357146 4.5394803 -59.8340945
O 5.8902408 5.4700017 -57.9372644
H 6.6083844 5.6589023 -57.3062582
N 5.3715908 1.2506011 -57.3572631
H 4.6553821 1.8125038 -56.8983762
C 5.0533527 -0.1448405 -57.6414244
H 5.9175855 -0.7539901 -57.3692877
C 4.7686066 -0.3651727 -59.1348527
O 3.9382459 0.3149866 -59.7426248
C 3.8790949 -0.6020956 -56.7719286
H 2.9852171 -0.0491227 -57.0590648
H 4.0992285 -0.4156114 -55.7198245
H 3.7121333 -1.6687299 -56.9224242
N 5.384449 -1.4132231 -59.6710028
H 5.9615485 -1.978817 -59.0585404
C 5.274073 -1.8707675 -61.0481607
H 4.52063 -1.2819514 -61.5709594
C 4.819635 -3.3368143 -61.111532
O 4.71484 -4.0327033 -60.0985856
C 6.6290324 -1.6468094 -61.7413333
H 6.7247861 -2.2962304 -62.6122492
H 7.4480343 -1.8571985 -61.0513851
S 6.7161264 0.0744788 -62.2988248
H 5.7073454 -0.0156779 -63.1767116
N 4.5804398 -3.8163232 -62.3306187
H 4.6770828 -3.1691214 -63.1064769
C 4.346574 -5.2294842 -62.6551505
H 4.459447 -5.8264837 -61.7510928

C 5.4032849 -5.7429896 -63.6329606
O 6.1897972 -4.9572822 -64.1649364
C 2.9059161 -5.4309908 -63.1662228
H 2.7672556 -6.4849372 -63.4143421
H 2.2061259 -5.1920836 -62.3647088
C 2.5629917 -4.5759187 -64.3962885
H 3.3538419 -4.6507815 -65.1381943
H 2.4774119 -3.5317996 -64.1082129
C 1.239358 -5.0199314 -65.0200982
H 0.4409992 -4.8935195 -64.2851521
H 1.2911458 -6.0804497 -65.2787751
N 0.9258565 -4.2126199 -66.212061
H 0.4081314 -3.3590145 -66.0351975
C 1.2259418 -4.5210722 -67.4631425
N 1.9688723 -5.5468659 -67.7776155
H 2.1152903 -5.8721481 -68.7219343
H 2.3570204 -6.1389378 -67.0480665
N 0.7801215 -3.779182 -68.4391839
H 0.1596895 -3.0162654 -68.2328983
H 0.9754474 -4.0537471 -69.3841574
N 5.3899017 -7.0384504 -63.929528
H 4.7383203 -7.6344439 -63.4363782
C 6.0942102 -7.5596355 -65.1036889
H 7.0977374 -7.1300138 -65.1006469
C 5.4232967 -7.0925274 -66.4226341
O 4.3354509 -6.5106139 -66.4170726
C 6.2677109 -9.0838745 -64.9594063
H 6.6936781 -9.2945219 -63.9792089
H 6.9785677 -9.4402086 -65.7029615
C 4.9912565 -9.885612 -65.1365704
O 4.2025192 -9.6370408 -66.0222303
N 4.7808317 -10.9165378 -64.3629691
H 3.901636 -11.390675 -64.4666077
H 5.4341878 -11.1196392 -63.6117621
N 6.087026 -7.301176 -67.5654737
H 6.97492 -7.7768733 -67.5279906
C 5.5686416 -6.8960675 -68.8824775
H 5.3268371 -5.8354456 -68.847767
C 4.2717675 -7.636028 -69.2757134
O 3.3960169 -7.0361793 -69.9063768
C 6.6966829 -7.1057223 -69.9118019
H 7.5465829 -6.4850685 -69.622889
H 7.0144693 -8.1489414 -69.8602459
C 6.3428838 -6.7968035 -71.3816833
H 5.5274226 -7.4439008 -71.7033055
C 5.9359276 -5.3387014 -71.5982807

H 6.7312629 -4.6712647 -71.263949
H 5.0185384 -5.1252485 -71.0539862
H 5.7487945 -5.1681951 -72.6580208
C 7.5553389 -7.0841982 -72.2684394
H 8.3866897 -6.4341584 -71.9947723
H 7.2949337 -6.9136733 -73.313153
H 7.8578121 -8.1251123 -72.1530906
N 4.1334798 -8.8979656 -68.8636108
H 4.8250579 -9.2749777 -68.2335885
C 3.0282568 -9.7990911 -69.2066128
H 2.994302 -9.8630138 -70.2915907
C 1.6602333 -9.3119172 -68.6927965
O 1.5945623 -8.516802 -67.7502102
C 3.3072741 -11.2092038 -68.6610331
H 2.5364612 -11.8929918 -69.0166306
H 3.2739869 -11.2045757 -67.5732222
O 4.5717364 -11.6737482 -69.1032695
H 4.7435367 -12.5260357 -68.6915725
N 0.5433211 -9.7783737 -69.2905212
C -0.7985269 -9.4881579 -68.7920022
H -0.974344 -8.4156383 -68.878059
C -0.9754439 -9.9212677 -67.3331253
O -0.527133 -10.9988821 -66.9390965
C -1.7659872 -10.2366409 -69.7171588
H -1.976375 -11.2318292 -69.3196156
H -2.693753 -9.6827823 -69.8650191
C 0.4554123 -10.5666366 -70.5124142
H 1.1583618 -10.2258453 -71.2721388
H 0.6242898 -11.6193873 -70.280253
C -0.973233 -10.3730318 -71.0152853
H -1.0393598 -9.4455678 -71.586019
H -1.3115491 -11.2200922 -71.6130074
N -1.6717958 -9.0991233 -66.547554
H -1.9774122 -8.2113845 -66.9104634
C -1.9426282 -9.3712957 -65.1363939
H -0.9869589 -9.4906116 -64.6280691
C -2.7599268 -10.6578647 -64.9656591
O -3.8400503 -10.7898855 -65.5414905
C -2.701309 -8.1901542 -64.5041285
H -2.7718945 -8.3380187 -63.4261289
H -3.7101012 -8.1606276 -64.9151862
O -2.0772418 -6.945159 -64.7763668
H -2.4140318 -6.2669389 -64.1668781
N -2.292544 -11.5662814 -64.1043462
H -1.3627723 -11.44079 -63.7365719
C -3.0207712 -12.7921241 -63.7457168

H -3.2244875 -13.323451 -64.6781217
C -4.3966937 -12.4684447 -63.1343671
O -4.6230048 -11.3825377 -62.579092
C -2.1096583 -13.72106 -62.9038127
H -1.910148 -14.6234283 -63.4834728
H -1.1418491 -13.2284864 -62.8004951
C -2.5148567 -14.1252411 -61.4840284
H -2.7946617 -13.2267165 -60.9280213
H -1.6416138 -14.553518 -60.9864657
C -3.6505368 -15.1492933 -61.4302847
O -4.2452488 -15.22482 -60.332582
O -4.0275463 -15.7087837 -62.4791995
N -5.3275999 -13.4242844 -63.2208885
H -5.0081732 -14.3620026 -63.4746751
C -6.7373853 -13.2558538 -62.8708644
H -7.1275333 -12.4809128 -63.5271843
C -6.946757 -12.7679238 -61.4410492
O -7.9117529 -12.0322895 -61.2615722
C -7.4812086 -14.5788163 -63.1367407
H -7.2969487 -14.8900304 -64.1668939
H -7.0568127 -15.3459934 -62.4846586
C -8.9995942 -14.5392484 -62.9048429
H -9.3885239 -15.5543231 -62.9959171
H -9.1947796 -14.2219061 -61.8805407
C -9.7553918 -13.6445641 -63.9063237
H -10.1326223 -14.2766743 -64.7127237
H -9.0952164 -12.9031723 -64.3565096
C -10.9343519 -12.935928 -63.2296966
H -11.4774207 -13.6713244 -62.6279936
H -11.6147183 -12.563259 -64.0023007
N -10.4659423 -11.8111498 -62.3804404
H -9.7675482 -12.1340372 -61.710333
H -11.2343898 -11.3862407 -61.8810379
H -10.0067443 -11.1094506 -62.9439105
N -6.0816248 -13.0886981 -60.4718269
H -5.3969747 -13.8110332 -60.6825751
C -6.0968226 -12.5397581 -59.1017552
H -7.1172031 -12.2357542 -58.8791371
C -5.2212816 -11.2922417 -58.8815129
O -5.4720267 -10.5585664 -57.9262851
C -5.7549487 -13.6167123 -58.0640669
H -6.3952429 -14.4869167 -58.2181221
H -5.9482444 -13.2182487 -57.0671658
O -4.3992676 -14.0098058 -58.1432093
H -4.343102 -14.5866227 -58.960205
N -4.2784466 -10.9636136 -59.7796263

H -4.191226 -11.5356199 -60.6116894
C -3.5645761 -9.675878 -59.7459851
H -3.2982635 -9.4612025 -58.710529
C -4.4764878 -8.5391038 -60.2170946
O -4.6424323 -7.5619692 -59.492832
C -2.2637066 -9.7208699 -60.5782299
H -1.5860132 -10.4576694 -60.1459091
H -2.5060962 -10.0411425 -61.5913691
C -1.54592 -8.348949 -60.6594275
H -2.2252017 -7.6004366 -61.0640231
C -1.049715 -7.869789 -59.2926931
H -0.3925492 -8.6162577 -58.8481829
H -1.895953 -7.6778135 -58.6345231
H -0.5004847 -6.9352805 -59.4146506
C -0.3461376 -8.4080241 -61.5969519
H 0.3865714 -9.1103831 -61.226032
H 0.1108929 -7.4212003 -61.6794127
H -0.6523785 -8.7230191 -62.5882788
N -5.1169134 -8.689943 -61.3849318
H -4.9457027 -9.5459942 -61.90868
C -6.0412299 -7.6860189 -61.9416269
H -5.4551489 -6.7857532 -62.1437637
C -7.1240799 -7.231914 -60.9375162
O -7.2297424 -6.0346225 -60.7273625
C -6.5934143 -8.1928127 -63.2907024
H -5.7661369 -8.2380254 -63.9984981
H -6.9541082 -9.2147834 -63.16253
C -7.7329938 -7.3712305 -63.9310273
H -7.8971692 -7.7500898 -64.9408023
H -8.6442331 -7.5659483 -63.3636415
C -7.5575872 -5.8442729 -64.0094301
H -7.4980538 -5.4352241 -63.0077227
H -8.4615509 -5.4163366 -64.4482493
C -6.3614685 -5.3390559 -64.8205827
H -6.662531 -5.2804452 -65.8695286
H -5.5195257 -6.0309988 -64.7300999
N -5.9571078 -4.0034871 -64.3263948
H -6.7679802 -3.395136 -64.17414
H -5.2630821 -3.5587778 -64.9038722
H -5.4837897 -4.0957187 -63.4295262
N -7.871645 -8.0931999 -60.2239082
C -8.873936 -7.6616363 -59.2573345
H -9.5234761 -6.9139405 -59.7159941
C -8.2649088 -7.0320422 -57.9992117
O -8.9337415 -6.198365 -57.3998078
C -9.7099499 -8.8997193 -58.9176722

H -10.0869565 -8.8805224 -57.8938592
H -10.5345133 -8.9914458 -59.6264198
C -7.9351771 -9.5147877 -60.3439147
H -8.4774898 -9.7379089 -61.262144
H -6.9490279 -9.9651637 -60.3563997
C -8.720291 -10.0246495 -59.1344312
H -9.2281338 -10.9685741 -59.3297095
H -8.0659944 -10.10908 -58.2666461
N -7.021786 -7.3606245 -57.6043131
H -6.4998668 -8.0322734 -58.1535313
C -6.3056253 -6.6353359 -56.5348897
H -6.9728801 -6.5111878 -55.6824392
C -5.9322881 -5.2270172 -57.0036012
O -6.1602659 -4.2700824 -56.2681912
C -5.0687729 -7.4390979 -56.0860549
H -5.3895788 -8.424192 -55.7418211
H -4.4305065 -7.5948128 -56.9556649
C -4.1992388 -6.7757217 -54.993123
H -3.2115868 -7.2341931 -55.0481208
H -4.0668317 -5.7140503 -55.2046993
C -4.7169933 -6.9550987 -53.5545555
H -4.8133604 -8.0239603 -53.351868
H -5.6999526 -6.4874112 -53.4551922
C -3.7314371 -6.3211879 -52.5553124
H -3.9467867 -5.2535484 -52.4822222
H -2.7197203 -6.4281609 -52.9491867
N -3.7880118 -6.9492504 -51.2117734
H -4.7008678 -6.8646728 -50.7950431
H -3.1294465 -6.5095671 -50.5629086
H -3.5314121 -7.9249485 -51.2652042
N -5.4101965 -5.0928218 -58.2204721
H -5.2513331 -5.9347831 -58.7634563
C -5.1139113 -3.8079675 -58.8626221
H -4.4172266 -3.2441876 -58.2414818
C -6.3827766 -2.9486339 -58.9977917
O -6.424878 -1.8461333 -58.4601641
C -4.4306352 -4.0925634 -60.2147685
H -3.4889227 -4.612916 -60.0251308
H -5.0681263 -4.750292 -60.8032861
C -4.1402352 -2.8329003 -61.0402102
H -3.5223312 -2.1630568 -60.4391233
H -5.0881717 -2.3339607 -61.2560169
C -3.4537445 -3.1310734 -62.3813936
O -3.6879035 -4.2298107 -62.9487496
O -2.7668995 -2.224337 -62.9038172
N -7.4664462 -3.5002571 -59.5499567

H -7.3734285 -4.4240226 -59.9575726
C -8.7116426 -2.7639545 -59.7964255
H -8.4353106 -1.8095062 -60.2480485
C -9.4638687 -2.4089424 -58.5074308
O -9.9528278 -1.288221 -58.3755535
C -9.6118555 -3.5466591 -60.7821039
H -9.0607949 -4.3693492 -61.2330581
H -10.4668038 -3.9705644 -60.2532929
C -10.1111674 -2.6392604 -61.9230753
H -10.7214885 -1.8416841 -61.4932013
H -10.7558622 -3.2309231 -62.5768496
C -8.9667387 -2.0313695 -62.7636765
O -8.2371866 -2.7994259 -63.4391515
O -8.8066446 -0.7847794 -62.7441526
N -9.47042 -3.2962327 -57.4999447
H -9.0417097 -4.20374 -57.6524768
C -10.1010431 -3.0170332 -56.1973756
H -11.0288266 -2.4764864 -56.3973745
C -9.2769901 -2.0761537 -55.3142795
O -9.8498703 -1.4157787 -54.4554675
C -10.4660358 -4.334596 -55.4944689
H -11.0058283 -4.9560219 -56.2112972
H -9.5566922 -4.8544857 -55.1890845
C -11.3704398 -4.1053174 -54.2696322
H -10.7558442 -3.807233 -53.4188719
H -12.0692675 -3.2953097 -54.4855386
C -12.2007088 -5.3376816 -53.8874421
H -12.8839548 -5.5640901 -54.7082912
H -12.7938671 -5.0865768 -53.0059178
C -11.3301939 -6.5636396 -53.5898349
H -10.6293076 -6.3072624 -52.7892185
H -10.7480368 -6.8020651 -54.4859351
N -12.1634946 -7.726805 -53.1936491
H -12.701381 -7.5095402 -52.3633894
H -11.5843428 -8.534113 -52.9993255
H -12.8073944 -7.9650457 -53.9371857
N -7.9716141 -1.9444161 -55.5661454
H -7.5758423 -2.5040646 -56.3110106
C -7.1488087 -0.8533673 -55.0040867
H -7.6222074 -0.5067725 -54.0856447
C -7.1064176 0.3871534 -55.9041949
O -6.4532318 1.363562 -55.5514579
C -5.7327836 -1.3132419 -54.6059369
H -5.1704231 -0.4493004 -54.2512706
C -5.7686133 -2.3392788 -53.4714731
H -6.3200593 -3.2274988 -53.7777969

H -6.2540956 -1.8980821 -52.6008782
H -4.7497511 -2.6174525 -53.2046511
O -5.0121386 -1.9015539 -55.6648057
H -5.5081724 -2.690785 -55.9458878
N -7.8166632 0.3654646 -57.0417343
H -8.3462446 -0.4726061 -57.2427193
C -7.8502309 1.3899778 -58.0923619
H -8.3461036 0.901136 -58.9292334
C -6.4594769 1.7892235 -58.633805
O -6.3001095 2.8344891 -59.2640218
C -8.7526115 2.5574542 -57.6322694
H -9.6187161 2.1305136 -57.1235023
H -8.1999693 3.1478449 -56.8997247
C -9.2803135 3.5032468 -58.7320415
H -8.4579137 4.0436594 -59.1952806
C -10.0659324 2.771923 -59.8264709
H -10.8596408 2.1741863 -59.3761475
H -9.4021933 2.1159166 -60.3858728
H -10.5024011 3.4948423 -60.5147045
C -10.2132689 4.5376623 -58.0986059
H -11.0746174 4.0426197 -57.649767
H -10.5543428 5.2422816 -58.85601
H -9.6730952 5.0878871 -57.3275553
N -5.4426723 0.9434196 -58.4481881
H -5.6678236 0.0355238 -58.0548346
C -4.0798935 1.185912 -58.9357536
H -3.8150141 2.2211407 -58.7113507
C -4.0051982 1.032849 -60.4518383
O -4.7678273 0.27695 -61.0434649
C -3.0949649 0.2568287 -58.2130025
H -3.5138127 -0.7490093 -58.1422002
H -2.1678002 0.2159683 -58.784929
C -2.795052 0.8141 -56.8097441
H -2.3119699 1.7866437 -56.9208895
H -3.7301621 0.9656837 -56.2690902
C -1.8893985 -0.0773955 -55.9502131
H -2.4545609 -0.9480272 -55.6154601
H -1.5950424 0.5033683 -55.0736083
C -0.6439782 -0.5463608 -56.7108241
H -0.3018319 0.268945 -57.3493463
H -0.9215663 -1.3830817 -57.3566641
N 0.4467047 -0.9339892 -55.7916971
H 0.7465728 -0.112789 -55.2498367
H 1.2748543 -1.2105249 -56.2979961
H 0.1611309 -1.6486113 -55.1440047
N -3.0487572 1.7262357 -61.0774219

H -2.4915023 2.3509615 -60.5015827
C -2.7384924 1.6213578 -62.5167704
H -3.2764944 0.767828 -62.9341887
C -1.2710996 1.2723612 -62.7035448
O -0.4066791 2.1504201 -62.6343866
C -3.1835379 2.9095297 -63.2545982
H -3.4195988 3.6894009 -62.5300726
H -2.3764105 3.2816492 -63.8842477
C -4.4157079 2.6815394 -64.1627795
H -4.7723658 3.6569256 -64.4963477
H -4.1117891 2.1108096 -65.0407656
C -5.5713926 1.9356428 -63.4710507
H -5.670715 2.3288555 -62.460506
H -5.3424209 0.8725846 -63.4125411
C -6.9189723 2.0674559 -64.1837241
H -6.9312356 1.427326 -65.0694601
H -7.0421595 3.108668 -64.4916204
N -8.0097774 1.7033435 -63.2510456
H -7.9605382 0.719054 -62.960874
H -8.919042 1.7965361 -63.6756161
H -7.9574248 2.2846703 -62.4272009
N -0.9851324 -0.0152724 -62.8685472
H -1.7507671 -0.6921228 -62.9343551
C 0.3622951 -0.5630426 -62.7563147
H 0.9805693 0.166144 -62.2307181
C 0.9623679 -0.7635467 -64.1569443
O 0.4536773 -1.5034089 -65.0035394
C 0.3662204 -1.8384641 -61.8795023
H -0.0630003 -2.6610733 -62.4548948
C 1.819163 -2.1735927 -61.5191769
H 2.2062048 -1.4454116 -60.8056847
H 2.4410637 -2.152216 -62.4072739
H 1.8767011 -3.1699076 -61.08193
C -0.4305112 -1.6738873 -60.5577608
H 0.0351023 -0.9039866 -59.9410057
H -1.442414 -1.3440835 -60.7786126
C -0.5557201 -2.9669112 -59.7395202
H 0.4082644 -3.2467547 -59.3150915
H -0.9233073 -3.7726035 -60.3764866
H -1.265722 -2.8133331 -58.927048
N 2.0820627 -0.0851478 -64.4029156
H 2.4655832 0.475942 -63.6575331
C 2.9068083 -0.2770199 -65.5962123
H 2.3018658 -0.6952979 -66.4020926
C 4.0396154 -1.2643317 -65.292497
O 4.4307098 -1.4269569 -64.1332412

C 3.4495725 1.0814523 -66.061202
H 4.0958151 0.9525892 -66.9322466
H 4.0432311 1.5292122 -65.261646
S 2.0709612 2.1846743 -66.4876026
H 1.3003227 1.8921115 -65.4342484
N 4.6448883 -1.8495044 -66.3268338
H 4.3466771 -1.6176306 -67.261478
C 5.9932657 -2.3856146 -66.1595621
H 6.0491473 -2.8706147 -65.1886549
C 7.0088041 -1.2347704 -66.1249314
O 6.737833 -0.132612 -66.6086652
C 6.3073177 -3.4524623 -67.2163512
H 5.4957053 -4.1766962 -67.2517385
H 7.2191217 -3.9815768 -66.9375472
C 6.5230252 -2.8364934 -68.5773457
O 7.5678347 -2.280934 -68.8635157
N 5.5086591 -2.8222407 -69.403895
H 5.6807276 -2.3748843 -70.3032807
H 4.6772054 -3.3460722 -69.2191437
N 8.1801732 -1.5131329 -65.5618081
H 8.3470506 -2.4508453 -65.2320084
C 9.2452561 -0.5331003 -65.3748175
H 8.8589516 0.286887 -64.7647094
C 9.6957767 0.098359 -66.7072817
O 9.7789872 1.3222783 -66.8014925
C 10.3873618 -1.2029495 -64.5780293
H 11.2574185 -0.549599 -64.5671677
H 10.0590078 -1.2929866 -63.5414857
C 10.8177317 -2.5865022 -65.0490904
C 11.9181375 -2.7365552 -65.9169673
H 12.4819757 -1.8755661 -66.2390297
C 12.3053206 -4.0136478 -66.363435
H 13.1452796 -4.1187607 -67.0335377
C 11.597087 -5.1547648 -65.9319127
O 11.9592972 -6.3881262 -66.3699191
H 12.7274721 -6.3595775 -66.9421564
C 10.5129368 -5.0118722 -65.040923
H 9.9908157 -5.8930459 -64.6965091
C 10.1282528 -3.7322316 -64.5973171
H 9.308215 -3.6379075 -63.8983316
N 9.914116 -0.6953223 -67.7660937
H 9.7185 -1.6813804 -67.6780474
C 10.4258322 -0.1969978 -69.0526211
H 11.3531195 0.3334907 -68.8684677
C 9.4744436 0.8147866 -69.7068052
O 9.9050969 1.903903 -70.0918449

C 10.7555701 -1.3866897 -69.9704476
H 11.4799712 -2.0264374 -69.4643361
H 9.8468928 -1.968167 -70.1290859
C 11.319076 -0.9912917 -71.3499645
H 10.5886703 -0.3825456 -71.8825983
C 12.633967 -0.2142643 -71.2488822
H 13.3625122 -0.779567 -70.6682336
H 12.462991 0.7532524 -70.7772734
H 13.0319279 -0.0350601 -72.2476385
C 11.5735498 -2.2547509 -72.1719436
H 12.3160219 -2.8816896 -71.6783682
H 11.9344282 -1.9816748 -73.1636924
H 10.6437644 -2.8131246 -72.2827093
N 8.1721243 0.5219023 -69.7300654
H 7.8781855 -0.3931025 -69.4008782
C 7.1627271 1.4694778 -70.216607
H 7.465206 1.8258281 -71.2019721
C 7.0485684 2.7218684 -69.344085
O 6.7977963 3.7974389 -69.8845881
C 5.8037654 0.7721156 -70.3591403
H 5.0142317 1.5249066 -70.3314356
H 5.6424419 0.0910384 -69.5229777
C 5.6722935 0.0300481 -71.6647418
N 5.9532087 -1.3209948 -71.8725283
C 5.7499517 -1.545272 -73.178947
H 5.897551 -2.4993019 -73.6695688
N 5.3500598 -0.4176791 -73.790252
H 5.1471925 -0.3273254 -74.7758818
C 5.2980715 0.5887433 -72.851667
H 5.0341347 1.6258243 -73.0140616
N 7.3164396 2.6332608 -68.0360767
H 7.5599259 1.7314697 -67.6420576
C 7.2614763 3.7968527 -67.1509479
H 6.3620564 4.3393293 -67.4291629
C 8.4261241 4.7738874 -67.3880553
O 8.1695259 5.9696373 -67.5199989
C 7.0948289 3.3363918 -65.6891448
H 6.3288467 2.5601209 -65.6683109
H 8.0271337 2.8928056 -65.3390609
C 6.6681347 4.4549309 -64.7117129
H 7.5189438 5.109165 -64.5258907
C 5.4997377 5.3156355 -65.2074584
H 4.6757884 4.679795 -65.5306189
H 5.8301918 5.9439793 -66.0348046
H 5.1677863 5.9791257 -64.4099764
C 6.2145039 3.8387536 -63.3851094

H 5.3016056 3.2618588 -63.5260214
H 6.0445249 4.6237662 -62.6487135
H 6.9882306 3.1722436 -63.0041928
N 9.6480168 4.2850584 -67.6533181
H 9.7997559 3.2846604 -67.5630426
C 10.7262591 5.1507473 -68.1801541
H 10.8557976 5.9933419 -67.499146
C 10.3952471 5.6900104 -69.5631661
O 10.7418516 6.824877 -69.8713061
C 12.0605472 4.4240518 -68.3016877
H 12.1166467 3.8610945 -69.2350101
H 12.121121 3.7293563 -67.4803064
O 13.1491082 5.3247477 -68.1948499
H 13.269569 5.8510611 -69.0056855
N 9.6793356 4.9112594 -70.3797769
H 9.4739327 3.9631262 -70.084374
C 9.1529553 5.3651188 -71.6612905
H 9.9800387 5.6169965 -72.3253587
H 8.5721268 4.5597261 -72.1096179
C 8.2436502 6.5889815 -71.5206075
O 8.4964877 7.5881455 -72.1829673
N 7.271716 6.5802667 -70.6017955
H 7.1008963 5.7179054 -70.0934582
C 6.3753098 7.7248641 -70.3564864
H 5.913338 8.0175862 -71.2986071
C 7.1064366 8.967594 -69.8279953
O 6.7901523 10.0960769 -70.2162708
C 5.2706065 7.3106618 -69.3736671
H 5.7289851 6.9026424 -68.4718638
H 4.7203521 8.2063148 -69.0832355
C 4.2527546 6.3259864 -69.9274463
C 3.9569052 5.1377987 -69.2312677
H 4.470828 4.90886 -68.3126257
C 2.9868958 4.2463839 -69.7247382
H 2.7656118 3.3367646 -69.1875547
C 2.2932897 4.5473782 -70.9100298
H 1.5351606 3.8746796 -71.2849157
C 2.5749518 5.7375725 -71.6014384
H 2.0297721 5.995317 -72.5016579
C 3.5527959 6.6201892 -71.1146242
H 3.7383022 7.5375435 -71.6559592
N 8.1333138 8.7500029 -69.0098282
H 8.2811018 7.8069615 -68.6678179
C 9.07562 9.7716512 -68.5730746
H 8.5547353 10.5922679 -68.0912188
C 9.8605926 10.3853852 -69.7513712

O 9.7566054 11.5847949 -70.0065046
C 9.9597189 9.1076336 -67.5219712
H 10.8339721 9.7058895 -67.3759088
H 10.2895408 8.1279684 -67.8552867
S 9.0625811 8.9471298 -65.9503501
H 9.8351265 9.8474364 -65.3019252
N 10.4805271 9.5504548 -70.5898632
H 10.5030738 8.5688311 -70.3340113
C 11.2121939 9.9579073 -71.8004392
H 12.0155281 10.6371849 -71.5108146
C 10.3189662 10.6842638 -72.8220458
O 10.6706195 11.7496569 -73.3218699
C 11.8287769 8.7012708 -72.4308687
H 11.0787146 8.1829872 -73.0301747
H 12.1581597 8.0264929 -71.641054
O 12.9535704 9.016732 -73.2287619
H 13.2151123 8.2260534 -73.7115009
N 9.0917743 10.1931924 -73.0341901
H 8.8462209 9.3288767 -72.56349
C 8.0758771 10.7997598 -73.908273
H 8.5230395 10.9933282 -74.8827677
C 7.5601152 12.1572996 -73.4059811
O 6.9709603 12.9057875 -74.1813275
C 6.8980951 9.8177845 -74.0665242
H 6.5963806 9.4797408 -73.0746319
H 6.0469815 10.3468555 -74.4990079
C 7.2010356 8.596359 -74.9559661
H 8.1359891 8.1311667 -74.6495739
C 6.078985 7.5667695 -74.8172612
H 5.1265064 8.0041828 -75.1217947
H 6.002105 7.244648 -73.7790986
H 6.2918725 6.6974123 -75.4373065
C 7.3195826 8.9722824 -76.4354788
H 6.4050387 9.4644876 -76.7684964
H 7.4802548 8.0744735 -77.031481
H 8.1643764 9.6422728 -76.586479
N 7.8083199 12.5056056 -72.1428918
H 8.3267403 11.8644839 -71.5540116
C 7.4700463 13.8201725 -71.5844624
H 6.6249218 14.2248713 -72.140381
C 8.5965015 14.8537852 -71.7283394
O 8.4412966 15.9715581 -71.2569983
C 7.0096966 13.6702722 -70.1392581
H 6.7278152 14.646009 -69.7435907
H 7.8241777 13.2648047 -69.5378654
O 5.8818799 12.8084382 -70.0899695

H 6.1934362 11.8939971 -70.2199382
N 9.6527789 14.5194905 -72.4726668
H 9.6609405 13.583641 -72.8564542
C 10.8450282 15.3307806 -72.7823003
H 11.3215018 14.7681748 -73.5857787
C 11.9223128 15.3449434 -71.6844862
O 12.7208264 16.2703293 -71.5460031
C 10.4885248 16.6983789 -73.4123986
H 10.3052822 17.4216129 -72.6154718
H 9.5630553 16.5886969 -73.9810148
C 11.5483248 17.2610639 -74.3780375
O 12.3958398 16.4764423 -74.8659835
O 11.4371665 18.4657644 -74.7049311
N 11.9834183 14.26542 -70.8998863
H 11.310854 13.5258562 -71.0438426
C 13.0028759 14.0912496 -69.8714545
H 13.306885 15.0728525 -69.5156226
C 14.2829326 13.4339092 -70.3807713
O 14.2769208 12.3465484 -70.963321
C 12.4106606 13.3349922 -68.6923727
H 12.0504773 12.3665639 -69.0383334
H 13.215576 13.1470367 -67.9823596
C 11.3089208 14.0511588 -67.9372688
C 10.4143586 13.2895275 -67.1652494
H 10.4207411 12.2129036 -67.2452246
C 9.622269 13.9020251 -66.183757
H 9.0420531 13.2912381 -65.5123696
C 9.6896914 15.2939169 -66.0117228
O 9.0688287 15.8810816 -64.9659394
H 8.6517885 15.2197967 -64.4135997
C 10.476267 16.0688853 -66.8777947
H 10.5008894 17.1334103 -66.7732363
C 11.2869235 15.4568867 -67.8389113
H 11.9309688 16.084664 -68.4366982
N 15.413457 14.0666489 -70.0648519
H 15.3470006 14.8954175 -69.4865486
C 16.7466287 13.6072225 -70.4479379
H 16.6439033 13.0148765 -71.3578561
C 17.3845581 12.7089696 -69.3861672
O 17.0498046 12.785335 -68.2019845
C 17.6340751 14.8021464 -70.8021027
H 17.0888067 15.4656922 -71.4759625
H 18.5250258 14.4458703 -71.3186039
O 18.0387931 15.5263655 -69.6592734
H 17.3011331 15.5682376 -69.0191956
N 18.3085085 11.8526167 -69.839663

H 18.540374 11.8840546 -70.8182419
C 19.0346307 10.8940314 -69.0057114
H 19.4675616 10.1472293 -69.6713046
C 18.1121144 10.1118048 -68.0476025
O 18.3664568 10.0593789 -66.8463913
C 20.2060694 11.6112569 -68.3167241
H 19.8233584 12.3411857 -67.6012615
H 20.822286 12.1241624 -69.055329
H 20.8190484 10.8836954 -67.7848943
N 17.0082147 9.5547067 -68.5680509
H 16.8205954 9.6340163 -69.5538031
C 16.1258953 8.7216485 -67.7571872
H 16.0455573 9.2076642 -66.7871337
C 16.7726785 7.3640399 -67.4612841
O 17.037384 6.5613184 -68.3551552
C 14.6792575 8.6592675 -68.293561
H 14.2122908 9.6269136 -68.1101076
H 14.1203773 7.931987 -67.704949
C 14.479784 8.3334479 -69.7606206
O 14.1690558 7.2130685 -70.1375173
N 14.5042034 9.3331716 -70.6191546
H 14.1908225 9.1433714 -71.5603483
H 14.5384129 10.2922772 -70.2963576
N 17.0392665 7.1424113 -66.177116
H 16.830673 7.8928146 -65.5251381
C 17.5157217 5.8978433 -65.5954028
H 17.9177789 5.2616605 -66.384763
C 16.3357946 5.207524 -64.9288189
O 15.6159122 5.8225069 -64.1466295
C 18.6212937 6.1737034 -64.5672921
H 18.1711266 6.6156535 -63.6769974
H 19.0702786 5.2249355 -64.2716113
C 19.7093446 7.0913046 -65.0380035
C 19.7931824 8.4044157 -64.7311366
H 19.0720161 8.9404149 -64.124824
N 20.8504143 8.9783652 -65.4059943
H 20.9953366 9.9773354 -65.4322483
C 21.4912532 8.0630289 -66.2111008
C 22.5607112 8.1678796 -67.1126483
H 23.0547657 9.1163029 -67.2628505
C 22.976687 7.0234162 -67.8144749
H 23.7994497 7.0845023 -68.5131482
C 22.3164877 5.7977862 -67.6094385
H 22.6337933 4.9207785 -68.1562422
C 21.2368826 5.7069224 -66.707767
H 20.7248034 4.766601 -66.5741431

C 20.7952953 6.8365602 -65.9837824
N 16.1848563 3.912817 -65.176291
H 16.821799 3.469261 -65.8205527
C 15.0796393 3.1104436 -64.6775838
H 14.54308 3.6333506 -63.8833365
C 15.6247004 1.7985474 -64.1325112
O 16.0577958 0.9402323 -64.9020713
C 14.131694 2.8887033 -65.8487103
H 13.64995 3.8395378 -66.0623428
H 14.7289034 2.6128478 -66.7202805
C 13.0901348 1.8002141 -65.5654381
H 13.4487714 1.0489044 -64.8679001
C 11.8698551 2.4330043 -64.9143906
H 11.3898081 3.1481945 -65.575671
H 12.1699034 2.9421886 -63.9979327
H 11.1508884 1.6662489 -64.6483251
C 12.8433832 1.0456084 -66.8611406
H 12.3756177 1.6885177 -67.5980209
H 12.2143535 0.1981272 -66.6495824
H 13.7858171 0.6635186 -67.2549475
N 15.5022455 1.6123315 -62.8227831
H 15.0176951 2.3212374 -62.2749974
C 15.9819101 0.4156113 -62.1354336
H 16.5767406 -0.1688088 -62.8309248
C 14.7948239 -0.412656 -61.638592
O 14.0877462 0.0314062 -60.7298387
C 16.9228566 0.7713672 -60.9737509
H 16.3487296 1.1324943 -60.119959
C 17.7625409 -0.435787 -60.5611816
H 18.3692173 -0.7740933 -61.4018301
H 17.1180753 -1.2500821 -60.2345302
H 18.4224701 -0.1512396 -59.7425426
O 17.83844 1.7718527 -61.3585353
H 17.3314862 2.5714806 -61.5236629
N 14.5757734 -1.6381667 -62.1522123
C 13.7832369 -2.6243789 -61.4326093
H 12.8078798 -2.2156713 -61.1651937
C 14.5459745 -3.0172483 -60.1610187
O 15.6270337 -3.5983735 -60.2304277
C 13.5987399 -3.7952014 -62.405327
H 13.5382697 -4.7567764 -61.8952256
H 12.6985773 -3.6284725 -62.9974839
C 15.1494966 -2.2202696 -63.3610773
H 14.6833669 -1.7644417 -64.236083
H 16.2312348 -2.0961457 -63.4033807
C 14.823558 -3.7137458 -63.3158808

H 14.6132543 -4.1093829 -64.3096971
H 15.6551597 -4.2543635 -62.8618106
N 13.9662979 -2.746838 -58.9908767
H 13.0661721 -2.2754632 -59.0090904
C 14.5182136 -3.1102436 -57.6789229
H 15.4998097 -3.5591397 -57.8229977
C 13.7014467 -4.2044761 -56.9428997
O 13.6238671 -4.1604091 -55.7146881
C 14.714941 -1.81596 -56.8514598
H 13.7756341 -1.2643591 -56.8431172
H 14.9534945 -2.0737327 -55.819355
C 15.8229594 -0.873091 -57.3541501
H 15.6404761 -0.6146515 -58.3959849
C 15.8120489 0.4127376 -56.5303236
H 16.028931 0.1980959 -55.4826737
H 14.8309305 0.8833728 -56.593059
H 16.5554212 1.1109018 -56.9145674
C 17.2136708 -1.5018124 -57.2217831
H 17.3755269 -1.8389114 -56.1975999
H 17.9750578 -0.7631874 -57.470298
H 17.3151205 -2.3433035 -57.9050428
N 13.0948036 -5.214763 -57.6151661
C 12.2053006 -6.1721577 -56.9514439
H 11.3888186 -5.644905 -56.4617025
C 12.9249789 -7.0386965 -55.9075064
O 12.3195788 -7.4085346 -54.9111168
C 11.6162382 -7.0363963 -58.0682371
H 11.3764525 -8.0460584 -57.7312578
H 10.7270492 -6.5549761 -58.4702904
C 13.2765494 -5.6211828 -59.0043124
H 12.7053484 -4.9577645 -59.6524598
H 14.33053 -5.6213029 -59.2835882
C 12.7120005 -7.0340446 -59.1282169
H 12.3119336 -7.2235629 -60.1252091
H 13.4796364 -7.7662537 -58.8743209
N 14.2193922 -7.3309605 -56.0875591
H 14.6758111 -6.9951267 -56.9187628
C 15.0103271 -8.0990672 -55.112049
H 14.434692 -8.9910839 -54.8655987
C 15.2070223 -7.332927 -53.7964256
O 15.0766997 -7.9116657 -52.717904
C 16.3497863 -8.552893 -55.7413196
H 16.1067111 -9.1238207 -56.6393262
C 17.2457079 -7.3784555 -56.1790683
H 17.6104389 -6.833258 -55.3064419
H 16.7096291 -6.695959 -56.8363059

H 18.1109972 -7.7622999 -56.7213065
C 17.1758962 -9.4774166 -54.8192359
H 18.0864798 -9.7670527 -55.3455807
H 17.4716894 -8.935527 -53.9193684
C 16.4479887 -10.7630839 -54.4053729
H 16.0929498 -11.2919093 -55.2902479
H 15.605773 -10.5327737 -53.7532482
H 17.1389073 -11.4070984 -53.8607548
N 15.3935547 -6.0141343 -53.87863
H 15.4044782 -5.57695 -54.785056
C 15.4989416 -5.1573755 -52.7005961
H 16.1260958 -5.6549509 -51.9609007
C 14.1275042 -4.979588 -52.0493859
O 14.0041547 -5.2410058 -50.8625342
C 16.1529791 -3.8128398 -53.0612849
H 15.5453294 -3.3096024 -53.8136461
H 16.1749842 -3.1849001 -52.1697456
C 17.5888162 -3.9565456 -53.5969701
H 17.9604463 -2.9636573 -53.8583359
H 17.5825816 -4.5500643 -54.513358
C 18.5239075 -4.5941443 -52.560969
O 18.5957494 -5.8428049 -52.4662396
O 19.14147 -3.8633975 -51.7551244
N 13.0718701 -4.7526557 -52.834066
H 13.246795 -4.5814834 -53.8192455
C 11.6674641 -4.7573474 -52.3915481
H 11.5110988 -3.9424476 -51.6838844
C 11.2638044 -6.055375 -51.6705936
O 10.7000723 -5.9916155 -50.5815018
C 10.8225721 -4.4658085 -53.6373247
H 11.2703018 -4.9725609 -54.4847155
H 10.8980451 -3.3996945 -53.8162933
C 9.3400869 -4.8422281 -53.6032018
O 8.5960027 -4.4336681 -52.6834481
O 8.8625046 -5.4577929 -54.5871172
N 11.6917214 -7.2238873 -52.1577427
H 12.1378338 -7.2256039 -53.0701622
C 11.4803921 -8.5084706 -51.4804795
H 10.4192842 -8.5902366 -51.2439924
C 12.2581264 -8.616735 -50.1576347
O 11.7173718 -9.1125472 -49.1729519
C 11.8590517 -9.6523727 -52.4351888
H 12.9441046 -9.6885066 -52.5437256
H 11.4310558 -9.4628766 -53.4176041
C 11.3583046 -11.0199242 -51.9562242
H 11.7482812 -11.2164564 -50.9570363

H 11.7634607 -11.7769659 -52.6281464
S 9.5513028 -11.2041661 -51.9296248
C 9.4280423 -12.9497713 -51.463499
H 8.3802362 -13.2482151 -51.4521366
H 9.8585708 -13.0948903 -50.4720088
H 9.9653893 -13.5614913 -52.1881289
N 13.4992124 -8.1155487 -50.1013757
H 13.8751445 -7.7085546 -50.9531837
C 14.3462607 -8.0992723 -48.8917458
H 14.3261654 -9.0880359 -48.4302966
C 13.8414349 -7.109273 -47.8321094
O 13.8696885 -7.4088854 -46.6438106
C 15.7803139 -7.7771225 -49.3386629
H 16.0728206 -8.4953921 -50.1072518
H 15.78842 -6.7853287 -49.7877521
C 16.8146607 -7.8333564 -48.2019743
H 16.5459291 -7.1366515 -47.4067893
H 16.8280532 -8.8437172 -47.7908348
C 18.2187971 -7.4873246 -48.7193819
H 18.4184365 -8.0829343 -49.6128579
H 18.9536731 -7.7499724 -47.9570584
C 18.3487833 -5.992413 -49.0458139
H 18.4319706 -5.4217815 -48.1165061
H 17.4451803 -5.6561907 -49.5619324
N 19.5031294 -5.7293883 -49.9309239
H 20.3954323 -5.9966135 -49.5549232
H 19.5174901 -4.749167 -50.2346865
H 19.3464276 -6.1609146 -50.848426
N 13.3322727 -5.9630881 -48.2710839
H 13.3916351 -5.7966786 -49.2700395
C 12.6752334 -4.9259791 -47.4652083
H 13.3066394 -4.639705 -46.6204001
C 11.3514425 -5.4659331 -46.907204
O 11.0801258 -5.3692947 -45.708615
C 12.463281 -3.6936637 -48.3766892
H 12.0638674 -4.0566426 -49.3241162
C 11.3939495 -2.735481 -47.8251092
H 11.5699158 -2.5482408 -46.7698325
H 10.4027027 -3.1721763 -47.9439616
H 11.4167651 -1.795219 -48.3641339
C 13.8090055 -3.0011842 -48.7056933
H 14.1943818 -2.5034339 -47.8276267
H 14.5553025 -3.7476582 -48.9678682
C 13.7415405 -1.9940272 -49.8643963
H 13.1388901 -1.128049 -49.5957665
H 13.3172396 -2.4665957 -50.7493734

H 14.7489847 -1.6485579 -50.0979748
N 10.5569209 -6.1251239 -47.7575478
H 10.8058184 -6.1502801 -48.7417588
C 9.3406062 -6.8155937 -47.3499945
H 8.687436 -6.0980664 -46.8515223
C 9.6451199 -7.9320667 -46.3426887
O 8.9789309 -7.9857565 -45.3181334
C 8.6090851 -7.3439192 -48.5900398
H 9.2347282 -8.0646011 -49.1166765
H 8.3743305 -6.5167723 -49.2610325
H 7.6825766 -7.8315087 -48.286617
N 10.6967849 -8.7327052 -46.5459848
H 11.1877227 -8.6812436 -47.4322827
C 11.1410652 -9.7413254 -45.5766499
H 10.3095113 -10.4188421 -45.3765601
C 11.5417736 -9.1181235 -44.2316211
O 11.0027769 -9.5243077 -43.2065835
C 12.2879134 -10.5821685 -46.1574739
H 13.1383181 -9.9410307 -46.3900585
H 11.9539012 -11.054893 -47.0827182
C 12.7241101 -11.6616207 -45.1622198
O 13.5224159 -11.3337599 -44.252894
O 12.2018212 -12.793503 -45.265959
N 12.3829782 -8.0769071 -44.2118859
H 12.8033958 -7.7736289 -45.0842588
C 12.8418208 -7.4756884 -42.9545116
H 13.20788 -8.2795145 -42.315361
C 11.7105569 -6.7843906 -42.179033
O 11.5105007 -7.0762136 -40.995856
C 14.0107534 -6.5158331 -43.2086705
H 14.8199205 -7.0620265 -43.6953675
H 13.6867322 -5.7238326 -43.8863098
C 14.5352889 -5.889822 -41.9505213
C 15.3513898 -6.4920806 -41.0560262
H 15.7532765 -7.4912347 -41.1718495
N 15.5335855 -5.6654725 -39.9629168
H 16.0897329 -5.9158626 -39.160673
C 14.8148192 -4.49376 -40.0872147
C 14.6321954 -3.3866221 -39.242576
H 15.1032791 -3.3500841 -38.2735638
C 13.819259 -2.3236695 -39.676404
H 13.6551516 -1.4597462 -39.0434416
C 13.2231146 -2.3758491 -40.9490175
H 12.61592 -1.5480294 -41.2963996
C 13.413713 -3.4925454 -41.785488
H 12.9690569 -3.4997165 -42.7718436

C 14.1985453 -4.5894945 -41.3712467
N 10.9043925 -5.9527161 -42.8492866
H 11.1134384 -5.7630804 -43.8240366
C 9.7438538 -5.2921988 -42.2212211
H 10.0844229 -4.773692 -41.3238945
C 8.6828352 -6.2978319 -41.7797415
O 8.1475363 -6.1684389 -40.6765001
C 9.0796136 -4.2487583 -43.1304959
H 8.177974 -3.8895772 -42.6371761
C 9.9720763 -3.0467225 -43.4104666
H 10.8669468 -3.3593763 -43.9445606
H 10.2661708 -2.5763767 -42.4714854
H 9.4314004 -2.3171273 -44.0117579
O 8.7181591 -4.8066588 -44.3726664
H 9.5496823 -4.9479732 -44.8577988
N 8.4358032 -7.3581704 -42.5633335
H 8.8963046 -7.4314515 -43.466883
C 7.5723311 -8.4495715 -42.1268535
H 6.6350766 -8.0155664 -41.7795424
C 8.1700975 -9.1517994 -40.9247922
O 7.4758181 -9.174459 -39.9330543
C 7.2448461 -9.4620492 -43.2330206
H 8.1639771 -9.7685181 -43.7320877
H 6.8107539 -10.3548548 -42.779457
C 6.239774 -8.9211213 -44.2597927
H 6.2060887 -9.6067299 -45.1067549
H 6.5649357 -7.9487869 -44.6272582
C 4.8301142 -8.7907822 -43.6962677
O 4.0185217 -9.6936336 -43.7800733
N 4.4373901 -7.6329512 -43.2092787
H 3.4549186 -7.5555601 -42.9669857
H 5.0762198 -6.8618807 -43.140931
N 9.4273277 -9.6006942 -40.9284614
H 9.9686587 -9.4839231 -41.780828
C 10.0214635 -10.412134 -39.8567427
H 9.3084547 -11.1990181 -39.609992
C 10.253828 -9.6601678 -38.5291229
O 10.2250454 -10.2893742 -37.4666527
C 11.2944858 -11.0964365 -40.3945858
H 11.8618845 -11.5134649 -39.5643313
H 11.9244235 -10.3674132 -40.904203
C 10.9694015 -12.2521054 -41.3360884
O 10.0165345 -12.9890257 -41.1245311
N 11.7728694 -12.4937022 -42.343769
H 11.4995019 -13.1733068 -43.0362986
H 12.499172 -11.8467642 -42.6449554

N 10.2929531 -8.3240194 -38.537167
H 10.3624872 -7.8437779 -39.4304161
C 10.2075646 -7.5261957 -37.311539
H 10.9370343 -7.9083139 -36.5955343
C 8.8173012 -7.6078404 -36.6366541
O 8.7224682 -7.6976754 -35.4119872
C 10.5911494 -6.0827475 -37.6571958
H 9.8848154 -5.6694188 -38.3793169
H 11.5923603 -6.0574983 -38.0909011
H 10.5793001 -5.471428 -36.7549126
N 7.7281362 -7.648853 -37.4093658
H 7.8464977 -7.6620426 -38.4175533
C 6.3615845 -7.7065981 -36.8837774
H 6.2525433 -6.8732437 -36.1891368
C 5.9991926 -8.9928891 -36.0905116
O 5.3798625 -8.8602907 -35.0312186
C 5.3875801 -7.4605357 -38.0417105
H 5.317507 -8.3466364 -38.6717451
H 5.7360643 -6.6360806 -38.6607618
H 4.3964641 -7.2320804 -37.651936
N 6.3537059 -10.227742 -36.5201682
C 6.0921003 -11.4546801 -35.7992746
H 5.0595046 -11.4728627 -35.4552056
C 7.0141917 -11.5868196 -34.5825291
O 6.5442037 -12.022288 -33.5387212
C 6.2944393 -12.5878507 -36.8162142
H 6.5796654 -13.5239315 -36.3340225
H 5.3790469 -12.7278175 -37.3944718
C 6.9710996 -10.6108314 -37.7595143
H 6.2311127 -10.4906961 -38.55036
H 7.8621115 -10.0271148 -37.9027458
C 7.3852351 -12.0701274 -37.7139111
H 7.3632158 -12.5333561 -38.7012058
H 8.3603036 -12.1870356 -37.2408137
N 8.2600128 -11.0993172 -34.6473682
H 8.5974122 -10.7351963 -35.529587
C 9.1638845 -11.0324667 -33.493428
H 9.2559856 -12.0264496 -33.0531509
C 8.6246533 -10.1030979 -32.3918366
O 8.5898426 -10.4804776 -31.2234502
C 10.5463395 -10.5906214 -33.9929249
H 10.9400176 -11.3588206 -34.6604347
H 10.4349591 -9.6795107 -34.5821898
C 11.5645544 -10.3336599 -32.8982995
C 12.1676299 -11.4139774 -32.226674
H 11.9007571 -12.4282594 -32.4853161

C 13.115578 -11.1764532 -31.2147571
H 13.5750579 -12.007349 -30.6985902
C 13.464218 -9.8578058 -30.873536
H 14.194 -9.6750549 -30.0974929
C 12.8624226 -8.7769348 -31.541472
H 13.1324783 -7.7636193 -31.281157
C 11.9107974 -9.0133361 -32.5496745
H 11.4482941 -8.1777601 -33.0568941
N 8.0603085 -8.9495652 -32.7631566
H 8.1434311 -8.665565 -33.7337643
C 7.4168677 -8.0211198 -31.8271501
H 7.9214617 -8.1082216 -30.8649975
C 5.9315139 -8.326438 -31.5456683
O 5.2535744 -7.4801421 -30.961817
C 7.5941536 -6.574922 -32.324633
H 7.124499 -6.4941971 -33.3064549
H 7.0538484 -5.9068789 -31.6550215
C 9.0136829 -6.0386909 -32.3987931
C 9.4463363 -5.359517 -33.5561945
H 8.7912065 -5.2787379 -34.4129759
C 10.7433545 -4.8150995 -33.6203321
H 11.075635 -4.3075551 -34.5142454
C 11.6148391 -4.9453172 -32.5180674
O 12.8717441 -4.435554 -32.5709646
H 13.0345853 -3.9454995 -33.3800472
C 11.174962 -5.6029107 -31.3511048
H 11.8453005 -5.6829437 -30.507972
C 9.8769556 -6.144496 -31.2893049
H 9.5563376 -6.6474406 -30.3872537
N 5.3871766 -9.481238 -31.956426
H 6.0033601 -10.1538206 -32.4006634
C 3.9274016 -9.6951489 -32.0875758
H 3.5788743 -9.0167038 -32.8656313
C 3.101727 -9.3795924 -30.8315989
O 1.9946272 -8.8568 -30.9597617
C 3.6184426 -11.1199467 -32.5819641
H 4.1373467 -11.2775613 -33.5237867
C 4.0262775 -12.229422 -31.6076246
H 3.4924745 -12.148523 -30.6645446
H 5.0959923 -12.1726016 -31.405786
H 3.820187 -13.1996915 -32.0601955
O 2.2341245 -11.2757929 -32.8256162
H 2.1031661 -11.0808149 -33.7675107
N 3.6312656 -9.6287895 -29.6257628
H 4.5649006 -10.0097299 -29.5872965
C 2.9405287 -9.3177676 -28.3687039

H 1.9221669 -9.6969241 -28.445873
C 2.8276563 -7.8017421 -28.1507249
O 1.7167741 -7.2833488 -28.1555535
C 3.6362754 -10.0368499 -27.1973246
H 3.6353806 -11.1098669 -27.3948457
H 4.6745595 -9.7054275 -27.1474315
C 2.978396 -9.7839054 -25.8233671
H 2.9966347 -8.7178673 -25.6005677
C 1.5310448 -10.2812004 -25.7697653
H 1.4817257 -11.3377576 -26.0293787
H 0.9096754 -9.6938931 -26.4446131
H 1.1425322 -10.1385847 -24.7604909
C 3.7720642 -10.514862 -24.7416685
H 3.7563083 -11.5903824 -24.9153329
H 3.3368301 -10.2987848 -23.7649057
H 4.8026104 -10.1599394 -24.7388168
N 3.9616382 -7.0915666 -28.1298216
H 4.8305866 -7.5964343 -28.1841824
C 4.0327294 -5.6286734 -27.9990707
H 3.5475118 -5.3373421 -27.066531
C 3.3223149 -4.9052738 -29.1588863
O 2.7887892 -3.8112358 -29.0000078
C 5.522688 -5.2420034 -27.9073454
H 5.9417744 -5.7125461 -27.0159418
H 6.0387796 -5.6508386 -28.77724
C 5.8148372 -3.7290084 -27.839953
H 5.4302001 -3.2507583 -28.740477
C 5.2052008 -3.0369301 -26.6265279
H 5.5270028 -3.5308572 -25.7079944
H 4.1165442 -3.0902357 -26.6670599
H 5.5002705 -1.9890255 -26.5938106
C 7.3263404 -3.5054407 -27.8182893
H 7.7487717 -3.9367071 -26.9075049
H 7.5460048 -2.4366408 -27.8259105
H 7.7977514 -3.9704279 -28.681937
N 3.3240981 -5.4922525 -30.3565642
H 3.8200164 -6.3736162 -30.4481514
C 2.6217824 -4.9657189 -31.5266407
H 2.8835245 -3.9148142 -31.6417372
C 1.0943839 -5.029999 -31.3514762
O 0.4108227 -4.0473604 -31.6319395
C 3.127867 -5.7370606 -32.7582961
H 4.209855 -5.607543 -32.8206349
H 2.9292394 -6.7975108 -32.6056175
C 2.5111026 -5.3318787 -34.1077532
H 1.4532277 -5.5947615 -34.1197899

C 2.6686143 -3.8435188 -34.4097948
H 3.7203921 -3.5715168 -34.3494023
H 2.0853599 -3.2554452 -33.704076
H 2.2981698 -3.6320755 -35.4139741
C 3.226156 -6.095398 -35.2194507
H 4.2560045 -5.7514367 -35.3113525
H 2.7083162 -5.937219 -36.1650524
H 3.2404669 -7.1588567 -34.9842874
N 0.5617721 -6.1321914 -30.8100106
H 1.1854655 -6.8955181 -30.565335
C -0.8774119 -6.2932057 -30.5294087
H -1.4450707 -5.8133694 -31.3273322
C -1.3387857 -5.5989339 -29.2472027
O -2.4972833 -5.2107496 -29.1799714
C -1.2259879 -7.7857952 -30.5059816
H -0.5395652 -8.3161481 -29.8425807
H -2.2423577 -7.9080194 -30.1304602
C -1.1550235 -8.3653732 -31.9235204
H -0.1595575 -8.2283015 -32.3457716
H -1.8592111 -7.825137 -32.5503861
C -1.517416 -9.8499764 -31.9535904
H -1.7789138 -10.1101266 -32.9786297
H -2.3926915 -10.0247935 -31.3275004
N -0.3815104 -10.6888054 -31.5472774
H 0.4686067 -10.5891001 -32.0922565
C -0.2819976 -11.5284697 -30.5397689
N 0.7987728 -12.2399241 -30.4453776
H 0.9213052 -12.9191248 -29.7195572
H 1.4926744 -12.1243647 -31.1788985
N -1.2140364 -11.691179 -29.6413828
H -2.0524106 -11.1461476 -29.7056909
H -1.0837421 -12.3425586 -28.8909044
N -0.4359914 -5.4137744 -28.291648
H 0.4565919 -5.878876 -28.4106519
C -0.6121594 -4.6637386 -27.0405899
H -1.5484464 -4.9448234 -26.5574101
C -0.6419177 -3.1442925 -27.2699944
O -1.4719593 -2.454789 -26.6849815
C 0.5604857 -5.0831334 -26.1452238
H 0.4619065 -6.1502756 -25.9371324
H 1.4675594 -4.941078 -26.7285774
C 0.7664583 -4.3591921 -24.8161269
H -0.1072872 -4.5150072 -24.1788573
H 0.8875321 -3.2879945 -24.9917974
C 2.035639 -4.9226683 -24.1581527
O 1.8929473 -5.6917077 -23.1831504

O 3.1384206 -4.6239782 -24.6732221
N 0.1917402 -2.6286023 -28.1822229
H 0.9142711 -3.2259607 -28.5641261
C 0.1832659 -1.2150978 -28.5658876
H 0.0037262 -0.6137299 -27.6722889
C -0.9246664 -0.8558517 -29.5768898
O -1.3427934 0.2986789 -29.6289793
C 1.5712314 -0.8568396 -29.111109
H 1.7755088 -1.4407739 -30.0082925
H 2.3275326 -1.0683264 -28.3535512
H 1.5991124 0.2052893 -29.3567835
N -1.3690744 -1.8151607 -30.399589
H -0.9570034 -2.7323259 -30.3107115
C -2.4122034 -1.6197109 -31.4130676
H -2.8401389 -0.6237023 -31.2850044
C -3.5838278 -2.5891446 -31.1989809
O -4.4984085 -2.2893246 -30.4387788
C -1.8158924 -1.6685056 -32.833723
H -1.3145361 -2.624073 -32.987596
H -2.63415 -1.6141187 -33.553035
C -0.8409829 -0.5505884 -33.1512774
C -1.3096435 0.649862 -33.7150379
H -2.3647469 0.7833783 -33.9051926
C 0.531843 -0.7025696 -32.8802269
H 0.8864084 -1.6149005 -32.4260483
C -0.4110316 1.6879453 -34.01348
H -0.7806818 2.6123909 -34.4308109
C 1.4300197 0.3388147 -33.1757978
H 2.4760809 0.2347951 -32.9481857
C 0.9602995 1.5313944 -33.7512922
H 1.6466328 2.3356076 -33.9776088
N -3.6205628 -3.7132496 -31.9236746
H -2.8479066 -3.9034345 -32.5436994
C -4.6046085 -4.7861899 -31.7255437
H -4.6477881 -5.0219769 -30.6606922
C -4.2176418 -6.0541322 -32.4919207
O -3.4466607 -6.006089 -33.4557384
C -6.0082382 -4.3507814 -32.1811528
H -6.3304691 -3.4699273 -31.624698
H -6.7175425 -5.152142 -31.9705056
O -6.0325689 -4.0692292 -33.5679786
H -5.899994 -3.1163384 -33.6518697
N -4.8741413 -7.1737827 -32.163381
H -5.4716506 -7.1487112 -31.3494907
C -4.8665038 -8.4037354 -32.9751168
H -3.8421984 -8.7622554 -33.024553

C -5.296619 -8.1459118 -34.4291164
O -4.6251069 -8.5946963 -35.3565651
C -5.7191938 -9.4992887 -32.2851475
H -5.2617698 -9.6962688 -31.3135578
C -7.1749464 -9.0643969 -32.0189038
H -7.7180113 -8.9252308 -32.9539572
H -7.2157387 -8.1483551 -31.4309694
H -7.6857446 -9.8419113 -31.4478146
C -5.7591062 -10.8389807 -33.0521625
H -6.3486961 -11.5529072 -32.4750014
H -6.2623358 -10.6958331 -34.0075932
C -4.3868889 -11.4732058 -33.3065014
H -3.86465 -11.6184485 -32.3614838
H -3.7929475 -10.8453952 -33.9693974
H -4.5241808 -12.4437534 -33.7840225
N -6.3197801 -7.3081212 -34.6459337
H -6.7867577 -6.9174776 -33.8407924
C -6.8108968 -6.9429317 -35.9822531
H -6.9295796 -7.8697976 -36.5438468
C -5.8010042 -6.0946841 -36.7703672
O -5.7534856 -6.2049544 -37.9993011
C -8.1929232 -6.2678168 -35.828368
H -8.8936588 -6.9935093 -35.4113768
H -8.1035006 -5.4462128 -35.1147961
C -8.795135 -5.6932521 -37.1238768
H -9.7559562 -5.2332397 -36.8894435
H -8.1411857 -4.8942535 -37.4703831
C -9.0162358 -6.7421972 -38.2346595
H -9.9843958 -7.2211919 -38.076221
H -8.2596339 -7.5233973 -38.1830747
C -8.9707821 -6.1540308 -39.6519625
H -9.7832162 -5.4324325 -39.7803459
H -9.1312743 -6.9749907 -40.3575952
N -7.6582374 -5.5172426 -39.9201159
H -7.5238449 -4.6979612 -39.3466649
H -7.5232324 -5.260878 -40.8933727
H -6.8930147 -6.1433565 -39.6825251
N -4.9831557 -5.2914209 -36.0868715
H -5.1114733 -5.2316311 -35.0833861
C -3.8621657 -4.5557308 -36.6755995
H -4.2125805 -3.9476157 -37.5101316
H -3.4303927 -3.8987074 -35.9202956
C -2.7652385 -5.4947951 -37.177779
O -2.4814048 -5.5085537 -38.373279
N -2.2677174 -6.3806718 -36.3070245
H -2.5945535 -6.342489 -35.3479413

C -1.2607822 -7.3958332 -36.6512525
H -0.3445124 -6.8925585 -36.961776
C -1.6939054 -8.2984128 -37.8176158
O -0.9311845 -8.4717807 -38.7673839
C -0.9587692 -8.2239316 -35.3945171
H -0.5126998 -7.5591821 -34.6545479
H -1.9003645 -8.5918536 -34.9852958
C -0.0205586 -9.403772 -35.5957312
C -0.476939 -10.7237244 -35.4080804
H -1.509788 -10.9114615 -35.1526382
C 0.4018223 -11.8087914 -35.5828742
H 0.0411055 -12.8215519 -35.4605813
C 1.7396798 -11.5814567 -35.9510346
H 2.4073452 -12.4177367 -36.1126058
C 2.197147 -10.2669613 -36.1448257
H 3.2167878 -10.0866761 -36.4561706
C 1.3158525 -9.1847377 -35.975059
H 1.6594669 -8.1812815 -36.1674815
N -2.9508894 -8.7602365 -37.8162996
H -3.5180262 -8.6013591 -36.9897401
C -3.5164419 -9.5705426 -38.9055008
H -2.8830213 -10.4489155 -39.0412821
C -3.4884375 -8.8191223 -40.2441416
O -3.0854456 -9.4032043 -41.2444796
C -4.9353343 -10.0606912 -38.5229828
H -5.4930771 -9.2156787 -38.1147458
C -5.7199256 -10.6008875 -39.7362459
H -5.1748012 -11.428586 -40.1934969
H -5.8704642 -9.8203191 -40.4808494
H -6.7059161 -10.950001 -39.4339102
C -4.8187313 -11.1576382 -37.4374046
H -4.3660146 -12.0503218 -37.8713733
H -4.1598068 -10.814504 -36.6407699
C -6.1553173 -11.5460725 -36.7923586
H -6.7955629 -12.0580817 -37.5094256
H -6.6616767 -10.6550823 -36.4198857
H -5.9674939 -12.2254155 -35.9610021
N -3.8323925 -7.5248185 -40.2879394
H -4.0656676 -7.0556824 -39.4218744
C -3.7691065 -6.745843 -41.5391746
H -4.3005425 -7.2994218 -42.3139513
C -2.3393446 -6.5519942 -42.0526116
O -2.1145754 -6.6247537 -43.2621331
C -4.4332741 -5.3718712 -41.3703427
H -3.8537605 -4.6109815 -41.8970195
H -4.4733346 -5.1006908 -40.3138927

O -5.7412933 -5.3987764 -41.9212897
H -5.6390313 -5.3843889 -42.8827614
N -1.3861787 -6.3126124 -41.1495095
H -1.6547347 -6.2826085 -40.1714205
C 0.0158965 -6.0425797 -41.481886
H 0.0577829 -5.3112942 -42.2894989
C 0.740751 -7.3012983 -41.9896076
O 1.4825599 -7.2298511 -42.9719905
C 0.6993255 -5.4537121 -40.2332656
H 0.5759614 -6.161702 -39.4119538
H 1.768021 -5.3583632 -40.4270006
C 0.1460188 -4.0790425 -39.7977291
H -0.9410754 -4.0968931 -39.8068985
C 0.5881962 -3.7787531 -38.3651539
H 1.6688825 -3.8436922 -38.2817429
H 0.1358835 -4.4987693 -37.6855946
H 0.2728934 -2.7773283 -38.0815491
C 0.5838749 -2.9467471 -40.7300837
H 1.6679734 -2.8669288 -40.751094
H 0.1714852 -2.0035095 -40.3724266
H 0.2150466 -3.1247907 -41.7383577
N 0.4550266 -8.4611916 -41.3890336
H -0.1379686 -8.4416052 -40.564342
C 0.9646168 -9.7595951 -41.8364496
H 2.0196674 -9.6433663 -42.0892642
C 0.2485841 -10.2596498 -43.1035624
O 0.8987943 -10.7616169 -44.0158858
C 0.8455888 -10.7511466 -40.6638834
H 1.3523783 -10.326376 -39.7955445
H -0.210257 -10.8640917 -40.4110157
C 1.4357562 -12.1477194 -40.9424942
H 0.9147976 -12.6038002 -41.7839374
C 2.9345315 -12.1074372 -41.2461209
H 3.4682867 -11.5756565 -40.4587843
H 3.1048157 -11.6059846 -42.1990644
H 3.3258448 -13.1209115 -41.3293987
C 1.228386 -13.036948 -39.7157139
H 1.7534838 -12.6171902 -38.8578697
H 1.6078131 -14.0384338 -39.9180822
H 0.1647686 -13.1057515 -39.4881301
N -1.0698244 -10.0583188 -43.2209953
H -1.5784126 -9.6705319 -42.4337335
C -1.8288138 -10.4543792 -44.4133578
H -1.6406428 -11.512455 -44.6026739
C -1.3632127 -9.6903915 -45.6534104
O -1.0336879 -10.3147171 -46.6612399

C -3.3368109 -10.2801862 -44.1613091
H -3.5384081 -9.2595103 -43.8397539
H -3.6148275 -10.9493552 -43.3461491
C -4.2354917 -10.6192243 -45.3629775
H -5.2387681 -10.8120393 -44.981554
H -3.8764807 -11.5469091 -45.8110439
S -4.390968 -9.3657168 -46.6781462
C -5.1799099 -8.0079527 -45.7729399
H -5.4383667 -7.2116522 -46.4701612
H -4.4931662 -7.6138861 -45.0242914
H -6.0870682 -8.3705426 -45.2881865
N -1.2836702 -8.3547994 -45.597801
H -1.5351508 -7.8696527 -44.7427603
C -0.9788531 -7.5879528 -46.8043397
H -1.457325 -8.1034868 -47.6360249
C 0.521404 -7.5756896 -47.1451478
O 0.8637811 -7.5102657 -48.3264741
C -1.6430715 -6.2032923 -46.748782
H -1.1355764 -5.5871202 -46.0089988
H -2.6802938 -6.3144597 -46.4385527
C -1.6394516 -5.5151375 -48.1094585
N -0.9709235 -4.389742 -48.2157069
H -0.4685428 -4.0199304 -47.4270163
H -0.9501478 -3.9346266 -49.1103779
O -2.2486604 -5.9590541 -49.0785727
N 1.416904 -7.7516498 -46.1678263
H 1.0980991 -7.8154289 -45.2078644
C 2.8375119 -7.9866019 -46.4363406
H 3.1969886 -7.2710589 -47.1758242
H 3.3911814 -7.8211393 -45.5225839
C 3.1279278 -9.4043741 -46.945777
O 3.8298141 -9.5493483 -47.9453507
N 2.447086 -10.4225332 -46.4101552
H 1.9315207 -10.2624063 -45.5490469
C 2.4854058 -11.7979805 -46.9139969
H 3.5152832 -12.1562071 -46.9087821
H 1.8931677 -12.4278665 -46.2504679
C 1.9239234 -11.9295401 -48.3328251
O 2.5932635 -12.4826701 -49.2022689
N 0.7929469 -11.2777152 -48.628854
H 0.267872 -10.8632915 -47.8641561
C 0.2683398 -11.1228987 -49.9938683
H 0.0573556 -12.1067264 -50.4128921
C 1.2826057 -10.4351154 -50.919821
O 1.4660857 -10.8427349 -52.0678355
C -1.03995 -10.3126457 -49.9511539

H -1.8060414 -10.9034882 -49.4476874
H -0.8743164 -9.4136616 -49.3588149
C -1.5543657 -9.8836418 -51.2950987
C -1.1187229 -8.7991881 -51.9758444
H -0.3563124 -8.1147227 -51.6133052
N -1.6791194 -8.7882528 -53.2363242
H -1.3736528 -8.1625098 -53.9676252
C -2.5438482 -9.8457334 -53.4186151
C -3.3411695 -10.2526051 -54.4992947
H -3.3476281 -9.6931163 -55.422582
C -4.1539985 -11.3894695 -54.358604
H -4.801853 -11.6994674 -55.1695271
C -4.1314936 -12.1172815 -53.1550939
H -4.7589197 -12.9925633 -53.044334
C -3.3101688 -11.7105559 -52.0842815
H -3.3046579 -12.2804389 -51.1652787
C -2.499787 -10.5592805 -52.1831049
N 1.96428 -9.3993704 -50.4253499
H 1.7809878 -9.1156849 -49.4709382
C 2.9636726 -8.654149 -51.2011413
H 2.5137098 -8.3680451 -52.1510126
C 4.1751778 -9.5252683 -51.5380323
O 4.584636 -9.5468988 -52.6991712
C 3.3562456 -7.3613529 -50.4714756
H 3.7076416 -7.5770001 -49.4645344
C 4.4220107 -6.5580398 -51.2039862
H 4.0784879 -6.3762013 -52.2179027
H 5.3645424 -7.1059384 -51.2128172
H 4.574357 -5.6062331 -50.6915057
O 2.2203371 -6.5288334 -50.4111686
H 1.6945377 -6.8265429 -49.6442254
N 4.6517677 -10.3483214 -50.5991615
H 4.2697926 -10.2894075 -49.6593257
C 5.6772822 -11.3580867 -50.8568759
H 6.5451977 -10.8700353 -51.3041686
C 5.1923812 -12.4341222 -51.8490049
O 5.8968642 -12.7354913 -52.8110289
C 6.1067911 -11.9647842 -49.5145566
H 5.2580115 -12.4435631 -49.0255728
H 6.499734 -11.1841373 -48.8617151
H 6.8863492 -12.7084471 -49.6781218
N 3.9534366 -12.9249302 -51.7086818
H 3.4267844 -12.6496469 -50.8836684
C 3.334672 -13.8997941 -52.6291921
H 3.9737587 -14.7791788 -52.6745089
C 3.2311635 -13.3581777 -54.0601941

O 3.501972 -14.0879971 -55.0132972
C 1.9611062 -14.3607064 -52.0920771
H 1.4005589 -13.4924383 -51.7508018
C 1.0987965 -15.094003 -53.1308964
H 1.6582642 -15.9270397 -53.5595551
H 0.8015677 -14.4086898 -53.9226587
H 0.1946828 -15.4764729 -52.6554527
C 2.1541222 -15.3188572 -50.9094302
H 2.6359165 -16.2380737 -51.2437664
H 1.18814 -15.559973 -50.4658729
H 2.7765934 -14.8574841 -50.1444956
N 2.9105554 -12.073782 -54.2549198
H 2.6628376 -11.510579 -53.4444772
C 2.8382215 -11.4848562 -55.603466
H 2.5477263 -12.2731829 -56.294507
C 4.1932573 -11.0523229 -56.1735234
O 4.3050569 -11.0014922 -57.3976465
C 1.7500749 -10.3972881 -55.6684607
H 1.973677 -9.6224873 -54.932472
H 1.7766916 -9.9369756 -56.657886
C 0.3181127 -10.9321047 -55.4242165
H 0.2277581 -11.265099 -54.390261
H -0.3748739 -10.1024204 -55.5677658
C -0.0987512 -12.0894509 -56.3560436
H 0.544746 -12.9544503 -56.17014
H 0.0153415 -11.7777132 -57.39766
C -1.5577073 -12.5187353 -56.1029819
H -2.2269318 -11.6802266 -56.3325773
H -1.6736438 -12.7561101 -55.0366384
N -1.920309 -13.7026939 -56.930816
H -1.8494992 -13.5031153 -57.9228767
H -2.903941 -13.9547134 -56.8042702
H -1.3493329 -14.5134468 -56.71273
N 5.239108 -10.8811623 -55.358782
H 5.0832237 -10.9063656 -54.3572045
C 6.6252368 -10.7490509 -55.8568326
H 6.6136699 -10.1725473 -56.7835268
C 7.23075 -12.111499 -56.2120809
O 7.8163342 -12.237301 -57.2830009
C 7.5248285 -9.9796415 -54.8686851
H 7.4639628 -10.4501565 -53.8877875
C 8.9905939 -9.944826 -55.3227672
H 9.0568977 -9.5997377 -56.3555774
H 9.4372889 -10.9357012 -55.2428839
H 9.5579733 -9.2635562 -54.6896495
C 7.0793244 -8.5186274 -54.7449023

H 7.2214826 -8.0117064 -55.6995769
H 7.6884329 -8.0243809 -53.9867521
H 6.0354079 -8.4688468 -54.4529373
N 7.0278598 -13.1595392 -55.4024952
H 6.5409429 -13.011564 -54.5230944
C 7.6271667 -14.4858677 -55.6559459
H 8.6634613 -14.3031979 -55.9438593
C 6.9906641 -15.2279986 -56.8419336
O 7.6716931 -15.9879257 -57.5225639
C 7.6575082 -15.3317962 -54.3638388
H 7.9124323 -14.6589608 -53.5431137
C 6.2996944 -15.9707711 -54.0400428
H 6.0816508 -16.7851927 -54.7321286
H 5.5262014 -15.2171777 -54.1221082
H 6.2972443 -16.3543042 -53.0201916
C 8.777694 -16.3911983 -54.4438692
H 8.5331507 -17.1303691 -55.2074872
H 9.7068405 -15.8989223 -54.7355244
C 9.0351373 -17.1189259 -53.1182629
H 8.173149 -17.7267722 -52.8436591
H 9.2387219 -16.3944539 -52.3289923
H 9.8999306 -17.7735446 -53.2299662
N 5.7387061 -14.9039989 -57.1868191
H 5.2127234 -14.3222172 -56.5485866
C 5.1062204 -15.3009733 -58.4534771
H 5.3874356 -16.3318692 -58.6680816
C 5.6030433 -14.4874707 -59.672003
O 5.0849158 -14.6599443 -60.773668
C 3.5735904 -15.2270815 -58.2905951
H 3.3168007 -14.2373792 -57.910768
H 3.1057086 -15.3358337 -59.269651
C 2.9775319 -16.3018904 -57.3585176
H 3.4899355 -16.2918303 -56.3990126
C 1.4957103 -15.9962792 -57.1210999
H 0.9610944 -15.9934025 -58.0710052
H 1.4152946 -15.0272253 -56.6348142
H 1.0717324 -16.755584 -56.4636815
C 3.0815974 -17.7101515 -57.947436
H 2.616133 -17.7444842 -58.9324024
H 2.5847047 -18.4241014 -57.2906702
H 4.1275589 -18.0041337 -58.0324987
N 6.5584693 -13.5695965 -59.4840437
H 6.9635998 -13.4805897 -58.559394
C 7.1419561 -12.7294521 -60.533019
H 8.0300305 -12.2428227 -60.1296168
H 7.4428545 -13.3604889 -61.3694312

C 6.2113295 -11.6372743 -61.0664694
O 6.4668133 -11.1007314 -62.1414297
N 5.0989407 -11.3521821 -60.3807085
H 4.9813281 -11.7997437 -59.4822739
C 4.011173 -10.5016823 -60.8819061
H 4.0372055 -10.488733 -61.9692609
C 4.1458654 -9.0360507 -60.4550618
O 4.0311999 -8.1487806 -61.2999311
C 2.6640571 -11.1114873 -60.4638078
H 1.8926477 -10.351034 -60.4842377
H 2.730934 -11.4831747 -59.4414438
C 2.2535011 -12.2657448 -61.3868724
H 1.4591474 -12.8339579 -60.902344
H 3.1069454 -12.932034 -61.5231857
S 1.6498686 -11.7327541 -63.0139356
C 1.6214996 -13.3280329 -63.8756168
H 1.2174771 -13.1904539 -64.8799789
H 0.9970689 -14.0358844 -63.3303468
H 2.6341481 -13.7267153 -63.9450518
N 4.4403578 -8.7747006 -59.1755103
H 4.5741277 -9.5745046 -58.565276
C 4.7721047 -7.4432695 -58.6461105
H 4.2329189 -6.6929209 -59.2287529
C 6.2628089 -7.1513591 -58.8255713
O 7.1132365 -8.0286716 -58.7190631
C 4.3124856 -7.3225627 -57.1776623
H 3.2218766 -7.3201566 -57.1538076
H 4.664482 -8.1940433 -56.6248092
C 4.8413101 -6.0586761 -56.4669405
H 5.9269408 -6.1201353 -56.3940316
H 4.5835999 -5.1738691 -57.0510368
C 4.2767258 -5.8952971 -55.0518431
H 4.2724669 -6.8599056 -54.5417714
H 3.2469937 -5.5465683 -55.1283597
C 5.0870934 -4.8828455 -54.2236432
H 4.4787623 -4.580427 -53.3658758
H 5.2958632 -3.991815 -54.8235343
N 6.3573003 -5.466491 -53.7188249
H 6.2395348 -6.3113772 -53.1868018
H 6.9070721 -4.8226841 -53.1439003
H 7.0461657 -5.6510471 -54.4521823
N 6.5831059 -5.8796803 -59.0166479
H 5.8235267 -5.2105063 -59.113986
C 7.9375657 -5.3227778 -59.0310328
H 8.6143732 -5.9856498 -58.4939189
C 7.9299697 -3.9753104 -58.3016906

O 6.8849011 -3.3299543 -58.2056804
C 8.3940536 -5.2114097 -60.498702
H 7.5989185 -4.7461053 -61.083704
C 9.6867695 -4.4393521 -60.7534294
H 10.461601 -4.7634189 -60.0624864
H 9.5092652 -3.3712309 -60.637248
H 10.0261984 -4.6212524 -61.7728531
O 8.6164189 -6.5095718 -60.9963761
H 8.1255677 -7.1366062 -60.4489059
N 9.0868513 -3.5097747 -57.8288001
H 9.9133854 -4.0836863 -57.9019445
C 9.2842826 -2.0970461 -57.484942
H 8.3344924 -1.5820776 -57.6283965
C 10.2397767 -1.4588863 -58.488381
O 11.1822076 -2.0878065 -58.9748673
C 9.6723067 -1.8988096 -55.9998239
H 9.4745394 -2.8358148 -55.4851521
C 11.1589555 -1.5434698 -55.8111287
H 11.3819425 -0.5661663 -56.2441685
H 11.7740098 -2.3086833 -56.2769605
H 11.4226248 -1.4985324 -54.7557414
C 8.7636256 -0.8305709 -55.3496209
H 8.9602225 0.1445593 -55.7991688
H 7.7226656 -1.0931424 -55.5430241
C 8.9349449 -0.7232145 -53.8274753
H 9.9006709 -0.2819557 -53.5826193
H 8.8562241 -1.7099889 -53.3708842
H 8.1532925 -0.0869728 -53.4164581
N 9.9517901 -0.2103244 -58.8094482
H 9.1682783 0.231535 -58.3373574
C 10.654951 0.6360141 -59.7608788
H 11.4393476 0.0802286 -60.2761536
C 11.3137825 1.7707113 -58.9834636
O 10.6744127 2.3280577 -58.0948429
C 9.5994166 1.0937026 -60.7929687
H 8.6089785 1.028997 -60.3467226
H 9.6158493 0.3827572 -61.6188321
C 9.6756245 2.4856417 -61.3678588
N 9.6560584 2.7683857 -62.7304314
C 9.6304442 4.1041051 -62.8320208
H 9.6677495 4.647484 -63.7669766
N 9.599696 4.6686446 -61.6145544
H 9.6754936 5.6576488 -61.4159251
C 9.6037052 3.6646771 -60.6808175
H 9.5692624 3.7963986 -59.6106405
N 12.5267095 2.1509732 -59.3842939

H 13.0058659 1.5797279 -60.0714761
C 13.1036285 3.4786336 -59.1419166
H 12.3791542 4.0801703 -58.6065819
C 13.4003518 4.137936 -60.4980748
O 13.9804095 3.4895145 -61.3751488
C 14.3486196 3.3947989 -58.2404563
H 14.0355429 3.0345998 -57.2602812
H 15.0443904 2.6671256 -58.661884
C 15.0998792 4.7290328 -58.0561606
H 15.5972644 4.9762276 -58.9978091
H 15.8861092 4.5829474 -57.3125827
C 14.2028056 5.9034645 -57.6227446
O 14.1451209 6.8963968 -58.3833069
O 13.5576779 5.8054743 -56.5593021
N 13.0453446 5.4126846 -60.6712948
H 12.6425479 5.9066621 -59.8759317
C 13.3632583 6.2062371 -61.8533931
H 14.1139036 5.6662568 -62.4167197
C 13.9496037 7.5757594 -61.5001384
O 13.3110616 8.3851175 -60.8299668
C 12.1357501 6.3519771 -62.752082
H 11.3121412 6.7960327 -62.1946084
H 11.8710934 5.3727228 -63.1411174
H 12.3807769 6.9994993 -63.5956822
N 15.109189 7.8745898 -62.0873356
H 15.4773862 7.2053572 -62.7563538
C 15.8432781 9.1316037 -61.9126432
H 15.2448618 9.8123746 -61.30726
C 16.0561624 9.7692783 -63.2824116
O 16.4188304 9.0839867 -64.2349318
C 17.1881392 8.8969502 -61.188147
H 17.8076121 8.2561194 -61.8169794
C 16.9764192 8.1765845 -59.8366834
H 16.3622953 8.7964619 -59.1838709
H 16.4432474 7.2414061 -60.0042166
C 17.9261826 10.2389898 -60.9959842
H 17.3004281 10.929227 -60.4298202
H 18.1645092 10.6855659 -61.9607414
H 18.8686102 10.0894265 -60.4738383
C 18.2749524 7.8099776 -59.110778
H 18.7708445 8.7035771 -58.7336403
H 18.9412565 7.2714474 -59.7855677
H 18.0370505 7.1684199 -58.2616609
N 15.8801458 11.0823937 -63.3805037
H 15.5331748 11.5872138 -62.5688742
C 15.9623305 11.817571 -64.6442913

H 16.7952455 11.4316431 -65.2351664
C 16.2306405 13.3158319 -64.4223685
O 16.001996 13.8407697 -63.3256233
C 14.6780471 11.5721577 -65.4405948
H 14.5614297 12.3592306 -66.1828294
H 14.8143697 10.6397859 -65.9771394
C 13.3981278 11.4233831 -64.5967226
H 13.2409175 12.3213791 -64.0054631
H 13.4843556 10.5777558 -63.912386
C 12.2010246 11.1445692 -65.486179
O 12.4220538 10.9836834 -66.7062945
O 11.0711677 11.0577901 -64.952999
N 16.7127389 14.0282412 -65.4533237
H 16.8577185 13.566903 -66.349382
C 16.9273486 15.4814323 -65.3898288
H 16.3871383 15.8476574 -64.5193038
C 16.3447654 16.2645858 -66.5817988
O 16.1558116 15.7419953 -67.6835179
C 18.4109936 15.8096423 -65.1287159
H 18.8261429 15.0930979 -64.4212154
H 18.4804463 16.7986531 -64.677849
C 19.2719235 15.817902 -66.3805842
O 19.9812442 14.8726451 -66.670495
N 19.2629312 16.8803734 -67.1552989
H 19.7115882 16.7659892 -68.0471681
H 18.6938739 17.6945154 -66.9492704
N 16.1171737 17.5560555 -66.3461034
H 16.3244087 17.9218972 -65.4216912
C 15.6409335 18.5213459 -67.3311437
H 15.1563779 17.9880668 -68.1515813
C 16.8207639 19.3019001 -67.9050657
O 17.814395 19.5196466 -67.2156087
C 14.6025547 19.470826 -66.7156745
H 15.0457816 19.9951369 -65.8663646
H 13.7532703 18.8943344 -66.3687152
C 14.1049765 20.4776596 -67.7549781
O 14.0497 21.6808959 -67.4357038
O 13.8939207 20.0301459 -68.9036003
N 16.6932574 19.7127438 -69.1587056
H 15.7361606 19.6535177 -69.5221682
C 17.6417525 20.4891035 -69.9664808
H 18.1709276 19.7718774 -70.5775385
C 18.7397635 21.2279062 -69.1911158
O 19.6163392 20.5303848 -68.6231238
O 18.7905354 22.4280729 -69.0989548
C 16.8603575 21.4153103 -70.9236658

H 16.3055296 22.1240006 -70.3041713
H 17.5800612 21.9809911 -71.516076
C 15.8643249 20.7213124 -71.8783228
H 15.1037594 20.1933291 -71.2956912
C 15.1458514 21.7861056 -72.7164382
H 15.8579213 22.3362843 -73.3312233
H 14.6219113 22.4788679 -72.0561909
H 14.4058411 21.3071409 -73.3609419
C 16.5435476 19.7199673 -72.8232848
H 17.329409 20.2094041 -73.398468
H 15.8002973 19.3095617 -73.5106352
H 16.9602655 18.8885803 -72.2557724

**Supplementary Data 7. Cartesian coordinates for perivine docked into TePeNMT+SAM.**

N -2.1385667 21.7753992 -15.9844882
H -2.8881317 22.3451992 -15.612073
H -2.4187043 21.4412796 -16.898962
H -1.952445 20.9940613 -15.3721405
C -0.8983927 22.6008534 -16.1048954
H -1.0805973 23.3575681 -16.8709468
C 0.33849 21.7897622 -16.5659561
O 1.2553163 22.327476 -17.1874333
C -0.6053818 23.3116885 -14.7771474
H 0.3668547 23.80077 -14.8479421
H -0.5567763 22.5711725 -13.9778747
C -1.639947 24.3835703 -14.4160168
H -1.396285 24.7758699 -13.426708
H -2.6273748 23.9290305 -14.3587376
S -1.7285502 25.7740286 -15.5733039
C -2.9534829 26.7844932 -14.6928753
H -3.1342375 27.7045542 -15.2488035
H -2.5798462 27.0332981 -13.6984196
H -3.8891334 26.2323095 -14.5953778
N 0.3673335 20.4886197 -16.2657559
H -0.4109708 20.1080689 -15.7405706
C 1.3938537 19.5859753 -16.7771625
H 1.3580658 19.6103407 -17.8672226
H 2.3753087 19.9422459 -16.4680576
C 1.2496557 18.1385969 -16.3224611
O 0.7986647 17.8537127 -15.2154758
N 1.7628938 17.2390447 -17.1522057
H 2.1121366 17.5432004 -18.0501647
C 2.0288584 15.8382482 -16.8338954
H 1.9028599 15.7027836 -15.7604402
C 3.4957816 15.5076051 -17.1394012
O 4.119702 16.1812749 -17.9609391
C 0.9990926 14.9223526 -17.5032517
H 1.2572628 13.8928438 -17.2749979
H 0.0423459 15.1358366 -17.026009
C 0.8149397 15.0455348 -19.020348
H 1.6157817 14.5050359 -19.5288645
H 0.8699863 16.0954136 -19.3176114
C -0.5554566 14.4754363 -19.4180847
O -1.297864 15.2053148 -20.1119062
O -0.875195 13.3478418 -18.9747065
N 4.1061184 14.7016468 -16.2632362
H 3.5442127 14.1102083 -15.6715657
C 5.5629684 14.6122266 -16.0983634

H 6.0446756 14.8186955 -17.0600251
C 5.9695982 13.2129132 -15.6736854
O 5.3463653 12.639753 -14.786269
C 6.0594875 15.5900786 -15.0182406
H 5.5966651 15.3285324 -14.0641537
H 7.1389711 15.4636292 -14.915048
C 5.7688522 17.0627619 -15.3210483
H 6.199365 17.3109927 -16.2919241
H 4.6914859 17.2284646 -15.351451
C 6.3694499 17.9615862 -14.2310701
H 7.4516987 17.8220036 -14.192695
H 5.9514537 17.6743696 -13.2638997
C 6.0413128 19.4366002 -14.4861744
H 6.4275873 20.029751 -13.6516003
H 4.9522862 19.5499365 -14.5026758
N 6.6253332 19.9107507 -15.7656037
H 7.6331593 19.818804 -15.7546612
H 6.3843249 20.8792368 -15.933991
H 6.2683797 19.3550584 -16.5356419
N 7.106455 12.7593095 -16.1705568
H 7.4956956 13.2759729 -16.9574346
C 7.5639831 11.383838 -16.027934
H 6.7220861 10.7521143 -15.7410336
C 8.6256294 11.3080201 -14.9245199
O 9.6775993 11.9537076 -14.9983873
C 8.0743326 10.8706348 -17.3888252
H 8.9570019 11.4416081 -17.6791626
H 8.3787829 9.8302572 -17.2664366
C 7.0169986 10.9370419 -18.5172246
H 6.1405201 10.371064 -18.1914962
H 7.409044 10.4393792 -19.4038394
C 6.5974706 12.3719046 -18.896483
O 7.4785217 13.2681533 -18.9084406
O 5.3745041 12.6072441 -18.9936412
N 8.3366279 10.5700175 -13.8495991
H 7.428984 10.1185224 -13.7997984
C 9.2537472 10.4628666 -12.7165948
H 9.411715 11.4704627 -12.3299929
C 10.6243327 9.9036971 -13.153139
O 10.6986872 8.9385328 -13.9069108
C 8.6037136 9.6350831 -11.6036772
H 8.4204313 8.619799 -11.9600852
H 7.655759 10.0896643 -11.3114457
H 9.2647124 9.595986 -10.737355
N 11.703524 10.5367378 -12.6721267
H 11.5296608 11.3030954 -12.0418231

C 13.1142625 10.3142897 -13.0702562
H 13.6516325 11.146085 -12.6173586
C 13.4243279 10.5119185 -14.5675031
O 14.4681525 10.0599575 -15.0339048
C 13.7765064 9.0619071 -12.4395625
H 14.845682 9.122573 -12.6445486
C 13.6265139 9.0675566 -10.9118445
H 12.5803642 8.9423089 -10.6318339
H 14.0018322 10.0070241 -10.506304
H 14.2061243 8.2486758 -10.485005
C 13.2962742 7.7038875 -12.9652798
H 12.2890587 7.4911381 -12.6080123
H 13.9637118 6.9192992 -12.6112727
H 13.2950871 7.7094786 -14.0550769
N 12.605294 11.3349009 -15.2378421
H 11.7369915 11.5814565 -14.7805248
C 12.7314974 11.833814 -16.6111805
H 11.8724211 12.4921825 -16.7495073
C 12.6154246 10.7757957 -17.7237248
O 13.1625965 9.6728211 -17.644985
C 13.9716254 12.728998 -16.7542726
H 14.876663 12.1276992 -16.6566322
H 13.9650252 13.5017634 -15.9855672
H 13.9695782 13.2008787 -17.7373648
N 11.9102878 11.1457732 -18.7984009
H 11.4901366 12.0606529 -18.8311174
C 11.6113386 10.2161504 -19.8849453
H 11.4683386 9.2575284 -19.3922061
C 12.7697246 10.041093 -20.8749029
O 13.4841971 10.9883786 -21.2103005
C 10.2956217 10.5834444 -20.6018401
H 9.5642602 10.8737195 -19.8541547
H 10.4613373 11.4376453 -21.2598264
C 9.7172444 9.4047201 -21.4196476
H 10.350257 9.2350742 -22.2929309
H 8.7362979 9.7002982 -21.7986705
C 9.5899082 8.089336 -20.6183328
O 8.4815808 7.7830257 -20.1380566
O 10.6205886 7.3866881 -20.4498584
N 12.8901662 8.8305339 -21.4204041
H 12.2017952 8.1247666 -21.1470783
C 13.7998552 8.4972353 -22.5092701
H 14.1771403 9.4236569 -22.9434863
C 13.0399482 7.741936 -23.6088019
O 12.2666108 6.8259826 -23.3460136
C 14.9936922 7.7225667 -21.9245386

H 15.5173435 8.3745069 -21.2235556
H 14.6001376 6.8863099 -21.3570609
C 16.0029798 7.189281 -22.9591753
H 15.4878043 6.568367 -23.6890142
C 16.7306364 8.3210255 -23.6896215
H 17.2504768 8.9574053 -22.9726436
H 16.0235264 8.9250527 -24.2541046
H 17.4549132 7.90003 -24.3877077
C 17.0509741 6.3236455 -22.2584349
H 17.5894864 6.9110609 -21.5149428
H 17.7563171 5.9395483 -22.9960493
H 16.5630007 5.4766885 -21.778768
N 13.2962296 8.1160457 -24.8611726
H 13.9627613 8.8548464 -25.0049569
C 12.7297022 7.4720004 -26.0467334
H 11.6552424 7.6561114 -26.0668372
C 12.963355 5.9483119 -26.0268958
O 14.0825531 5.4964434 -25.7932219
C 13.3669937 8.1475173 -27.2671617
H 13.1403425 9.2139322 -27.2266732
H 14.4513269 8.049724 -27.187207
C 12.9656811 7.6189319 -28.6307211
C 12.1494424 8.3971049 -29.4761888
H 11.769383 9.3526089 -29.1454154
C 11.8565891 7.9520163 -30.7798108
H 11.2562024 8.555452 -31.4472005
C 12.3673248 6.7178848 -31.2374061
O 12.1160781 6.2857259 -32.4969706
H 11.5991475 6.9488647 -33.0185918
C 13.1697677 5.9340538 -30.3847744
H 13.5492799 4.9896701 -30.7459872
C 13.4827043 6.3907892 -29.091865
H 14.1273567 5.791667 -28.4621886
N 11.8951892 5.1788313 -26.2599184
H 11.0304658 5.6427895 -26.4800882
C 11.8426092 3.7036502 -26.2137578
H 10.7811114 3.4599108 -26.2631227
C 12.3180866 3.0493723 -24.893297
O 12.588105 1.8486882 -24.8229072
C 12.4769853 3.1028593 -27.4818833
H 13.5610504 3.0598399 -27.3602348
H 12.2582179 3.7523311 -28.3311342
C 11.9188234 1.7098089 -27.8129528
O 10.6734795 1.5608478 -27.7641107
O 12.7303627 0.8270589 -28.169355
N 12.3786583 3.8077385 -23.7892188

H 12.1567416 4.7960764 -23.8565606
C 12.7236866 3.2518266 -22.4780661
H 13.5786598 2.5902866 -22.6181944
C 11.5733328 2.4149249 -21.9209309
O 10.5345585 2.9233827 -21.5059758
C 13.1586237 4.3717878 -21.5322934
H 14.0067916 4.8753226 -21.9927235
H 12.3398627 5.0838206 -21.4074609
C 13.5708913 3.8411374 -20.1519667
H 12.6903613 3.4453154 -19.6452956
H 14.3110593 3.0473288 -20.2626016
C 14.1595742 4.9730826 -19.3074994
H 13.477531 5.8273082 -19.3386531
H 15.1255757 5.265841 -19.7198963
C 14.3246592 4.5251755 -17.8558798
H 14.9647772 3.6393135 -17.8237746
H 13.3343658 4.2459127 -17.4769636
N 14.894404 5.6206571 -17.034387
H 15.8026652 5.9022239 -17.3743342
H 14.9585333 5.3479973 -16.0632593
H 14.281514 6.4323719 -17.0835563
N 11.8145313 1.1129027 -21.796752
H 12.636759 0.7428888 -22.2469702
C 10.915173 0.206206 -21.075752
H 9.9261211 0.3116122 -21.5233934
C 10.7879049 0.6022152 -19.5986625
O 11.7565775 1.0128309 -18.9601126
C 11.3345322 -1.270057 -21.22834
H 10.6296722 -1.8783407 -20.6606041
C 11.2418011 -1.7098514 -22.6941182
H 11.9608367 -1.1650701 -23.3087444
H 10.2395993 -1.5174528 -23.0780593
H 11.4468521 -2.7773516 -22.7753339
C 12.7458793 -1.5768076 -20.7095885
H 13.4907476 -1.0077613 -21.2656222
H 12.9570051 -2.6395233 -20.8284199
H 12.8214824 -1.3296274 -19.6506744
N 9.6090573 0.3641778 -19.0190877
H 8.8636369 0.042936 -19.61778
C 9.3030641 0.4995601 -17.5765515
H 8.2498576 0.2378134 -17.4888618
C 9.3464409 1.9102799 -16.9589934
O 9.2888891 2.018324 -15.7386553
C 10.0341763 -0.545724 -16.7016046
H 9.6693134 -0.4676256 -15.6770594
C 9.7794235 -1.9797159 -17.169852

H 10.215899 -2.1450359 -18.1540049
H 8.7079161 -2.1740235 -17.2032576
H 10.239578 -2.6699839 -16.4624216
O 11.4299275 -0.3613229 -16.691399
H 11.6667592 0.1678767 -17.4727227
N 9.2779939 2.9909397 -17.7450031
H 9.3809 2.8980378 -18.7455964
C 8.8924366 4.3186524 -17.2226461
H 9.484174 4.5459203 -16.3361664
C 7.3950813 4.3363773 -16.8420819
O 6.54833 4.1563548 -17.7157283
C 9.1737252 5.4060425 -18.2724269
H 8.8412563 6.3707971 -17.8859208
H 8.6096819 5.1895781 -19.1816057
O 10.5517975 5.4957348 -18.5958262
H 10.5894134 6.1754387 -19.3178923
N 7.0367162 4.4791013 -15.5515566
H 7.7637049 4.4744686 -14.8485576
C 5.6335974 4.372326 -15.0850094
H 5.0070584 4.6318068 -15.9400818
C 5.0839267 5.3356751 -13.9496445
O 4.1802507 4.8921545 -13.2441249
C 5.3501324 2.8742188 -14.7954481
H 5.6601937 2.2734319 -15.6495461
H 4.2770818 2.7385852 -14.6680957
C 6.0481724 2.3356799 -13.5563096
O 6.9355157 2.9397106 -12.9802825
N 5.6839717 1.1536912 -13.1130988
H 6.1721809 0.8278604 -12.2984982
H 4.9434468 0.6468731 -13.5561149
N 5.4209234 6.6581869 -13.7954432
H 6.1313111 7.0377768 -14.4037847
C 4.6727308 7.6690857 -12.8730958
H 3.6722803 7.2716519 -12.7031805
H 5.1665915 7.6425688 -11.902207
C 4.4605241 9.2667223 -13.2399755
O 5.4811072 9.6663177 -13.812121
N 3.2536409 10.1109433 -12.9872607
H 2.6474233 9.6614132 -12.3206553
C 2.3771526 11.4343133 -13.6922496
H 3.1658733 12.0715482 -14.0792353
C 1.0505375 12.8198719 -12.978889
O 0.738086 12.1140384 -12.0071596
C 1.9540762 10.4352165 -14.9544058
H 0.9329988 10.1097298 -14.746017
C 2.7417697 9.1177858 -15.5124699

H 3.7821969 9.3932306 -15.6797871
H 2.6711718 8.3582525 -14.7352443
C 1.9181844 11.302737 -16.1856385
H 2.9290002 11.590086 -16.4899791
H 1.3394983 12.1952761 -15.9637221
H 1.4171187 10.8031861 -17.0150758
C 2.3461044 8.3062146 -16.7841532
H 1.2838958 8.043668 -16.7411571
H 2.9411495 7.3919643 -16.8207085
H 2.5328117 8.8854332 -17.695951
N 0.2333932 14.4003043 -13.1834479
H 0.7437184 14.7653437 -13.9704046
C -1.2123799 15.955379 -12.9597387
H -1.0028175 16.3292374 -13.960482
C -2.5312399 15.1695335 -13.1010222
H -2.3258053 14.1092915 -13.2484796
H -3.0618878 15.2689053 -12.1555772
C -3.4175086 15.6977339 -14.2585728
H -3.2432872 16.7657087 -14.3985759
C -3.0783778 14.9714564 -15.5675978
H -3.2811317 13.903087 -15.4809576
H -2.0269104 15.1037593 -15.8111044
H -3.6757972 15.3801559 -16.3812018
C -4.9110054 15.512951 -13.9555661
H -5.1400087 14.4586235 -13.8201664
H -5.5040244 15.9052164 -14.7807417
H -5.172629 16.0607214 -13.0540348
C -2.0596333 17.9254177 -12.3430272
O -1.3832877 18.2475852 -11.374691
N -3.271554 19.2249776 -12.5340521
H -3.8017849 18.9469697 -13.3433045
C -4.1853409 20.9390259 -11.9780666
H -4.1201797 20.8378689 -10.8941409
C -5.8142116 21.9798425 -12.0685304
O -6.4577917 21.990306 -13.1238488
C -3.3553264 22.2522131 -12.3368714
H -2.7508435 21.9810248 -13.199742
H -3.9431639 23.1195955 -12.6376299
C -2.3935577 22.6639575 -11.2141059
H -1.5875927 23.2551301 -11.6539897
H -1.9417954 21.7737149 -10.7710014
C -3.1027948 23.4921229 -10.1365626
O -4.1338848 23.0043489 -9.6189633
O -2.6283562 24.6174618 -9.8712537
N -6.4475046 23.0719258 -11.2698861
H -5.8681384 23.316696 -10.4654281

C -7.9445496 23.8116708 -11.2481781
H -8.5313975 23.1447577 -11.8786809
C -8.4505552 25.3183528 -11.7282815
O -7.9562339 26.3541144 -11.2608184
C -8.5827225 23.7319299 -9.8142739
H -8.3098143 22.7644516 -9.3910452
H -8.1237068 24.4929663 -9.1820639
C -10.1128287 23.8553926 -9.699581
H -10.4295997 24.8665053 -9.9608487
H -10.3884539 23.7049408 -8.6526486
C -10.8425833 22.8299823 -10.5754585
O -10.9583405 23.1023724 -11.793048
O -11.2574013 21.7859035 -10.0265513
N -9.602907 25.5151302 -12.4609812
H -10.1970189 24.6947314 -12.6038089
C -10.0093518 26.7968804 -13.139427
H -9.6033406 27.6153906 -12.5465371
C -11.5373224 27.0770783 -13.3129338
O -12.3430523 26.1660776 -13.4651874
C -9.3158021 26.7963794 -14.5218177
H -8.2652178 26.5407625 -14.3723088
H -9.7616841 25.9988297 -15.1187812
C -9.3656862 28.1006173 -15.3396433
H -10.3980501 28.3527717 -15.5746801
C -8.7151959 29.2767986 -14.6053596
H -7.6855865 29.0272619 -14.3458923
H -9.2616884 29.5103425 -13.6946444
H -8.721151 30.1580805 -15.2468748
C -8.6202634 27.902131 -16.6598573
H -7.5714266 27.6723762 -16.4704162
H -8.6904418 28.808201 -17.2612677
H -9.0736317 27.0808258 -17.214682
N -11.9412789 28.3572898 -13.4297045
H -11.2467902 29.0828588 -13.3712445
C -13.3321225 28.7589476 -13.708797
H -13.9922764 28.0787705 -13.1669539
C -13.6973559 28.668285 -15.20173
O -13.2212429 29.4599915 -16.0166546
C -13.5976231 30.1744778 -13.1694646
H -13.3339687 30.2032292 -12.1112647
H -12.9526566 30.8857408 -13.6884859
C -15.0439728 30.6091374 -13.3172145
C -16.0007793 30.2075446 -12.3654889
H -15.7026398 29.6095012 -11.5164327
C -17.3497155 30.5734271 -12.5229045
H -18.084102 30.2600018 -11.7942258

C -17.7459906 31.3401123 -13.632773
H -18.784204 31.6156339 -13.7569758
C -16.7930155 31.742311 -14.5848431
H -17.0984245 32.326439 -15.4416514
C -15.4440982 31.3771052 -14.4276294
H -14.7165289 31.6734974 -15.1713742
N -14.6326318 27.780819 -15.5396291
H -14.9521194 27.1150618 -14.8471413
C -15.2390964 27.6886577 -16.8666391
H -15.735075 28.6312197 -17.0994544
H -14.4624243 27.5134405 -17.6111349
C -16.2666536 26.5596064 -16.957317
O -16.3953891 25.7524785 -16.0401148
N -16.9764725 26.4878918 -18.0819378
H -16.8289854 27.176356 -18.8044725
C -17.9575033 25.4408391 -18.3831007
H -17.7899232 24.5827215 -17.7304426
C -17.7792012 24.9927546 -19.8397808
O -17.7959146 25.8188285 -20.750926
C -19.3742105 25.9740925 -18.1133224
H -19.4072767 26.3626889 -17.0943744
H -19.5911057 26.7896238 -18.8053619
C -20.4418095 24.8784784 -18.2588574
H -20.2170614 24.0779836 -17.5490798
H -20.3816661 24.4589508 -19.2655484
C -21.873747 25.3850101 -18.0153021
O -22.0329252 26.4946208 -17.4568844
O -22.8044396 24.6367793 -18.3888393
N -17.5576422 23.6953514 -20.0649298
H -17.6282623 23.0479037 -19.293315
C -17.3269048 23.1256682 -21.396853
H -17.6295227 23.8524687 -22.1539443
C -18.173276 21.8705255 -21.6133325
O -18.4492201 21.1344351 -20.6665456
C -15.8303326 22.833808 -21.5971231
H -15.5031065 22.1074362 -20.8521036
H -15.2664698 23.755845 -21.4523805
C -15.5235885 22.3038595 -22.9769314
N -16.0573017 22.8083599 -24.1656877
C -15.6125519 22.0028993 -25.140073
H -15.854249 22.1181749 -26.1888167
N -14.8449621 21.02673 -24.6277346
H -14.429788 20.2755673 -25.1579691
C -14.7816493 21.1953241 -23.2621255
H -14.2791179 20.561661 -22.5447457
N -18.5185024 21.5818415 -22.8701808

H -18.1432063 22.1738025 -23.6009505
C -19.2288161 20.3661674 -23.257331
H -19.9116933 20.1067755 -22.4457962
C -18.2431347 19.2015468 -23.4437471
O -17.5034608 19.1367934 -24.4310812
C -20.0753688 20.6536823 -24.5101802
H -20.7622502 21.4721288 -24.2876809
H -19.4111275 20.9769363 -25.3134978
C -20.8899068 19.4416545 -25.0031907
H -20.2061397 18.632868 -25.252551
C -21.8900833 18.9305006 -23.9653095
H -22.5664824 19.7301243 -23.6638583
H -21.3716057 18.543429 -23.0898099
H -22.4721546 18.1090986 -24.387707
C -21.6558409 19.8178908 -26.2704486
H -22.3670983 20.6156827 -26.0549008
H -22.1966695 18.9460329 -26.6408658
H -20.9593117 20.149503 -27.0399621
N -18.305586 18.2264973 -22.5363312
H -18.9848484 18.3084584 -21.7879638
C -17.4987295 17.0022879 -22.56464
H -16.4445225 17.2767669 -22.5352472
C -17.7136686 16.1851165 -23.8479637
O -16.7324037 15.7061853 -24.4089253
C -17.8183622 16.1676056 -21.3118248
H -17.6392762 16.7679502 -20.4189096
H -17.1541192 15.3031549 -21.279748
C -19.2469521 15.6902349 -21.2993199
N -20.3555822 16.5291505 -21.1770985
C -21.409785 15.8002387 -21.5780685
H -22.404937 16.1941265 -21.7444035
N -21.0189659 14.5690479 -21.940586
H -21.5461555 13.999855 -22.611493
C -19.6585309 14.4792161 -21.7690455
H -19.0246639 13.6887216 -22.1457658
N -18.9342631 16.1416122 -24.3967885
H -19.7139306 16.4659898 -23.8383305
C -19.2306073 15.4622993 -25.6698397
H -18.9191364 14.4191048 -25.5943322
C -18.4740459 16.0915514 -26.8531044
O -17.994577 15.3816032 -27.732199
C -20.7422481 15.4827005 -25.9605596
H -20.9140981 15.0341016 -26.9402223
H -21.0845436 16.5175317 -26.0055241
C -21.5715667 14.7083567 -24.9337949
O -21.1731487 13.5796127 -24.57134

O -22.5721805 15.2767049 -24.4473625
N -18.2496141 17.4112082 -26.8323886
H -18.5716361 17.9329869 -26.0288096
C -17.4531297 18.1047217 -27.8489587
H -17.7977212 17.8135617 -28.8424323
H -17.5879439 19.1789559 -27.7304308
C -15.956921 17.7942072 -27.7454374
O -15.2888172 17.6280449 -28.7627028
N -15.439025 17.6201509 -26.5247796
H -16.0609943 17.7043393 -25.7294654
C -14.0743246 17.1266898 -26.3031017
H -13.3826248 17.7038566 -26.9186169
C -13.9256189 15.6534948 -26.724003
O -12.9564171 15.2990751 -27.3953963
C -13.702554 17.3422062 -24.8291262
H -13.7609572 18.4063683 -24.6038623
H -14.431762 16.8391893 -24.1953113
C -12.3186415 16.8515113 -24.4543145
C -11.189853 17.6482056 -24.7279804
H -11.3040053 18.609059 -25.2098121
C -9.9051194 17.1911998 -24.3777829
H -9.0312386 17.792005 -24.5808524
C -9.7460546 15.9368061 -23.7535044
O -8.5019852 15.5094553 -23.4137942
H -8.5099345 14.6569228 -22.9779248
C -10.8775819 15.1370117 -23.4808996
H -10.7593813 14.1747477 -23.0085044
C -12.161335 15.5959257 -23.8337612
H -13.0275541 14.9793858 -23.6353157
N -14.9268359 14.8201123 -26.423841
H -15.7014839 15.176822 -25.8741997
C -14.9828394 13.4197593 -26.8408227
H -14.0769896 12.9181517 -26.4986893
C -15.0310294 13.2711907 -28.3685764
O -14.316041 12.4302505 -28.9112801
C -16.1878629 12.7409781 -26.1724631
H -16.0987507 12.8425083 -25.0898935
H -17.1015123 13.2506904 -26.4752072
C -16.3194695 11.2716038 -26.5208461
C -15.679925 10.2960364 -25.7306851
H -15.1296517 10.5903453 -24.8487413
C -15.7427155 8.9385754 -26.1018896
H -15.2470437 8.1817352 -25.5143806
C -16.4327316 8.5582085 -27.2723703
O -16.4601685 7.2539103 -27.6505408
H -16.8895214 7.1591939 -28.5014005

C -17.087672 9.5346431 -28.0532165
H -17.603424 9.2540987 -28.9595907
C -17.035941 10.887789 -27.6716777
H -17.5222426 11.6367727 -28.2851494
N -15.7781968 14.1209507 -29.0785084
H -16.4213335 14.735133 -28.58639
C -15.8302485 14.0991621 -30.5458371
H -16.023302 13.0723458 -30.857743
C -14.4935853 14.5210283 -31.1785911
O -13.967663 13.81329 -32.0333803
C -16.994066 14.9750367 -31.0424011
H -17.90762 14.6865268 -30.519727
H -16.7819786 16.0183928 -30.8031416
C -17.238316 14.8508303 -32.5555673
H -16.3290961 15.1177248 -33.0992785
H -18.0038016 15.5779021 -32.8375267
C -17.6986865 13.4395358 -32.9540066
O -18.9264795 13.2539991 -33.097649
O -16.8225467 12.5548429 -33.0888592
N -13.837982 15.5729289 -30.6701081
H -14.2892367 16.1201696 -29.9440593
C -12.4898729 15.9648231 -31.1351682
H -12.5346458 16.1337827 -32.2113024
C -11.4694612 14.8360737 -30.9194148
O -10.6875397 14.5295424 -31.8238719
C -12.0510367 17.2817301 -30.4618222
H -12.2135606 17.2075124 -29.385962
C -10.5745282 17.6261432 -30.7001145
H -10.360449 17.6415137 -31.7693089
H -9.9326989 16.8899406 -30.2164525
H -10.3471552 18.6032195 -30.2737645
C -12.8768666 18.454303 -31.009272
H -12.6844 18.5843449 -32.0745506
H -12.6143381 19.3701809 -30.4802161
H -13.9403605 18.2682826 -30.8668087
N -11.5418689 14.1369177 -29.7810758
H -12.2068198 14.4377171 -29.0741423
C -10.7457172 12.934077 -29.5229422
H -9.6880299 13.1653715 -29.6509979
H -10.9128238 12.6231638 -28.4919327
C -11.1015855 11.7534031 -30.4368387
O -10.2052046 11.0525872 -30.9091058
N -12.3806552 11.5703557 -30.7665667
H -13.0669852 12.1981408 -30.3596135
C -12.8830576 10.4904933 -31.6335773
H -12.4536585 9.5465298 -31.299618

C -12.4546029 10.6930136 -33.0856704
O -11.8379689 9.7967565 -33.662114
C -14.4115139 10.3664975 -31.5108397
H -14.8837716 11.3195435 -31.7430037
C -15.0075931 9.281508 -32.405127
H -14.497612 8.3332624 -32.2385487
H -14.9184434 9.5764776 -33.4497101
H -16.0681325 9.1738969 -32.1753859
O -14.7191285 9.9990169 -30.1858573
H -14.6611214 10.8041515 -29.6445041
N -12.6061399 11.9050074 -33.6278135
H -13.1277568 12.5991368 -33.0943308
C -12.1108057 12.2899864 -34.96069
H -12.5626445 11.6330904 -35.7039491
C -10.5924574 12.1086371 -35.0606404
O -10.1118234 11.5017037 -36.0194828
C -12.5253388 13.7419652 -35.2846723
H -12.2047179 14.3950538 -34.4716908
C -11.9014053 14.2583629 -36.5896139
H -12.1485647 13.587198 -37.4129321
H -10.8182307 14.3273054 -36.4922998
H -12.2846337 15.2548982 -36.8119629
C -14.0473735 13.8569901 -35.4397281
H -14.390893 13.2481522 -36.2760117
H -14.3288191 14.8958381 -35.6094773
H -14.5551193 13.5168148 -34.5364626
N -9.8327613 12.5183483 -34.0376294
H -10.2691124 13.0158676 -33.2677802
C -8.3866883 12.2947826 -33.9964652
H -7.94803 12.713808 -34.9032965
C -8.0253687 10.7951613 -33.964585
O -7.1750255 10.3535353 -34.7385592
C -7.808095 13.0498299 -32.7940855
H -8.2322251 12.6640796 -31.8665257
H -8.0425363 14.1119177 -32.8768195
H -6.7249933 12.9258191 -32.7693519
N -8.7383063 9.9928143 -33.1690346
H -9.4399586 10.4216383 -32.5757625
C -8.5261026 8.5373236 -33.0519002
H -7.476906 8.3564513 -32.8190178
C -8.836892 7.7959867 -34.3557814
O -8.0721417 6.9214099 -34.7582541
C -9.3671494 7.960609 -31.9009351
H -10.4270834 8.1175921 -32.1029293
C -9.1210937 6.4714297 -31.6602582
H -8.0569718 6.2924769 -31.5037895

H -9.4663804 5.8883818 -32.5134126
H -9.6722286 6.1524028 -30.7752524
O -9.0287859 8.6085484 -30.6970051
H -9.4300372 9.4946216 -30.711502
N -9.8765834 8.2071453 -35.0894348
H -10.4751493 8.9342993 -34.7066042
C -10.218649 7.6484065 -36.4087586
H -10.3078933 6.565189 -36.3149444
C -9.0961985 7.9031619 -37.4291875
O -8.8107229 7.0319327 -38.2457573
C -11.5883493 8.2058925 -36.8724934
H -11.5800876 9.2895272 -36.7418409
C -11.8649231 7.9080697 -38.3605028
H -11.8182208 6.8336763 -38.5433383
H -11.1330206 8.4106419 -38.9928817
H -12.8465962 8.2794081 -38.6492608
C -12.7194695 7.6055139 -36.002287
H -12.8725628 6.5606242 -36.275951
H -12.4291906 7.6247014 -34.9527788
C -14.0547499 8.3529914 -36.1158443
H -14.469557 8.2558855 -37.1180749
H -13.9111854 9.4081356 -35.8800284
H -14.765007 7.9268607 -35.4065953
N -8.3937191 9.0388911 -37.3367375
H -8.6846259 9.7176446 -36.6427142
C -7.265239 9.3812153 -38.2229573
H -7.5533014 9.1179589 -39.2408252
C -5.9497281 8.6286735 -37.9283472
O -4.9741865 8.7465544 -38.6840646
C -7.0390962 10.899473 -38.1899562
H -8.0014138 11.4136505 -38.2093944
H -6.4800146 11.1924024 -39.0771632
O -6.2990547 11.3175345 -37.05805
H -6.6658562 10.914714 -36.2473093
N -5.8900761 7.8921141 -36.8100548
H -6.7249579 7.8243468 -36.2397125
C -4.6565738 7.3302429 -36.256341
H -3.8362244 8.0007177 -36.5123651
C -4.2800623 5.9408626 -36.8074929
O -3.1352983 5.5205919 -36.6491747
C -4.7709993 7.3240925 -34.7269169
H -5.5236596 6.6026775 -34.4107831
H -5.0474193 8.3167635 -34.3677705
H -3.8122191 7.0434425 -34.2890576
N -5.1800992 5.2659515 -37.5310566
H -6.0659794 5.7270073 -37.7010793

C -4.9863126 3.9392569 -38.1512939
H -5.0038412 3.1815968 -37.3671395
C -3.6467069 3.7571432 -38.8886477
O -2.9731921 2.7386014 -38.7264792
C -6.1697208 3.6740353 -39.1048049
H -7.0402153 3.4328757 -38.4935022
H -5.9499646 2.7970377 -39.7150907
C -6.5637775 4.8197352 -40.0201111
N -5.7699764 5.9207787 -40.3641479
C -6.5674691 6.752385 -41.049963
H -6.277414 7.7318812 -41.4028678
N -7.7926363 6.2244398 -41.1826337
H -8.5854946 6.6964794 -41.5940588
C -7.8129371 5.0107498 -40.5332027
H -8.6720512 4.3747877 -40.3736965
N -3.1845526 4.7824287 -39.6094995
H -3.8397485 5.5490973 -39.7402001
C -1.9094941 4.7790629 -40.3503428
H -1.9058746 3.9210138 -41.0259016
C -0.6755178 4.6000095 -39.4590681
O 0.3500762 4.1523355 -39.9634635
C -1.8108003 6.0655976 -41.1818839
H -1.9802055 6.9149351 -40.5182176
H -0.8101594 6.155225 -41.6088882
C -2.8366963 6.0737128 -42.3295833
H -3.7805566 5.6467642 -41.9998432
H -2.4675716 5.4691415 -43.158521
C -3.090666 7.5062857 -42.7975055
H -2.21557 7.8834898 -43.328123
H -3.2604295 8.1357064 -41.9227019
N -4.2898586 7.6146451 -43.6353146
H -5.0985667 8.0752973 -43.2163844
C -4.3632322 7.5927761 -44.9465972
N -3.4118636 7.0991039 -45.680707
H -3.2172373 7.5904414 -46.5573116
H -2.6560279 6.644648 -45.2205878
N -5.4028833 8.1065841 -45.5234574
H -6.0523492 8.6180683 -44.9202338
H -5.4423454 8.1830156 -46.5185052
N -0.7546542 4.8641941 -38.1528525
H -1.6478001 5.1514024 -37.7665882
C 0.3441051 4.6164829 -37.2169392
H 1.2323201 5.1264964 -37.5885598
C 0.6996833 3.1210416 -37.0906775
O 1.8637562 2.7984179 -36.8587498
C -0.0133202 5.2297426 -35.858726

H -0.9017486 4.748926 -35.4486878
H -0.2065543 6.2971976 -35.9704315
H 0.8169899 5.0909097 -35.1647239
N -0.2423564 2.2117153 -37.3765593
H -1.1831707 2.5200405 -37.5942453
C 0.0441412 0.7786215 -37.4595675
H 0.5673747 0.4724402 -36.5515695
C 0.9552045 0.4341165 -38.6550826
O 1.7906982 -0.4590169 -38.5465271
C -1.2872208 0.0221727 -37.522223
H -1.8167315 0.2672544 -38.4447155
H -1.9079066 0.2944651 -36.668599
H -1.1024213 -1.0504495 -37.4916568
N 0.8781965 1.1977947 -39.7544587
H 0.2102942 1.9555281 -39.7611379
C 1.8112123 1.0873253 -40.8905783
H 2.0039892 0.0301488 -41.0804667
C 3.1532747 1.7347408 -40.5479506
O 4.1918602 1.1211358 -40.7786454
C 1.2242149 1.7004207 -42.1796402
H 1.0712918 2.7691655 -42.0362597
C 2.1680245 1.5163423 -43.3761676
H 2.3959256 0.458749 -43.5151264
H 3.0968456 2.0616945 -43.2111974
H 1.7047064 1.9050683 -44.2829471
C -0.1270984 1.0684292 -42.5415657
H -0.0134202 -0.0094525 -42.6618157
H -0.4995357 1.4948188 -43.4727939
H -0.8607535 1.2658962 -41.7604837
N 3.1497115 2.9075864 -39.8983128
H 2.2584563 3.3646896 -39.7477886
C 4.3829764 3.5769137 -39.4271644
H 5.0316617 3.7532182 -40.287047
C 5.1714679 2.6870662 -38.4572727
O 6.3914124 2.6466122 -38.5342007
C 4.0854947 4.9447377 -38.7734971
H 3.4492565 4.7868126 -37.9038288
C 5.3579813 5.667574 -38.3077099
H 6.0473034 5.7700222 -39.1479344
H 5.8520541 5.0988091 -37.5198768
H 5.1091859 6.6515776 -37.9116408
C 3.3657123 5.9094878 -39.7292705
H 4.0142422 6.146524 -40.5715909
H 3.1117469 6.8290265 -39.2030367
H 2.4524687 5.4645623 -40.1099666
N 4.5099699 1.8691594 -37.6261136

H 3.501018 1.9661057 -37.5679173
C 5.1768327 0.8861925 -36.7517641
H 5.9189905 1.4157616 -36.1524491
C 5.9655105 -0.1805465 -37.5278367
O 6.9655235 -0.6580613 -37.005066
C 4.1151826 0.2654933 -35.821984
H 3.6872668 1.0584642 -35.2070059
H 3.3115988 -0.1409395 -36.4366679
C 4.6026419 -0.8663175 -34.8987054
H 4.9701494 -1.6946528 -35.5041773
H 3.7425868 -1.2253198 -34.3369632
C 5.691348 -0.4487287 -33.8979481
H 5.305648 0.3329929 -33.2411631
H 6.5347315 -0.0299299 -34.4515579
N 6.1852786 -1.5988657 -33.1065998
H 7.0302672 -2.023504 -33.4565587
C 5.6931212 -2.0929878 -31.9796119
N 4.6137617 -1.6238221 -31.4121932
H 4.2496792 -2.0315972 -30.5644291
H 4.1500759 -0.8388203 -31.8268032
N 6.2826626 -3.0960025 -31.392016
H 7.158683 -3.4460252 -31.7437805
H 5.9220268 -3.4384356 -30.5210906
N 5.5130667 -0.5786026 -38.7196586
H 4.6989518 -0.1098716 -39.0946508
C 6.155844 -1.6215128 -39.5477068
H 6.7171868 -2.2989085 -38.9044692
C 7.1870424 -1.0471198 -40.5141252
O 8.1672071 -1.7115576 -40.8379074
C 5.0729423 -2.429844 -40.2970744
H 4.4719049 -1.7392024 -40.8919679
C 5.6505611 -3.4778044 -41.2668926
H 6.3652394 -4.1194898 -40.7503349
H 6.1577817 -2.9777053 -42.0922651
H 4.8537515 -4.0898952 -41.6888069
C 4.1272498 -3.1381766 -39.3001663
H 3.4521743 -2.3981917 -38.8745679
H 3.519856 -3.8567523 -39.8472324
C 4.8022508 -3.8790266 -38.1377464
H 5.1620025 -3.1674568 -37.3956499
H 5.6444629 -4.4611102 -38.5038849
H 4.0797422 -4.5405924 -37.6627072
N 6.9634562 0.1827065 -40.9660073
H 6.1059518 0.645179 -40.6903566
C 7.8918005 0.9019409 -41.8307362
H 8.3560989 0.1968688 -42.5199967

C 9.0541694 1.5083209 -41.0392823
O 10.1751688 1.4175308 -41.5113396
C 7.0910835 1.9372281 -42.6425295
H 6.4755174 2.5141194 -41.9494156
C 8.015055 2.9405889 -43.3353853
H 8.8170607 2.4130698 -43.8517664
H 8.4436482 3.6079035 -42.5859543
H 7.4537262 3.5413332 -44.0376991
C 6.1329575 1.2547909 -43.6511673
H 5.3388 0.7519175 -43.1009127
H 5.6559528 2.0240523 -44.2518133
C 6.7607953 0.2206656 -44.5975052
H 7.0017951 -0.6915304 -44.0477814
H 7.6692296 0.6123557 -45.0487245
H 6.0481605 -0.0291608 -45.3837918
N 8.798992 1.9819869 -39.8195457
H 7.8269675 1.9509433 -39.5448266
C 9.6149082 2.8267209 -38.9312898
H 9.2170332 2.5797365 -37.9459199
C 9.3250434 4.3409519 -39.0306445
O 8.5238253 4.7993794 -39.8542185
C 11.099113 2.4020543 -38.8279642
H 11.7082873 3.1408157 -39.3434734
H 11.2510327 1.4269978 -39.2904088
C 11.5965131 2.3038259 -37.3833269
O 11.4492724 3.3205405 -36.6703128
O 12.1249463 1.2330349 -37.0094698
N 9.8588958 5.1058401 -38.0754761
H 10.516748 4.6466534 -37.4396315
C 9.3325992 6.4041998 -37.6512809
H 8.3526531 6.2042132 -37.2199303
C 9.1395831 7.4234129 -38.7896545
O 10.0494829 7.7492656 -39.5515223
C 10.2024572 6.9760756 -36.5200309
H 10.3373002 6.2059069 -35.7593743
H 11.1814432 7.2466722 -36.9189787
C 9.5601829 8.2096529 -35.8657287
H 8.5933201 7.9247972 -35.443284
H 9.3792816 8.9717041 -36.6261681
C 10.4478246 8.8008571 -34.7647909
O 10.8020372 8.0456266 -33.8314376
O 10.7428137 10.012847 -34.8410345
N 7.9356494 8.0039627 -38.8202964
H 7.2747294 7.6949909 -38.1276181
C 7.4979327 9.066309 -39.7246913
H 6.4475177 9.2219085 -39.4784392

C 7.4607978 8.7250306 -41.2328113
O 7.2182063 9.6195186 -42.0501456
C 8.189382 10.3821855 -39.339387
H 9.2533861 10.3291555 -39.5703739
H 8.0772996 10.5585665 -38.2684137
H 7.7299772 11.2135753 -39.8729389
N 7.5456185 7.4423073 -41.6155564
H 7.7541 6.7354562 -40.9135248
C 7.344726 6.9801426 -42.997353
H 6.9802142 7.8289962 -43.5714683
C 6.2228111 5.9398072 -43.1423331
O 5.8136015 5.298277 -42.1811172
C 8.7040589 6.5417867 -43.5937623
H 9.3769214 6.2789443 -42.781539
H 8.5857406 5.6690598 -44.2357264
C 9.2911549 7.7100855 -44.425368
H 9.1907816 8.6410518 -43.8675988
C 10.7676023 7.5379856 -44.7641775
H 10.9094523 6.6899523 -45.4326336
H 11.347273 7.4053009 -43.8535032
H 11.1408564 8.4293545 -45.2677901
C 8.5546316 7.8484516 -45.7653518
H 8.6226347 6.9085588 -46.3141622
H 9.0244539 8.6267692 -46.3653165
H 7.5122898 8.121164 -45.6183874
N 5.6747873 5.8134553 -44.3622881
H 6.051826 6.3868802 -45.1032308
C 4.5891328 4.8649351 -44.6980872
H 4.5631062 4.1139725 -43.9051263
C 4.8447935 4.0459821 -45.973334
O 3.9017478 3.6498304 -46.6520897
C 3.2105171 5.5603428 -44.666108
H 2.4533252 4.7738203 -44.6831793
H 3.1054279 6.0864467 -43.7153432
C 2.9161759 6.5454592 -45.8179066
H 3.1261552 6.0697121 -46.7735315
H 3.5448542 7.4315362 -45.7241825
C 1.4308664 6.9484155 -45.7776237
H 1.2590212 7.5693356 -44.8966521
H 0.8212512 6.0482975 -45.6740764
N 0.981203 7.6811794 -46.9780385
H 0.7524881 8.6673509 -46.8923419
C 0.6228584 7.177358 -48.1443321
N 0.8186153 5.929421 -48.4685675
H 0.5752193 5.6132175 -49.3870961
H 1.388143 5.3577715 -47.8677134

N 0.0406873 7.949872 -49.0092621
H -0.2933387 8.855375 -48.6610897
H -0.2623347 7.6163466 -49.9005672
N 6.1190758 3.8578277 -46.3209588
H 6.8301691 4.1363819 -45.6635574
C 6.5958218 3.2428633 -47.5626859
H 5.830005 2.5719029 -47.9542416
C 7.8916732 2.4367347 -47.343168
O 8.5668267 2.64766 -46.3401455
C 6.8674228 4.3491416 -48.5851797
H 7.7970464 4.8464069 -48.304794
H 7.032437 3.8770175 -49.5533758
C 5.793543 5.4093771 -48.7466088
C 4.5694425 5.071977 -49.3430479
H 4.3617759 4.0434651 -49.5976387
C 3.6563468 6.0786253 -49.6922932
H 2.7837729 5.8310304 -50.2772817
C 3.9157208 7.4151039 -49.3467661
H 3.2349086 8.196869 -49.6514519
C 5.0976506 7.7468283 -48.6644446
H 5.3156042 8.7854837 -48.4512446
C 6.0470079 6.7471097 -48.3865945
H 6.9997683 7.0245668 -47.966154
N 8.2414517 1.5676536 -48.3001291
H 7.6597218 1.5224626 -49.118932
C 9.3979198 0.6578697 -48.2914649
H 9.1016447 -0.2646165 -47.791358
C 10.6602547 1.1969945 -47.5742744
O 11.1786285 2.245083 -47.9427162
C 9.7087369 0.3224341 -49.75895
H 9.967396 1.234773 -50.3010873
H 8.8483582 -0.1504811 -50.2328802
H 10.5608699 -0.3531544 -49.8204047
N 11.1058984 0.4683517 -46.5404187
H 10.6020477 -0.3756197 -46.3181145
C 12.3498292 0.6411544 -45.7620579
H 12.0842348 0.2321049 -44.78769
C 12.7677959 2.0582833 -45.3295639
O 13.9257997 2.2966503 -44.965395
C 13.488643 -0.2542563 -46.2966031
H 14.4106292 0.3125754 -46.3885813
H 13.233996 -0.5667301 -47.3062372
C 13.7404825 -1.5012609 -45.4212878
O 12.782663 -1.9446545 -44.7479203
O 14.8558588 -2.0616043 -45.4835954
N 11.7676098 2.9221305 -45.1175289

H 10.8663841 2.6705479 -45.5032949
C 11.8434355 4.1137479 -44.2565654
H 10.9702733 4.7131008 -44.4953178
C 13.0474841 5.0013958 -44.5690478
O 13.2566867 5.4146228 -45.7033659
C 11.6839069 3.7215239 -42.7654516
H 10.7042933 3.2772 -42.7433744
C 12.7172875 2.6995277 -42.243958
H 13.7411237 3.021697 -42.4103113
H 12.5627012 1.7344082 -42.7242798
H 12.5786212 2.5252549 -41.1832717
C 11.6078686 4.8907969 -41.7666777
H 12.5088229 4.963613 -41.1569353
H 10.7748965 4.7152793 -41.0967139
H 11.4403098 5.837781 -42.2673589
N 13.7997814 5.3551081 -43.5357305
H 13.5427582 4.9757027 -42.6392564
C 14.9722896 6.1999847 -43.5773004
H 14.7513394 7.1316738 -44.1038787
C 16.1325397 5.5277022 -44.3124657
O 16.971215 6.2121722 -44.8791594
C 15.3363167 6.5059358 -42.1167684
H 14.5993989 7.1982355 -41.7073855
H 15.2826306 5.5845827 -41.5328418
C 16.7149966 7.0988042 -41.9408267
C 16.9780411 8.3984896 -42.403626
H 16.1908624 8.9809401 -42.8654804
C 17.7536204 6.3218351 -41.3924162
H 17.5600729 5.3123153 -41.0611771
C 18.2759221 8.9240865 -42.299878
H 18.4802673 9.9218835 -42.6403416
C 19.0546992 6.8452967 -41.3078177
H 19.8555147 6.2433117 -40.90308
C 19.314826 8.149128 -41.7601691
H 20.3159865 8.5514257 -41.6963891
N 16.2058513 4.196207 -44.3268838
H 15.3924863 3.6449569 -44.0809636
C 17.3365519 3.5317345 -44.9880782
H 18.247824 4.0148928 -44.6328091
C 17.3362498 3.7162578 -46.5202865
O 18.411362 3.6889561 -47.1212283
C 17.4139431 2.0551039 -44.544089
H 17.2640812 2.0186564 -43.4640312
C 18.7685605 1.4097662 -44.8475106
H 18.7772956 0.3945679 -44.4540022
H 19.5699411 1.9849547 -44.3903993

H 18.9200973 1.3641078 -45.9264033
O 16.4573901 1.2286298 -45.1599023
H 15.5577683 1.5969757 -45.0914068
N 16.1818698 4.0690221 -47.1078517
H 15.3465728 4.1192239 -46.5422669
C 16.0387294 4.3659216 -48.539581
H 16.9320402 3.9392428 -48.9945931
C 16.0907108 5.8966401 -49.0803814
O 16.0794458 5.9767522 -50.2908172
C 14.8556147 3.4957892 -49.0773819
H 13.9798023 4.1366509 -49.1933825
H 14.5968292 2.7346661 -48.3378472
C 15.1275989 2.7525254 -50.4024108
O 14.1400315 2.2286328 -50.965854
O 16.307982 2.6299838 -50.8096799
N 16.2900343 7.1238414 -48.4203981
H 16.4497442 7.0131143 -47.4277553
C 16.1613041 8.688626 -48.8951202
H 15.5643498 8.6270493 -49.8051702
C 17.4543756 9.8556596 -49.3389581
O 18.4787587 9.2741032 -49.0634247
C 15.1839224 9.2297703 -47.7945255
H 14.6070166 8.4267525 -47.3302822
H 14.4664933 9.8869038 -48.2885934
C 15.9020718 10.0680588 -46.713811
O 16.3872989 11.1886415 -47.009515
O 15.9809798 9.6471257 -45.537687
N 17.6582507 11.2710311 -50.0187799
H 16.7432841 11.6473397 -50.215925
C 18.9196021 12.4999217 -50.4996121
H 19.4239745 12.7072473 -49.557024
C 19.8910586 11.6284874 -51.3214552
H 19.3040143 10.926469 -51.916006
H 20.4598406 12.2416738 -52.0211191
C 20.8996192 10.8577388 -50.4511839
H 20.4908512 10.6758339 -49.4595413
H 21.799341 11.4600084 -50.3269145
C 21.2793501 9.5187183 -51.0747255
O 22.2065281 9.4215954 -51.8603748
N 20.5592735 8.459898 -50.7803827
H 20.794535 7.5910496 -51.2218482
H 19.7724068 8.5434667 -50.1355936
C 19.092366 14.182846 -51.3913657
O 18.0971829 14.350783 -52.0856275
N 20.1085432 15.3856543 -51.5493382
H 20.9260687 15.212857 -50.986138

C 20.2948739 16.9461427 -52.3131094
H 20.2227503 16.7196393 -53.3770916
C 21.4078328 18.3179152 -52.2406162
O 22.5468109 18.183726 -51.7988277
C 19.0686253 17.7464106 -51.9697146
H 19.2338989 18.2728335 -51.0391949
H 18.2005998 17.0932304 -51.9204808
H 18.8667864 18.4802587 -52.7438545
N 21.1702238 19.6749261 -52.6514902
H 20.2345851 19.8379585 -52.9935248
C 22.0024176 21.0115075 -52.4479166
H 22.8683875 20.7431257 -51.8441611
C 21.2832516 22.1579803 -51.6588977
O 20.098186 22.3880363 -51.8856444
C 22.5113401 21.7661131 -53.7349204
H 21.6390627 22.0773934 -54.3111731
H 23.0283835 22.6774737 -53.4283394
C 23.4507222 21.0203275 -54.6889031
H 22.9115143 20.1707727 -55.1106251
H 23.7236569 21.6915994 -55.5045992
C 24.7266258 20.5101083 -54.0066815
H 24.4530278 19.8220364 -53.2042071
H 25.2791937 21.3544152 -53.5879837
C 25.5984771 19.7681775 -55.0268131
H 25.909443 20.4728439 -55.8051529
H 24.9888066 18.9920389 -55.5006992
N 26.7854235 19.1515691 -54.3842775
H 27.3653918 19.8580064 -53.9512649
H 27.3408378 18.6543916 -55.0695705
H 26.4940335 18.4923601 -53.6723818
N 22.0141776 22.9790716 -50.8776924
H 22.9848334 22.7561847 -50.7481297
C 21.4363151 24.0595014 -50.0406279
H 20.6651534 23.5880877 -49.4217957
C 20.6932301 25.1571828 -50.8434591
O 21.1123859 25.4928957 -51.9501055
C 22.4954675 24.6021902 -49.0576117
H 21.9681595 25.0697176 -48.2281703
H 23.0446883 23.7611733 -48.6298199
C 23.4967813 25.6136439 -49.6395768
H 24.1308012 25.1185841 -50.3761876
H 22.9616084 26.426131 -50.1294037
C 24.3600993 26.2058559 -48.5122502
H 25.0695733 25.4548231 -48.159759
H 23.7129745 26.4715583 -47.6719088
C 25.1007745 27.47991 -48.9414417

H 25.6487684 27.8643149 -48.0751497
H 24.3538345 28.231996 -49.2207631
N 26.033051 27.2429212 -50.0717632
H 26.729265 26.5557402 -49.8134378
H 26.5058074 28.1019389 -50.3270792
H 25.5216444 26.9054137 -50.8778729
N 19.6653497 25.7902966 -50.2502701
C 18.5260935 26.4199977 -50.9302971
H 17.683172 26.2611953 -50.25573
C 17.9666354 25.9230463 -52.2825018
O 17.2120042 26.6858309 -52.8855268
C 18.8245867 27.9163968 -50.8994079
H 19.6064095 28.1600803 -51.6210341
H 17.9311673 28.5198536 -51.0658878
C 19.9982801 26.7186724 -49.1631419
H 19.6497361 26.3469128 -48.204326
H 21.0706298 26.8855675 -49.103357
C 19.3411503 28.0678737 -49.4726293
H 18.5032263 28.2293829 -48.7937525
H 20.0551531 28.8885143 -49.395854
N 18.1444302 24.6585658 -52.7170806
H 18.7575415 24.0313991 -52.2081714
C 17.2497154 24.0944877 -53.7603078
H 17.1809009 24.8305569 -54.5644565
C 15.8108259 23.9536811 -53.2184155
O 15.6116559 23.7733122 -52.0168994
C 17.7905208 22.7718561 -54.367871
H 18.2164828 22.1394345 -53.5915213
H 16.9504542 22.2274091 -54.799801
C 18.8275157 22.9966047 -55.490398
H 19.7490875 23.3914824 -55.0618205
H 18.4292468 23.7618966 -56.1540239
C 19.1471149 21.7322105 -56.331149
H 18.3107319 21.0347568 -56.2604309
H 20.0375569 21.2460875 -55.9308658
N 19.3493069 22.0762304 -57.7577649
H 19.2235844 23.0639289 -58.0004847
C 19.3902273 21.3013719 -58.8329725
N 19.5688706 20.0105413 -58.789513
H 19.5868354 19.4947518 -59.6476506
H 19.6328429 19.5190873 -57.9186205
N 19.2411043 21.8219077 -60.01574
H 18.7989704 22.7503794 -60.07748
H 19.2658129 21.2691169 -60.8598066
N 14.821947 23.9635607 -54.1171727
H 15.0640462 24.1295175 -55.0795782

C 13.4070457 23.7053603 -53.8023649
H 13.2318405 23.8314889 -52.7303655
C 13.0148305 22.2690653 -54.1901151
O 13.2650034 21.8554559 -55.3268438
C 12.5042714 24.70102 -54.554355
H 11.4657565 24.4018961 -54.4176821
H 12.7363243 24.6603459 -55.6166903
C 12.5918951 26.14774 -54.1111584
O 13.1869008 26.5106661 -53.1113018
N 11.9857542 27.0281791 -54.8750885
H 12.0663734 27.990201 -54.6088372
H 11.474667 26.7304553 -55.7012412
N 12.3104354 21.5618859 -53.3039725
H 12.1319232 21.9882772 -52.3959269
C 11.7798987 20.2088839 -53.5218785
H 12.1354009 19.8236533 -54.4750295
C 10.2499869 20.2006078 -53.5304971
O 9.636071 20.6928764 -52.5837385
C 12.2939867 19.2814334 -52.4104324
H 12.0003572 19.7111831 -51.4550013
H 13.3811827 19.2399207 -52.459261
C 11.7331357 17.8531895 -52.451972
H 10.6473428 17.8812916 -52.5280063
H 11.9562163 17.3702826 -51.5054196
S 12.3827862 16.813984 -53.7779941
C 13.8931229 16.2105975 -52.9728887
H 14.4627188 15.602152 -53.6752431
H 13.6285852 15.6018071 -52.1075691
H 14.5036321 17.0516099 -52.6454799
N 9.6561252 19.476022 -54.4810838
H 10.2356146 19.1014098 -55.228018
C 8.2618632 19.0294284 -54.4110718
H 7.7432147 19.5950109 -53.6411374
C 8.2012683 17.5367186 -54.0391612
O 8.6091906 16.6916482 -54.8315145
C 7.5848191 19.3377223 -55.7563497
H 7.6533027 20.404727 -55.9472126
H 8.1407942 18.8235216 -56.5354205
C 6.1094319 18.9161937 -55.8471269
H 6.0223168 17.8621598 -55.5987058
C 5.2008171 19.7105303 -54.9111302
H 5.2842959 20.7773402 -55.1160026
H 5.4789999 19.5191252 -53.8783082
H 4.1706631 19.3882301 -55.0433692
C 5.5958432 19.1157678 -57.2735893
H 5.6369121 20.1715894 -57.5396105

H 4.5696247 18.7588489 -57.3505647
H 6.2159663 18.5490014 -57.9673902
N 7.650806 17.207511 -52.8705797
H 7.2809473 17.9542867 -52.2913677
C 7.3710433 15.8279812 -52.4464654
H 8.0060151 15.1508264 -53.0126345
C 5.9027388 15.4793458 -52.7611271
O 4.9709185 16.075372 -52.2065813
C 7.75986 15.6405981 -50.9667627
H 7.1203425 16.2573012 -50.3370099
H 8.7862314 15.9905422 -50.8370111
C 7.6824443 14.1863395 -50.4760692
O 6.8708591 13.4090086 -51.019936
O 8.4087115 13.8653462 -49.5055914
N 5.6864177 14.5711102 -53.7189983
H 6.4974395 14.0713565 -54.0777662
C 4.3447916 14.1842873 -54.1843736
H 3.6345714 14.9349283 -53.8406842
C 3.9351687 12.8544228 -53.5605881
O 4.3232699 11.7831128 -54.0192536
C 4.2336867 14.1575708 -55.7201635
H 4.8568936 13.3737071 -56.1278179
C 2.7974381 13.8629456 -56.1479142
H 2.1426664 14.6095088 -55.703318
H 2.4969621 12.8683791 -55.8186253
H 2.7229068 13.8922377 -57.2343704
C 4.6461988 15.4961607 -56.3443168
H 4.0765308 16.3124174 -55.897042
H 4.4772019 15.482627 -57.4198195
H 5.7094427 15.6583488 -56.1699027
N 3.0257182 12.9205146 -52.5909151
H 2.717312 13.8404901 -52.2952782
C 2.6511322 11.7941466 -51.7378099
H 1.5919382 11.8736081 -51.495824
H 2.826334 10.8515972 -52.2505211
C 3.4378837 11.764253 -50.4292475
O 3.7783718 10.6859284 -49.9559292
N 3.6725557 12.9262538 -49.8125931
H 3.3218807 13.7634739 -50.2589932
C 4.5639542 13.1282033 -48.6574106
H 5.5763021 12.8435635 -48.9560907
C 4.2394702 12.317265 -47.379758
O 4.9944892 12.3378076 -46.4012804
C 4.557135 14.6320349 -48.3562011
H 4.9178299 15.1702423 -49.2356233
H 5.2327209 14.8285762 -47.5267004

S 2.8894722 15.2074225 -47.9074698
H 2.7061254 14.4032963 -46.8479299
N 3.0995178 11.6227276 -47.3420058
H 2.5537956 11.6005599 -48.1895828
C 2.6973692 10.7695608 -46.2284903
H 1.7296098 10.3306215 -46.4382888
H 3.4252333 9.964224 -46.1301078
C 2.5964225 11.5335775 -44.9096616
O 2.0756425 12.6458024 -44.8617237
N 3.1475683 10.9405853 -43.8409104
H 3.5995258 10.050913 -43.9813798
C 3.2644505 11.5676352 -42.5110625
H 2.5363332 12.3721322 -42.4922009
C 4.6024855 12.3215117 -42.2989558
O 4.9330438 12.7035656 -41.1837733
C 2.7685976 10.6219858 -41.3876643
H 3.329892 9.6866489 -41.3966698
H 2.9110664 11.1133101 -40.4234859
C 1.2482459 10.3497499 -41.5881163
H 0.7919812 11.3134804 -41.7977461
H 1.1127473 9.7337692 -42.4784474
C 0.4227669 9.7161131 -40.4391496
H 0.7314226 10.1512193 -39.4901884
H 0.5960085 8.639396 -40.4025918
C -1.0758004 10.0217816 -40.7095582
H -1.41602 9.4220592 -41.5561837
H -1.1388727 11.0716698 -41.0127608
N -2.0107233 9.8625097 -39.5598994
H -2.0996365 8.9127489 -39.2334567
H -2.9406933 10.1949484 -39.8135329
H -1.7309402 10.4529809 -38.7839952
N 5.3030928 12.6456133 -43.3932679
H 4.9509657 12.3158047 -44.2839729
C 6.263606 13.7539353 -43.4685514
H 6.339384 14.0373157 -44.5168193
H 5.8664254 14.6025745 -42.9108326
C 7.6966603 13.5217271 -42.9844836
O 8.5229441 14.4189115 -43.1446022
N 8.052298 12.3272621 -42.5032093
H 7.3432049 11.6119669 -42.3818921
C 9.4128024 12.0200965 -42.040962
H 9.6646061 12.6654394 -41.1981862
H 9.4474178 10.9834761 -41.7123813
C 10.481707 12.1834795 -43.1192423
O 11.5222019 12.7773601 -42.8556136
N 10.1750316 11.7843984 -44.3589289

H 9.2846575 11.3332788 -44.4955374
C 11.0186339 12.0413754 -45.5425943
H 11.9980506 11.588156 -45.3790969
C 11.2701527 13.5389522 -45.7486745
O 12.4236188 13.9518113 -45.8260238
C 10.4258214 11.3832015 -46.8083101
H 10.5905208 10.3099959 -46.7269139
C 8.9312833 11.6033541 -47.0463739
H 8.7099762 12.6632397 -47.1442538
H 8.3393666 11.1827496 -46.2344072
H 8.6361123 11.1131843 -47.9758533
O 11.0895781 11.8306891 -47.9597574
H 10.484578 12.3844576 -48.4701324
N 10.2252136 14.371186 -45.663101
H 9.3074535 13.9722939 -45.5446904
C 10.2857366 15.8236971 -45.8547703
H 10.7235008 16.0265557 -46.8268567
C 11.1106349 16.537419 -44.7650356
O 11.8863183 17.4526382 -45.0535196
C 8.8657689 16.3970441 -45.8856242
H 8.8986116 17.4030429 -46.3069571
H 8.4867539 16.4648006 -44.8668808
S 7.7163401 15.3759861 -46.8510702
H 8.4533831 15.1086649 -47.9512639
N 10.9632892 16.0999518 -43.508704
H 10.2825801 15.3646959 -43.3383485
C 11.7666912 16.5933009 -42.3777083
H 11.7210304 17.682167 -42.3558897
C 13.2337723 16.2149077 -42.5646649
O 14.1041332 17.0794444 -42.4631432
C 11.2181531 16.0596538 -41.0385742
H 11.0788588 14.9803609 -41.1088303
C 12.1583111 16.3454275 -39.8608403
H 12.413162 17.4044263 -39.8365978
H 13.0713173 15.7583224 -39.9544092
H 11.6747709 16.0658247 -38.9240883
C 9.8619149 16.7101419 -40.7311641
H 9.9738266 17.7906836 -40.6408162
H 9.465485 16.313601 -39.7953368
H 9.147258 16.4904463 -41.5236473
N 13.5086787 14.9554922 -42.9189931
H 12.7374487 14.300883 -43.0121126
C 14.8616324 14.4600437 -43.1763215
H 15.4605874 14.5795315 -42.2714439
C 15.5717593 15.257738 -44.2626702
O 16.6984701 15.6980688 -44.0520232

C 14.8119003 12.9696725 -43.5454865
H 15.47017 12.778969 -44.3885111
H 13.8159203 12.6800994 -43.8734669
C 15.2350372 12.1081566 -42.3887418
N 16.5503927 11.9810507 -41.9354251
C 16.4753185 11.2196844 -40.8299497
H 17.3253826 10.9192111 -40.2306378
N 15.2029599 10.8627996 -40.5836198
H 14.890693 10.2906702 -39.8085624
C 14.4037633 11.4233475 -41.5538219
H 13.3257702 11.3738953 -41.6274226
N 14.9078435 15.5075468 -45.387301
H 13.989292 15.091668 -45.5152898
C 15.5112271 16.2350314 -46.5011762
H 16.4944499 15.8005517 -46.6810838
C 15.788866 17.702243 -46.1455768
O 16.9395239 18.1374501 -46.2012692
C 14.6897345 16.0179734 -47.7882799
H 15.1327128 16.6236197 -48.5603665
C 14.7908711 14.5234165 -48.182398
H 14.1962195 13.8987777 -47.5198436
H 15.8220997 14.1781548 -48.1262874
H 14.4372692 14.3716629 -49.2011021
C 13.2366266 16.4975189 -47.7320752
H 12.7646062 16.0195718 -46.8976035
H 13.2059957 17.5711979 -47.5585925
C 12.4407379 16.1837724 -49.0033466
H 12.2244151 15.1164961 -49.062049
H 13.0189562 16.48687 -49.8741816
H 11.4970485 16.7230964 -48.9888032
N 14.8187234 18.4245126 -45.5793613
H 13.9008259 18.0120409 -45.4473034
C 15.0314858 19.8172963 -45.1857533
H 15.4248899 20.3646793 -46.0422485
C 16.0654803 19.9815299 -44.052507
O 16.775709 20.9872676 -44.0197683
C 13.6739593 20.4215669 -44.8288207
H 13.2487701 19.8884107 -43.9775321
H 12.9997992 20.3346726 -45.682005
H 13.7950387 21.4777533 -44.588073
N 16.1945128 19.0031707 -43.1417661
H 15.5550477 18.2125874 -43.1960906
C 17.1426889 19.0652184 -42.0165262
H 17.2682905 20.115006 -41.7540704
C 18.5555244 18.5806105 -42.3633946
O 19.5022617 19.2022987 -41.8920977

C 16.5356863 18.3408313 -40.8014493
H 15.5546198 18.7709715 -40.5929376
H 16.3868088 17.2899072 -41.0560764
C 17.3783728 18.4041283 -39.5137768
H 18.2761985 17.8007418 -39.655826
H 16.803368 17.9420041 -38.7110996
C 17.8159365 19.8120997 -39.0658109
H 18.4907804 20.2290409 -39.816017
H 18.3932875 19.7217441 -38.1440225
N 16.6897875 20.7491832 -38.8609668
H 16.5665419 21.4502419 -39.5716769
C 15.8807065 20.8193544 -37.8169131
N 14.9853566 21.7589121 -37.7435133
H 14.3645694 21.8061651 -36.956886
H 14.8726153 22.4170928 -38.4946404
N 15.9472023 19.9843426 -36.8189475
H 16.6241288 19.2455779 -36.8502505
H 15.2949912 20.0477172 -36.0589826
N 18.7300829 17.5290883 -43.1764286
H 17.9068943 17.0488226 -43.5306661
C 20.0684893 17.0390531 -43.5746573
H 20.7065707 16.9291063 -42.6964889
C 20.7865705 18.0002679 -44.5366372
O 22.0134416 18.0053269 -44.5586972
C 19.9813346 15.699411 -44.3219547
H 19.304658 15.8747004 -45.1616404
H 20.9671976 15.5209637 -44.7569981
C 19.5628823 14.3578569 -43.6784448
H 20.3290906 14.0366348 -42.971993
H 18.6129878 14.4437749 -43.1562454
C 19.4543242 13.3380914 -44.844005
H 20.418487 13.3271226 -45.354375
H 18.7167883 13.694725 -45.5669937
C 19.1362141 11.8730891 -44.4980398
H 19.490443 11.6480878 -43.4892399
H 19.686472 11.2395421 -45.2001729
N 17.6955314 11.5486037 -44.6343123
H 17.1272324 12.0169392 -43.9401613
H 17.4806865 10.5531329 -44.5600195
H 17.3202713 11.7308229 -45.5705452
N 20.0478271 18.6624719 -45.4364741
H 19.046923 18.5093921 -45.436498
C 20.6358853 19.3309775 -46.6085273
H 21.6749389 19.5607893 -46.3684505
C 20.0541746 20.7091617 -46.9622581
O 20.3152965 21.2210728 -48.0515675

C 20.6868561 18.354292 -47.7957786
H 21.0473326 18.879063 -48.6814314
H 21.4566581 17.6177888 -47.5613333
C 19.4373287 17.602644 -48.1667651
C 19.394892 16.1940631 -48.1271936
H 20.196105 15.6434021 -47.6560693
C 18.3451897 15.5113149 -48.7728262
H 18.2923324 14.4372359 -48.7484014
C 17.3896775 16.2497781 -49.5044677
O 16.5031357 15.6745726 -50.3589183
H 16.8175994 14.8393012 -50.723428
C 17.3695605 17.6457992 -49.4114359
H 16.6398896 18.1806343 -49.9827419
C 18.3848271 18.32481 -48.7295873
H 18.4231516 19.3990316 -48.7319403
N 19.3517895 21.3539171 -46.024161
H 19.1651048 20.8775033 -45.1539639
C 19.0384741 22.7911583 -46.0750765
H 18.3539854 22.9771147 -45.2457682
C 18.2660689 23.2374827 -47.3416898
O 18.3498463 24.3844502 -47.7803859
C 20.3254224 23.5929801 -45.7682459
H 21.0190659 23.4837652 -46.6030665
H 20.8007875 23.1682607 -44.8823023
C 20.0831759 25.0814136 -45.500918
O 19.0655738 25.4065578 -44.8420731
O 20.8655128 25.9146396 -46.0124582
N 17.4548563 22.3556176 -47.9295338
H 17.4238769 21.424189 -47.5465261
C 16.4823397 22.7281121 -48.9710723
H 16.9464183 23.4477338 -49.6473765
C 15.24281 23.3977265 -48.3725054
O 14.9522023 23.2457856 -47.1836795
C 16.0422905 21.5162724 -49.8130752
H 15.2603537 21.8499514 -50.4920367
C 17.2011516 21.0558416 -50.7125561
H 18.1143336 20.9147098 -50.1404907
H 17.4026366 21.8357731 -51.4439333
H 16.943333 20.1436236 -51.2459316
C 15.398919 20.4182088 -48.9405732
H 16.1665696 19.8238888 -48.4571155
H 14.7955406 20.8598812 -48.151009
C 14.4493492 19.5350595 -49.7478417
H 14.8862927 19.2531794 -50.7036183
H 13.5268763 20.0795328 -49.9194258
H 14.2201392 18.6408149 -49.1834476

N 14.4360211 24.010578 -49.2375354
H 14.7402083 24.0446147 -50.2085445
C 12.9993631 24.158699 -49.0076405
H 12.8175786 24.2573771 -47.9397338
C 12.2662528 22.9046082 -49.5103378
O 12.6107608 22.3634176 -50.5604784
C 12.5234879 25.4517901 -49.6859483
H 13.0406288 26.283309 -49.2048236
H 12.830272 25.44303 -50.731898
C 11.0112581 25.7338392 -49.6079951
H 10.8575365 26.7986213 -49.7828324
H 10.658697 25.5126149 -48.6002629
C 10.1424015 24.9839405 -50.6247752
O 10.5934761 24.3664138 -51.5757024
N 8.8434225 24.9954737 -50.4307784
H 8.2650384 24.4828781 -51.0906918
H 8.4331033 25.5005896 -49.6701504
N 11.2359635 22.4621827 -48.7844512
H 11.0578115 22.9049529 -47.8891113
C 10.3429926 21.383246 -49.214284
H 10.5949631 21.0920705 -50.235015
C 8.8711568 21.8264889 -49.2272665
O 8.3637704 22.3460571 -48.2286926
C 10.5732244 20.1618264 -48.309774
H 10.3332883 20.4190762 -47.2777976
H 11.6227969 19.8672222 -48.3609254
S 9.5400098 18.7650291 -48.8429061
H 8.3402798 19.3259352 -48.6493615
N 8.1565084 21.4877271 -50.299685
H 8.6580875 21.1050509 -51.0997332
C 6.6915622 21.4723115 -50.3796701
H 6.2622286 21.8728319 -49.4665579
C 6.2201094 20.031677 -50.5303368
O 6.4399128 19.4158451 -51.5664035
C 6.1943466 22.3420754 -51.5434893
H 6.7074003 22.060723 -52.4640156
C 4.6826819 22.2560971 -51.7615391
H 4.1511206 22.4652032 -50.8329504
H 4.408085 21.2625488 -52.117419
H 4.3832472 22.9787312 -52.5207642
O 6.4597851 23.693421 -51.2481149
H 6.0035097 23.908674 -50.423727
N 5.5750189 19.4858299 -49.5003679
H 5.4489048 20.0421054 -48.6609263
C 4.8934822 18.1915638 -49.5707935
H 5.3805521 17.5583986 -50.3075649

H 4.9561912 17.6920084 -48.6044006
C 3.4178491 18.3381219 -49.9451647
O 2.7499778 19.2457229 -49.4508098
N 2.8759735 17.4192718 -50.7447628
H 3.489359 16.728848 -51.172119
C 1.4218424 17.2768095 -50.9304672
H 0.9352328 17.8212694 -50.1261785
C 0.9560668 15.8305307 -50.7697592
O 1.6406314 14.8966804 -51.1867693
C 0.9136146 17.8905378 -52.2565359
H -0.1448096 17.6360649 -52.3435319
C 1.6387051 17.3248981 -53.5004349
H 2.6843131 17.6363338 -53.4916327
H 1.6106378 16.2363657 -53.4623999
C 0.9961788 19.4249524 -52.1858548
H 2.0365676 19.7521869 -52.1926341
H 0.5247676 19.7783184 -51.266841
H 0.4642133 19.8755679 -53.0218273
C 1.0052803 17.7519525 -54.8316522
H 1.1088672 18.8260369 -54.9781292
H -0.0515642 17.481976 -54.8436029
H 1.510511 17.2397663 -55.65089
N -0.2477491 15.6495421 -50.2260329
H -0.7465903 16.4668413 -49.8863459
C -0.9656272 14.3704334 -50.212057
H -0.6470092 13.784003 -51.0743246
C -2.4736961 14.6046495 -50.3361702
O -2.9923013 15.6108318 -49.855645
C -0.6345655 13.5626088 -48.9469876
H -1.035592 14.063649 -48.0667914
H 0.4483293 13.4718768 -48.8486547
O -1.2039403 12.2647033 -49.0661658
H -0.8230701 11.6633302 -48.3915009
N -3.189996 13.6624306 -50.9545782
H -2.693636 12.8593069 -51.3085801
C -4.6570956 13.7038033 -51.1109514
H -4.9453094 14.6681594 -51.5324737
C -5.3659256 13.5807981 -49.7505522
O -6.4750307 14.0756237 -49.5707605
C -5.0838402 12.5810381 -52.0946599
H -4.5868338 11.656628 -51.7934535
C -6.60297 12.3174183 -52.0927149
H -7.1427278 13.2367823 -52.3255887
H -6.9278242 11.951214 -51.1188279
H -6.8636599 11.5560362 -52.8261546
C -4.6265523 12.9470809 -53.5247128

H -5.2699191 13.744181 -53.8956398
H -3.6030739 13.3215718 -53.5000957
C -4.6555746 11.788384 -54.5300713
H -5.6777838 11.458972 -54.7113472
H -4.0625365 10.9539342 -54.1543668
H -4.2321241 12.1260136 -55.4769672
N -4.7271601 12.913471 -48.788543
H -3.775143 12.6272275 -48.9723444
C -5.2999645 12.5558781 -47.4919011
H -6.3581933 12.3426238 -47.6238231
C -5.12327 13.6827605 -46.4548777
O -3.9915082 13.9538372 -46.0403971
C -4.6452439 11.2446981 -47.0565748
H -3.587913 11.2809028 -47.3174804
H -5.1004324 10.4159779 -47.6001787
O -4.7366918 11.0128571 -45.6723862
H -5.6758702 11.0025864 -45.3613888
N -6.2181431 14.2734505 -45.9265974
C -6.1448261 15.2535177 -44.8392723
H -5.5037262 16.0768951 -45.1469994
C -5.5994107 14.6593758 -43.5365247
O -4.990551 15.3639187 -42.7414758
C -7.5767455 15.7683423 -44.6357585
H -8.056633 15.2366621 -43.8112928
H -7.5919532 16.8425494 -44.4489233
C -7.6005661 14.1249832 -46.3585443
H -7.6892693 14.0050037 -47.4364082
H -8.051605 13.2737373 -45.845865
C -8.2932888 15.4153499 -45.9362787
H -8.111043 16.1900229 -46.680453
H -9.3633916 15.2698172 -45.7853592
N -5.798928 13.3603836 -43.3244477
H -6.3334307 12.8472974 -44.0199305
C -5.3819279 12.5909009 -42.1533687
H -5.5086175 13.2194111 -41.2710266
C -3.889537 12.1953296 -42.1819728
O -3.2267231 12.2061379 -41.1417984
C -6.3412377 11.3905832 -41.9922851
H -7.3113502 11.7703061 -41.6675729
H -5.9575394 10.7498656 -41.1971473
C -6.5663179 10.5388814 -43.2538189
O -6.908747 11.078591 -44.3329274
O -6.3508923 9.3107568 -43.1812859
N -3.3008267 11.9683216 -43.3619274
H -3.8866016 11.9470974 -44.1933319
C -1.8430178 11.9831623 -43.5618261

H -1.3723892 11.257173 -42.9150736
C -1.2618897 13.3502526 -43.1982805
O -0.4067809 13.4546671 -42.3135894
C -1.4776396 11.6187214 -45.0103698
H -2.1218696 12.1563712 -45.7031098
H -0.4492902 11.9202012 -45.2048059
C -1.5826308 10.1109484 -45.2530965
H -2.542401 9.7508069 -44.8866851
H -0.8029225 9.6052696 -44.6769365
C -1.4265517 9.7523301 -46.7342513
O -2.1887154 8.9014817 -47.2378949
O -0.4387212 10.1870761 -47.3704002
N -1.8238071 14.3981459 -43.7998905
H -2.5652821 14.2183388 -44.4716944
C -1.3629664 15.7743681 -43.644951
H -0.2966211 15.7971953 -43.8619811
C -1.5011144 16.2671664 -42.1931932
O -0.5669668 16.8774303 -41.6842395
C -2.0937697 16.6432986 -44.6956286
H -3.1616424 16.4331521 -44.6208048
C -1.6453572 16.3062192 -46.1415492
H -1.8060816 15.2459139 -46.3228022
H -2.2857106 16.8442621 -46.8391984
C -1.9169274 18.1412204 -44.4519846
H -0.8624135 18.3648704 -44.4776171
H -2.351247 18.4235725 -43.4927015
H -2.4173335 18.7036014 -45.2397571
C -0.1850993 16.6247081 -46.5047295
H 0.502714 16.0852862 -45.8550906
H 0.005957 17.6948585 -46.4390428
H -0.0012286 16.3148047 -47.5317343
N -2.5671139 15.9256739 -41.4689525
H -3.3267024 15.4447242 -41.9411177
C -2.7735512 16.3191514 -40.0675664
H -2.7786111 17.4067291 -40.0129945
C -1.6436206 15.8404344 -39.1348382
O -1.1884872 16.6049745 -38.2795479
C -4.1712491 15.8244989 -39.6523631
H -4.8964345 16.3669418 -40.2600162
H -4.276831 14.7663523 -39.8970248
C -4.5547229 16.0502673 -38.1793378
H -5.6347657 16.1922723 -38.1337433
H -4.0852274 16.9595111 -37.8033673
C -4.2106249 14.87867 -37.2602295
O -3.478791 13.9624486 -37.5900405
N -4.7524033 14.838826 -36.0651956

H -4.5608134 14.0108061 -35.5281391
H -5.3835633 15.5549238 -35.7602172
N -1.1004638 14.6424705 -39.3708469
H -1.5013084 14.0764096 -40.106404
C 0.0418472 14.1322373 -38.6116889
H -0.0436926 14.4501244 -37.5713118
C 1.3850099 14.6518156 -39.1239408
O 2.2660308 14.8908551 -38.3035319
C 0.0435491 12.6104591 -38.6527647
H 1.0465421 12.2488474 -38.4138007
H -0.2262749 12.3017169 -39.6598596
S -1.1117245 11.9445062 -37.4241918
H -2.0986728 12.845829 -37.6105998
N 1.5575739 14.8170369 -40.4417058
H 0.8054631 14.5690666 -41.0762493
C 2.7701594 15.422793 -40.987119
H 3.6444812 14.9045317 -40.5883586
C 2.8696392 16.8863013 -40.5636503
O 3.5847836 17.2487937 -39.6283336
C 2.7907183 15.3258836 -42.5238653
H 1.9156234 15.7985274 -42.9702768
H 2.7822393 14.2994419 -42.8465516
H 3.7036186 15.7788831 -42.9060338
N 2.1002966 17.7180647 -41.2791163
H 1.1970245 17.3057366 -41.4945638
C 2.6117307 18.5517644 -42.3861192
H 3.1341734 17.8723516 -43.0516016
C 1.3970127 19.0600589 -43.2242848
H 1.358215 18.4262558 -44.0974844
H 0.5018309 18.6692998 -42.7733033
C 1.1166488 20.5727843 -43.5444475
H 1.2915895 21.151315 -42.6635274
H 1.8411256 20.9728381 -44.2381746
C -0.3230066 20.9360951 -43.9914732
H -1.0227159 20.3720331 -43.3859128
H -0.4699022 20.6178342 -45.0179328
C -0.6963571 22.4298742 -43.8025919
H 0.1995555 22.9755746 -43.4898631
H -1.4260614 22.4979107 -42.9928553
N -1.2635441 23.0865246 -45.012343
H -0.5895041 23.0906335 -45.7804691
H -1.4992876 24.0597524 -44.8902245
H -2.0834365 22.6400856 -45.4285437
C 3.7642777 19.4606364 -41.9383207
O 4.6934606 18.9814363 -41.3095185
N 3.6311882 20.7771193 -41.9989438

H 2.8876535 21.1417414 -42.5604266
C 4.0788258 21.6000937 -40.8865894
H 5.1237588 21.3834285 -40.6673247
C 3.9999911 23.0820359 -41.3327063
H 3.0493725 23.2473995 -41.8409777
H 4.7944901 23.2717171 -42.0553756
C 4.1026739 24.1179475 -40.2399803
N 3.0918861 25.0331692 -39.9409579
C 3.4223691 25.5589047 -38.7527186
H 2.8010558 26.2448663 -38.192991
N 4.5752577 25.0358408 -38.3042514
H 4.8930887 25.1113105 -37.3429746
C 5.0364284 24.1390869 -39.2428601
H 5.8685618 23.4582498 -39.1352582
C 3.2420288 21.199852 -39.6385361
O 3.0797663 22.025223 -38.7500341
N 2.530232 20.0383135 -39.640792
H 2.865367 19.2416506 -40.1623869
C 1.4823966 19.8353562 -38.7041374
H 1.2481924 20.7922532 -38.2384218
C 2.0858318 19.0933861 -37.5361898
O 3.067777 19.6165891 -37.0050026
C 0.1413169 19.4928001 -39.3442083
H 0.1442248 18.4588372 -39.6817178
H -0.6141341 19.569777 -38.5611991
C -0.267998 20.4123699 -40.5175301
H 0.1183068 19.9509151 -41.3678312
C -1.7776922 20.4353344 -40.70414
H -2.2492346 20.8376265 -39.8066394
H -2.1313518 19.4171078 -40.8620015
H -2.0806715 21.041301 -41.5528518
C 0.1944014 21.8747087 -40.5369619
H 1.2344464 22.003532 -40.7787416
H 0.035205 22.3050513 -39.5470494
H -0.3843236 22.4742456 -41.2283432
N 1.4878269 18.020571 -37.0275607
H 0.680745 17.619781 -37.490007
C 1.8370793 17.567107 -35.682473
H 1.6480608 18.3923588 -34.994206
C 3.3255548 17.1909504 -35.5105984
O 3.975249 17.6990186 -34.5966676
C 0.897452 16.4265003 -35.2938252
H 1.0844826 15.5580817 -35.9233497
H -0.1416476 16.7397962 -35.4098183
H 1.0723946 16.150676 -34.2533891
N 3.895938 16.3935091 -36.4204664

H 3.3273487 16.0340792 -37.1822979
C 5.2891756 15.9425756 -36.3358269
H 5.4806631 15.6036171 -35.3165186
C 6.3276383 17.0397494 -36.6405866
O 7.4910534 16.9063308 -36.2670223
C 5.4618095 14.744717 -37.274927
H 5.2733635 15.0489023 -38.3063018
H 4.7617187 13.9547081 -37.0015401
H 6.47971 14.3605827 -37.1961688
N 5.9285687 18.1328717 -37.2982429
H 4.962632 18.1795496 -37.6016175
C 6.8314934 19.2418052 -37.635999
H 7.8429605 18.8505207 -37.6832907
C 6.844892 20.3829303 -36.6121256
O 7.5699517 21.3645417 -36.8076774
C 6.4883151 19.7654549 -39.0249614
H 5.4763035 20.1549587 -38.9918722
H 6.5452826 18.940506 -39.7349417
O 7.3909096 20.7802827 -39.4271918
H 7.5897174 21.3108912 -38.6363837
N 6.0301942 20.3258281 -35.5583977
H 5.4633655 19.4990513 -35.4131305
C 5.9102691 21.4436743 -34.6242259
H 5.8569889 22.3420848 -35.2375565
C 7.1639527 21.593333 -33.7492034
O 7.6308825 20.6298769 -33.149975
C 4.6035516 21.3149437 -33.8160339
H 3.9610644 20.5586151 -34.2705869
H 4.8261419 20.983383 -32.8007128
C 3.8146552 22.6339158 -33.7630863
H 4.3761121 23.361526 -33.1761685
H 2.8597389 22.454182 -33.2696363
C 3.5503843 23.2164719 -35.1512038
O 4.0029425 24.3035214 -35.4804174
N 2.9488639 22.4629638 -36.0411139
H 2.950617 22.7735293 -37.0102779
H 2.7918841 21.4817724 -35.8719266
N 7.7210904 22.8081187 -33.6914249
H 7.2987774 23.5567177 -34.2190106
C 8.9972279 23.0788989 -33.0131334
H 9.0701395 24.1482844 -32.8145984
H 9.0214978 22.5526099 -32.0586359
C 10.2493908 22.6746477 -33.805537
O 11.3286693 22.5814544 -33.2268775
N 10.1271567 22.3925806 -35.1093067
H 9.2142375 22.4390954 -35.5407744

C 11.2828804 22.1739636 -35.9765919
H 12.1470555 21.9182832 -35.3639204
C 11.6311631 23.4425166 -36.7703543
O 10.8468171 23.8739616 -37.6062429
C 11.0248787 21.0051895 -36.9419587
H 10.2282802 21.2982624 -37.627223
H 11.9359448 20.8776523 -37.5163873
C 10.6521745 19.6372191 -36.3484215
H 9.6886744 19.7031776 -35.8480046
C 10.554009 18.6256642 -37.4922608
H 11.5421073 18.4392663 -37.908598
H 9.9033941 19.0025286 -38.2786216
H 10.1402736 17.6890333 -37.1141196
C 11.700169 19.1190278 -35.363519
H 12.6761902 19.0615675 -35.8409176
H 11.4049478 18.1266817 -35.0192001
H 11.7434391 19.7788958 -34.4960605
N 12.8948299 23.8661966 -36.7091582
H 13.3352564 23.7065076 -35.812184
C 13.4155394 25.0179326 -37.4790076
H 12.6681205 25.8145446 -37.4282954
C 13.6732908 24.7387133 -38.9836209
O 14.3679382 25.4938641 -39.6582383
C 14.7076514 25.5493859 -36.8171534
H 15.5330047 24.8870948 -37.0856383
H 14.9254605 26.5347873 -37.2302733
C 14.686988 25.6625318 -35.2845744
H 15.5866248 26.1902592 -34.9596257
H 14.7320807 24.6561241 -34.8586402
C 13.4378158 26.3886491 -34.7758366
O 13.4280188 27.6368107 -34.819975
O 12.4900572 25.6633847 -34.4019596
N 13.2648524 23.5772005 -39.5105643
H 12.5354699 23.085907 -39.0068225
C 13.5570177 23.1795007 -40.8976494
H 14.5254876 23.6009869 -41.1761233
C 12.542468 23.7627457 -41.8860466
O 11.3445727 23.7642826 -41.6133993
C 13.6717175 21.6505914 -41.0511484
H 14.6737933 21.3409764 -40.7633607
H 13.5409867 21.3814689 -42.0977122
C 12.6807254 20.8409029 -40.234541
O 13.0819143 19.9571612 -39.4998137
N 11.4063889 21.1652132 -40.2593715
H 10.7663929 20.6171536 -39.7086994
H 11.0835354 21.9909526 -40.7547637

N 13.0133136 24.1123106 -43.0888082
H 14.003369 23.9861169 -43.252692
C 12.2559559 24.8063065 -44.14478
H 11.634147 25.5703974 -43.6725806
C 11.2900693 23.8636724 -44.8836361
O 11.4794884 23.5296958 -46.0528063
C 13.2402964 25.5207442 -45.0935189
H 13.7974318 24.7585126 -45.6299273
H 12.6747639 26.1075278 -45.8195301
C 14.2367 26.4457212 -44.363787
H 13.6782029 27.2076148 -43.8182901
H 14.8081668 25.8651489 -43.6389009
C 15.2530414 27.1445263 -45.2802328
H 15.9662655 27.6665972 -44.6388995
H 14.7410762 27.8844194 -45.8974409
C 16.0017683 26.1590706 -46.1895801
H 15.4383225 26.0365783 -47.1185131
H 16.0485386 25.1791219 -45.7028254
N 17.3833513 26.6013149 -46.4784672
H 17.4501689 27.5257558 -46.8646949
H 17.8394313 25.9286781 -47.0934372
H 17.9672793 26.5200239 -45.6378488
N 10.2586788 23.4106932 -44.1767312
H 10.1841574 23.7178249 -43.2124954
C 9.2046284 22.5279462 -44.6902692
H 9.4646982 22.2318761 -45.7066837
C 7.8664666 23.2561846 -44.773848
O 7.5173305 24.0732073 -43.9268036
C 9.0728533 21.2239852 -43.8778855
H 8.316634 20.6091786 -44.3642886
C 10.3766276 20.4230278 -43.8853184
H 11.1563743 20.9672999 -43.3545985
H 10.6901088 20.2389319 -44.9118237
H 10.2204614 19.4614051 -43.3955968
C 8.6444678 21.4437578 -42.4225151
H 9.3296493 22.1190396 -41.9159669
H 8.6253233 20.491972 -41.8909169
H 7.6444986 21.8751438 -42.3818422
N 7.0848025 22.9020958 -45.7835561
H 7.4542566 22.2468272 -46.4660422
C 5.688535 23.2901281 -45.9603944
H 5.2592379 23.5642908 -44.9971347
C 4.9368154 22.090787 -46.5162855
O 5.5175932 21.2799915 -47.2402207
C 5.572196 24.4861637 -46.9093264
H 6.1315988 25.3254024 -46.4927753

H 4.5253632 24.7775338 -47.0057321
O 6.0920421 24.166681 -48.1936715
H 6.9019288 23.6412406 -48.069159
N 3.6649475 21.9328174 -46.1572492
H 3.1924113 22.6509743 -45.6290061
C 2.8540294 20.891737 -46.7718369
H 3.2603949 20.7420939 -47.768523
C 1.4232672 21.3469321 -46.9993605
O 0.9655742 22.2461308 -46.2958749
C 2.8723023 19.5311039 -46.0727033
H 1.8998189 19.3593649 -45.6756641
H 2.831625 18.8431272 -46.9167144
C 4.0238134 18.9905664 -45.2169582
C 4.3071869 17.6253821 -45.3860567
H 3.6888653 17.0224533 -46.0309537
C 4.8919789 19.744078 -44.3979068
H 4.7530479 20.7996534 -44.2590226
C 5.3992881 17.0235059 -44.752768
H 5.5957283 15.9801922 -44.9285487
C 5.9619852 19.1264943 -43.7177794
H 6.5828412 19.700868 -43.0462872
C 6.2154413 17.761582 -43.8888187
H 7.0295482 17.2877361 -43.3583586
N 0.7049211 20.6843602 -47.9071269
H 1.1580329 19.9917536 -48.4999263
C -0.7426341 20.8387991 -48.0115702
H -1.0824204 20.9641876 -46.9898499
C -1.5361428 19.617803 -48.4852348
O -0.9924186 18.6595985 -49.0384563
C -1.11455 22.1121096 -48.8003637
H -1.7026334 21.8618073 -49.6844732
H -0.2187602 22.6483505 -49.119513
C -1.9431406 22.9948283 -47.8556361
O -2.8487189 22.4398679 -47.1824651
O -1.4945848 24.1167787 -47.540539
N -2.8489941 19.6723543 -48.2388324
H -3.2118883 20.5386921 -47.8451067
C -3.8093695 18.7281656 -48.8164363
H -3.3973647 17.7249245 -48.7310293
C -3.9734597 19.0752603 -50.2936677
O -4.3973474 20.175993 -50.6415666
C -5.1732061 18.7435119 -48.0992768
H -5.6730347 19.6950951 -48.2825975
C -6.0637773 17.6068255 -48.6193746
H -5.6052813 16.639463 -48.4108445
H -6.2151491 17.7031978 -49.6940242

H -7.0395154 17.6576898 -48.1394967
C -5.0359728 18.5600012 -46.5826229
H -4.4763312 17.6507279 -46.373113
H -6.0235974 18.4938287 -46.1264681
H -4.5200769 19.4177832 -46.1531065
N -3.6215795 18.1436875 -51.1741785
H -3.2879382 17.2507786 -50.8282856
C -3.6115106 18.3903012 -52.6077545
H -4.5005105 18.9612831 -52.879691
H -2.7342712 18.9845956 -52.864576
C -3.5764643 17.1062873 -53.4192378
O -2.760003 16.2192524 -53.1788795
N -4.4347018 17.0516582 -54.4326559
H -5.0732818 17.8204822 -54.5581563
C -4.3864658 16.05468 -55.4956002
H -4.0915978 15.0912157 -55.0883004
C -3.3390088 16.4662952 -56.5310674
O -3.4495113 17.5140222 -57.164842
C -5.7890218 15.9175663 -56.0942179
H -6.1264015 16.9003061 -56.4288134
H -6.4684305 15.5787826 -55.3090581
C -5.884318 14.9468145 -57.2730016
O -4.8383242 14.4918336 -57.7921059
O -7.0322555 14.7422459 -57.7062923
N -2.3237965 15.6294866 -56.7222922
H -2.3348741 14.7515165 -56.2290766
C -1.1959614 15.917138 -57.5968154
H -0.8042434 16.9026035 -57.3349828
C -1.5477935 15.9520276 -59.0940706
O -0.7986989 16.5414193 -59.8719076
C -0.1261968 14.8713849 -57.2991538
H -0.4332321 13.8964363 -57.6832249
H 0.0211865 14.8034629 -56.2217764
H 0.8044678 15.171372 -57.7763393
N -2.6894224 15.3899282 -59.5099429
H -3.3057519 14.9434825 -58.8268714
C -3.1680469 15.5081525 -60.8889374
H -2.2977069 15.4764396 -61.5426236
C -3.8714604 16.8578481 -61.1353749
O -3.8881519 17.3427345 -62.2717246
C -4.0589673 14.2899356 -61.2019821
H -3.5800509 13.3944349 -60.8022451
H -5.0094949 14.4145144 -60.6792547
C -4.3336983 14.054691 -62.7013104
H -4.8066073 14.9365606 -63.1323418
C -3.0607759 13.732287 -63.4923323

H -2.5395826 12.8897156 -63.037984
H -2.3987076 14.5952794 -63.5160859
H -3.3198034 13.4669686 -64.5171777
C -5.2831101 12.8667808 -62.868187
H -4.8269732 11.9632398 -62.4652122
H -5.5106869 12.7195113 -63.9246257
H -6.2135557 13.0676634 -62.3361046
N -4.3438173 17.5317856 -60.0767708
H -4.239743 17.0913676 -59.1689351
C -5.1392953 18.7670931 -60.1508346
H -5.3101043 19.014173 -61.1954728
C -4.4652452 20.0192241 -59.553186
O -4.7247452 21.1197762 -60.036646
C -6.5275032 18.4864972 -59.5562995
H -7.1041324 19.4099779 -59.5149897
H -6.4332717 18.0886791 -58.546576
C -7.2894454 17.5033969 -60.4354308
O -7.5563357 17.7986585 -61.5883171
N -7.5704357 16.2932746 -60.0008698
H -8.0536807 15.6856149 -60.632247
H -7.3300548 15.9419432 -59.0695682
N -3.5291447 19.8666245 -58.6129897
H -3.3778296 18.9328279 -58.2480178
C -2.7161973 20.948246 -58.0426512
H -3.3974747 21.6988151 -57.6398965
C -1.8381347 21.6244684 -59.108357
O -1.3603305 20.9679296 -60.0376461
C -1.8708616 20.3801509 -56.8857777
H -1.2155043 19.5960797 -57.2688558
H -2.5403692 19.9339514 -56.1488536
C -1.0113161 21.4298223 -56.1713507
H -0.2443703 21.7912918 -56.8567282
H -0.5059511 20.9491715 -55.3327881
S -1.9402459 22.8506813 -55.5394919
C -0.5701828 23.8292679 -54.8690719
H -0.9613234 24.74235 -54.4197375
H 0.1193026 24.0926016 -55.6715556
H -0.0410525 23.252321 -54.1098706
N -1.6015408 22.9369568 -58.9811233
H -1.9423219 23.4054305 -58.1486339
C -0.8192637 23.7390643 -59.9395053
H -0.2576772 23.064157 -60.5851844
C 0.2282464 24.5957342 -59.2270807
O 0.004776 25.0706294 -58.1183271
C -1.7509353 24.6002576 -60.822006
H -2.2485006 25.3361505 -60.1875853

H -1.1470927 25.1432575 -61.5512182
C -2.8355023 23.8041089 -61.576147
H -3.4692309 24.5095959 -62.1148217
H -3.4670243 23.293783 -60.85025
C -2.285634 22.7794283 -62.5838831
H -1.469829 22.2166229 -62.1362081
H -1.8727529 23.3160784 -63.440655
N -3.3480238 21.8600974 -63.0454371
H -4.1592275 22.2924263 -63.4494333
C -3.3712553 20.539867 -62.9367282
N -4.3632524 19.8552986 -63.436589
H -4.394121 18.8635073 -63.2050823
H -5.2093213 20.3214733 -63.7056537
N -2.4470361 19.8412767 -62.3456859
H -1.8072108 20.3142319 -61.7116381
H -2.5589807 18.8412342 -62.2636225
N 1.3477479 24.8147152 -59.9086339
H 1.3980397 24.4714973 -60.8561952
C 2.4801481 25.6382775 -59.4817666
H 2.171737 26.310206 -58.6796175
C 2.9540436 26.4820188 -60.6742369
O 2.5493109 26.2124622 -61.8048482
C 3.6101897 24.7291168 -58.9715134
H 3.9556652 24.0994097 -59.7921348
H 4.4519802 25.3525014 -58.6710504
C 3.2414508 23.8502424 -57.790578
C 3.2255769 24.3988016 -56.4940977
H 3.4578802 25.4439975 -56.3432923
C 2.944317 22.484486 -57.9805326
H 2.957988 22.0603603 -58.9749413
C 2.9118273 23.5871558 -55.3886793
H 2.893409 23.9989979 -54.390721
C 2.6376701 21.6672874 -56.8746023
H 2.4174114 20.6212776 -57.0198648
C 2.6196309 22.2218223 -55.5761608
O 2.3179465 21.4526177 -54.4997302
H 2.18435 20.5319836 -54.724826
N 3.8334992 27.4628261 -60.4509498
H 4.2082822 27.584594 -59.5219213
C 4.4939213 28.1620494 -61.5616058
H 3.7309422 28.5488971 -62.2384471
C 5.3882736 27.2020676 -62.3485732
O 6.0176283 26.3114741 -61.7616727
C 5.3217809 29.3477141 -61.0436929
H 6.091829 28.9980095 -60.3569351
H 4.6612574 30.029898 -60.5071673

O 5.9393037 30.0538829 -62.1032955
H 6.7243398 29.5613144 -62.4093461
N 5.5820288 27.4940815 -63.6317522
H 5.0106585 28.1958685 -64.0746187
C 6.7307802 26.9874338 -64.3760478
H 6.6366855 25.9085016 -64.4254029
C 8.028407 27.3558119 -63.6304401
O 8.1187891 28.4400932 -63.0417593
C 6.7479946 27.5450176 -65.8100117
H 7.719706 27.3149662 -66.2505881
H 6.6485008 28.6315395 -65.7731873
C 5.6672195 26.9697377 -66.7358325
O 4.772257 26.2412801 -66.2508656
O 5.7522914 27.2543315 -67.9504335
N 9.0046514 26.4438149 -63.5827792
H 8.8767 25.5757138 -64.0999271
C 10.2928452 26.6834173 -62.9168859
H 10.9628395 25.863104 -63.1596956
H 10.726691 27.5994604 -63.3176861
C 10.2581107 26.8168125 -61.3836782
O 11.1929088 27.3706979 -60.812188
N 9.2068629 26.3463782 -60.7000323
H 8.4713915 25.8971 -61.2280263
C 9.0731089 26.4374344 -59.2283582
H 9.1851045 27.4791121 -58.9258279
C 10.1069491 25.6136815 -58.4432941
O 10.4836519 25.994552 -57.3288833
C 7.6897912 25.9495096 -58.7796666
H 7.6527518 25.9341602 -57.6888603
H 7.5223822 24.9374057 -59.148062
O 6.6580577 26.7898418 -59.2502181
H 6.5249451 26.5892104 -60.1998395
N 10.5092653 24.4553591 -58.9775317
H 10.148396 24.2047745 -59.8925566
C 11.2703098 23.4359545 -58.2511685
H 11.53296 23.831252 -57.2725145
C 12.5826537 23.0539197 -58.9343996
O 12.7237566 23.0892687 -60.153064
C 10.4016771 22.1945978 -58.0069762
H 10.1678104 21.7295215 -58.9665107
H 10.9725225 21.4720044 -57.4234963
C 9.1174901 22.4918067 -57.2616053
C 9.1440574 22.7882535 -55.8861933
H 10.0770687 22.7663086 -55.3446341
C 7.8965917 22.5106125 -57.9559699
H 7.8822044 22.2996273 -59.0114969

C 7.9532069 23.1130515 -55.2127899
H 7.9771022 23.3415634 -54.1554797
C 6.7052395 22.8277978 -57.284616
H 5.7704859 22.841606 -57.8236995
C 6.7337942 23.135257 -55.9133376
H 5.8189192 23.3828901 -55.3936062
N 13.517583 22.5839715 -58.1188466
H 13.3072881 22.5350507 -57.1278463
C 14.8179288 22.0601079 -58.5444263
H 15.0314648 22.3351386 -59.5799209
C 14.8820061 20.5283609 -58.4571595
O 15.7122647 19.8905221 -59.1081171
C 15.861854 22.6970743 -57.6264017
H 15.4958654 22.6849266 -56.6000881
H 16.7664742 22.1004187 -57.6568063
C 16.1785143 24.1454572 -58.0345006
H 16.4935678 24.6931106 -57.143149
H 15.2907343 24.6425045 -58.4312527
C 17.305602 24.1476694 -59.067901
O 17.1190339 23.5958259 -60.1692975
O 18.44763 24.4781666 -58.6847995
N 14.0008482 19.9431131 -57.6415347
H 13.375073 20.5442361 -57.11928
C 13.8557196 18.5135042 -57.3961515
H 14.3000178 17.9505765 -58.2132188
C 12.3562485 18.1809218 -57.2916426
O 11.6055412 18.9392478 -56.6695756
C 14.5613807 18.1437631 -56.0764476
H 13.9703084 18.5856413 -55.2756477
H 14.5331341 17.0604963 -55.9537592
C 16.0075358 18.6382877 -55.8749913
H 16.0774433 19.7022195 -56.0961422
C 16.4272853 18.4705187 -54.4170553
H 16.4803628 17.4129 -54.1558347
H 15.7119752 18.972793 -53.7650525
H 17.3987076 18.9361346 -54.2814031
C 17.0207935 17.8886108 -56.7338335
H 16.9321508 16.814618 -56.5697651
H 18.0321975 18.2073195 -56.4892937
H 16.8337156 18.1120355 -57.7796173
N 11.9268476 17.0254245 -57.802589
H 12.5667476 16.4679935 -58.3580903
C 10.6134922 16.4408486 -57.4727742
H 10.200647 16.9817986 -56.6240596
C 10.8460628 14.9974708 -57.0338666
O 11.4968357 14.2196056 -57.7343886

C 9.584129 16.5453119 -58.623281
H 9.8891 15.8453796 -59.384544
C 9.5544193 17.9798466 -59.2141413
H 9.2019235 18.6905423 -58.4697116
H 10.5636784 18.2729503 -59.500348
C 8.1844474 16.0604285 -58.2049168
H 7.7911132 16.6907992 -57.4099126
H 8.2346799 15.0332518 -57.8434264
H 7.5072373 16.083636 -59.0554359
C 8.7221937 18.1222675 -60.4791129
H 7.6661354 18.1114382 -60.2270789
H 8.9737782 17.3097286 -61.1536164
H 8.9770545 19.0592215 -60.9698104
N 10.3564225 14.6574374 -55.8469035
H 9.7941865 15.3397182 -55.3446681
C 10.5610505 13.361721 -55.2088406
H 11.1527113 12.721799 -55.8623133
C 9.2265463 12.6740554 -54.9607565
O 8.2716464 13.3041168 -54.5053329
C 11.33476 13.5316808 -53.9000093
H 12.3374437 13.8914798 -54.1303004
H 10.8329836 14.2807638 -53.2847405
C 11.4560679 12.2414929 -53.107116
C 12.2584577 11.1893237 -53.5913009
H 12.8093437 11.3079028 -54.5115507
C 12.3285074 9.9712448 -52.892086
H 12.935959 9.1595666 -53.2719667
C 11.6002187 9.80203 -51.7022888
H 11.6597189 8.8679635 -51.1608939
C 10.7885537 10.8448729 -51.2227153
H 10.2173261 10.7141468 -50.3149552
C 10.7115946 12.0603749 -51.9250526
H 10.0647863 12.8469327 -51.5572318
N 9.1602094 11.3777451 -55.2613215
H 9.967209 10.9333375 -55.6892098
C 7.9328766 10.5948167 -55.1189933
H 7.301889 11.0701356 -54.3646456
C 8.255057 9.1831792 -54.6477118
O 9.0891373 8.509012 -55.2435789
C 7.1597415 10.5440188 -56.4471444
H 7.7114147 9.9222563 -57.1494066
C 5.8095002 9.8629732 -56.2113163
H 5.4896513 10.0212756 -55.1882349
H 5.9140703 8.7885214 -56.3956002
H 5.0585275 10.2781186 -56.8705113
C 7.0334598 11.9183157 -57.1168015

H 6.7794035 12.6694729 -56.3779092
H 6.2773547 11.9109892 -57.8926942
H 7.9869524 12.1928344 -57.5734035
N 7.5304832 8.6922801 -53.6430702
H 6.8760449 9.3036927 -53.1726615
C 7.6681163 7.3252057 -53.1358409
H 8.2068826 6.7406371 -53.8801015
C 6.2876119 6.6673452 -52.9862792
O 5.4097723 7.1751835 -52.2984167
C 8.5623854 7.3067255 -51.8731256
H 9.5579647 7.6240205 -52.1890165
C 8.094785 8.2976582 -50.7891014
H 7.0661867 8.0877915 -50.5071326
H 8.1462975 9.3195321 -51.1659953
H 8.7354529 8.2405992 -49.9112774
C 8.6896144 5.865025 -51.3446308
H 7.7397634 5.5693468 -50.9079072
H 8.8984866 5.198318 -52.1818034
C 9.804598 5.6643442 -50.3096542
H 9.5898272 6.2220642 -49.3977556
H 10.7584064 5.9929064 -50.7211156
H 9.8773923 4.6049462 -50.0589894
N 6.0735872 5.5791504 -53.7322323
H 6.8322139 5.3057652 -54.3480851
C 4.8587987 4.7465541 -53.7745885
H 4.9169297 4.1923917 -54.7089597
C 3.516118 5.5057891 -53.8614048
O 2.5191161 5.1118456 -53.2532807
C 4.8607276 3.7088065 -52.6438668
H 4.6254463 4.2273965 -51.7263386
H 4.0580549 2.9937707 -52.8329263
C 6.1573545 2.9237375 -52.4338794
H 5.9667513 2.1123083 -51.7318169
H 6.9089625 3.5794581 -51.9982039
C 6.7097361 2.3381834 -53.7201912
O 7.8204849 2.6367907 -54.124086
N 5.9596121 1.5408023 -54.4434662
H 6.3860314 1.1601171 -55.2659575
H 4.9443976 1.5731702 -54.3602218
N 3.4922299 6.609794 -54.6151327
H 4.3734201 6.9183661 -54.9925454
C 2.3102518 7.4619524 -54.7883182
H 1.523502 7.1422596 -54.1042785
C 1.7272705 7.3825057 -56.209906
O 0.5213259 7.2247492 -56.3878293
C 2.7044591 8.890572 -54.3960494

H 3.4554001 9.2689719 -55.0839756
H 3.1257726 8.8895343 -53.3873539
S 1.267988 9.9956678 -54.455393
H 1.9581013 11.1116626 -54.1764562
N 2.5785643 7.3956426 -57.2408166
H 3.5667076 7.4519611 -57.0465178
C 2.1412431 7.4998195 -58.6453044
H 1.3706166 8.2684934 -58.6961616
C 1.455654 6.224998 -59.1639819
O 0.6846533 6.2812467 -60.1185169
C 3.3271499 7.9979964 -59.498981
H 4.2271993 7.4604443 -59.1935633
C 3.5120954 9.5030321 -59.2234801
H 2.7433385 10.070109 -59.7525158
H 3.3888159 9.7067547 -58.1618482
C 3.1360484 7.7830679 -61.0060698
H 2.1735433 8.1823495 -61.3276595
H 3.2063335 6.7193164 -61.2171017
H 3.9295564 8.2650958 -61.5743967
C 4.8911054 10.0097662 -59.6504475
H 5.0099305 9.9628237 -60.7314375
H 5.6735898 9.4236341 -59.1685517
H 4.9797716 11.0455733 -59.3443017
N 1.5717582 5.1041732 -58.4564899
H 2.1827045 5.1206104 -57.6529454
C 0.7328439 3.9149034 -58.651388
H 0.8947388 3.5462889 -59.6639284
C -0.7823536 4.1812482 -58.5331897
O -1.5480582 3.5308124 -59.2486143
C 1.1982313 2.8047712 -57.6992863
H 0.5310809 1.9524817 -57.8022436
H 2.1807305 2.4791663 -58.0396875
C 1.2976814 3.1948038 -56.2119626
H 0.3012642 3.1688227 -55.7657597
H 1.6944627 4.205053 -56.0979221
C 2.2384522 2.2254211 -55.4939914
O 3.4720441 2.4194285 -55.590334
O 1.7426194 1.2099946 -54.9540701
N -1.2002262 5.2068174 -57.7761104
H -0.5129244 5.7123035 -57.2254629
C -2.598939 5.6605511 -57.6700557
H -3.2650489 4.7962077 -57.7183515
C -3.0249164 6.6303356 -58.7861373
O -4.212663 6.7435325 -59.0897442
C -2.8084363 6.3407486 -56.3074555
H -3.8525083 6.6461592 -56.2269157

H -2.1951503 7.2411331 -56.2540386
C -2.494111 5.4648105 -55.1222584
N -3.2748369 4.3995001 -54.6813414
C -1.402258 5.5724542 -54.3123075
H -0.6083186 6.2963122 -54.4092074
C -2.6439835 3.8919381 -53.61534
H -2.9894624 3.0311948 -53.0587202
N -1.5076655 4.5702295 -53.3754836
H -0.8044458 4.3204002 -52.6953078
N -2.0794667 7.3391896 -59.4136958
H -1.1158875 7.1556197 -59.1719981
C -2.3616997 8.3287098 -60.4667779
H -3.2276859 8.9173499 -60.1602128
C -2.7464924 7.5875773 -61.7548098
O -2.001621 6.7304906 -62.2333305
C -1.169923 9.3091347 -60.6337605
H -0.2399088 8.7535995 -60.5132491
C -1.1908147 10.461597 -59.5993055
H -0.3002541 11.0742561 -59.749503
H -2.0610367 11.0934907 -59.7853356
C -1.1248122 9.968303 -62.0256172
H -2.0519598 10.5058078 -62.2273489
H -0.9656117 9.221869 -62.8030621
H -0.2929621 10.6711498 -62.0761009
C -1.2165483 10.0459058 -58.1225757
H -0.4028053 9.3553923 -57.9105932
H -2.1699243 9.5788426 -57.8756608
H -1.0975652 10.9321865 -57.4986943
N -3.9191623 7.9002296 -62.3131057
H -4.4943561 8.5906527 -61.8567087
C -4.4811881 7.1647627 -63.4544377
H -4.2521094 6.1048541 -63.3300885
C -3.8496852 7.5951677 -64.7889643
O -3.2727482 6.7640869 -65.4867586
C -6.0132403 7.3088003 -63.4796484
H -6.2694267 8.3537053 -63.6622205
H -6.4039576 6.7184002 -64.3096062
C -6.7166646 6.8608593 -62.1861027
H -7.7887787 7.0203155 -62.3018223
H -6.3872767 7.4812947 -61.3521355
C -6.4970237 5.3889815 -61.8459751
O -6.9995508 4.4963847 -62.5114945
N -5.7572628 5.0902401 -60.8042215
H -5.833909 4.14822 -60.4307188
H -5.3482376 5.8097848 -60.2156199
N -3.8569681 8.8981386 -65.0946658

H -4.3202539 9.5439063 -64.4785248
C -3.1401507 9.4583446 -66.2477274
H -3.0411782 8.6833109 -67.0107969
C -1.7236527 9.8698317 -65.8282051
O -1.4618108 10.9868417 -65.364135
C -3.9305454 10.5951731 -66.9132344
H -4.0322571 11.4261952 -66.2131145
H -4.9304066 10.2295566 -67.1538309
C -3.2659619 11.0967388 -68.2087631
O -2.0957811 10.7292587 -68.4770125
O -3.9030893 11.919111 -68.8986723
N -0.803794 8.9111941 -65.9612056
H -1.116004 8.0117478 -66.302686
C 0.6148988 9.1220086 -65.6716635
H 0.701243 9.6227611 -64.7065084
C 1.2656105 10.0741342 -66.6689175
O 2.098196 10.8648421 -66.2441968
C 1.3529267 7.7757301 -65.5926566
H 1.1126767 7.1689474 -66.4679944
H 2.4297305 7.955249 -65.5762918
C 0.9574017 7.0366377 -64.3062496
H 1.2856483 7.6316669 -63.4540904
H -0.1280695 6.9419732 -64.2625537
C 1.5717334 5.633152 -64.2128231
H 2.6407791 5.6991469 -64.4166036
H 1.1128656 4.9710833 -64.9484462
C 1.3819517 5.0770151 -62.7971826
H 1.7874912 5.8138986 -62.1066302
H 1.9538723 4.1524789 -62.6816544
N -0.0351696 4.8430507 -62.4374898
H -0.6226182 5.656999 -62.6128357
H -0.1159741 4.6848988 -61.4413045
H -0.410129 4.0257107 -62.9061462
N 0.857613 10.1083443 -67.9375912
H 0.0306158 9.5949737 -68.2147783
C 1.4627276 11.0419099 -68.8908006
H 2.5495178 10.9536548 -68.8456466
C 1.0955327 12.4971915 -68.5945729
O 1.9570158 13.368256 -68.6695282
C 1.0240044 10.7936445 -70.3246517
H -0.0383867 11.0268319 -70.4208665
H 1.5987591 11.5133366 -70.9013528
C 1.2870139 9.4334093 -70.9591885
H 0.6207275 8.6875652 -70.5204886
H 2.321262 9.1555211 -70.7495981
C 1.0670261 9.5267931 -72.4818624

O 1.5837226 8.6346019 -73.1863309
O 0.449561 10.5278113 -72.9382914
N -0.1651418 12.7909911 -68.2489103
H -0.8538143 12.0316837 -68.2624408
C -0.597046 14.1400191 -67.8500139
H -0.3606733 14.8368522 -68.6493023
C 0.1788132 14.6128788 -66.6331437
O 0.6875716 15.7306213 -66.6664212
C -2.1126775 14.1466946 -67.5941196
H -2.6227997 14.0571914 -68.5530931
H -2.3576696 13.272807 -66.9910062
C -2.6896842 15.3612165 -66.8459386
H -3.7652708 15.2087699 -66.7430743
H -2.2667664 15.3935762 -65.8416272
C -2.4521123 16.7168604 -67.5216241
H -2.901966 16.7154243 -68.5155877
H -1.3820194 16.9074539 -67.6060141
C -3.0951271 17.8042082 -66.6541248
H -2.7560098 17.6658543 -65.6223159
H -4.1813943 17.6762267 -66.6795216
N -2.71647 19.159312 -67.1134731
H -1.700728 19.2804625 -67.0277463
H -3.1123092 19.8764223 -66.5258363
H -2.9676407 19.3212056 -68.075571
N 0.3204954 13.766832 -65.6119406
H -0.1301998 12.8600258 -65.6626692
C 1.1372004 14.0988161 -64.4489276
H 0.7681291 15.0311553 -64.0186157
C 2.6010132 14.3488103 -64.8440332
O 3.1203401 15.4231067 -64.5515805
C 1.0048208 13.0052398 -63.3833035
H -0.0274297 12.9715737 -63.0338871
H 1.2416141 12.0362933 -63.8251743
C 1.9196494 13.2544928 -62.2026704
C 1.5902525 14.2330469 -61.2449909
H 0.6507624 14.7645696 -61.3091331
C 2.5054578 14.5589645 -60.2283914
H 2.2645 15.3361693 -59.5185997
C 3.7477584 13.9046379 -60.163313
H 4.4610224 14.1635952 -59.3939693
C 4.075634 12.9324819 -61.1222156
H 5.0420339 12.4538226 -61.0965527
C 3.1634757 12.6011592 -62.1355357
H 3.4384687 11.8742898 -62.8856969
N 3.2219607 13.4431795 -65.6129092
H 2.7236004 12.5935768 -65.8627977

C 4.6285742 13.5613066 -66.0272132
H 5.2190656 13.6913037 -65.136371
C 4.8554892 14.8103973 -66.9071792
O 5.7538651 15.5982238 -66.6248052
C 5.1469566 12.2423806 -66.6375441
H 4.5264298 12.0049949 -67.503892
C 6.6112232 12.3569321 -67.0848273
H 7.294004 12.1712733 -66.2581363
H 6.8394987 13.3368053 -67.4929083
H 6.78462 11.6171695 -67.8603109
C 5.0988351 11.0728698 -65.6156212
H 5.8917595 11.1882859 -64.8758281
H 4.1709873 11.0890031 -65.0529447
C 5.2309691 9.6913749 -66.2729081
H 6.195366 9.5950969 -66.7695518
H 4.4343643 9.5515472 -67.0036455
H 5.1519131 8.9179254 -65.5089451
N 3.9636351 15.1292163 -67.8520424
H 3.2265652 14.4600239 -68.0547949
C 4.0218868 16.3871815 -68.6226211
H 5.0042765 16.4596209 -69.0901891
C 3.8797531 17.6361744 -67.751329
O 4.3853571 18.6900604 -68.1393617
C 2.9423549 16.3991231 -69.7126516
H 1.9845357 16.1193492 -69.2705717
H 2.8543825 17.4158555 -70.0989016
C 3.2701847 15.4763868 -70.8965743
H 3.2834725 14.434007 -70.5792135
H 4.2600221 15.7274747 -71.2810722
C 2.2462379 15.6651832 -72.0260354
H 2.5661878 15.0769262 -72.8877034
H 2.2537104 16.7118842 -72.3344896
N 0.873631 15.2917612 -71.613203
H 0.3736673 15.9475456 -71.0440993
C 0.2606047 14.1451865 -71.8617678
N 0.8324717 13.2109762 -72.562001
H 0.4215544 12.2750905 -72.7048818
H 1.7817286 13.3170273 -72.8573687
N -0.9394563 13.9204432 -71.4047869
H -1.4254517 14.5628763 -70.815964
H -1.3235025 12.9947997 -71.5411287
N 3.1468849 17.5726836 -66.640555
H 2.7147858 16.6893069 -66.3908726
C 3.0422072 18.6861717 -65.6891141
H 2.9536692 19.6115987 -66.2544648
C 4.3172776 18.8331359 -64.8542892

O 4.818539 19.9522103 -64.7316767
C 1.7891423 18.5422215 -64.8007602
H 1.0555546 17.9044931 -65.2924164
H 2.0612306 18.0606817 -63.8602092
C 1.1195113 19.8957002 -64.5028273
H 1.8473703 20.5638183 -64.0369374
H 0.3048766 19.7346467 -63.7929669
C 0.5491297 20.5310302 -65.7790337
O -0.2282403 19.8629453 -66.4986209
O 0.9457575 21.6510811 -66.1609826
N 4.94132 17.725144 -64.4234338
H 4.4998686 16.8199543 -64.5638902
C 6.2140599 17.8019982 -63.6909489
H 6.0524835 18.5815138 -62.9562327
C 7.3890477 18.3481159 -64.518059
O 8.2732855 18.9822581 -63.9445966
C 6.5604161 16.5360761 -62.8697259
H 7.6036421 16.612572 -62.5826921
C 5.687129 16.5745079 -61.5928064
H 4.6438896 16.3754152 -61.8451636
H 5.7360653 17.5498829 -61.1123979
H 6.0256541 15.83006 -60.8757306
C 6.3611885 15.1622226 -63.5084856
H 5.298497 14.9474787 -63.5301493
H 6.74844 15.1843201 -64.5161972
C 7.0610066 14.0140301 -62.7739185
H 6.6684855 13.8981371 -61.7667884
H 8.122155 14.2216184 -62.7084251
H 6.9110273 13.0840706 -63.3228865
N 7.30301 18.2991724 -65.852446
H 6.594574 17.6978391 -66.2541671
C 8.2430857 18.9945263 -66.7521705
H 9.256973 18.7089679 -66.4899572
C 8.1690656 20.51075 -66.5818082
O 9.1990222 21.1611148 -66.460308
C 7.977772 18.6399987 -68.229651
H 6.9892592 19.0220523 -68.484297
C 8.9845327 19.2914653 -69.1874602
H 10.0014391 18.9979181 -68.9194354
H 8.9126701 20.3778295 -69.1491886
H 8.7886603 18.966429 -70.2109033
C 7.96589 17.1408008 -68.5243393
H 8.943109 16.8146886 -68.8779901
H 7.2357508 16.9438971 -69.3046957
H 7.7171778 16.551408 -67.6508645
N 6.9615765 21.0893303 -66.532496

H 6.1488449 20.486938 -66.5022801
C 6.780615 22.5474602 -66.4135413
H 7.4330732 23.0541204 -67.1286945
C 7.2202667 23.0717325 -65.0569848
O 7.912679 24.0815246 -64.9821131
C 5.3159302 22.9327156 -66.6760503
H 4.6831159 22.049465 -66.5844177
H 4.9780527 23.653042 -65.9286919
C 5.1585947 23.5769347 -68.0589042
H 5.5905341 22.9257093 -68.8152424
H 5.7275276 24.5057287 -68.0837541
C 3.6905468 23.8966561 -68.3952165
H 3.6380314 24.2920585 -69.4108383
H 3.3606373 24.6917945 -67.7218206
N 2.7656957 22.7454938 -68.2527494
H 2.0283473 22.845749 -67.5579178
C 2.88091 21.5274568 -68.7454514
N 3.8693124 21.140862 -69.4991559
H 4.0949378 20.1565639 -69.4957941
H 4.5709032 21.8214323 -69.7161739
N 1.9903342 20.6334323 -68.4586807
H 1.2955849 20.8980357 -67.7519289
H 2.1477213 19.6774308 -68.6872359
N 6.8385406 22.3850171 -63.9793491
H 6.2678414 21.5594362 -64.1188139
C 7.0762139 22.8882716 -62.6154744
H 6.7536171 23.9302454 -62.5989197
C 8.5529194 22.901266 -62.209056
O 8.8936518 23.5075295 -61.1960446
C 6.2302789 22.1490035 -61.5659052
H 6.3002834 22.7158685 -60.6375336
C 4.7406745 22.1370212 -61.9312552
H 4.5608851 21.5072413 -62.7992222
H 4.4104188 23.1524728 -62.155835
H 4.1599278 21.7513496 -61.0951958
C 6.7187884 20.740197 -61.2639461
H 6.946144 20.2365441 -62.1944734
H 5.9697426 20.1820863 -60.7022476
H 7.6299397 20.7966649 -60.6732389
N 9.4330305 22.2598839 -62.9727844
H 9.1190088 21.8290885 -63.831774
C 10.8586508 22.2157185 -62.6987607
H 11.0205601 22.3896924 -61.6385775
C 11.6587618 23.2613323 -63.4943908
O 11.250341 23.686733 -64.571033
C 11.3162717 20.8126336 -63.009183

H 11.2222436 20.673128 -64.0870845
H 10.6948542 20.0962774 -62.4706286
H 12.34619 20.6925253 -62.698318
N 12.8048385 23.6814376 -62.9622398
H 13.0708624 23.3246349 -62.0518115
C 13.7495271 24.5578181 -63.6509041
H 13.1778 25.2862108 -64.2238724
C 14.6719354 23.7794981 -64.6223379
O 14.975169 22.6149417 -64.3663321
C 14.5534374 25.3133525 -62.584314
H 15.1349368 24.6050227 -61.9900518
H 13.8784704 25.8645982 -61.9278724
H 15.2402197 26.0126728 -63.0612602
N 15.2562672 24.4346856 -65.6428965
C 16.3457623 23.8565942 -66.4329207
H 15.9645131 23.0084207 -66.9972777
C 17.5084646 23.3942187 -65.5437926
O 17.9666931 24.1420647 -64.6815619
C 16.7604415 24.9539896 -67.41675
H 17.5396302 25.5804047 -66.9808329
H 17.0916402 24.5356973 -68.368371
C 15.4800201 25.7739039 -67.5748403
H 14.8102338 25.2746548 -68.2775496
H 15.6876092 26.7949285 -67.8961341
C 14.8755827 25.7311526 -66.1737705
H 13.7917232 25.8368128 -66.2352599
H 15.3069372 26.5190763 -65.555338
N 17.9761557 22.1537945 -65.7184936
H 17.5578649 21.5750566 -66.4370437
C 18.9691881 21.5330127 -64.828799
H 19.4528712 20.7173447 -65.3653614
H 19.7268063 22.271958 -64.5676816
C 18.3945524 20.9579177 -63.5231969
O 19.1431088 20.4234825 -62.6929978
N 17.0763422 20.9995032 -63.325819
H 16.4871733 21.5126475 -63.9742723
C 16.4200515 20.282351 -62.2438022
H 16.8992307 20.5608404 -61.3042546
C 16.5430015 18.7673145 -62.4295642
O 16.5979498 18.2344241 -63.5433115
C 14.9511917 20.6876016 -62.1594518
H 14.4759675 20.4721613 -63.1160417
H 14.876111 21.7519627 -61.935609
H 14.4412873 20.1255796 -61.3775577
N 16.4551063 18.0541021 -61.3125826
H 16.2855776 18.5690471 -60.4511779

C 16.3207142 16.6066863 -61.2672556
H 16.667364 16.1436285 -62.1925943
C 14.8430527 16.4876811 -61.1714432
O 14.2703098 16.4918358 -60.0760996
C 17.1025975 16.0055349 -60.0967373
H 16.7267382 16.4254048 -59.1674263
H 16.9332383 14.927969 -60.0696969
C 18.6151246 16.2789271 -60.2035818
H 18.7934009 17.3000794 -60.5397025
H 19.0498173 15.6006874 -60.9384452
C 19.3309473 16.1098317 -58.8681021
O 19.8209138 17.063498 -58.2767724
N 19.4280626 14.9016131 -58.3528361
H 19.9160454 14.8206799 -57.47876
H 19.0269415 14.1074667 -58.8221429
N 14.2900884 16.7928196 -62.3621341
H 14.9082487 16.9578584 -63.1449686
C 12.8660165 16.8921841 -62.5786515
H 12.50313 17.8129391 -62.118628
C 12.3945187 16.8195691 -64.0583579
H 11.5182164 16.1806944 -64.1205188
C 11.927267 18.2080647 -64.4477732
H 12.7140434 18.9289267 -64.2293846
H 11.0102521 18.4103486 -63.8975376
H 11.6871596 18.2879955 -65.5028421
C 13.3973882 16.3601632 -65.1198995
H 13.1242213 16.743578 -66.1014841
H 14.3632553 16.7589904 -64.8585091
C 13.4802412 14.8541873 -65.2635705
H 12.5137973 14.4524802 -65.5470749
H 13.7913259 14.4259101 -64.3196536
H 14.1809111 14.6061911 -66.0551717
C 12.2832975 15.764129 -61.7971147
O 11.2494559 16.0014477 -61.1925983
N 12.9953431 14.6160479 -61.6922139
H 13.8487719 14.4634771 -62.2140222
C 12.4409562 13.5648977 -60.9104669
H 11.9661348 14.0429408 -60.056893
C 13.3950247 12.5287825 -60.3251963
O 14.3898634 12.1321749 -60.9337501
C 11.3641755 12.9163869 -61.7760491
H 11.69877 11.9263678 -62.0871113
C 10.1448767 12.7675339 -60.8818847
H 9.7797381 13.7482455 -60.5753726
H 10.389334 12.200937 -59.985975
H 9.3515675 12.2659959 -61.4236928

C 10.8142608 13.647437 -63.0167973
H 10.2651247 14.5505195 -62.7809551
H 10.1246692 12.9691349 -63.5158578
H 11.6027534 13.8396563 -63.7244018
N 12.992268 12.0624941 -59.1407394
H 12.2290494 12.5696896 -58.7005972
C 13.3517678 10.7991302 -58.4875107
H 13.8691189 10.1454328 -59.1901295
C 12.022295 10.1409558 -58.0722694
O 11.2124517 10.7801555 -57.3905926
C 14.2732964 11.0619765 -57.2714117
H 13.7556098 11.7467959 -56.5983282
C 14.5539489 9.7552096 -56.5040559
H 15.098085 9.0547942 -57.1393494
H 13.6255968 9.2720965 -56.1977078
H 15.1326487 9.9473315 -55.6028041
C 15.5967541 11.7351388 -57.7104154
H 16.1322448 11.0671286 -58.3834904
H 15.3704105 12.6510699 -58.2557262
C 16.5299331 12.1281646 -56.5575202
H 16.9353557 11.2398739 -56.0737671
H 15.9862613 12.7290053 -55.8275692
H 17.3598219 12.7148318 -56.9523203
N 11.7580335 8.9046503 -58.5018071
H 12.4971308 8.4049395 -59.0006782
C 10.4983895 8.1867901 -58.2471085
H 9.9675434 8.6814518 -57.4396159
C 10.7986262 6.7492106 -57.8252037
O 11.102295 5.9443846 -58.7003625
C 9.5576584 8.1569015 -59.488233
H 10.0657489 7.6118162 -60.2850846
C 9.2280089 9.5527545 -60.0531933
H 8.7489912 10.1650888 -59.2892799
H 10.1684049 10.018405 -60.3410172
C 8.2538598 7.3944316 -59.1507915
H 7.707404 7.913199 -58.3643917
H 8.4738782 6.3822778 -58.8197286
H 7.6197814 7.2880813 -60.0276344
C 8.3419087 9.5094014 -61.311817
H 7.3388524 9.1658198 -61.0701747
H 8.7868381 8.8466241 -62.055071
H 8.2449684 10.4990386 -61.7479271
N 10.3778145 6.3647875 -56.6173089
H 10.0867171 7.0875804 -55.969359
C 10.0771669 4.965996 -56.2829504
H 10.6161371 4.3342597 -56.978693

C 8.5735844 4.707958 -56.4325046
O 7.7547345 5.5273144 -56.0055968
C 10.6004787 4.5910992 -54.8968489
H 10.1061673 5.1945444 -54.1366728
H 11.6735419 4.7889482 -54.8607758
O 10.3828632 3.2140938 -54.6418497
H 9.4472056 3.0366627 -54.4338089
N 8.1607001 3.6309068 -57.1055167
H 8.8769937 2.9804449 -57.4273788
C 6.7371884 3.300688 -57.3620578
H 6.1596131 3.5083897 -56.4621738
C 6.5491433 1.8064488 -57.6597888
O 7.4787363 1.1227106 -58.0834072
C 6.1822375 4.201798 -58.4902398
H 6.9381585 4.3099942 -59.2694911
C 4.8723056 3.766805 -59.1432156
H 4.1171477 3.6139869 -58.3752925
H 5.0235349 2.8460121 -59.7068042
H 4.5357146 4.5394803 -59.8340945
O 5.8902408 5.4700017 -57.9372644
H 6.6083844 5.6589023 -57.3062582
N 5.3715908 1.2506011 -57.3572631
H 4.6553821 1.8125038 -56.8983762
C 5.0533527 -0.1448405 -57.6414244
H 5.9175855 -0.7539901 -57.3692877
C 4.7686066 -0.3651727 -59.1348527
O 3.9382459 0.3149866 -59.7426248
C 3.8790949 -0.6020956 -56.7719286
H 2.9852171 -0.0491227 -57.0590648
H 4.0992285 -0.4156114 -55.7198245
H 3.7121333 -1.6687299 -56.9224242
N 5.384449 -1.4132231 -59.6710028
H 5.9615485 -1.978817 -59.0585404
C 5.274073 -1.8707675 -61.0481607
H 4.52063 -1.2819514 -61.5709594
C 4.819635 -3.3368143 -61.111532
O 4.71484 -4.0327033 -60.0985856
C 6.6290324 -1.6468094 -61.7413333
H 6.7247861 -2.2962304 -62.6122492
H 7.4480343 -1.8571985 -61.0513851
S 6.7161264 0.0744788 -62.2988248
H 5.7073454 -0.0156779 -63.1767116
N 4.5804398 -3.8163232 -62.3306187
H 4.6770828 -3.1691214 -63.1064769
C 4.346574 -5.2294842 -62.6551505
H 4.459447 -5.8264837 -61.7510928

C 5.4032849 -5.7429896 -63.6329606
O 6.1897972 -4.9572822 -64.1649364
C 2.9059161 -5.4309908 -63.1662228
H 2.7672556 -6.4849372 -63.4143421
H 2.2061259 -5.1920836 -62.3647088
C 2.5629917 -4.5759187 -64.3962885
H 3.3538419 -4.6507815 -65.1381943
H 2.4774119 -3.5317996 -64.1082129
C 1.239358 -5.0199314 -65.0200982
H 0.4409992 -4.8935195 -64.2851521
H 1.2911458 -6.0804497 -65.2787751
N 0.9258565 -4.2126199 -66.212061
H 0.4081314 -3.3590145 -66.0351975
C 1.2259418 -4.5210722 -67.4631425
N 1.9688723 -5.5468659 -67.7776155
H 2.1152903 -5.8721481 -68.7219343
H 2.3570204 -6.1389378 -67.0480665
N 0.7801215 -3.779182 -68.4391839
H 0.1596895 -3.0162654 -68.2328983
H 0.9754474 -4.0537471 -69.3841574
N 5.3899017 -7.0384504 -63.929528
H 4.7383203 -7.6344439 -63.4363782
C 6.0942102 -7.5596355 -65.1036889
H 7.0977374 -7.1300138 -65.1006469
C 5.4232967 -7.0925274 -66.4226341
O 4.3354509 -6.5106139 -66.4170726
C 6.2677109 -9.0838745 -64.9594063
H 6.6936781 -9.2945219 -63.9792089
H 6.9785677 -9.4402086 -65.7029615
C 4.9912565 -9.885612 -65.1365704
O 4.2025192 -9.6370408 -66.0222303
N 4.7808317 -10.9165378 -64.3629691
H 3.901636 -11.390675 -64.4666077
H 5.4341878 -11.1196392 -63.6117621
N 6.087026 -7.301176 -67.5654737
H 6.97492 -7.7768733 -67.5279906
C 5.5686416 -6.8960675 -68.8824775
H 5.3268371 -5.8354456 -68.847767
C 4.2717675 -7.636028 -69.2757134
O 3.3960169 -7.0361793 -69.9063768
C 6.6966829 -7.1057223 -69.9118019
H 7.5465829 -6.4850685 -69.622889
H 7.0144693 -8.1489414 -69.8602459
C 6.3428838 -6.7968035 -71.3816833
H 5.5274226 -7.4439008 -71.7033055
C 5.9359276 -5.3387014 -71.5982807

H 6.7312629 -4.6712647 -71.263949
H 5.0185384 -5.1252485 -71.0539862
H 5.7487945 -5.1681951 -72.6580208
C 7.5553389 -7.0841982 -72.2684394
H 8.3866897 -6.4341584 -71.9947723
H 7.2949337 -6.9136733 -73.313153
H 7.8578121 -8.1251123 -72.1530906
N 4.1334798 -8.8979656 -68.8636108
H 4.8250579 -9.2749777 -68.2335885
C 3.0282568 -9.7990911 -69.2066128
H 2.994302 -9.8630138 -70.2915907
C 1.6602333 -9.3119172 -68.6927965
O 1.5945623 -8.516802 -67.7502102
C 3.3072741 -11.2092038 -68.6610331
H 2.5364612 -11.8929918 -69.0166306
H 3.2739869 -11.2045757 -67.5732222
O 4.5717364 -11.6737482 -69.1032695
H 4.7435367 -12.5260357 -68.6915725
N 0.5433211 -9.7783737 -69.2905212
C -0.7985269 -9.4881579 -68.7920022
H -0.974344 -8.4156383 -68.878059
C -0.9754439 -9.9212677 -67.3331253
O -0.527133 -10.9988821 -66.9390965
C -1.7659872 -10.2366409 -69.7171588
H -1.976375 -11.2318292 -69.3196156
H -2.693753 -9.6827823 -69.8650191
C 0.4554123 -10.5666366 -70.5124142
H 1.1583618 -10.2258453 -71.2721388
H 0.6242898 -11.6193873 -70.280253
C -0.973233 -10.3730318 -71.0152853
H -1.0393598 -9.4455678 -71.586019
H -1.3115491 -11.2200922 -71.6130074
N -1.6717958 -9.0991233 -66.547554
H -1.9774122 -8.2113845 -66.9104634
C -1.9426282 -9.3712957 -65.1363939
H -0.9869589 -9.4906116 -64.6280691
C -2.7599268 -10.6578647 -64.9656591
O -3.8400503 -10.7898855 -65.5414905
C -2.701309 -8.1901542 -64.5041285
H -2.7718945 -8.3380187 -63.4261289
H -3.7101012 -8.1606276 -64.9151862
O -2.0772418 -6.945159 -64.7763668
H -2.4140318 -6.2669389 -64.1668781
N -2.292544 -11.5662814 -64.1043462
H -1.3627723 -11.44079 -63.7365719
C -3.0207712 -12.7921241 -63.7457168

H -3.2244875 -13.323451 -64.6781217
C -4.3966937 -12.4684447 -63.1343671
O -4.6230048 -11.3825377 -62.579092
C -2.1096583 -13.72106 -62.9038127
H -1.910148 -14.6234283 -63.4834728
H -1.1418491 -13.2284864 -62.8004951
C -2.5148567 -14.1252411 -61.4840284
H -2.7946617 -13.2267165 -60.9280213
H -1.6416138 -14.553518 -60.9864657
C -3.6505368 -15.1492933 -61.4302847
O -4.2452488 -15.22482 -60.332582
O -4.0275463 -15.7087837 -62.4791995
N -5.3275999 -13.4242844 -63.2208885
H -5.0081732 -14.3620026 -63.4746751
C -6.7373853 -13.2558538 -62.8708644
H -7.1275333 -12.4809128 -63.5271843
C -6.946757 -12.7679238 -61.4410492
O -7.9117529 -12.0322895 -61.2615722
C -7.4812086 -14.5788163 -63.1367407
H -7.2969487 -14.8900304 -64.1668939
H -7.0568127 -15.3459934 -62.4846586
C -8.9995942 -14.5392484 -62.9048429
H -9.3885239 -15.5543231 -62.9959171
H -9.1947796 -14.2219061 -61.8805407
C -9.7553918 -13.6445641 -63.9063237
H -10.1326223 -14.2766743 -64.7127237
H -9.0952164 -12.9031723 -64.3565096
C -10.9343519 -12.935928 -63.2296966
H -11.4774207 -13.6713244 -62.6279936
H -11.6147183 -12.563259 -64.0023007
N -10.4659423 -11.8111498 -62.3804404
H -9.7675482 -12.1340372 -61.710333
H -11.2343898 -11.3862407 -61.8810379
H -10.0067443 -11.1094506 -62.9439105
N -6.0816248 -13.0886981 -60.4718269
H -5.3969747 -13.8110332 -60.6825751
C -6.0968226 -12.5397581 -59.1017552
H -7.1172031 -12.2357542 -58.8791371
C -5.2212816 -11.2922417 -58.8815129
O -5.4720267 -10.5585664 -57.9262851
C -5.7549487 -13.6167123 -58.0640669
H -6.3952429 -14.4869167 -58.2181221
H -5.9482444 -13.2182487 -57.0671658
O -4.3992676 -14.0098058 -58.1432093
H -4.343102 -14.5866227 -58.960205
N -4.2784466 -10.9636136 -59.7796263

H -4.191226 -11.5356199 -60.6116894
C -3.5645761 -9.675878 -59.7459851
H -3.2982635 -9.4612025 -58.710529
C -4.4764878 -8.5391038 -60.2170946
O -4.6424323 -7.5619692 -59.492832
C -2.2637066 -9.7208699 -60.5782299
H -1.5860132 -10.4576694 -60.1459091
H -2.5060962 -10.0411425 -61.5913691
C -1.54592 -8.348949 -60.6594275
H -2.2252017 -7.6004366 -61.0640231
C -1.049715 -7.869789 -59.2926931
H -0.3925492 -8.6162577 -58.8481829
H -1.895953 -7.6778135 -58.6345231
H -0.5004847 -6.9352805 -59.4146506
C -0.3461376 -8.4080241 -61.5969519
H 0.3865714 -9.1103831 -61.226032
H 0.1108929 -7.4212003 -61.6794127
H -0.6523785 -8.7230191 -62.5882788
N -5.1169134 -8.689943 -61.3849318
H -4.9457027 -9.5459942 -61.90868
C -6.0412299 -7.6860189 -61.9416269
H -5.4551489 -6.7857532 -62.1437637
C -7.1240799 -7.231914 -60.9375162
O -7.2297424 -6.0346225 -60.7273625
C -6.5934143 -8.1928127 -63.2907024
H -5.7661369 -8.2380254 -63.9984981
H -6.9541082 -9.2147834 -63.16253
C -7.7329938 -7.3712305 -63.9310273
H -7.8971692 -7.7500898 -64.9408023
H -8.6442331 -7.5659483 -63.3636415
C -7.5575872 -5.8442729 -64.0094301
H -7.4980538 -5.4352241 -63.0077227
H -8.4615509 -5.4163366 -64.4482493
C -6.3614685 -5.3390559 -64.8205827
H -6.662531 -5.2804452 -65.8695286
H -5.5195257 -6.0309988 -64.7300999
N -5.9571078 -4.0034871 -64.3263948
H -6.7679802 -3.395136 -64.17414
H -5.2630821 -3.5587778 -64.9038722
H -5.4837897 -4.0957187 -63.4295262
N -7.871645 -8.0931999 -60.2239082
C -8.873936 -7.6616363 -59.2573345
H -9.5234761 -6.9139405 -59.7159941
C -8.2649088 -7.0320422 -57.9992117
O -8.9337415 -6.198365 -57.3998078
C -9.7099499 -8.8997193 -58.9176722

H -10.0869565 -8.8805224 -57.8938592
H -10.5345133 -8.9914458 -59.6264198
C -7.9351771 -9.5147877 -60.3439147
H -8.4774898 -9.7379089 -61.262144
H -6.9490279 -9.9651637 -60.3563997
C -8.720291 -10.0246495 -59.1344312
H -9.2281338 -10.9685741 -59.3297095
H -8.0659944 -10.10908 -58.2666461
N -7.021786 -7.3606245 -57.6043131
H -6.4998668 -8.0322734 -58.1535313
C -6.3056253 -6.6353359 -56.5348897
H -6.9728801 -6.5111878 -55.6824392
C -5.9322881 -5.2270172 -57.0036012
O -6.1602659 -4.2700824 -56.2681912
C -5.0687729 -7.4390979 -56.0860549
H -5.3895788 -8.424192 -55.7418211
H -4.4305065 -7.5948128 -56.9556649
C -4.1992388 -6.7757217 -54.993123
H -3.2115868 -7.2341931 -55.0481208
H -4.0668317 -5.7140503 -55.2046993
C -4.7169933 -6.9550987 -53.5545555
H -4.8133604 -8.0239603 -53.351868
H -5.6999526 -6.4874112 -53.4551922
C -3.7314371 -6.3211879 -52.5553124
H -3.9467867 -5.2535484 -52.4822222
H -2.7197203 -6.4281609 -52.9491867
N -3.7880118 -6.9492504 -51.2117734
H -4.7008678 -6.8646728 -50.7950431
H -3.1294465 -6.5095671 -50.5629086
H -3.5314121 -7.9249485 -51.2652042
N -5.4101965 -5.0928218 -58.2204721
H -5.2513331 -5.9347831 -58.7634563
C -5.1139113 -3.8079675 -58.8626221
H -4.4172266 -3.2441876 -58.2414818
C -6.3827766 -2.9486339 -58.9977917
O -6.424878 -1.8461333 -58.4601641
C -4.4306352 -4.0925634 -60.2147685
H -3.4889227 -4.612916 -60.0251308
H -5.0681263 -4.750292 -60.8032861
C -4.1402352 -2.8329003 -61.0402102
H -3.5223312 -2.1630568 -60.4391233
H -5.0881717 -2.3339607 -61.2560169
C -3.4537445 -3.1310734 -62.3813936
O -3.6879035 -4.2298107 -62.9487496
O -2.7668995 -2.224337 -62.9038172
N -7.4664462 -3.5002571 -59.5499567

H -7.3734285 -4.4240226 -59.9575726
C -8.7116426 -2.7639545 -59.7964255
H -8.4353106 -1.8095062 -60.2480485
C -9.4638687 -2.4089424 -58.5074308
O -9.9528278 -1.288221 -58.3755535
C -9.6118555 -3.5466591 -60.7821039
H -9.0607949 -4.3693492 -61.2330581
H -10.4668038 -3.9705644 -60.2532929
C -10.1111674 -2.6392604 -61.9230753
H -10.7214885 -1.8416841 -61.4932013
H -10.7558622 -3.2309231 -62.5768496
C -8.9667387 -2.0313695 -62.7636765
O -8.2371866 -2.7994259 -63.4391515
O -8.8066446 -0.7847794 -62.7441526
N -9.47042 -3.2962327 -57.4999447
H -9.0417097 -4.20374 -57.6524768
C -10.1010431 -3.0170332 -56.1973756
H -11.0288266 -2.4764864 -56.3973745
C -9.2769901 -2.0761537 -55.3142795
O -9.8498703 -1.4157787 -54.4554675
C -10.4660358 -4.334596 -55.4944689
H -11.0058283 -4.9560219 -56.2112972
H -9.5566922 -4.8544857 -55.1890845
C -11.3704398 -4.1053174 -54.2696322
H -10.7558442 -3.807233 -53.4188719
H -12.0692675 -3.2953097 -54.4855386
C -12.2007088 -5.3376816 -53.8874421
H -12.8839548 -5.5640901 -54.7082912
H -12.7938671 -5.0865768 -53.0059178
C -11.3301939 -6.5636396 -53.5898349
H -10.6293076 -6.3072624 -52.7892185
H -10.7480368 -6.8020651 -54.4859351
N -12.1634946 -7.726805 -53.1936491
H -12.701381 -7.5095402 -52.3633894
H -11.5843428 -8.534113 -52.9993255
H -12.8073944 -7.9650457 -53.9371857
N -7.9716141 -1.9444161 -55.5661454
H -7.5758423 -2.5040646 -56.3110106
C -7.1488087 -0.8533673 -55.0040867
H -7.6222074 -0.5067725 -54.0856447
C -7.1064176 0.3871534 -55.9041949
O -6.4532318 1.363562 -55.5514579
C -5.7327836 -1.3132419 -54.6059369
H -5.1704231 -0.4493004 -54.2512706
C -5.7686133 -2.3392788 -53.4714731
H -6.3200593 -3.2274988 -53.7777969

H -6.2540956 -1.8980821 -52.6008782
H -4.7497511 -2.6174525 -53.2046511
O -5.0121386 -1.9015539 -55.6648057
H -5.5081724 -2.690785 -55.9458878
N -7.8166632 0.3654646 -57.0417343
H -8.3462446 -0.4726061 -57.2427193
C -7.8502309 1.3899778 -58.0923619
H -8.3461036 0.901136 -58.9292334
C -6.4594769 1.7892235 -58.633805
O -6.3001095 2.8344891 -59.2640218
C -8.7526115 2.5574542 -57.6322694
H -9.6187161 2.1305136 -57.1235023
H -8.1999693 3.1478449 -56.8997247
C -9.2803135 3.5032468 -58.7320415
H -8.4579137 4.0436594 -59.1952806
C -10.0659324 2.771923 -59.8264709
H -10.8596408 2.1741863 -59.3761475
H -9.4021933 2.1159166 -60.3858728
H -10.5024011 3.4948423 -60.5147045
C -10.2132689 4.5376623 -58.0986059
H -11.0746174 4.0426197 -57.649767
H -10.5543428 5.2422816 -58.85601
H -9.6730952 5.0878871 -57.3275553
N -5.4426723 0.9434196 -58.4481881
H -5.6678236 0.0355238 -58.0548346
C -4.0798935 1.185912 -58.9357536
H -3.8150141 2.2211407 -58.7113507
C -4.0051982 1.032849 -60.4518383
O -4.7678273 0.27695 -61.0434649
C -3.0949649 0.2568287 -58.2130025
H -3.5138127 -0.7490093 -58.1422002
H -2.1678002 0.2159683 -58.784929
C -2.795052 0.8141 -56.8097441
H -2.3119699 1.7866437 -56.9208895
H -3.7301621 0.9656837 -56.2690902
C -1.8893985 -0.0773955 -55.9502131
H -2.4545609 -0.9480272 -55.6154601
H -1.5950424 0.5033683 -55.0736083
C -0.6439782 -0.5463608 -56.7108241
H -0.3018319 0.268945 -57.3493463
H -0.9215663 -1.3830817 -57.3566641
N 0.4467047 -0.9339892 -55.7916971
H 0.7465728 -0.112789 -55.2498367
H 1.2748543 -1.2105249 -56.2979961
H 0.1611309 -1.6486113 -55.1440047
N -3.0487572 1.7262357 -61.0774219

H -2.4915023 2.3509615 -60.5015827
C -2.7384924 1.6213578 -62.5167704
H -3.2764944 0.767828 -62.9341887
C -1.2710996 1.2723612 -62.7035448
O -0.4066791 2.1504201 -62.6343866
C -3.1835379 2.9095297 -63.2545982
H -3.4195988 3.6894009 -62.5300726
H -2.3764105 3.2816492 -63.8842477
C -4.4157079 2.6815394 -64.1627795
H -4.7723658 3.6569256 -64.4963477
H -4.1117891 2.1108096 -65.0407656
C -5.5713926 1.9356428 -63.4710507
H -5.670715 2.3288555 -62.460506
H -5.3424209 0.8725846 -63.4125411
C -6.9189723 2.0674559 -64.1837241
H -6.9312356 1.427326 -65.0694601
H -7.0421595 3.108668 -64.4916204
N -8.0097774 1.7033435 -63.2510456
H -7.9605382 0.719054 -62.960874
H -8.919042 1.7965361 -63.6756161
H -7.9574248 2.2846703 -62.4272009
N -0.9851324 -0.0152724 -62.8685472
H -1.7507671 -0.6921228 -62.9343551
C 0.3622951 -0.5630426 -62.7563147
H 0.9805693 0.166144 -62.2307181
C 0.9623679 -0.7635467 -64.1569443
O 0.4536773 -1.5034089 -65.0035394
C 0.3662204 -1.8384641 -61.8795023
H -0.0630003 -2.6610733 -62.4548948
C 1.819163 -2.1735927 -61.5191769
H 2.2062048 -1.4454116 -60.8056847
H 2.4410637 -2.152216 -62.4072739
H 1.8767011 -3.1699076 -61.08193
C -0.4305112 -1.6738873 -60.5577608
H 0.0351023 -0.9039866 -59.9410057
H -1.442414 -1.3440835 -60.7786126
C -0.5557201 -2.9669112 -59.7395202
H 0.4082644 -3.2467547 -59.3150915
H -0.9233073 -3.7726035 -60.3764866
H -1.265722 -2.8133331 -58.927048
N 2.0820627 -0.0851478 -64.4029156
H 2.4655832 0.475942 -63.6575331
C 2.9068083 -0.2770199 -65.5962123
H 2.3018658 -0.6952979 -66.4020926
C 4.0396154 -1.2643317 -65.292497
O 4.4307098 -1.4269569 -64.1332412

C 3.4495725 1.0814523 -66.061202
H 4.0958151 0.9525892 -66.9322466
H 4.0432311 1.5292122 -65.261646
S 2.0709612 2.1846743 -66.4876026
H 1.3003227 1.8921115 -65.4342484
N 4.6448883 -1.8495044 -66.3268338
H 4.3466771 -1.6176306 -67.261478
C 5.9932657 -2.3856146 -66.1595621
H 6.0491473 -2.8706147 -65.1886549
C 7.0088041 -1.2347704 -66.1249314
O 6.737833 -0.132612 -66.6086652
C 6.3073177 -3.4524623 -67.2163512
H 5.4957053 -4.1766962 -67.2517385
H 7.2191217 -3.9815768 -66.9375472
C 6.5230252 -2.8364934 -68.5773457
O 7.5678347 -2.280934 -68.8635157
N 5.5086591 -2.8222407 -69.403895
H 5.6807276 -2.3748843 -70.3032807
H 4.6772054 -3.3460722 -69.2191437
N 8.1801732 -1.5131329 -65.5618081
H 8.3470506 -2.4508453 -65.2320084
C 9.2452561 -0.5331003 -65.3748175
H 8.8589516 0.286887 -64.7647094
C 9.6957767 0.098359 -66.7072817
O 9.7789872 1.3222783 -66.8014925
C 10.3873618 -1.2029495 -64.5780293
H 11.2574185 -0.549599 -64.5671677
H 10.0590078 -1.2929866 -63.5414857
C 10.8177317 -2.5865022 -65.0490904
C 11.9181375 -2.7365552 -65.9169673
H 12.4819757 -1.8755661 -66.2390297
C 12.3053206 -4.0136478 -66.363435
H 13.1452796 -4.1187607 -67.0335377
C 11.597087 -5.1547648 -65.9319127
O 11.9592972 -6.3881262 -66.3699191
H 12.7274721 -6.3595775 -66.9421564
C 10.5129368 -5.0118722 -65.040923
H 9.9908157 -5.8930459 -64.6965091
C 10.1282528 -3.7322316 -64.5973171
H 9.308215 -3.6379075 -63.8983316
N 9.914116 -0.6953223 -67.7660937
H 9.7185 -1.6813804 -67.6780474
C 10.4258322 -0.1969978 -69.0526211
H 11.3531195 0.3334907 -68.8684677
C 9.4744436 0.8147866 -69.7068052
O 9.9050969 1.903903 -70.0918449

C 10.7555701 -1.3866897 -69.9704476
H 11.4799712 -2.0264374 -69.4643361
H 9.8468928 -1.968167 -70.1290859
C 11.319076 -0.9912917 -71.3499645
H 10.5886703 -0.3825456 -71.8825983
C 12.633967 -0.2142643 -71.2488822
H 13.3625122 -0.779567 -70.6682336
H 12.462991 0.7532524 -70.7772734
H 13.0319279 -0.0350601 -72.2476385
C 11.5735498 -2.2547509 -72.1719436
H 12.3160219 -2.8816896 -71.6783682
H 11.9344282 -1.9816748 -73.1636924
H 10.6437644 -2.8131246 -72.2827093
N 8.1721243 0.5219023 -69.7300654
H 7.8781855 -0.3931025 -69.4008782
C 7.1627271 1.4694778 -70.216607
H 7.465206 1.8258281 -71.2019721
C 7.0485684 2.7218684 -69.344085
O 6.7977963 3.7974389 -69.8845881
C 5.8037654 0.7721156 -70.3591403
H 5.0142317 1.5249066 -70.3314356
H 5.6424419 0.0910384 -69.5229777
C 5.6722935 0.0300481 -71.6647418
N 5.9532087 -1.3209948 -71.8725283
C 5.7499517 -1.545272 -73.178947
H 5.897551 -2.4993019 -73.6695688
N 5.3500598 -0.4176791 -73.790252
H 5.1471925 -0.3273254 -74.7758818
C 5.2980715 0.5887433 -72.851667
H 5.0341347 1.6258243 -73.0140616
N 7.3164396 2.6332608 -68.0360767
H 7.5599259 1.7314697 -67.6420576
C 7.2614763 3.7968527 -67.1509479
H 6.3620564 4.3393293 -67.4291629
C 8.4261241 4.7738874 -67.3880553
O 8.1695259 5.9696373 -67.5199989
C 7.0948289 3.3363918 -65.6891448
H 6.3288467 2.5601209 -65.6683109
H 8.0271337 2.8928056 -65.3390609
C 6.6681347 4.4549309 -64.7117129
H 7.5189438 5.109165 -64.5258907
C 5.4997377 5.3156355 -65.2074584
H 4.6757884 4.679795 -65.5306189
H 5.8301918 5.9439793 -66.0348046
H 5.1677863 5.9791257 -64.4099764
C 6.2145039 3.8387536 -63.3851094

H 5.3016056 3.2618588 -63.5260214
H 6.0445249 4.6237662 -62.6487135
H 6.9882306 3.1722436 -63.0041928
N 9.6480168 4.2850584 -67.6533181
H 9.7997559 3.2846604 -67.5630426
C 10.7262591 5.1507473 -68.1801541
H 10.8557976 5.9933419 -67.499146
C 10.3952471 5.6900104 -69.5631661
O 10.7418516 6.824877 -69.8713061
C 12.0605472 4.4240518 -68.3016877
H 12.1166467 3.8610945 -69.2350101
H 12.121121 3.7293563 -67.4803064
O 13.1491082 5.3247477 -68.1948499
H 13.269569 5.8510611 -69.0056855
N 9.6793356 4.9112594 -70.3797769
H 9.4739327 3.9631262 -70.084374
C 9.1529553 5.3651188 -71.6612905
H 9.9800387 5.6169965 -72.3253587
H 8.5721268 4.5597261 -72.1096179
C 8.2436502 6.5889815 -71.5206075
O 8.4964877 7.5881455 -72.1829673
N 7.271716 6.5802667 -70.6017955
H 7.1008963 5.7179054 -70.0934582
C 6.3753098 7.7248641 -70.3564864
H 5.913338 8.0175862 -71.2986071
C 7.1064366 8.967594 -69.8279953
O 6.7901523 10.0960769 -70.2162708
C 5.2706065 7.3106618 -69.3736671
H 5.7289851 6.9026424 -68.4718638
H 4.7203521 8.2063148 -69.0832355
C 4.2527546 6.3259864 -69.9274463
C 3.9569052 5.1377987 -69.2312677
H 4.470828 4.90886 -68.3126257
C 2.9868958 4.2463839 -69.7247382
H 2.7656118 3.3367646 -69.1875547
C 2.2932897 4.5473782 -70.9100298
H 1.5351606 3.8746796 -71.2849157
C 2.5749518 5.7375725 -71.6014384
H 2.0297721 5.995317 -72.5016579
C 3.5527959 6.6201892 -71.1146242
H 3.7383022 7.5375435 -71.6559592
N 8.1333138 8.7500029 -69.0098282
H 8.2811018 7.8069615 -68.6678179
C 9.07562 9.7716512 -68.5730746
H 8.5547353 10.5922679 -68.0912188
C 9.8605926 10.3853852 -69.7513712

O 9.7566054 11.5847949 -70.0065046
C 9.9597189 9.1076336 -67.5219712
H 10.8339721 9.7058895 -67.3759088
H 10.2895408 8.1279684 -67.8552867
S 9.0625811 8.9471298 -65.9503501
H 9.8351265 9.8474364 -65.3019252
N 10.4805271 9.5504548 -70.5898632
H 10.5030738 8.5688311 -70.3340113
C 11.2121939 9.9579073 -71.8004392
H 12.0155281 10.6371849 -71.5108146
C 10.3189662 10.6842638 -72.8220458
O 10.6706195 11.7496569 -73.3218699
C 11.8287769 8.7012708 -72.4308687
H 11.0787146 8.1829872 -73.0301747
H 12.1581597 8.0264929 -71.641054
O 12.9535704 9.016732 -73.2287619
H 13.2151123 8.2260534 -73.7115009
N 9.0917743 10.1931924 -73.0341901
H 8.8462209 9.3288767 -72.56349
C 8.0758771 10.7997598 -73.908273
H 8.5230395 10.9933282 -74.8827677
C 7.5601152 12.1572996 -73.4059811
O 6.9709603 12.9057875 -74.1813275
C 6.8980951 9.8177845 -74.0665242
H 6.5963806 9.4797408 -73.0746319
H 6.0469815 10.3468555 -74.4990079
C 7.2010356 8.596359 -74.9559661
H 8.1359891 8.1311667 -74.6495739
C 6.078985 7.5667695 -74.8172612
H 5.1265064 8.0041828 -75.1217947
H 6.002105 7.244648 -73.7790986
H 6.2918725 6.6974123 -75.4373065
C 7.3195826 8.9722824 -76.4354788
H 6.4050387 9.4644876 -76.7684964
H 7.4802548 8.0744735 -77.031481
H 8.1643764 9.6422728 -76.586479
N 7.8083199 12.5056056 -72.1428918
H 8.3267403 11.8644839 -71.5540116
C 7.4700463 13.8201725 -71.5844624
H 6.6249218 14.2248713 -72.140381
C 8.5965015 14.8537852 -71.7283394
O 8.4412966 15.9715581 -71.2569983
C 7.0096966 13.6702722 -70.1392581
H 6.7278152 14.646009 -69.7435907
H 7.8241777 13.2648047 -69.5378654
O 5.8818799 12.8084382 -70.0899695

H 6.1934362 11.8939971 -70.2199382
N 9.6527789 14.5194905 -72.4726668
H 9.6609405 13.583641 -72.8564542
C 10.8450282 15.3307806 -72.7823003
H 11.3215018 14.7681748 -73.5857787
C 11.9223128 15.3449434 -71.6844862
O 12.7208264 16.2703293 -71.5460031
C 10.4885248 16.6983789 -73.4123986
H 10.3052822 17.4216129 -72.6154718
H 9.5630553 16.5886969 -73.9810148
C 11.5483248 17.2610639 -74.3780375
O 12.3958398 16.4764423 -74.8659835
O 11.4371665 18.4657644 -74.7049311
N 11.9834183 14.26542 -70.8998863
H 11.310854 13.5258562 -71.0438426
C 13.0028759 14.0912496 -69.8714545
H 13.306885 15.0728525 -69.5156226
C 14.2829326 13.4339092 -70.3807713
O 14.2769208 12.3465484 -70.963321
C 12.4106606 13.3349922 -68.6923727
H 12.0504773 12.3665639 -69.0383334
H 13.215576 13.1470367 -67.9823596
C 11.3089208 14.0511588 -67.9372688
C 10.4143586 13.2895275 -67.1652494
H 10.4207411 12.2129036 -67.2452246
C 9.622269 13.9020251 -66.183757
H 9.0420531 13.2912381 -65.5123696
C 9.6896914 15.2939169 -66.0117228
O 9.0688287 15.8810816 -64.9659394
H 8.6517885 15.2197967 -64.4135997
C 10.476267 16.0688853 -66.8777947
H 10.5008894 17.1334103 -66.7732363
C 11.2869235 15.4568867 -67.8389113
H 11.9309688 16.084664 -68.4366982
N 15.413457 14.0666489 -70.0648519
H 15.3470006 14.8954175 -69.4865486
C 16.7466287 13.6072225 -70.4479379
H 16.6439033 13.0148765 -71.3578561
C 17.3845581 12.7089696 -69.3861672
O 17.0498046 12.785335 -68.2019845
C 17.6340751 14.8021464 -70.8021027
H 17.0888067 15.4656922 -71.4759625
H 18.5250258 14.4458703 -71.3186039
O 18.0387931 15.5263655 -69.6592734
H 17.3011331 15.5682376 -69.0191956
N 18.3085085 11.8526167 -69.839663

H 18.540374 11.8840546 -70.8182419
C 19.0346307 10.8940314 -69.0057114
H 19.4675616 10.1472293 -69.6713046
C 18.1121144 10.1118048 -68.0476025
O 18.3664568 10.0593789 -66.8463913
C 20.2060694 11.6112569 -68.3167241
H 19.8233584 12.3411857 -67.6012615
H 20.822286 12.1241624 -69.055329
H 20.8190484 10.8836954 -67.7848943
N 17.0082147 9.5547067 -68.5680509
H 16.8205954 9.6340163 -69.5538031
C 16.1258953 8.7216485 -67.7571872
H 16.0455573 9.2076642 -66.7871337
C 16.7726785 7.3640399 -67.4612841
O 17.037384 6.5613184 -68.3551552
C 14.6792575 8.6592675 -68.293561
H 14.2122908 9.6269136 -68.1101076
H 14.1203773 7.931987 -67.704949
C 14.479784 8.3334479 -69.7606206
O 14.1690558 7.2130685 -70.1375173
N 14.5042034 9.3331716 -70.6191546
H 14.1908225 9.1433714 -71.5603483
H 14.5384129 10.2922772 -70.2963576
N 17.0392665 7.1424113 -66.177116
H 16.830673 7.8928146 -65.5251381
C 17.5157217 5.8978433 -65.5954028
H 17.9177789 5.2616605 -66.384763
C 16.3357946 5.207524 -64.9288189
O 15.6159122 5.8225069 -64.1466295
C 18.6212937 6.1737034 -64.5672921
H 18.1711266 6.6156535 -63.6769974
H 19.0702786 5.2249355 -64.2716113
C 19.7093446 7.0913046 -65.0380035
C 19.7931824 8.4044157 -64.7311366
H 19.0720161 8.9404149 -64.124824
N 20.8504143 8.9783652 -65.4059943
H 20.9953366 9.9773354 -65.4322483
C 21.4912532 8.0630289 -66.2111008
C 22.5607112 8.1678796 -67.1126483
H 23.0547657 9.1163029 -67.2628505
C 22.976687 7.0234162 -67.8144749
H 23.7994497 7.0845023 -68.5131482
C 22.3164877 5.7977862 -67.6094385
H 22.6337933 4.9207785 -68.1562422
C 21.2368826 5.7069224 -66.707767
H 20.7248034 4.766601 -66.5741431

C 20.7952953 6.8365602 -65.9837824
N 16.1848563 3.912817 -65.176291
H 16.821799 3.469261 -65.8205527
C 15.0796393 3.1104436 -64.6775838
H 14.54308 3.6333506 -63.8833365
C 15.6247004 1.7985474 -64.1325112
O 16.0577958 0.9402323 -64.9020713
C 14.131694 2.8887033 -65.8487103
H 13.64995 3.8395378 -66.0623428
H 14.7289034 2.6128478 -66.7202805
C 13.0901348 1.8002141 -65.5654381
H 13.4487714 1.0489044 -64.8679001
C 11.8698551 2.4330043 -64.9143906
H 11.3898081 3.1481945 -65.575671
H 12.1699034 2.9421886 -63.9979327
H 11.1508884 1.6662489 -64.6483251
C 12.8433832 1.0456084 -66.8611406
H 12.3756177 1.6885177 -67.5980209
H 12.2143535 0.1981272 -66.6495824
H 13.7858171 0.6635186 -67.2549475
N 15.5022455 1.6123315 -62.8227831
H 15.0176951 2.3212374 -62.2749974
C 15.9819101 0.4156113 -62.1354336
H 16.5767406 -0.1688088 -62.8309248
C 14.7948239 -0.412656 -61.638592
O 14.0877462 0.0314062 -60.7298387
C 16.9228566 0.7713672 -60.9737509
H 16.3487296 1.1324943 -60.119959
C 17.7625409 -0.435787 -60.5611816
H 18.3692173 -0.7740933 -61.4018301
H 17.1180753 -1.2500821 -60.2345302
H 18.4224701 -0.1512396 -59.7425426
O 17.83844 1.7718527 -61.3585353
H 17.3314862 2.5714806 -61.5236629
N 14.5757734 -1.6381667 -62.1522123
C 13.7832369 -2.6243789 -61.4326093
H 12.8078798 -2.2156713 -61.1651937
C 14.5459745 -3.0172483 -60.1610187
O 15.6270337 -3.5983735 -60.2304277
C 13.5987399 -3.7952014 -62.405327
H 13.5382697 -4.7567764 -61.8952256
H 12.6985773 -3.6284725 -62.9974839
C 15.1494966 -2.2202696 -63.3610773
H 14.6833669 -1.7644417 -64.236083
H 16.2312348 -2.0961457 -63.4033807
C 14.823558 -3.7137458 -63.3158808

H 14.6132543 -4.1093829 -64.3096971
H 15.6551597 -4.2543635 -62.8618106
N 13.9662979 -2.746838 -58.9908767
H 13.0661721 -2.2754632 -59.0090904
C 14.5182136 -3.1102436 -57.6789229
H 15.4998097 -3.5591397 -57.8229977
C 13.7014467 -4.2044761 -56.9428997
O 13.6238671 -4.1604091 -55.7146881
C 14.714941 -1.81596 -56.8514598
H 13.7756341 -1.2643591 -56.8431172
H 14.9534945 -2.0737327 -55.819355
C 15.8229594 -0.873091 -57.3541501
H 15.6404761 -0.6146515 -58.3959849
C 15.8120489 0.4127376 -56.5303236
H 16.028931 0.1980959 -55.4826737
H 14.8309305 0.8833728 -56.593059
H 16.5554212 1.1109018 -56.9145674
C 17.2136708 -1.5018124 -57.2217831
H 17.3755269 -1.8389114 -56.1975999
H 17.9750578 -0.7631874 -57.470298
H 17.3151205 -2.3433035 -57.9050428
N 13.0948036 -5.214763 -57.6151661
C 12.2053006 -6.1721577 -56.9514439
H 11.3888186 -5.644905 -56.4617025
C 12.9249789 -7.0386965 -55.9075064
O 12.3195788 -7.4085346 -54.9111168
C 11.6162382 -7.0363963 -58.0682371
H 11.3764525 -8.0460584 -57.7312578
H 10.7270492 -6.5549761 -58.4702904
C 13.2765494 -5.6211828 -59.0043124
H 12.7053484 -4.9577645 -59.6524598
H 14.33053 -5.6213029 -59.2835882
C 12.7120005 -7.0340446 -59.1282169
H 12.3119336 -7.2235629 -60.1252091
H 13.4796364 -7.7662537 -58.8743209
N 14.2193922 -7.3309605 -56.0875591
H 14.6758111 -6.9951267 -56.9187628
C 15.0103271 -8.0990672 -55.112049
H 14.434692 -8.9910839 -54.8655987
C 15.2070223 -7.332927 -53.7964256
O 15.0766997 -7.9116657 -52.717904
C 16.3497863 -8.552893 -55.7413196
H 16.1067111 -9.1238207 -56.6393262
C 17.2457079 -7.3784555 -56.1790683
H 17.6104389 -6.833258 -55.3064419
H 16.7096291 -6.695959 -56.8363059

H 18.1109972 -7.7622999 -56.7213065
C 17.1758962 -9.4774166 -54.8192359
H 18.0864798 -9.7670527 -55.3455807
H 17.4716894 -8.935527 -53.9193684
C 16.4479887 -10.7630839 -54.4053729
H 16.0929498 -11.2919093 -55.2902479
H 15.605773 -10.5327737 -53.7532482
H 17.1389073 -11.4070984 -53.8607548
N 15.3935547 -6.0141343 -53.87863
H 15.4044782 -5.57695 -54.785056
C 15.4989416 -5.1573755 -52.7005961
H 16.1260958 -5.6549509 -51.9609007
C 14.1275042 -4.979588 -52.0493859
O 14.0041547 -5.2410058 -50.8625342
C 16.1529791 -3.8128398 -53.0612849
H 15.5453294 -3.3096024 -53.8136461
H 16.1749842 -3.1849001 -52.1697456
C 17.5888162 -3.9565456 -53.5969701
H 17.9604463 -2.9636573 -53.8583359
H 17.5825816 -4.5500643 -54.513358
C 18.5239075 -4.5941443 -52.560969
O 18.5957494 -5.8428049 -52.4662396
O 19.14147 -3.8633975 -51.7551244
N 13.0718701 -4.7526557 -52.834066
H 13.246795 -4.5814834 -53.8192455
C 11.6674641 -4.7573474 -52.3915481
H 11.5110988 -3.9424476 -51.6838844
C 11.2638044 -6.055375 -51.6705936
O 10.7000723 -5.9916155 -50.5815018
C 10.8225721 -4.4658085 -53.6373247
H 11.2703018 -4.9725609 -54.4847155
H 10.8980451 -3.3996945 -53.8162933
C 9.3400869 -4.8422281 -53.6032018
O 8.5960027 -4.4336681 -52.6834481
O 8.8625046 -5.4577929 -54.5871172
N 11.6917214 -7.2238873 -52.1577427
H 12.1378338 -7.2256039 -53.0701622
C 11.4803921 -8.5084706 -51.4804795
H 10.4192842 -8.5902366 -51.2439924
C 12.2581264 -8.616735 -50.1576347
O 11.7173718 -9.1125472 -49.1729519
C 11.8590517 -9.6523727 -52.4351888
H 12.9441046 -9.6885066 -52.5437256
H 11.4310558 -9.4628766 -53.4176041
C 11.3583046 -11.0199242 -51.9562242
H 11.7482812 -11.2164564 -50.9570363

H 11.7634607 -11.7769659 -52.6281464
S 9.5513028 -11.2041661 -51.9296248
C 9.4280423 -12.9497713 -51.463499
H 8.3802362 -13.2482151 -51.4521366
H 9.8585708 -13.0948903 -50.4720088
H 9.9653893 -13.5614913 -52.1881289
N 13.4992124 -8.1155487 -50.1013757
H 13.8751445 -7.7085546 -50.9531837
C 14.3462607 -8.0992723 -48.8917458
H 14.3261654 -9.0880359 -48.4302966
C 13.8414349 -7.109273 -47.8321094
O 13.8696885 -7.4088854 -46.6438106
C 15.7803139 -7.7771225 -49.3386629
H 16.0728206 -8.4953921 -50.1072518
H 15.78842 -6.7853287 -49.7877521
C 16.8146607 -7.8333564 -48.2019743
H 16.5459291 -7.1366515 -47.4067893
H 16.8280532 -8.8437172 -47.7908348
C 18.2187971 -7.4873246 -48.7193819
H 18.4184365 -8.0829343 -49.6128579
H 18.9536731 -7.7499724 -47.9570584
C 18.3487833 -5.992413 -49.0458139
H 18.4319706 -5.4217815 -48.1165061
H 17.4451803 -5.6561907 -49.5619324
N 19.5031294 -5.7293883 -49.9309239
H 20.3954323 -5.9966135 -49.5549232
H 19.5174901 -4.749167 -50.2346865
H 19.3464276 -6.1609146 -50.848426
N 13.3322727 -5.9630881 -48.2710839
H 13.3916351 -5.7966786 -49.2700395
C 12.6752334 -4.9259791 -47.4652083
H 13.3066394 -4.639705 -46.6204001
C 11.3514425 -5.4659331 -46.907204
O 11.0801258 -5.3692947 -45.708615
C 12.463281 -3.6936637 -48.3766892
H 12.0638674 -4.0566426 -49.3241162
C 11.3939495 -2.735481 -47.8251092
H 11.5699158 -2.5482408 -46.7698325
H 10.4027027 -3.1721763 -47.9439616
H 11.4167651 -1.795219 -48.3641339
C 13.8090055 -3.0011842 -48.7056933
H 14.1943818 -2.5034339 -47.8276267
H 14.5553025 -3.7476582 -48.9678682
C 13.7415405 -1.9940272 -49.8643963
H 13.1388901 -1.128049 -49.5957665
H 13.3172396 -2.4665957 -50.7493734

H 14.7489847 -1.6485579 -50.0979748
N 10.5569209 -6.1251239 -47.7575478
H 10.8058184 -6.1502801 -48.7417588
C 9.3406062 -6.8155937 -47.3499945
H 8.687436 -6.0980664 -46.8515223
C 9.6451199 -7.9320667 -46.3426887
O 8.9789309 -7.9857565 -45.3181334
C 8.6090851 -7.3439192 -48.5900398
H 9.2347282 -8.0646011 -49.1166765
H 8.3743305 -6.5167723 -49.2610325
H 7.6825766 -7.8315087 -48.286617
N 10.6967849 -8.7327052 -46.5459848
H 11.1877227 -8.6812436 -47.4322827
C 11.1410652 -9.7413254 -45.5766499
H 10.3095113 -10.4188421 -45.3765601
C 11.5417736 -9.1181235 -44.2316211
O 11.0027769 -9.5243077 -43.2065835
C 12.2879134 -10.5821685 -46.1574739
H 13.1383181 -9.9410307 -46.3900585
H 11.9539012 -11.054893 -47.0827182
C 12.7241101 -11.6616207 -45.1622198
O 13.5224159 -11.3337599 -44.252894
O 12.2018212 -12.793503 -45.265959
N 12.3829782 -8.0769071 -44.2118859
H 12.8033958 -7.7736289 -45.0842588
C 12.8418208 -7.4756884 -42.9545116
H 13.20788 -8.2795145 -42.315361
C 11.7105569 -6.7843906 -42.179033
O 11.5105007 -7.0762136 -40.995856
C 14.0107534 -6.5158331 -43.2086705
H 14.8199205 -7.0620265 -43.6953675
H 13.6867322 -5.7238326 -43.8863098
C 14.5352889 -5.889822 -41.9505213
C 15.3513898 -6.4920806 -41.0560262
H 15.7532765 -7.4912347 -41.1718495
N 15.5335855 -5.6654725 -39.9629168
H 16.0897329 -5.9158626 -39.160673
C 14.8148192 -4.49376 -40.0872147
C 14.6321954 -3.3866221 -39.242576
H 15.1032791 -3.3500841 -38.2735638
C 13.819259 -2.3236695 -39.676404
H 13.6551516 -1.4597462 -39.0434416
C 13.2231146 -2.3758491 -40.9490175
H 12.61592 -1.5480294 -41.2963996
C 13.413713 -3.4925454 -41.785488
H 12.9690569 -3.4997165 -42.7718436

C 14.1985453 -4.5894945 -41.3712467
N 10.9043925 -5.9527161 -42.8492866
H 11.1134384 -5.7630804 -43.8240366
C 9.7438538 -5.2921988 -42.2212211
H 10.0844229 -4.773692 -41.3238945
C 8.6828352 -6.2978319 -41.7797415
O 8.1475363 -6.1684389 -40.6765001
C 9.0796136 -4.2487583 -43.1304959
H 8.177974 -3.8895772 -42.6371761
C 9.9720763 -3.0467225 -43.4104666
H 10.8669468 -3.3593763 -43.9445606
H 10.2661708 -2.5763767 -42.4714854
H 9.4314004 -2.3171273 -44.0117579
O 8.7181591 -4.8066588 -44.3726664
H 9.5496823 -4.9479732 -44.8577988
N 8.4358032 -7.3581704 -42.5633335
H 8.8963046 -7.4314515 -43.466883
C 7.5723311 -8.4495715 -42.1268535
H 6.6350766 -8.0155664 -41.7795424
C 8.1700975 -9.1517994 -40.9247922
O 7.4758181 -9.174459 -39.9330543
C 7.2448461 -9.4620492 -43.2330206
H 8.1639771 -9.7685181 -43.7320877
H 6.8107539 -10.3548548 -42.779457
C 6.239774 -8.9211213 -44.2597927
H 6.2060887 -9.6067299 -45.1067549
H 6.5649357 -7.9487869 -44.6272582
C 4.8301142 -8.7907822 -43.6962677
O 4.0185217 -9.6936336 -43.7800733
N 4.4373901 -7.6329512 -43.2092787
H 3.4549186 -7.5555601 -42.9669857
H 5.0762198 -6.8618807 -43.140931
N 9.4273277 -9.6006942 -40.9284614
H 9.9686587 -9.4839231 -41.780828
C 10.0214635 -10.412134 -39.8567427
H 9.3084547 -11.1990181 -39.609992
C 10.253828 -9.6601678 -38.5291229
O 10.2250454 -10.2893742 -37.4666527
C 11.2944858 -11.0964365 -40.3945858
H 11.8618845 -11.5134649 -39.5643313
H 11.9244235 -10.3674132 -40.904203
C 10.9694015 -12.2521054 -41.3360884
O 10.0165345 -12.9890257 -41.1245311
N 11.7728694 -12.4937022 -42.343769
H 11.4995019 -13.1733068 -43.0362986
H 12.499172 -11.8467642 -42.6449554

N 10.2929531 -8.3240194 -38.537167
H 10.3624872 -7.8437779 -39.4304161
C 10.2075646 -7.5261957 -37.311539
H 10.9370343 -7.9083139 -36.5955343
C 8.8173012 -7.6078404 -36.6366541
O 8.7224682 -7.6976754 -35.4119872
C 10.5911494 -6.0827475 -37.6571958
H 9.8848154 -5.6694188 -38.3793169
H 11.5923603 -6.0574983 -38.0909011
H 10.5793001 -5.471428 -36.7549126
N 7.7281362 -7.648853 -37.4093658
H 7.8464977 -7.6620426 -38.4175533
C 6.3615845 -7.7065981 -36.8837774
H 6.2525433 -6.8732437 -36.1891368
C 5.9991926 -8.9928891 -36.0905116
O 5.3798625 -8.8602907 -35.0312186
C 5.3875801 -7.4605357 -38.0417105
H 5.317507 -8.3466364 -38.6717451
H 5.7360643 -6.6360806 -38.6607618
H 4.3964641 -7.2320804 -37.651936
N 6.3537059 -10.227742 -36.5201682
C 6.0921003 -11.4546801 -35.7992746
H 5.0595046 -11.4728627 -35.4552056
C 7.0141917 -11.5868196 -34.5825291
O 6.5442037 -12.022288 -33.5387212
C 6.2944393 -12.5878507 -36.8162142
H 6.5796654 -13.5239315 -36.3340225
H 5.3790469 -12.7278175 -37.3944718
C 6.9710996 -10.6108314 -37.7595143
H 6.2311127 -10.4906961 -38.55036
H 7.8621115 -10.0271148 -37.9027458
C 7.3852351 -12.0701274 -37.7139111
H 7.3632158 -12.5333561 -38.7012058
H 8.3603036 -12.1870356 -37.2408137
N 8.2600128 -11.0993172 -34.6473682
H 8.5974122 -10.7351963 -35.529587
C 9.1638845 -11.0324667 -33.493428
H 9.2559856 -12.0264496 -33.0531509
C 8.6246533 -10.1030979 -32.3918366
O 8.5898426 -10.4804776 -31.2234502
C 10.5463395 -10.5906214 -33.9929249
H 10.9400176 -11.3588206 -34.6604347
H 10.4349591 -9.6795107 -34.5821898
C 11.5645544 -10.3336599 -32.8982995
C 12.1676299 -11.4139774 -32.226674
H 11.9007571 -12.4282594 -32.4853161

C 13.115578 -11.1764532 -31.2147571
H 13.5750579 -12.007349 -30.6985902
C 13.464218 -9.8578058 -30.873536
H 14.194 -9.6750549 -30.0974929
C 12.8624226 -8.7769348 -31.541472
H 13.1324783 -7.7636193 -31.281157
C 11.9107974 -9.0133361 -32.5496745
H 11.4482941 -8.1777601 -33.0568941
N 8.0603085 -8.9495652 -32.7631566
H 8.1434311 -8.665565 -33.7337643
C 7.4168677 -8.0211198 -31.8271501
H 7.9214617 -8.1082216 -30.8649975
C 5.9315139 -8.326438 -31.5456683
O 5.2535744 -7.4801421 -30.961817
C 7.5941536 -6.574922 -32.324633
H 7.124499 -6.4941971 -33.3064549
H 7.0538484 -5.9068789 -31.6550215
C 9.0136829 -6.0386909 -32.3987931
C 9.4463363 -5.359517 -33.5561945
H 8.7912065 -5.2787379 -34.4129759
C 10.7433545 -4.8150995 -33.6203321
H 11.075635 -4.3075551 -34.5142454
C 11.6148391 -4.9453172 -32.5180674
O 12.8717441 -4.435554 -32.5709646
H 13.0345853 -3.9454995 -33.3800472
C 11.174962 -5.6029107 -31.3511048
H 11.8453005 -5.6829437 -30.507972
C 9.8769556 -6.144496 -31.2893049
H 9.5563376 -6.6474406 -30.3872537
N 5.3871766 -9.481238 -31.956426
H 6.0033601 -10.1538206 -32.4006634
C 3.9274016 -9.6951489 -32.0875758
H 3.5788743 -9.0167038 -32.8656313
C 3.101727 -9.3795924 -30.8315989
O 1.9946272 -8.8568 -30.9597617
C 3.6184426 -11.1199467 -32.5819641
H 4.1373467 -11.2775613 -33.5237867
C 4.0262775 -12.229422 -31.6076246
H 3.4924745 -12.148523 -30.6645446
H 5.0959923 -12.1726016 -31.405786
H 3.820187 -13.1996915 -32.0601955
O 2.2341245 -11.2757929 -32.8256162
H 2.1031661 -11.0808149 -33.7675107
N 3.6312656 -9.6287895 -29.6257628
H 4.5649006 -10.0097299 -29.5872965
C 2.9405287 -9.3177676 -28.3687039

H 1.9221669 -9.6969241 -28.445873
C 2.8276563 -7.8017421 -28.1507249
O 1.7167741 -7.2833488 -28.1555535
C 3.6362754 -10.0368499 -27.1973246
H 3.6353806 -11.1098669 -27.3948457
H 4.6745595 -9.7054275 -27.1474315
C 2.978396 -9.7839054 -25.8233671
H 2.9966347 -8.7178673 -25.6005677
C 1.5310448 -10.2812004 -25.7697653
H 1.4817257 -11.3377576 -26.0293787
H 0.9096754 -9.6938931 -26.4446131
H 1.1425322 -10.1385847 -24.7604909
C 3.7720642 -10.514862 -24.7416685
H 3.7563083 -11.5903824 -24.9153329
H 3.3368301 -10.2987848 -23.7649057
H 4.8026104 -10.1599394 -24.7388168
N 3.9616382 -7.0915666 -28.1298216
H 4.8305866 -7.5964343 -28.1841824
C 4.0327294 -5.6286734 -27.9990707
H 3.5475118 -5.3373421 -27.066531
C 3.3223149 -4.9052738 -29.1588863
O 2.7887892 -3.8112358 -29.0000078
C 5.522688 -5.2420034 -27.9073454
H 5.9417744 -5.7125461 -27.0159418
H 6.0387796 -5.6508386 -28.77724
C 5.8148372 -3.7290084 -27.839953
H 5.4302001 -3.2507583 -28.740477
C 5.2052008 -3.0369301 -26.6265279
H 5.5270028 -3.5308572 -25.7079944
H 4.1165442 -3.0902357 -26.6670599
H 5.5002705 -1.9890255 -26.5938106
C 7.3263404 -3.5054407 -27.8182893
H 7.7487717 -3.9367071 -26.9075049
H 7.5460048 -2.4366408 -27.8259105
H 7.7977514 -3.9704279 -28.681937
N 3.3240981 -5.4922525 -30.3565642
H 3.8200164 -6.3736162 -30.4481514
C 2.6217824 -4.9657189 -31.5266407
H 2.8835245 -3.9148142 -31.6417372
C 1.0943839 -5.029999 -31.3514762
O 0.4108227 -4.0473604 -31.6319395
C 3.127867 -5.7370606 -32.7582961
H 4.209855 -5.607543 -32.8206349
H 2.9292394 -6.7975108 -32.6056175
C 2.5111026 -5.3318787 -34.1077532
H 1.4532277 -5.5947615 -34.1197899

C 2.6686143 -3.8435188 -34.4097948
H 3.7203921 -3.5715168 -34.3494023
H 2.0853599 -3.2554452 -33.704076
H 2.2981698 -3.6320755 -35.4139741
C 3.226156 -6.095398 -35.2194507
H 4.2560045 -5.7514367 -35.3113525
H 2.7083162 -5.937219 -36.1650524
H 3.2404669 -7.1588567 -34.9842874
N 0.5617721 -6.1321914 -30.8100106
H 1.1854655 -6.8955181 -30.565335
C -0.8774119 -6.2932057 -30.5294087
H -1.4450707 -5.8133694 -31.3273322
C -1.3387857 -5.5989339 -29.2472027
O -2.4972833 -5.2107496 -29.1799714
C -1.2259879 -7.7857952 -30.5059816
H -0.5395652 -8.3161481 -29.8425807
H -2.2423577 -7.9080194 -30.1304602
C -1.1550235 -8.3653732 -31.9235204
H -0.1595575 -8.2283015 -32.3457716
H -1.8592111 -7.825137 -32.5503861
C -1.517416 -9.8499764 -31.9535904
H -1.7789138 -10.1101266 -32.9786297
H -2.3926915 -10.0247935 -31.3275004
N -0.3815104 -10.6888054 -31.5472774
H 0.4686067 -10.5891001 -32.0922565
C -0.2819976 -11.5284697 -30.5397689
N 0.7987728 -12.2399241 -30.4453776
H 0.9213052 -12.9191248 -29.7195572
H 1.4926744 -12.1243647 -31.1788985
N -1.2140364 -11.691179 -29.6413828
H -2.0524106 -11.1461476 -29.7056909
H -1.0837421 -12.3425586 -28.8909044
N -0.4359914 -5.4137744 -28.291648
H 0.4565919 -5.878876 -28.4106519
C -0.6121594 -4.6637386 -27.0405899
H -1.5484464 -4.9448234 -26.5574101
C -0.6419177 -3.1442925 -27.2699944
O -1.4719593 -2.454789 -26.6849815
C 0.5604857 -5.0831334 -26.1452238
H 0.4619065 -6.1502756 -25.9371324
H 1.4675594 -4.941078 -26.7285774
C 0.7664583 -4.3591921 -24.8161269
H -0.1072872 -4.5150072 -24.1788573
H 0.8875321 -3.2879945 -24.9917974
C 2.035639 -4.9226683 -24.1581527
O 1.8929473 -5.6917077 -23.1831504

O 3.1384206 -4.6239782 -24.6732221
N 0.1917402 -2.6286023 -28.1822229
H 0.9142711 -3.2259607 -28.5641261
C 0.1832659 -1.2150978 -28.5658876
H 0.0037262 -0.6137299 -27.6722889
C -0.9246664 -0.8558517 -29.5768898
O -1.3427934 0.2986789 -29.6289793
C 1.5712314 -0.8568396 -29.111109
H 1.7755088 -1.4407739 -30.0082925
H 2.3275326 -1.0683264 -28.3535512
H 1.5991124 0.2052893 -29.3567835
N -1.3690744 -1.8151607 -30.399589
H -0.9570034 -2.7323259 -30.3107115
C -2.4122034 -1.6197109 -31.4130676
H -2.8401389 -0.6237023 -31.2850044
C -3.5838278 -2.5891446 -31.1989809
O -4.4984085 -2.2893246 -30.4387788
C -1.8158924 -1.6685056 -32.833723
H -1.3145361 -2.624073 -32.987596
H -2.63415 -1.6141187 -33.553035
C -0.8409829 -0.5505884 -33.1512774
C -1.3096435 0.649862 -33.7150379
H -2.3647469 0.7833783 -33.9051926
C 0.531843 -0.7025696 -32.8802269
H 0.8864084 -1.6149005 -32.4260483
C -0.4110316 1.6879453 -34.01348
H -0.7806818 2.6123909 -34.4308109
C 1.4300197 0.3388147 -33.1757978
H 2.4760809 0.2347951 -32.9481857
C 0.9602995 1.5313944 -33.7512922
H 1.6466328 2.3356076 -33.9776088
N -3.6205628 -3.7132496 -31.9236746
H -2.8479066 -3.9034345 -32.5436994
C -4.6046085 -4.7861899 -31.7255437
H -4.6477881 -5.0219769 -30.6606922
C -4.2176418 -6.0541322 -32.4919207
O -3.4466607 -6.006089 -33.4557384
C -6.0082382 -4.3507814 -32.1811528
H -6.3304691 -3.4699273 -31.624698
H -6.7175425 -5.152142 -31.9705056
O -6.0325689 -4.0692292 -33.5679786
H -5.899994 -3.1163384 -33.6518697
N -4.8741413 -7.1737827 -32.163381
H -5.4716506 -7.1487112 -31.3494907
C -4.8665038 -8.4037354 -32.9751168
H -3.8421984 -8.7622554 -33.024553

C -5.296619 -8.1459118 -34.4291164
O -4.6251069 -8.5946963 -35.3565651
C -5.7191938 -9.4992887 -32.2851475
H -5.2617698 -9.6962688 -31.3135578
C -7.1749464 -9.0643969 -32.0189038
H -7.7180113 -8.9252308 -32.9539572
H -7.2157387 -8.1483551 -31.4309694
H -7.6857446 -9.8419113 -31.4478146
C -5.7591062 -10.8389807 -33.0521625
H -6.3486961 -11.5529072 -32.4750014
H -6.2623358 -10.6958331 -34.0075932
C -4.3868889 -11.4732058 -33.3065014
H -3.86465 -11.6184485 -32.3614838
H -3.7929475 -10.8453952 -33.9693974
H -4.5241808 -12.4437534 -33.7840225
N -6.3197801 -7.3081212 -34.6459337
H -6.7867577 -6.9174776 -33.8407924
C -6.8108968 -6.9429317 -35.9822531
H -6.9295796 -7.8697976 -36.5438468
C -5.8010042 -6.0946841 -36.7703672
O -5.7534856 -6.2049544 -37.9993011
C -8.1929232 -6.2678168 -35.828368
H -8.8936588 -6.9935093 -35.4113768
H -8.1035006 -5.4462128 -35.1147961
C -8.795135 -5.6932521 -37.1238768
H -9.7559562 -5.2332397 -36.8894435
H -8.1411857 -4.8942535 -37.4703831
C -9.0162358 -6.7421972 -38.2346595
H -9.9843958 -7.2211919 -38.076221
H -8.2596339 -7.5233973 -38.1830747
C -8.9707821 -6.1540308 -39.6519625
H -9.7832162 -5.4324325 -39.7803459
H -9.1312743 -6.9749907 -40.3575952
N -7.6582374 -5.5172426 -39.9201159
H -7.5238449 -4.6979612 -39.3466649
H -7.5232324 -5.260878 -40.8933727
H -6.8930147 -6.1433565 -39.6825251
N -4.9831557 -5.2914209 -36.0868715
H -5.1114733 -5.2316311 -35.0833861
C -3.8621657 -4.5557308 -36.6755995
H -4.2125805 -3.9476157 -37.5101316
H -3.4303927 -3.8987074 -35.9202956
C -2.7652385 -5.4947951 -37.177779
O -2.4814048 -5.5085537 -38.373279
N -2.2677174 -6.3806718 -36.3070245
H -2.5945535 -6.342489 -35.3479413

C -1.2607822 -7.3958332 -36.6512525
H -0.3445124 -6.8925585 -36.961776
C -1.6939054 -8.2984128 -37.8176158
O -0.9311845 -8.4717807 -38.7673839
C -0.9587692 -8.2239316 -35.3945171
H -0.5126998 -7.5591821 -34.6545479
H -1.9003645 -8.5918536 -34.9852958
C -0.0205586 -9.403772 -35.5957312
C -0.476939 -10.7237244 -35.4080804
H -1.509788 -10.9114615 -35.1526382
C 0.4018223 -11.8087914 -35.5828742
H 0.0411055 -12.8215519 -35.4605813
C 1.7396798 -11.5814567 -35.9510346
H 2.4073452 -12.4177367 -36.1126058
C 2.197147 -10.2669613 -36.1448257
H 3.2167878 -10.0866761 -36.4561706
C 1.3158525 -9.1847377 -35.975059
H 1.6594669 -8.1812815 -36.1674815
N -2.9508894 -8.7602365 -37.8162996
H -3.5180262 -8.6013591 -36.9897401
C -3.5164419 -9.5705426 -38.9055008
H -2.8830213 -10.4489155 -39.0412821
C -3.4884375 -8.8191223 -40.2441416
O -3.0854456 -9.4032043 -41.2444796
C -4.9353343 -10.0606912 -38.5229828
H -5.4930771 -9.2156787 -38.1147458
C -5.7199256 -10.6008875 -39.7362459
H -5.1748012 -11.428586 -40.1934969
H -5.8704642 -9.8203191 -40.4808494
H -6.7059161 -10.950001 -39.4339102
C -4.8187313 -11.1576382 -37.4374046
H -4.3660146 -12.0503218 -37.8713733
H -4.1598068 -10.814504 -36.6407699
C -6.1553173 -11.5460725 -36.7923586
H -6.7955629 -12.0580817 -37.5094256
H -6.6616767 -10.6550823 -36.4198857
H -5.9674939 -12.2254155 -35.9610021
N -3.8323925 -7.5248185 -40.2879394
H -4.0656676 -7.0556824 -39.4218744
C -3.7691065 -6.745843 -41.5391746
H -4.3005425 -7.2994218 -42.3139513
C -2.3393446 -6.5519942 -42.0526116
O -2.1145754 -6.6247537 -43.2621331
C -4.4332741 -5.3718712 -41.3703427
H -3.8537605 -4.6109815 -41.8970195
H -4.4733346 -5.1006908 -40.3138927

O -5.7412933 -5.3987764 -41.9212897
H -5.6390313 -5.3843889 -42.8827614
N -1.3861787 -6.3126124 -41.1495095
H -1.6547347 -6.2826085 -40.1714205
C 0.0158965 -6.0425797 -41.481886
H 0.0577829 -5.3112942 -42.2894989
C 0.740751 -7.3012983 -41.9896076
O 1.4825599 -7.2298511 -42.9719905
C 0.6993255 -5.4537121 -40.2332656
H 0.5759614 -6.161702 -39.4119538
H 1.768021 -5.3583632 -40.4270006
C 0.1460188 -4.0790425 -39.7977291
H -0.9410754 -4.0968931 -39.8068985
C 0.5881962 -3.7787531 -38.3651539
H 1.6688825 -3.8436922 -38.2817429
H 0.1358835 -4.4987693 -37.6855946
H 0.2728934 -2.7773283 -38.0815491
C 0.5838749 -2.9467471 -40.7300837
H 1.6679734 -2.8669288 -40.751094
H 0.1714852 -2.0035095 -40.3724266
H 0.2150466 -3.1247907 -41.7383577
N 0.4550266 -8.4611916 -41.3890336
H -0.1379686 -8.4416052 -40.564342
C 0.9646168 -9.7595951 -41.8364496
H 2.0196674 -9.6433663 -42.0892642
C 0.2485841 -10.2596498 -43.1035624
O 0.8987943 -10.7616169 -44.0158858
C 0.8455888 -10.7511466 -40.6638834
H 1.3523783 -10.326376 -39.7955445
H -0.210257 -10.8640917 -40.4110157
C 1.4357562 -12.1477194 -40.9424942
H 0.9147976 -12.6038002 -41.7839374
C 2.9345315 -12.1074372 -41.2461209
H 3.4682867 -11.5756565 -40.4587843
H 3.1048157 -11.6059846 -42.1990644
H 3.3258448 -13.1209115 -41.3293987
C 1.228386 -13.036948 -39.7157139
H 1.7534838 -12.6171902 -38.8578697
H 1.6078131 -14.0384338 -39.9180822
H 0.1647686 -13.1057515 -39.4881301
N -1.0698244 -10.0583188 -43.2209953
H -1.5784126 -9.6705319 -42.4337335
C -1.8288138 -10.4543792 -44.4133578
H -1.6406428 -11.512455 -44.6026739
C -1.3632127 -9.6903915 -45.6534104
O -1.0336879 -10.3147171 -46.6612399

C -3.3368109 -10.2801862 -44.1613091
H -3.5384081 -9.2595103 -43.8397539
H -3.6148275 -10.9493552 -43.3461491
C -4.2354917 -10.6192243 -45.3629775
H -5.2387681 -10.8120393 -44.981554
H -3.8764807 -11.5469091 -45.8110439
S -4.390968 -9.3657168 -46.6781462
C -5.1799099 -8.0079527 -45.7729399
H -5.4383667 -7.2116522 -46.4701612
H -4.4931662 -7.6138861 -45.0242914
H -6.0870682 -8.3705426 -45.2881865
N -1.2836702 -8.3547994 -45.597801
H -1.5351508 -7.8696527 -44.7427603
C -0.9788531 -7.5879528 -46.8043397
H -1.457325 -8.1034868 -47.6360249
C 0.521404 -7.5756896 -47.1451478
O 0.8637811 -7.5102657 -48.3264741
C -1.6430715 -6.2032923 -46.748782
H -1.1355764 -5.5871202 -46.0089988
H -2.6802938 -6.3144597 -46.4385527
C -1.6394516 -5.5151375 -48.1094585
N -0.9709235 -4.389742 -48.2157069
H -0.4685428 -4.0199304 -47.4270163
H -0.9501478 -3.9346266 -49.1103779
O -2.2486604 -5.9590541 -49.0785727
N 1.416904 -7.7516498 -46.1678263
H 1.0980991 -7.8154289 -45.2078644
C 2.8375119 -7.9866019 -46.4363406
H 3.1969886 -7.2710589 -47.1758242
H 3.3911814 -7.8211393 -45.5225839
C 3.1279278 -9.4043741 -46.945777
O 3.8298141 -9.5493483 -47.9453507
N 2.447086 -10.4225332 -46.4101552
H 1.9315207 -10.2624063 -45.5490469
C 2.4854058 -11.7979805 -46.9139969
H 3.5152832 -12.1562071 -46.9087821
H 1.8931677 -12.4278665 -46.2504679
C 1.9239234 -11.9295401 -48.3328251
O 2.5932635 -12.4826701 -49.2022689
N 0.7929469 -11.2777152 -48.628854
H 0.267872 -10.8632915 -47.8641561
C 0.2683398 -11.1228987 -49.9938683
H 0.0573556 -12.1067264 -50.4128921
C 1.2826057 -10.4351154 -50.919821
O 1.4660857 -10.8427349 -52.0678355
C -1.03995 -10.3126457 -49.9511539

H -1.8060414 -10.9034882 -49.4476874
H -0.8743164 -9.4136616 -49.3588149
C -1.5543657 -9.8836418 -51.2950987
C -1.1187229 -8.7991881 -51.9758444
H -0.3563124 -8.1147227 -51.6133052
N -1.6791194 -8.7882528 -53.2363242
H -1.3736528 -8.1625098 -53.9676252
C -2.5438482 -9.8457334 -53.4186151
C -3.3411695 -10.2526051 -54.4992947
H -3.3476281 -9.6931163 -55.422582
C -4.1539985 -11.3894695 -54.358604
H -4.801853 -11.6994674 -55.1695271
C -4.1314936 -12.1172815 -53.1550939
H -4.7589197 -12.9925633 -53.044334
C -3.3101688 -11.7105559 -52.0842815
H -3.3046579 -12.2804389 -51.1652787
C -2.499787 -10.5592805 -52.1831049
N 1.96428 -9.3993704 -50.4253499
H 1.7809878 -9.1156849 -49.4709382
C 2.9636726 -8.654149 -51.2011413
H 2.5137098 -8.3680451 -52.1510126
C 4.1751778 -9.5252683 -51.5380323
O 4.584636 -9.5468988 -52.6991712
C 3.3562456 -7.3613529 -50.4714756
H 3.7076416 -7.5770001 -49.4645344
C 4.4220107 -6.5580398 -51.2039862
H 4.0784879 -6.3762013 -52.2179027
H 5.3645424 -7.1059384 -51.2128172
H 4.574357 -5.6062331 -50.6915057
O 2.2203371 -6.5288334 -50.4111686
H 1.6945377 -6.8265429 -49.6442254
N 4.6517677 -10.3483214 -50.5991615
H 4.2697926 -10.2894075 -49.6593257
C 5.6772822 -11.3580867 -50.8568759
H 6.5451977 -10.8700353 -51.3041686
C 5.1923812 -12.4341222 -51.8490049
O 5.8968642 -12.7354913 -52.8110289
C 6.1067911 -11.9647842 -49.5145566
H 5.2580115 -12.4435631 -49.0255728
H 6.499734 -11.1841373 -48.8617151
H 6.8863492 -12.7084471 -49.6781218
N 3.9534366 -12.9249302 -51.7086818
H 3.4267844 -12.6496469 -50.8836684
C 3.334672 -13.8997941 -52.6291921
H 3.9737587 -14.7791788 -52.6745089
C 3.2311635 -13.3581777 -54.0601941

O 3.501972 -14.0879971 -55.0132972
C 1.9611062 -14.3607064 -52.0920771
H 1.4005589 -13.4924383 -51.7508018
C 1.0987965 -15.094003 -53.1308964
H 1.6582642 -15.9270397 -53.5595551
H 0.8015677 -14.4086898 -53.9226587
H 0.1946828 -15.4764729 -52.6554527
C 2.1541222 -15.3188572 -50.9094302
H 2.6359165 -16.2380737 -51.2437664
H 1.18814 -15.559973 -50.4658729
H 2.7765934 -14.8574841 -50.1444956
N 2.9105554 -12.073782 -54.2549198
H 2.6628376 -11.510579 -53.4444772
C 2.8382215 -11.4848562 -55.603466
H 2.5477263 -12.2731829 -56.294507
C 4.1932573 -11.0523229 -56.1735234
O 4.3050569 -11.0014922 -57.3976465
C 1.7500749 -10.3972881 -55.6684607
H 1.973677 -9.6224873 -54.932472
H 1.7766916 -9.9369756 -56.657886
C 0.3181127 -10.9321047 -55.4242165
H 0.2277581 -11.265099 -54.390261
H -0.3748739 -10.1024204 -55.5677658
C -0.0987512 -12.0894509 -56.3560436
H 0.544746 -12.9544503 -56.17014
H 0.0153415 -11.7777132 -57.39766
C -1.5577073 -12.5187353 -56.1029819
H -2.2269318 -11.6802266 -56.3325773
H -1.6736438 -12.7561101 -55.0366384
N -1.920309 -13.7026939 -56.930816
H -1.8494992 -13.5031153 -57.9228767
H -2.903941 -13.9547134 -56.8042702
H -1.3493329 -14.5134468 -56.71273
N 5.239108 -10.8811623 -55.358782
H 5.0832237 -10.9063656 -54.3572045
C 6.6252368 -10.7490509 -55.8568326
H 6.6136699 -10.1725473 -56.7835268
C 7.23075 -12.111499 -56.2120809
O 7.8163342 -12.237301 -57.2830009
C 7.5248285 -9.9796415 -54.8686851
H 7.4639628 -10.4501565 -53.8877875
C 8.9905939 -9.944826 -55.3227672
H 9.0568977 -9.5997377 -56.3555774
H 9.4372889 -10.9357012 -55.2428839
H 9.5579733 -9.2635562 -54.6896495
C 7.0793244 -8.5186274 -54.7449023

H 7.2214826 -8.0117064 -55.6995769
H 7.6884329 -8.0243809 -53.9867521
H 6.0354079 -8.4688468 -54.4529373
N 7.0278598 -13.1595392 -55.4024952
H 6.5409429 -13.011564 -54.5230944
C 7.6271667 -14.4858677 -55.6559459
H 8.6634613 -14.3031979 -55.9438593
C 6.9906641 -15.2279986 -56.8419336
O 7.6716931 -15.9879257 -57.5225639
C 7.6575082 -15.3317962 -54.3638388
H 7.9124323 -14.6589608 -53.5431137
C 6.2996944 -15.9707711 -54.0400428
H 6.0816508 -16.7851927 -54.7321286
H 5.5262014 -15.2171777 -54.1221082
H 6.2972443 -16.3543042 -53.0201916
C 8.777694 -16.3911983 -54.4438692
H 8.5331507 -17.1303691 -55.2074872
H 9.7068405 -15.8989223 -54.7355244
C 9.0351373 -17.1189259 -53.1182629
H 8.173149 -17.7267722 -52.8436591
H 9.2387219 -16.3944539 -52.3289923
H 9.8999306 -17.7735446 -53.2299662
N 5.7387061 -14.9039989 -57.1868191
H 5.2127234 -14.3222172 -56.5485866
C 5.1062204 -15.3009733 -58.4534771
H 5.3874356 -16.3318692 -58.6680816
C 5.6030433 -14.4874707 -59.672003
O 5.0849158 -14.6599443 -60.773668
C 3.5735904 -15.2270815 -58.2905951
H 3.3168007 -14.2373792 -57.910768
H 3.1057086 -15.3358337 -59.269651
C 2.9775319 -16.3018904 -57.3585176
H 3.4899355 -16.2918303 -56.3990126
C 1.4957103 -15.9962792 -57.1210999
H 0.9610944 -15.9934025 -58.0710052
H 1.4152946 -15.0272253 -56.6348142
H 1.0717324 -16.755584 -56.4636815
C 3.0815974 -17.7101515 -57.947436
H 2.616133 -17.7444842 -58.9324024
H 2.5847047 -18.4241014 -57.2906702
H 4.1275589 -18.0041337 -58.0324987
N 6.5584693 -13.5695965 -59.4840437
H 6.9635998 -13.4805897 -58.559394
C 7.1419561 -12.7294521 -60.533019
H 8.0300305 -12.2428227 -60.1296168
H 7.4428545 -13.3604889 -61.3694312

C 6.2113295 -11.6372743 -61.0664694
O 6.4668133 -11.1007314 -62.1414297
N 5.0989407 -11.3521821 -60.3807085
H 4.9813281 -11.7997437 -59.4822739
C 4.011173 -10.5016823 -60.8819061
H 4.0372055 -10.488733 -61.9692609
C 4.1458654 -9.0360507 -60.4550618
O 4.0311999 -8.1487806 -61.2999311
C 2.6640571 -11.1114873 -60.4638078
H 1.8926477 -10.351034 -60.4842377
H 2.730934 -11.4831747 -59.4414438
C 2.2535011 -12.2657448 -61.3868724
H 1.4591474 -12.8339579 -60.902344
H 3.1069454 -12.932034 -61.5231857
S 1.6498686 -11.7327541 -63.0139356
C 1.6214996 -13.3280329 -63.8756168
H 1.2174771 -13.1904539 -64.8799789
H 0.9970689 -14.0358844 -63.3303468
H 2.6341481 -13.7267153 -63.9450518
N 4.4403578 -8.7747006 -59.1755103
H 4.5741277 -9.5745046 -58.565276
C 4.7721047 -7.4432695 -58.6461105
H 4.2329189 -6.6929209 -59.2287529
C 6.2628089 -7.1513591 -58.8255713
O 7.1132365 -8.0286716 -58.7190631
C 4.3124856 -7.3225627 -57.1776623
H 3.2218766 -7.3201566 -57.1538076
H 4.664482 -8.1940433 -56.6248092
C 4.8413101 -6.0586761 -56.4669405
H 5.9269408 -6.1201353 -56.3940316
H 4.5835999 -5.1738691 -57.0510368
C 4.2767258 -5.8952971 -55.0518431
H 4.2724669 -6.8599056 -54.5417714
H 3.2469937 -5.5465683 -55.1283597
C 5.0870934 -4.8828455 -54.2236432
H 4.4787623 -4.580427 -53.3658758
H 5.2958632 -3.991815 -54.8235343
N 6.3573003 -5.466491 -53.7188249
H 6.2395348 -6.3113772 -53.1868018
H 6.9070721 -4.8226841 -53.1439003
H 7.0461657 -5.6510471 -54.4521823
N 6.5831059 -5.8796803 -59.0166479
H 5.8235267 -5.2105063 -59.113986
C 7.9375657 -5.3227778 -59.0310328
H 8.6143732 -5.9856498 -58.4939189
C 7.9299697 -3.9753104 -58.3016906

O 6.8849011 -3.3299543 -58.2056804
C 8.3940536 -5.2114097 -60.498702
H 7.5989185 -4.7461053 -61.083704
C 9.6867695 -4.4393521 -60.7534294
H 10.461601 -4.7634189 -60.0624864
H 9.5092652 -3.3712309 -60.637248
H 10.0261984 -4.6212524 -61.7728531
O 8.6164189 -6.5095718 -60.9963761
H 8.1255677 -7.1366062 -60.4489059
N 9.0868513 -3.5097747 -57.8288001
H 9.9133854 -4.0836863 -57.9019445
C 9.2842826 -2.0970461 -57.484942
H 8.3344924 -1.5820776 -57.6283965
C 10.2397767 -1.4588863 -58.488381
O 11.1822076 -2.0878065 -58.9748673
C 9.6723067 -1.8988096 -55.9998239
H 9.4745394 -2.8358148 -55.4851521
C 11.1589555 -1.5434698 -55.8111287
H 11.3819425 -0.5661663 -56.2441685
H 11.7740098 -2.3086833 -56.2769605
H 11.4226248 -1.4985324 -54.7557414
C 8.7636256 -0.8305709 -55.3496209
H 8.9602225 0.1445593 -55.7991688
H 7.7226656 -1.0931424 -55.5430241
C 8.9349449 -0.7232145 -53.8274753
H 9.9006709 -0.2819557 -53.5826193
H 8.8562241 -1.7099889 -53.3708842
H 8.1532925 -0.0869728 -53.4164581
N 9.9517901 -0.2103244 -58.8094482
H 9.1682783 0.231535 -58.3373574
C 10.654951 0.6360141 -59.7608788
H 11.4393476 0.0802286 -60.2761536
C 11.3137825 1.7707113 -58.9834636
O 10.6744127 2.3280577 -58.0948429
C 9.5994166 1.0937026 -60.7929687
H 8.6089785 1.028997 -60.3467226
H 9.6158493 0.3827572 -61.6188321
C 9.6756245 2.4856417 -61.3678588
N 9.6560584 2.7683857 -62.7304314
C 9.6304442 4.1041051 -62.8320208
H 9.6677495 4.647484 -63.7669766
N 9.599696 4.6686446 -61.6145544
H 9.6754936 5.6576488 -61.4159251
C 9.6037052 3.6646771 -60.6808175
H 9.5692624 3.7963986 -59.6106405
N 12.5267095 2.1509732 -59.3842939

H 13.0058659 1.5797279 -60.0714761
C 13.1036285 3.4786336 -59.1419166
H 12.3791542 4.0801703 -58.6065819
C 13.4003518 4.137936 -60.4980748
O 13.9804095 3.4895145 -61.3751488
C 14.3486196 3.3947989 -58.2404563
H 14.0355429 3.0345998 -57.2602812
H 15.0443904 2.6671256 -58.661884
C 15.0998792 4.7290328 -58.0561606
H 15.5972644 4.9762276 -58.9978091
H 15.8861092 4.5829474 -57.3125827
C 14.2028056 5.9034645 -57.6227446
O 14.1451209 6.8963968 -58.3833069
O 13.5576779 5.8054743 -56.5593021
N 13.0453446 5.4126846 -60.6712948
H 12.6425479 5.9066621 -59.8759317
C 13.3632583 6.2062371 -61.8533931
H 14.1139036 5.6662568 -62.4167197
C 13.9496037 7.5757594 -61.5001384
O 13.3110616 8.3851175 -60.8299668
C 12.1357501 6.3519771 -62.752082
H 11.3121412 6.7960327 -62.1946084
H 11.8710934 5.3727228 -63.1411174
H 12.3807769 6.9994993 -63.5956822
N 15.109189 7.8745898 -62.0873356
H 15.4773862 7.2053572 -62.7563538
C 15.8432781 9.1316037 -61.9126432
H 15.2448618 9.8123746 -61.30726
C 16.0561624 9.7692783 -63.2824116
O 16.4188304 9.0839867 -64.2349318
C 17.1881392 8.8969502 -61.188147
H 17.8076121 8.2561194 -61.8169794
C 16.9764192 8.1765845 -59.8366834
H 16.3622953 8.7964619 -59.1838709
H 16.4432474 7.2414061 -60.0042166
C 17.9261826 10.2389898 -60.9959842
H 17.3004281 10.929227 -60.4298202
H 18.1645092 10.6855659 -61.9607414
H 18.8686102 10.0894265 -60.4738383
C 18.2749524 7.8099776 -59.110778
H 18.7708445 8.7035771 -58.7336403
H 18.9412565 7.2714474 -59.7855677
H 18.0370505 7.1684199 -58.2616609
N 15.8801458 11.0823937 -63.3805037
H 15.5331748 11.5872138 -62.5688742
C 15.9623305 11.817571 -64.6442913

H 16.7952455 11.4316431 -65.2351664
C 16.2306405 13.3158319 -64.4223685
O 16.001996 13.8407697 -63.3256233
C 14.6780471 11.5721577 -65.4405948
H 14.5614297 12.3592306 -66.1828294
H 14.8143697 10.6397859 -65.9771394
C 13.3981278 11.4233831 -64.5967226
H 13.2409175 12.3213791 -64.0054631
H 13.4843556 10.5777558 -63.912386
C 12.2010246 11.1445692 -65.486179
O 12.4220538 10.9836834 -66.7062945
O 11.0711677 11.0577901 -64.952999
N 16.7127389 14.0282412 -65.4533237
H 16.8577185 13.566903 -66.349382
C 16.9273486 15.4814323 -65.3898288
H 16.3871383 15.8476574 -64.5193038
C 16.3447654 16.2645858 -66.5817988
O 16.1558116 15.7419953 -67.6835179
C 18.4109936 15.8096423 -65.1287159
H 18.8261429 15.0930979 -64.4212154
H 18.4804463 16.7986531 -64.677849
C 19.2719235 15.817902 -66.3805842
O 19.9812442 14.8726451 -66.670495
N 19.2629312 16.8803734 -67.1552989
H 19.7115882 16.7659892 -68.0471681
H 18.6938739 17.6945154 -66.9492704
N 16.1171737 17.5560555 -66.3461034
H 16.3244087 17.9218972 -65.4216912
C 15.6409335 18.5213459 -67.3311437
H 15.1563779 17.9880668 -68.1515813
C 16.8207639 19.3019001 -67.9050657
O 17.814395 19.5196466 -67.2156087
C 14.6025547 19.470826 -66.7156745
H 15.0457816 19.9951369 -65.8663646
H 13.7532703 18.8943344 -66.3687152
C 14.1049765 20.4776596 -67.7549781
O 14.0497 21.6808959 -67.4357038
O 13.8939207 20.0301459 -68.9036003
N 16.6932574 19.7127438 -69.1587056
H 15.7361606 19.6535177 -69.5221682
C 17.6417525 20.4891035 -69.9664808
H 18.1709276 19.7718774 -70.5775385
C 18.7397635 21.2279062 -69.1911158
O 19.6163392 20.5303848 -68.6231238
O 18.7905354 22.4280729 -69.0989548
C 16.8603575 21.4153103 -70.9236658

H 16.3055296 22.1240006 -70.3041713
H 17.5800612 21.9809911 -71.516076
C 15.8643249 20.7213124 -71.8783228
H 15.1037594 20.1933291 -71.2956912
C 15.1458514 21.7861056 -72.7164382
H 15.8579213 22.3362843 -73.3312233
H 14.6219113 22.4788679 -72.0561909
H 14.4058411 21.3071409 -73.3609419
C 16.5435476 19.7199673 -72.8232848
H 17.329409 20.2094041 -73.398468
H 15.8002973 19.3095617 -73.5106352
H 16.9602655 18.8885803 -72.2557724
